# Genetic characterization of outbred Sprague Dawley rats and utility for genome-wide association studies

**DOI:** 10.1101/412924

**Authors:** Alexander F. Gileta, Christopher J. Fitzpatrick, Apurva S. Chitre, Celine L. St. Pierre, Elizabeth V. Joyce, Rachael J. Maguire, Africa M. McLeod, Natalia M. Gonzales, April E. Williams, Jonathan D. Morrow, Terry E. Robinson, Shelly B. Flagel, Abraham A. Palmer

## Abstract

Sprague Dawley (**SD**) rats are among the most widely used outbred laboratory rat populations. Despite this, the genetic characteristics of SD rats have not been clearly described, and SD rats are rarely used for experiments aimed at exploring genotype-phenotype relationships. In order to use SD rats to perform a genome-wide association study (**GWAS**), we collected behavioral data from 4,625 SD rats that were predominantly obtained from two commercial vendors, Charles River Laboratories and Harlan Sprague Dawley Inc. Using double-digest genotyping-by-sequencing (**ddGBS**), we obtained dense, high-quality genotypes at 291,438 SNPs across 4,061 rats. This genetic data allowed us to characterize the variation present in Charles River vs. Harlan SD rats. We found that the two populations are highly diverged (F_ST_ > 0.4). Furthermore, even for rats obtained from the same vendor, there was strong population structure across breeding facilities and even between rooms at the same facility. We performed multiple separate GWAS by fitting a linear mixed model that accounted population structure and using meta-analysis to jointly analyze all cohorts. Our study examined Pavlovian conditioned approach (**PavCA**) behavior, which assesses the propensity for rats to attribute incentive salience to reward-associated cues. We identified 46 significant associations for the various metrics used to define PavCA. The surprising degree of population structure among SD rats from different sources has important implications for their use in both genetic and non-genetic studies.

**Author Summary:** Outbred Sprague Dawley rats are among the most commonly used rats for neuroscience, physiology and pharmacological research; in the year 2020, 4,188 publications contained the keyword “Sprague Dawley”. Rats identified as “Sprague Dawley” are sold by several commercial vendors, including Charles River Laboratories and Harlan Sprague Dawley Inc. (now Envigo). Despite their widespread use, little is known about the genetic diversity of SD. We genotyped more than 4,000 SD rats, which we used for a genome-wide association study (**GWAS**) and to characterize genetic differences between SD rats from Charles River Laboratories and Harlan. Our analysis revealed extensive population structure both between and within vendors. The GWAS for Pavlovian conditioned approach (**PavCA**) identified a number of genome-wide significant loci for that complex behavioral trait. Our results demonstrate that, despite sharing an identical name, SD rats that are obtained from different vendors are very different. Future studies should carefully define the exact source of SD rats being used and may exploit their genetic diversity for genetic studies of complex traits.

## Introduction

Rats are among the most commonly used organisms for experimental psychology and biomedical research. Whereas research using mice makes extensive use of inbred strains, research conducted in rats typically uses commercially available *outbred* rats. Sprague Dawley (SD) rats are very commonly used; we identified 4,188 publications in the year 2020 that contained the key word “Sprague Dawley”. Rats identified as “Sprague Dawley” are distributed by several vendors. Each vendor has multiple breeding locations, and each breeding location has one or more rooms or “barriers facilities” which may represent population isolates. Prior studies have identified numerous physiological differences between SD rats obtained from different vendors [1–5]. Despite these observations, many researchers may assume that SD rats obtained from different vendors, breeding facilities or barrier facilities are largely interchangeable. There has been little research into the genetic diversity and population structure that exists among commercially available outbred rats [6]. Prior rat genetic studies have used F_2_ and more complex, multi-parental crosses of inbred strains for quantitative trait loci (QTL) mapping [7–9] and GWAS [10–13]; however, we are unaware of any such studies that have employed commercially available outbred rats. We and others have demonstrated the potential benefits and challenges associated with the use of commercially available outbred mice for GWAS [14–16], suggesting that similar studies in rats might also be of value.

SD rats originated in 1925 at the Sprague-Dawley Animal Company (Madison, WI), where they were created from a cross between a hooded male hybrid of unknown origin and an albino Wistar female [17]. In 1950, Charles River Inc. began to distribute SD rats commercially. In 1980, Harlan Inc. (now Envigo, Inc.) also began to distribute SD rats after their acquisition of Sprague-Dawley, Inc. [18]. In 1992, Charles River reestablished a foundation colony of SD rats, using 100 breeder pairs from various existing colonies [19]. The resulting litters were used to populate SD colonies globally and have since been bred using a mating system that minimizes inbreeding. Every 3 years, Charles River replaces 25% of their male breeders in each production colony with rats from a single foundation colony. Charles River also replaces 5% of their foundation colony breeding pairs with rats from the production colonies on a yearly basis. These practices are intended to reduce genetic drift between the production colonies [20]. Harlan also reported using a rotational breeding system to limit inbreeding [21]; however, more detailed information was not publicly available. Since Harlan’s acquisition by Envigo, the process has become more transparent [22]. Envigo follows a Poiley rotational breeding scheme [23], whereby animals are cycled through different sections of the colony with each generation, reducing genetic drift and inbreeding.

We used SD rats from multiple vendors, breeding locations, and barrier facilities to elucidate the genetic background of SD and to perform a GWAS of a complex behavior. DNA samples and phenotypic information were obtained from rats used in multiple studies as part of an unrelated Program Project grant (P01DA031656) concerned with individual variation in the propensity to attribute incentive value to food and drug cues [24,25]. All rats were first screened for Pavlovian conditioned approach behavior (PavCA) [26], which provides an index of the degree to which a reward cue has been attributed with incentive salience. Although the genetic analyses reported here were not part of the original design, our study took advantage of the opportunities afforded by that large, behavioral study. We extracted genomic DNA from available tissue samples and then applied double digest genotype-by-sequencing (ddGBS) to obtain dense genotypes for 4,625 SD rats. We used these genotypes to genetically characterize different populations of SD. Next, we used those genotype data and behavioral phenotypic data to perform what we believe is the largest rodent GWAS ever undertaken. Our results provide insights about the population structure and suitability of SD for GWAS and the genetic underpinnings of Pavlovian conditioned approach.

## Results

### Phenotype

The final quality controlled dataset consisted of 4,061 genotyped male SD rats that were also phenotyped for Pavlovian conditioned approach; 2,281 from Harlan and 1,780 from Charles River, from 5 and 4 different breeding locations, respectively. As noted previously [26], we found that the metrics used to describe performance in PavCA are highly correlated (S1 Fig). Additionally, several of the base and composite PavCA metrics had tail-heavy distributions due to biased responding in sign- and goal-tracking from the animals during the testing time periods (S2 Fig). For this reason, we chose to quantile normalize all measurements prior to mapping. The PavCA index score [26], which has been used previously to categorize rats into sign-trackers (**STs**), goal-trackers (**GTs**), and intermediate responders (**IRs**), showed the expected divergence and stabilization for STs and GTs over the five days of testing (Fig 1A) [26].

**Fig 1.**
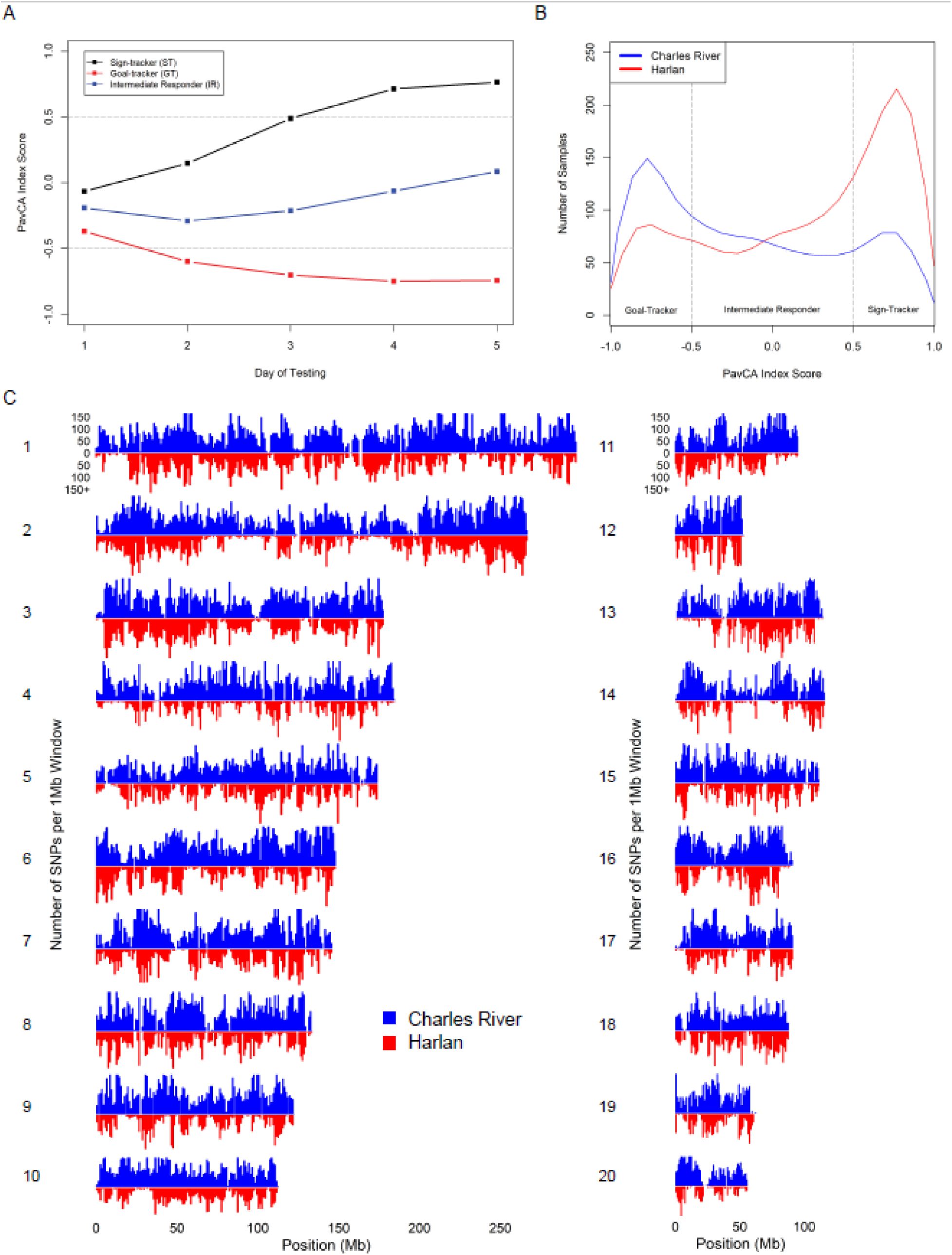
PavCA index score progression across days and distribution between Charles River and Harlan. (A) Samples were classified as ST, GT, or IR based on their average day 4 and 5 PavCA index score. PavCA index scores for each day of training (1-5) were averaged across all STs, GTs, and IRs. (B) Density curves of average day 4 and 5 PavCA index score in Harlan (n=2,281) vs. Charles River (n=1,780) SD rats. (C) Filtered SNP density per 1Mb window in Charles River vs. Harlan samples for all 20 autosomes.

Charles River and Harlan rats had significantly different average PavCA index score distributions (Fig 1B; Welch’s 2-sample t-test, p-value < 2.2×10^−16^), with Charles River rats biased towards goal-tracking and Harlan rats biased towards sign-tracking. We divided the samples further into the breeding locations that the rats originated from to investigate intra-vendor behavioral variance (S1 Table). The differences in PavCA index score between breeding locations within vendor were smaller, but still significant (S3 Fig) [6].

### Genotyping and genetic characterization of SD rats

Variant calling was performed on the full sample of genotyped SD rats. Subsequent quality and minor allele frequency filtering of the variants was initially completed for each vendor separately, under the notion that the observed differences in PavCA performance between vendors may reflect underlying genetic variation. We identified more single nucleotide polymorphisms (SNPs) for the rats from Charles River compared to Harlan (214,309 vs 114,568; S2 Table). Fig 1C compares the distribution of SNPs across each chromosome for each vendor. There were some regions where Harlan had few SNPs, but Charles River had many, and other regions where both Harlan and Charles River had few SNPs. The observed jagged distributions were expected due to the use of double digest genotyping-by-sequencing (ddGBS) to capture the genotypes genome-wide (S1 Text). After combining the two SNP sets, we identified a total of 234,887 unique, high-quality, bi-allelic SNPs. Using 381 samples with two replicates of sequencing, we were able to evaluate our genotyping accuracy by looking at the discordance between calls made in the replicates. The discordance rate between replicates was calculated to be ∼1.7%. However, since discordance putatively reflects random errors in either replicate, it is expected to be approximately twice the true error rate, suggesting that our genotypes were >99% accurate (0.85% error rate; although we cannot exclude the possibility that certain errors were consistent across samples).

In addition to the variants described above, which were obtained using ANGSD/Beagle, we also used STITCH as an alternative genotyping pipeline [27]. This approach identified a larger total number of variants (S2 Table); however, after pruning SNPs with high linkage disequilibrium (LD; r^2^ ≥ 0.95), STITCH produced fewer SNPs compared to ANGSD/Beagle. Preliminary GWAS suggested the two SNP sets produced broadly similar results. Ultimately, we chose to use the ANGSD/Beagle variants for all subsequent analyses.

To further investigate potential genetic divergence between Charles River and Harlan, we performed principal component analysis (PCA) on a set of 4,502 LD-pruned (r^2^ < 0.5) SNPs with MAF > 0.05 across both vendor populations (Fig 2B). The first PC corresponded to vendor (Charles River or Harlan) and accounted for ∼33.7% of the variance. The second PC accounted for just ∼0.9% of the variance and reflected population structure within Harlan SD rats. To investigate the within-vendor structure further (Fig 2A), we performed PCA on the samples from each vendor separately using the same set of SNPs. Fig 2C and 2D show evidence of substructure at both the level of breeding location (i.e. the city) and barrier facility (i.e., the segregated breeding areas within the building). Interestingly, rats from some barrier facilities showed greater divergence from barrier facilities within the same breeding location than barriers at other locations.

**Fig 2.**
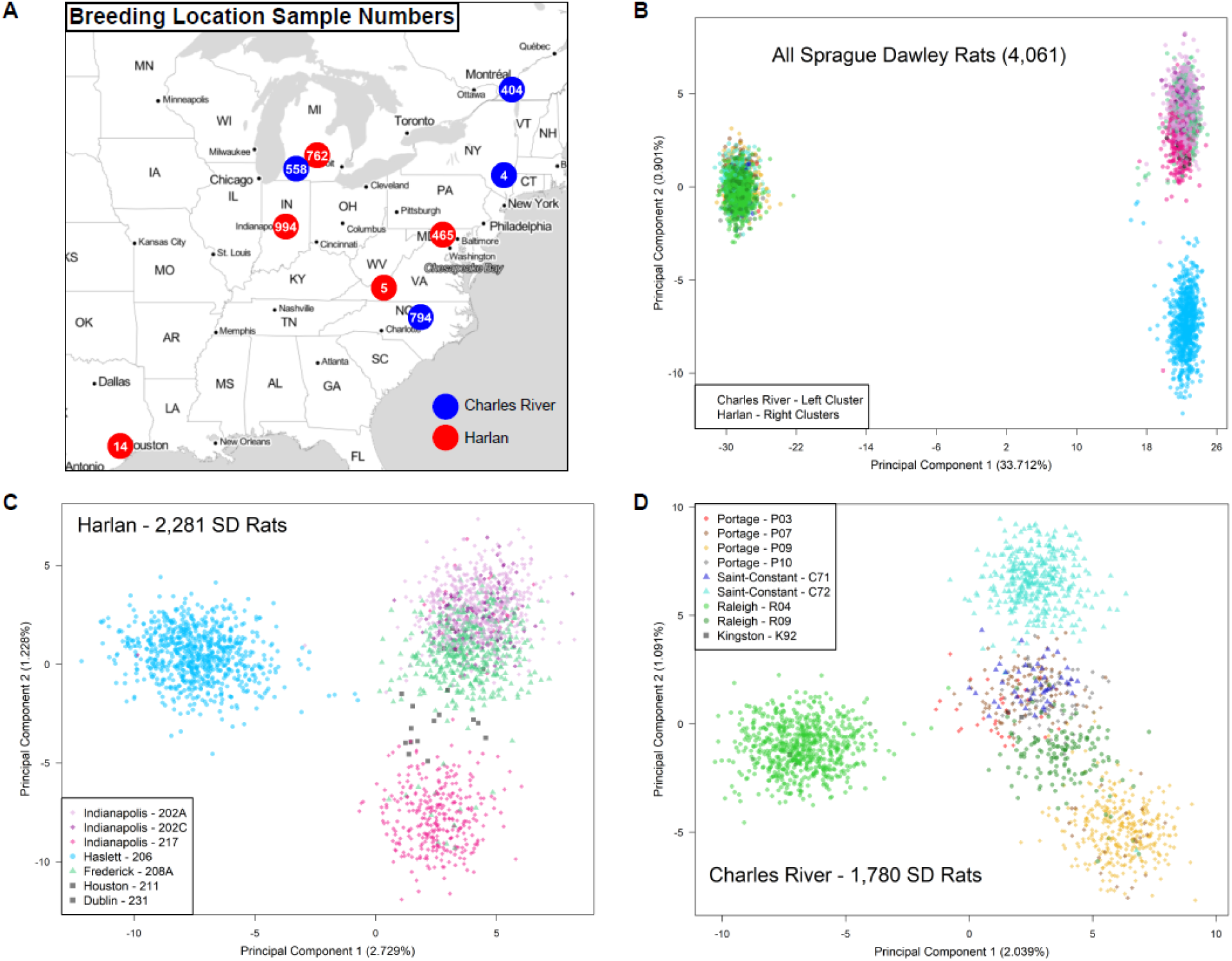
Genetic architecture of SD rats from Charles River vs. Harlan. (A) Map of the nine vendor breeding locations and the number of SD rats obtained from each location. (B) A summary of the genetic data from all 4,061 SD rats based on principal components 1 and 2 from PCA. Each point represents a sample. The left cluster is composed of samples from Charles River and the right clusters are composed of samples from Harlan. (C-D) Repeated PCA analyses on subsets of the samples from Harlan and Charles River, colored by barrier facility of origin.

Due to the stark clustering of samples within and across certain barriers (Fig 2C & Fig 2D), we opted to divide the sample set up based on PCA clustering for all subsequent analyses, including the GWAS for PavCA. This heuristic sample division resulted in seven subgroups, each containing genetically-similar samples: SD rats originating from Harlan 202A/202C/208A, Harlan 206, Harlan 217, Charles River R09/P03/P07/P10/C71/K92, Charles River R04, Charles River P09, and Charles River C72. Variant filtering steps were reapplied to each subgroup individually to reach final sets of SNPs (S4 Table). A more lenient minor allele frequency (MAF) threshold was applied to permit more variants to pass filtering.

We observed a large difference in the minor allele frequency (MAF) distributions for subgroups of Charles River vs Harlan origin (Fig 3A), with Charles River rats harboring more variation and a far greater proportion of SNPs with high MAF (>0.05). This observation could reflect the fact that Charles River has adhered to their International Genetics Standardization Protocol for 25+ years, whereas Harlan appears to have focused on maintaining diversity within breeding colonies and may have allowed for a moderate degree of drift between them. We went on to examine the levels of linkage disequilibrium (LD) in the subgroups by constructing LD decay curves (Fig 3B). These curves show the rate at which LD between two genetic loci dissipates as a function of the distance between the loci. Harlan rats had more extensive LD compared to Charles River. For contrast, we included the decay curve for Swiss Webster (CFW) mice, a commercially available outbred mouse population that has been successfully used for GWAS in the past [14,15].

**Fig 3.**
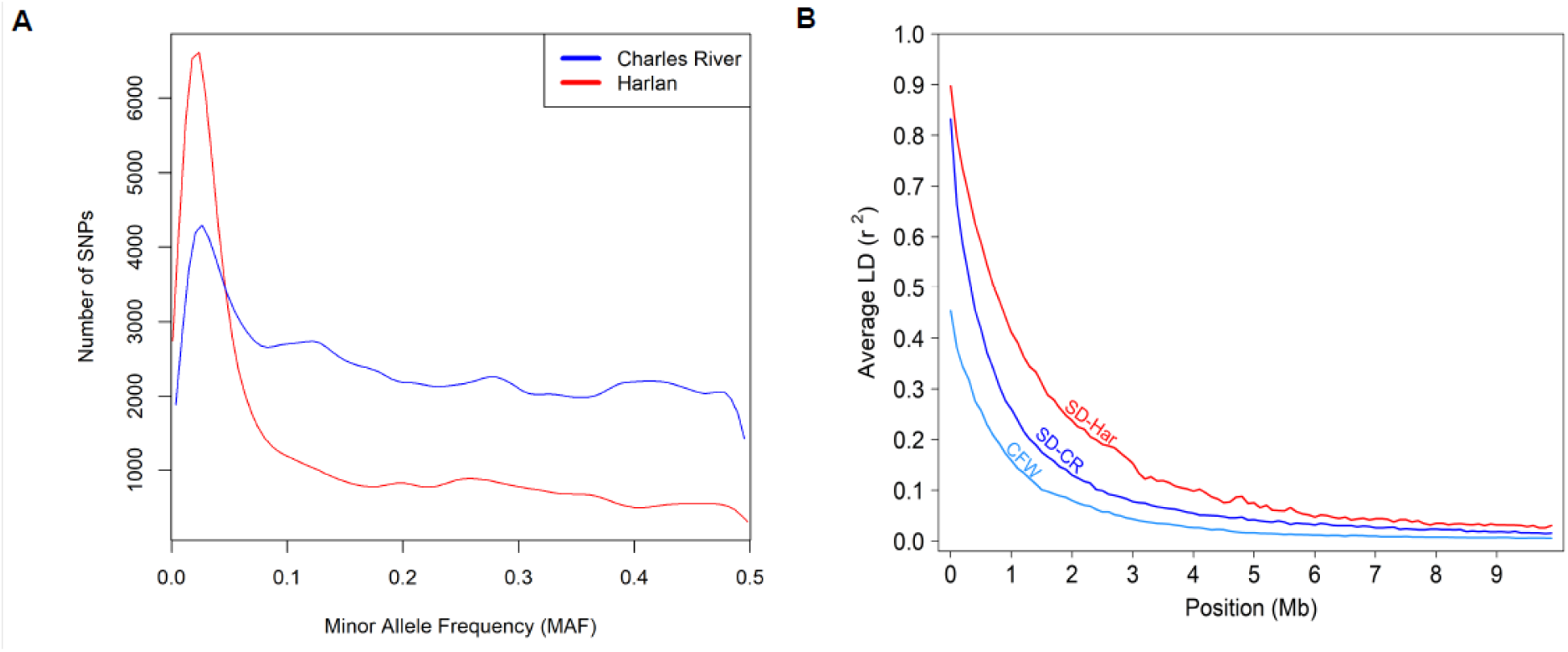
SNP MAF distributions and comparison of linkage disequilibrium decay rates. (A) Density curves of minor allele frequencies for 214,309 SNPs in Charles River and 114,568 SNPs in Harlan, after removing SNPs with MAF < 0.01. (B) Linkage disequilibrium decay rates in SD rats from both vendors and outbred Swiss Webster (CFW) mice.

The fixation index (F_ST_) is a statistic widely employed by population geneticists to measure the level of structure in populations [28]. It is calculated using the variance in allele frequencies among populations; values closer to 0 indicate genetic homogeneity, and values closer to 1 indicate genetic differentiation. We calculated the pairwise F_ST_ between all seven subgroups (Table 1). F_ST_ values between vendors were ∼0.423, which is over 2-fold greater than corresponding estimates for major human lineages [29], whereas the values for different subgroups within a vendor were substantially lower.

**Table 1.**
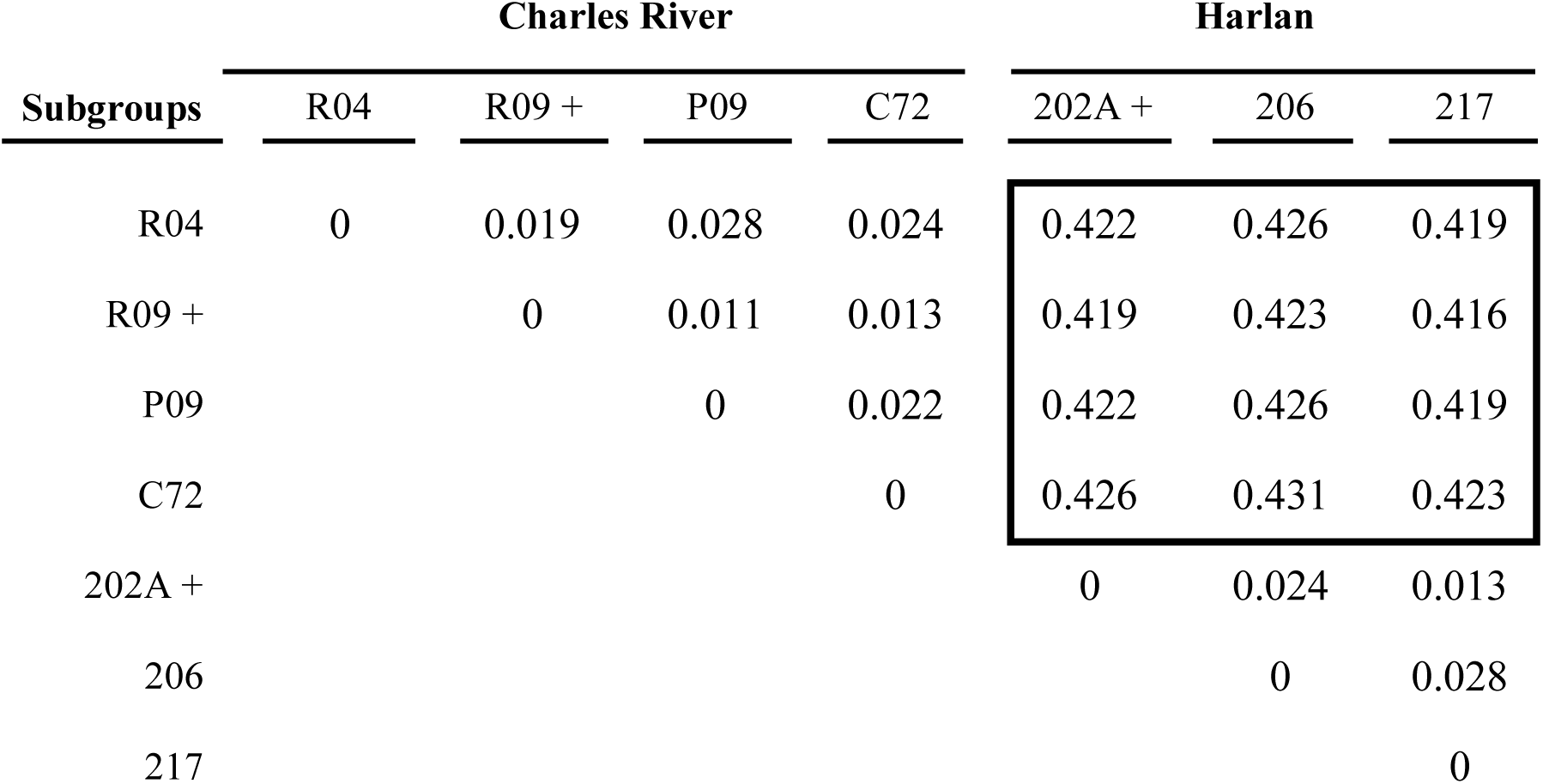
Pairwise F_ST_ statistics for Harlan and Charles River breeding locations.

| Subgroups | Charles River |  |  |  | Harlan |  |  |
| --- | --- | --- | --- | --- | --- | --- | --- |
|  | R04 | R09 + | P09 | C72 | 202A + | 206 | 217 |
| R04 | 0 | 0.019 | 0.028 | 0.024 | 0.422 | 0.426 | 0.419 |
| R09 + |  | 0 | 0.011 | 0.013 | 0.419 | 0.423 | 0.416 |
| P09 |  |  | 0 | 0.022 | 0.422 | 0.426 | 0.419 |
| C72 |  |  |  | 0 | 0.426 | 0.431 | 0.423 |
| 202A + |  |  |  |  | 0 | 0.024 | 0.013 |
| 206 |  |  |  |  |  | 0 | 0.028 |
| 217 |  |  |  |  |  |  | 0 |

We speculated that some of the rats in our sample might also share close genetic relationships with one another. We used *plink 1.9* [30,31] to estimate the pairwise proportions of identity-by-descent (IBD; Panels A and C in S4 Fig), which showed that while most rats were only distantly related, a subset shared closer familial relationships. We removed several rats that showed high levels of relatedness with many other samples (presumably due to technical error), as well as any with unreasonably high levels of IBD (S5 Table; Panels B and D in S4 Fig).

### SNP heritability and genome-wide association analyses

Although evidence from selective breeding studies has suggested that behavior in the Pavlovian conditioned approach procedure is heritable [32], we are not aware of any specific heritability estimates for PavCA. We used GCTA to calculate the proportion of the variance in the base and composite PavCA metrics that could be explained by the set of SNPs unique to each subgroup and the within-vendor SNP sets (S6 Table). Due to the low subgroup sample sizes, the heritability estimates generally only reached significance when considering the vendor as a whole. The SNP heritability estimates for all PavCA metrics in Harlan ranged from ∼4-11%, whereas they were ∼4-21% for Charles River; on average the estimated heritability was about ∼1.9-fold greater in Charles River. Importantly, some of the highest heritabilities were for metrics used to designate sign-trackers vs. goal-trackers, such as the average of day 4 and day 5 response bias, probability difference, and PavCA index score. However, even the heritability estimates from Charles River were lower than SNP heritability estimates for many other behavioral traits [12,14,15,33].

Next, we performed GWAS for various metrics derived from Pavlovian conditioned approach by using GEMMA to fit a linear mixed model that allowed us to account for population structure with a relatedness matrix. Initially, we planned on performing the GWAS separately for rats from Charles River vs. Harlan to maximize sample size; however, we observed marked inflation in our Q-Q plots for a number of PavCA metrics (S5 Fig). This pattern of inflation could reflect true signal that is amplified by the large LD-blocks characteristic of small breeding populations -- if a locus shows significant association with a trait, all SNPs co-occurring in the same LD-block will have inflated test statistics, thereby artificially enriching the Q-Q plot with mid-range p-values. However, inflation in a Q-Q plot might also reflect a failure to account for population structure Given the degree of the observed inflation, we suspected that the LMM might have been insufficient to account for the residual subgroup structure within each vendor. Therefore, we chose to perform individual GWAS on each subgroup followed by meta-analysis of the results.

For each of the seven subgroups, all 11 PavCA metrics were examined for all five days of training. Although PavCA data are often analyzed such that rats are categorized as sign-trackers, goal-trackers, and intermediate responders, based on their PavCA performance on days 4 and 5 [26], we chose to analyze behavior on all 5 days since those results could yield insights into loci involved in learning this complex behavior. Significance thresholds were determined using MultiTrans, a parametric bootstrapping resampling approach [34]. Genome-wide significant results from the individual subgroup GWAS are available in S1 File, and the Manhattan plots for each analysis can be viewed in S2 File. The sample sizes for the subgroup analyses were small and the GWAS yielded a marginal number of genome-wide significant results. In total, we discovered six independent genome-wide significant hits across five of the seven subgroups, representing PavCA metrics from days 3, 4, and 5 of training. All six significant hits were unique to the subgroup they were identified in. This could be due to incomplete power, highly variable allele frequencies between the groups, or the existence of unique modifier loci. Alternatively, these hits may be artifactual and occurred by chance due to the number of analyses that were run. Importantly though, we did not observe any significant inflation in the Q-Q plots for the GWAS in these 7 subgroups (S3 File). To further assess how well we controlled population structure, we used LD score regression [35]. We found that the average deviation from the expected intercept was only 0.027 for the subgroups (S7 Table), indicating that dividing the samples into seven subgroups was sufficient to control population structure.

To increase our power and improve our ability to identify true signals of association, we meta-analyzed the results for all seven subgroups together, as well as in smaller sets of subgroups derived from either Charles River or Harlan. The meta-analyses were run with MR-MEGA [36], a program originally designed to accomplish trans-ethnic meta-analyses in populations with potentially heterogeneous allelic effects, which is conceptually similar to this dataset. Genome-wide significant results from the meta-analyses are presented in S1 File, Manhattan plots in S4 File, and Q-Q plots in S5 File. There were five genome-wide significant loci identified in the Harlan-specific meta-analyses, two in the Charles River-specific meta-analyses, and five in the Harlan plus Charles River meta-analyses. Due to the sparsity of SNP genotypes assayed by the ddGBS approach, these loci are generally composed of only a handful of SNPs and their boundaries are not well defined. Notably, of the seven loci identified in the vendor-specific meta-analyses, three were replicated in the Harlan plus Charles River meta-analyses. The four loci that did not replicate were composed of SNPs that were unique to either Charles River or Harlan and therefore could not have been replicated in the Harlan plus Charles River meta-analyses.

One noteworthy finding was a locus identified in both the Charles River-specific and all-subgroup meta-analyses. The two significant SNPs are located on chromosome 9 and are associated with the training day 5 PavCA index score (Fig 4), which measures a rat’s overall propensity of a rat to attribute motivational value to a conditioned stimulus, and the day 5 probability difference, which directly measures the observed skew in responding to a conditioned stimulus vs. a reward delivery cup (S4 File, page 35). Interestingly, the directions of effect for the associated alleles were contrasting between the two vendors. In Charles River subgroups, the associated allele had a negative effect on the trait, whereas in Harlan subgroups, the effect was positive. This observation emphasizes the necessity of using a meta-analysis approach that accounts for heterogenous allelic effects, as the association likely would not have replicated in the broader subgroup meta-analysis with standard meta-analysis approaches. Presumably the effects are opposite because the relationship between the SNP and the causal allele is reversed in SD rats from Harlan relative to Charles River.

**Fig 4.**
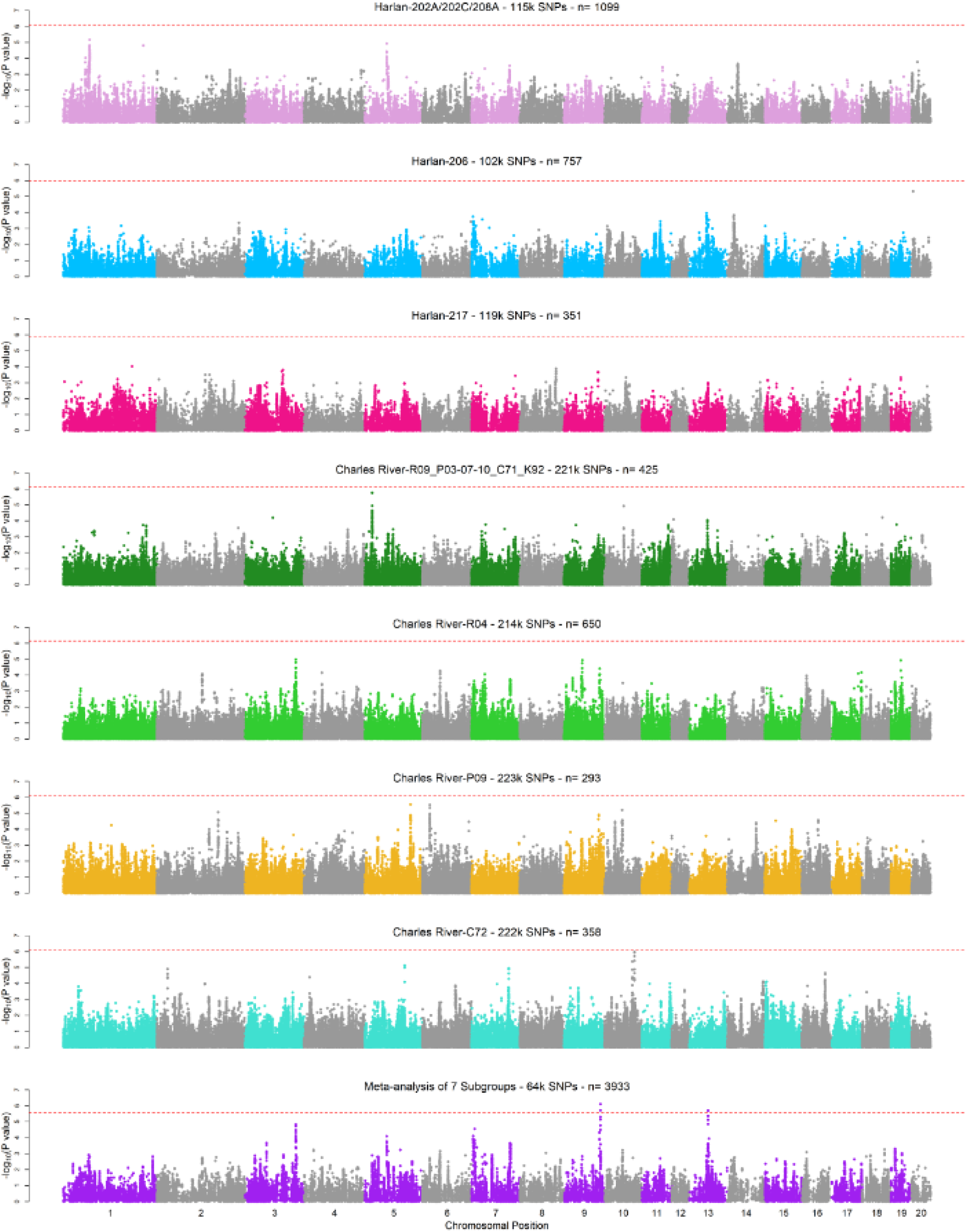
Manhattan plots for the GWAS on PavCA Index Score on day 5 in each subgroup and the meta-analysis. The titles above each of the eight Manhattan plots indicate the population, SNP count, and sample size for each depicted GWAS. The y-axis is the −log_10_ of the p-value from the likelihood-ratio test performed by GEMMA. The x-axis is the genomic coordinate of the variants within each chromosome, in ascending order of chromosome.

Although the gene nearest the most associated SNP is frequently not the causal gene; we examined the nearest gene to the chromosome 9 associated SNPs, which was *Efna5*, located ∼500kb downstream. The gene *Efna5* codes for an ephrin-A ligand for EphA receptor tyrosine kinases and is expressed in the brain. Ephrin-A5 has long been appreciated for its role in guidance of retinal, cortical, and hippocampal axons during development [37–39]. However, this gene continues to be expressed in the adult brain and has been shown to contribute to synaptic plasticity of the mature hippocampus and long-term potentiation [38]. Experiments in mice have shown that increased activation of EphA5, the receptor for Ephrin-A5, can lead to improved memory [40] and that knocking out *Efna5* reduces anxious and aggressive behaviors [41,42].

In addition to *Efna5*, another late-stage PavCA training day association was identified on chromosome 1 and was linked to the average latency to a lever contact (S4 File, page 4). The associated SNP is located in an intron of *Park2*, a large 1.4Mb gene. This gene is most frequently thought of as the causal locus for autosomal recessive Parkinson’s disease in humans. However, in rodents, *Park2* knockout lines do not display the same patterns of motor impairment and neurodegeneration [43]. *Park2* rat knockout lines show reduced expression of dopaminergic receptors, such as trace-amine receptor TAAR-1 and post-synaptic dopamine D2 receptor, in the striatum [44]. Dopaminergic signaling is paramount in the establishment of incentive salience in the Pavlovian conditioned approach paradigm [45,46], making this association an attractive target for further investigation.

## Discussion

Numerous behavioral and physiological studies use outbred Sprague Dawley rats obtained from commercial breeders [47]. The vast majority of these studies are not designed with genetics in mind. While SD and other outbred rats have been used for genetic selection [48–50], ours is the first to explore the use of SD for GWAS. We adapted our genotyping methods to SD and used them to densely genotype more than 4,000 SD rats. These data allowed us to characterize the genetic background of SD rats and to perform GWAS for Pavlovian conditioned approach behavior. This represents the largest rodent GWAS ever undertaken, and the first performed using a commercially available outbred rat population.

We found dramatic genetic differences between SD rats obtained from Harlan versus Charles River. F_ST_ estimates show that SD rats from Harlan and Charles River are more diverged than the major human ancestry groups [29] and nearly as diverged as some subspecies of mice [51]. There was also strong evidence of population structure among the various breeding locations and barrier facilities for each vendor. We found that SD rats from both Harlan and Charles River both showed a rapid decay in LD. However, at the time this study was performed, SD from Charles River appear to have more polymorphisms and more favorable MAF and LD profiles, suggesting that they would be preferable for future GWAS.

Although this study is the first to carefully document population structure within SD rats, it is not the first to highlight phenotypic differences among SD from different vendors. In 1973, Prejean et al. reported that the incidence of endocrine tumors varied among SD rats from different vendors [1]. Then Clark et al. (1991) [52] reported differences in noradrenergic neural projections among SD rats from different vendors. Subsequently, Turnbull & Rivier (1999) [4] reported vendor-specific differences in the response to inflammatory stimuli. Then Fuller et al. (2001) [53] reported vendor-specific differences in hypoxic response among SD rats. Even more recently, there have been additional publications reporting differences for a variety of traits among SD rats obtained from different vendors [2,5,54,55], and even suggesting that these phenotypic differences may extend to differences among a single vendor’s breeding facility [3]. Our own studies have previously reported both behavioral and genetic differences among SD rats obtained from different sources [6], observations that are much more comprehensively explored in the present dataset. Specifically, in addition to wide-spread genetic differences, we have also shown that the SD rats obtained from Harlan for these studies show a much higher proclivity to become sign-trackers compared to SD rats from Charles River. However, neither these prior publications nor the current one can differentiate between two possibilities: that the observed behavioral differences are the result of the different environment in which these animals are raised versus the genetic difference that we have clearly demonstrated. This question could be addressed by future studies in which SD rats are bred in the same facility and the offspring tested in the same manner.

Using genome-wide genotype data, we have provided the first quantitative estimates of the SNP heritability of the component measurements of PavCA. The highest heritabilities (16-20% in Charles River) were seen for measures typically used to assess the propensity to attribute incentive salience to reward cues, including the averages of the day 4 and day 5 PavCA index score, response bias, and latency score. Previous work had shown that SD rats selectively bred for ∼15 generations for high or low responses to a novel environment were also highly divergent for behaviors in PavCA [32]. Those selection studies demonstrate that SD rats have alleles that influence PavCA. The present results are consistent with this conclusion. We obtained lower heritability estimates for SD rats from Harlan compared to Charles River, further emphasizing the genetic differences between SD rats from the two vendors.

As this was the first genome-wide association study to examine PavCA, we had to choose from among many possible ways of summarizing the phenotype. Ultimately, we decided that the analyses should be comprehensive. Therefore, we ran GWAS for all possible PavCA component metrics since it was unclear which had the most predictive power with regard to the overarching behavioral paradigm. In the individual subgroup GWAS, which had limited power due to sample size, we identified only six genome-wide significant loci (alpha = 0.05) despite performing 385 total tests (5 training days x 11 PavCA metrics x 7 groups). Additionally, despite high correlations between many of the metrics, some of the loci we detected were not found for any other PavCA metric or training day. Furthermore, nearly half of the QTL identified in the subgroup and meta-analysis GWAS occurred on earlier training days (days 1-3), while the literature surrounding PavCA focuses on behavior after training (days 4/5). Fortunately, we did observe replication of several associations in the meta-analyses across multiple PavCA metrics (S1 File).

Utilizing the cost-efficient ddGBS to genotype SD rats allowed us to gather variant data on several thousand samples. However, the power of this approach relies on the reasonably large LD blocks that still exist in most commercially-available outbred rodent populations (Figure 3B). Due to the sparsity and uneven distribution of genotypes assayed by ddGBS (Figure 1C), it is difficult to pinpoint the exact source of the signals discovered via GWAS. Although the majority of identified variants do not have clear functional implications, it is possible that these loci are tightly linked to variants that influence the expression or function of nearby protein-coding genes. Among the 46 significant associations from the full sample set meta-analyses (S1 File), we have highlighted two loci. The first locus was discovered on chromosome 9 and was associated with the PavCA index score on the final day of testing, when the rats’ behaviors have typically plateaued (Figure 1A). This locus was located ∼500kb downstream of *Efna5* which codes for an ephrin-A ligand for EphA receptor tyrosine kinases and has known ties learning and memory through its effects on hippocampal development and synaptic plasticity [37–39]. The second was an intronic variant in *Park2*. In rats, knocking out this gene has been shown to reduce expression of dopaminergic receptors, notably TAAR-1, a post-synaptic dopamine D2 receptor. Seeing as dopamine is essential for the attribution incentive salience to reward-associated cues [45,46,56], this result is an enticing one for future follow-up studies.

Because our project used samples from an ongoing, non-genetically focused project, we inherited a design that used Sprague Dawley from many different sources. The full impact of this was not clear until we completed genotyping the rats. The population structure that we encountered was similar to human genetic studies that include individuals from multiple ancestries, and we sought to address it by analyzing each group separately (with an LMM) and then combining the results using meta-analysis (e.g. [57]). In humans, different ancestry groups (e.g. East Asian vs. European) often do not share the same causal variants. Similarly, our power was likely reduced because causal variants were not shared between the subgroups. Modifiers of causal variants might also be dissimilar between the various subgroups, further hindering our meta-analyses. [58]

Regardless of whether differences in SD rats obtained from different vendors are due to genetic or environmental differences, our results demonstrate the need for much greater care in the use of SD rats. Even when the primary research question is non-genetic, researchers should carefully consider whether to use rats from a single vendor and/or breeding facility. Ignoring the genetic background of test subjects can easily confound experimental results. The exact sources (vendor, breeding facility, barrier within each breeding facility) should also be carefully documented and clearly reporting in the results section. Failure to do so can lead to problems with replication, since observations made in SD rats from one source may not extend to SD rats from other sources. In addition to non-replication, a more subtle consequence of using SD rats from multiple vendors or breeding facilities is that spurious correlations can occur. For example, if SD rats from Charles River were higher for traits A and B, compared to SD rats from Harlan, a heterogeneous cohort of SD rats from both vendors would show a significant positive correlation between traits A and B. Such a correlation may not be due to any shared biological mechanisms but could instead be the result of either genetic population structure or environmental differences between SD rats from the two vendors. Thus, even studies that are not intended to address genetics questions could lead to incorrect inferences because of genetic differences among SD rats obtained from different sources.

Our study is the first to use SD rats for GWAS of a complex trait and is the largest rodent GWAS ever undertaken. We have exposed extensive population structure among SD rats that has important implications for genetic and non-genetic uses of SD rats. Based on our data, we suggest that Charles River SD rats have greater genetic diversity, finer scale LD and more favorable MAF, and thus are an attractive choice for future GWAS using outbred, commercially available rats. Although our GWAS was hampered by population structure, our results have identified several genome-wide significant loci that provide the first insights into the genetic basis for PavCA, a trait that is relevant to understating motivational processes, especially in the context of substance abuse.

## Data Availability

Data files that were too large to be included as supplemental content are to be uploaded to a central online repository through the UCSD Library System: https://doi.org/10.6075/J0XS5V8K. Data will be available after publication. The DOI will be able to be followed for access to (1) a VCF file containing all raw SNP calls with dosage data in the original set of 4,625 Sprague Dawley rats, (2) filtered variants in BIM/BED/FAM format for each of the seven SD subgroups, and (3) summary files for the GWAS results for all 55 metric x day combinations for all analyses, individual subgroups and meta-analyses. In addition, FASTQ files from the original ddGBS and light WGS sequencing runs will be submitted to the NCBI SRA. Please contact the authors for more information on accessing this sequencing data. These data, along with the supplemental files and detailed methods included, are sufficient to replicate the analyses performed in this publication.

## Methods

### Sprague Dawley samples

Tissue samples from 5,206 male Sprague Dawley rats were obtained, predominantly from Charles River and Harlan, with a few samples from Taconic. A subset of 4,625 of these rats went on to be genotyped by ddGBS and/or WGS. After sample filtering, a final set of 4,061 genotyped SD rats were used for the population genetic and association analyses. S1 Table lists the number of many samples that came from each vendor, breeding location, and barrier facility. Detailed information about these 4,061 rats is available in S6 File. Behavioral testing was performed between February 2012 and August 2015 as part of work for multiple studies [25,59–65,65–71]. All experiments were approved by the University of Michigan IACUC. Housing, feeding, lighting and other relevant environmental conditions have been previously described. Following sacrifice at the University of Michigan, tissue samples were shipped to the University of Chicago; subsequent processing of those samples is described in the following sections.

### Pavlovian conditioned approach

Pavlovian conditioned approach procedures have been thoroughly described previously [72,73] as a means to assess the tendency to attribute incentive motivational value or incentive salience to a cue that has been repeatedly paired with a noncontingent reward. Briefly, rats are placed into a testing chamber in which an illuminated lever (conditioned stimulus; CS) enters the chamber and after 8 seconds the lever-CS retracts and a food pellet (unconditioned stimulus; US) is immediately delivered into an adjacent food cup. Rats are scored for their three possible responses to the lever-CS entering the cage: approach and interact with the lever, approach and interact with the food receptacle (magazine), or make neither approach. Conditioned responses are captured during the 8-second period during which the lever-CS has entered the chamber, but before the food reward enters the magazine. The following measures are obtained in automated fashion: the number of lever contacts as measured by lever depressions, number of magazine entries as measured by infrared sensor in the food receptacle, and the latency to both during the 8-second lever-CS presentation. The rats are tested in this manner with 25 trials per session, and one session is conducted per day for 5 consecutive days. For the purposes of this project, the number of lever contacts and magazine entries are summed across all 25 trials within a given session, and the latencies are averaged across 25 trials within a session.

Along with response counts and latencies, three additional measurements are recorded: 1) the proportion of trials in a session during which a rat made a lever contact (“probability” of lever press), 2) the proportion of trials during which they made a magazine entry (“probability” of magazine entry), and 3) the number of non-CS (NCS) magazine entries that occurred outside of the 8 second trials (when the cue was not present during the intertrial interval). We also calculated composite scores to categorize rats as sign-trackers (ST; defined as rats that preferentially interacted with the lever-CS), goal-trackers (GT; defined as rats that preferentially interacted with the food magazine), and intermediate responders (IR; rats that vacillated between sign- and goal-tracking behavior) [26]. These scores include: response bias ([lever presses – magazine entries]/[lever presses + magazine entries]), latency score ([average magazine entry latency – average lever press latency]/8), and probability difference ([lever press probability – magazine entry probability]). The PavCA index score is the average of the response bias, latency score, and probability difference. A value of [−1, −0.5] for the PavCA index score indicates a GT, (−0.5, 0.5) indicates an IR, and [0.5, 1] indicates a ST. We performed a Welch’s 2-sample t-test to show that the PavCA index score distributions differed significantly between Charles River and Harlan SD rats (t = 20.161, df = 3908.1, p-value < 2.2×10^−16^). In summary, 11 PavCA metrics were available for analysis, each of which we measured on days 1, 2, 3, 4, and 5 (S8 Table). Phenotypic data is available in S6 File.

### Double digest genotype-by-sequencing (ddGBS)

To obtain genotypes, we used ddGBS, a genotyping method that reduces the complexity of the genome by only sequencing regions proximal to restriction enzyme cut sites [6,74]. We have recently described the technical aspects of this protocol in detail [75]. The ddGBS protocol used in this paper is a synthesis of the GBS approach described in Graboski et al. [76] and used more recently by Parker et al. [14] and Gonzales et al. [33], and an analogous approach known as double digest restriction associated DNA sequencing (ddRADseq) [77]. The full protocol is available in S1 Text.

DNA was extracted from rat tails using the PureLink® Genomic DNA kit. DNA purity was assayed using a Nanodrop 8000 (260/280 ≥ 1.8) and DNA integrity by gel electrophoresis (minimal smearing). Genomic DNA was then digested using two restriction enzymes: *PstI* (6-bp recognition site) and *NlaIII* (4-bp recognition site). Adapter oligos were annealed to overhangs left by *Pst1* and *NlaIII*. The *PstI* adapters contained 48 unique 4-8 bp in-line indexes [14,33,76]. A Y-adapter was annealed to the *NlaIII* cut sites, which controlled the direction of the first round of PCR amplification and thus ensured that the library was primarily composed of fragments with one of each of the adapters. Post-annealing, sets of 24 individual sample libraries were quantified and pooled. Pooled libraries were PCR amplified for 12 cycles, size-selected for 300-450bp using the Pippin Prep, and quality checked by Agilent Bioanalyzer (peak range ∼ 300-500bp and conc. ≥ 20nM). Sequences for the 48 barcoded adapters, Y-adapter, and PCR primers are provided in S2 Text.

Sequencing of pooled libraries was performed by Beckman Coulter Genomics (now GENEWIZ). Sequencing was carried out on the Illumina HiSeq 2500 using v4 chemistry and 125-bp single-end reads. Each lane consisted of a pool of 24 samples, resulting in an average of 8.9 million reads per sample. A total of 4,608 unique ddGBS sample libraries were sequenced. Of these samples, 381 were sequenced twice, resulting in two sets of sequencing data for each sample from the same library prep that were used for to check genotype concordance (S3 Table).

### Light whole-genome sequencing (WGS)

To discover new variants and support imputation, we performed low-coverage whole-genome sequencing of 80 SD rats from this same cohort. The rats were selected to represent sign-trackers, goal-trackers, and intermediate responders from each of the major barriers within the 6 major subgroups of Harlan and Charles River. Sample libraries were prepared using the Illumina TruSeq® PCR-Free Library Prep kit and quality checked using an Agilent Bioanalyzer and qPCR on an Applied Biosystems StepOne Real-Time PCR System to ensure they met Illumina quality standards. Sample pooling (10 samples per pool) was performed by Beckman Coulter Genomics. Each pooled library was sequenced on two lanes of an Illumina 2500 flow cell with 125-bp single-end reads, resulting in an average of 51.6 million reads per sample per two lanes. Assuming that the rat genome (rn6) is ∼2.87Gb, this provided about 180x coverage of the rat genome, or about 2.25x coverage per rat.

### ddGBS Sequence data processing

We have recently described the bioinformatic steps that we use for ddGBS in detail [75]. We follow an analogous approach in this paper, though we deviate at the imputation step due to our use of SD instead of HS rats. Briefly, raw reads from ddGBS were demultiplexed using FASTX Barcode Splitter [78], allowing for 1 mismatch. After demultiplexing, barcodes were trimmed by *cutadapt* [79]. Any reads not matching a sample’s barcode within 1-bp were filtered out. We removed 316 samples for which there were less than 4 million reads, leaving 4,292 samples with ddGBS data. We also used *cutadapt* to trim low-quality base pairs (phred quality score < 20) at the ends of the reads, and to remove 3’ adapter sequences. Reads trimmed to less than 25-bp were discarded. Next, all reads were aligned to the rat reference genome assembly (rn6) using *bwa* [80]. All ddGBS reads were realigned at known indel sites by GATK’s IndelRealigner [81]. Because of a lack of SD-specific variant data, we used variant data from 42 whole-genome sequenced rat strains and substrains [82] as the reference set for indels. We then used GATK to perform base quality score recalibration (BQSR) on the BAM files, using data (SNP & indel) from the 42 rat genomes as the “known” set of variants. For the ddGBS samples that were sequenced twice (381 remaining after filtering for read count), we performed all quality control and variant calling steps in parallel, since our goal was to compare calls made in these samples as a means of estimating the genotyping error rate.

### Light WGS data processing

Raw reads from WGS were processed in an analogous manner to the ddGBS data (detailed above) through the alignment step. After alignment, duplicates reads were removed using *picard* [83]. Reads were then realigned and underwent BQSR. The final WGS BAM files from each lane (2 files per sample) were merged. The WGS BAM files for the 63 samples that had undergone both ddGBS and WGS were then merged.

### Variant discovery and imputation – ANGSD/Beagle

We found that GATK’s HaplotypeCaller tool [81] was ineffectual at making high-confidence SNP calls in our dataset, likely due to the unusual distribution of reads produced by ddGBS. Instead, we used the Samtools variant calling model [84] as implemented by ANGSD [85] to estimate genotype likelihoods from the mapped ddGBS reads. Likelihoods were obtained in 10Mb chunks of the genome, which were subsequently concatenated. Major and minor alleles were inferred from the data based on allele frequency estimates made from the genotype likelihoods. The likelihoods were only estimated at sites where at least 100 samples had reads. ddGBS data results in low call rates at many loci. However, we retained these loci because we anticipated they would be useful for imputation. Next, we used the ANGSD genotype likelihoods to impute missing genotypes (that is for SNPs where only a portion of the rats had genotyping information) using Beagle [86,87], which produced a VCF file containing hard genotype calls (0,1, or 2), dosages ([0,2]), and posterior probabilities for each genotype ([0,1]) for 2,274,118 biallelic SNPs in 4,309 rats (ddGBS+WGS). This is the unfiltered set of SNPs and samples we moved forward with for all subsequent steps. We elected not to pursue variant quality score recalibration using the GATK VariantRecalibrator algorithm [88], because we did not have the required “truth” SNP set. Due to the poorly understood population history of SD rats, it was unclear whether the variation present in the 42 rat genomes would be representative of the variation present in our sample. Using the 42 genomes as a reference for the VariantRecalibrator may also have negatively impacted the calling of novel SD variants.

### STITCH (Sequencing To Imputation Through Constructing Haplotypes)

In addition to variant calling and imputation using ANGSD/Beagle, we also explored the use of STITCH [27], since it is specifically designed for low-coverage WGS data lacking haplotype reference panels. However, ddGBS data is higher coverage and sparser than the input for which STITCH was designed. We used the set of alignment files and known variant sites from the 42 genomes [82], as described above. STITCH queries the user for the number of ancestral haplotypes that exist in the population (K). Due to our lack of knowledge about the founder population, we ran STITCH on a single chromosome using 5 different values of K: 2, 3, 4, 5, and 6. We found that K values of 3, 4, and 5 worked similarly well and chose K=4 to maximize the number of SNPs called and minimize error rate as ascertained by comparison of genotype calls between duplicate samples. STITCH yielded 8,691,886 biallelic SNPs, vastly more than were called with ANGSD/Beagle. However, after applying filters for dosage r^2^/INFO score ≥ 0.9, MAF ≥ 0.01, HWE p-value ≤ 1×10^−7^, as well as removing sites in near perfect pairwise LD > 0.95, we found that the genotypes from STITCH contained fewer SNPs compared to the comparably filtered output from ANGSD/BEAGLE (see S2 Table). For this reason, we did not use the SNP calls made by STITCH in any of the analyses presented in this paper.

### Genotype concordance check

Whereas some of our past projects that used GBS were able to determine the accuracy of GBS genotypes by comparing them to genotypes obtained from SNP microarrays, we did not have microarray-based genotypes for this cohort. Instead, we relied on the remaining 381 duplicate samples whose genotypes were called in parallel. To estimate genotyping accuracy, we compared the rate of concordance of hard genotype calls among the duplicate samples. We first filtered variants by dosage r^2^ (DR^2^), a measure of the accuracy of the genotype imputation performed by Beagle. We tested three different DR^2^ thresholds (≥0.7, ≥0.8, ≥0.9). We then removed variants with MAFs < 1% or that violated Hardy-Weinberg equilibrium at a threshold of 1×10^−7^ in either vendor population. Concordance rates were checked by two methods: 1) by using hard calls in the RAW format from *plink 1.9* and dividing the number of times a genotype call matched between duplicate samples by the total number of SNPs and 2) by taking the mean Pearson correlation of the dosages of the duplicate samples. The results are presented in S3 Table. Similar error estimates were obtained by the hard call and dosage approaches. We chose to move forward with the DR^2^ threshold of 0.9 for all subsequent analyses, which yielded an error rate of ∼0.85%.

### Post-genotyping sample filtering

We removed female rats (n=77) and rats from Taconic Farms (n=4) since they made up a very small fraction of the total sample. We also removed rats that showed poor clustering in the PCA analysis, described below. We filtered out individuals with unusually high or low rates of heterozygosity and high degrees of relatedness as detailed below. Lastly, we excluded a small set of duplicate samples (n=7) and samples missing phenotype data for mapping (n=10). All filters and sample numbers are listed in S5 Table. After these steps, 4,061 unique male SD rats from Charles River (n=1,780) and Harlan (n=2,281) remained.

### Principal component analysis, identity-by-descent, and heterozygosity

We performed principal component analysis (PCA) on the cohort of 4,228 samples filtered for low read count, having been received from Taconic, and females. PCA was performed on genotype calls in R using the *prcomp* function in R [89] on a set of variants pruned for SNPs with MAF ≤ in the combined sample set, SNPs in pairwise LD > 0.5, and SNPs violating HWE at a p-value < 1×10^−7^ in either Charles River or Harlan. The first PC clearly separated animals from Harlan and Charles River; however, there was a set of 54 rats that did not visually cluster as expected at the level of vendor (data not shown). These animals were removed from all subsequent analyses (we assumed they reflected inaccurate records, sample mix-ups, or some other technical problems).

With this further reduced set of 4,174 rats, we continued on to assess the genetic relationships among the rats in our sample. SD rats were ordered in multiple batches over several years, and we suspected some of these rats would be closely related (siblings, cousins, etc.). We reapplied the variant filters used for PCA and utilized the *--genome* function in *plink 1.9* on the resulting SNP set to estimate (the proportion of genotypes predicted to be identical by descent), for all pairs of samples. Panels A and C in S4 Fig show that while most of the animals were unrelated, there were a significant subset of closely related pairs, as well as some pairs with exceedingly high IBD1 rates in Harlan.

We used the *plink 1.9* function --*het* to examine possible inflation or deflation of heterozygosity rates in our samples. Panels A and C in S6 Fig show that a handful of samples in both populations with uncharacteristically high (> 0.25) or low (< 0.25) rates of heterozygosity. We filtered out 34 such samples, as we found they drove the majority of the anomalous signal in our pre-filtering plots in S4 Fig. We also removed 12 samples with more than 30 close relations (defined as [Inline1] ≥ 0.1875) and 32 samples that had a [Inline1] ≥ 0.6 with another sample (likely sample contamination/mix-up). After applying these sample filters, we were left with Panels B and D in S4 Fig and S6 Fig.

After reaching our final set of 4,061 rats, we reapplied the SNP filters on the reduced sample set, resulting in 4,502 SNPs that were polymorphic in both populations. We ran PCA on these SNPs as described above. We then repeated PCA on the samples from Charles River and Harlan separately to examine substructure within each vendor population. Using the results from the vendor-level PCA analysis, new groupings of barrier facilities were determined based on genetic similarity. Barrier facilities that clustered most highly in the PCA were grouped together for subsequent analysis.

### Final variant filtering and minor allele frequency spectrum

After establishing the final cohort of 4,061 rats, we sought to establish a final set of SNPs to be used for the association analyses. Starting from the initial full set of 2,274,118 SNPs, we first removed sites with DR^2^ < 0.9. Then, based on the PCA results, we separated the samples into seven subgroups, each containing genetically-similar samples that clustered highly: SD rats originating from Harlan 202A/202C/208A (Har 202A), Harlan 206, Harlan 217, Charles River R09/P03/P07/P10/C71/K92 (Charles River R09), Charles River R04, Charles River P09, and Charles River C72. These seven groupings were used for all subsequent variant filtering performed in *plink 1.9*, which converts the posterior probabilities we received from Beagle to hard genotype calls, so long as the probability is ≥ 0.9. In each cluster, we removed SNPs with MAF < 0.0001 using the *–maf* option in *plink 1.9*. Lastly, we used the *--hardy* function in *plink 1.9* [90] to perform tests for SNP Hardy-Weinberg equilibrium and removed all SNPs with an HWE p-value < 1×10^−7^ in any of the seven groups. The final counts of samples and SNPs in each barrier grouping are available in S3 Table. We used the *--freq* option in *plink 1.9* to estimate the MAFs for these final filtered sets SNPs. The distributions are show in (Fig 3A).

Taking the union of these seven sets of SNPs, there were a total of 291,438 polymorphic sites passing our filters across all subgroups. Our final set of filtered SNPs contained 75,405 novel SNPs that had not be previously reported by Hermsen et al. [82]. We also compared our final set of SNPs to the most recent dbSNP release for rats (Build 149, November 7, 2016) and found that 178,545 of the 291,438 SNPs we discovered were not present in the current dbSNP build. See the Data Availability section for more information on accessing this data.

### Fixation Index (F_ST_)

To quantify the population divergence between SD rats from Harlan and Charles River, we computed F_ST_ for each SNP in the union set using *smartpca* within the *EIGENSOFT* package [29,91,92]. Due to the substructure within vendor and vendor location that we saw in our PCA analyses, we chose to estimate the pairwise F_ST_ between all seven sample subgroups defined above.

### Linkage Disequilibrium

We plotted the decay of linkage disequilibrium (LD) using the *--r^2^* utility in *plink 1.9* and the procedures described in Parker et. al 2016 [14]. Briefly, each population’s curve included all SNPs with MAF > 20% and pairwise LD comparisons were restricted to SNPs with allele frequencies within 5% of each other. An average r^2^ estimate was obtained using 10,000 randomly selected SNP pairs from each 100kb interval for the distance between two SNPs, starting with 0-100kb and end at 9.9-10Mb. The procedure was repeated for cluster grouping. The CFW mice curve was downloaded from a repository for Parker et al. (http://datadryad.org/resource/doi:10.5061/dryad.2rs41) [14] and used for comparison as another commercially available, outbred rodent stock.

### LMM covariates and phenotype data pre-processing

To select covariates for the GWAS, we performed univariate linear regression for each potential covariate for each PavCA metric. This was done separately for rats from each genetically-similar cluster defined previously. Any covariates that accounted for 1% or more of the variance for at least one PavCA metric were passed into multivariate model selection with the R package *leaps* [93]. Model selection with *leaps* was performed for all metrics for all days of testing, as well as the average of days 4 and 5. Out of the 66 models, all covariates that had surpassed the 1% threshold to reach this step were ultimately selected in at least 40% of the *leaps* models. The covariates included in the for the downstream GWAS analyses were rat’s age at testing, housing (binary – single or multiple), light cycle (binary – standard or reverse), and a set of binary ‘indicator’ variables to model the effects of different experimenters/technicians responsible for the phenotypic testing (S9 Table). All covariates were included in the LMMs for association testing by GEMMA, rather than being regressed out prior to GWAS.

Many of the PavCA metrics were exceedingly non-normally distributed. In most cases, this was expected due to how the behaviors were measured and defined. For example, the rats only had a window of 8 seconds in each trial during which to contact the lever and/or make a nose poke into the food magazine. All values for “average latency to lever press” or “average latency to magazine entry” were therefore necessarily between 0 and 8 seconds. As is typical for latency traits, many of the values were near 0 or exactly 8. Similarly, the “probability” of lever press or magazine entry were very skewed towards the limits 0 and 1, especially after conditioned responding had been established on the later test days of training. Given these unusual distributions, we chose to quantile normalize all metrics within each subgroup prior to association testing, accepting a possible loss in power since samples with identical values are ranked randomly during the quantile normalization procedure.

### SNP-based heritabilities

Heritabilities were estimated separately for each vendor and subgroup using their sets of filtered SNPs. We used the SNP sets to construct genetic relationship matrices (GRMs) for each cluster using GCTA [94]. We then used the restricted maximum likelihood (REML) approach within GCTA on the GRMs, covariates, and quantile normalized PavCA data to calculate the SNP-based heritabilities for each metric on each testing day.

### GWAS

We used GEMMA [95], which implements an LMM for GWAS analysis. We included a GRM as a random term to account for relatedness and population structure. Though beneficial for preventing false positive associations, GRMs can also reduce power to detect QTLs in populations with greater levels of LD; this is due to proximal contamination [96,97]. To avoid this reduction in power, we used the leave-one-chromosome-out (LOCO) approach when constructing the GRMs for each cluster of samples [33,98,99]. As described above, we selected covariates that were included as fixed effects in our model (S9 Table). Any samples missing measurements for a given metric were removed from that analysis. For all GWAS, genotypes were represented as dosages (continuous [0,2]) in lieu of ‘hard’ genotype calls [0,1,2] to account for uncertainty in the genotype calls. Reported p-values come from the likelihood-ratio test (LRT) performed by GEMMA. Results were plotted using a custom R script.

### GWAS meta-analysis

We performed meta-analyses of the GWAS results across all seven subgroups, across only the four Charles River subgroups, and across only the three Harlan subgroups. Due to the significant genetic differentiation between certain subgroups in this meta-analysis, it is possible that tested loci would have differing effect sizes and directions of effect. To account for this and maximize our power, we utilized the software MR-MEGA, originally designed to perform trans-ethnic meta-regression in humans [36]. A large number of the SNPs called in each subgroup were either unique to that singular group or a subset of the groups, making them of low utility in a meta-analysis of all seven subgroups. Therefore, the seven subgroup meta-analyses were performed on the intersection SNPs set for all seven clusters, amounting to 64,442 SNPs. We repeated the meta-analyses for the only the subgroups derived from Charles River (R04, R09, P09, C72) and only those from Harlan (202A, 206, 217) to allow for more SNPs to pass through to the meta-analysis. There were a total of 198,817 shared SNPs between the Charles River subgroups, but only 83,120 shared between the Harlan subgroups. Genome-wide significant results from these meta-analyses are presented in S1 File. For all significant hits, we also report the directionality and strength of effect of the SNPs in all the separate subgroups GWAS.

### P-value significance thresholds

In human GWAS, 5×10^−8^ is widely used as a significance threshold [100]. However, model organisms have vastly different levels of LD, meaning that the effective number of independent tests differs between studies. Therefore, many prior studies have used permutation testing [101,102], where phenotype data is shuffled with respect to fixed genotypes. The computational load of such methods becomes intractable for studies with large sample sizes and several traits being mapped [96]. Thus, we used the sequence of SLIDE [103,104] and MultiTrans [34] to obtain significance thresholds. We used separate thresholds for each subgroup because the number of SNPs, the LD structure, and the marker-based heritabilities were different, all of which affect the estimated number of independent tests performed as part of the GWAS. An advantage of this approach is that only one threshold was needed per cluster for all normalized metrics. We used a sliding window of 1000 SNPs and sampled from the multivariate normal 10 million times to obtain a 0.05 significance threshold. The subgroup-level −log_10_(p) thresholds for Charles River R04, R09, P09, and C72 were determined to be 6.14, 6.13, 6.11, and 6.11, respectively. Similarly, the Harlan 202A, 206, and 217 thresholds were 6.09, 6.00, and 5.89, respectively. Since there is no clear extension of this approach for a meta-analysis with largely varying sets of SNPs and LD patterns, we ultimately obtained the meta-analysis thresholds by taking the subgroup from each vendor with the lowest level of LD (Charles River R04 & Harlan 202A) and filtering their SNP set to contain only the SNPs included in each meta-analysis. We then reran MultiTrans using these pruned sets of SNPs, providing us with a −log_10_ threshold of 5.574 for the over-arching meta-analyses, 5.574 for the within-Harlan meta-analyses, and 5.993 for within-Charles River.

### LD score regression

We utilized LD score regression to more rigorously test whether after dividing samples up into seven subgroups, we were still observing inflation of the GWAS test statistics. LDSC is software designed to determine if observed inflation is due to cryptic relatedness and population stratification or true polygenic signal spread across regions of high LD [35]. Since there is no genetic map available for Sprague Dawley rats, we used a map created for heterogeneous stock (HS) rats [105], a similarly outbred population that shares some ancestry with SD. An LD window size of 5 centimorgans was used, corresponding to the point at which SD LD decayed sufficiently in most subgroups for loci to be considered unlinked. The results of running LDSC for all GWAS in each subgroup are available in S7 Table.

## Supporting information

S1_File - All Significant GWAS Hits

S1_Text - ddGBS Protocol

S2_File - PavCA GWAS Plots Subpops Days 1-5

S2_Text - ddGBS Primer and Adapter Sequences

S3_File - Subpop QQ-Plots

S4_File - PavCA GWAS Plots Meta-Analyses Days 1-5

S5_File - Meta QQ-Plots

S6_File - 4061 SD Phenotypic Data

S6_Table - Heritabilities

S7_Table - LDSC_Intercepts_Subpops

## Acknowledgements

We would like to thank John Novembre, Mark Abney, and Dan Nicolae at the University of Chicago for their advice concerning the statistical analyses involved in this publication.

## Supporting information

**S1 Fig.**
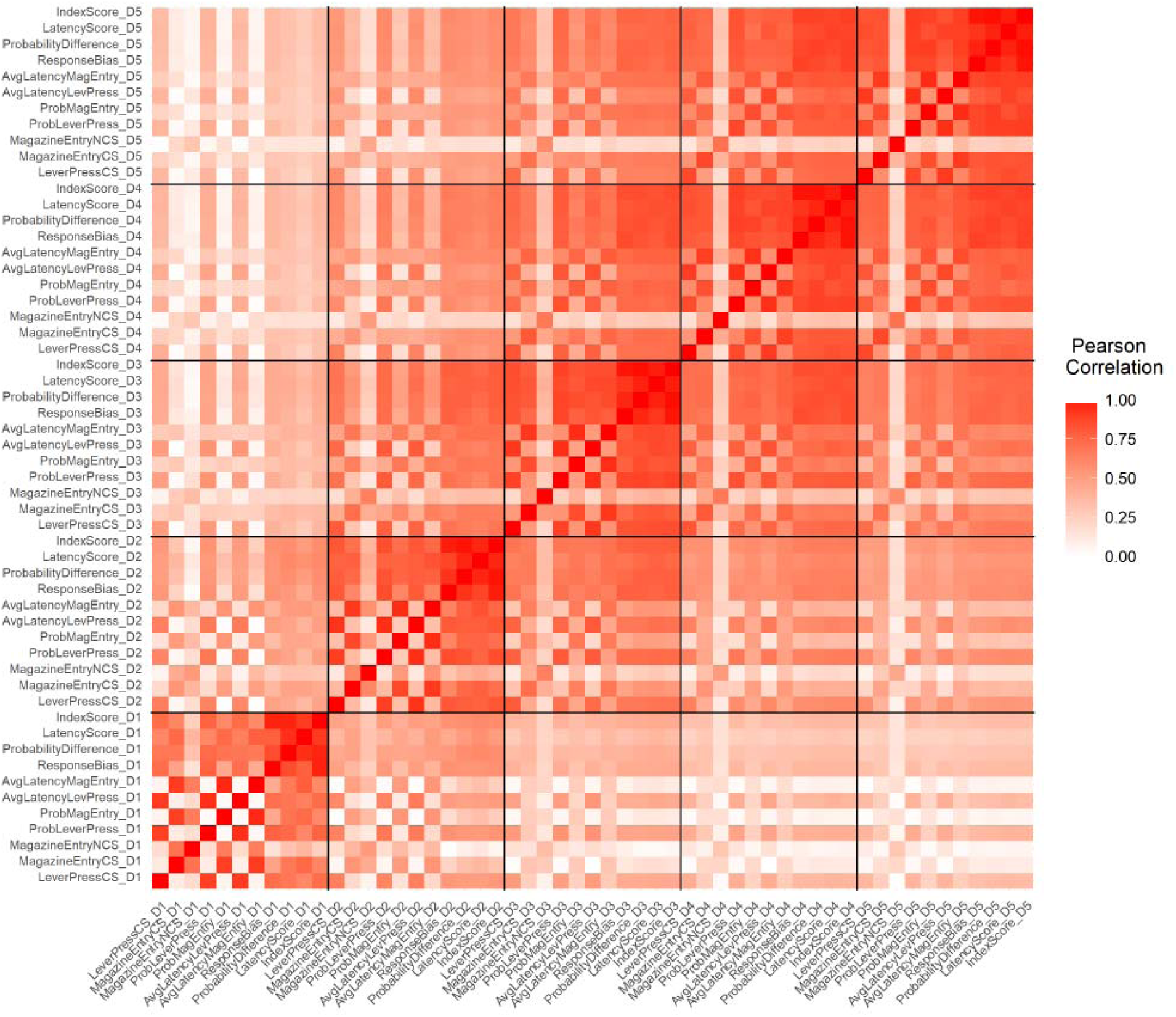
Correlation heatmap of PavCA metrics across days 1-5. The heatmap displays the absolute value of the Pearson correlation coefficient between each pair of PavCA metrics across all 5 days of testing.

**S2 Fig.**
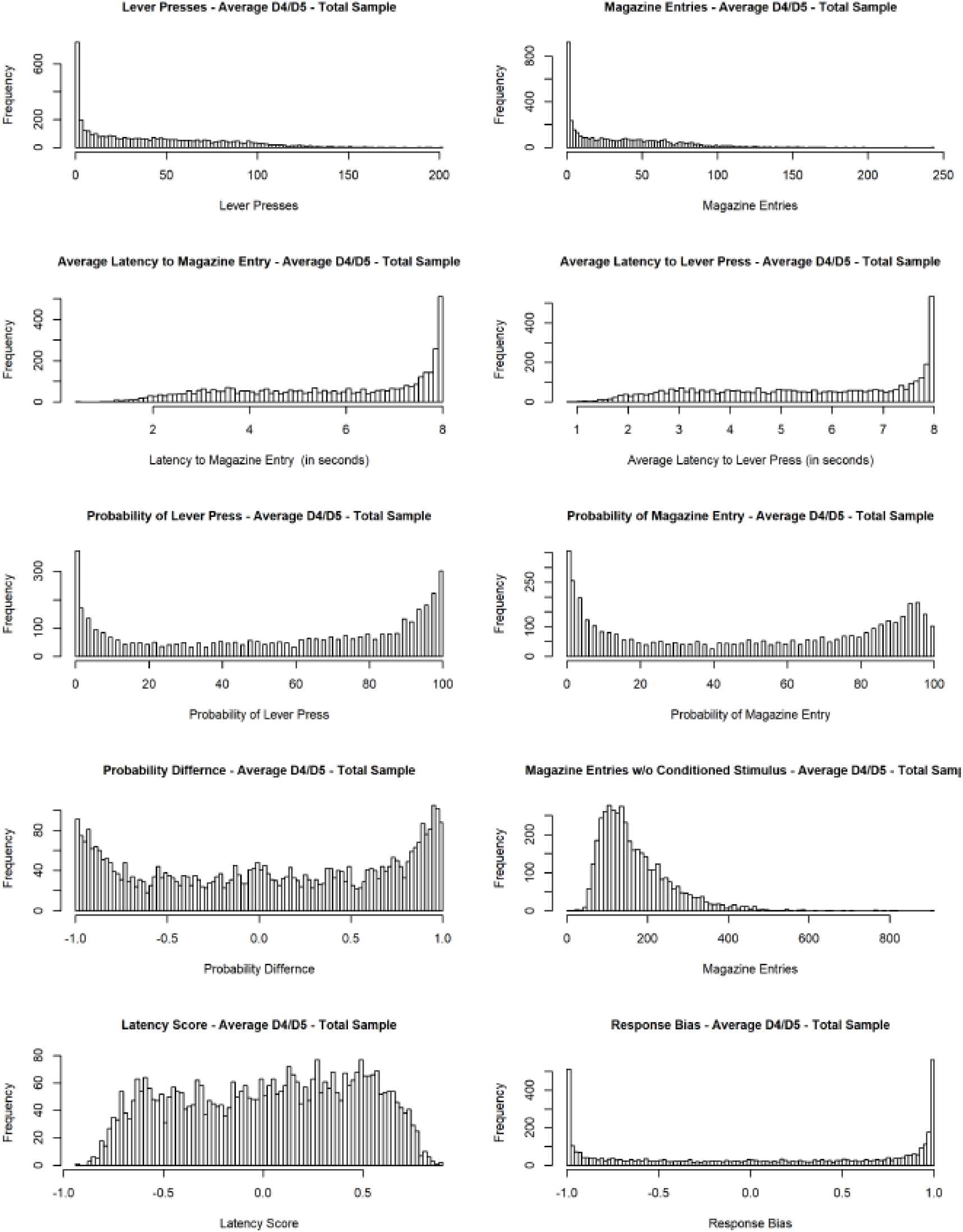
Distributions of the average of day 4 and day 5 measurements for 10 PavCA metrics. Histograms were constructed using measurements from the combined Harlan and Charles River sample set. Not shown is the PacCA index score.

**S1 Table.** Sample origins for all 4,061 SD rats in final filtered set.

|  | # Samples<br>(Barrier Facility) | # Samples<br>(Breeding Location) |
| --- | --- | --- |
| **Total samples post-filtering** | 4,061 | 4,061 |
| Harlan – Dublin, VA – Barrier 231 | 5 | 5 |
| Harlan – Frederick, MD – Barrier 208A | 465 | 465 |
| Harlan – Haslett, MI – Barrier 206 | 762 | 762 |
| Harlan – Houston, TX – Barrier 211 | 14 | 14 |
| Harlan – Indianapolis, IN – Barrier 202A | 546 | 994 |
| Harlan – Indianapolis, IN – Barrier 202C | 97 |  |
| Harlan – Indianapolis, IN – Barrier 217 | 351 |  |
| Charles River – Kingston, NY – Barrier K92 | 4 | 4 |
| Charles River – Portage, MI – Barrier P03 | 32 | 558 |
| Charles River – Portage, MI – Barrier P07 | 121 |  |
| Charles River – Portage, MI – Barrier P09 | 327 |  |
| Charles River – Portage, MI – Barrier P10 | 78 |  |
| Charles River – Raleigh, NC – Barrier R04 | 650 | 794 |
| Charles River – Raleigh, NC – Barrier R09 | 144 |  |
| Charles River – Saint Constant, CAN – Barrier C71 | 46 | 404 |
| Charles River – Saint Constant, CAN – Barrier C72 | 358 |  |
| Unknown Barrier | 61 | 61 |

**S3 Fig.**
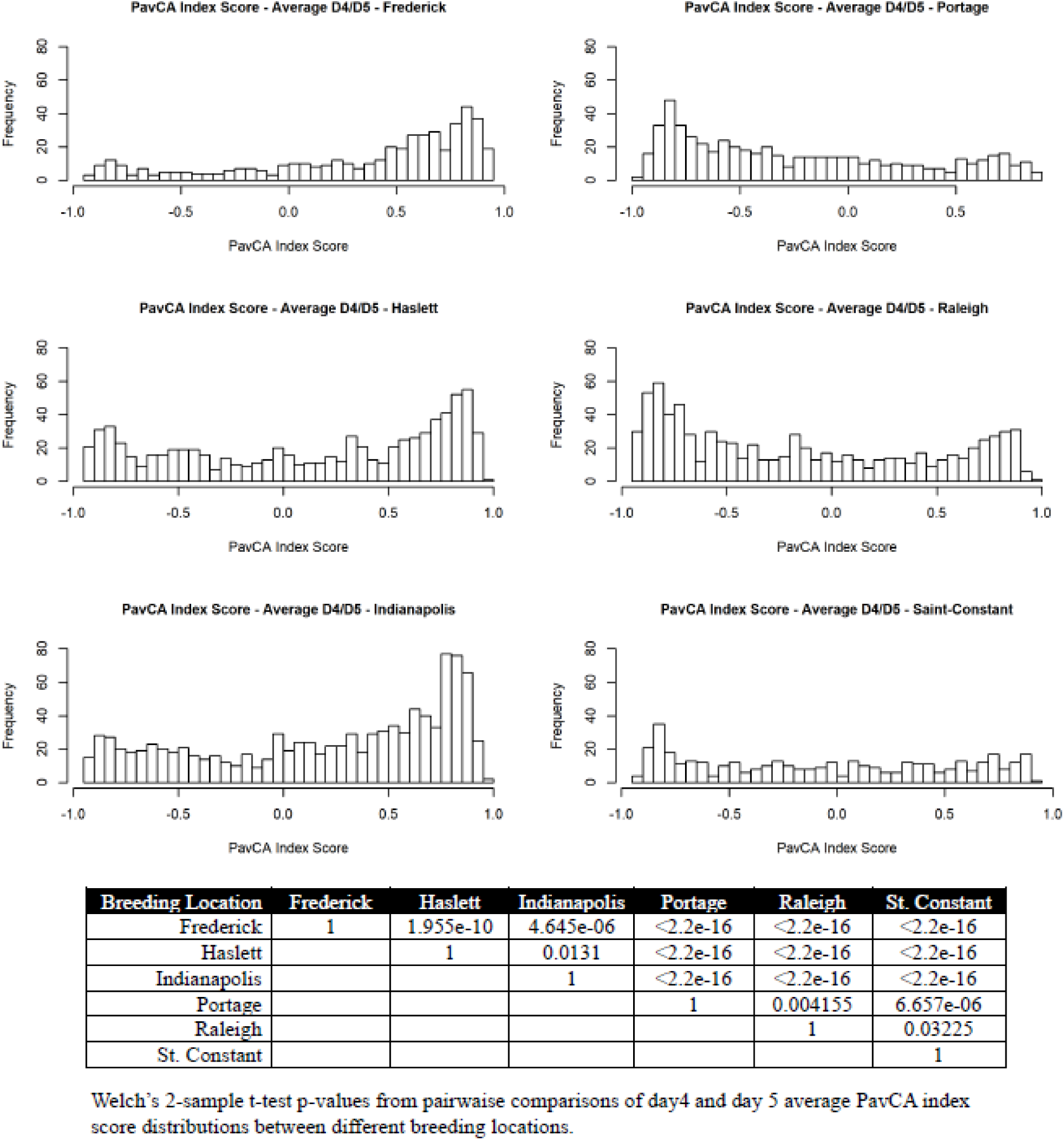
Distributions of the average of day 4 and day 5 PavCA index scores for 6 major breeding locations. Sample numbers for each breeding location can be found in S1 Table. The three locations in the left column are from Harlan, and the three locations in the right column belong to Charles River.

**S2 Table.** List of variant filtering steps and the numbers of SNPs remaining after each step for both ANGSD/Beagle and STITCH. The values highlighted in green are the final SNP totals used for GWAS in Charles River and Harlan. The values highlighted in orange are the counts we used as criteria for choosing ANGSD/Beagle over STITCH.

|  | ANGSD/Beagle |  | STITCH |  |
| --- | --- | --- | --- | --- |
| Sequential Filtering Steps | Number of SNPs Harlan | Number of SNPs Charles River | Number of SNPs Harlan | Number of SNPs Charles River |
| Total called/imputed | 2,274,118 | 2,274,118 | 8,691,886 | 8,691,886 |
| SNPs with dosage $r^2 \geq 0.9$ | 305,779 | 305,779 | 1,992,919 | 1,992,919 |
| SNPs with MAF $\geq 0.01$ | 235,583 | 128,572 | 1,058,812 | 1,843,240 |
| SNPs with HWE p-value $< 1 \times 10^{-7}$ | 214,309 | 114,568 | 930,685 | 1,718,475 |
| SNPs after LD pruning $r^2 < 0.95$ | 44,721 | 93,692 | 25,009 | 75,503 |
| Union set of SNPs | 234,887 |  |  |  |
| Novel SNPs compared to dbSNP (not found in Build 149 - Nov 7, 2016) | 195,299 |  |  |  |
| Novel SNPs compared to 42 genomes ref (not found in Hermesen et al. 2015) | 105,638 |  |  |  |

**S1 Text. ddGBS protocol used to sequence SD rats.**

**S3 Table.** Concordance and error rates for ANGSD/Beagle genotypes at different dosage r2 thresholds. Rates of concordance and genotyping error were calculated by comparing genotypes for 381 duplicate samples called in parallel. Each of the two replicates of the sample was assumed to contribute half the discordant genotypes. Therefore, the per sample genotyping error rate was calculated as half of the observed rate of discordance.

|  | **DR<sup>2</sup>≥0.9** | **DR<sup>2</sup>≥0.8** | **DR<sup>2</sup>≥0.7** |
| --- | --- | --- | --- |
| Total called and imputed by ANGSD/Beagle | 2,274,118 | 2,274,118 | 2,274,118 |
| SNPs passing dosage $r^2$ filter | 305,779 | 427,123 | 541,622 |
| SNPs passing MAF ≥ 0.01 filter | 244,586 | 326,550 | 393,569 |
| SNPs passing HWE $10^{-7}$ filter | 204,104 | 259,111 | 305,660 |
| Mean Pearson **correlation of dosages** (n=381) | 0.983 | 0.969 | 0.957 |
| **Rate of concordance** of hard calls (n=381) | 0.983 | 0.972 | 0.963 |
| **Rate of discordance** (1 – concordance) | 0.017 | 0.028 | 0.037 |
| **Error rate** (Rate of discordance/2)*100 | 0.85% | 1.4% | 1.85% |

**S4 Table.**
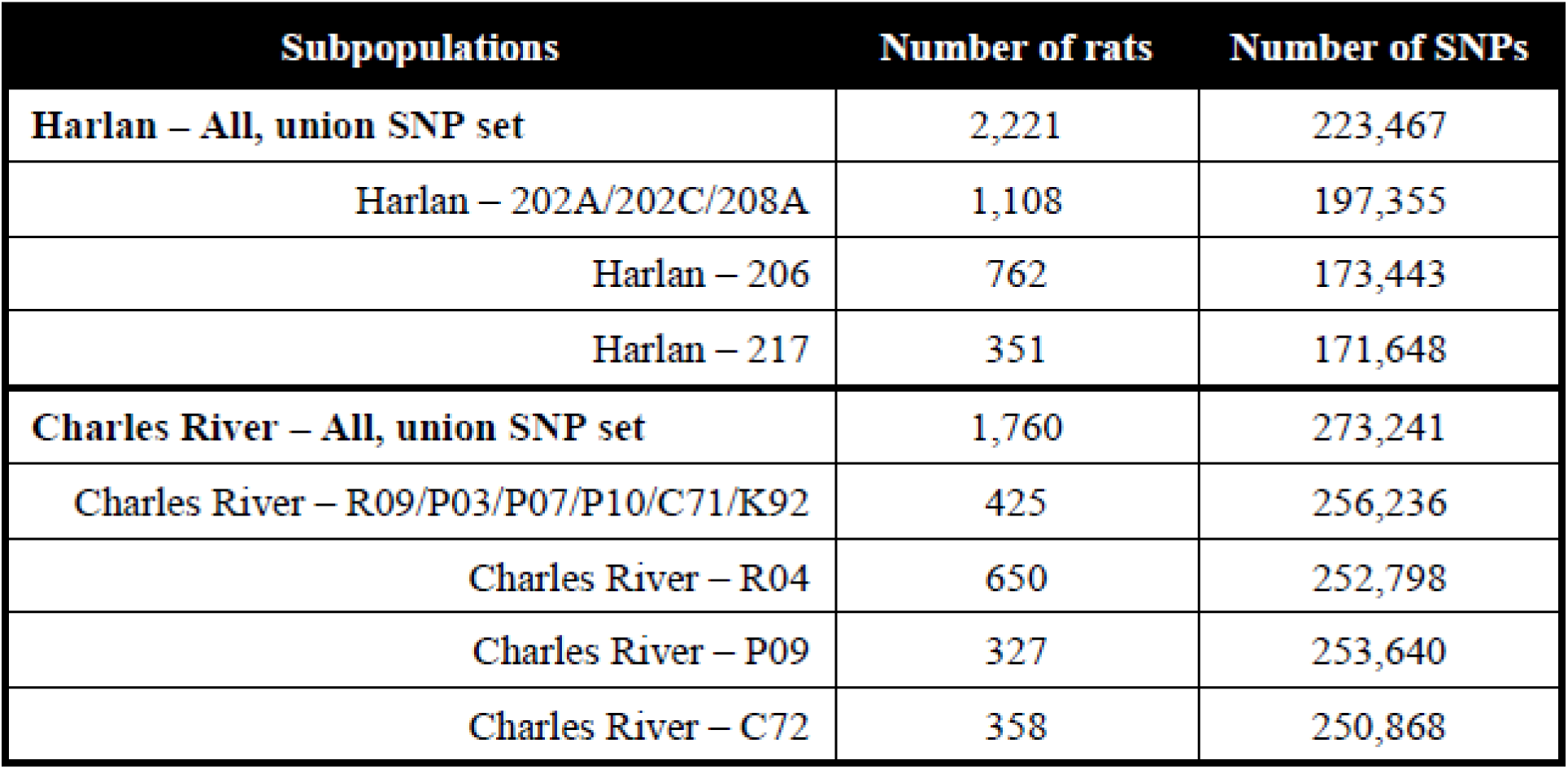
List of filtered variant counts and sample numbers in subpopulations.

| **Subpopulations** | **Number of rats** | **Number of SNPs** |
| --- | --- | --- |
| **Harlan – All, union SNP set** | 2,221 | 223,467 |
| Harlan – 202A/202C/208A | 1,108 | 197,355 |
| Harlan – 206 | 762 | 173,443 |
| Harlan – 217 | 351 | 171,648 |
| **Charles River – All, union SNP set** | 1,760 | 273,241 |
| Charles River – R09/P03/P07/P10/C71/K92 | 425 | 256,236 |
| Charles River – R04 | 650 | 252,798 |
| Charles River – P09 | 327 | 253,640 |
| Charles River – C72 | 358 | 250,868 |

**S4 Fig.**
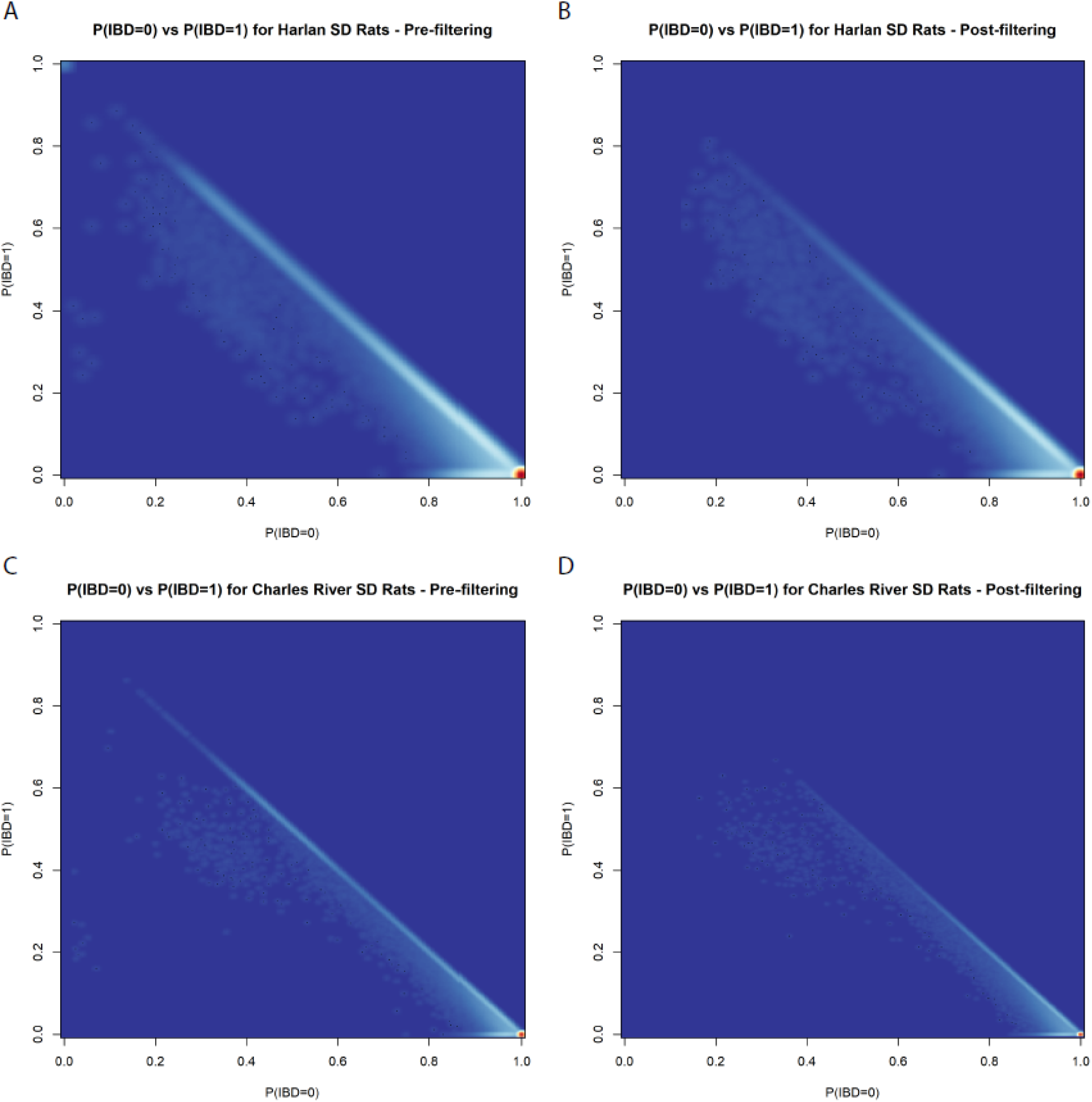
Heatmaps of pairwise identity-by-descent pre- and post-filtering. Panels A and C show pre-filtering values of P(IBD) = 0 plotted against P(IBD) = 1 for Harlan and Charles River. Unrelated samples cluster in the lower right corner. Samples along the diagonal have 2^nd^ and 3^rd^ degree levels of relatedness, while those clustering around (0.5,0.25) are full-siblings. Panels B and D demonstrate that the sample filtering steps removed several spurious relations from the sample.

**S5 Table.** List of sample filtering criteria and number of samples removed.

|  | **Number of Samples** |  |
| --- | --- | --- |
| Total sequenced by ddGBS + WGS | 4,625 (17 WGS only) |  |
| Low read count in FASTQ (< 4 million reads)<br>**performed prior to variant calling | 316 |  |
| Females samples (only females in sample set) | 77 |  |
| Purchased from Taconic | 4 |  |
| Lack of phenotype information | 10 |  |
| Poor clustering in principal component analysis | 54 |  |
| Putative sample mix-up | 18 |  |
| Duplicate sample | 7 |  |
| Inflated/deflated heterozygosity | 34 |  |
| $\geq 30$ samples with $\hat{\pi} \geq 0.1875$ in sample | 12 | |
| $\hat{\pi} > 0.6$ with another sample | 32 | |
| **Final Sample Set (Harlan + Charles River)** | **4,061** |  |
|  | **CR – 1,780** | **Har – 2,281** |

**S6 Table. Heritability estimates in Charles River, Harlan, and their respective subgroups for 55 mapped PavCA metrics.**

**S5 Fig.**
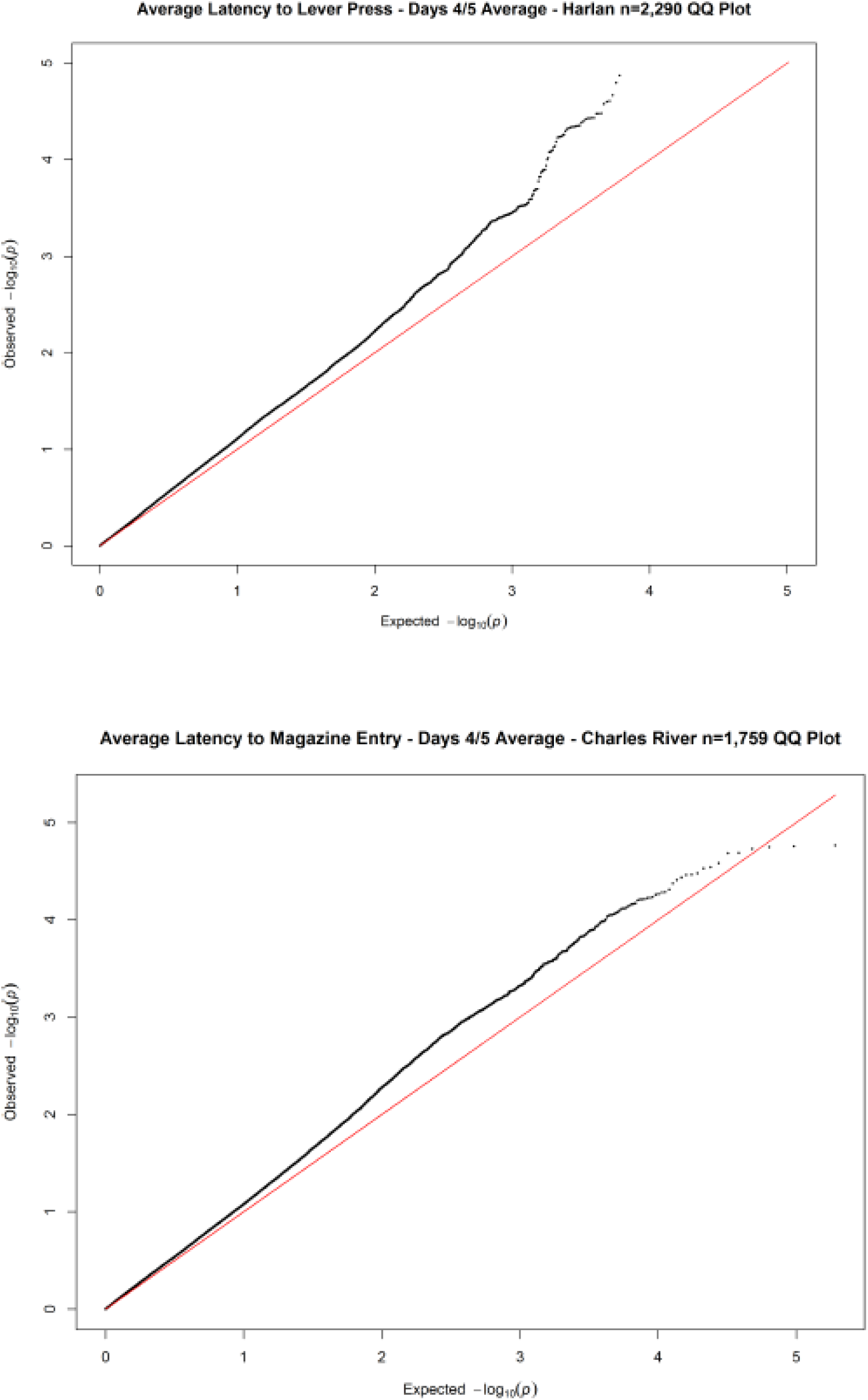
QQ-plot inflation observed in vendor-level GWAS.

**S1 File. Table of all significant PavCA GWAS hits.** For each significant GWAS association, the table contains the position, allele, allele frequency, PavCA metric, sample size, direction of effect, meta-analysis p-value, percent variance explained, effect size, and standard error. In addition, the values for all the independent GWAS that went into the meta-analyses are provided.

**S2 File. Stacked subgroup Manhattan plots for all analyzed metrics/traits.** Each page contains vertically stacked Manhattan plots with the GWAS results for all PavCA metrics in all seven sample subgroups. The plots are in the following order from top to bottom: Harlan 202A/202C/208A (115k SNPs), Harlan 206 (102k SNPs), Harlan 217 (115k SNPs), Charles River R09/P03/P07/P10 (221k SNPs), Charles River R04 (214k SNPs), Charles River P09 (223k SNPs), and Charles River C72 (222k SNPs).

**S3 File. Q-Q plots for all analyzed metrics/traits.** Each page contains Q-Q plots for the GWAS of a given day/metric for the seven subgroups.

**S4 File. Stacked meta-analyses Manhattan plots for all analyzed metrics/traits.** Each page contains vertically stacked Manhattan plots with the GWAS results for all PavCA metrics in all three meta-analysis: Charles River plus Harlan (64k), Charles River only (198k SNPs), and Harlan only (83k SNPs).

**S5 File. Q-Q plots for all metrics/trait meta-analyses.** Each page contains Q-Q plots for the meta-analyses of the GWAS results for a given day/metric across (1) all seven subgroups, (2) Charles River subgroups, and (3) Harlan subgroups.

**S2 Text. Sequences for 48 barcoded adapters, Y-adapter, and PCR primers.**

**S8 Table.** List of all PavCA metrics collected on SD rats. There are 11 total metrics across 5 days of training. The first 7 metrics are direct measurements made during the training periods. The following 3 metrics are calculated from the base measurements. The final metric, PavCA index score, is a composite score from the previous 3 metrics.

| Day 1<br>25 8-second trials | Day 2<br>25 8-second trials | Day 3<br>25 8-second trials | Day 4<br>25 8-second trials | Day 5<br>25 8-second trials | Data Type |
| --- | --- | --- | --- | --- | --- |
| Lever Presses | Lever Presses | Lever Presses | Lever Presses | Lever Presses | Count<br>(summed over trials) |
| Magazine Entries CS | Magazine Entries CS | Magazine Entries CS | Magazine Entries CS | Magazine Entries CS | Count<br>(summed over trials) |
| Magazine Entries NCS | Magazine Entries NCS | Magazine Entries NCS | Magazine Entries NCS | Magazine Entries NCS | Count<br>(summed over trials) |
| Latency to Lever Press | Latency to Lever Press | Latency to Lever Press | Latency to Lever Press | Latency to Lever Press | Continuous [0,8] secs<br>(averaged over trials) |
| Latency to Magazine Entry | Latency to Magazine Entry | Latency to Magazine Entry | Latency to Magazine Entry | Latency to Magazine Entry | Continuous [0,8] secs<br>(averaged over trials) |
| Probability of Lever Press | Probability of Lever Press | Probability of Lever Press | Probability of Lever Press | Probability of Lever Press | Proportion [0-1]<br>(0/25 to 25/25 trials) |
| Probability of Mag Entry | Probability of Mag Entry | Probability of Mag Entry | Probability of Mag Entry | Probability of Mag Entry | Proportion [0-1]<br>(0/25 to 25/25 trials) |
| Response Bias | Response Bias | Response Bias | Response Bias | Response Bias | Continuous [-1,1] |
| Latency Score | Latency Score | Latency Score | Latency Score | Latency Score | Continuous [-1,1] |
| Probability Difference | Probability Difference | Probability Difference | Probability Difference | Probability Difference | Continuous [-1,1] |
| PavCA Index Score | PavCA Index Score | PavCA Index Score | PavCA Index Score | PavCA Index Score | Continuous [-1,1] |

**S9 Table.** List of covariates used for the GWAS LMMs for Harlan and Charles River. The first three covariates were used in both Charles River and Harlan analyses. The remaining covariates were unique to each population.

| Har 202A | Har 206 | Har 217 | CR R04 | CR R09 | CR P09 | CR C72 |
| --- | --- | --- | --- | --- | --- | --- |
| Age of rat in days at start of training (continuous integer) |  |  |  |  |  |  |
| Housing condition (binary – single or multiple rats per chamber) |  |  |  |  |  |  |
| Light cycle (binary – standard or reverse 12h lighting) |  |  |  |  |  |  |
| AAK | CJF | AK | AAK | CJF | AK | AAK |
| BFS.AA | JC.JDM | BFS | CJF | CJF.JDM | BFS.AA | BFS |
| CJF | JJ | BTS | CJF.JDM | EGO | EGO | BFS.AA |
| CJF.JDM | KP | CJF | JJ |  | LMY | CJF.JDM |
| EGO |  | CJF.JDM |  |  | MLR.LMF |  |
| KP |  | JC.JDM |  |  |  |  |
| LMY |  | TW.KP |  |  |  |  |

**S6 Fig.**
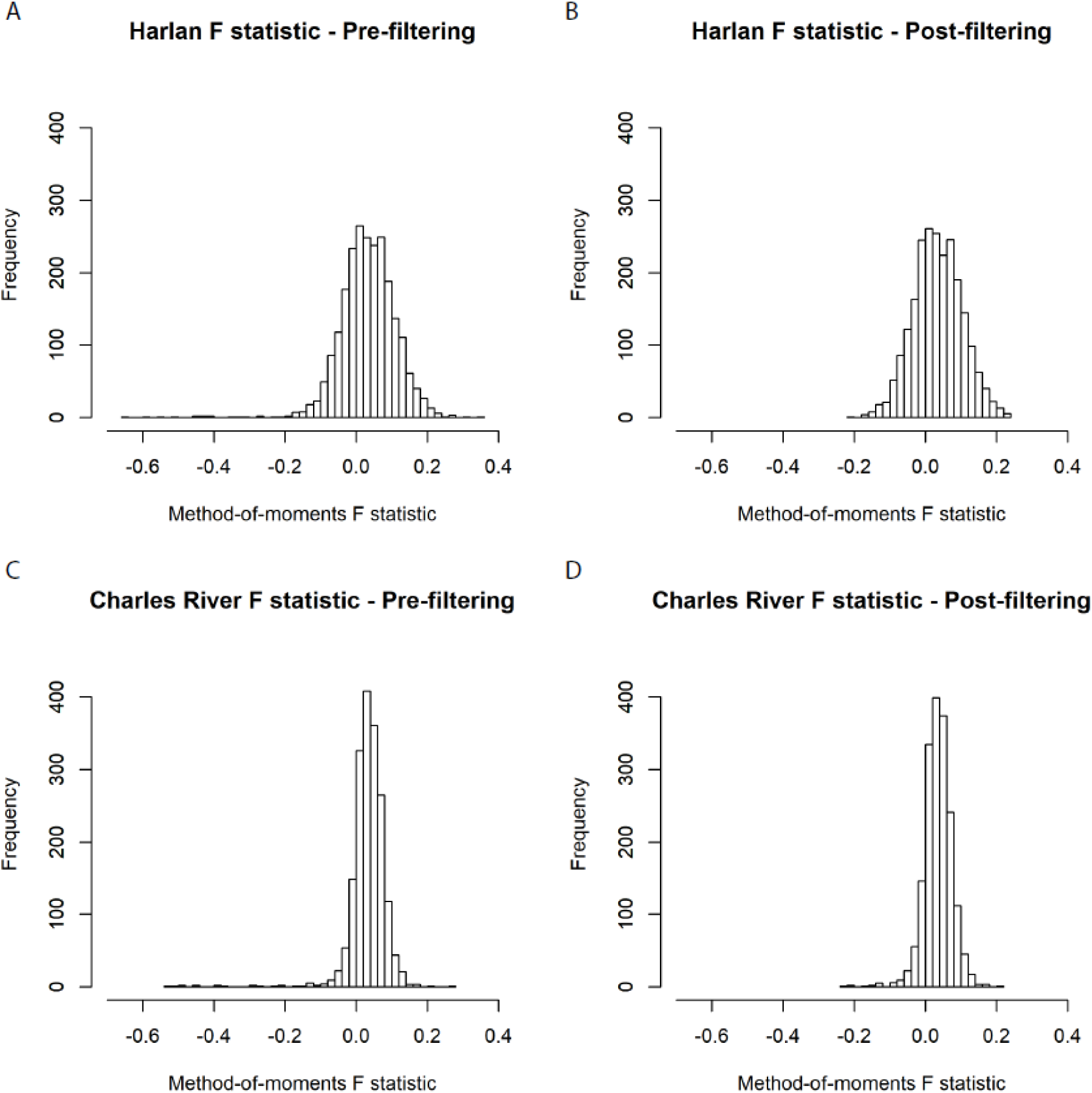
Pre- and post-filtering distributions of heterozygosity for Harlan and Charles River. Panels A and C show pre-filtering distributions of heterozygosity in Harlan and Charles River, as measured by the method-of-moments F coefficient. Panels B and D show the same distributions post-filtering. A value above 0 indicates a deflation of heterozygosity, whereas a value below 0 would be an inflation.

**S6 File. Spreadsheet with detailed sample sequencing and phenotype information.**

