## Supplementary material for "Genetic characterization of outbred Sprague Dawley rats and utility for genome-wide association studies": S1_Text - ddGBS Protocol

### Double Digest GBS Library Prep 08/23/2018

### Plate Setups

Prior to beginning the protocol, be sure to quantify all DNA samples. Because the position of the adapters is fixed in the plate (see below), you should randomize the order of your samples. To do this, list all of the sample IDs in column A of an excel spreadsheet. In column B, use the function **=rand()** to generate a random number for each entry in column A. Then sort the entries from highest to lowest using column B. Now create a third column listing the adapter wells A1-D12 so that each of your samples is assigned to an adapter. Organize your sample tubes correspondingly before plating out the DNA.

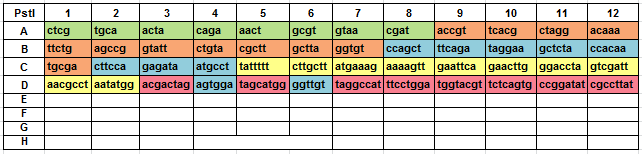

### Day 1: Plate out Sample DNA and adapters

Add samples and adapters to plates. Print out the sample-plating design and mark the samples off as you add them. Remember to order the sample tubes according to their plating position.

(**NOTE:** If you are starting a set of libraries for multiple plates of sequencing, choose one sample that will be present on all plates in at least 1 library, and additionally, pick about 2-3% of your total samples to be sequenced to have replicates across plates or within the same plate. This is meant to help with data analysis.)

**1. Plate out DNA samples.** 1µg of DNA per sample is needed: if samples are at various concentrations you will need to adjust accordingly. It does not matter whether you choose to add DNA or adapters to the plate first.

**2. Transfer specified amount of PstI working adapter stock** **to each plate (~0.2pmol). Be sure to mix the adapter plate well first by pipetting up and down 15-20 times.** Ensure that each adapter is matched to the correct corresponding DNA sample according to plate schematic. (NOTE: The average molecular weight per base pair of an adapter in the plate is 307.5 g/mol. Using this value and the concentration (ng/µL) of the adapters in the plate (as quantified by PicoGreen) you can calculate the volume of adapter stock necessary to obtain 0.2pmol per well.)

**3. Vortex and transfer 4 ul of (1.0µM ~ 4 picomoles) NlaIII working y-adapter stock (Adapter 2)** **to each well.** Use a strip tube and a multichannel to do this more quickly. If there are only the 10uM or 100uM stocks available, make the necessary dilution to 1uM to perform this step.

**4. Cover with Airpore tape and briefly spin down the plate.**

**5. Set at 37^o^C in thermocycler until liquid has completely evaporated**. This can take up to a day; it’s best to do this overnight. You may also leave the plate in a standard incubator; remember to adjust the temperature.

### Day 2: PstI Digestion and Ligation

Ligation should proceed immediately after digestion, so if you can only do one plate at a time, each plate should be prepped through the ligation before starting the digest for the next plate.

The following digestion, ligation and PCR master mixes assume a sample number of 96. Adjust the reagents accordingly for different sample sizes. When making the mixes, it is best to make enough for all of your samples plus at least 5. Thus, for the mixes below, the reagent volumes are calculated using a factor of 105 and not 96.

**1. Make PstI Digestion Master Mix**. Prepare **on ice** in a 5mL microcentrifuge tube. Pre-thaw NEB buffer and ensure that there is no precipitate present. Mix by vortexing.

| Reagent | Volume/Rxn |  | Total Rxns | **Volume in MasterMix** | Total Volume for **13 Rxns** | Total Volume for **52 rxns** |
| --- | --- | --- | --- | --- | --- | --- |
| **dH20** | 11.0 uL | **x** | 102 | **1122 uL** | 143 uL | 572 uL |
| **CutSmart Buffer** | 7.5 uL | **x** | 102 | **765 uL** | 97.5 uL | 390 uL |
| **PstI-HF** | 0.5 uL | **x** | 102 | **51 uL** | 6.5 uL | 26 uL |
| **NlaIII** | 1.0 uL | **x** | 102 | **102 uL** | 13 uL | 52 uL |
| **TOTAL** | 20.0 uL | **x** | 102 | **2040 uL** | 260 uL | 1040 uL |

**2. Add PstI Digestion Master Mix.** Transfer master mix to a multi-channel reservoir. Use a multi-channel pipettor to transfer **20** **µL** of the mix to each well.

**3. Cover with adhesive PCR film and vortex gently before spinning down.**

**4. Incubate at 37 degrees in the thermocycler for 2 hrs; hold at 4 degrees until ligation.**

**5. Transfer plate to ice.**

**6. Prepare Ligation Master Mix.** Prepare mix in 2mL Eppendorf tube (or 15-50mL Falcon tube depending on the sample number and final mix volume) **on ice.**

| Reagent | Volume/Rxn |  | Total Rxns | **Volume in MasterMix** | Total  Volume for **13 Rxns** | Total Volume for **52 rxns** |
| --- | --- | --- | --- | --- | --- | --- |
| **dH20** | 23.4 uL | **x** | 102 | 2389 uL | 304.2 uL | 1216.8 uL |
| **10X Ligation Buffer** | 5.0 uL | **x** | 102 | 510 uL | 65.0 uL | 260 uL |
| **T4 DNA Ligase** | 1.6 uL | **x** | 102 | 163.2 uL | 20.8 uL | 83.2 uL |
| **TOTAL** | 30.0 uL | **x** | 102 | 3,062 uL | 390 uL | 1560 uL |

7**. Add mix to multichannel reservoir** (or for a small sample size, use a 12 well strip) while **on ice**. Add **30 uL** Ligation Mix to each well of the reaction plate using a multi-channel pipettor.

**8. Cover with adhesive PCR film, vortex, and spin down briefly.**

**9. Program the thermocycler and incubate** at 16 degrees for 60 minutes (optimum for T4 ligase) and then 80 degrees for 30 minutes (kills Pst1). Hold at 4 degrees.

**10. Transfer plate to -20 freezer** (if have to stop at this step for the day—most likely will have to) Can also stop after the filtering step or the PicoGreen step.

### Day 3: Qiagen MinElute 96-Well Purification

**1. Thaw samples from Day 1 and spin down (if necessary).**

**2. Prepare the vacuum manifold.** Put a waste tray inside the base and place the MinElute 96 UF PCR purification plate on top of the vacuum manifold.

**3. Pipet the PCR samples into the MinElute plate**, marking which wells have been used. Volume in the reaction wells is roughly 50uL. It is important not to get any liquid on the sides of the purification plate wells; add directly to center of membrane*.*

**4. Apply vacuum.** Maintain at -800 mbar for 5 minutes or until the wells are completely dry. Switch off vacuum and disconnect tubing to break the seal.

**5. Wash with 50 uL dH20.** Add 50 uL deionized water to each well, apply vacuum, and maintain at -800 mbar or until wells are completely dry. Switch off vacuum and disconnect tubing to break the seal.

**6. Remove the plate from the vacuum manifold.** Carefully tap the plate on a stack of clean Kimwipes to remove any liquid. If a large amount of liquid remains, it needs to be vacuum treated longer.

**7. Elute DNA from the membrane.** Add 27 uL Elution Buffer to each well, then switch pipette to 25uL as to prevent air bubbles from accumulating during mixing. Mix by pipetting up and down 20+ times or more. For easier recovery of eluate, the plate can be placed at an angle on the vacuum manifold. Aspirate the full 25uL and put in a new plate or tubes.

**8. Transfer samples to a new PCR plate.** Check the wells of the purification plate to make sure no DNA was left behind.

**Quantification with PicoGreen**

**Notes before starting:**

- Everything below is listed in 96-well format and has 200ul final volumes
- **Bring Quanit-iT PicoGreen to room temperature before opening the vial**
- Use molecular biology-grade water (DNase-free) for any necessary dilutions
- PicoGreen is light-sensitive, so keep everything containing PicoGreen protected from light using aluminum.
- PICOGREEN SHOULD BE DONE IN DUPLICATES FOR THIS STEP, WITH EACH SAMPLE QUANITIFIED TWICE. We do this because it’s such a sensitive measurement with small volumes that there can be a lot of variability between measurements. You will use an average moving forward.

**Low-Range Readings (40 pg/ul to 50 ng/ul)**

1. **Prepare Tubes for "standard curve"**
   a. Thaw DNA standard provided with PicoGreen kit (100 ug/mL = 100 ng/ul)
   b. Label 12 1.5 mL Eppendorf tubes 1-12
   c. Follow the below instructions for filling the 12 tubes with 1xTE and for transferring volumes in the subsequent steps below.

| **Low Range Reading Table** | | | | | | | |
| --- | --- | --- | --- | --- | --- | --- | --- |
|  | **One Plate** | | **Two Plates** | | **Three Plates** | |  |
| **Tube** | **1xTE Vol** | **n-1 Tube Vol** | **1xTE Vol** | **n-1 Tube Vol** | **1xTE Vol** | **n-1 Tube Vol** | ***[DNA]ng/uL*** |
| **1** | 597 | 3 | 995 | 5 | 1393 | 7 | *50* |
| **2** | 360 | 240 | 600 | 400 | 840 | 560 | *20* |
| **3** | 300 | 300 | 500 | 500 | 700 | 700 | *10* |
| **4** | 300 | 300 | 500 | 500 | 700 | 700 | *5* |
| **5** | 300 | 300 | 500 | 500 | 700 | 700 | *2.5* |
| **6** | 300 | 300 | 500 | 500 | 700 | 700 | *1.25* |
| **7** | 300 | 300 | 500 | 500 | 700 | 700 | *0.625* |
| **8** | 300 | 300 | 500 | 500 | 700 | 700 | *0.3125* |
| **9** | 300 | 300 | 500 | 500 | 700 | 700 | *0.15625* |
| **10** | 300 | 300 | 500 | 500 | 700 | 700 | *0.078125* |
| **11** | 300 | 300 | 500 | 500 | 700 | 700 | *0.0390625* |
| **12** | 300 | 0 | 500 | 0 | 700 | 0 | *0* |

d. Set aside Tube 12
e. Add *3/5/or7* ul of 100ng/ul standard DNA (200x dilution = 0.5 ng/ul) to Tube 1 and vortex 15s
f. To Tube 2, add 240/400/or560 from Tube 1 (2.5x dilution = 0.2 ng/ul) and vortex 15 sec.
g. Transfer 300/500/or700 ul from Tube 2 to Tube 3 and vortex 15 sec.
h. Continue transferring 300ul and vortexing for the dilutions for Tubes 4-11. Do NOT add DNA to Tube 12.
i. Final DNA concentrations are in the Low-Range Table below

2. Transfer 100ul of each DNA standard Dilution to the appropriate well (usually A1-A12) in Grenier flat clear-bottomed 96-well plate.
3. Transfer 99ul of 1x TE to the remaining open wells of the fluorometer plates
4. Transfer 1.0ul of the sample DNA to the appropriate wells
5. **Prepare 1:200 dilution PicoGreen**
   a. Add *Y*/2 uL PicoGreen to [(100**Y*) - *Y*/2] uL of 1xTE and mix well
   b. *Y* = number of wells that will be used for the fluorometer - include "standard curve" wells and a few extra. Full plate ~100 wells
6. Add 100ul of 1:200 dilution PicoGreen to each well
7. Pipet up and down 10x with 100ul multichannel pipetor
8. Cover to protect from light and incubate at RT for 5 minutes.
9. Measure fluorescence on Tecan
   a. Use the standard template provided by the iControl program for PicoGreen dsDNA quantification and highlight the active wells
10. Analyze/calculate DNA concentrations using Excel
    a. Need to verify the R^2 value of the standard curve is at least 0.98. This can be done in excel.
       i. May want to use standard curve readings that are close to the range of the samples (ex: for low-concentrations samples, don't use wells A1-A4 for the standard curve)
    b. To get the concentrations, use the equation for the line provided by Excel from graphing the standard (concentrations from the final column above) against the PicoGreen readings for the standard. Then for each measurement that’s not part of the standard, subtract the y-intercept and divide by the slope to get the concentration in ng/uL.
    c. Average the duplicates/triplicates of each well for the final concentration.

### Pool Samples to Make Library

The goal for this step is normalize the concentration of each sample in the library. It is sufficient to only normalize the quantity in grams from each sample, as it is necessary to concentrate the sample using the vacuum plate after this step. Use the PicoGreen results for this step.

**1. Find the sample with the lowest concentration in each planned pool and multiply by 22uL to get total ng of smallest sample in each pool.** We estimate that 22uL is about the most you can reliably expect to recover from each well of the vacuum plate, so we use it as the maximum volume for the lowest concentration sample. This means that 22uL of this sample will move into your final pool.

**Volume to add per sample =** *([Minimum sample Conc] * 22uL) / [Current sample Conc]*

**2. For the remainder of the samples in each pool, divide the total ng of the lowest concentration sample from the last step by the concentration of the current sample, resulting in the volume of that sample to add to the final pool.**

**3. Combine calculated volumes in a single 1.5mL microcentrifuge tube per library (12/24/48)**

### Concentrate Library for Size Selection

**1.** **Sum all volumes added to final pool and divide final volume by 2 (for 12 samples/pool) or 4 (for 24 samples/pool, etc.) to get volume to add to each well on the vacuum plate.**

**2.** **Add library to vacuum wells, turn on vacuum to -800 mbar, and wait until wells are completely dry.**

**3.** Turn off the vacuum and detach the hose. **Re-suspend the DNA in 20-25uL of elution buffer, pipetting up and down 20+ times to recover it all.** With 12 samples per pool, re-suspending in 25uL of elution buffer in each well will result in slightly less than 50uL to move forward to the next step (which is more than enough, so you can also elute with 20uL of elution buffer per well instead). *With 24 samples per pool, you will need to perform 2 rounds of vacuum concentration to get a small final volume, using 4 wells and then 2 wells.* And so on for 48.

**4.** When finished, quantify the library using the Nanodrop and record the concentration and quality.

Pippen Prep works best with <5,000ng DNA. **If total [conc] will be higher, dilute with more EB first.**

**Pippin Prep Size Fractionation**

**Follow Quick Guide for Use of Pippin Prep.** (Make sure to allow the marker L tube to come to room temperature over a 30 minute period before use). 30uL of the library will be used for this step. But no more than 5,000ng. **For instace, if the library is 200ng/uL….5000/200 = 25uL library + 5 EB + 10uL loading mix.** Also, load the elution modules in the Pippin Prep gels with 45uL electrophoresis buffer, rather than 40uL as told.

### PCR Amplification

50ul PCR for each sample, starting with **10ul of post-Pippin product**, there should be enough volume for 4 PCR reactions per library. It is acceptable if the final PCR reaction is a bit short on DNA template from the Pippen Prep. If you are planning on multiplexing more than 48 samples per library for sequencing, you will need to make 2 separate PCR master mixes for 96 samples, 3 for 144, and 4 for 192. Each master mix will use a different PCR primer 2 with a unique multiplexing index that will be included in Read 3 on the Illumina HiSeq machine. We have made 4 such primers with 8 bp index sequences, but more could be used.

**1. Prepare PCR Master Mix**. Use 2mL Eppendorf tube or larger and be sure to prepare **on ice**. Vortex the mix.

| Reagent | Volume/Rxn |  | Total Rxns | Volume in Master Mix | Volume for **5 Rxns** |
| --- | --- | --- | --- | --- | --- |
| **dH20** | 13.0 uL | **x** | 18 uL | 234 uL | 65 uL |
| **NEB 2X Taq Master Mix** | 25.0 uL | **x** | 18 uL | 450 uL | 125 uL |
| **PCR primer 1 (10uM)** | 1.0 uL | **x** | 18 uL | 18 uL | 5 uL |
| **PCR primer 2 (indexed) (10uM)** | 1.0 uL | **x** | 18 uL | 18 uL | 5 uL |
| **TOTAL** | 40.0 uL | **x** | 18 uL | 720 uL |  |

**2. Prepare adequate PCR strip tubes or a PCR plate on ice and at 40uL of the mix to each well**

**3. Add 10ul of the purified post-ligation product to the appropriate wells in the PCR plate**

**4. After transfer is complete, cover plate with adhesive PCR film. Vortex the plate and spin down.** Be sure seal forms over each well.

**5. Run 12-cycle PCR.** This is already in the thermocycler.

| 1. 68 | 5 min |
| --- | --- |
| 2. 95 | 60s |
| 3. 95 | 30s |
| 4. 65 | 30s |
| 5. 68 | 30s |
| 6. Go to 3 11x | (12 total) |
| 7. 68 | 5 min |
| 8. 4 | hold |

**Clean up PCR Reactions**

**1. Follow Invitrogen PureLink Quick PCR Purification Kit instructions**. Use 1 column from kit for every 2 PCR reactions. For instance, add 200uL of Buffer 2 to each PCR reaction, and then combine 2x250uL per spin column when transferring (from the same library of course). This should mean 2 spin columns per library

**2. Elute with 50uL of Elution Buffer and pool all tubes containing the same library.**

**3. Concentrate the pooled library using the vacuum plate. 2** PCR rxns x 50uL = 100uL, use one vacuum well with 100uL each. Turn on vacuum, dry down, re-suspend with 50uL Elution Buffer per well

**Quantify and Bioanalyze the final pooled libraries**

Quantify each library using the NanoDrop and record the concentration & quality. If looking to pool multiple sets of 48 samples, use the concentrations in a similar manner to initial sample pooling and combine in equal quantities based on nanograms, not volume. Continue on to concentrate these libraries with the vacuum manifold to the desired final volume.

Follow Bioanalyzer protocol on card. Use DNA 1,000 chips and reagents. If gel is more than a month or two old, make new gel matrix. You will need to manually integrate over the curve to get the final library concentration for sequencing. The library should show a strong peak over the range you size selected for on the Pippin Prep.

**Supplemental Notes**

**1. Removal of adapter dimers (optional).** Depending on the Bioanalyzer results, it may be necessary to run the final pooled libraries through another round of purification to remove adapter dimers. If peaks around 150bp appear prominently in the Bioanalyzer results, this step is necessary. This step requires AmPure XP beads. Prior to use, ensure the beads are vortexed thoroughly and kept on ice.

1. Transfer entire library volume to .3mL PCR tube

2. To each library, add .9 volumes of XP beads. For example, if the final library volume is 50uL, add 45uL of XP beads to the library tube.

3. Mix the beads and libraries by pipetting.

4. Allow the mix to incubate at RT for 5 minutes.

5. Transfer tubes to magnetic plate; let the beads collect for 5 minutes or until the samples appear clear.

6. Remove and discard supernatant.

7. Remove tubes from magnet and wash the beads in each tube with 100uL nuclease free water. Mix beads in water by pipetting.

8. Transfer tubes to magnetic plate and incubate until liquid appears clear. Remove and discard supernatant.

9. Repeat steps 7-8.

10. Remove tubes from magnet and add 55uL of EB. Mix with beads by pipetting.

11. Incubate at RT for 2 minutes.

12. Transfer tubes to magnetic plate and incubate until liquid appears clear.

13. Transfer supernatant to new 1.5mL eppendorf tube labeled with library name.

14. Save the PCR tubes with the used XP beads in case the elution step was unsuccessful.

15. Run the final libraries through the nanodrop and the Bioanalyzer.

**General Comments**

--The protocol works best with a large number of samples. I typically do 96 at a time, but this is by no means a taxing performance. With help and proper planning, 400 samples or more can be prepped in a week.

--The goal before starting the protocol should be organization of the samples such that when performing the preps you never have to think of the plate design.

--Typically, I plate the samples on a Friday, allowing them to dry down over the weekend. I start the protocol the following week. Once plating is complete, the protocol can be finished in 2 days. However, I usually bioanalyze the samples on the third day so as not to be rushed.

--The limiting reagents and products for the protocol are adapters, T4 DNA ligase, Taq Master Mix, PCR primers, and vacuum purification plates. Prior to beginning the protocol, ensure that these reagents and products are present in sufficient quantity for the number of samples you will be preparing. If any are lacking, it is best to wait until you have obtained enough of the missing products to begin.

--Be sure to save the remaining post-ligation/pre-PCR samples and the PCR samples. The final concentrations of the PCR samples should be around 20ng/uL. You should have enough of the PCR samples to make multiples of each library. The pre-PCR samples should have a remaining volume of 15uL. As such, if an error occurs in the pooling step or the PCR, the libraries can be reattempted from the saved PCR and pre-PCR samples, respectively, without the need to redo the entire procedure. Store these samples in a clearly labeled box in the -20C.
