## Supplementary material for "Genetic characterization of outbred Sprague Dawley rats and utility for genome-wide association studies": S2_File - PavCA GWAS Plots Subpops Days 1-5

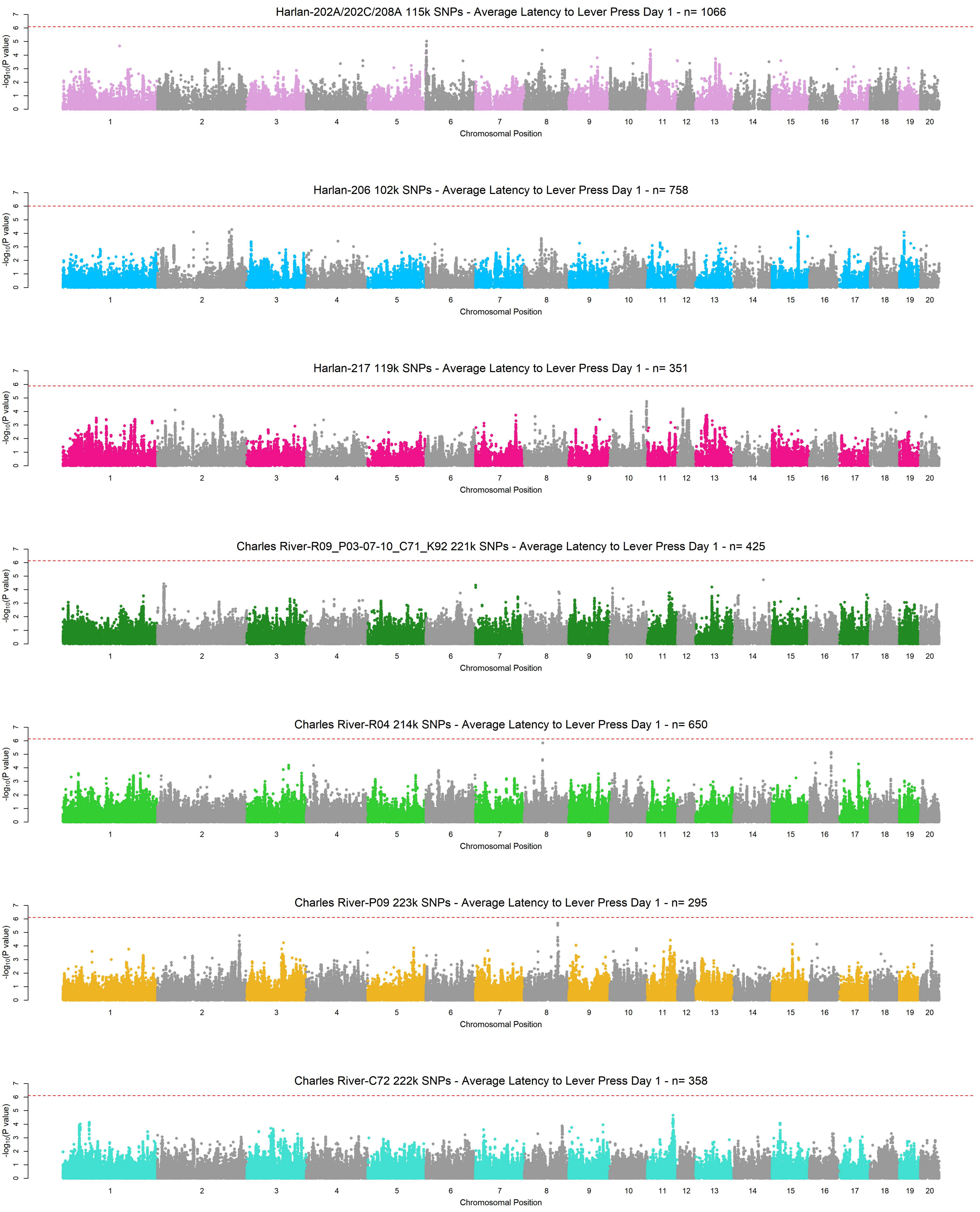
Harlan-202A/202C/208A 115k SNPs - Average Latency to Lever Press Day 1 - n= 1066
Harlan-206 102k SNPs - Average Latency to Lever Press Day 1 - n= 758
Harlan-217 119k SNPs - Average Latency to Lever Press Day 1 - n= 351
Charles River-R09_P03-07-10_C71_K92 221k SNPs - Average Latency to Lever Press Day 1 - n= 425
Charles River-R04 214k SNPs - Average Latency to Lever Press Day 1 - n= 650
Charles River-P09 223k SNPs - Average Latency to Lever Press Day 1 - n= 295
Charles River-C72 222k SNPs - Average Latency to Lever Press Day 1 - n= 358
$-\log_{10}$(P value)
Chromosomal Position
1 2 3 4 5 6 7 8 9 10 11 12 13 14 15 16 17 18 19 20
0 1 2 3 4 5 6 7

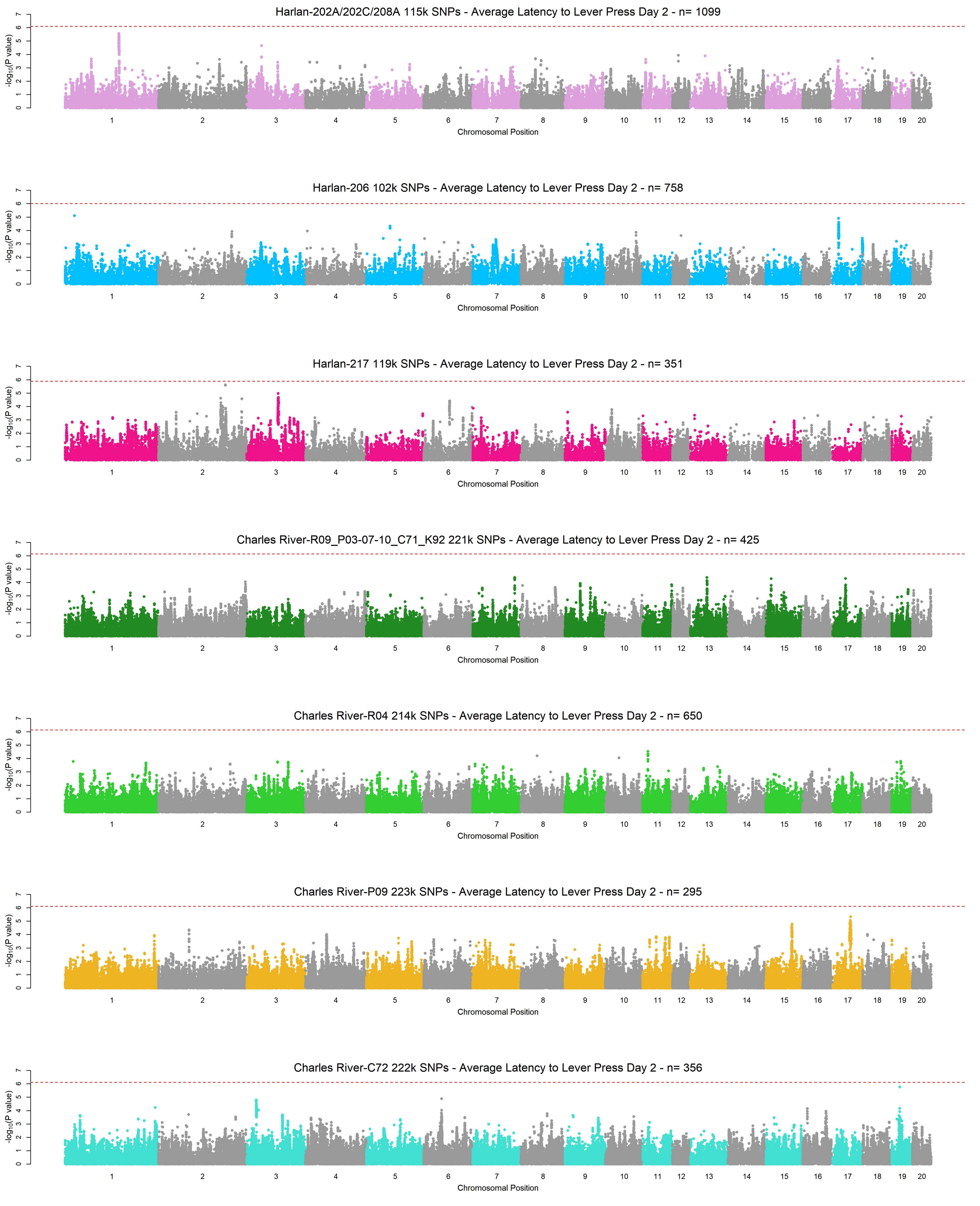
Harlan-202A/202C/208A 115k SNPs - Average Latency to Lever Press Day 2 - n= 1099
Harlan-206 102k SNPs - Average Latency to Lever Press Day 2 - n= 758
Harlan-217 119k SNPs - Average Latency to Lever Press Day 2 - n= 351
Charles River-R09_P03-07-10_C71_K92 221k SNPs - Average Latency to Lever Press Day 2 - n= 425
Charles River-R04 214k SNPs - Average Latency to Lever Press Day 2 - n= 650
Charles River-P09 223k SNPs - Average Latency to Lever Press Day 2 - n= 295
Charles River-C72 222k SNPs - Average Latency to Lever Press Day 2 - n= 356
-log10(P value)
0 1 2 3 4 5 6 7
1 2 3 4 5 6 7 8 9 10 11 12 13 14 15 16 17 18 19 20
Chromosomal Position

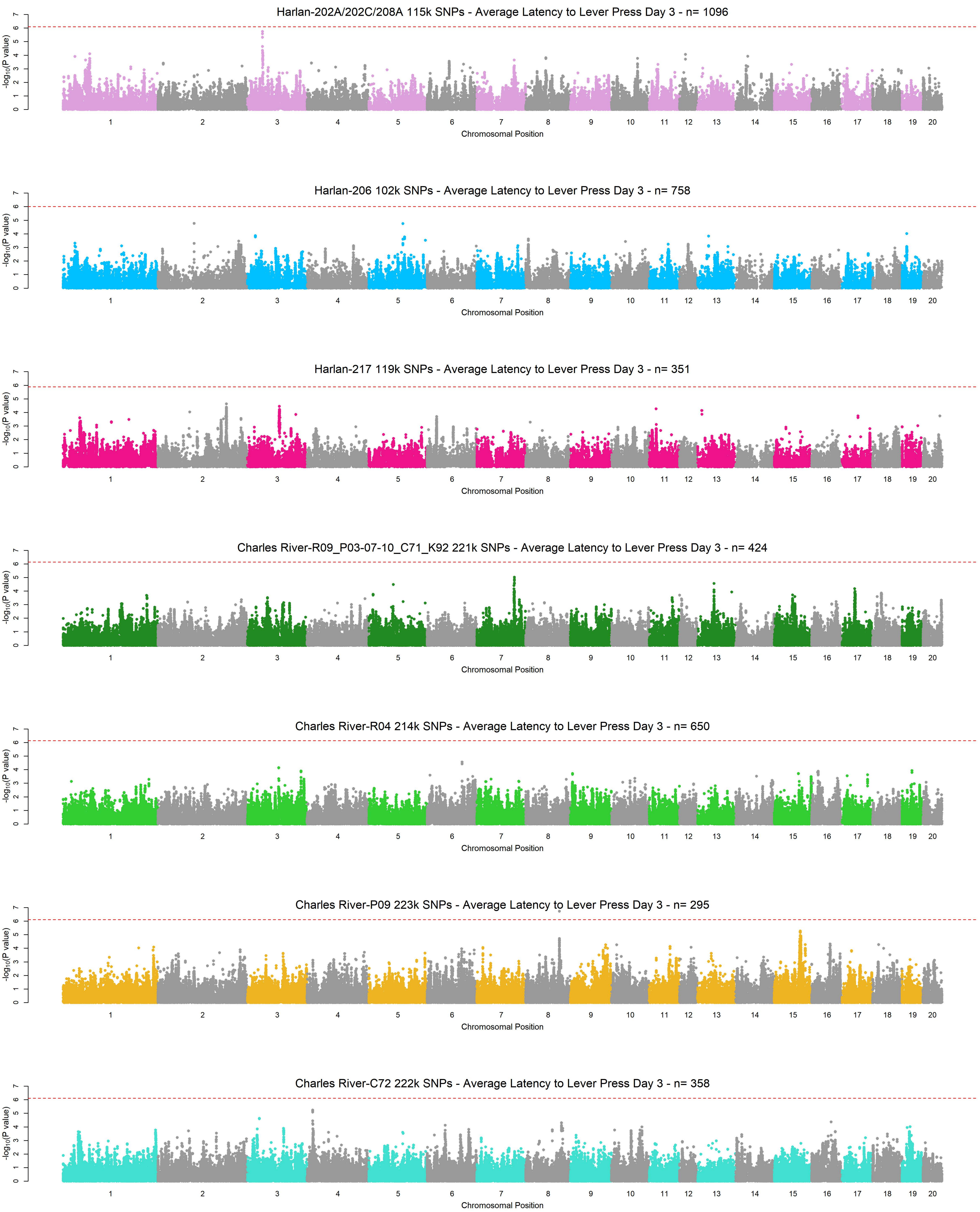
Harlan-202A/202C/208A 115k SNPs - Average Latency to Lever Press Day 3 - n= 1096
Harlan-206 102k SNPs - Average Latency to Lever Press Day 3 - n= 758
Harlan-217 119k SNPs - Average Latency to Lever Press Day 3 - n= 351
Charles River-R09_P03-07-10_C71_K92 221k SNPs - Average Latency to Lever Press Day 3 - n= 424
Charles River-R04 214k SNPs - Average Latency to Lever Press Day 3 - n= 650
Charles River-P09 223k SNPs - Average Latency to Lever Press Day 3 - n= 295
Charles River-C72 222k SNPs - Average Latency to Lever Press Day 3 - n= 358
$-\log_{10}$(P value)
Chromosomal Position
1 2 3 4 5 6 7 8 9 10 11 12 13 14 15 16 17 18 19 20

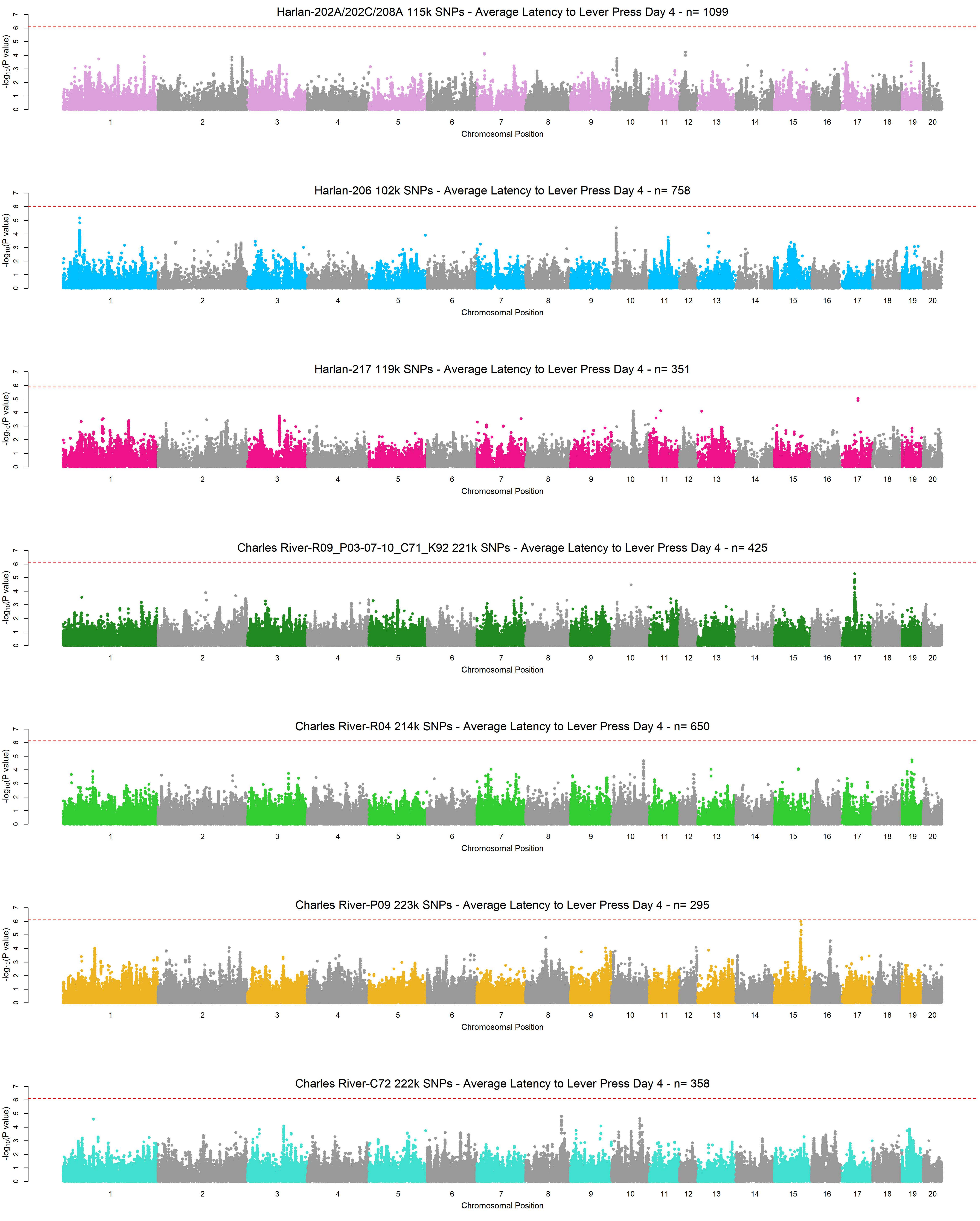
Harlan-202A/202C/208A 115k SNPs - Average Latency to Lever Press Day 4 - n= 1099
Harlan-206 102k SNPs - Average Latency to Lever Press Day 4 - n= 758
Harlan-217 119k SNPs - Average Latency to Lever Press Day 4 - n= 351
Charles River-R09_P03-07-10_C71_K92 221k SNPs - Average Latency to Lever Press Day 4 - n= 425
Charles River-R04 214k SNPs - Average Latency to Lever Press Day 4 - n= 650
Charles River-P09 223k SNPs - Average Latency to Lever Press Day 4 - n= 295
Charles River-C72 222k SNPs - Average Latency to Lever Press Day 4 - n= 358
$-\log_{10}$(P value)
Chromosomal Position

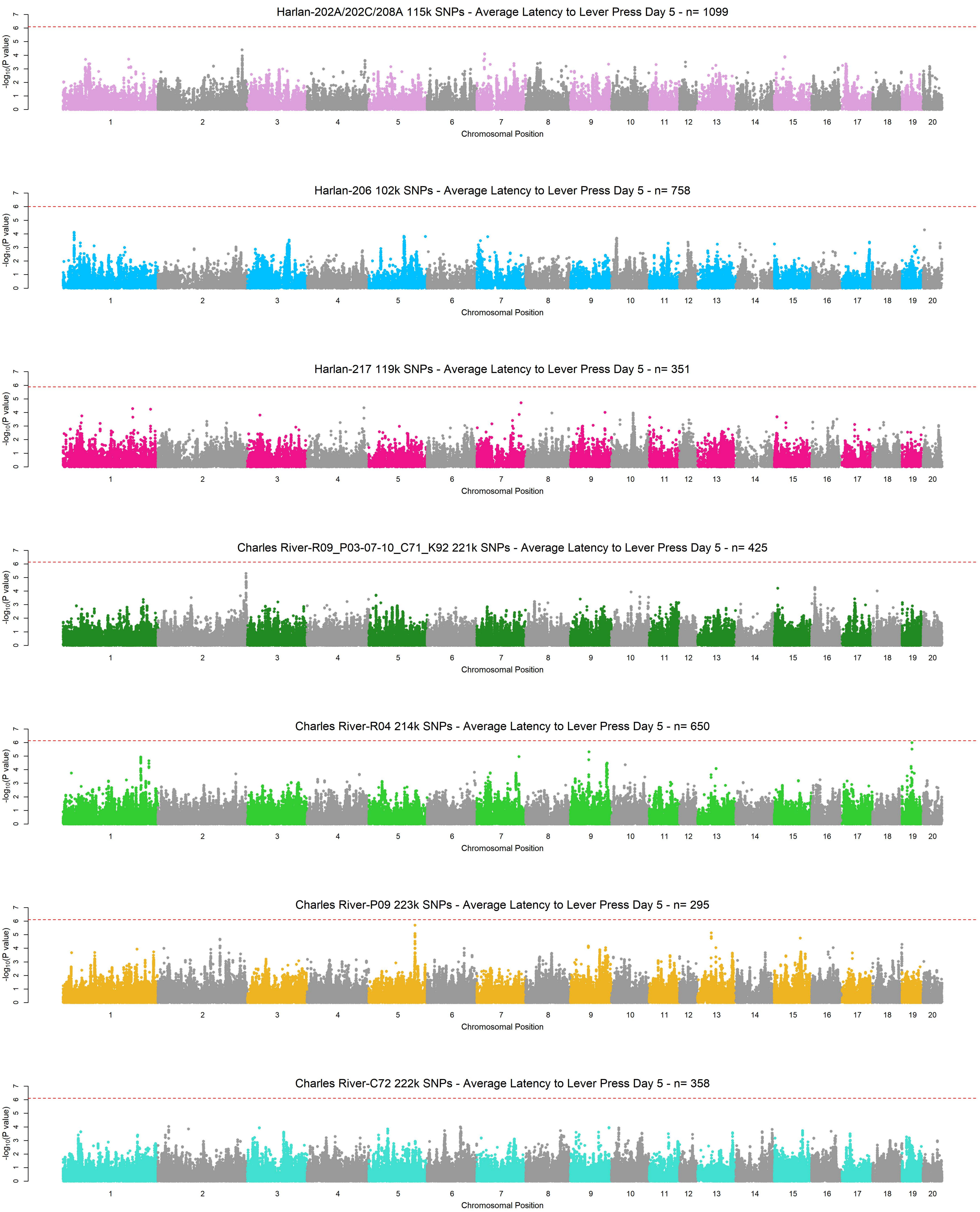
Harlan-202A/202C/208A 115k SNPs - Average Latency to Lever Press Day 5 - n= 1099
Harlan-206 102k SNPs - Average Latency to Lever Press Day 5 - n= 758
Harlan-217 119k SNPs - Average Latency to Lever Press Day 5 - n= 351
Charles River-R09_P03-07-10_C71_K92 221k SNPs - Average Latency to Lever Press Day 5 - n= 425
Charles River-R04 214k SNPs - Average Latency to Lever Press Day 5 - n= 650
Charles River-P09 223k SNPs - Average Latency to Lever Press Day 5 - n= 295
Charles River-C72 222k SNPs - Average Latency to Lever Press Day 5 - n= 358
-log10(P value)
Chromosomal Position
1 2 3 4 5 6 7 8 9 10 11 12 13 14 15 16 17 18 19 20
0 1 2 3 4 5 6 7

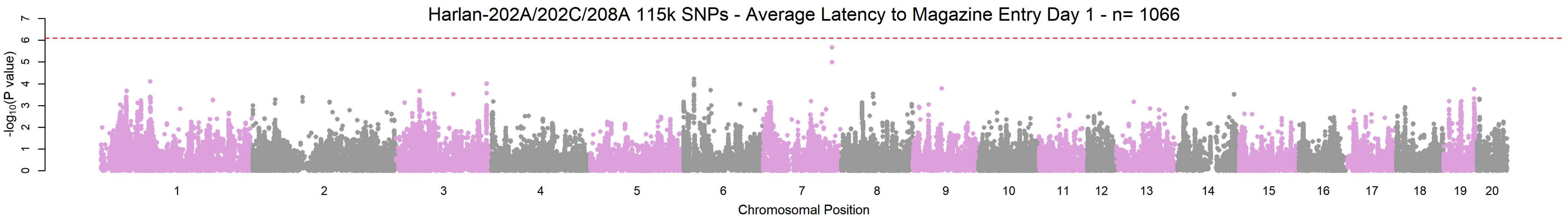
Harlan-202A/202C/208A 115k SNPs - Average Latency to Magazine Entry Day 1 - n= 1066
-log10(P value)
0 1 2 3 4 5 6 7
1 2 3 4 5 6 7 8 9 10 11 12 13 14 15 16 17 18 19 20
Chromosomal Position

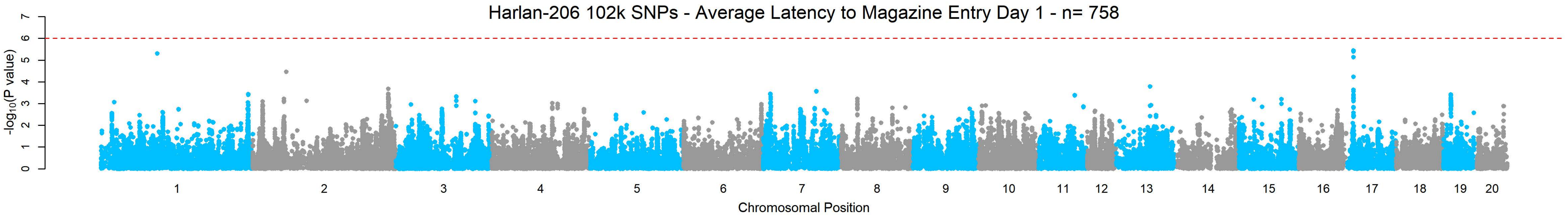
Harlan-206 102k SNPs - Average Latency to Magazine Entry Day 1 - n= 758
-log10(P value)
0 1 2 3 4 5 6 7
1 2 3 4 5 6 7 8 9 10 11 12 13 14 15 16 17 18 19 20
Chromosomal Position

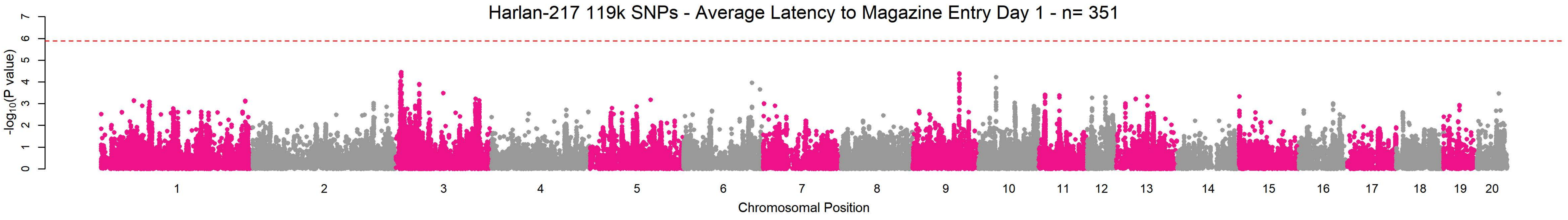
Harlan-217 119k SNPs - Average Latency to Magazine Entry Day 1 - n= 351
-log10(P value)
0 1 2 3 4 5 6 7
1 2 3 4 5 6 7 8 9 10 11 12 13 14 15 16 17 18 19 20
Chromosomal Position

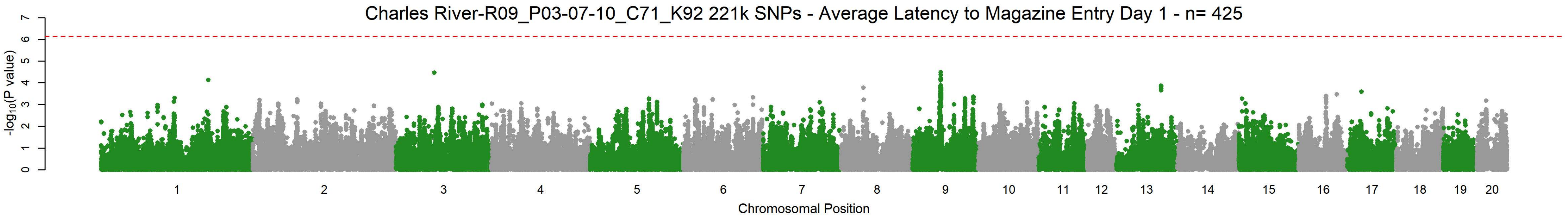
Charles River-R09_P03-07-10_C71_K92 221k SNPs - Average Latency to Magazine Entry Day 1 - n= 425
-log10(P value)
0 1 2 3 4 5 6 7
1 2 3 4 5 6 7 8 9 10 11 12 13 14 15 16 17 18 19 20
Chromosomal Position

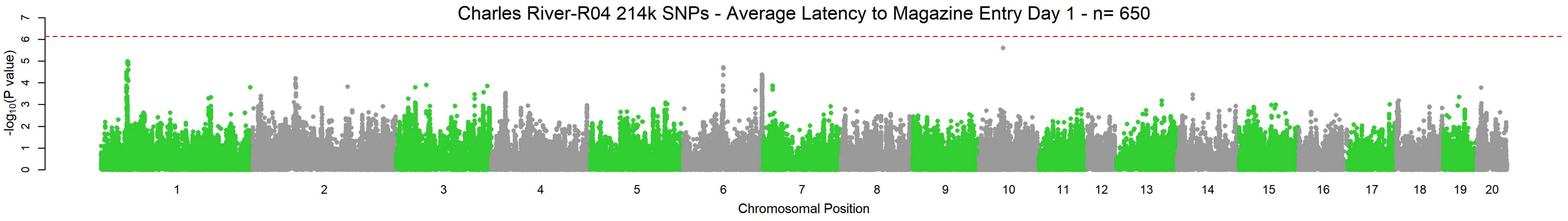
Charles River-R04 214k SNPs - Average Latency to Magazine Entry Day 1 - n= 650
-log10(P value)
0 1 2 3 4 5 6 7
1 2 3 4 5 6 7 8 9 10 11 12 13 14 15 16 17 18 19 20
Chromosomal Position

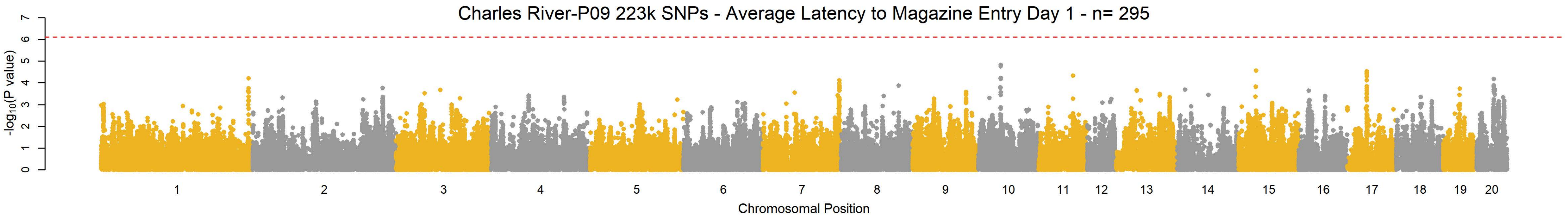
Charles River-P09 223k SNPs - Average Latency to Magazine Entry Day 1 - n= 295
-log10(P value)
0 1 2 3 4 5 6 7
1 2 3 4 5 6 7 8 9 10 11 12 13 14 15 16 17 18 19 20
Chromosomal Position

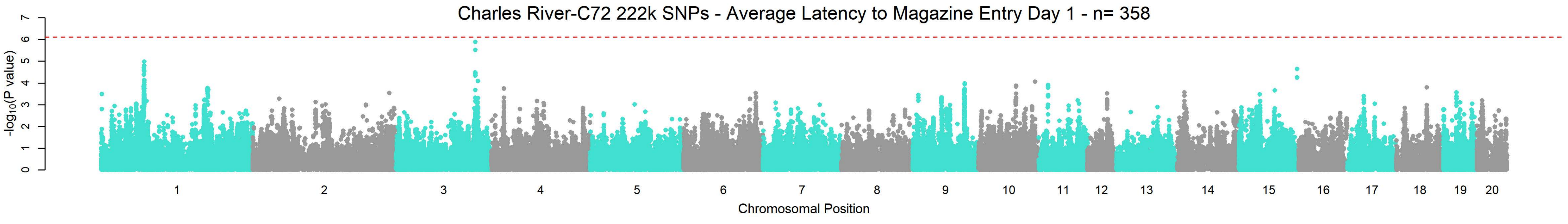
Charles River-C72 222k SNPs - Average Latency to Magazine Entry Day 1 - n= 358
-log10(P value)
0 1 2 3 4 5 6 7
1 2 3 4 5 6 7 8 9 10 11 12 13 14 15 16 17 18 19 20
Chromosomal Position

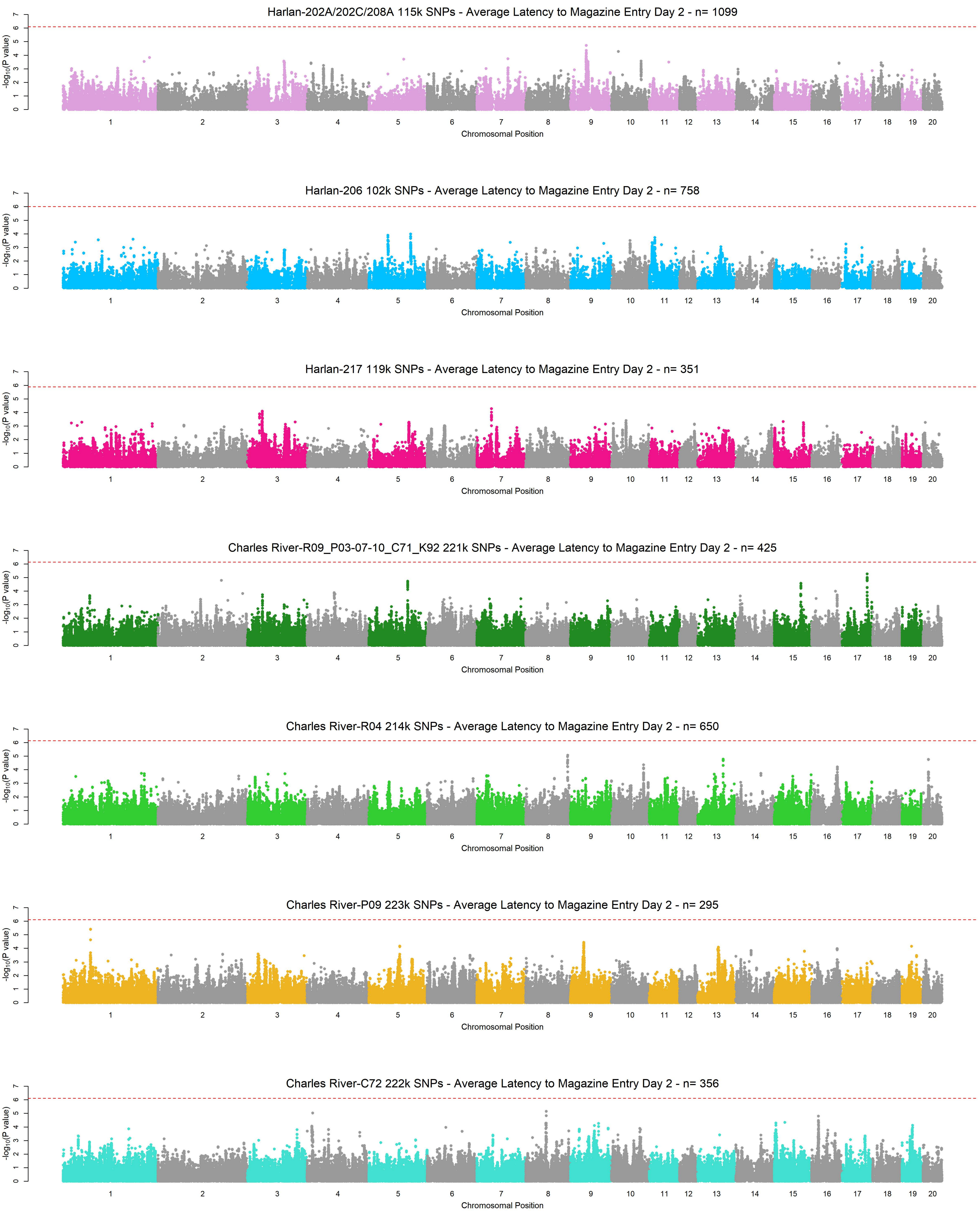
Harlan-202A/202C/208A 115k SNPs - Average Latency to Magazine Entry Day 2 - n= 1099
Harlan-206 102k SNPs - Average Latency to Magazine Entry Day 2 - n= 758
Harlan-217 119k SNPs - Average Latency to Magazine Entry Day 2 - n= 351
Charles River-R09_P03-07-10_C71_K92 221k SNPs - Average Latency to Magazine Entry Day 2 - n= 425
Charles River-R04 214k SNPs - Average Latency to Magazine Entry Day 2 - n= 650
Charles River-P09 223k SNPs - Average Latency to Magazine Entry Day 2 - n= 295
Charles River-C72 222k SNPs - Average Latency to Magazine Entry Day 2 - n= 356
-log10(P value)
Chromosomal Position
1 2 3 4 5 6 7 8 9 10 11 12 13 14 15 16 17 18 19 20

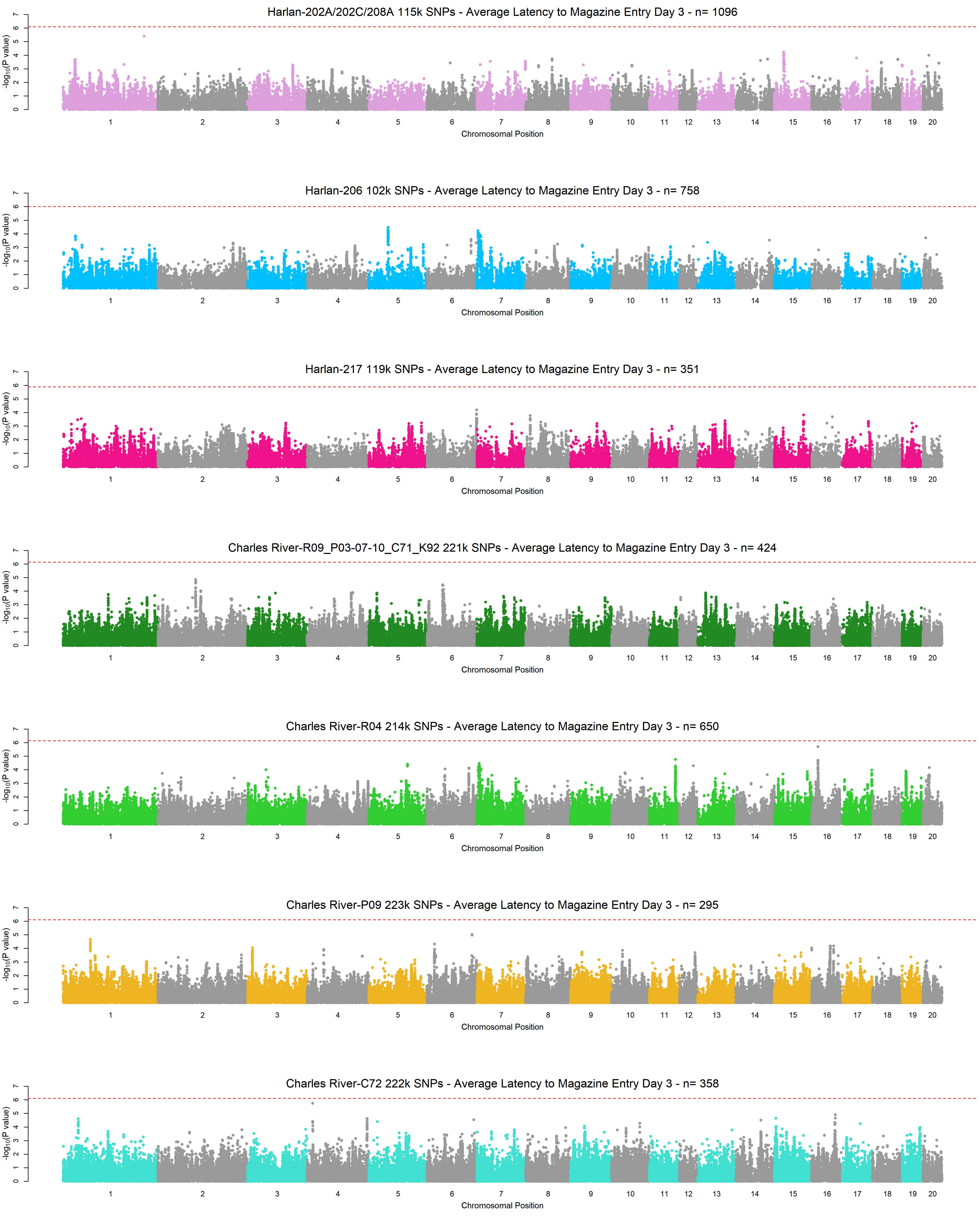
Harlan-202A/202C/208A 115k SNPs - Average Latency to Magazine Entry Day 3 - n= 1096
Harlan-206 102k SNPs - Average Latency to Magazine Entry Day 3 - n= 758
Harlan-217 119k SNPs - Average Latency to Magazine Entry Day 3 - n= 351
Charles River-R09_P03-07-10_C71_K92 221k SNPs - Average Latency to Magazine Entry Day 3 - n= 424
Charles River-R04 214k SNPs - Average Latency to Magazine Entry Day 3 - n= 650
Charles River-P09 223k SNPs - Average Latency to Magazine Entry Day 3 - n= 295
Charles River-C72 222k SNPs - Average Latency to Magazine Entry Day 3 - n= 358
-log10(P value)
0 1 2 3 4 5 6 7
1 2 3 4 5 6 7 8 9 10 11 12 13 14 15 16 17 18 19 20
Chromosomal Position

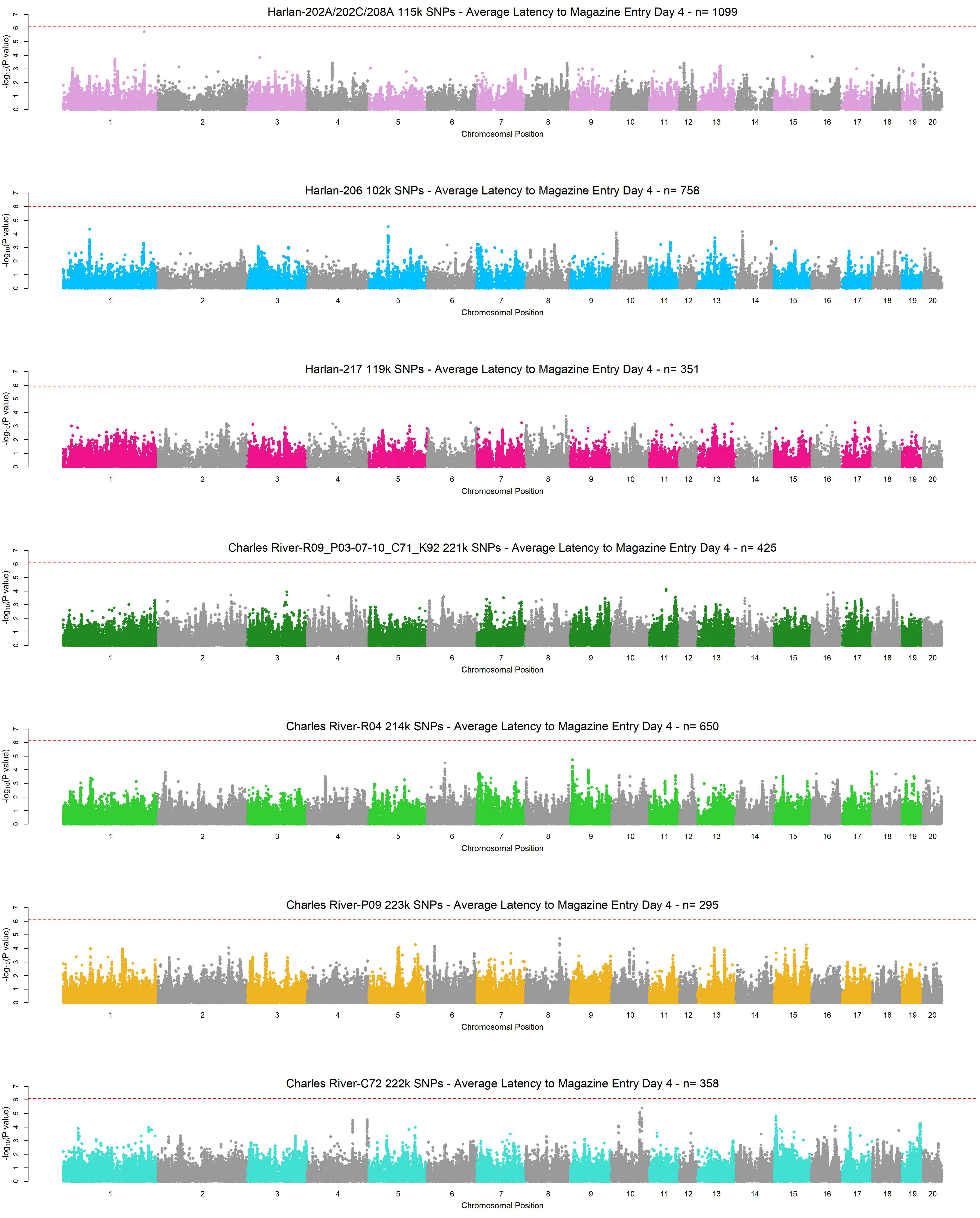
Harlan-202A/202C/208A 115k SNPs - Average Latency to Magazine Entry Day 4 - n= 1099
Harlan-206 102k SNPs - Average Latency to Magazine Entry Day 4 - n= 758
Harlan-217 119k SNPs - Average Latency to Magazine Entry Day 4 - n= 351
Charles River-R09_P03-07-10_C71_K92 221k SNPs - Average Latency to Magazine Entry Day 4 - n= 425
Charles River-R04 214k SNPs - Average Latency to Magazine Entry Day 4 - n= 650
Charles River-P09 223k SNPs - Average Latency to Magazine Entry Day 4 - n= 295
Charles River-C72 222k SNPs - Average Latency to Magazine Entry Day 4 - n= 358
-log10(P value)
Chromosomal Position
1 2 3 4 5 6 7 8 9 10 11 12 13 14 15 16 17 18 19 20
0 1 2 3 4 5 6 7

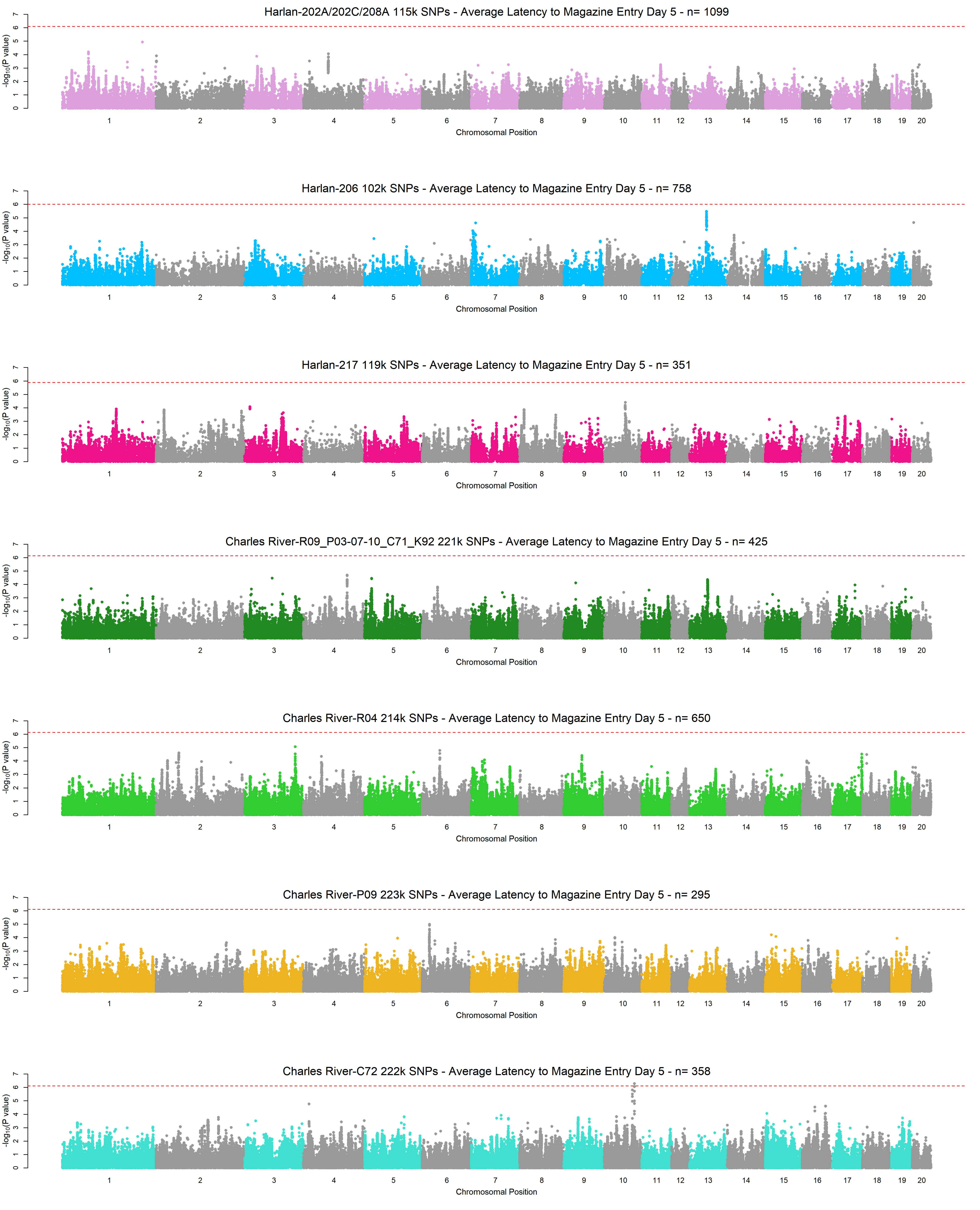

Harlan-202A/202C/208A 115k SNPs - Average Latency to Magazine Entry Day 5 - n= 1099
Harlan-206 102k SNPs - Average Latency to Magazine Entry Day 5 - n= 758
Harlan-217 119k SNPs - Average Latency to Magazine Entry Day 5 - n= 351
Charles River-R09_P03-07-10_C71_K92 221k SNPs - Average Latency to Magazine Entry Day 5 - n= 425
Charles River-R04 214k SNPs - Average Latency to Magazine Entry Day 5 - n= 650
Charles River-P09 223k SNPs - Average Latency to Magazine Entry Day 5 - n= 295
Charles River-C72 222k SNPs - Average Latency to Magazine Entry Day 5 - n= 358
$-\log_{10}$(P value)
Chromosomal Position
1 2 3 4 5 6 7 8 9 10 11 12 13 14 15 16 17 18 19 20

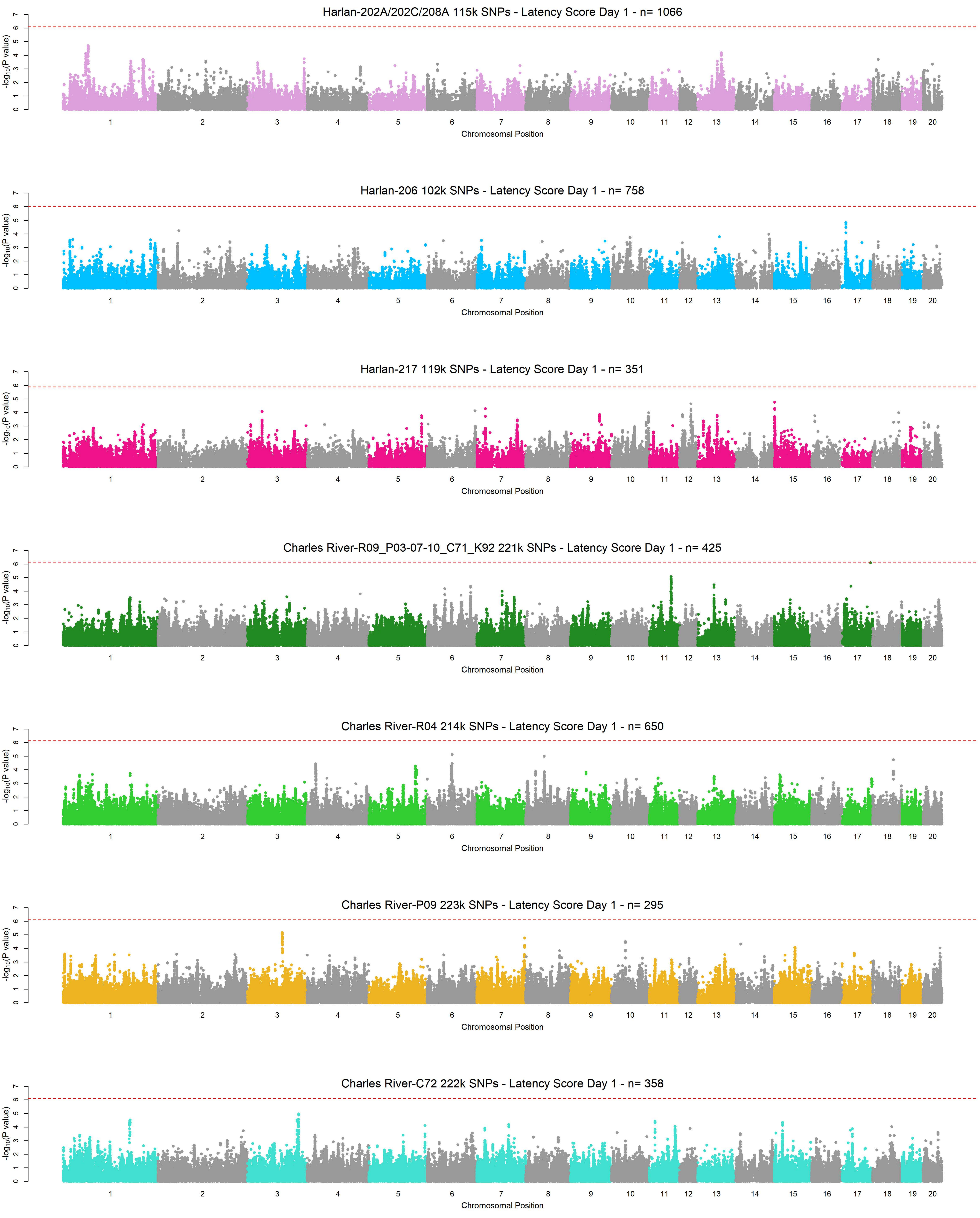
Harlan-202A/202C/208A 115k SNPs - Latency Score Day 1 - n= 1066
Harlan-206 102k SNPs - Latency Score Day 1 - n= 758
Harlan-217 119k SNPs - Latency Score Day 1 - n= 351
Charles River-R09_P03-07-10_C71_K92 221k SNPs - Latency Score Day 1 - n= 425
Charles River-R04 214k SNPs - Latency Score Day 1 - n= 650
Charles River-P09 223k SNPs - Latency Score Day 1 - n= 295
Charles River-C72 222k SNPs - Latency Score Day 1 - n= 358
-log10(P value)
0 1 2 3 4 5 6 7
1 2 3 4 5 6 7 8 9 10 11 12 13 14 15 16 17 18 19 20
Chromosomal Position

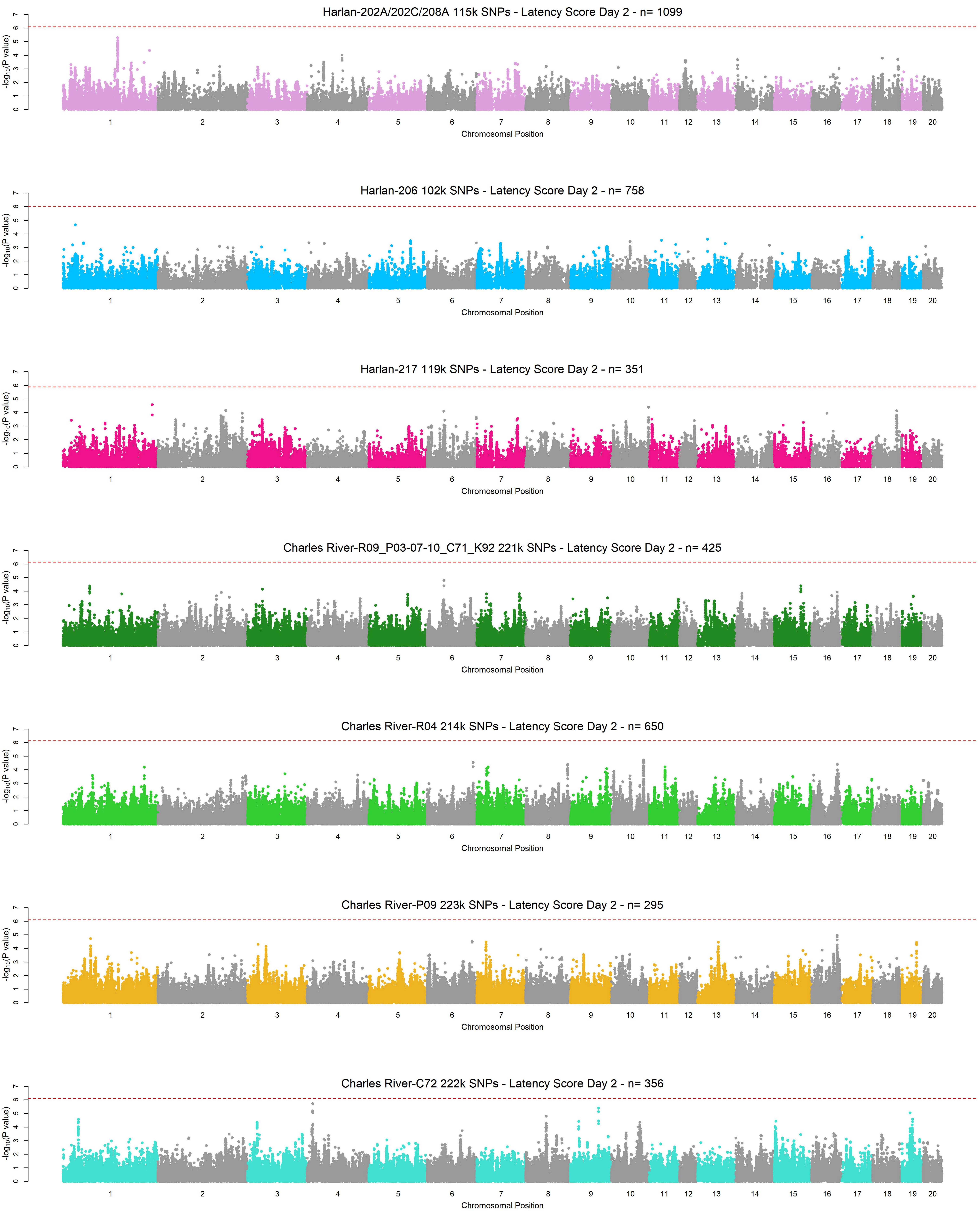
Harlan-202A/202C/208A 115k SNPs - Latency Score Day 2 - n= 1099
Harlan-206 102k SNPs - Latency Score Day 2 - n= 758
Harlan-217 119k SNPs - Latency Score Day 2 - n= 351
Charles River-R09_P03-07-10_C71_K92 221k SNPs - Latency Score Day 2 - n= 425
Charles River-R04 214k SNPs - Latency Score Day 2 - n= 650
Charles River-P09 223k SNPs - Latency Score Day 2 - n= 295
Charles River-C72 222k SNPs - Latency Score Day 2 - n= 356
$-\log_{10}$(P value)
0 1 2 3 4 5 6 7
1 2 3 4 5 6 7 8 9 10 11 12 13 14 15 16 17 18 19 20
Chromosomal Position

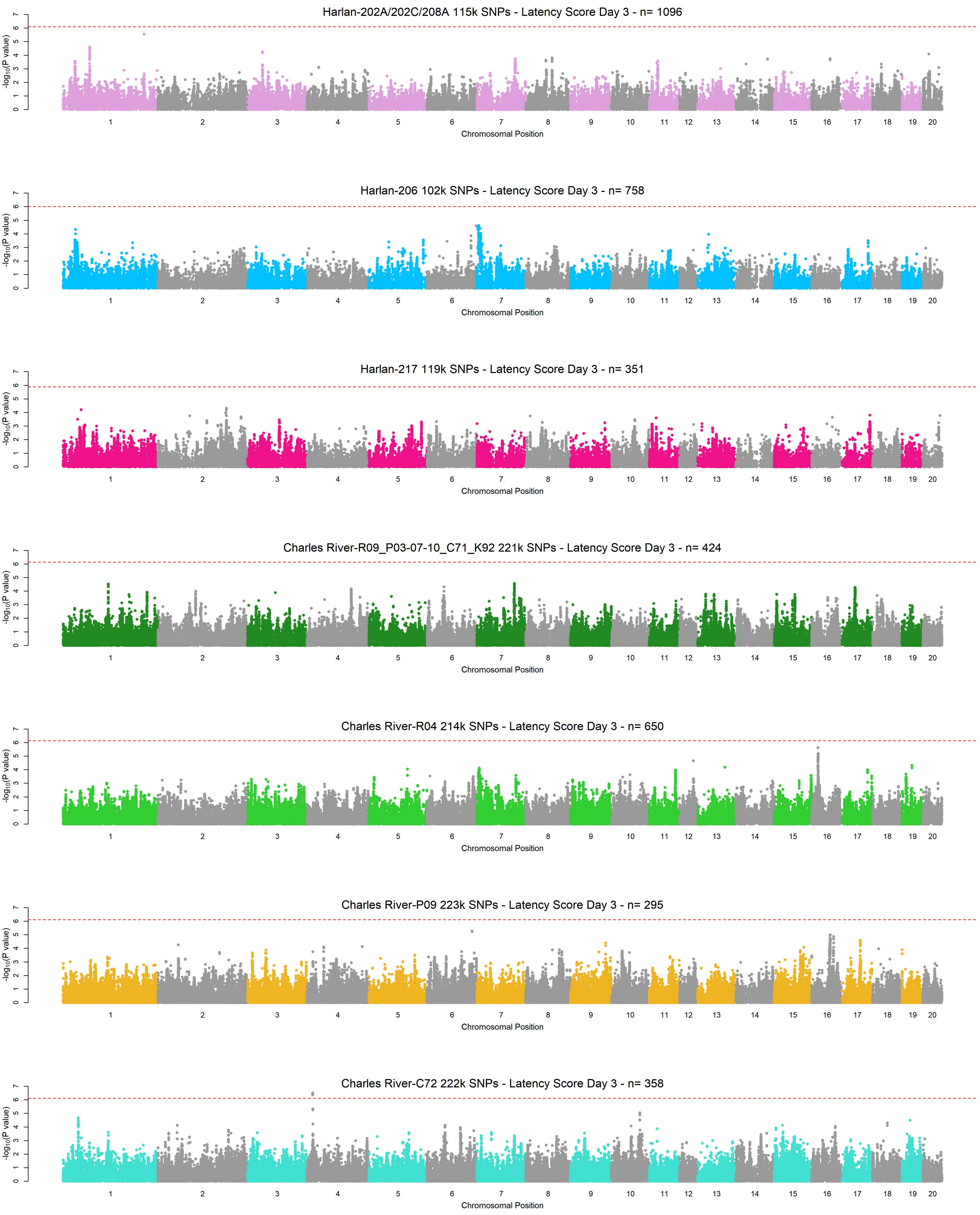
Harlan-202A/202C/208A 115k SNPs - Latency Score Day 3 - n= 1096
Harlan-206 102k SNPs - Latency Score Day 3 - n= 758
Harlan-217 119k SNPs - Latency Score Day 3 - n= 351
Charles River-R09_P03-07-10_C71_K92 221k SNPs - Latency Score Day 3 - n= 424
Charles River-R04 214k SNPs - Latency Score Day 3 - n= 650
Charles River-P09 223k SNPs - Latency Score Day 3 - n= 295
Charles River-C72 222k SNPs - Latency Score Day 3 - n= 358
-log10(P value)
Chromosomal Position
1 2 3 4 5 6 7 8 9 10 11 12 13 14 15 16 17 18 19 20
0 1 2 3 4 5 6 7

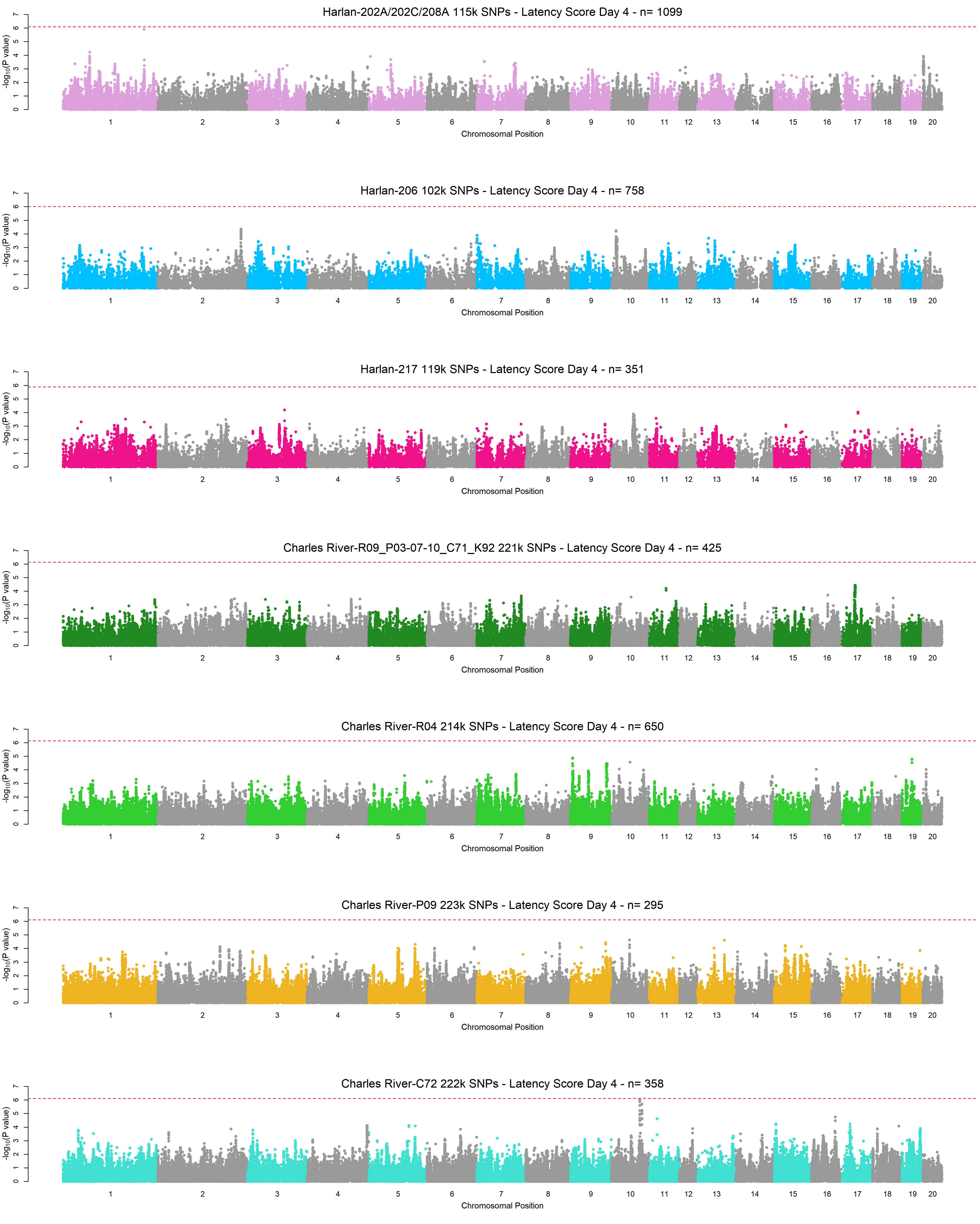

Harlan-202A/202C/208A 115k SNPs - Latency Score Day 4 - n= 1099
Harlan-206 102k SNPs - Latency Score Day 4 - n= 758
Harlan-217 119k SNPs - Latency Score Day 4 - n= 351
Charles River-R09_P03-07-10_C71_K92 221k SNPs - Latency Score Day 4 - n= 425
Charles River-R04 214k SNPs - Latency Score Day 4 - n= 650
Charles River-P09 223k SNPs - Latency Score Day 4 - n= 295
Charles River-C72 222k SNPs - Latency Score Day 4 - n= 358
$-\log_{10}$(P value)
Chromosomal Position
0 1 2 3 4 5 6 7
1 2 3 4 5 6 7 8 9 10 11 12 13 14 15 16 17 18 19 20

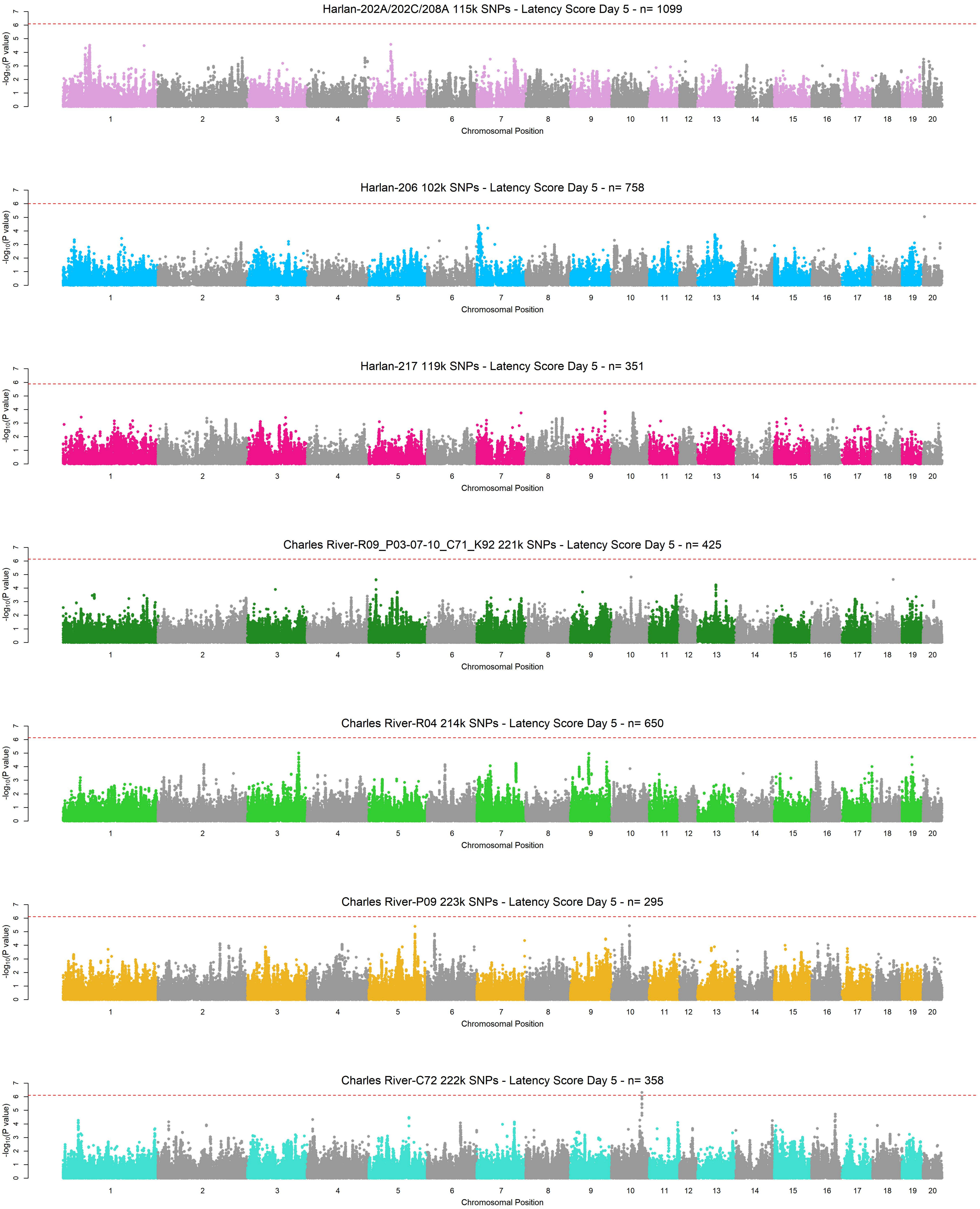
Harlan-202A/202C/208A 115k SNPs - Latency Score Day 5 - n= 1099
Harlan-206 102k SNPs - Latency Score Day 5 - n= 758
Harlan-217 119k SNPs - Latency Score Day 5 - n= 351
Charles River-R09_P03-07-10_C71_K92 221k SNPs - Latency Score Day 5 - n= 425
Charles River-R04 214k SNPs - Latency Score Day 5 - n= 650
Charles River-P09 223k SNPs - Latency Score Day 5 - n= 295
Charles River-C72 222k SNPs - Latency Score Day 5 - n= 358
-log10(P value)
Chromosomal Position
1 2 3 4 5 6 7 8 9 10 11 12 13 14 15 16 17 18 19 20
0 1 2 3 4 5 6 7

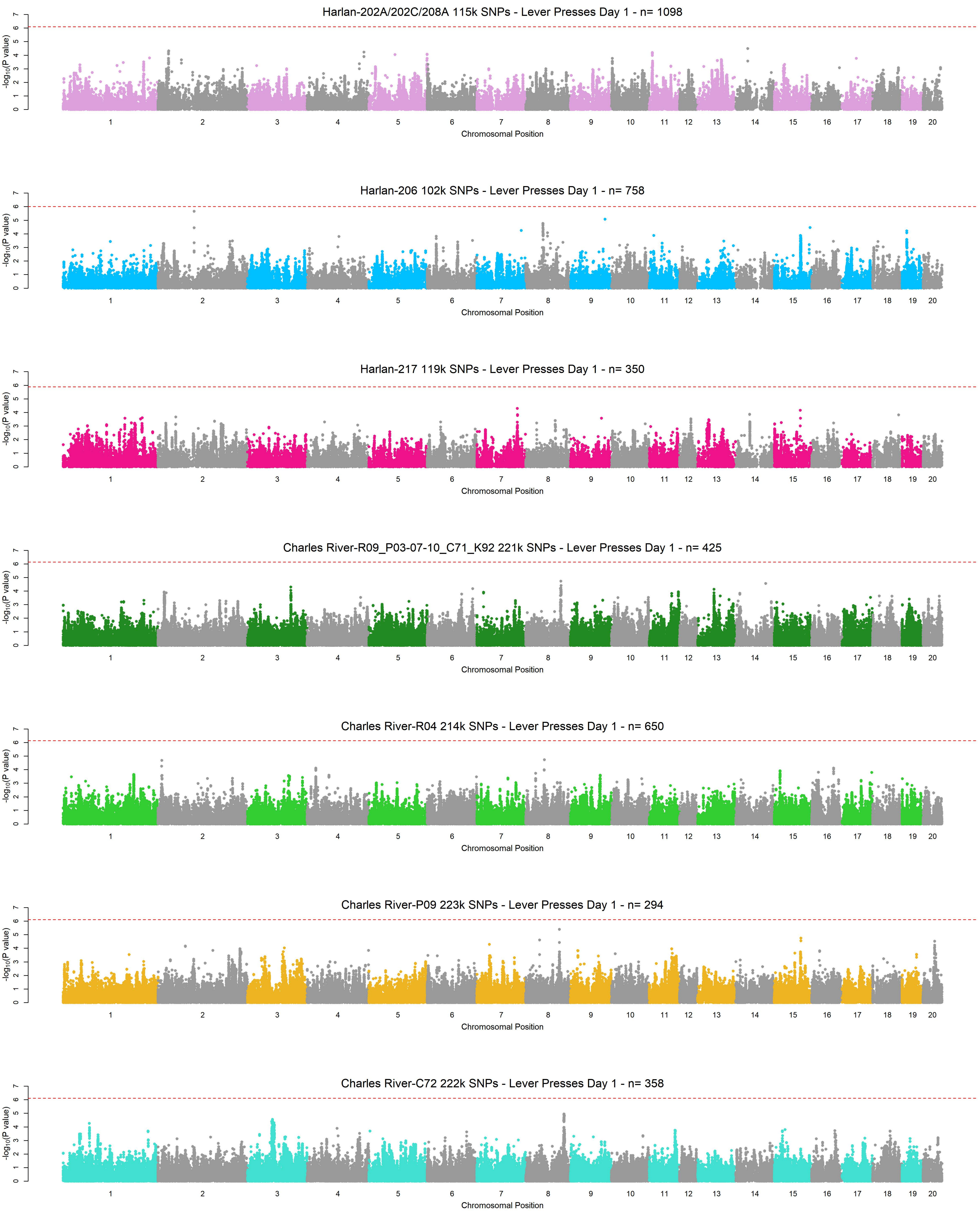
Harlan-202A/202C/208A 115k SNPs - Lever Presses Day 1 - n= 1098
Harlan-206 102k SNPs - Lever Presses Day 1 - n= 758
Harlan-217 119k SNPs - Lever Presses Day 1 - n= 350
Charles River-R09_P03-07-10_C71_K92 221k SNPs - Lever Presses Day 1 - n= 425
Charles River-R04 214k SNPs - Lever Presses Day 1 - n= 650
Charles River-P09 223k SNPs - Lever Presses Day 1 - n= 294
Charles River-C72 222k SNPs - Lever Presses Day 1 - n= 358
-log10(P value)
Chromosomal Position
1 2 3 4 5 6 7 8 9 10 11 12 13 14 15 16 17 18 19 20
0 1 2 3 4 5 6 7

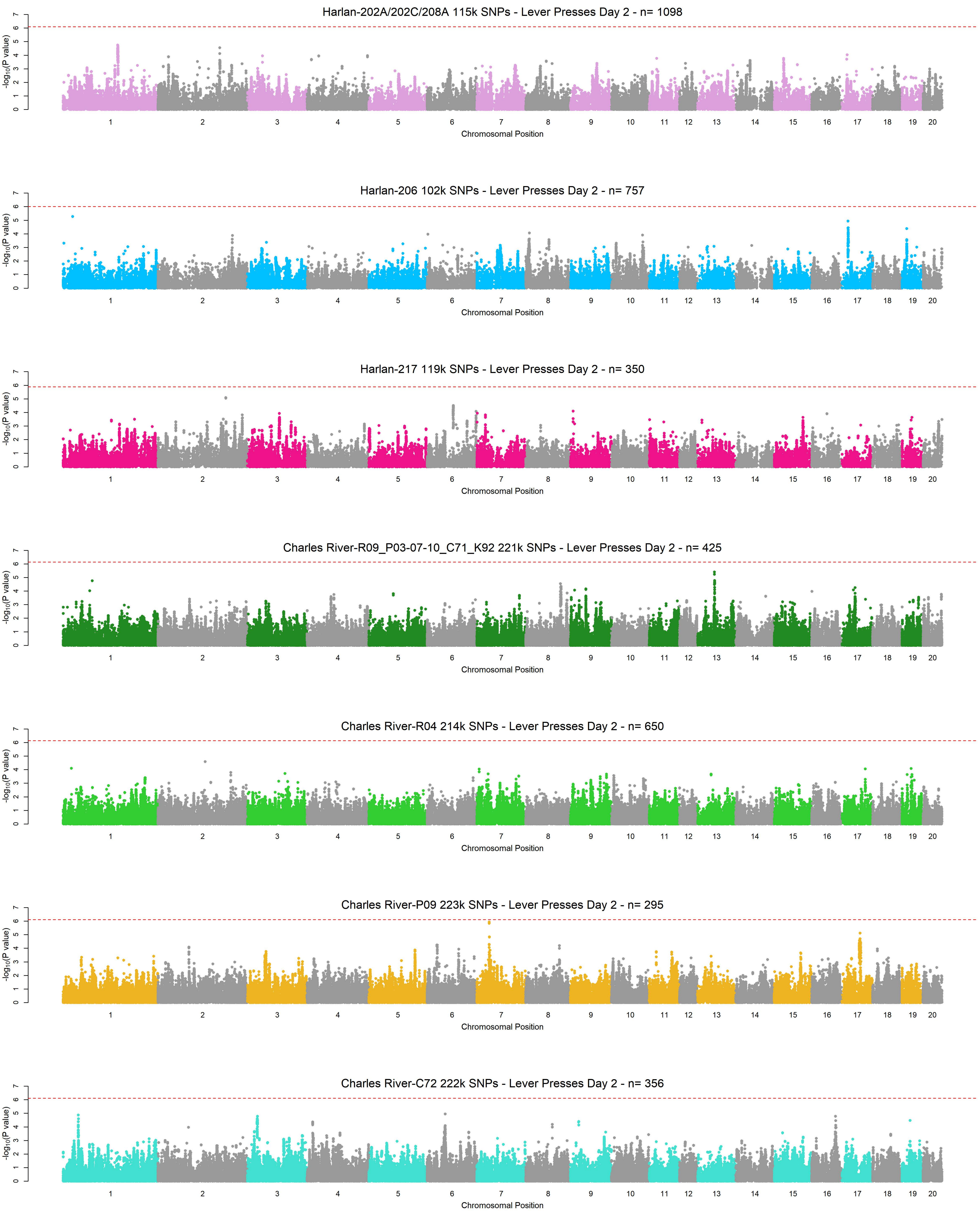
Harlan-202A/202C/208A 115k SNPs - Lever Presses Day 2 - n= 1098
Harlan-206 102k SNPs - Lever Presses Day 2 - n= 757
Harlan-217 119k SNPs - Lever Presses Day 2 - n= 350
Charles River-R09_P03-07-10_C71_K92 221k SNPs - Lever Presses Day 2 - n= 425
Charles River-R04 214k SNPs - Lever Presses Day 2 - n= 650
Charles River-P09 223k SNPs - Lever Presses Day 2 - n= 295
Charles River-C72 222k SNPs - Lever Presses Day 2 - n= 356
-log10(P value)
Chromosomal Position
0 1 2 3 4 5 6 7
1 2 3 4 5 6 7 8 9 10 11 12 13 14 15 16 17 18 19 20

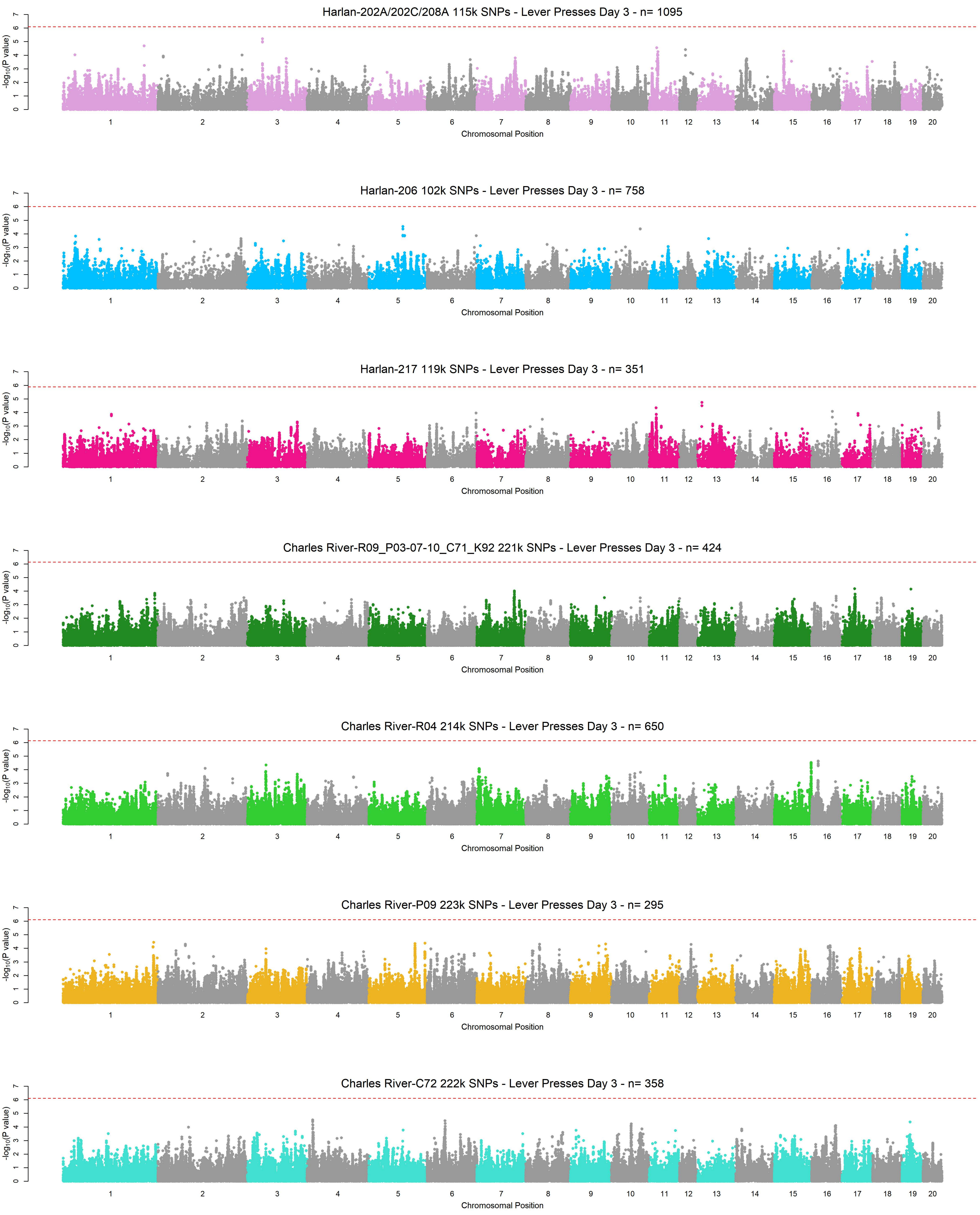
Harlan-202A/202C/208A 115k SNPs - Lever Presses Day 3 - n= 1095
Harlan-206 102k SNPs - Lever Presses Day 3 - n= 758
Harlan-217 119k SNPs - Lever Presses Day 3 - n= 351
Charles River-R09_P03-07-10_C71_K92 221k SNPs - Lever Presses Day 3 - n= 424
Charles River-R04 214k SNPs - Lever Presses Day 3 - n= 650
Charles River-P09 223k SNPs - Lever Presses Day 3 - n= 295
Charles River-C72 222k SNPs - Lever Presses Day 3 - n= 358
$-\log_{10}$(P value)
Chromosomal Position
1 2 3 4 5 6 7 8 9 10 11 12 13 14 15 16 17 18 19 20
0 1 2 3 4 5 6 7

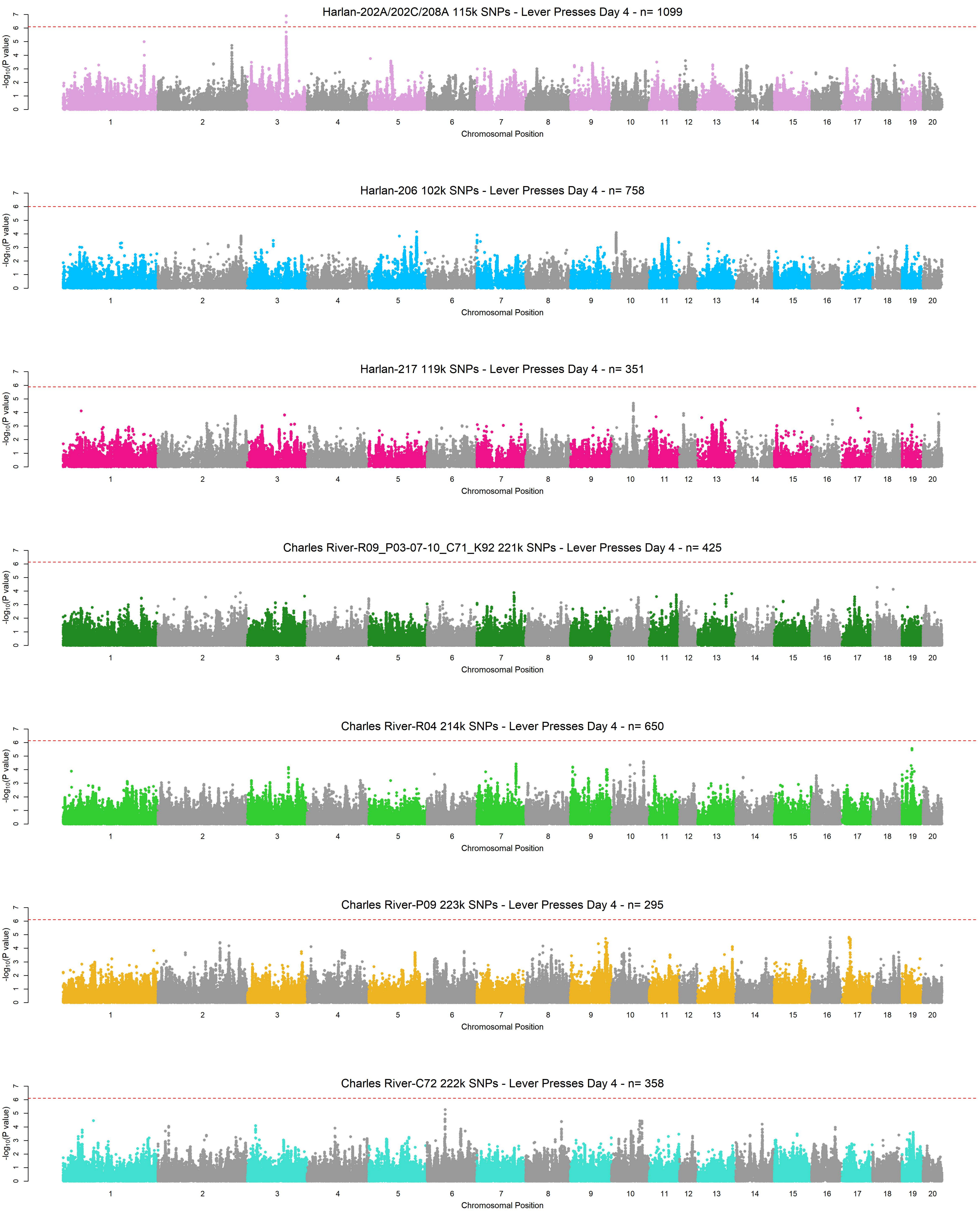

Harlan-202A/202C/208A 115k SNPs - Lever Presses Day 4 - n= 1099
Harlan-206 102k SNPs - Lever Presses Day 4 - n= 758
Harlan-217 119k SNPs - Lever Presses Day 4 - n= 351
Charles River-R09_P03-07-10_C71_K92 221k SNPs - Lever Presses Day 4 - n= 425
Charles River-R04 214k SNPs - Lever Presses Day 4 - n= 650
Charles River-P09 223k SNPs - Lever Presses Day 4 - n= 295
Charles River-C72 222k SNPs - Lever Presses Day 4 - n= 358
-log10(P value)
Chromosomal Position
1 2 3 4 5 6 7 8 9 10 11 12 13 14 15 16 17 18 19 20

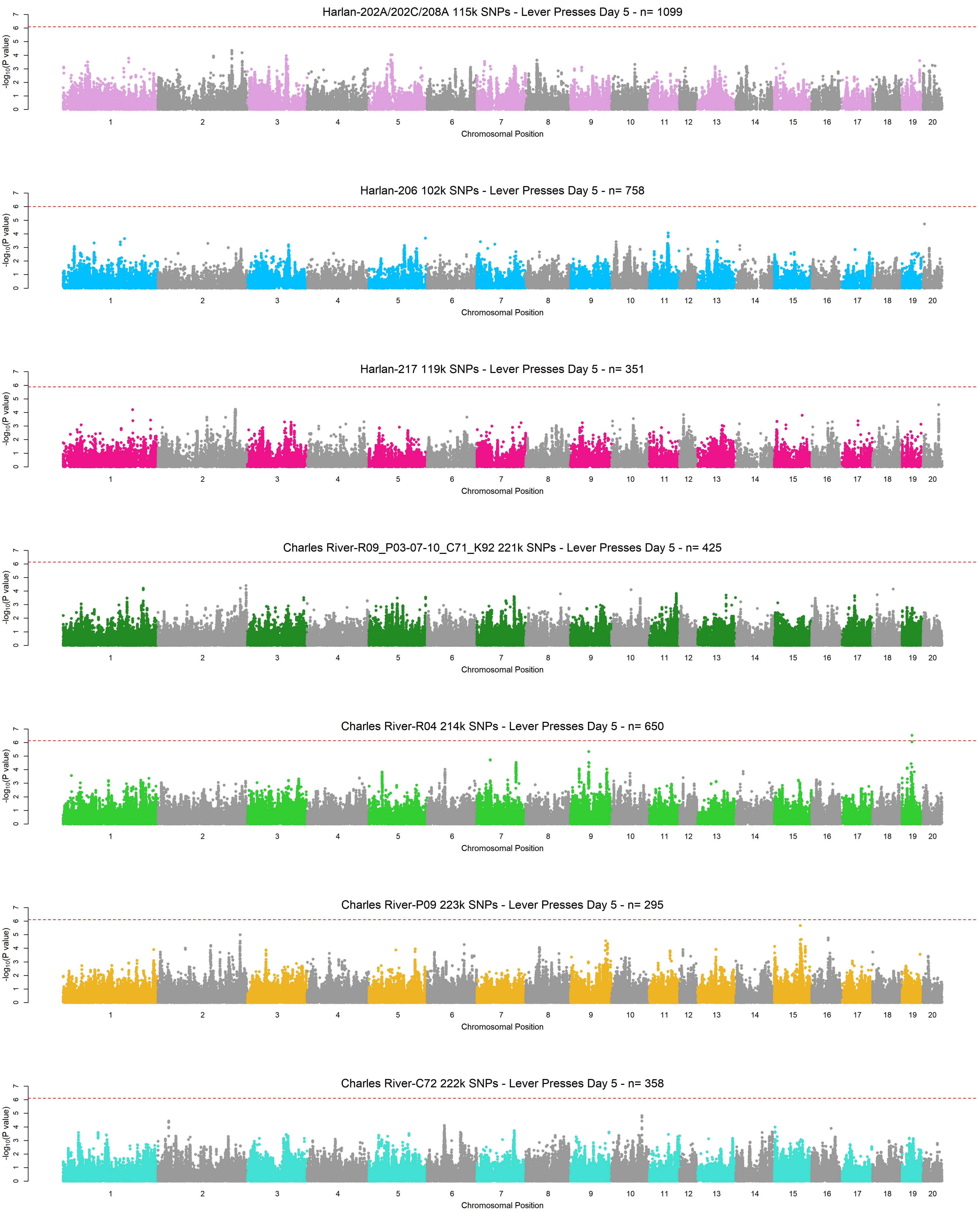

Harlan-202A/202C/208A 115k SNPs - Lever Presses Day 5 - n= 1099
Harlan-206 102k SNPs - Lever Presses Day 5 - n= 758
Harlan-217 119k SNPs - Lever Presses Day 5 - n= 351
Charles River-R09_P03-07-10_C71_K92 221k SNPs - Lever Presses Day 5 - n= 425
Charles River-R04 214k SNPs - Lever Presses Day 5 - n= 650
Charles River-P09 223k SNPs - Lever Presses Day 5 - n= 295
Charles River-C72 222k SNPs - Lever Presses Day 5 - n= 358
-log10(P value)
Chromosomal Position
1 2 3 4 5 6 7 8 9 10 11 12 13 14 15 16 17 18 19 20

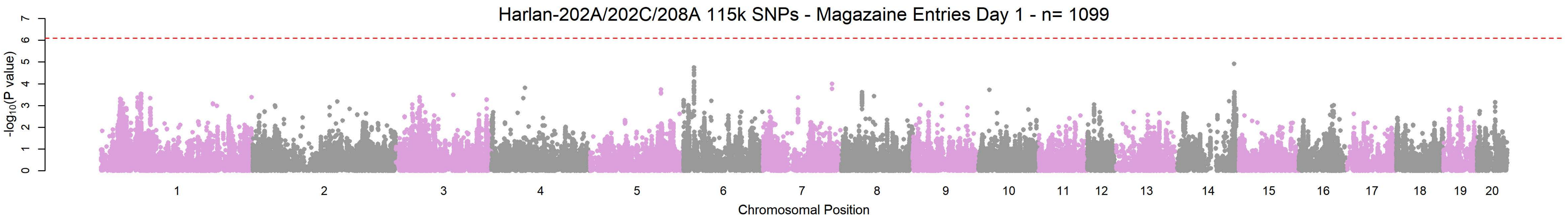
Harlan-202A/202C/208A 115k SNPs - Magazaine Entries Day 1 - n= 1099
-log10(P value)
Chromosomal Position

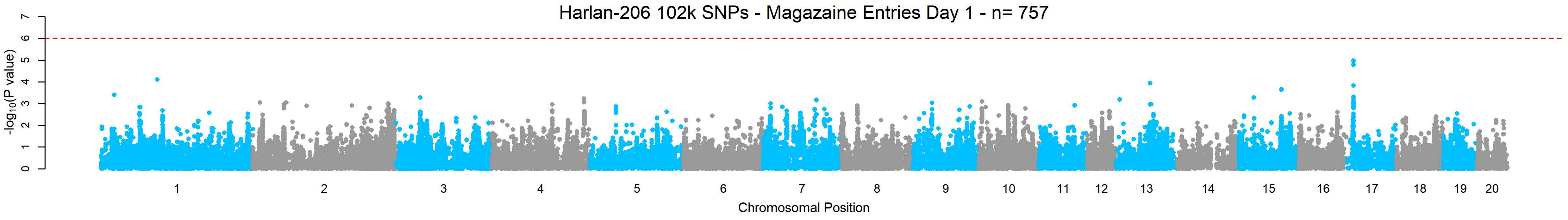
Harlan-206 102k SNPs - Magazaine Entries Day 1 - n= 757
-log10(P value)
Chromosomal Position

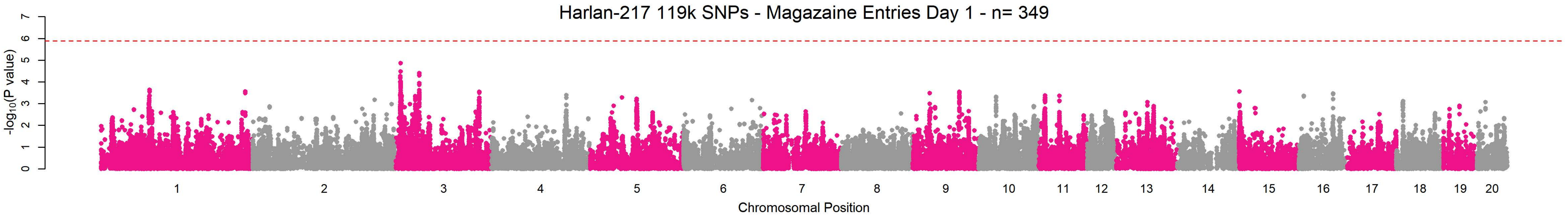
Harlan-217 119k SNPs - Magazaine Entries Day 1 - n= 349
-log10(P value)
Chromosomal Position

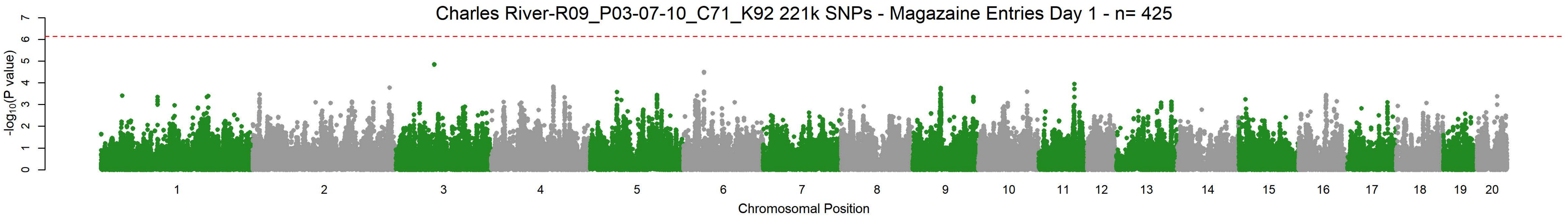
Charles River-R09_P03-07-10_C71_K92 221k SNPs - Magazaine Entries Day 1 - n= 425
-log10(P value)
Chromosomal Position

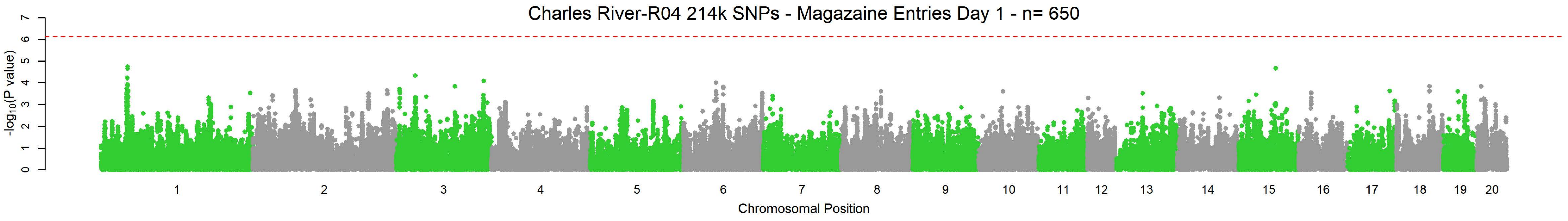
Charles River-R04 214k SNPs - Magazaine Entries Day 1 - n= 650
-log10(P value)
Chromosomal Position

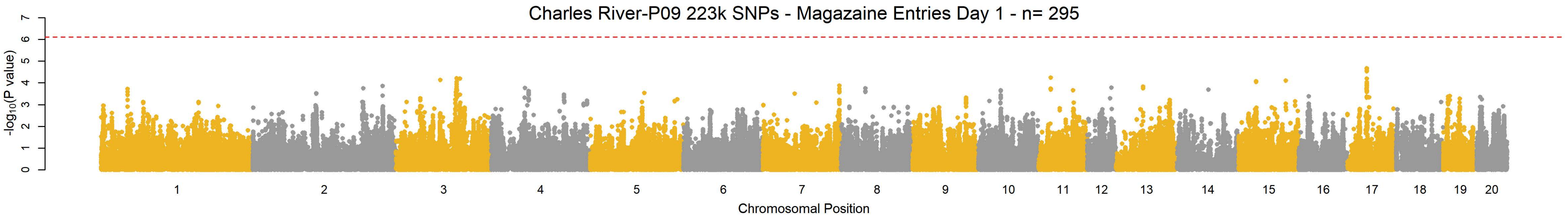
Charles River-P09 223k SNPs - Magazaine Entries Day 1 - n= 295
-log10(P value)
Chromosomal Position

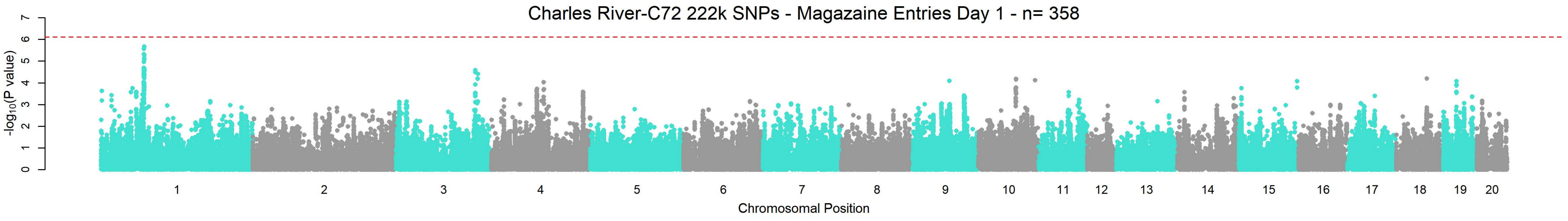
Charles River-C72 222k SNPs - Magazaine Entries Day 1 - n= 358
-log10(P value)
Chromosomal Position

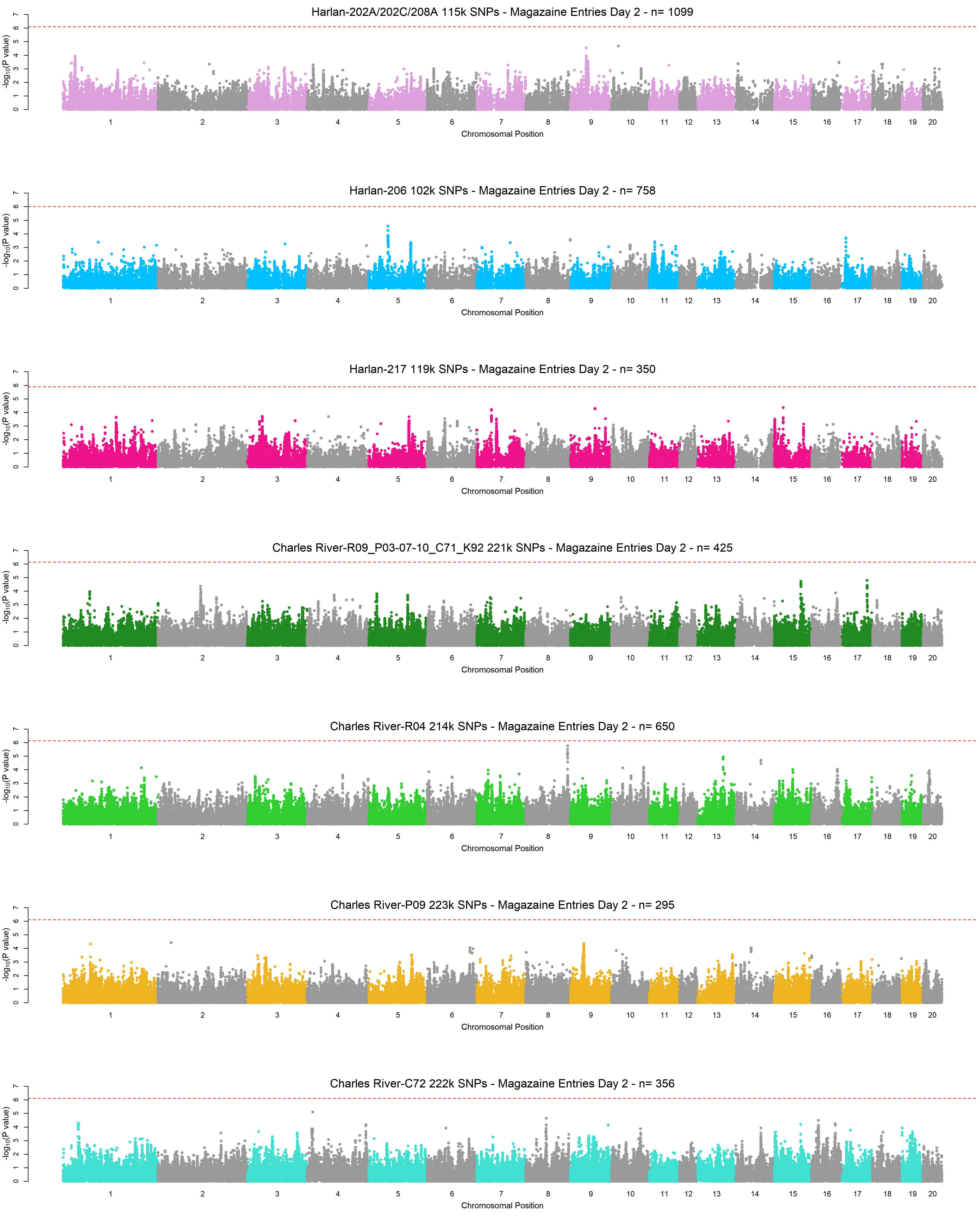
Harlan-202A/202C/208A 115k SNPs - Magazaine Entries Day 2 - n= 1099
Harlan-206 102k SNPs - Magazaine Entries Day 2 - n= 758
Harlan-217 119k SNPs - Magazaine Entries Day 2 - n= 350
Charles River-R09_P03-07-10_C71_K92 221k SNPs - Magazaine Entries Day 2 - n= 425
Charles River-R04 214k SNPs - Magazaine Entries Day 2 - n= 650
Charles River-P09 223k SNPs - Magazaine Entries Day 2 - n= 295
Charles River-C72 222k SNPs - Magazaine Entries Day 2 - n= 356
$-\log_{10}$(P value)
Chromosomal Position
1 2 3 4 5 6 7 8 9 10 11 12 13 14 15 16 17 18 19 20

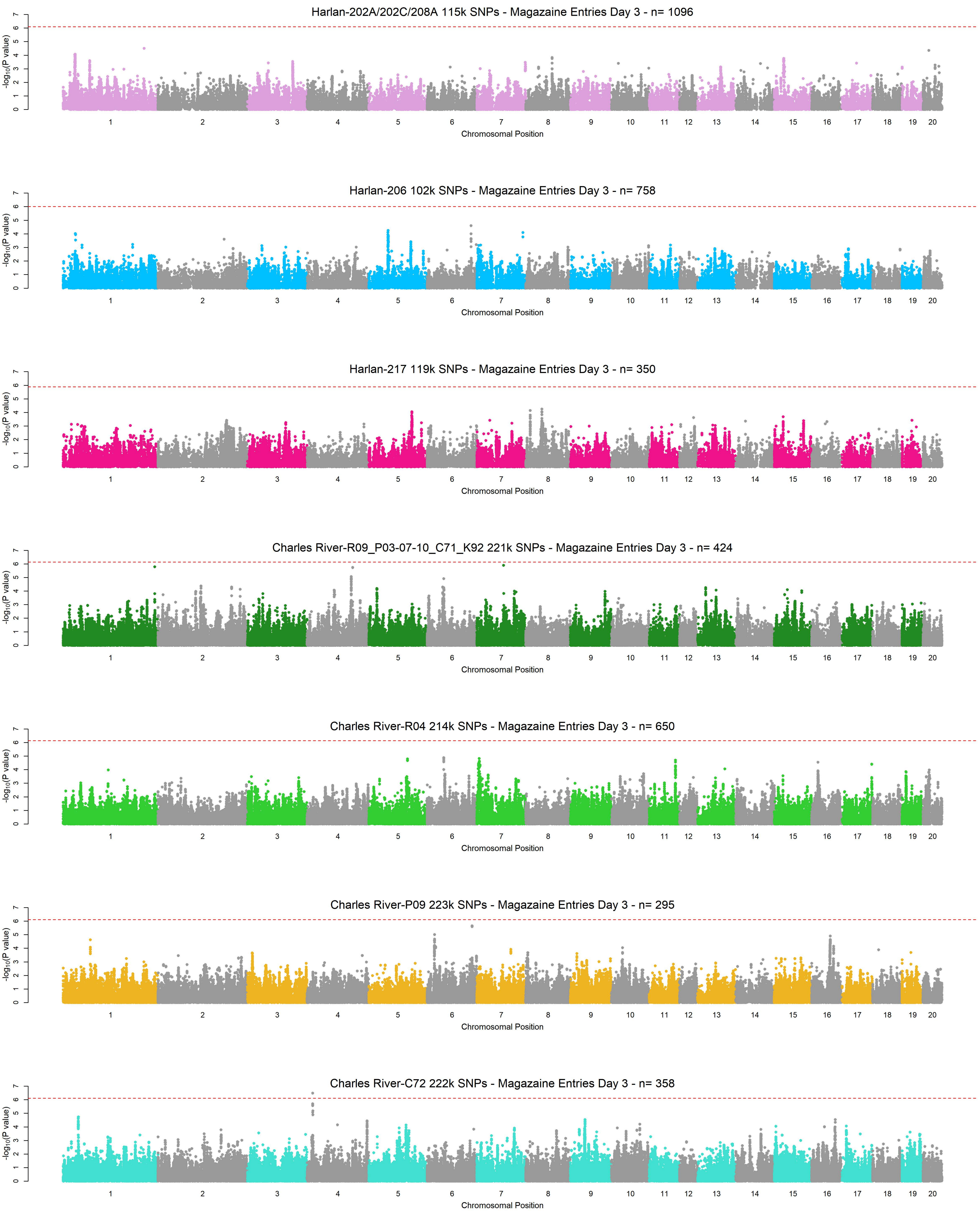
Harlan-202A/202C/208A 115k SNPs - Magazaine Entries Day 3 - n= 1096
Harlan-206 102k SNPs - Magazaine Entries Day 3 - n= 758
Harlan-217 119k SNPs - Magazaine Entries Day 3 - n= 350
Charles River-R09_P03-07-10_C71_K92 221k SNPs - Magazaine Entries Day 3 - n= 424
Charles River-R04 214k SNPs - Magazaine Entries Day 3 - n= 650
Charles River-P09 223k SNPs - Magazaine Entries Day 3 - n= 295
Charles River-C72 222k SNPs - Magazaine Entries Day 3 - n= 358
-log10(P value)
Chromosomal Position
1 2 3 4 5 6 7 8 9 10 11 12 13 14 15 16 17 18 19 20

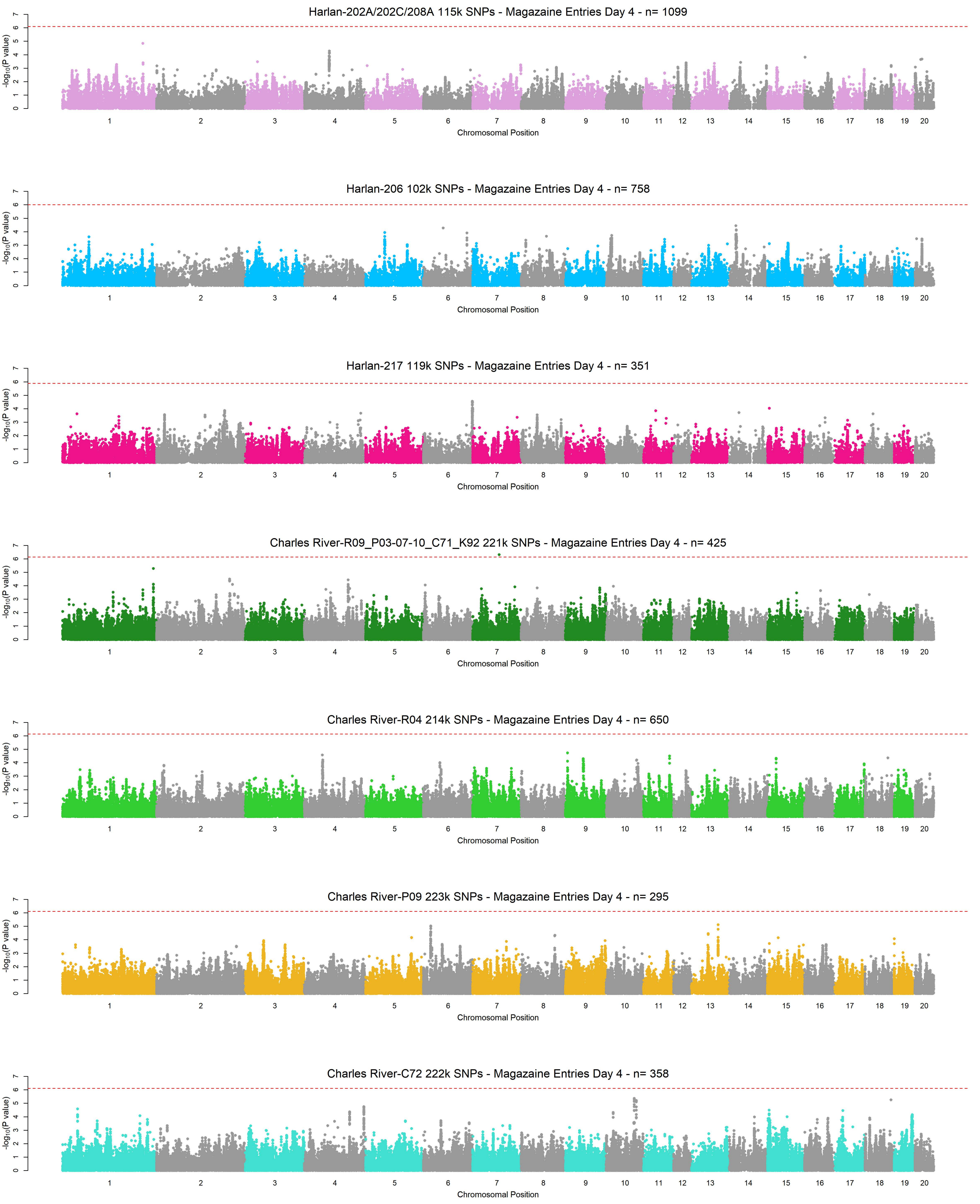
Harlan-202A/202C/208A 115k SNPs - Magazaine Entries Day 4 - n= 1099
Harlan-206 102k SNPs - Magazaine Entries Day 4 - n= 758
Harlan-217 119k SNPs - Magazaine Entries Day 4 - n= 351
Charles River-R09_P03-07-10_C71_K92 221k SNPs - Magazaine Entries Day 4 - n= 425
Charles River-R04 214k SNPs - Magazaine Entries Day 4 - n= 650
Charles River-P09 223k SNPs - Magazaine Entries Day 4 - n= 295
Charles River-C72 222k SNPs - Magazaine Entries Day 4 - n= 358
-log10(P value)
Chromosomal Position
1 2 3 4 5 6 7 8 9 10 11 12 13 14 15 16 17 18 19 20
0 1 2 3 4 5 6 7

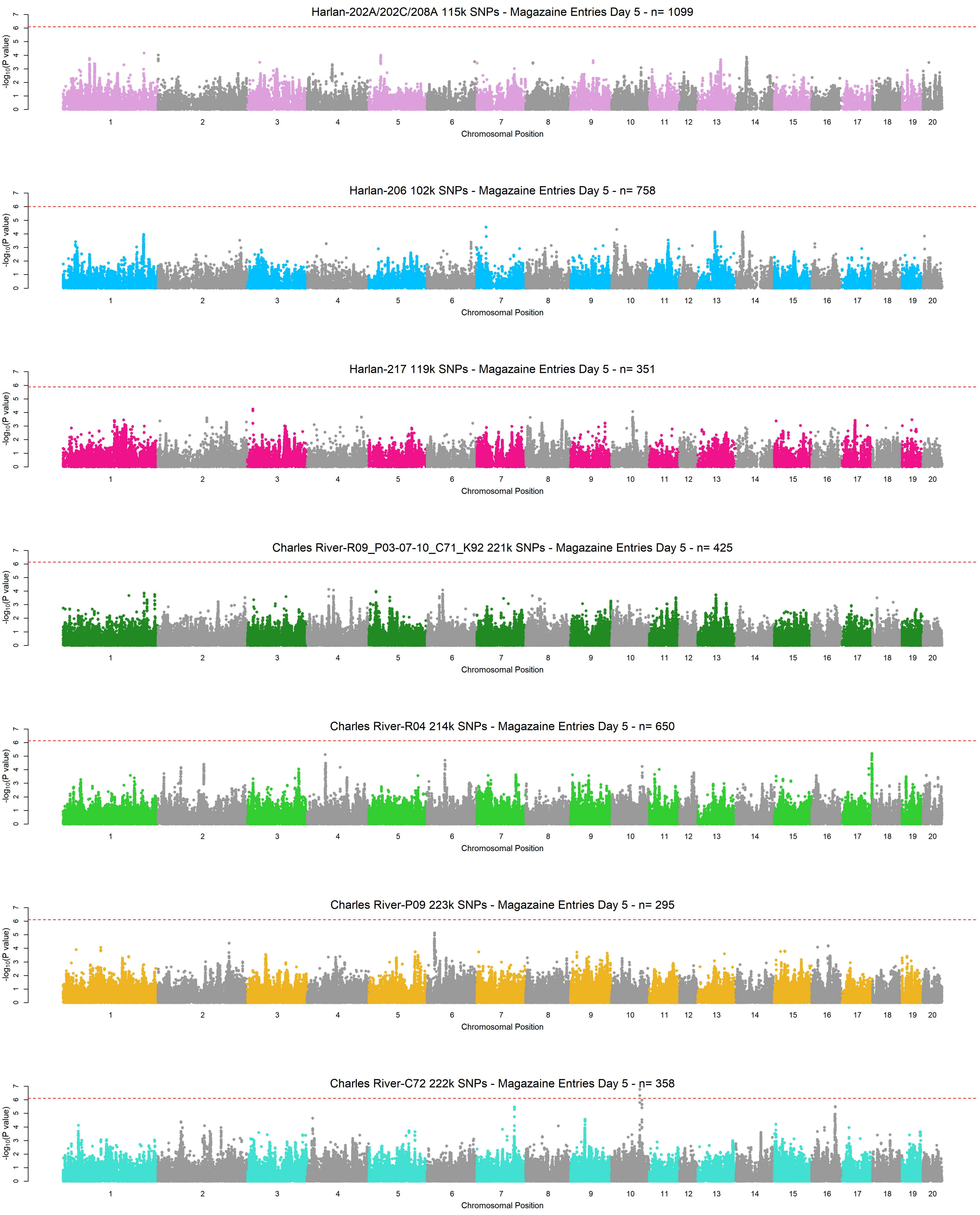
Harlan-202A/202C/208A 115k SNPs - Magazaine Entries Day 5 - n= 1099
Harlan-206 102k SNPs - Magazaine Entries Day 5 - n= 758
Harlan-217 119k SNPs - Magazaine Entries Day 5 - n= 351
Charles River-R09_P03-07-10_C71_K92 221k SNPs - Magazaine Entries Day 5 - n= 425
Charles River-R04 214k SNPs - Magazaine Entries Day 5 - n= 650
Charles River-P09 223k SNPs - Magazaine Entries Day 5 - n= 295
Charles River-C72 222k SNPs - Magazaine Entries Day 5 - n= 358
-log10(P value)
0 1 2 3 4 5 6 7
1 2 3 4 5 6 7 8 9 10 11 12 13 14 15 16 17 18 19 20
Chromosomal Position

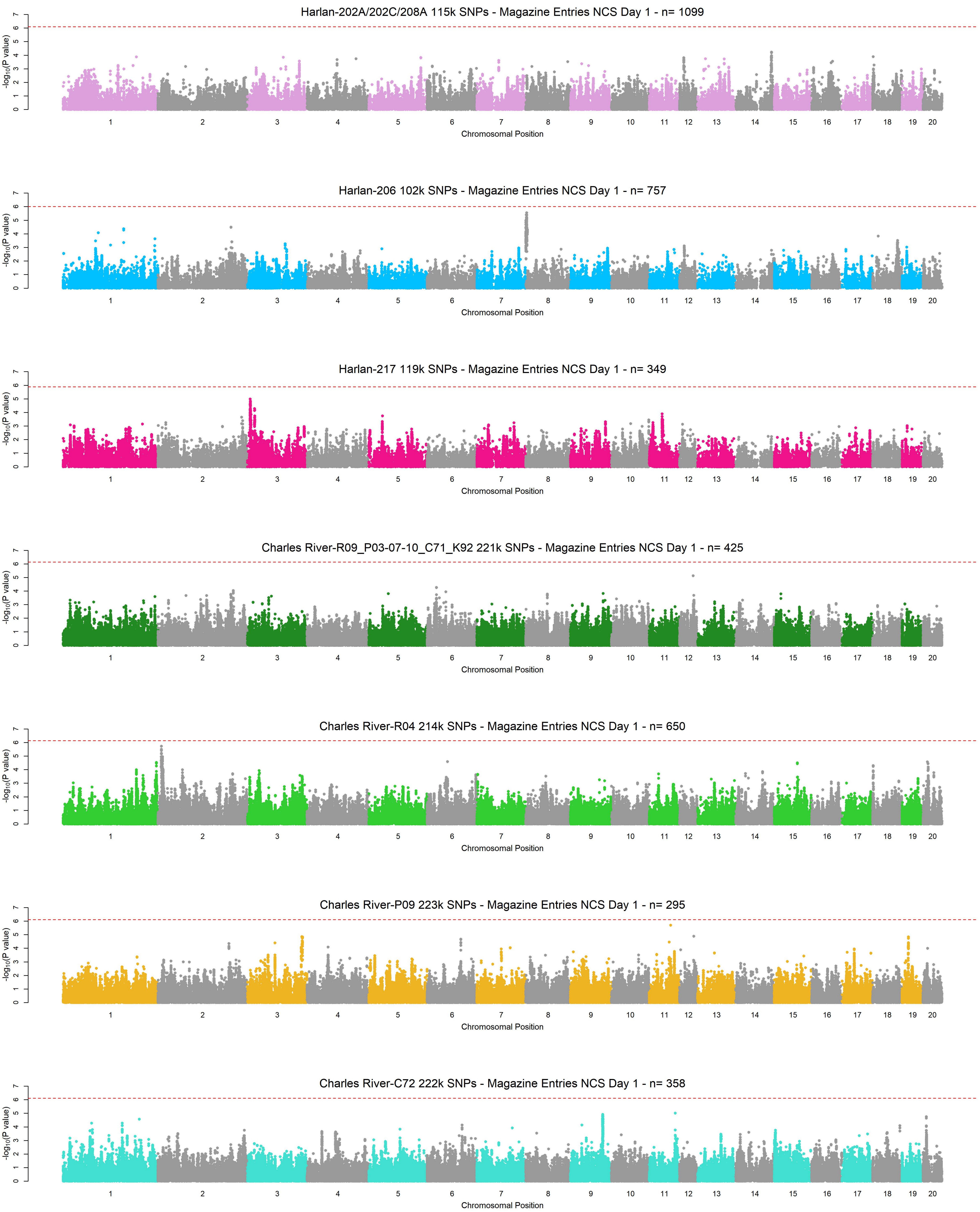
Harlan-202A/202C/208A 115k SNPs - Magazine Entries NCS Day 1 - n= 1099
Harlan-206 102k SNPs - Magazine Entries NCS Day 1 - n= 757
Harlan-217 119k SNPs - Magazine Entries NCS Day 1 - n= 349
Charles River-R09_P03-07-10_C71_K92 221k SNPs - Magazine Entries NCS Day 1 - n= 425
Charles River-R04 214k SNPs - Magazine Entries NCS Day 1 - n= 650
Charles River-P09 223k SNPs - Magazine Entries NCS Day 1 - n= 295
Charles River-C72 222k SNPs - Magazine Entries NCS Day 1 - n= 358
-log10(P value)
Chromosomal Position
1 2 3 4 5 6 7 8 9 10 11 12 13 14 15 16 17 18 19 20

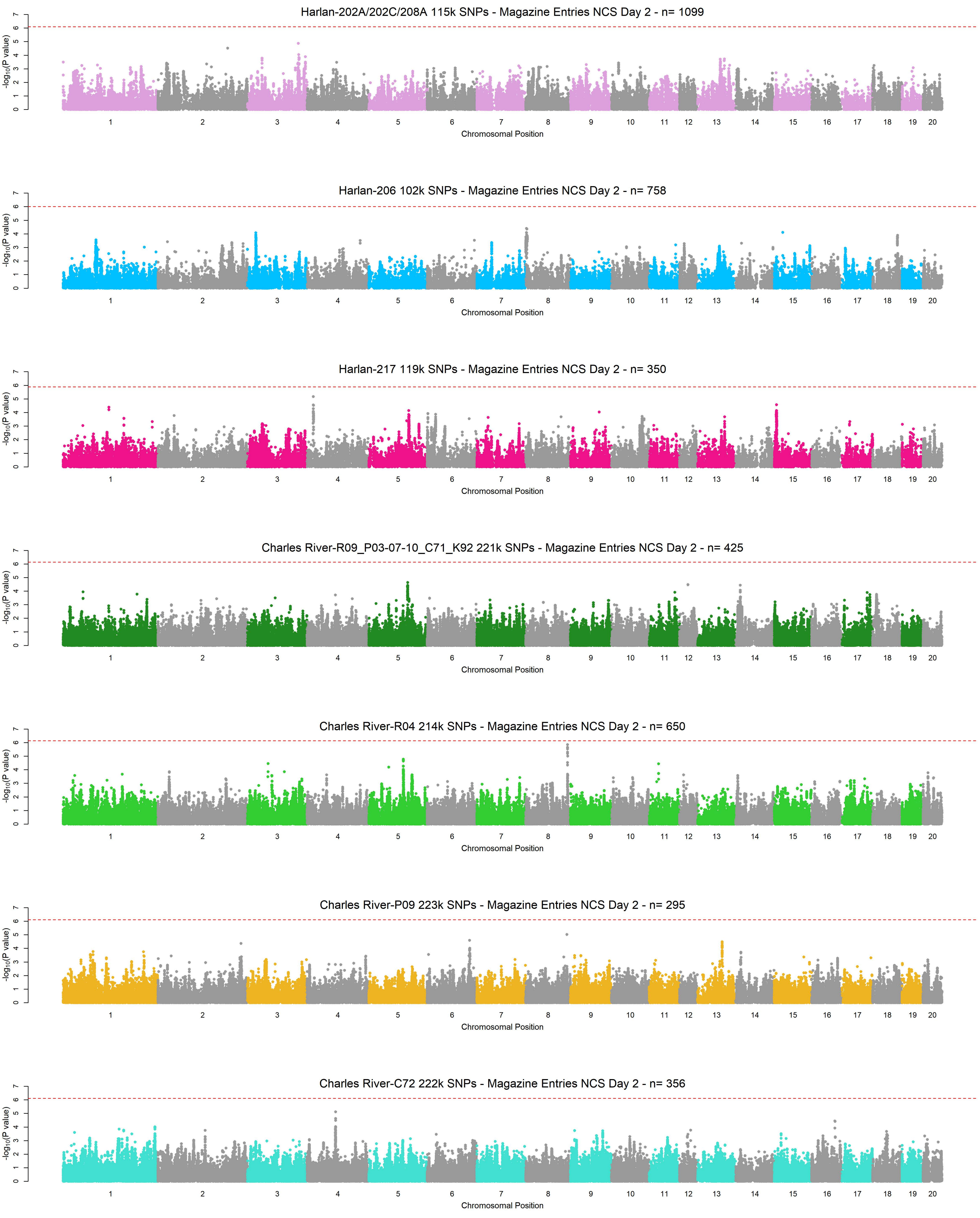
Harlan-202A/202C/208A 115k SNPs - Magazine Entries NCS Day 2 - n= 1099
Harlan-206 102k SNPs - Magazine Entries NCS Day 2 - n= 758
Harlan-217 119k SNPs - Magazine Entries NCS Day 2 - n= 350
Charles River-R09_P03-07-10_C71_K92 221k SNPs - Magazine Entries NCS Day 2 - n= 425
Charles River-R04 214k SNPs - Magazine Entries NCS Day 2 - n= 650
Charles River-P09 223k SNPs - Magazine Entries NCS Day 2 - n= 295
Charles River-C72 222k SNPs - Magazine Entries NCS Day 2 - n= 356
-log10(P value)
Chromosomal Position

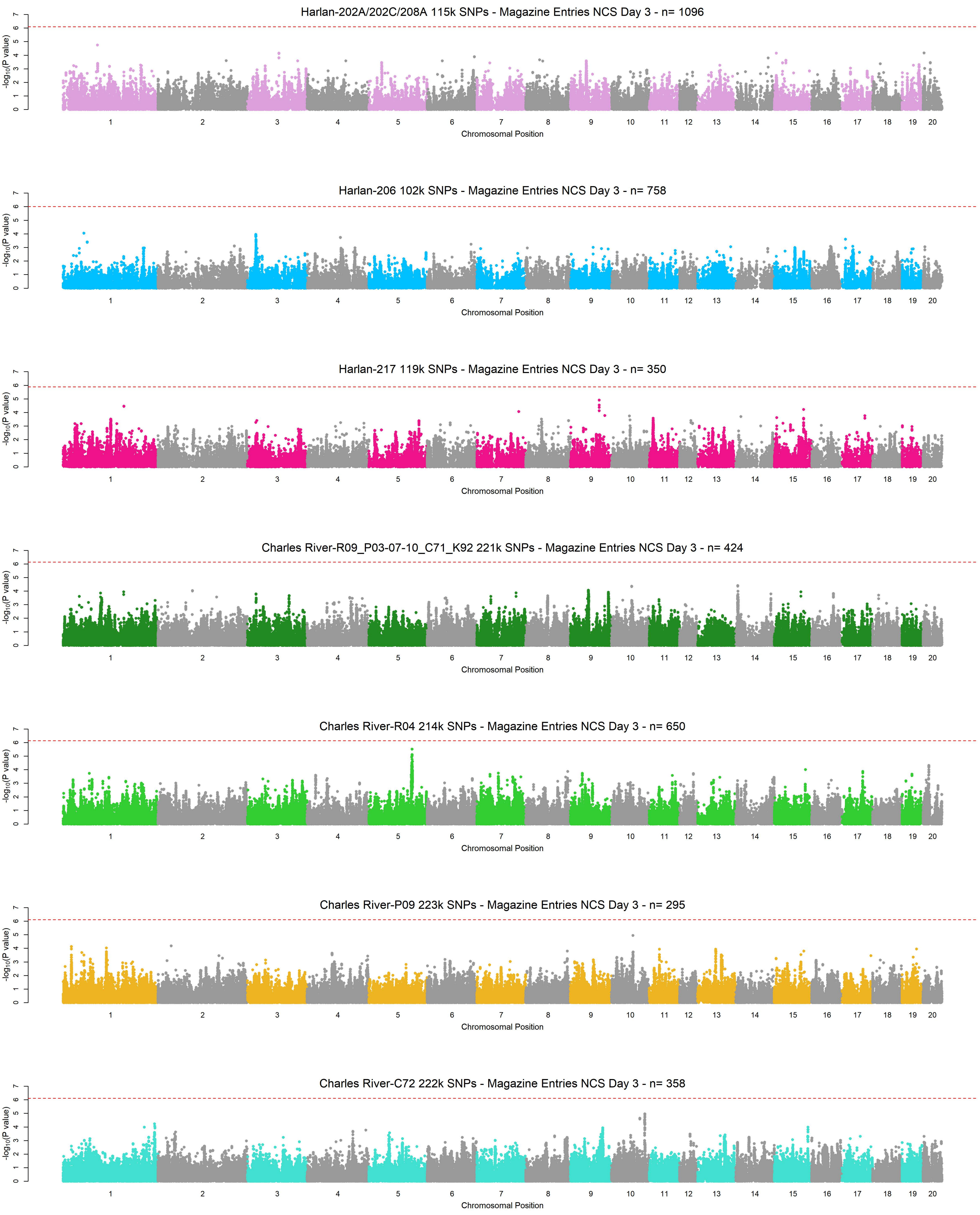
Harlan-202A/202C/208A 115k SNPs - Magazine Entries NCS Day 3 - n= 1096
Harlan-206 102k SNPs - Magazine Entries NCS Day 3 - n= 758
Harlan-217 119k SNPs - Magazine Entries NCS Day 3 - n= 350
Charles River-R09_P03-07-10_C71_K92 221k SNPs - Magazine Entries NCS Day 3 - n= 424
Charles River-R04 214k SNPs - Magazine Entries NCS Day 3 - n= 650
Charles River-P09 223k SNPs - Magazine Entries NCS Day 3 - n= 295
Charles River-C72 222k SNPs - Magazine Entries NCS Day 3 - n= 358
-log10(P value)
Chromosomal Position
1 2 3 4 5 6 7 8 9 10 11 12 13 14 15 16 17 18 19 20

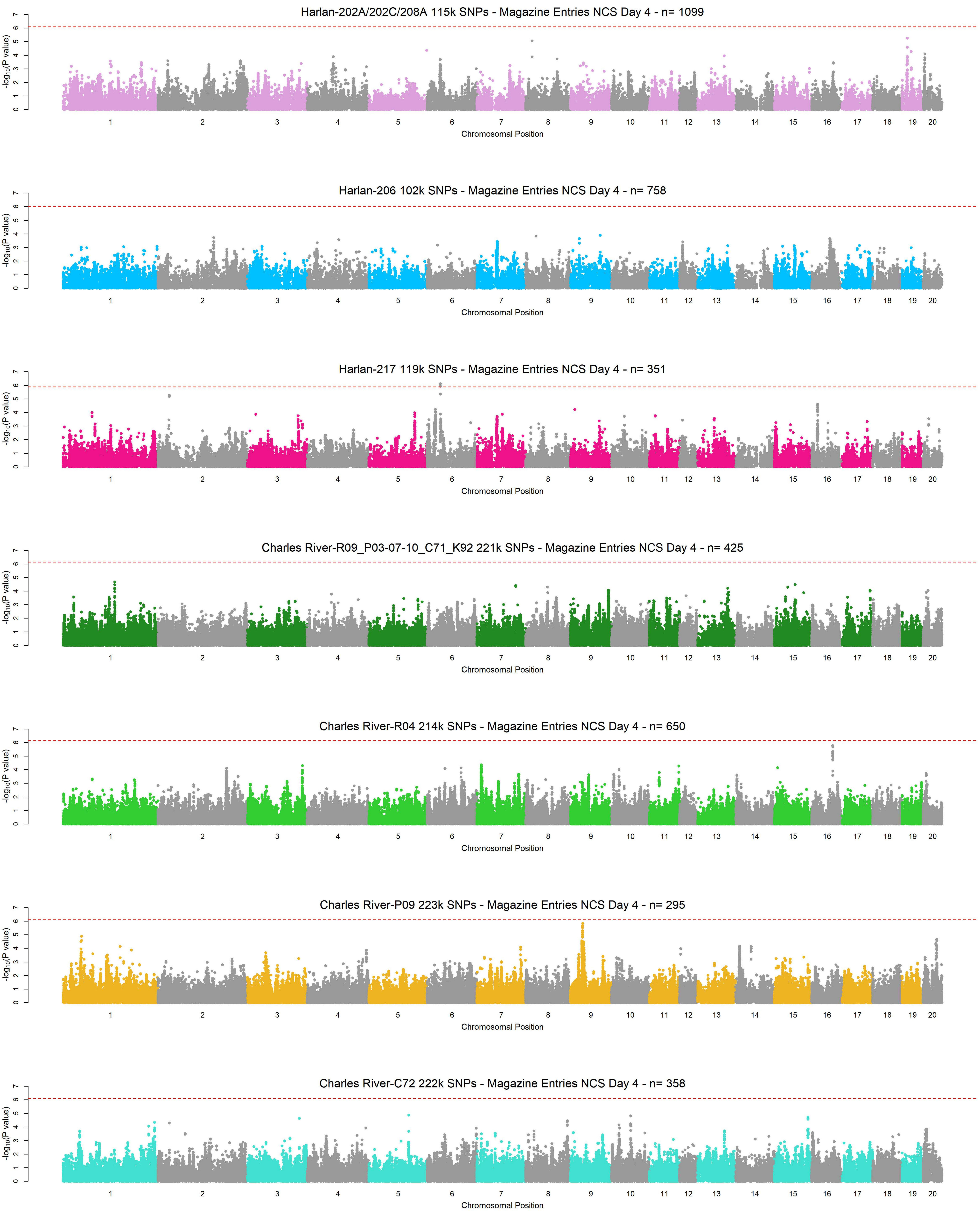
Harlan-202A/202C/208A 115k SNPs - Magazine Entries NCS Day 4 - n= 1099
Harlan-206 102k SNPs - Magazine Entries NCS Day 4 - n= 758
Harlan-217 119k SNPs - Magazine Entries NCS Day 4 - n= 351
Charles River-R09_P03-07-10_C71_K92 221k SNPs - Magazine Entries NCS Day 4 - n= 425
Charles River-R04 214k SNPs - Magazine Entries NCS Day 4 - n= 650
Charles River-P09 223k SNPs - Magazine Entries NCS Day 4 - n= 295
Charles River-C72 222k SNPs - Magazine Entries NCS Day 4 - n= 358
-log10(P value)
Chromosomal Position

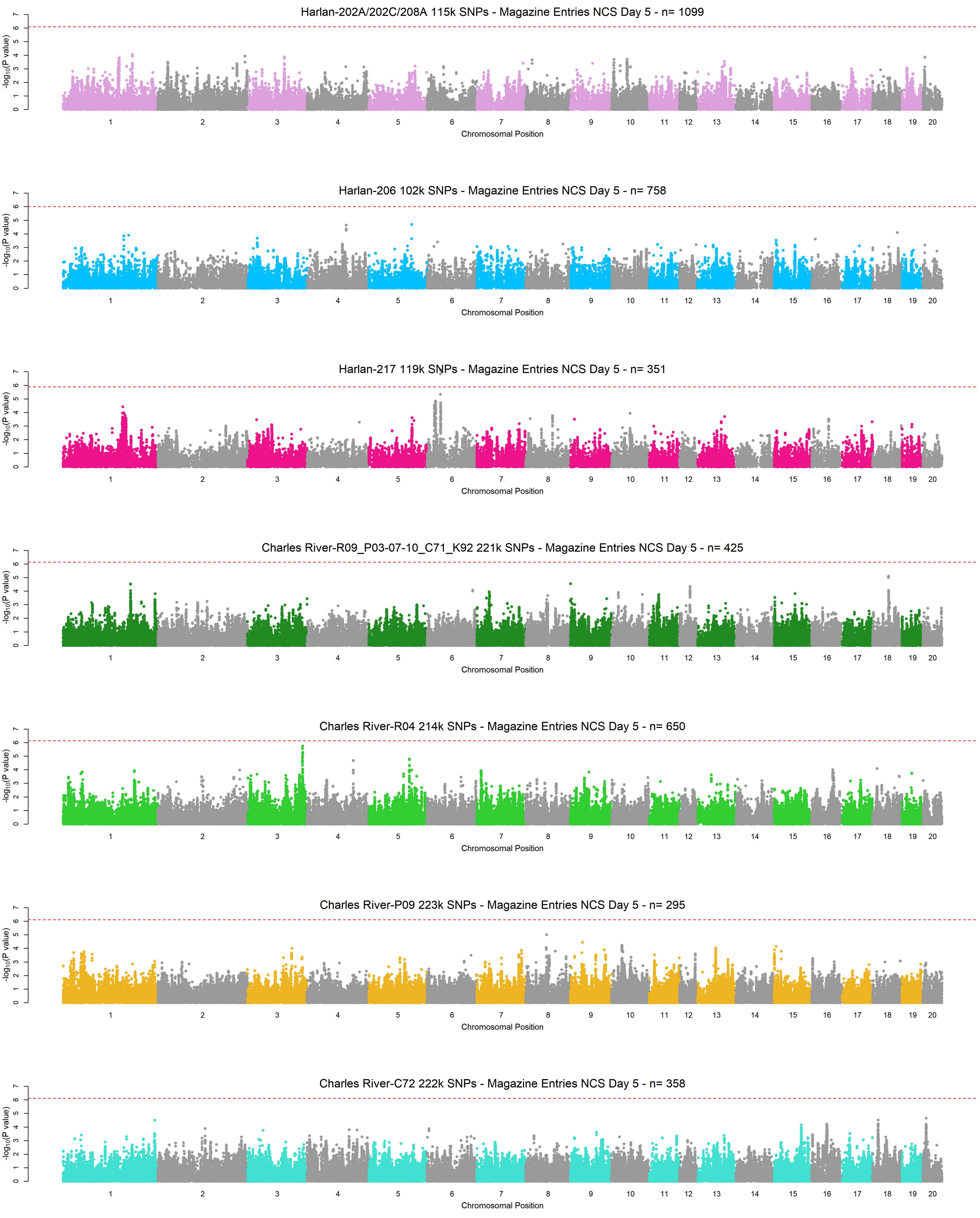
Harlan-202A/202C/208A 115k SNPs - Magazine Entries NCS Day 5 - n= 1099
Harlan-206 102k SNPs - Magazine Entries NCS Day 5 - n= 758
Harlan-217 119k SNPs - Magazine Entries NCS Day 5 - n= 351
Charles River-R09_P03-07-10_C71_K92 221k SNPs - Magazine Entries NCS Day 5 - n= 425
Charles River-R04 214k SNPs - Magazine Entries NCS Day 5 - n= 650
Charles River-P09 223k SNPs - Magazine Entries NCS Day 5 - n= 295
Charles River-C72 222k SNPs - Magazine Entries NCS Day 5 - n= 358
$-\log_{10}$(P value)
0 1 2 3 4 5 6 7
1 2 3 4 5 6 7 8 9 10 11 12 13 14 15 16 17 18 19 20
Chromosomal Position

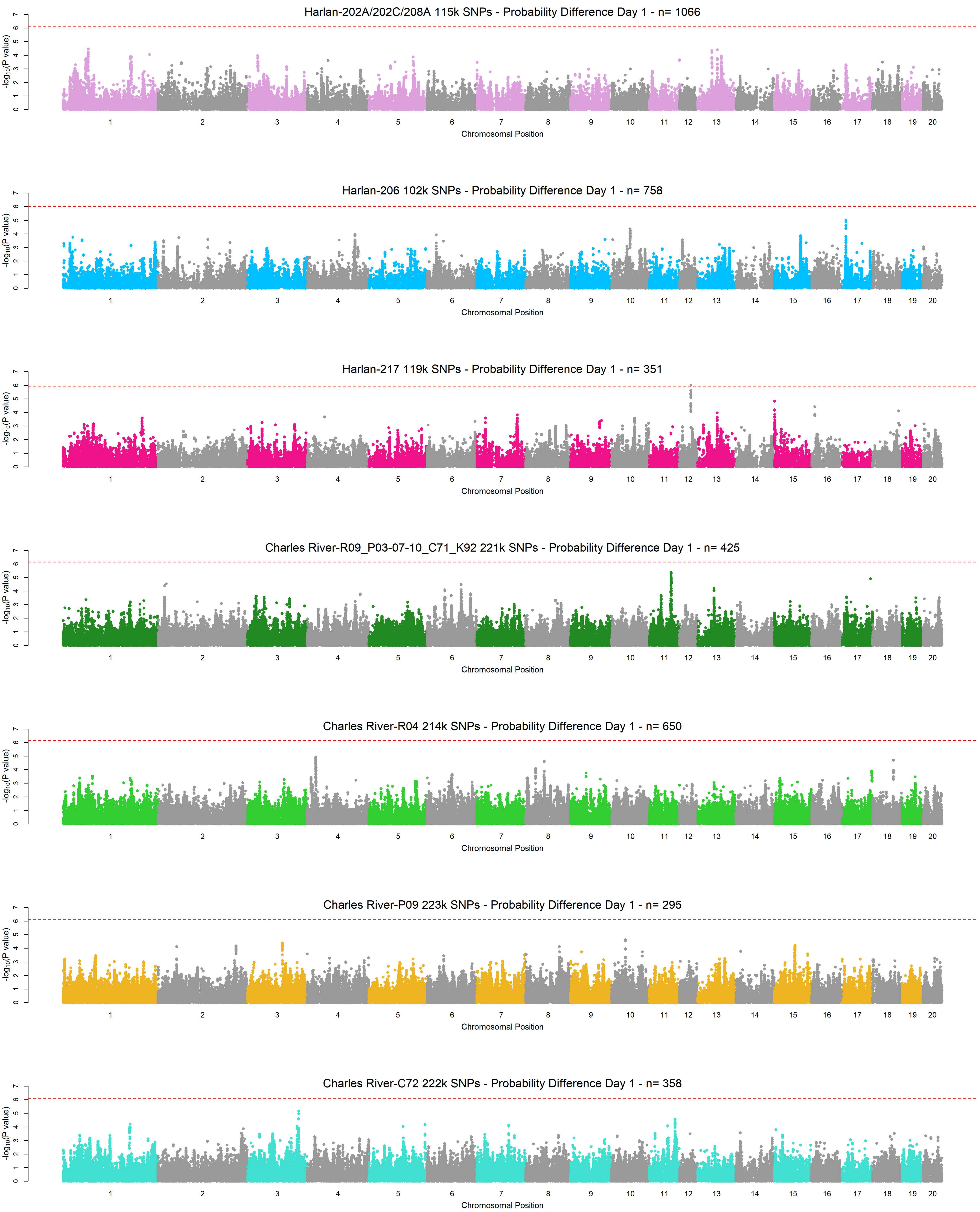
Harlan-202A/202C/208A 115k SNPs - Probability Difference Day 1 - n= 1066
Harlan-206 102k SNPs - Probability Difference Day 1 - n= 758
Harlan-217 119k SNPs - Probability Difference Day 1 - n= 351
Charles River-R09_P03-07-10_C71_K92 221k SNPs - Probability Difference Day 1 - n= 425
Charles River-R04 214k SNPs - Probability Difference Day 1 - n= 650
Charles River-P09 223k SNPs - Probability Difference Day 1 - n= 295
Charles River-C72 222k SNPs - Probability Difference Day 1 - n= 358
-log10(P value)
Chromosomal Position

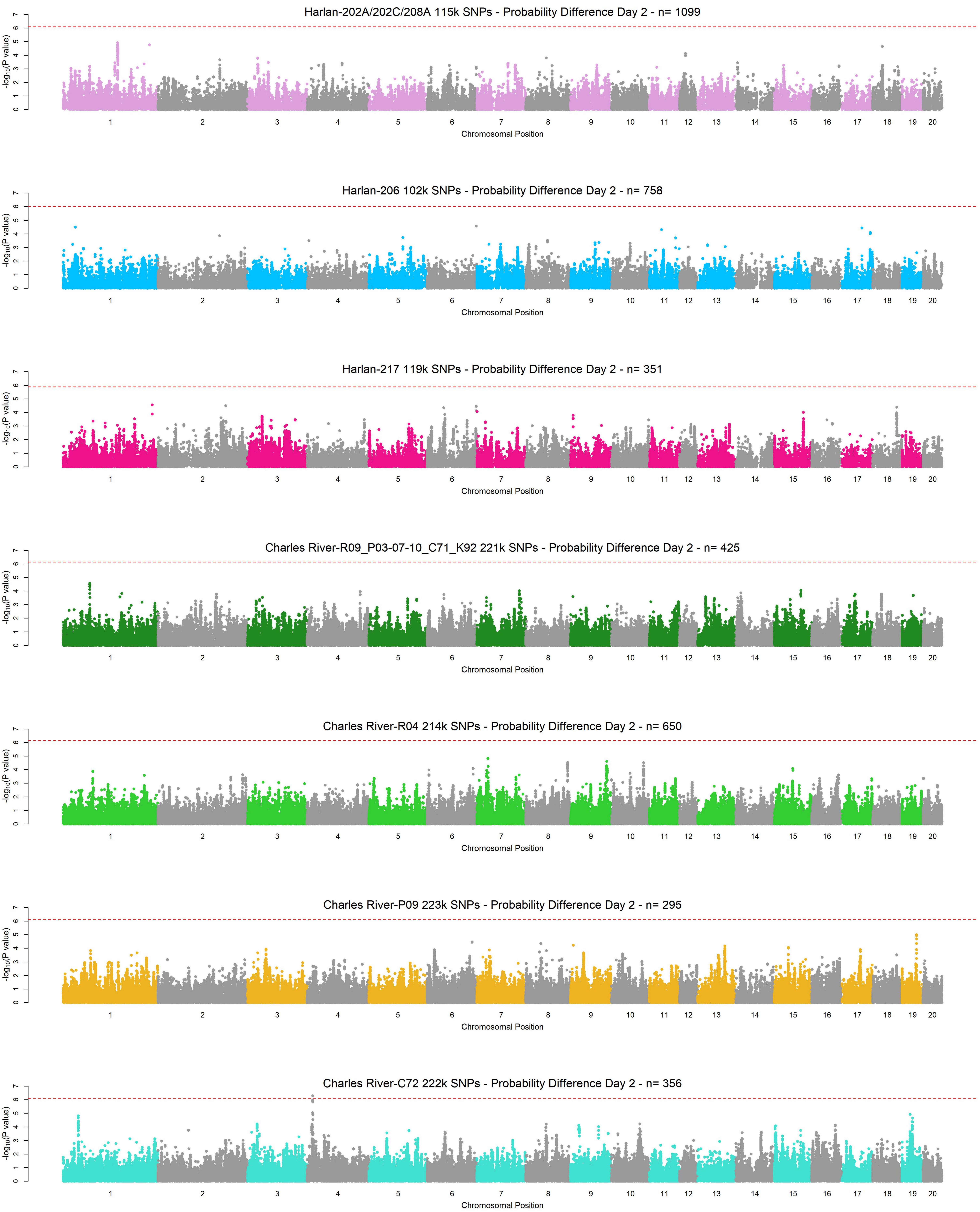
Harlan-202A/202C/208A 115k SNPs - Probability Difference Day 2 - n= 1099
Harlan-206 102k SNPs - Probability Difference Day 2 - n= 758
Harlan-217 119k SNPs - Probability Difference Day 2 - n= 351
Charles River-R09_P03-07-10_C71_K92 221k SNPs - Probability Difference Day 2 - n= 425
Charles River-R04 214k SNPs - Probability Difference Day 2 - n= 650
Charles River-P09 223k SNPs - Probability Difference Day 2 - n= 295
Charles River-C72 222k SNPs - Probability Difference Day 2 - n= 356
-log10(P value)
Chromosomal Position
1 2 3 4 5 6 7 8 9 10 11 12 13 14 15 16 17 18 19 20

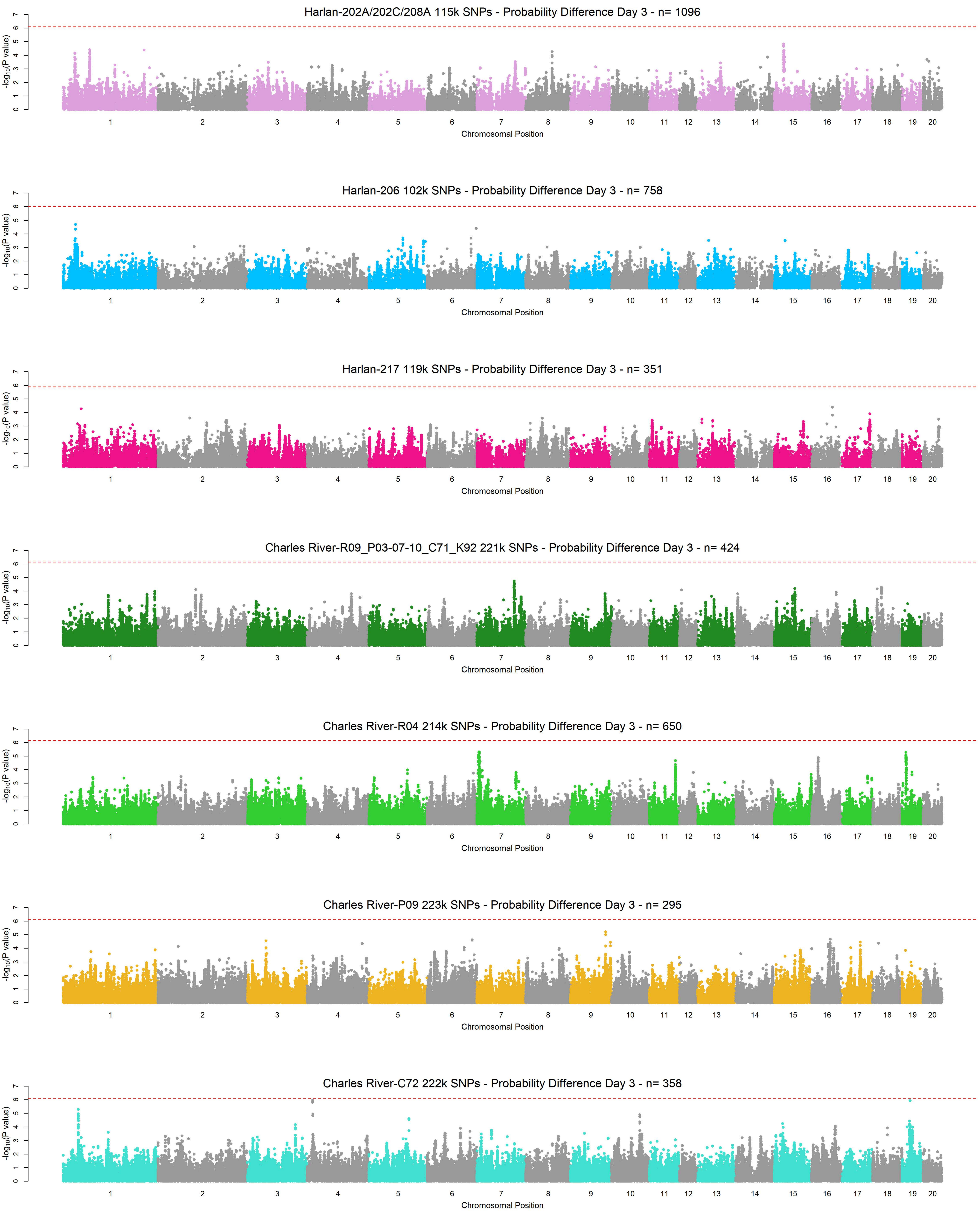
Harlan-202A/202C/208A 115k SNPs - Probability Difference Day 3 - n= 1096
Harlan-206 102k SNPs - Probability Difference Day 3 - n= 758
Harlan-217 119k SNPs - Probability Difference Day 3 - n= 351
Charles River-R09_P03-07-10_C71_K92 221k SNPs - Probability Difference Day 3 - n= 424
Charles River-R04 214k SNPs - Probability Difference Day 3 - n= 650
Charles River-P09 223k SNPs - Probability Difference Day 3 - n= 295
Charles River-C72 222k SNPs - Probability Difference Day 3 - n= 358
-log10(P value)
Chromosomal Position
1 2 3 4 5 6 7 8 9 10 11 12 13 14 15 16 17 18 19 20

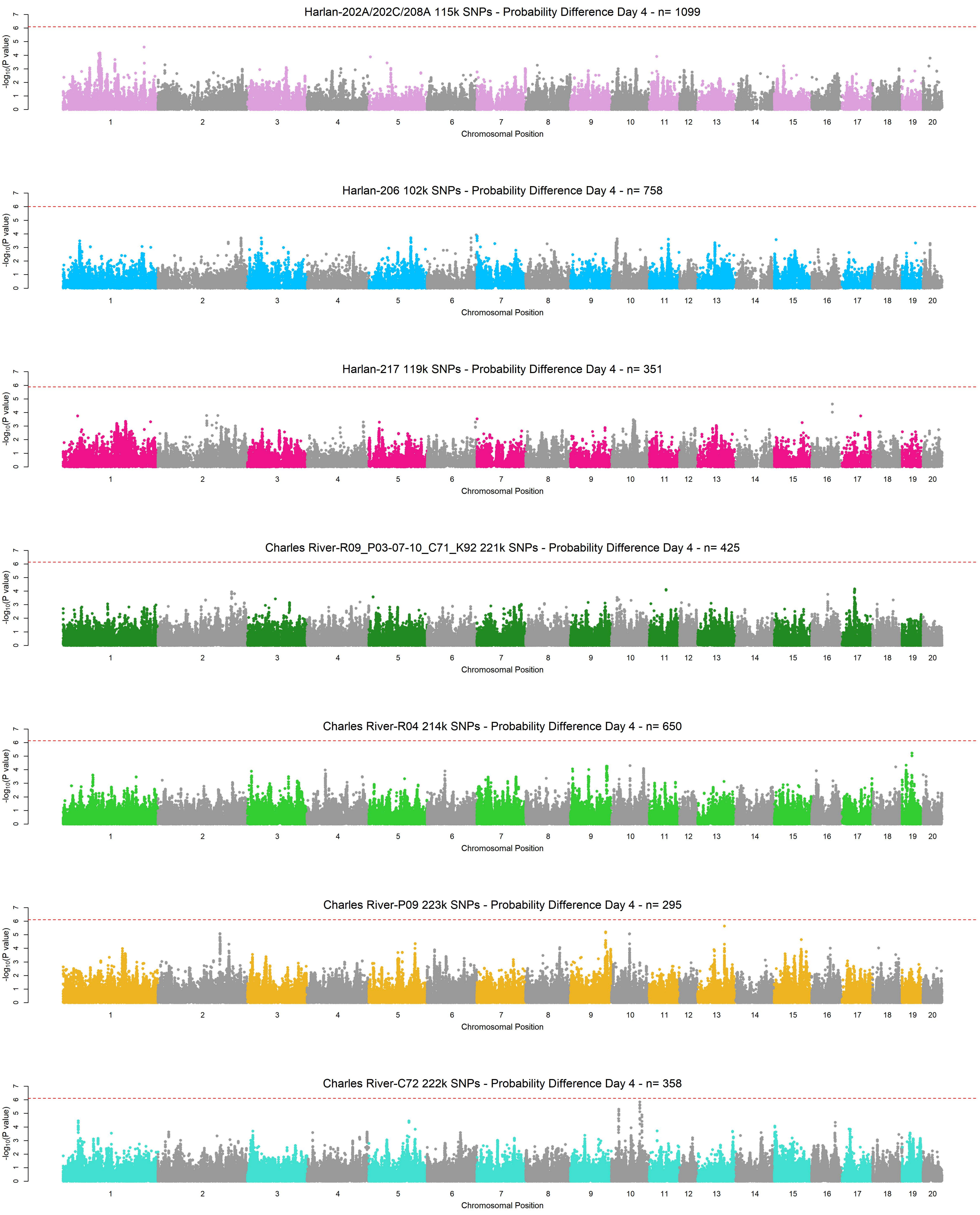
Harlan-202A/202C/208A 115k SNPs - Probability Difference Day 4 - n= 1099
Harlan-206 102k SNPs - Probability Difference Day 4 - n= 758
Harlan-217 119k SNPs - Probability Difference Day 4 - n= 351
Charles River-R09_P03-07-10_C71_K92 221k SNPs - Probability Difference Day 4 - n= 425
Charles River-R04 214k SNPs - Probability Difference Day 4 - n= 650
Charles River-P09 223k SNPs - Probability Difference Day 4 - n= 295
Charles River-C72 222k SNPs - Probability Difference Day 4 - n= 358
-log10(P value)
0 1 2 3 4 5 6 7
1 2 3 4 5 6 7 8 9 10 11 12 13 14 15 16 17 18 19 20
Chromosomal Position

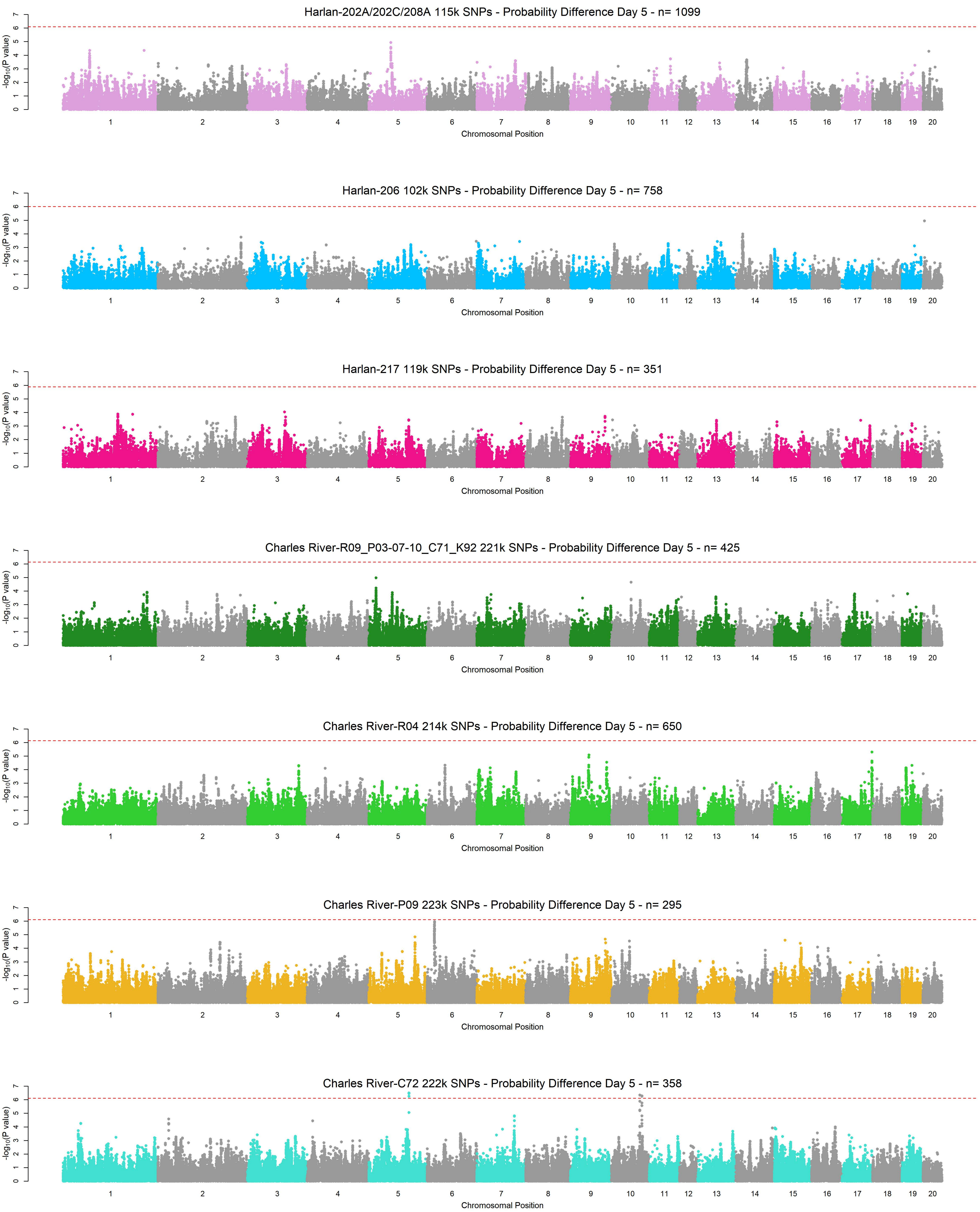

Harlan-202A/202C/208A 115k SNPs - Probability Difference Day 5 - n= 1099
Harlan-206 102k SNPs - Probability Difference Day 5 - n= 758
Harlan-217 119k SNPs - Probability Difference Day 5 - n= 351
Charles River-R09_P03-07-10_C71_K92 221k SNPs - Probability Difference Day 5 - n= 425
Charles River-R04 214k SNPs - Probability Difference Day 5 - n= 650
Charles River-P09 223k SNPs - Probability Difference Day 5 - n= 295
Charles River-C72 222k SNPs - Probability Difference Day 5 - n= 358
-log10(P value)
Chromosomal Position

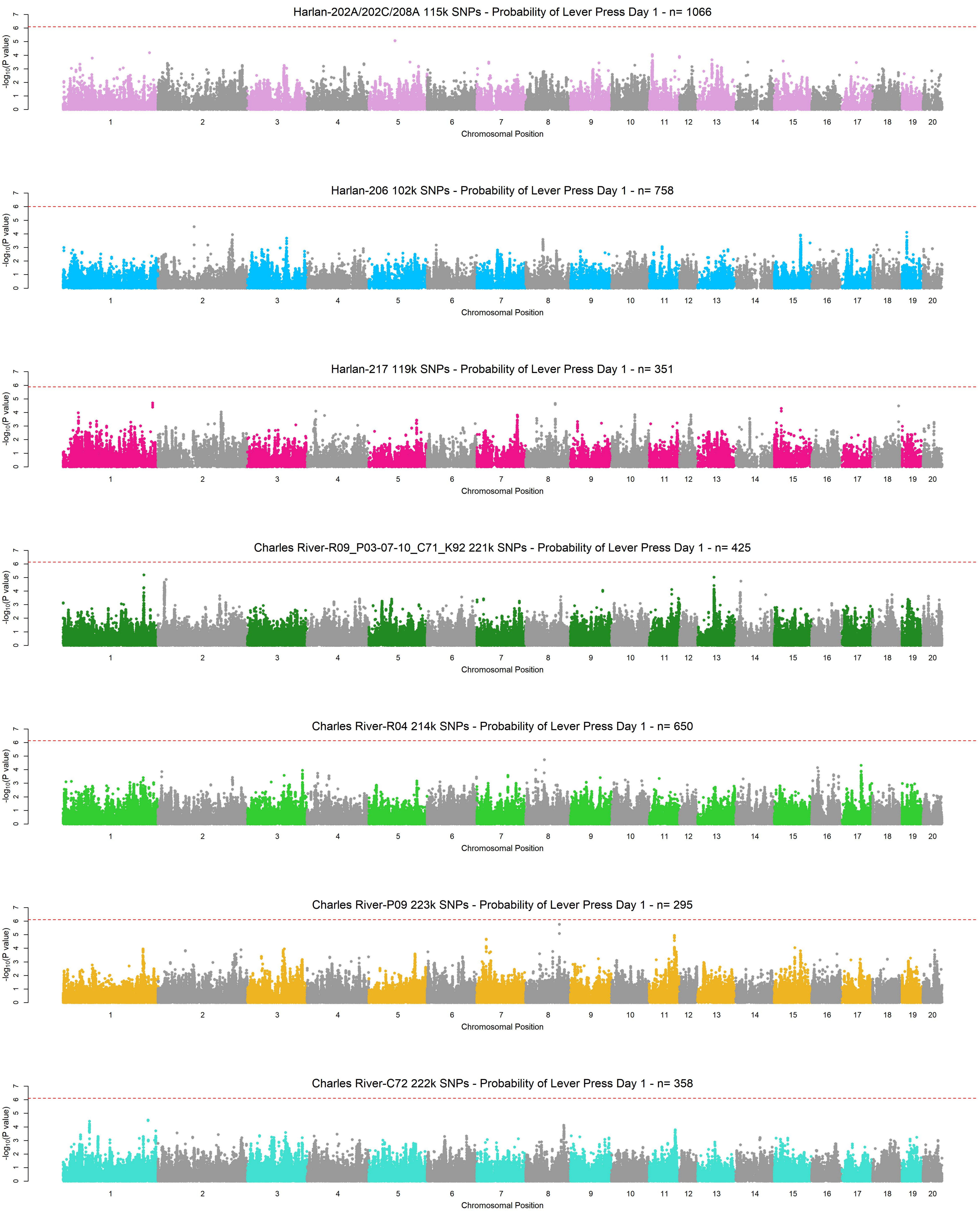
Harlan-202A/202C/208A 115k SNPs - Probability of Lever Press Day 1 - n= 1066
Harlan-206 102k SNPs - Probability of Lever Press Day 1 - n= 758
Harlan-217 119k SNPs - Probability of Lever Press Day 1 - n= 351
Charles River-R09_P03-07-10_C71_K92 221k SNPs - Probability of Lever Press Day 1 - n= 425
Charles River-R04 214k SNPs - Probability of Lever Press Day 1 - n= 650
Charles River-P09 223k SNPs - Probability of Lever Press Day 1 - n= 295
Charles River-C72 222k SNPs - Probability of Lever Press Day 1 - n= 358
-log10(P value)
Chromosomal Position
1 2 3 4 5 6 7 8 9 10 11 12 13 14 15 16 17 18 19 20
0 1 2 3 4 5 6 7

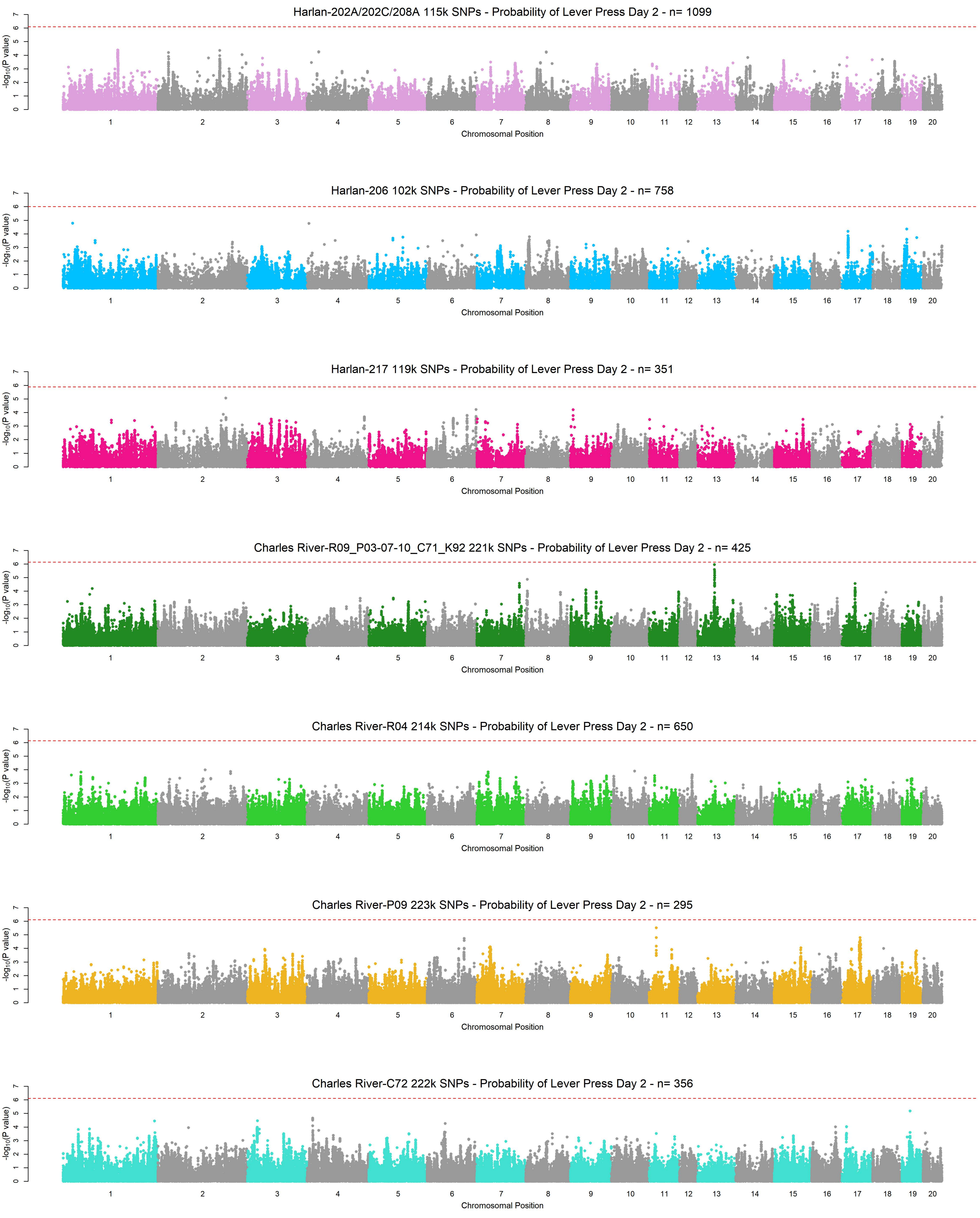
Harlan-202A/202C/208A 115k SNPs - Probability of Lever Press Day 2 - n= 1099
Harlan-206 102k SNPs - Probability of Lever Press Day 2 - n= 758
Harlan-217 119k SNPs - Probability of Lever Press Day 2 - n= 351
Charles River-R09_P03-07-10_C71_K92 221k SNPs - Probability of Lever Press Day 2 - n= 425
Charles River-R04 214k SNPs - Probability of Lever Press Day 2 - n= 650
Charles River-P09 223k SNPs - Probability of Lever Press Day 2 - n= 295
Charles River-C72 222k SNPs - Probability of Lever Press Day 2 - n= 356
-log10(P value)
0 1 2 3 4 5 6 7
1 2 3 4 5 6 7 8 9 10 11 12 13 14 15 16 17 18 19 20
Chromosomal Position

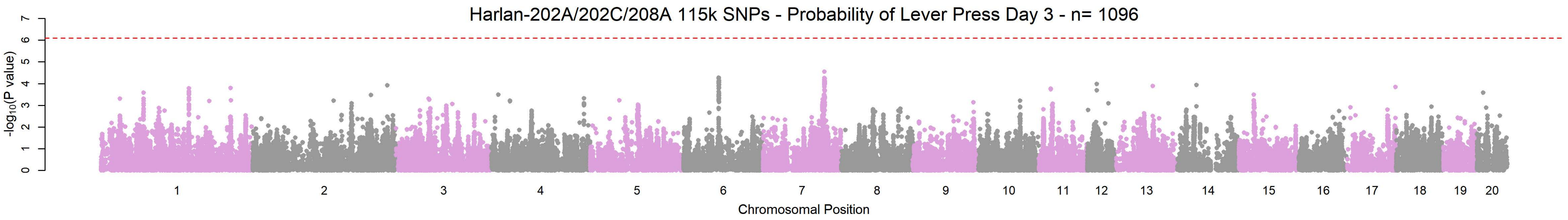
Harlan-202A/202C/208A 115k SNPs - Probability of Lever Press Day 3 - n= 1096
$-\log_{10}$(P value)
0 1 2 3 4 5 6 7
1 2 3 4 5 6 7 8 9 10 11 12 13 14 15 16 17 18 19 20
Chromosomal Position

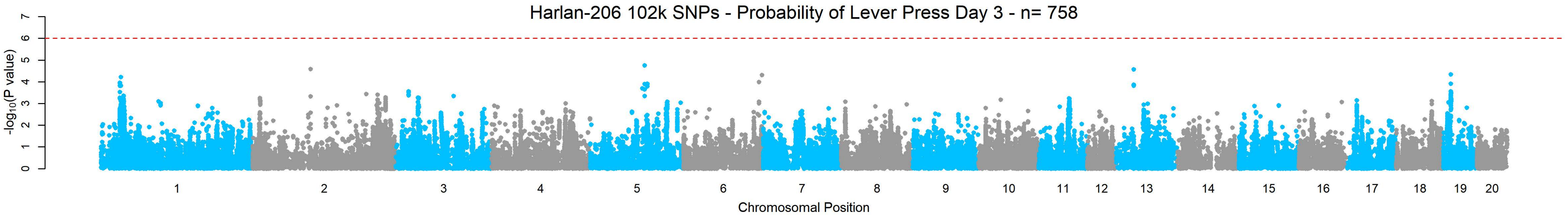
Harlan-206 102k SNPs - Probability of Lever Press Day 3 - n= 758
$-\log_{10}$(P value)
0 1 2 3 4 5 6 7
1 2 3 4 5 6 7 8 9 10 11 12 13 14 15 16 17 18 19 20
Chromosomal Position

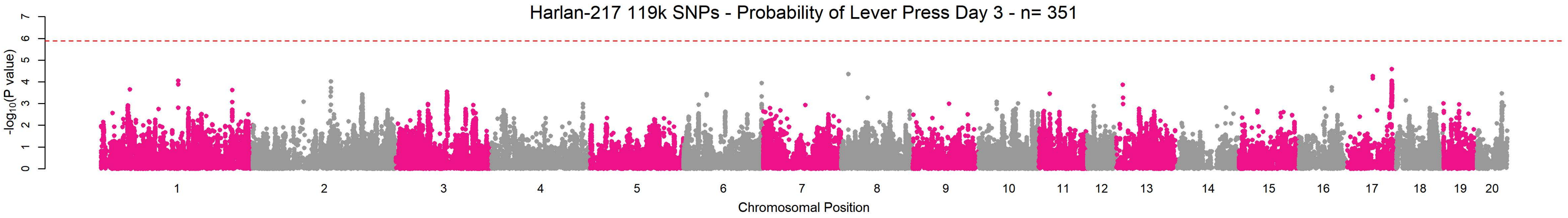
Harlan-217 119k SNPs - Probability of Lever Press Day 3 - n= 351
$-\log_{10}$(P value)
0 1 2 3 4 5 6 7
1 2 3 4 5 6 7 8 9 10 11 12 13 14 15 16 17 18 19 20
Chromosomal Position

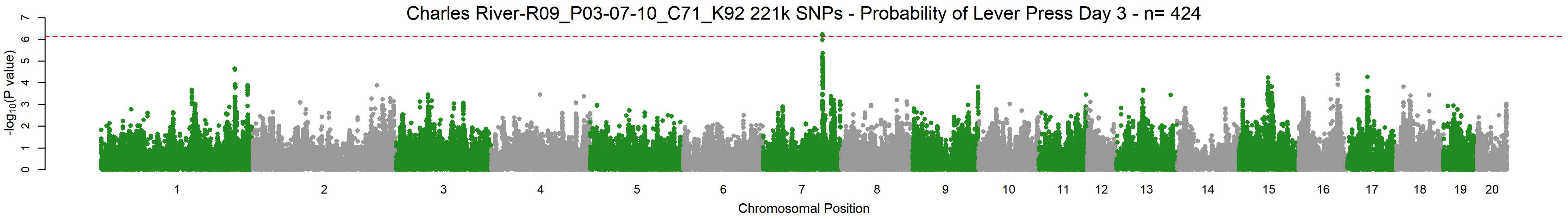
Charles River-R09_P03-07-10_C71_K92 221k SNPs - Probability of Lever Press Day 3 - n= 424
$-\log_{10}$(P value)
0 1 2 3 4 5 6 7
1 2 3 4 5 6 7 8 9 10 11 12 13 14 15 16 17 18 19 20
Chromosomal Position

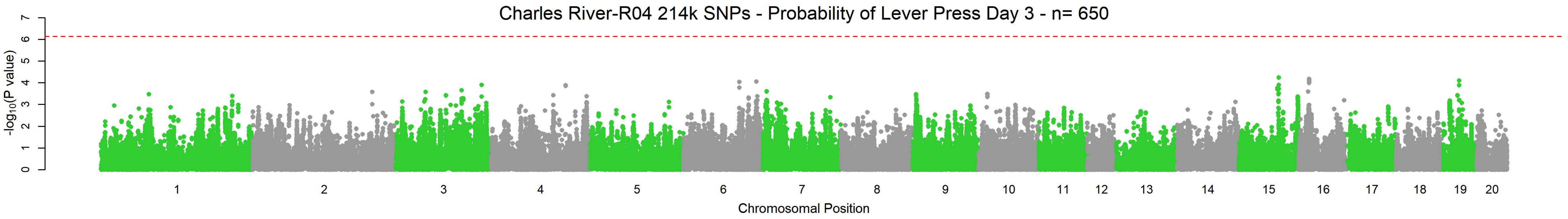
Charles River-R04 214k SNPs - Probability of Lever Press Day 3 - n= 650
$-\log_{10}$(P value)
0 1 2 3 4 5 6 7
1 2 3 4 5 6 7 8 9 10 11 12 13 14 15 16 17 18 19 20
Chromosomal Position

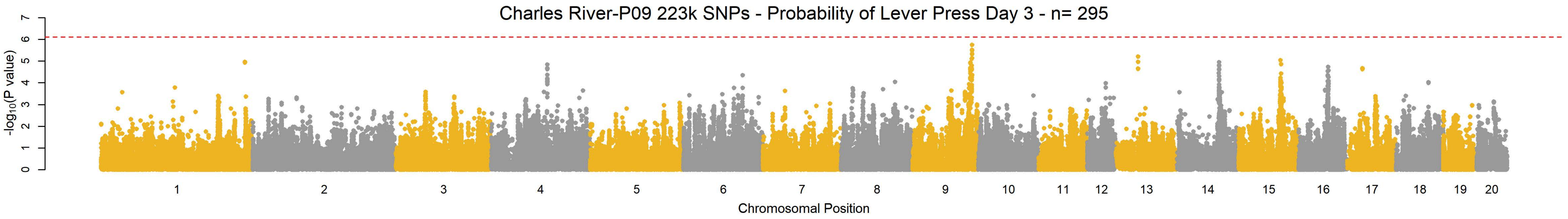
Charles River-P09 223k SNPs - Probability of Lever Press Day 3 - n= 295
$-\log_{10}$(P value)
0 1 2 3 4 5 6 7
1 2 3 4 5 6 7 8 9 10 11 12 13 14 15 16 17 18 19 20
Chromosomal Position

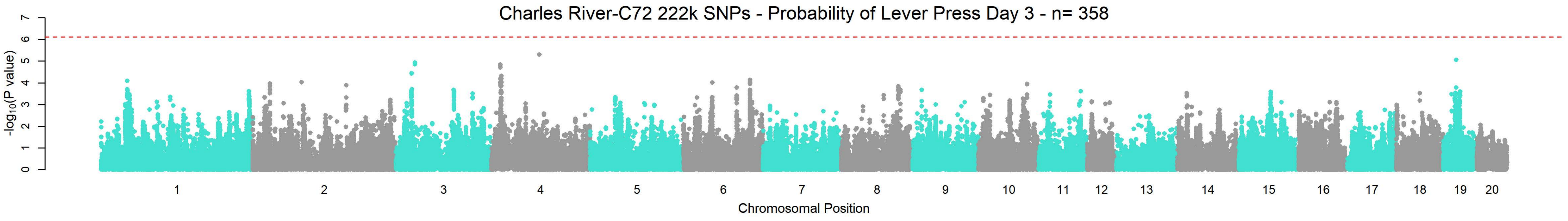
Charles River-C72 222k SNPs - Probability of Lever Press Day 3 - n= 358
$-\log_{10}$(P value)
0 1 2 3 4 5 6 7
1 2 3 4 5 6 7 8 9 10 11 12 13 14 15 16 17 18 19 20
Chromosomal Position

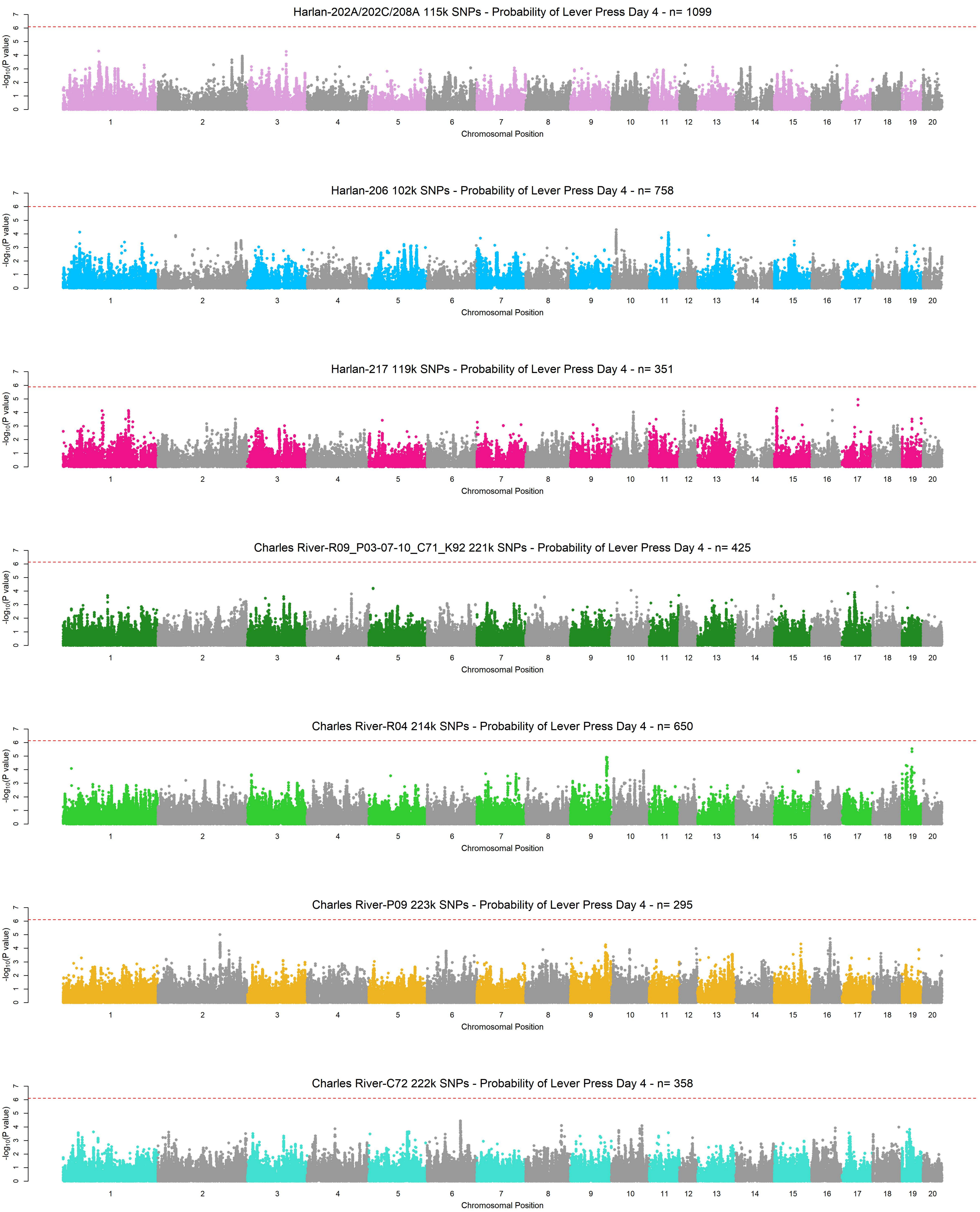
Harlan-202A/202C/208A 115k SNPs - Probability of Lever Press Day 4 - n= 1099
-log10(P value)
0 1 2 3 4 5 6 7
1 2 3 4 5 6 7 8 9 10 11 12 13 14 15 16 17 18 19 20
Chromosomal Position
Harlan-206 102k SNPs - Probability of Lever Press Day 4 - n= 758
-log10(P value)
0 1 2 3 4 5 6 7
1 2 3 4 5 6 7 8 9 10 11 12 13 14 15 16 17 18 19 20
Chromosomal Position
Harlan-217 119k SNPs - Probability of Lever Press Day 4 - n= 351
-log10(P value)
0 1 2 3 4 5 6 7
1 2 3 4 5 6 7 8 9 10 11 12 13 14 15 16 17 18 19 20
Chromosomal Position
Charles River-R09_P03-07-10_C71_K92 221k SNPs - Probability of Lever Press Day 4 - n= 425
-log10(P value)
0 1 2 3 4 5 6 7
1 2 3 4 5 6 7 8 9 10 11 12 13 14 15 16 17 18 19 20
Chromosomal Position
Charles River-R04 214k SNPs - Probability of Lever Press Day 4 - n= 650
-log10(P value)
0 1 2 3 4 5 6 7
1 2 3 4 5 6 7 8 9 10 11 12 13 14 15 16 17 18 19 20
Chromosomal Position
Charles River-P09 223k SNPs - Probability of Lever Press Day 4 - n= 295
-log10(P value)
0 1 2 3 4 5 6 7
1 2 3 4 5 6 7 8 9 10 11 12 13 14 15 16 17 18 19 20
Chromosomal Position
Charles River-C72 222k SNPs - Probability of Lever Press Day 4 - n= 358
-log10(P value)
0 1 2 3 4 5 6 7
1 2 3 4 5 6 7 8 9 10 11 12 13 14 15 16 17 18 19 20
Chromosomal Position

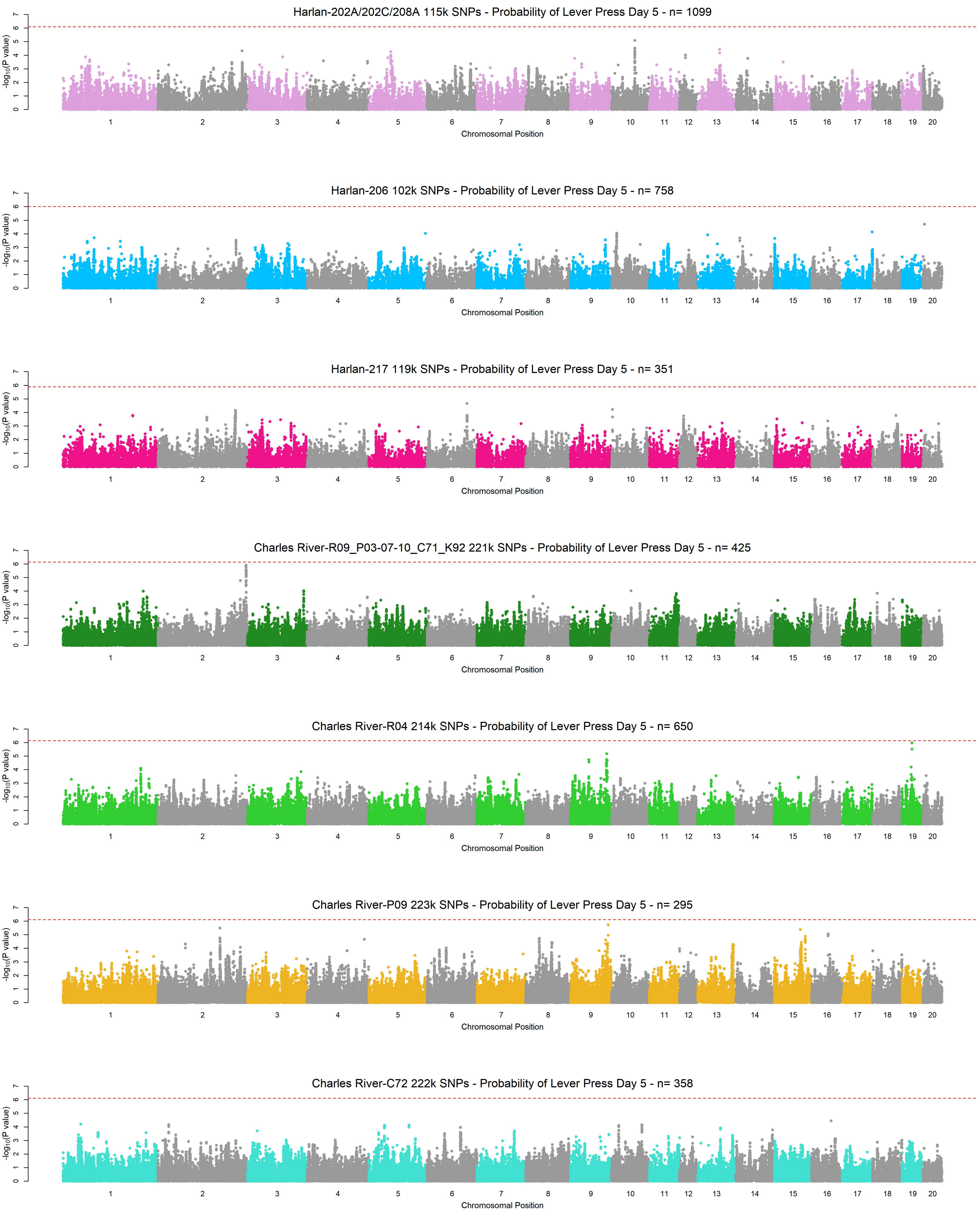
Harlan-202A/202C/208A 115k SNPs - Probability of Lever Press Day 5 - n= 1099
Harlan-206 102k SNPs - Probability of Lever Press Day 5 - n= 758
Harlan-217 119k SNPs - Probability of Lever Press Day 5 - n= 351
Charles River-R09_P03-07-10_C71_K92 221k SNPs - Probability of Lever Press Day 5 - n= 425
Charles River-R04 214k SNPs - Probability of Lever Press Day 5 - n= 650
Charles River-P09 223k SNPs - Probability of Lever Press Day 5 - n= 295
Charles River-C72 222k SNPs - Probability of Lever Press Day 5 - n= 358
-log10(P value)
Chromosomal Position
1 2 3 4 5 6 7 8 9 10 11 12 13 14 15 16 17 18 19 20
0 1 2 3 4 5 6 7

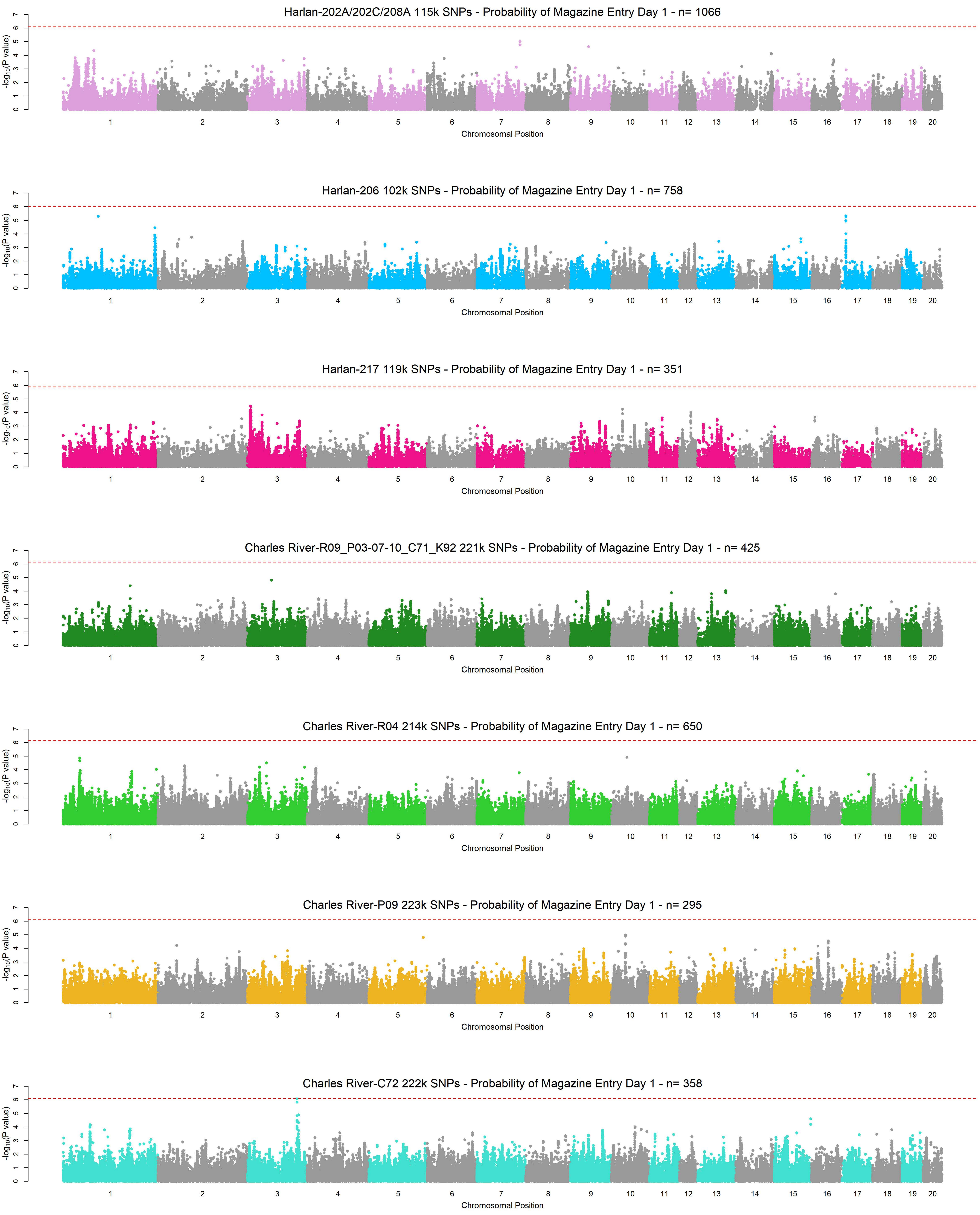
Harlan-202A/202C/208A 115k SNPs - Probability of Magazine Entry Day 1 - n= 1066
Harlan-206 102k SNPs - Probability of Magazine Entry Day 1 - n= 758
Harlan-217 119k SNPs - Probability of Magazine Entry Day 1 - n= 351
Charles River-R09_P03-07-10_C71_K92 221k SNPs - Probability of Magazine Entry Day 1 - n= 425
Charles River-R04 214k SNPs - Probability of Magazine Entry Day 1 - n= 650
Charles River-P09 223k SNPs - Probability of Magazine Entry Day 1 - n= 295
Charles River-C72 222k SNPs - Probability of Magazine Entry Day 1 - n= 358
-log10(P value)
Chromosomal Position
0 1 2 3 4 5 6 7
1 2 3 4 5 6 7 8 9 10 11 12 13 14 15 16 17 18 19 20

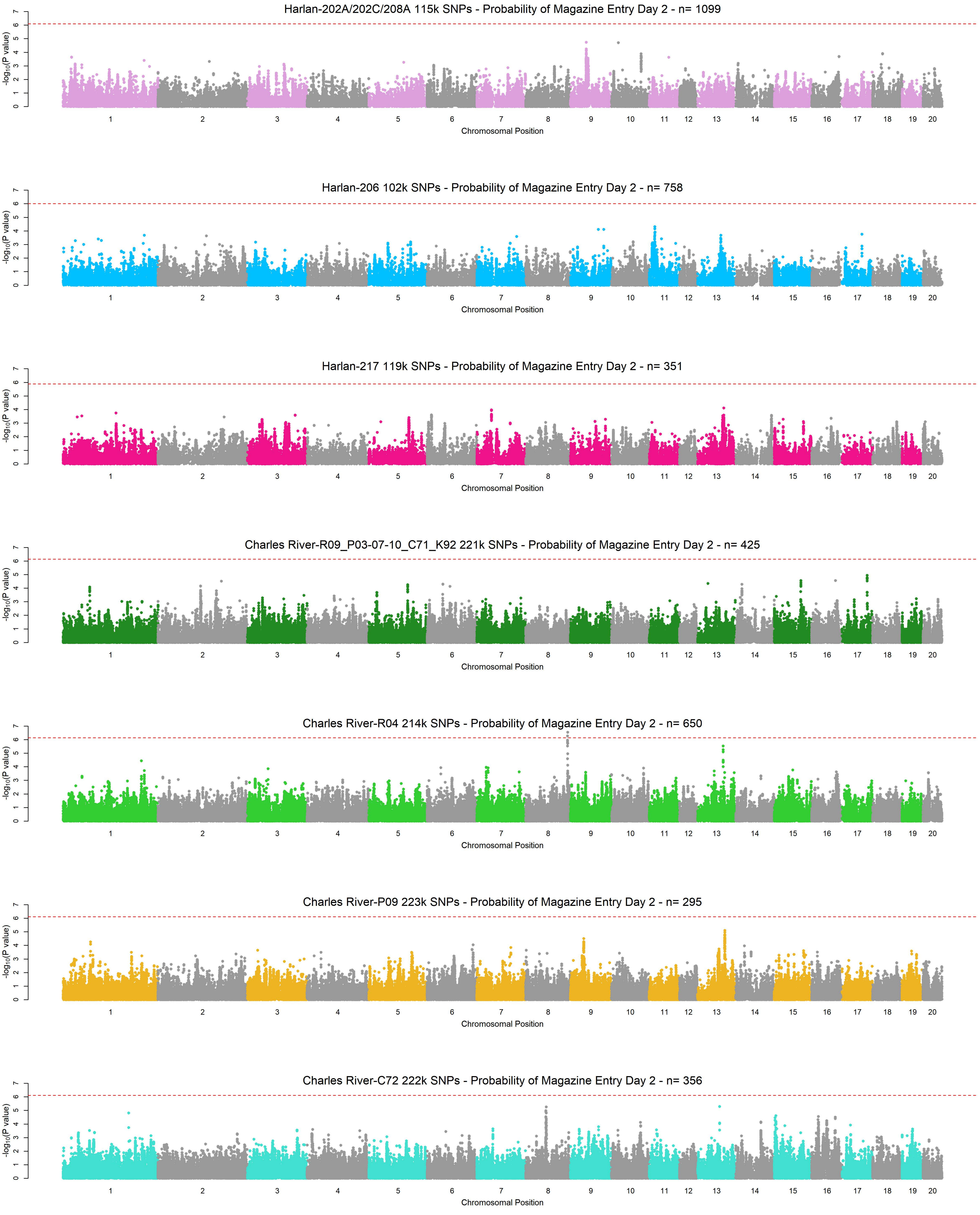
Harlan-202A/202C/208A 115k SNPs - Probability of Magazine Entry Day 2 - n= 1099
Harlan-206 102k SNPs - Probability of Magazine Entry Day 2 - n= 758
Harlan-217 119k SNPs - Probability of Magazine Entry Day 2 - n= 351
Charles River-R09_P03-07-10_C71_K92 221k SNPs - Probability of Magazine Entry Day 2 - n= 425
Charles River-R04 214k SNPs - Probability of Magazine Entry Day 2 - n= 650
Charles River-P09 223k SNPs - Probability of Magazine Entry Day 2 - n= 295
Charles River-C72 222k SNPs - Probability of Magazine Entry Day 2 - n= 356
$-\log_{10}$(P value)
Chromosomal Position
1 2 3 4 5 6 7 8 9 10 11 12 13 14 15 16 17 18 19 20
0 1 2 3 4 5 6 7

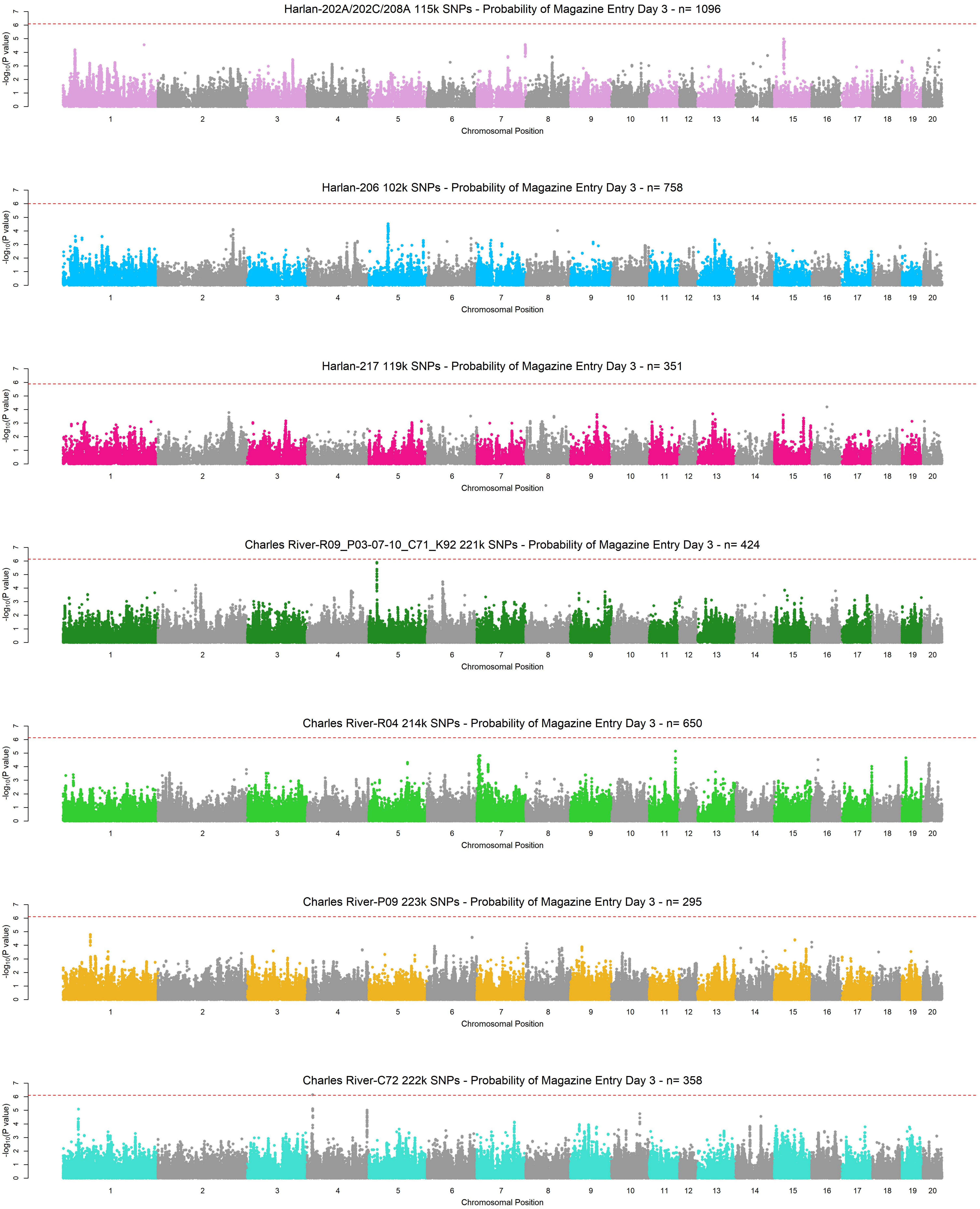
Harlan-202A/202C/208A 115k SNPs - Probability of Magazine Entry Day 3 - n= 1096
Harlan-206 102k SNPs - Probability of Magazine Entry Day 3 - n= 758
Harlan-217 119k SNPs - Probability of Magazine Entry Day 3 - n= 351
Charles River-R09_P03-07-10_C71_K92 221k SNPs - Probability of Magazine Entry Day 3 - n= 424
Charles River-R04 214k SNPs - Probability of Magazine Entry Day 3 - n= 650
Charles River-P09 223k SNPs - Probability of Magazine Entry Day 3 - n= 295
Charles River-C72 222k SNPs - Probability of Magazine Entry Day 3 - n= 358
-log10(P value)
0 1 2 3 4 5 6 7
1 2 3 4 5 6 7 8 9 10 11 12 13 14 15 16 17 18 19 20
Chromosomal Position

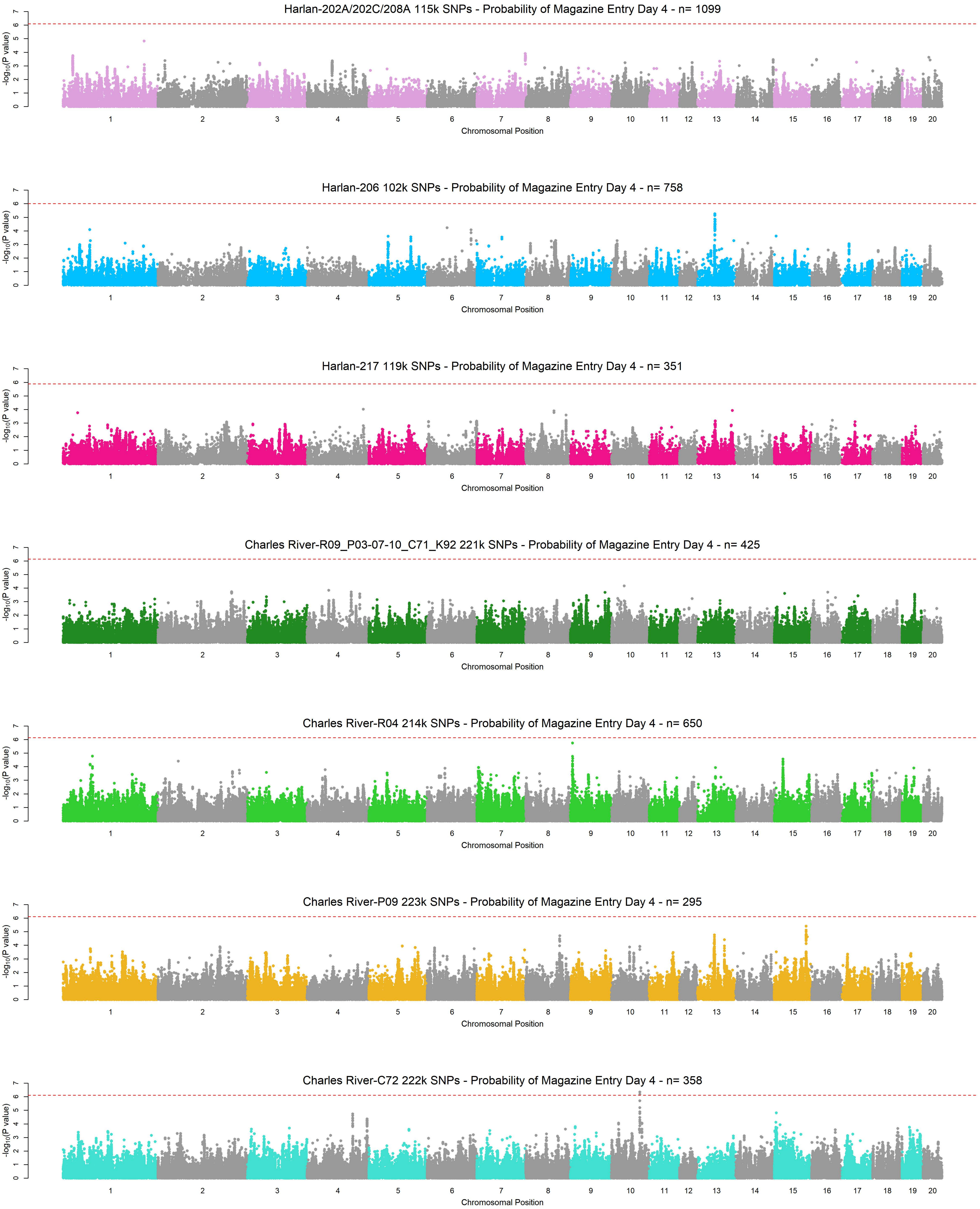
Harlan-202A/202C/208A 115k SNPs - Probability of Magazine Entry Day 4 - n= 1099
Harlan-206 102k SNPs - Probability of Magazine Entry Day 4 - n= 758
Harlan-217 119k SNPs - Probability of Magazine Entry Day 4 - n= 351
Charles River-R09_P03-07-10_C71_K92 221k SNPs - Probability of Magazine Entry Day 4 - n= 425
Charles River-R04 214k SNPs - Probability of Magazine Entry Day 4 - n= 650
Charles River-P09 223k SNPs - Probability of Magazine Entry Day 4 - n= 295
Charles River-C72 222k SNPs - Probability of Magazine Entry Day 4 - n= 358
$-\log_{10}$(P value)
Chromosomal Position
1 2 3 4 5 6 7 8 9 10 11 12 13 14 15 16 17 18 19 20

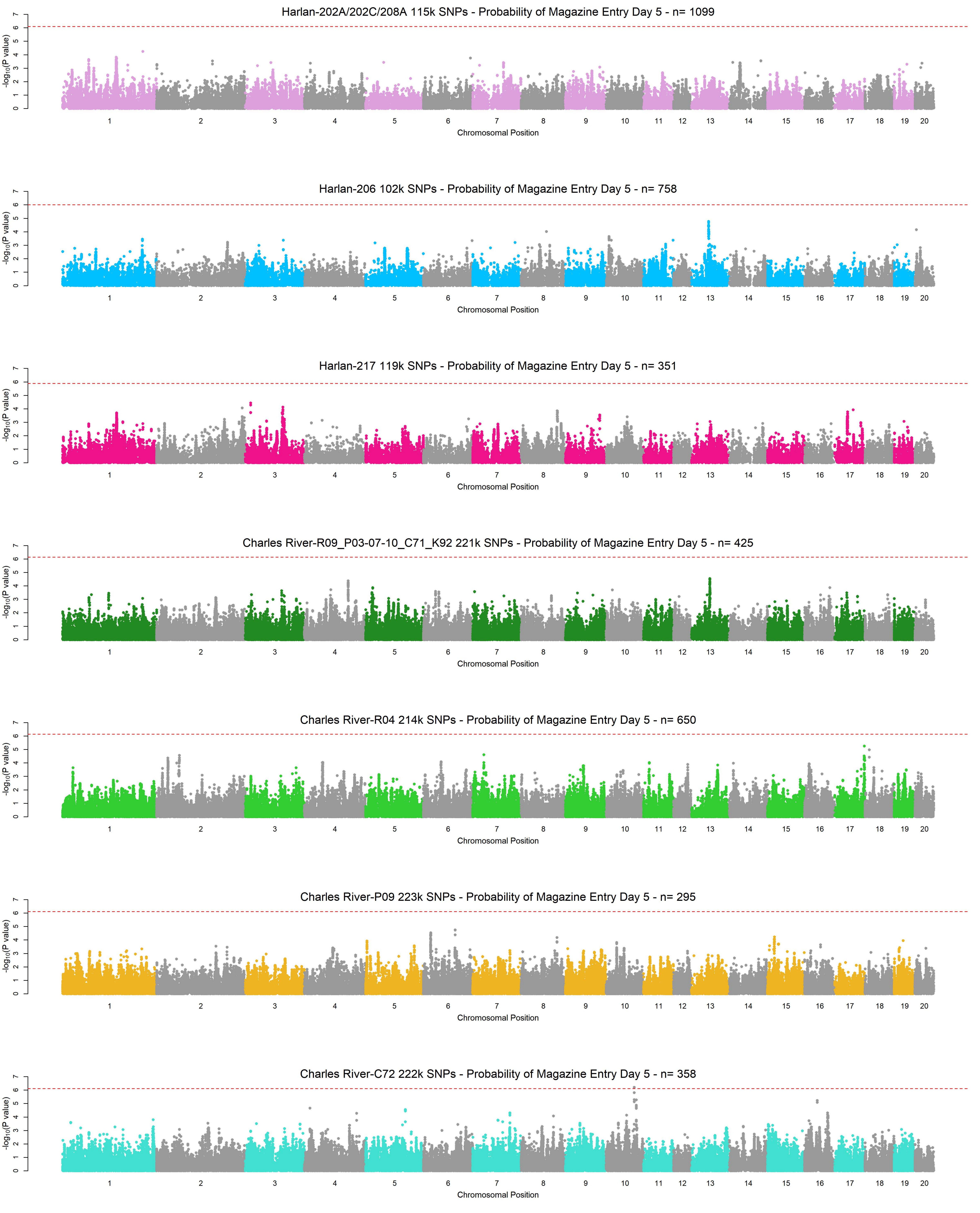
Harlan-202A/202C/208A 115k SNPs - Probability of Magazine Entry Day 5 - n= 1099
Harlan-206 102k SNPs - Probability of Magazine Entry Day 5 - n= 758
Harlan-217 119k SNPs - Probability of Magazine Entry Day 5 - n= 351
Charles River-R09_P03-07-10_C71_K92 221k SNPs - Probability of Magazine Entry Day 5 - n= 425
Charles River-R04 214k SNPs - Probability of Magazine Entry Day 5 - n= 650
Charles River-P09 223k SNPs - Probability of Magazine Entry Day 5 - n= 295
Charles River-C72 222k SNPs - Probability of Magazine Entry Day 5 - n= 358
-log10(P value)
Chromosomal Position
1 2 3 4 5 6 7 8 9 10 11 12 13 14 15 16 17 18 19 20

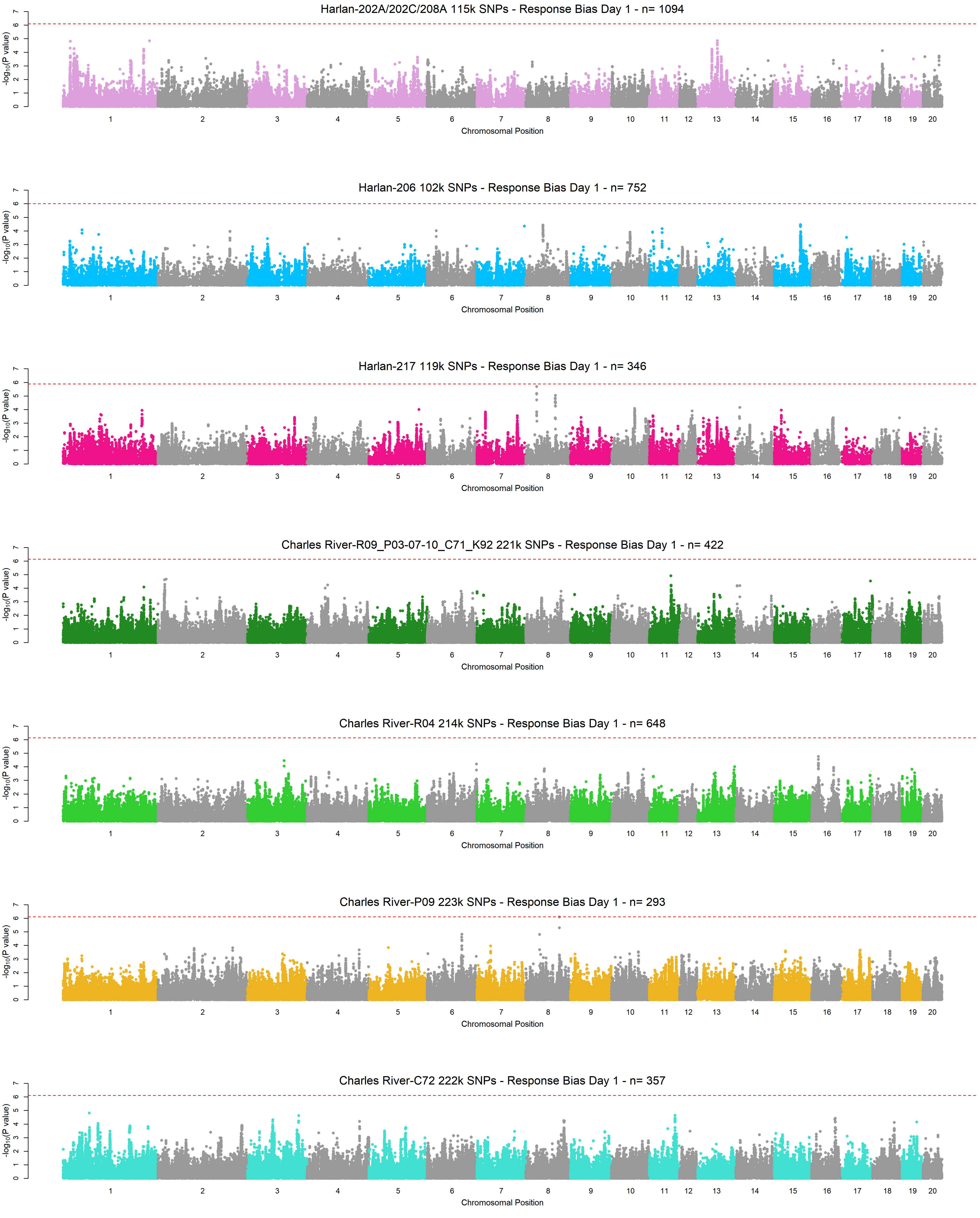
Harlan-202A/202C/208A 115k SNPs - Response Bias Day 1 - n= 1094
Harlan-206 102k SNPs - Response Bias Day 1 - n= 752
Harlan-217 119k SNPs - Response Bias Day 1 - n= 346
Charles River-R09_P03-07-10_C71_K92 221k SNPs - Response Bias Day 1 - n= 422
Charles River-R04 214k SNPs - Response Bias Day 1 - n= 648
Charles River-P09 223k SNPs - Response Bias Day 1 - n= 293
Charles River-C72 222k SNPs - Response Bias Day 1 - n= 357
-log10(P value)
Chromosomal Position
1 2 3 4 5 6 7 8 9 10 11 12 13 14 15 16 17 18 19 20
0 1 2 3 4 5 6 7

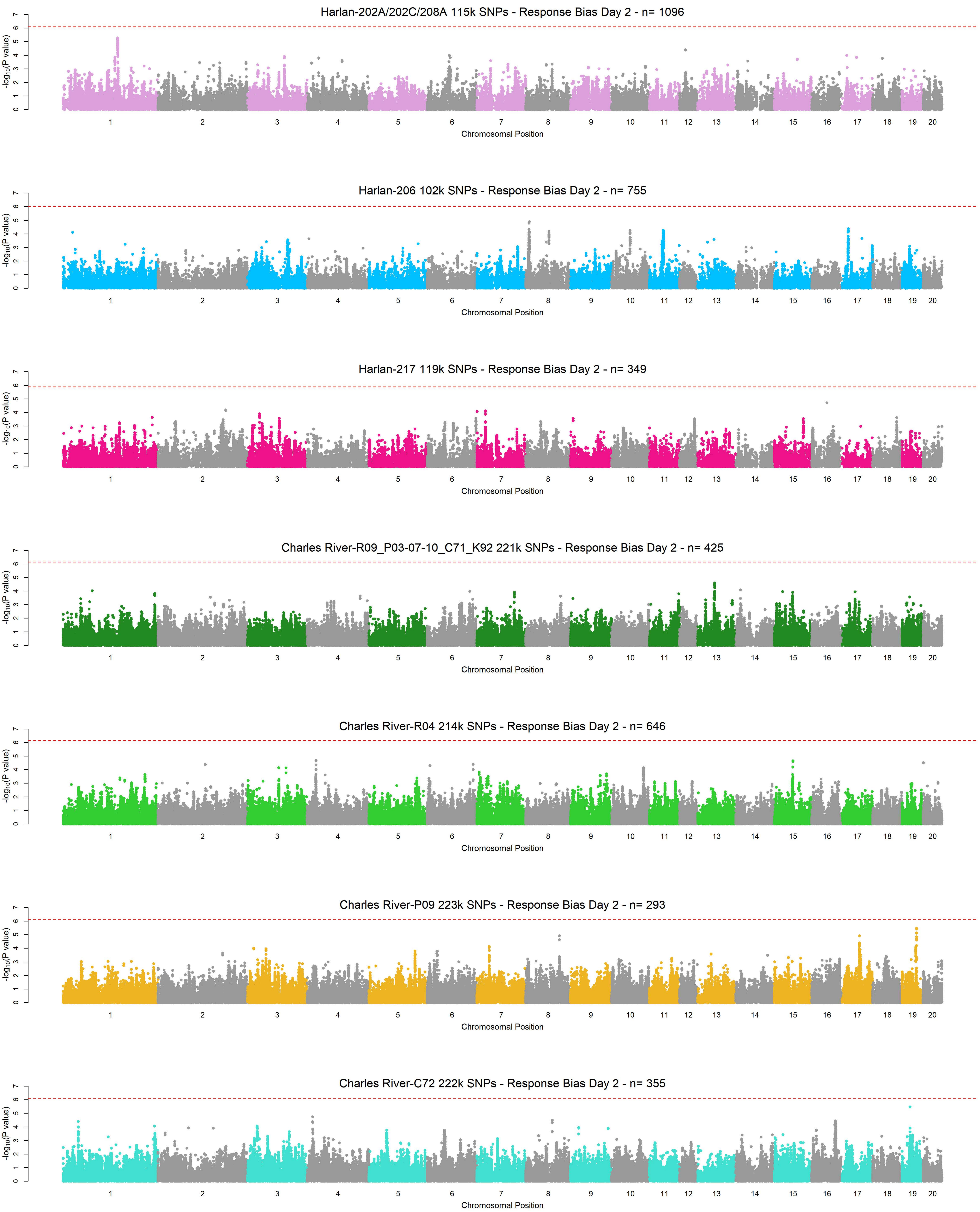
Harlan-202A/202C/208A 115k SNPs - Response Bias Day 2 - n= 1096
Harlan-206 102k SNPs - Response Bias Day 2 - n= 755
Harlan-217 119k SNPs - Response Bias Day 2 - n= 349
Charles River-R09_P03-07-10_C71_K92 221k SNPs - Response Bias Day 2 - n= 425
Charles River-R04 214k SNPs - Response Bias Day 2 - n= 646
Charles River-P09 223k SNPs - Response Bias Day 2 - n= 293
Charles River-C72 222k SNPs - Response Bias Day 2 - n= 355
-log10(P value)
0 1 2 3 4 5 6 7
1 2 3 4 5 6 7 8 9 10 11 12 13 14 15 16 17 18 19 20
Chromosomal Position

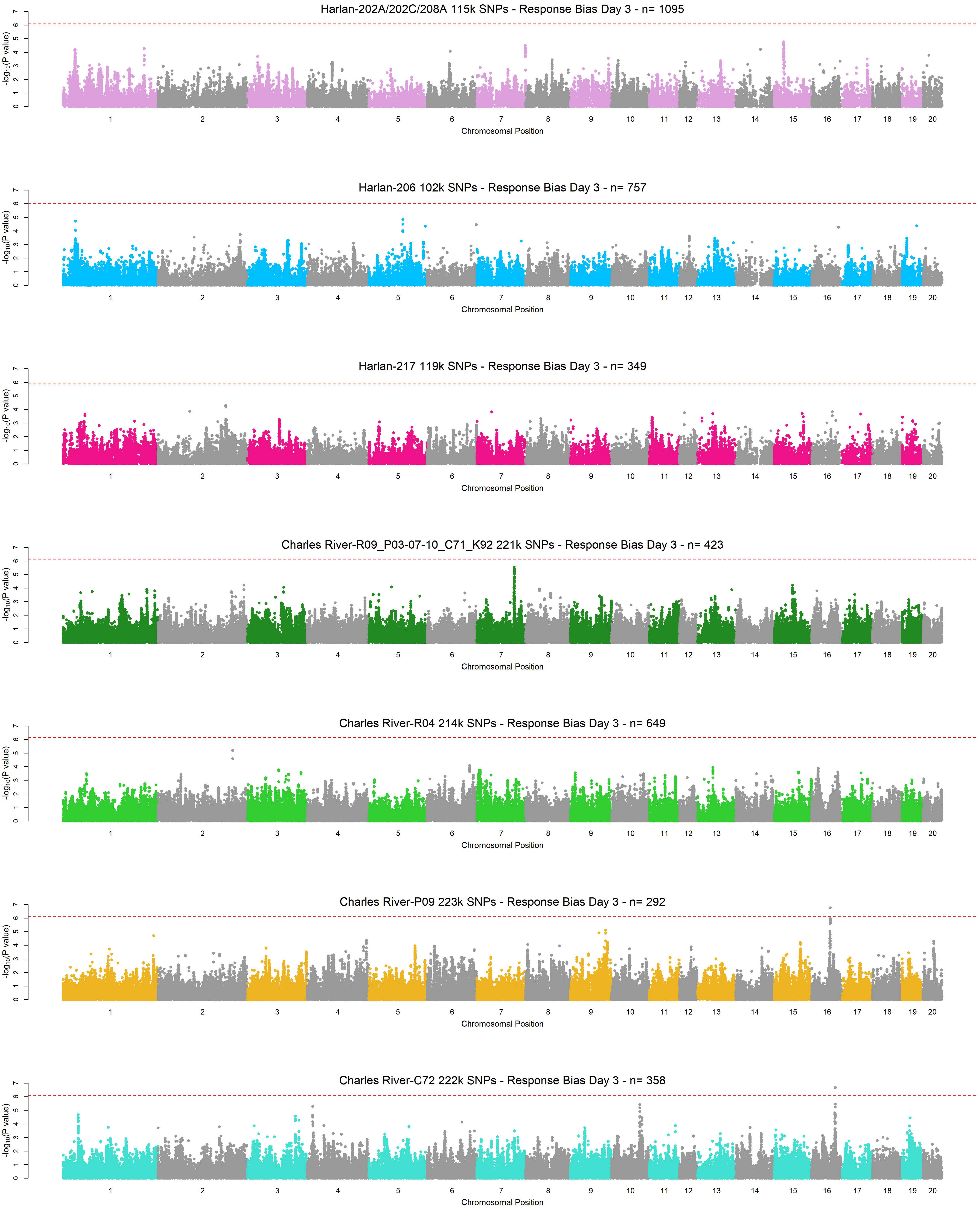
Harlan-202A/202C/208A 115k SNPs - Response Bias Day 3 - n= 1095
Harlan-206 102k SNPs - Response Bias Day 3 - n= 757
Harlan-217 119k SNPs - Response Bias Day 3 - n= 349
Charles River-R09_P03-07-10_C71_K92 221k SNPs - Response Bias Day 3 - n= 423
Charles River-R04 214k SNPs - Response Bias Day 3 - n= 649
Charles River-P09 223k SNPs - Response Bias Day 3 - n= 292
Charles River-C72 222k SNPs - Response Bias Day 3 - n= 358
-log10(P value)
Chromosomal Position
1 2 3 4 5 6 7 8 9 10 11 12 13 14 15 16 17 18 19 20
0 1 2 3 4 5 6 7

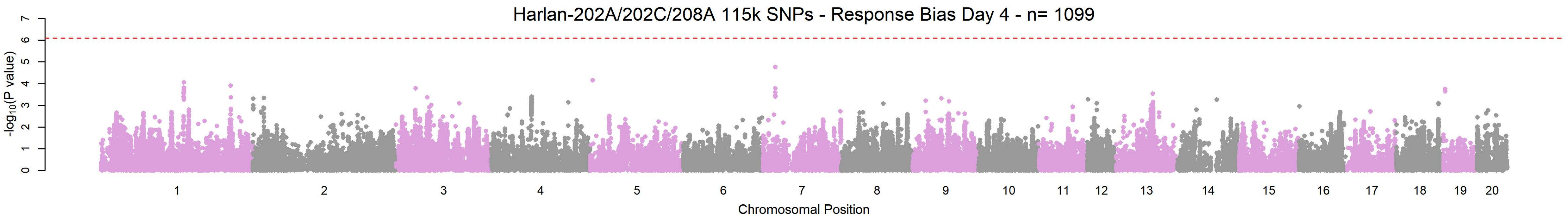
Harlan-202A/202C/208A 115k SNPs - Response Bias Day 4 - n= 1099
-log10(P value)
0 1 2 3 4 5 6 7
1 2 3 4 5 6 7 8 9 10 11 12 13 14 15 16 17 18 19 20
Chromosomal Position

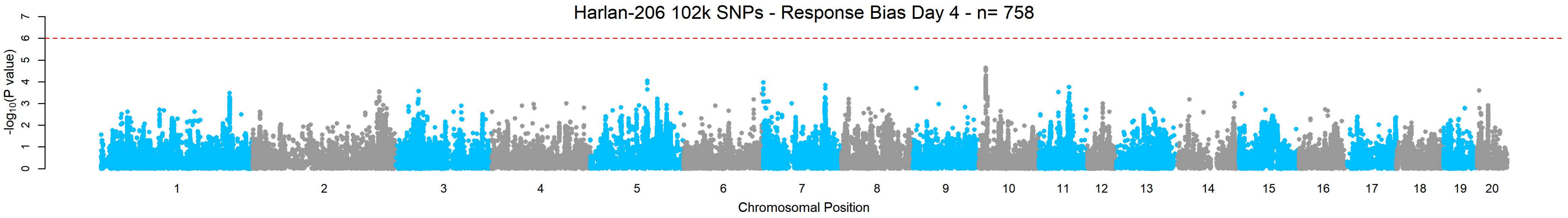
Harlan-206 102k SNPs - Response Bias Day 4 - n= 758
-log10(P value)
0 1 2 3 4 5 6 7
1 2 3 4 5 6 7 8 9 10 11 12 13 14 15 16 17 18 19 20
Chromosomal Position

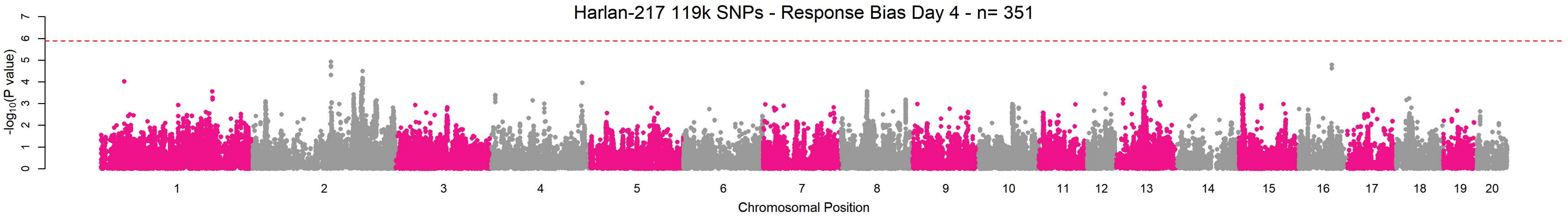
Harlan-217 119k SNPs - Response Bias Day 4 - n= 351
-log10(P value)
0 1 2 3 4 5 6 7
1 2 3 4 5 6 7 8 9 10 11 12 13 14 15 16 17 18 19 20
Chromosomal Position

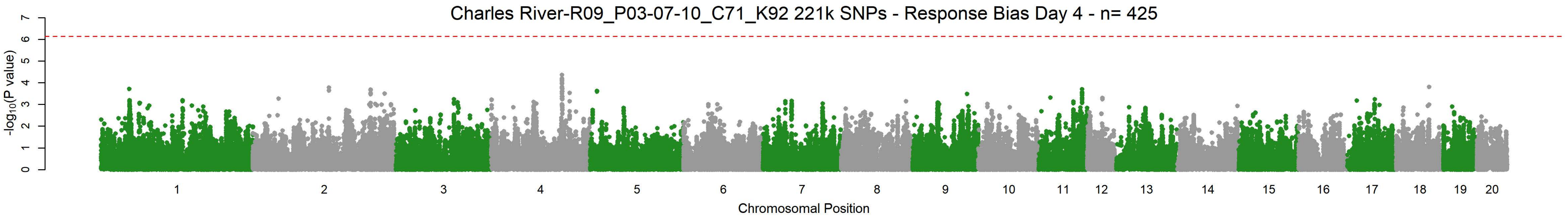
Charles River-R09_P03-07-10_C71_K92 221k SNPs - Response Bias Day 4 - n= 425
-log10(P value)
0 1 2 3 4 5 6 7
1 2 3 4 5 6 7 8 9 10 11 12 13 14 15 16 17 18 19 20
Chromosomal Position

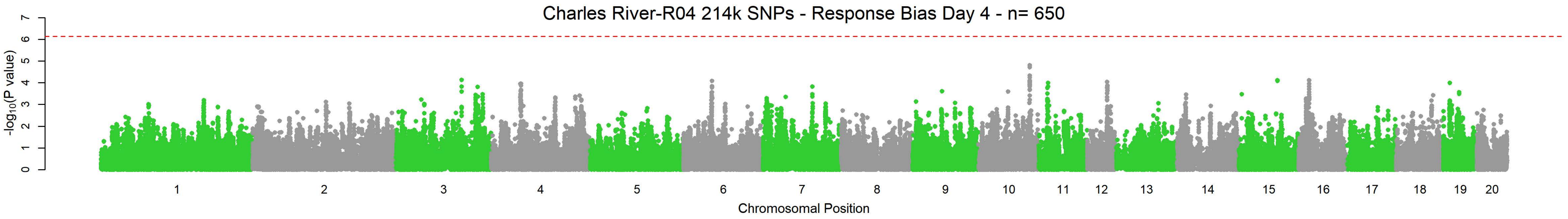
Charles River-R04 214k SNPs - Response Bias Day 4 - n= 650
-log10(P value)
0 1 2 3 4 5 6 7
1 2 3 4 5 6 7 8 9 10 11 12 13 14 15 16 17 18 19 20
Chromosomal Position

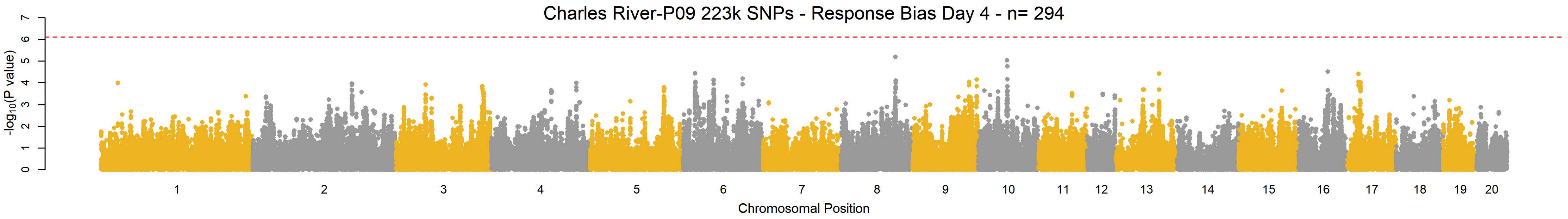
Charles River-P09 223k SNPs - Response Bias Day 4 - n= 294
-log10(P value)
0 1 2 3 4 5 6 7
1 2 3 4 5 6 7 8 9 10 11 12 13 14 15 16 17 18 19 20
Chromosomal Position

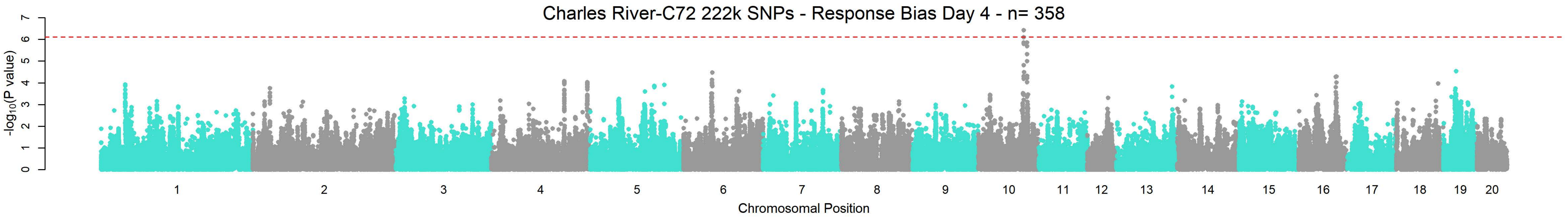
Charles River-C72 222k SNPs - Response Bias Day 4 - n= 358
-log10(P value)
0 1 2 3 4 5 6 7
1 2 3 4 5 6 7 8 9 10 11 12 13 14 15 16 17 18 19 20
Chromosomal Position

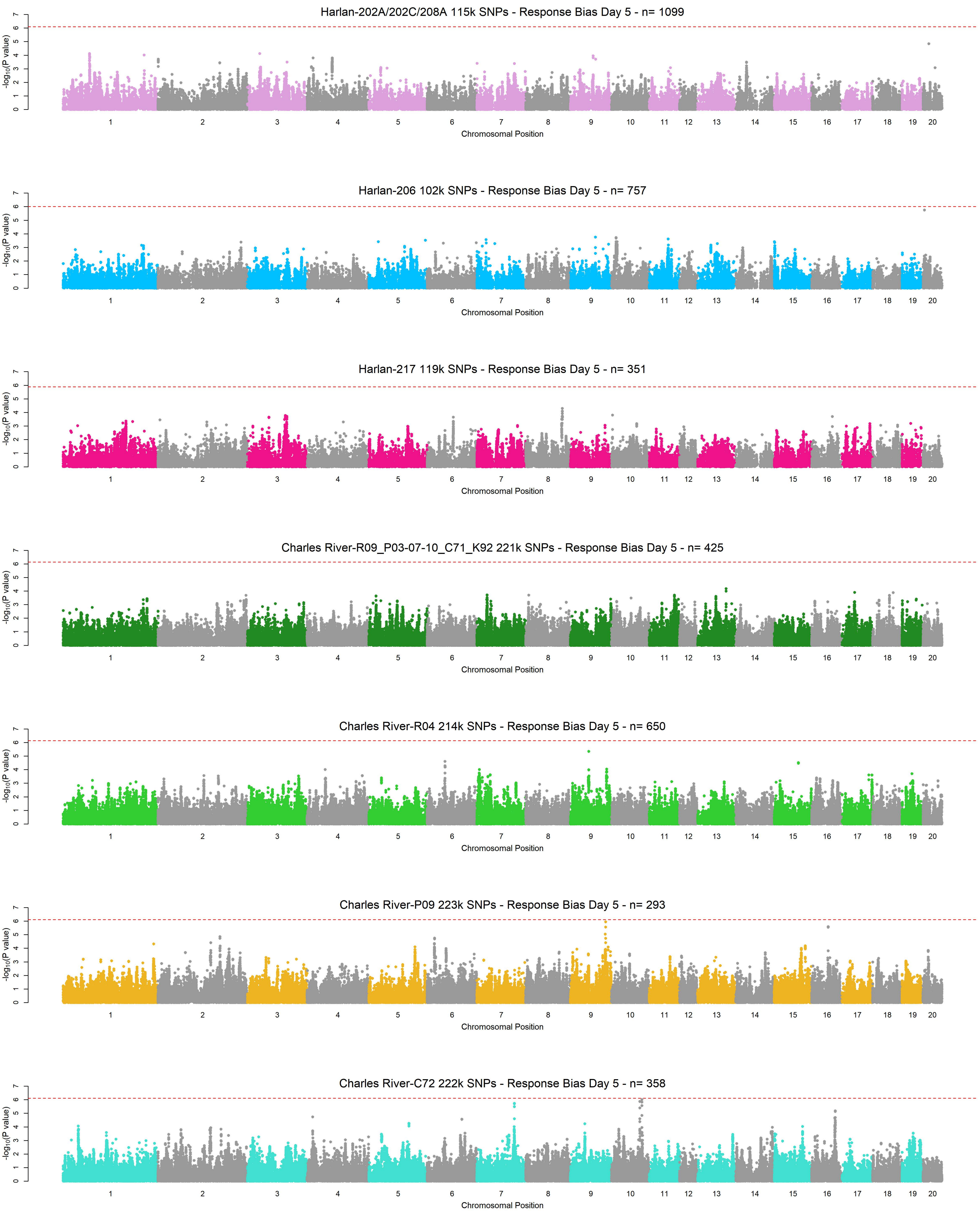
Harlan-202A/202C/208A 115k SNPs - Response Bias Day 5 - n= 1099
Harlan-206 102k SNPs - Response Bias Day 5 - n= 757
Harlan-217 119k SNPs - Response Bias Day 5 - n= 351
Charles River-R09_P03-07-10_C71_K92 221k SNPs - Response Bias Day 5 - n= 425
Charles River-R04 214k SNPs - Response Bias Day 5 - n= 650
Charles River-P09 223k SNPs - Response Bias Day 5 - n= 293
Charles River-C72 222k SNPs - Response Bias Day 5 - n= 358
$-\log_{10}$(P value)
0 1 2 3 4 5 6 7
1 2 3 4 5 6 7 8 9 10 11 12 13 14 15 16 17 18 19 20
Chromosomal Position

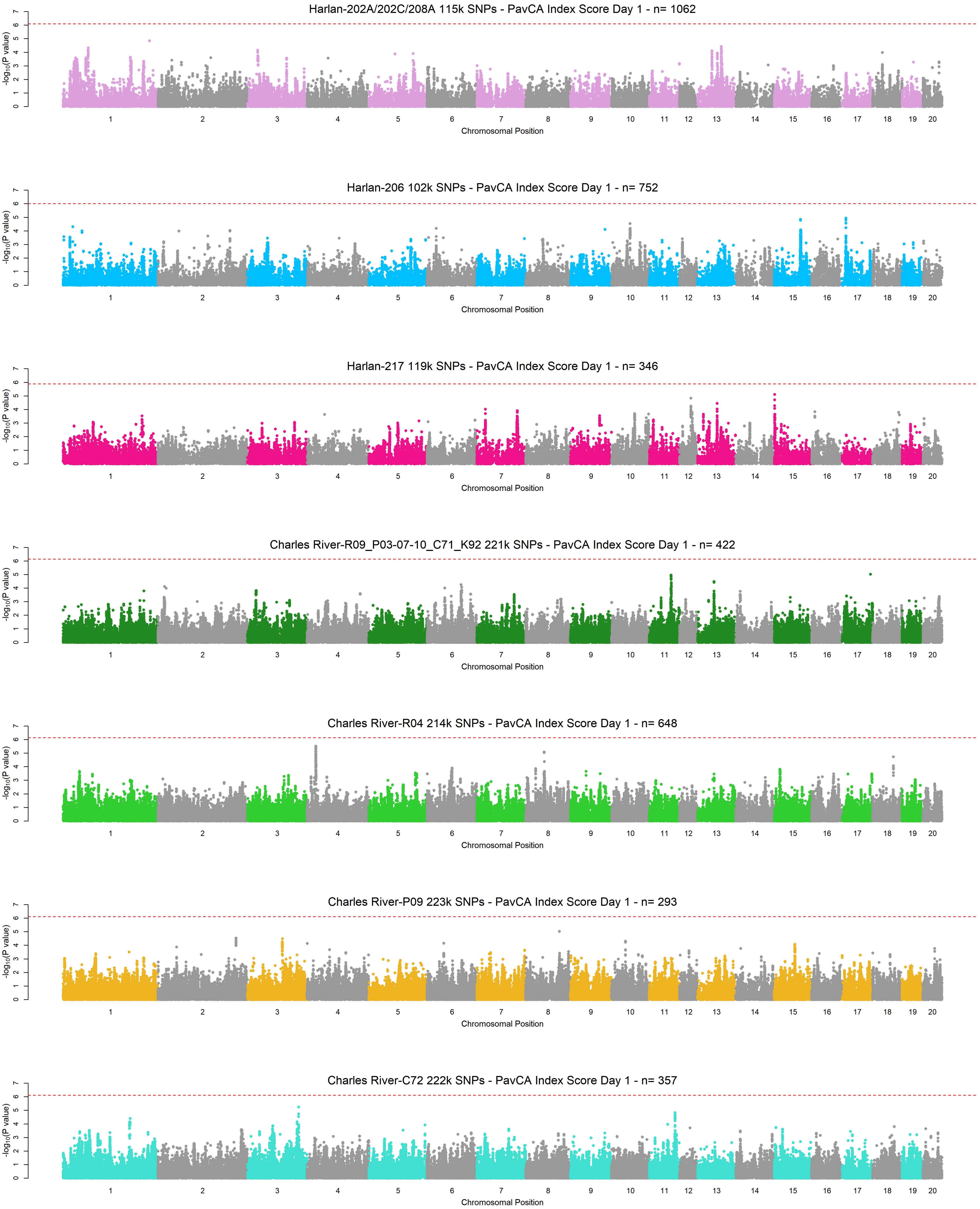
Harlan-202A/202C/208A 115k SNPs - PavCA Index Score Day 1 - n= 1062
Harlan-206 102k SNPs - PavCA Index Score Day 1 - n= 752
Harlan-217 119k SNPs - PavCA Index Score Day 1 - n= 346
Charles River-R09_P03-07-10_C71_K92 221k SNPs - PavCA Index Score Day 1 - n= 422
Charles River-R04 214k SNPs - PavCA Index Score Day 1 - n= 648
Charles River-P09 223k SNPs - PavCA Index Score Day 1 - n= 293
Charles River-C72 222k SNPs - PavCA Index Score Day 1 - n= 357
-log10(P value)
Chromosomal Position
1 2 3 4 5 6 7 8 9 10 11 12 13 14 15 16 17 18 19 20

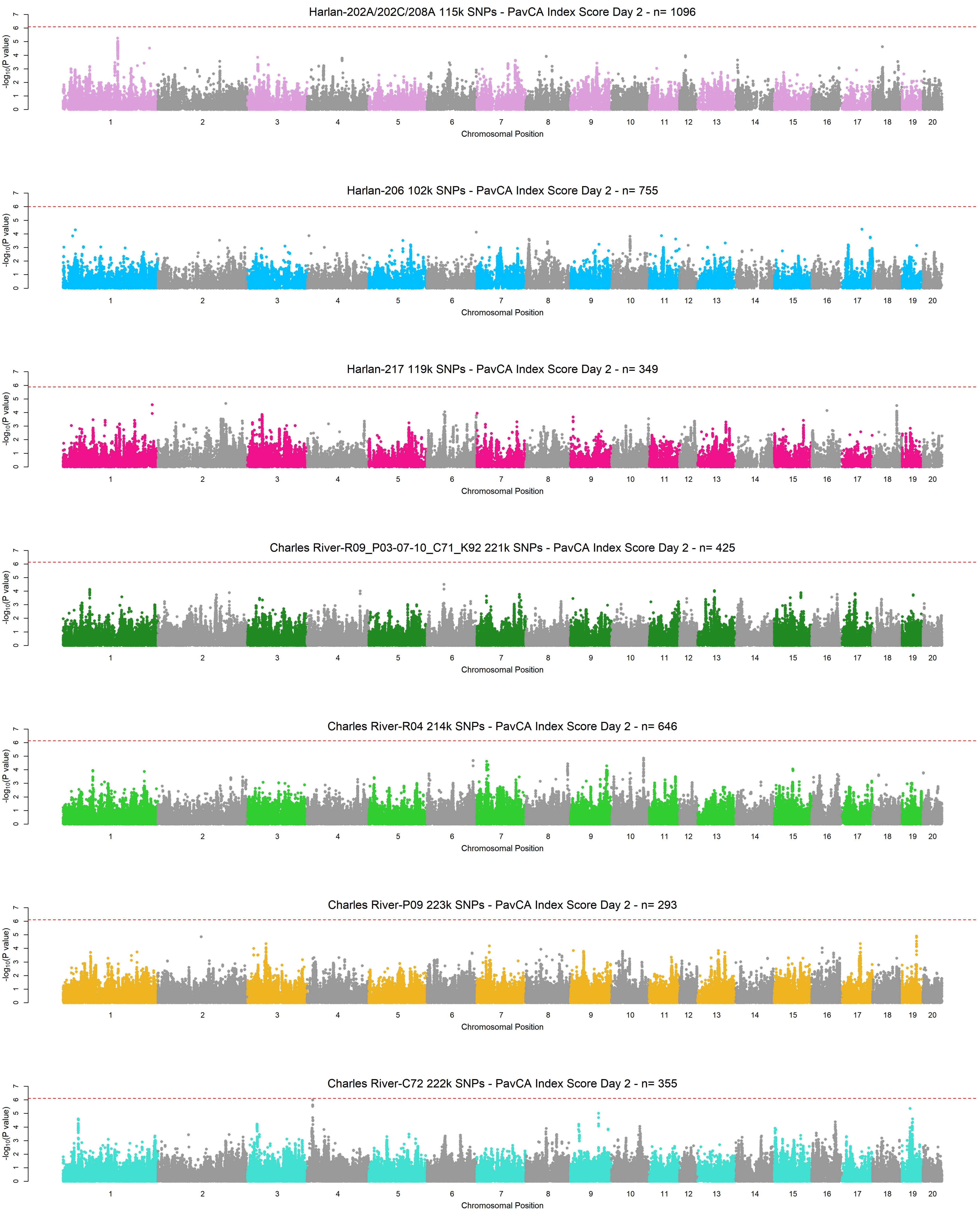

Harlan-202A/202C/208A 115k SNPs - PavCA Index Score Day 2 - n= 1096
Harlan-206 102k SNPs - PavCA Index Score Day 2 - n= 755
Harlan-217 119k SNPs - PavCA Index Score Day 2 - n= 349
Charles River-R09_P03-07-10_C71_K92 221k SNPs - PavCA Index Score Day 2 - n= 425
Charles River-R04 214k SNPs - PavCA Index Score Day 2 - n= 646
Charles River-P09 223k SNPs - PavCA Index Score Day 2 - n= 293
Charles River-C72 222k SNPs - PavCA Index Score Day 2 - n= 355
-log10(P value)
Chromosomal Position
1 2 3 4 5 6 7 8 9 10 11 12 13 14 15 16 17 18 19 20

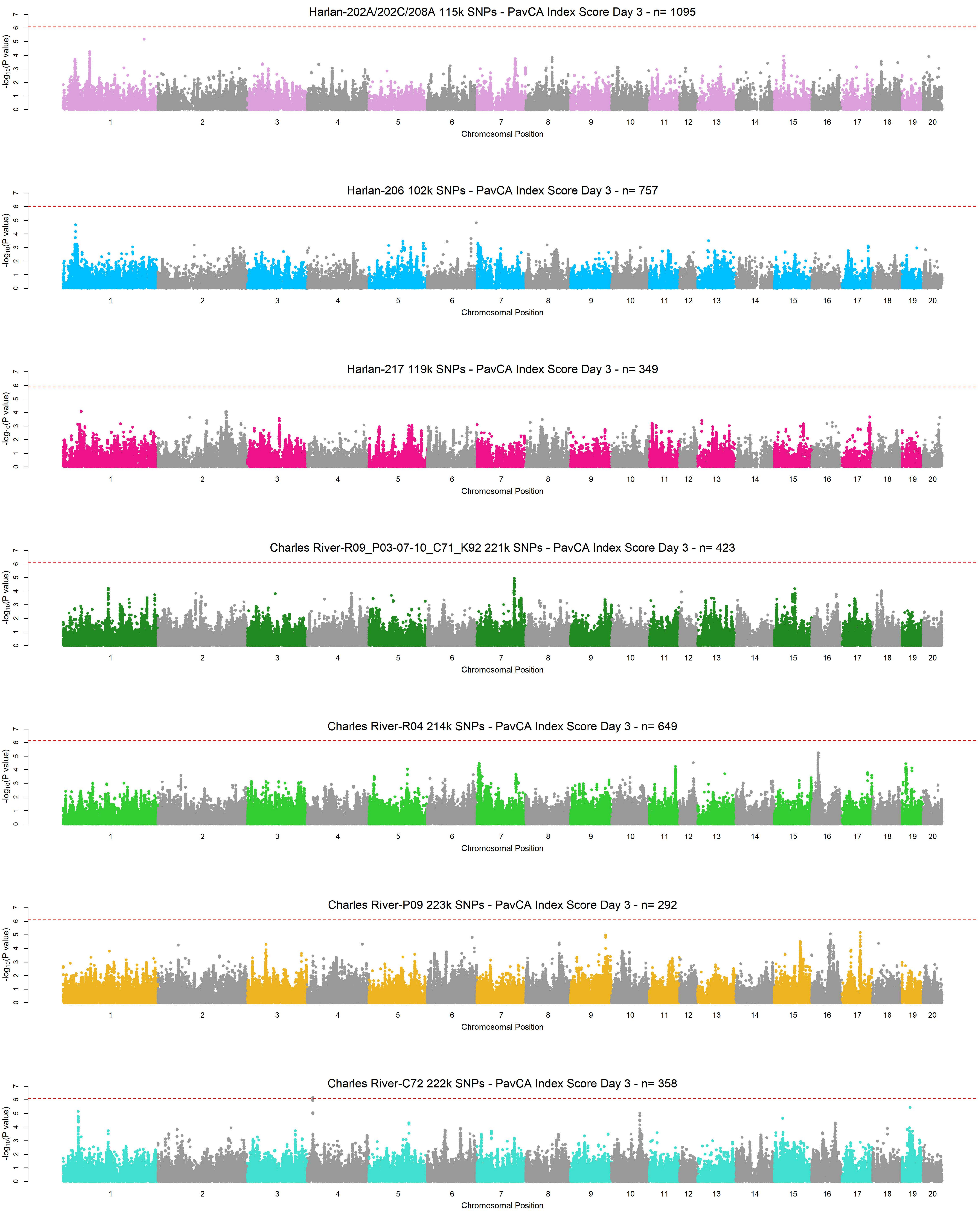
Harlan-202A/202C/208A 115k SNPs - PavCA Index Score Day 3 - n= 1095
Harlan-206 102k SNPs - PavCA Index Score Day 3 - n= 757
Harlan-217 119k SNPs - PavCA Index Score Day 3 - n= 349
Charles River-R09_P03-07-10_C71_K92 221k SNPs - PavCA Index Score Day 3 - n= 423
Charles River-R04 214k SNPs - PavCA Index Score Day 3 - n= 649
Charles River-P09 223k SNPs - PavCA Index Score Day 3 - n= 292
Charles River-C72 222k SNPs - PavCA Index Score Day 3 - n= 358
-log10(P value)
Chromosomal Position
1 2 3 4 5 6 7 8 9 10 11 12 13 14 15 16 17 18 19 20

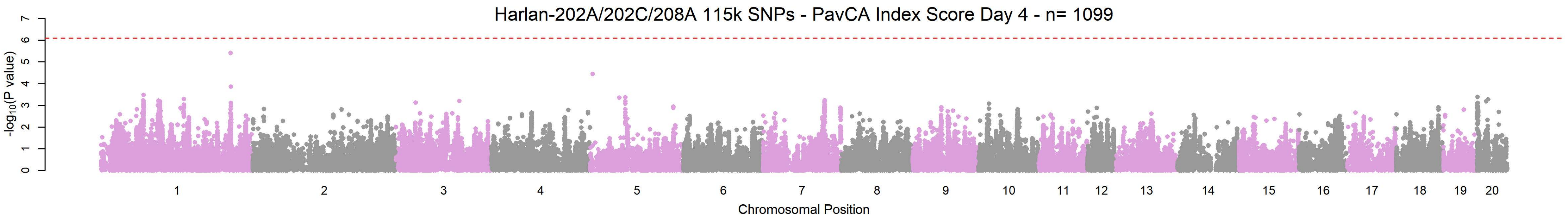
Harlan-202A/202C/208A 115k SNPs - PavCA Index Score Day 4 - n= 1099
-log10(P value)
Chromosomal Position

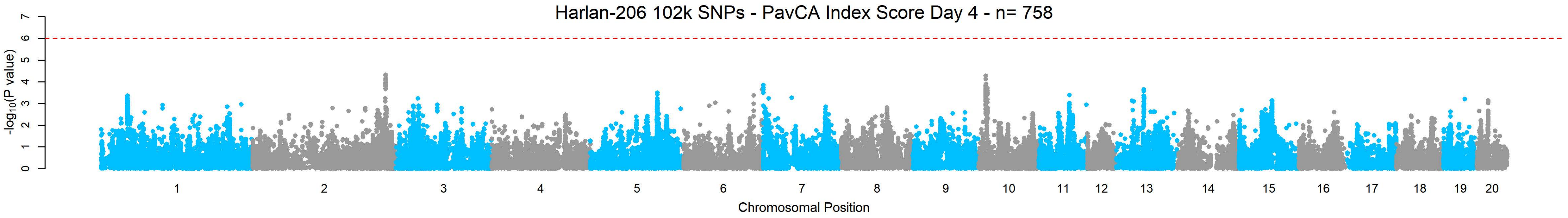
Harlan-206 102k SNPs - PavCA Index Score Day 4 - n= 758
-log10(P value)
Chromosomal Position

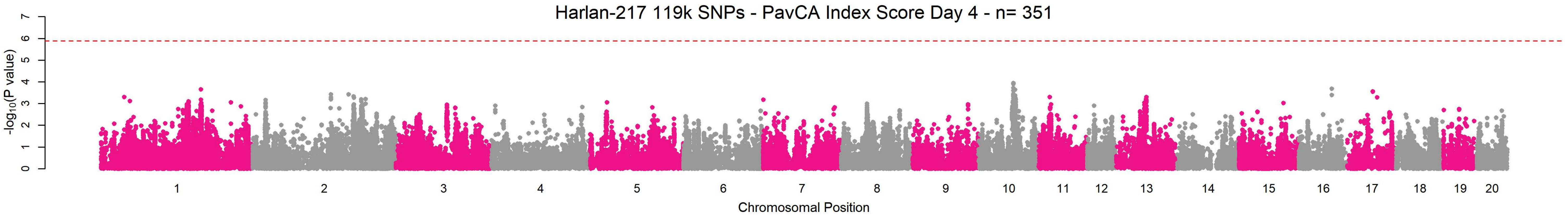
Harlan-217 119k SNPs - PavCA Index Score Day 4 - n= 351
-log10(P value)
Chromosomal Position

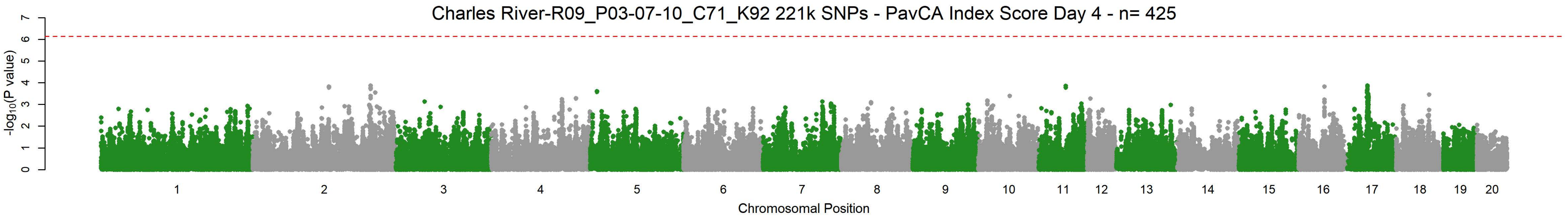
Charles River-R09_P03-07-10_C71_K92 221k SNPs - PavCA Index Score Day 4 - n= 425
-log10(P value)
Chromosomal Position

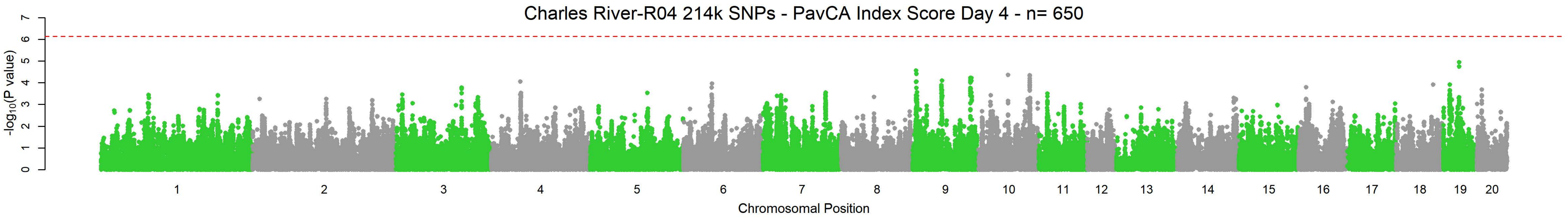
Charles River-R04 214k SNPs - PavCA Index Score Day 4 - n= 650
-log10(P value)
Chromosomal Position

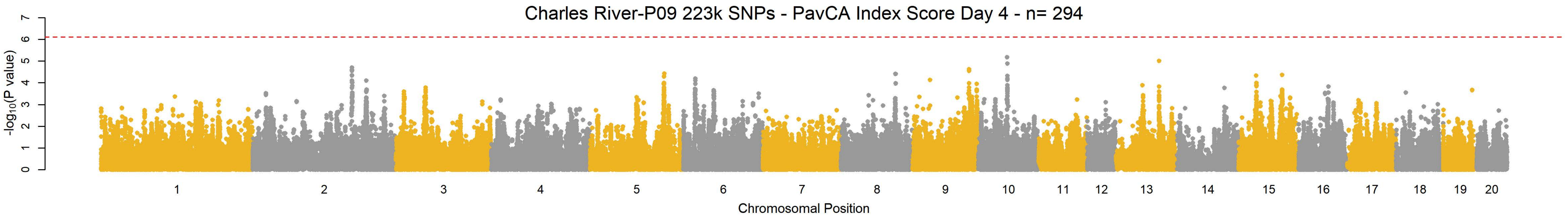
Charles River-P09 223k SNPs - PavCA Index Score Day 4 - n= 294
-log10(P value)
Chromosomal Position

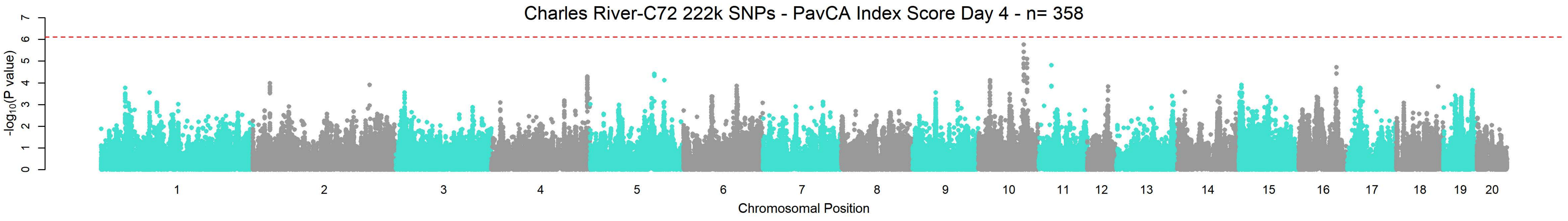
Charles River-C72 222k SNPs - PavCA Index Score Day 4 - n= 358
-log10(P value)
Chromosomal Position

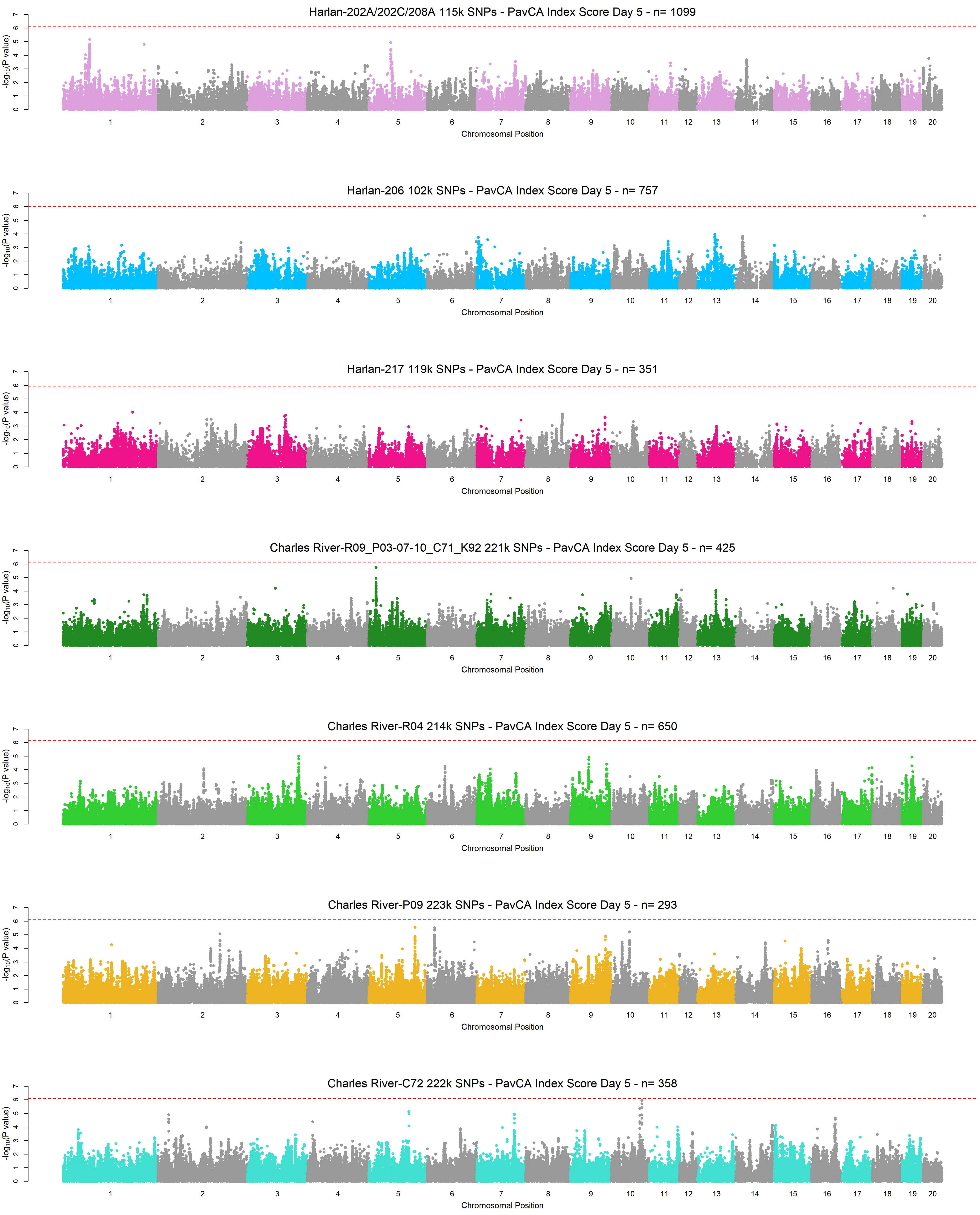
Harlan-202A/202C/208A 115k SNPs - PavCA Index Score Day 5 - n= 1099
Harlan-206 102k SNPs - PavCA Index Score Day 5 - n= 757
Harlan-217 119k SNPs - PavCA Index Score Day 5 - n= 351
Charles River-R09_P03-07-10_C71_K92 221k SNPs - PavCA Index Score Day 5 - n= 425
Charles River-R04 214k SNPs - PavCA Index Score Day 5 - n= 650
Charles River-P09 223k SNPs - PavCA Index Score Day 5 - n= 293
Charles River-C72 222k SNPs - PavCA Index Score Day 5 - n= 358
-log10(P value)
Chromosomal Position
1 2 3 4 5 6 7 8 9 10 11 12 13 14 15 16 17 18 19 20
