## Supplementary material for "Genetic characterization of outbred Sprague Dawley rats and utility for genome-wide association studies": S2_Text - ddGBS Primer and Adapter Sequences

**ddGBS Adapter/Primer Sequence List**

**ddGBS PCR Primer 1**

**Seq:** 5'-AAT GAT ACG GCG ACC ACC GAG ATC TAC ACT CTT TCC CTA CAC GAC GCT CTT CCG ATC T-3'
Amount: 100nmoles
Modifications: HPLC Purification

**ddGBS PCR Primer 2**

**Seq:** 5'-CAA GCA GAA GAC GGC ATA CGA GAT CGG TCT CGG CAT TCC TGC TGA A-3'
Amount: 100nmoles
Modifications: HPLC Purification

**NlaIII Y-Adapter**

**Top:
Seq:** 5’-/5Phos/A*GA TCG GAA GAG CGG GGA CTT TAA GC-3'
Amount: 100nmoles
Modifications: HPLC Purification, 5’ Phosphorylation, Phosphorothioate Bond

**Bottom:
Seq:** 5'-GAT CGG TCT CGG CAT TCC TGC TGA ACC GCT CTT CCG ATC TCA T*G-3'
Amount: 100nmoles
Modifications: HPLC Purification, Phosphorothioate Bond

**Pst1 Barcode Adapters**

| **Well** | **Barcode** | **Strand** | **Sequence** |
| --- | --- | --- | --- |
| A01 | CTCG | F | ACA CTC TTT CCC TAC ACG ACG CTC TTC CGA TCT CTC GTG CA |
|  |  | R | CGA GAG ATC GGA AGA GCG TCG TGT AGG GAA AGA GTG T |
| A02 | TGCA | F | ACA CTC TTT CCC TAC ACG ACG CTC TTC CGA TCT TGC ATG CA |
|  |  | R | TGC AAG ATC GGA AGA GCG TCG TGT AGG GAA AGA GTG T |
| A03 | ACTA | F | ACA CTC TTT CCC TAC ACG ACG CTC TTC CGA TCT ACT ATG CA |
|  |  | R | TAG TAG ATC GGA AGA GCG TCG TGT AGG GAA AGA GTG T |
| A04 | CAGA | F | ACA CTC TTT CCC TAC ACG ACG CTC TTC CGA TCT CAG ATG CA |
|  |  | R | TCT GAG ATC GGA AGA GCG TCG TGT AGG GAA AGA GTG T |
| A05 | AACT | F | ACA CTC TTT CCC TAC ACG ACG CTC TTC CGA TCT AAC TTG CA |
|  |  | R | AGT TAG ATC GGA AGA GCG TCG TGT AGG GAA AGA GTG T |
| A06 | GCGT | F | ACA CTC TTT CCC TAC ACG ACG CTC TTC CGA TCT GCG TTG CA |
|  |  | R | ACG CAG ATC GGA AGA GCG TCG TGT AGG GAA AGA GTG T |
| A07 | GTAA | F | ACA CTC TTT CCC TAC ACG ACG CTC TTC CGA TCT GTA ATG CA |
|  |  | R | TTA CAG ATC GGA AGA GCG TCG TGT AGG GAA AGA GTG T |
| A08 | CGAT | F | ACA CTC TTT CCC TAC ACG ACG CTC TTC CGA TCT CGA TTG CA |
|  |  | R | ATC GAG ATC GGA AGA GCG TCG TGT AGG GAA AGA GTG T |
| A09 | ACCGT | F | ACA CTC TTT CCC TAC ACG ACG CTC TTC CGA TCT ACC GTT GCA |
|  |  | R | ACG GTA GAT CGG AAG AGC GTC GTG TAG GGA AAG AGT GT |
| A10 | TCACG | F | ACA CTC TTT CCC TAC ACG ACG CTC TTC CGA TCT TCA CGT GCA |
|  |  | R | CGT GAA GAT CGG AAG AGC GTC GTG TAG GGA AAG AGT GT |
| A11 | CTAGG | F | ACA CTC TTT CCC TAC ACG ACG CTC TTC CGA TCT CTA GGT GCA |
|  |  | R | CCT AGA GAT CGG AAG AGC GTC GTG TAG GGA AAG AGT GT |
| A12 | ACAAA | F | ACA CTC TTT CCC TAC ACG ACG CTC TTC CGA TCT ACA AAT GCA |
|  |  | R | TTT GTA GAT CGG AAG AGC GTC GTG TAG GGA AAG AGT GT |
| B01 | TTCTG | F | ACA CTC TTT CCC TAC ACG ACG CTC TTC CGA TCT TTC TGT GCA |
|  |  | R | CAG AAA GAT CGG AAG AGC GTC GTG TAG GGA AAG AGT GT |
| B02 | AGCCG | F | ACA CTC TTT CCC TAC ACG ACG CTC TTC CGA TCT AGC CGT GCA |
|  |  | R | CGG CTA GAT CGG AAG AGC GTC GTG TAG GGA AAG AGT GT |
| B03 | GTATT | F | ACA CTC TTT CCC TAC ACG ACG CTC TTC CGA TCT GTA TTT GCA |
|  |  | R | AAT ACA GAT CGG AAG AGC GTC GTG TAG GGA AAG AGT GT |
| B04 | CTGTA | F | ACA CTC TTT CCC TAC ACG ACG CTC TTC CGA TCT CTG TAT GCA |
|  |  | R | TAC AGA GAT CGG AAG AGC GTC GTG TAG GGA AAG AGT GT |
| B05 | CGCTT | F | ACA CTC TTT CCC TAC ACG ACG CTC TTC CGA TCT CGC TTT GCA |
|  |  | R | AAG CGA GAT CGG AAG AGC GTC GTG TAG GGA AAG AGT GT |
| B06 | GCTTA | F | ACA CTC TTT CCC TAC ACG ACG CTC TTC CGA TCT GCT TAT GCA |
|  |  | R | TAA GCA GAT CGG AAG AGC GTC GTG TAG GGA AAG AGT GT |
| B07 | GGTGT | F | ACA CTC TTT CCC TAC ACG ACG CTC TTC CGA TCT GGT GTT GCA |
|  |  | R | ACA CCA GAT CGG AAG AGC GTC GTG TAG GGA AAG AGT GT |
| B08 | CCAGCT | F | ACA CTC TTT CCC TAC ACG ACG CTC TTC CGA TCT CCA GCT TGC A |
|  |  | R | AGC TGG AGA TCG GAA GAG CGT CGT GTA GGG AAA GAG TGT |
| B09 | TTCAGA | F | ACA CTC TTT CCC TAC ACG ACG CTC TTC CGA TCT TTC AGA TGC A |
|  |  | R | TCT GAA AGA TCG GAA GAG CGT CGT GTA GGG AAA GAG TGT |
| B10 | TAGGAA | F | ACA CTC TTT CCC TAC ACG ACG CTC TTC CGA TCT TAG GAA TGC A |
|  |  | R | TTC CTA AGA TCG GAA GAG CGT CGT GTA GGG AAA GAG TGT |
| B11 | GCTCTA | F | ACA CTC TTT CCC TAC ACG ACG CTC TTC CGA TCT GCT CTA TGC A |
|  |  | R | TAG AGC AGA TCG GAA GAG CGT CGT GTA GGG AAA GAG TGT |
| B12 | CCACAA | F | ACA CTC TTT CCC TAC ACG ACG CTC TTC CGA TCT CCA CAA TGC A |
|  |  | R | TTG TGG AGA TCG GAA GAG CGT CGT GTA GGG AAA GAG TGT |
| C01 | TGCGA | F | ACA CTC TTT CCC TAC ACG ACG CTC TTC CGA TCT TGC GAT GCA |
|  |  | R | TCG CAA GAT CGG AAG AGC GTC GTG TAG GGA AAG AGT GT |
| C02 | CTTCCA | F | ACA CTC TTT CCC TAC ACG ACG CTC TTC CGA TCT CTT CCA TGC A |
|  |  | R | TGG AAG AGA TCG GAA GAG CGT CGT GTA GGG AAA GAG TGT |
| C03 | GAGATA | F | ACA CTC TTT CCC TAC ACG ACG CTC TTC CGA TCT GAG ATA TGC A |
|  |  | R | TAT CTC AGA TCG GAA GAG CGT CGT GTA GGG AAA GAG TGT |
| C04 | ATGCCT | F | ACA CTC TTT CCC TAC ACG ACG CTC TTC CGA TCT ATG CCT TGC A |
|  |  | R | AGG CAT AGA TCG GAA GAG CGT CGT GTA GGG AAA GAG TGT |
| C05 | TATTTTT | F | ACA CTC TTT CCC TAC ACG ACG CTC TTC CGA TCT TAT TTT TTG CA |
|  |  | R | AAA AAT AAG ATC GGA AGA GCG TCG TGT AGG GAA AGA GTG T |
| C06 | CTTGCTT | F | ACA CTC TTT CCC TAC ACG ACG CTC TTC CGA TCT CTT GCT TTG CA |
|  |  | R | AAG CAA GAG ATC GGA AGA GCG TCG TGT AGG GAA AGA GTG T |
| C07 | ATGAAAG | F | ACA CTC TTT CCC TAC ACG ACG CTC TTC CGA TCT ATG AAA GTG CA |
|  |  | R | CTT TCA TAG ATC GGA AGA GCG TCG TGT AGG GAA AGA GTG T |
| C08 | AAAAGTT | F | ACA CTC TTT CCC TAC ACG ACG CTC TTC CGA TCT AAA AGT TTG CA |
|  |  | R | AAC TTT TAG ATC GGA AGA GCG TCG TGT AGG GAA AGA GTG T |
| C09 | GAATTCA | F | ACA CTC TTT CCC TAC ACG ACG CTC TTC CGA TCT GAA TTC ATG CA |
|  |  | R | TGA ATT CAG ATC GGA AGA GCG TCG TGT AGG GAA AGA GTG T |
| C10 | GAACTTG | F | ACA CTC TTT CCC TAC ACG ACG CTC TTC CGA TCT GAA CTT GTG CA |
|  |  | R | CAA GTT CAG ATC GGA AGA GCG TCG TGT AGG GAA AGA GTG T |
| C11 | GGACCTA | F | ACA CTC TTT CCC TAC ACG ACG CTC TTC CGA TCT GGA CCT ATG CA |
|  |  | R | TAG GTC CAG ATC GGA AGA GCG TCG TGT AGG GAA AGA GTG T |
| C12 | GTCGATT | F | ACA CTC TTT CCC TAC ACG ACG CTC TTC CGA TCT GTC GAT TTG CA |
|  |  | R | AAT CGA CAG ATC GGA AGA GCG TCG TGT AGG GAA AGA GTG T |
| D01 | AACGCCT | F | ACA CTC TTT CCC TAC ACG ACG CTC TTC CGA TCT AAC GCC TTG CA |
|  |  | R | AGG CGT TAG ATC GGA AGA GCG TCG TGT AGG GAA AGA GTG T |
| D02 | AATATGG | F | ACA CTC TTT CCC TAC ACG ACG CTC TTC CGA TCT AAT ATG GTG CA |
|  |  | R | CCA TAT TAG ATC GGA AGA GCG TCG TGT AGG GAA AGA GTG T |
| D03 | ACGACTAG | F | ACA CTC TTT CCC TAC ACG ACG CTC TTC CGA TCT ACG ACT AGT GCA |
|  |  | R | CTA GTC GTA GAT CGG AAG AGC GTC GTG TAG GGA AAG AGT GT |
| D04 | AGTGGA | F | ACA CTC TTT CCC TAC ACG ACG CTC TTC CGA TCT AGT GGA TGC A |
|  |  | R | TCC ACT AGA TCG GAA GAG CGT CGT GTA GGG AAA GAG TGT |
| D05 | TAGCATGG | F | ACA CTC TTT CCC TAC ACG ACG CTC TTC CGA TCT TAG CAT GGT GCA |
|  |  | R | CCA TGC TAA GAT CGG AAG AGC GTC GTG TAG GGA AAG AGT GT |
| D06 | GGTTGT | F | ACA CTC TTT CCC TAC ACG ACG CTC TTC CGA TCT GGT TGT TGC A |
|  |  | R | ACA ACC AGA TCG GAA GAG CGT CGT GTA GGG AAA GAG TGT |
| D07 | TAGGCCAT | F | ACA CTC TTT CCC TAC ACG ACG CTC TTC CGA TCT TAG GCC ATT GCA |
|  |  | R | ATG GCC TAA GAT CGG AAG AGC GTC GTG TAG GGA AAG AGT GT |
| D08 | TTCCTGGA | F | ACA CTC TTT CCC TAC ACG ACG CTC TTC CGA TCT TTC CTG GAT GCA |
|  |  | R | TCC AGG AAA GAT CGG AAG AGC GTC GTG TAG GGA AAG AGT GT |
| D09 | TGGTACGT | F | ACA CTC TTT CCC TAC ACG ACG CTC TTC CGA TCT TGG TAC GTT GCA |
|  |  | R | ACG TAC CAA GAT CGG AAG AGC GTC GTG TAG GGA AAG AGT GT |
| D10 | TCTCAGTG | F | ACA CTC TTT CCC TAC ACG ACG CTC TTC CGA TCT TCT CAG TGT GCA |
|  |  | R | CAC TGA GAA GAT CGG AAG AGC GTC GTG TAG GGA AAG AGT GT |
| D11 | CCGGATAT | F | ACA CTC TTT CCC TAC ACG ACG CTC TTC CGA TCT CCG GAT ATT GCA |
|  |  | R | ATA TCC GGA GAT CGG AAG AGC GTC GTG TAG GGA AAG AGT GT |
| D12 | CGCCTTAT | F | ACA CTC TTT CCC TAC ACG ACG CTC TTC CGA TCT CGC CTT ATT GCA |
|  |  | R | ATA AGG CGA GAT CGG AAG AGC GTC GTG TAG GGA AAG AGT GT |
| F01 | Common | F | GAT CGG AAG AGC GGT TCA GCA GGA ATG CCG AG |
|  |  | R | CTC GGC ATT CCT GCT GAA CCG CTC TTC CGA TCT GCA |
| F02 | Common | F | GAT CGG AAG AGC GGT TCA GCA GGA ATG CCG AG |
|  |  | R | CTC GGC ATT CCT GCT GAA CCG CTC TTC CGA TCT GCA |
| F03 | Common | F | GAT CGG AAG AGC GGT TCA GCA GGA ATG CCG AG |
|  |  | R | CTC GGC ATT CCT GCT GAA CCG CTC TTC CGA TCT GCA |
