## Supplementary material for "Genetic characterization of outbred Sprague Dawley rats and utility for genome-wide association studies": S3_File - Subpop QQ-Plots

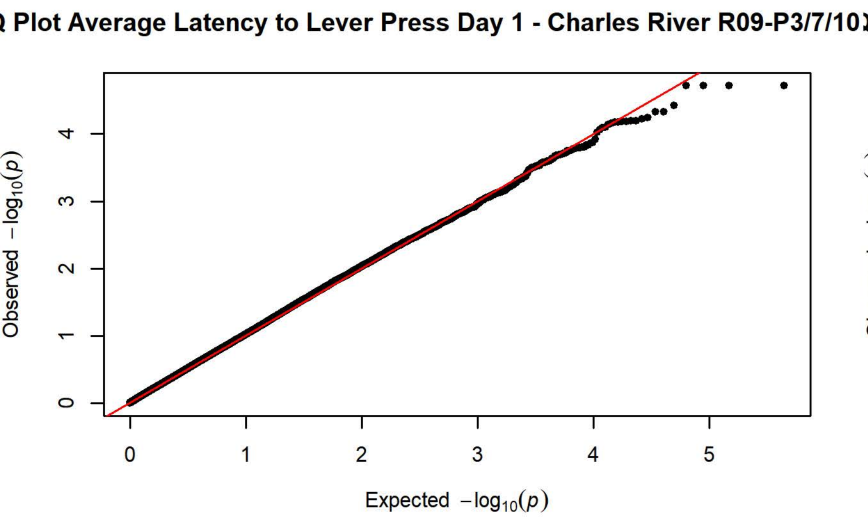
Q Plot Average Latency to Lever Press Day 1 - Charles River R09-P3/7/10
Observed −log10(p)
Expected −log10(p)

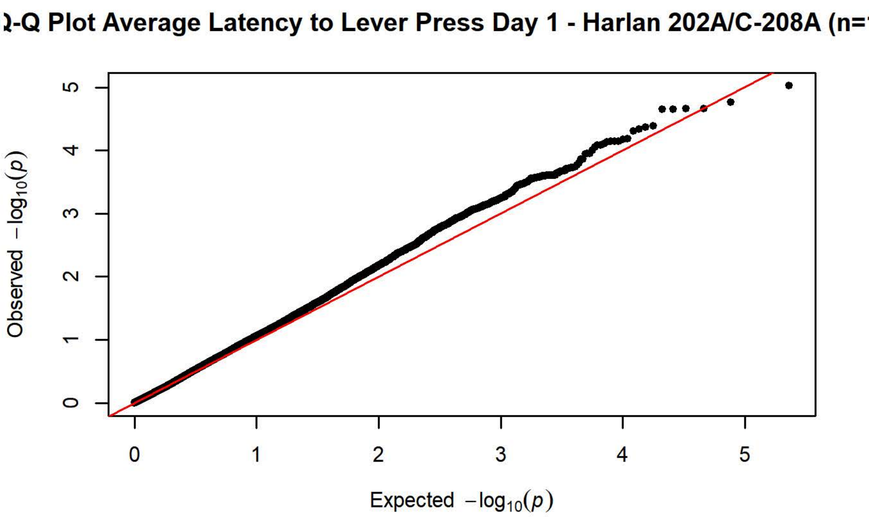
Q-Q Plot Average Latency to Lever Press Day 1 - Harlan 202A/C-208A (n=
Observed −log10(p)
Expected −log10(p)

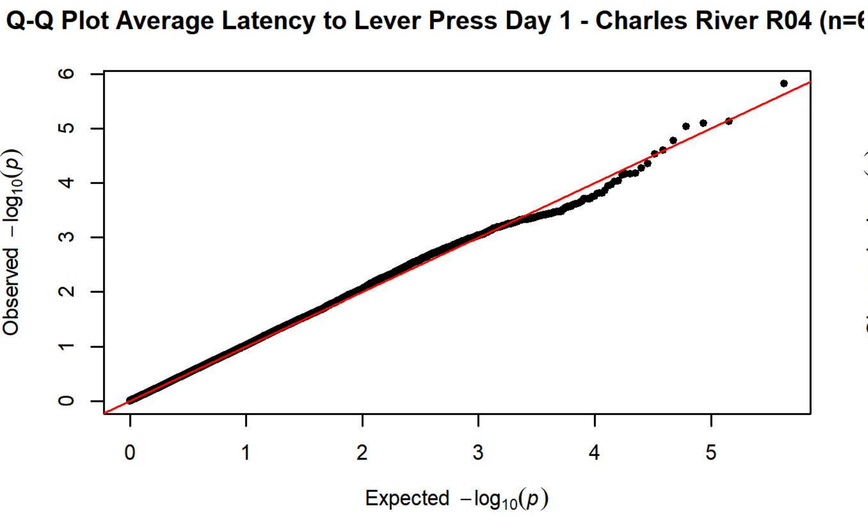
Q-Q Plot Average Latency to Lever Press Day 1 - Charles River R04 (n=
Observed −log10(p)
Expected −log10(p)

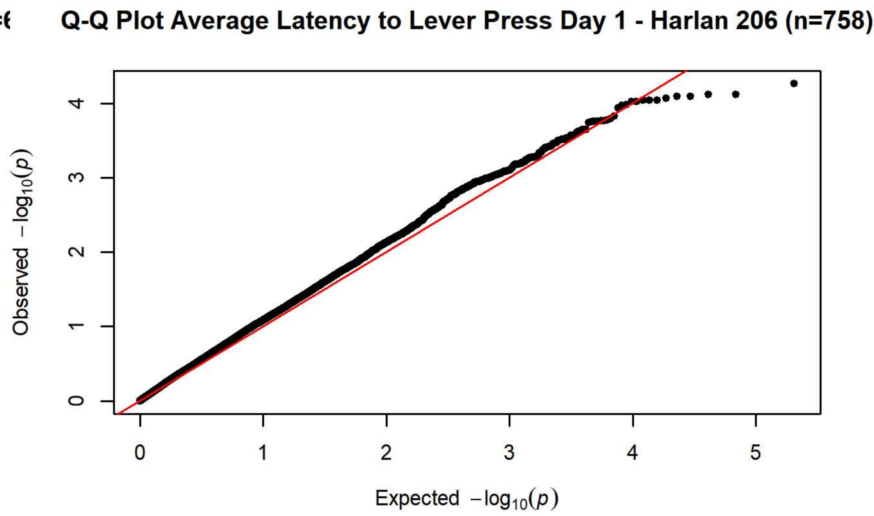
Q-Q Plot Average Latency to Lever Press Day 1 - Harlan 206 (n=758)
Observed −log10(p)
Expected −log10(p)

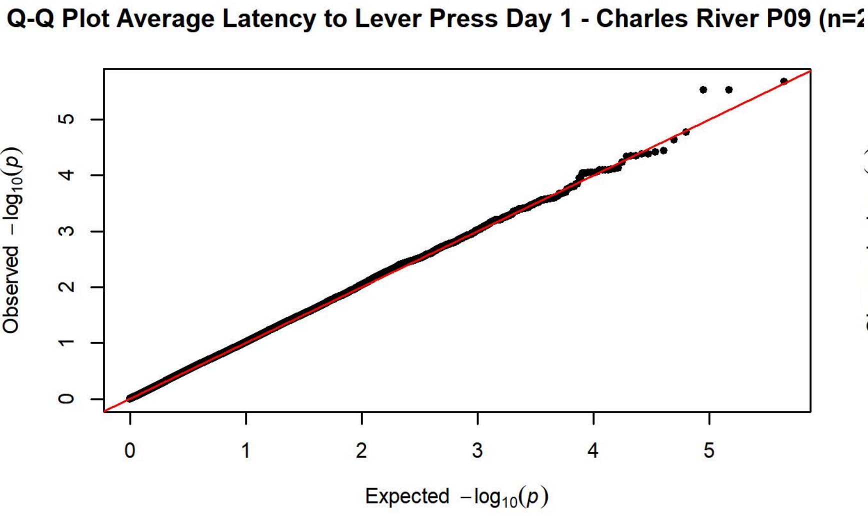
Q-Q Plot Average Latency to Lever Press Day 1 - Charles River P09 (n=
Observed −log10(p)
Expected −log10(p)

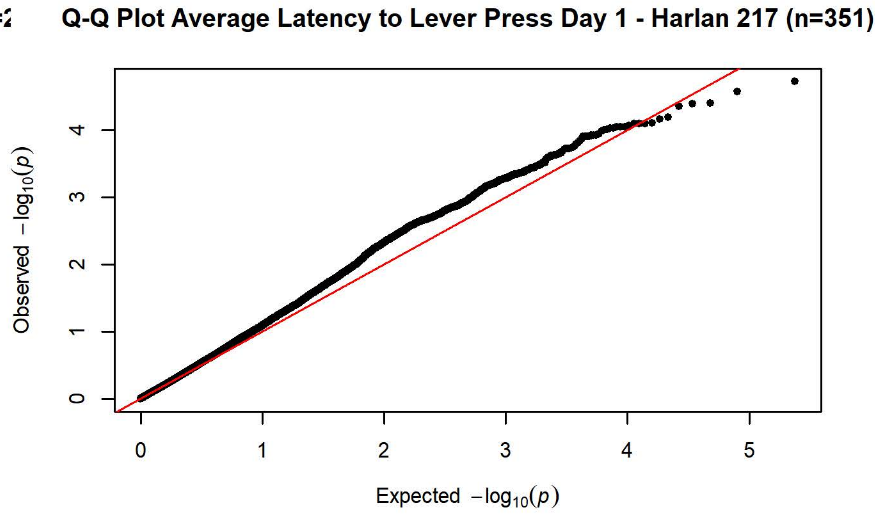
Q-Q Plot Average Latency to Lever Press Day 1 - Harlan 217 (n=351)
Observed −log10(p)
Expected −log10(p)

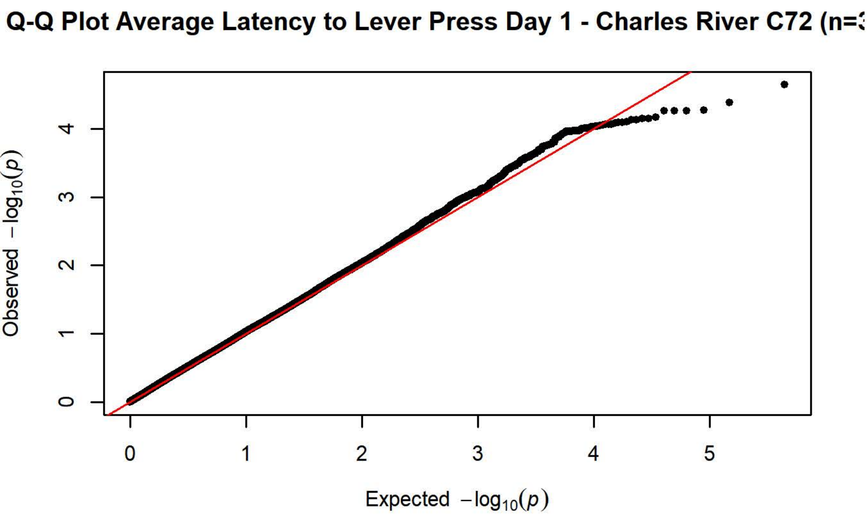
Q-Q Plot Average Latency to Lever Press Day 1 - Charles River C72 (n=
Observed −log10(p)
Expected −log10(p)

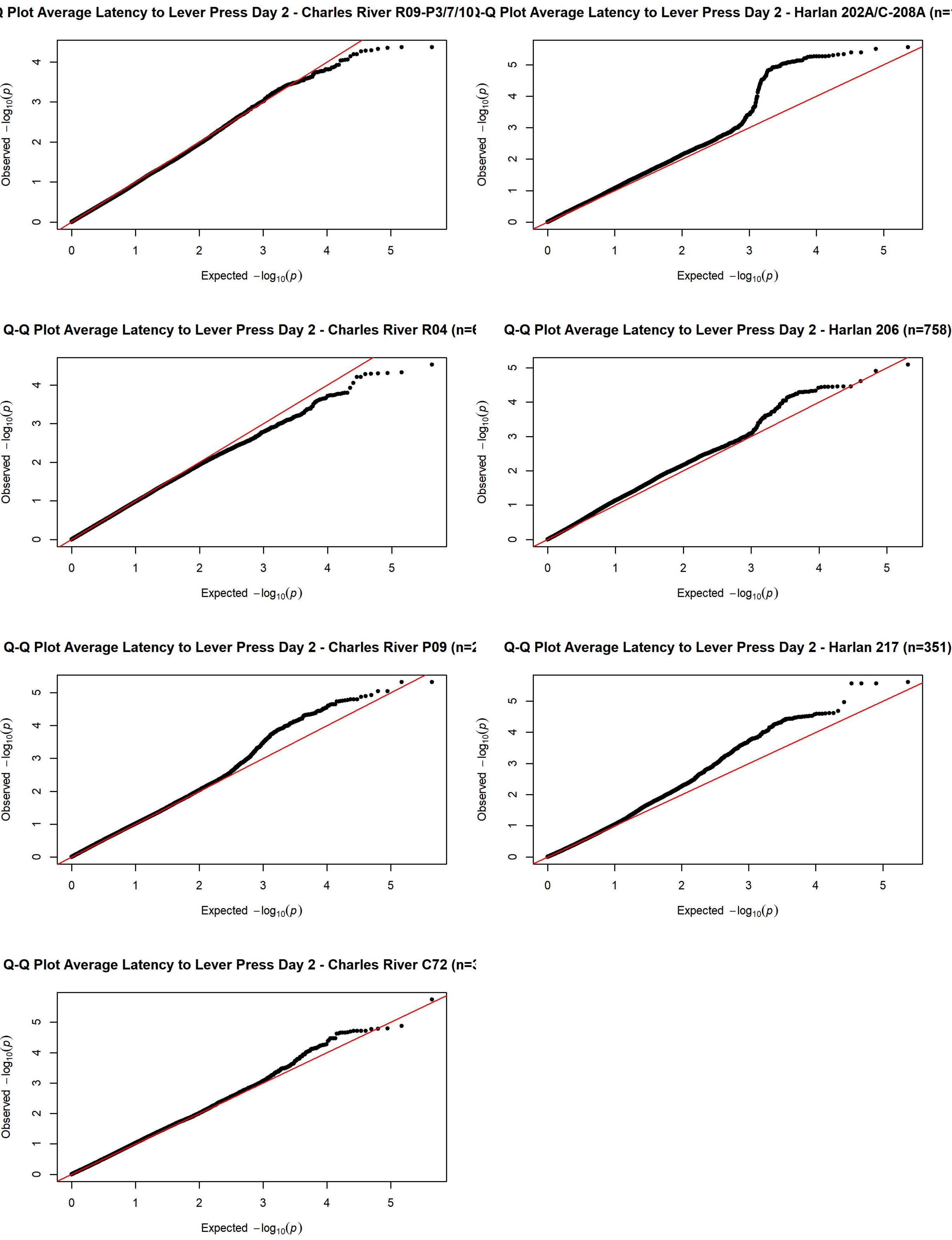

Q Plot Average Latency to Lever Press Day 2 - Charles River R09-P3/7/10
Q-Q Plot Average Latency to Lever Press Day 2 - Harlan 202A/C-208A (n=
Q-Q Plot Average Latency to Lever Press Day 2 - Charles River R04 (n=6
Q-Q Plot Average Latency to Lever Press Day 2 - Harlan 206 (n=758)
Q-Q Plot Average Latency to Lever Press Day 2 - Charles River P09 (n=2
Q-Q Plot Average Latency to Lever Press Day 2 - Harlan 217 (n=351)
Q-Q Plot Average Latency to Lever Press Day 2 - Charles River C72 (n=3
Observed $-\log_{10}(p)$
Expected $-\log_{10}(p)$

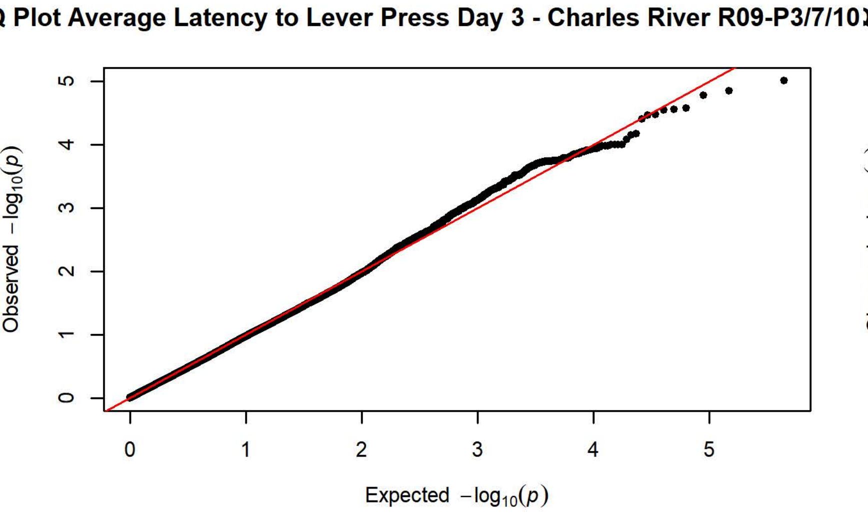
Q Plot Average Latency to Lever Press Day 3 - Charles River R09-P3/7/10
Observed $-\log_{10}(p)$
Expected $-\log_{10}(p)$

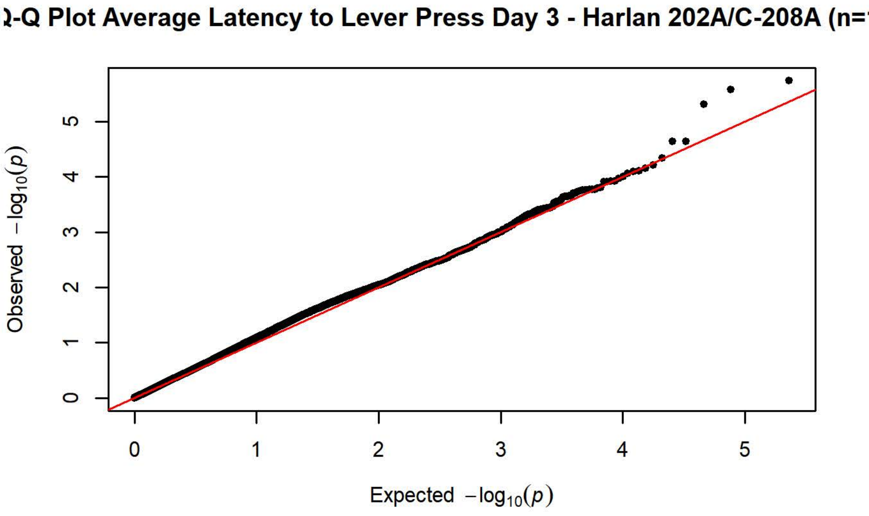
-Q Plot Average Latency to Lever Press Day 3 - Harlan 202A/C-208A (n=
Observed $-\log_{10}(p)$
Expected $-\log_{10}(p)$

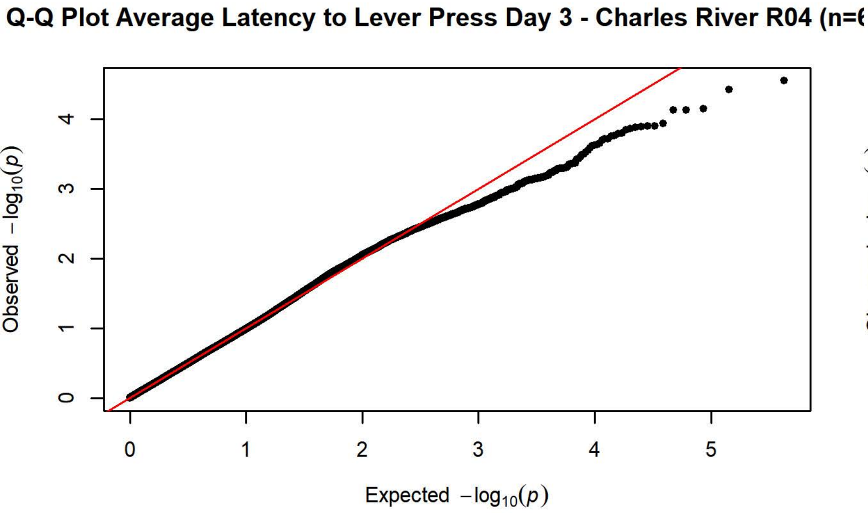
Q-Q Plot Average Latency to Lever Press Day 3 - Charles River R04 (n=
Observed $-\log_{10}(p)$
Expected $-\log_{10}(p)$

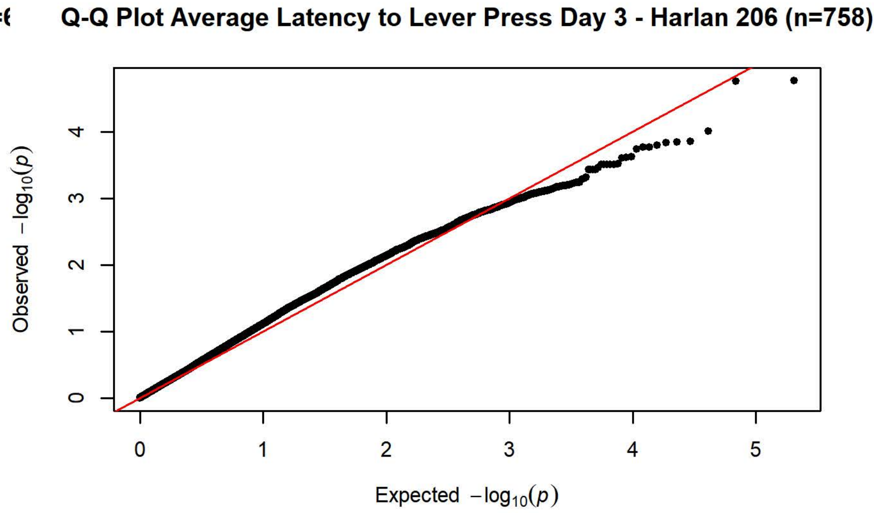
Q-Q Plot Average Latency to Lever Press Day 3 - Harlan 206 (n=758)
Observed $-\log_{10}(p)$
Expected $-\log_{10}(p)$

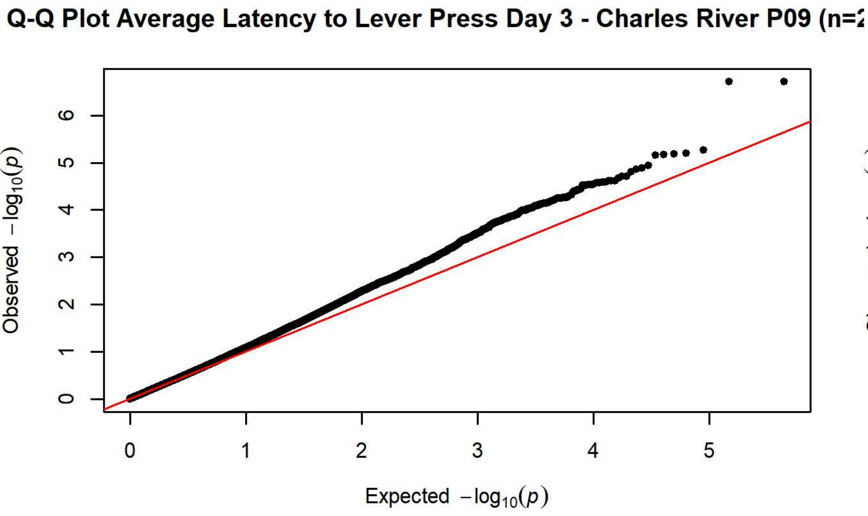
Q-Q Plot Average Latency to Lever Press Day 3 - Charles River P09 (n=
Observed $-\log_{10}(p)$
Expected $-\log_{10}(p)$

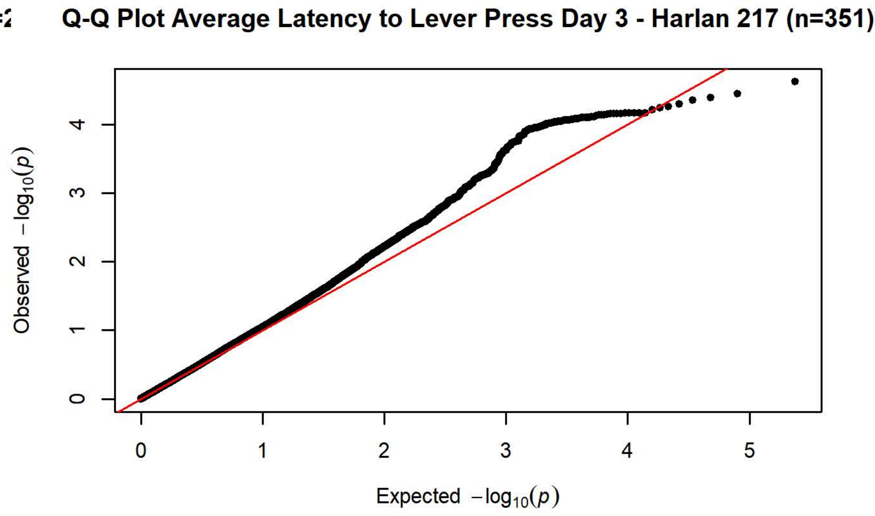
Q-Q Plot Average Latency to Lever Press Day 3 - Harlan 217 (n=351)
Observed $-\log_{10}(p)$
Expected $-\log_{10}(p)$

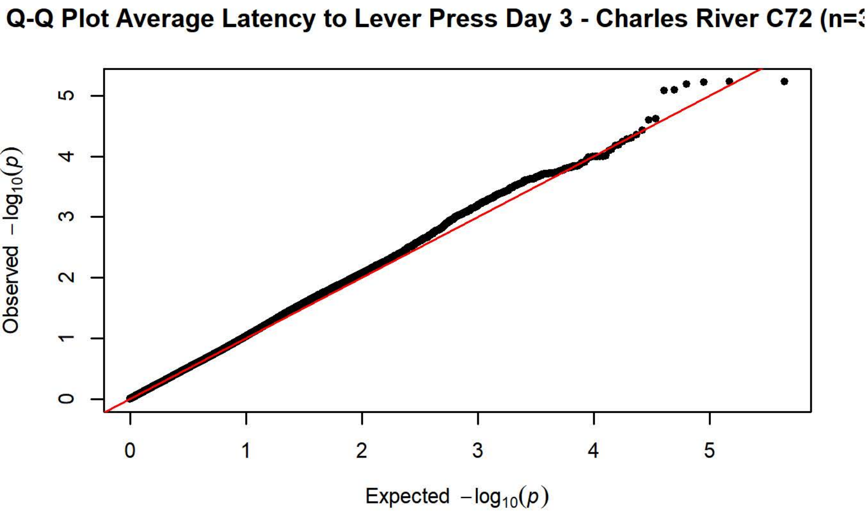
Q-Q Plot Average Latency to Lever Press Day 3 - Charles River C72 (n=
Observed $-\log_{10}(p)$
Expected $-\log_{10}(p)$

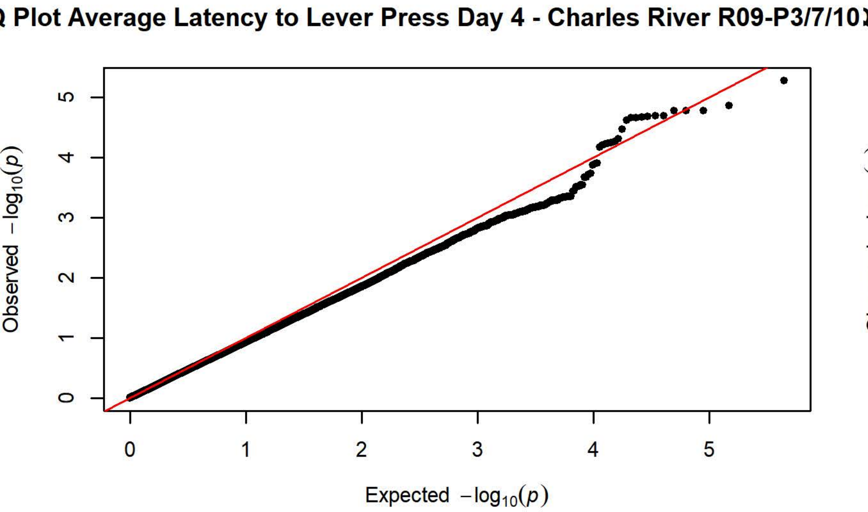
Q Plot Average Latency to Lever Press Day 4 - Charles River R09-P3/7/10
Observed −log10(p)
Expected −log10(p)

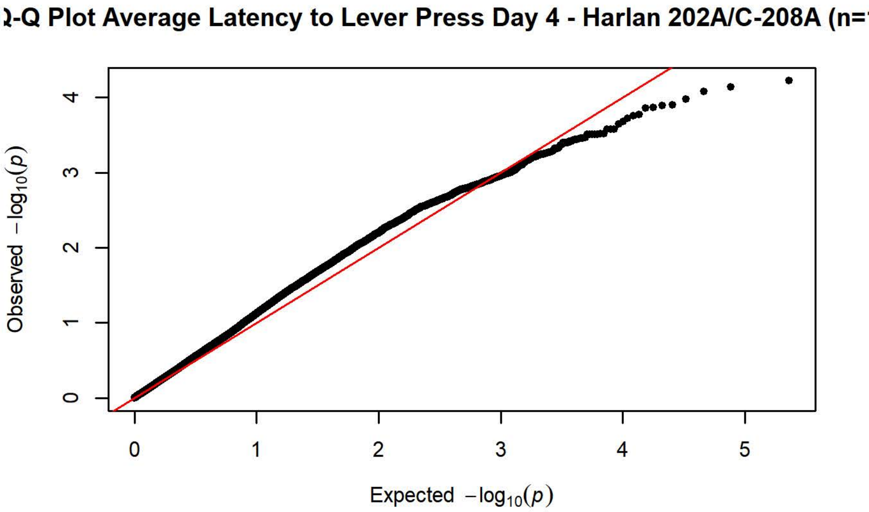
-Q Plot Average Latency to Lever Press Day 4 - Harlan 202A/C-208A (n=
Observed −log10(p)
Expected −log10(p)

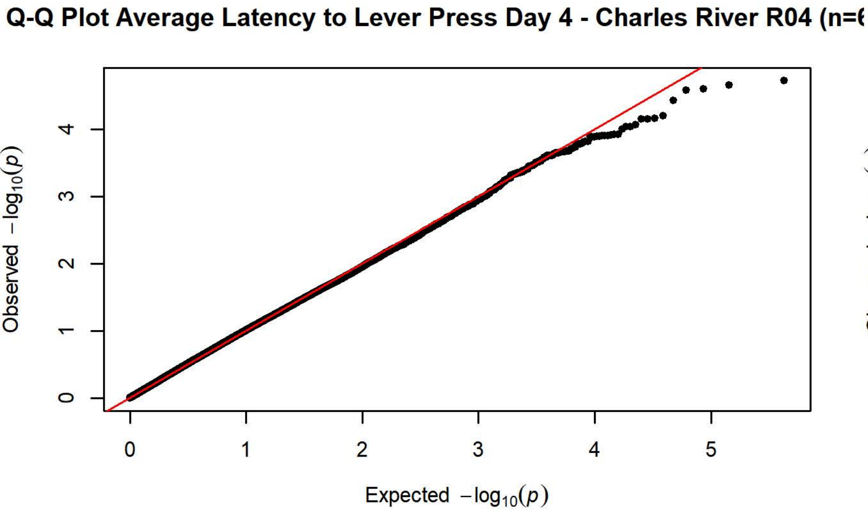
Q-Q Plot Average Latency to Lever Press Day 4 - Charles River R04 (n=
Observed −log10(p)
Expected −log10(p)

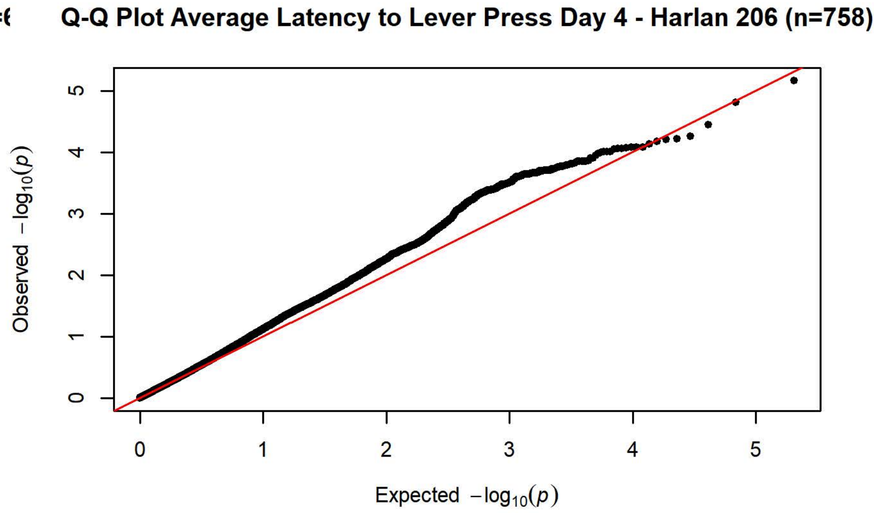
Q-Q Plot Average Latency to Lever Press Day 4 - Harlan 206 (n=758)
Observed −log10(p)
Expected −log10(p)

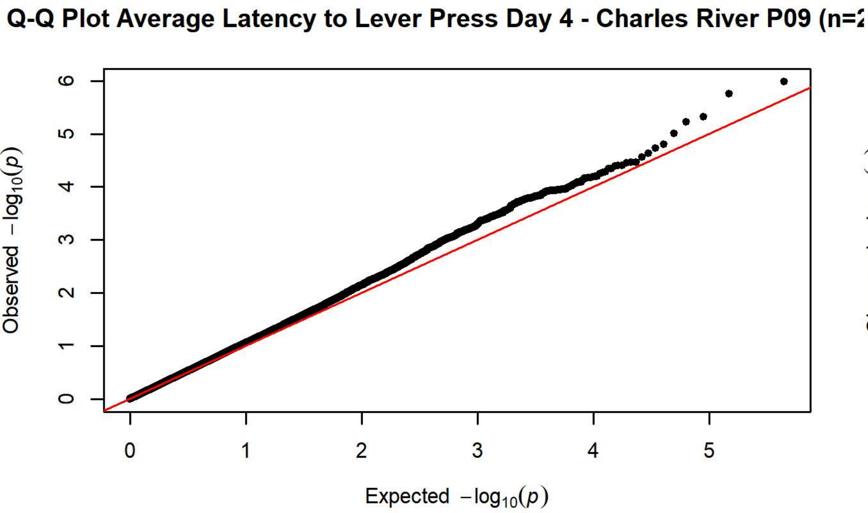
Q-Q Plot Average Latency to Lever Press Day 4 - Charles River P09 (n=
Observed −log10(p)
Expected −log10(p)

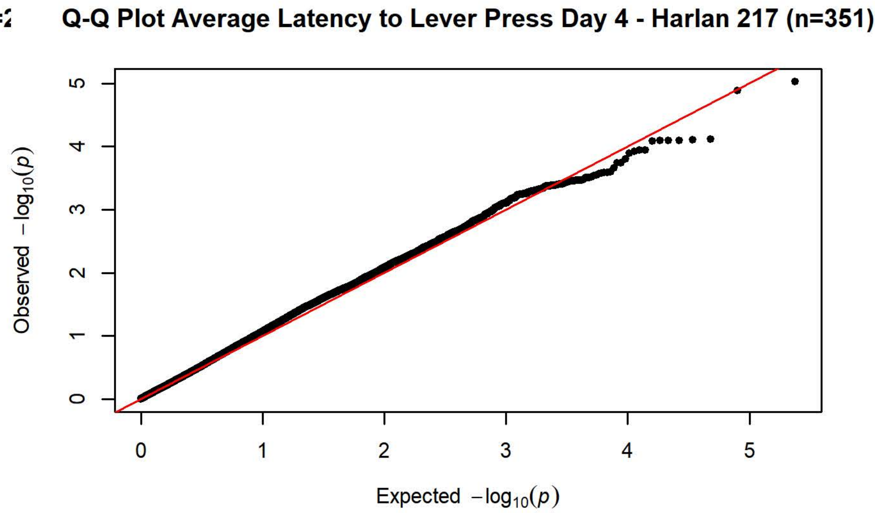
Q-Q Plot Average Latency to Lever Press Day 4 - Harlan 217 (n=351)
Observed −log10(p)
Expected −log10(p)

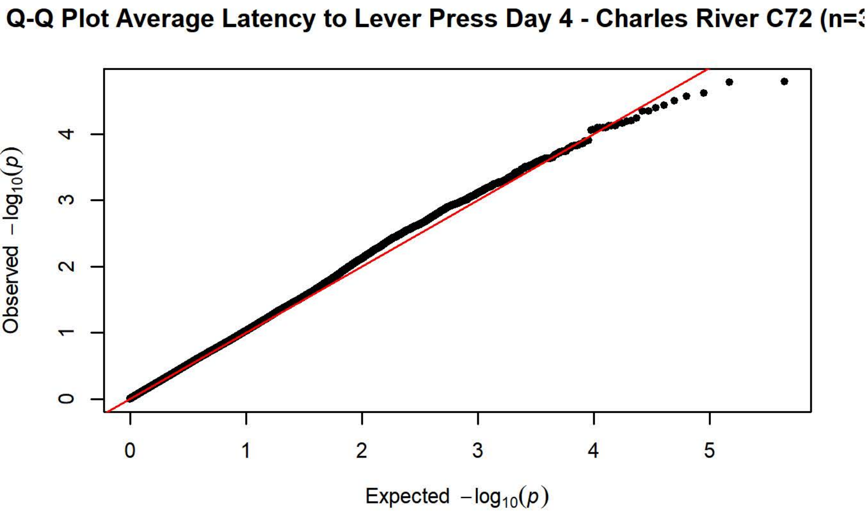
Q-Q Plot Average Latency to Lever Press Day 4 - Charles River C72 (n=
Observed −log10(p)
Expected −log10(p)

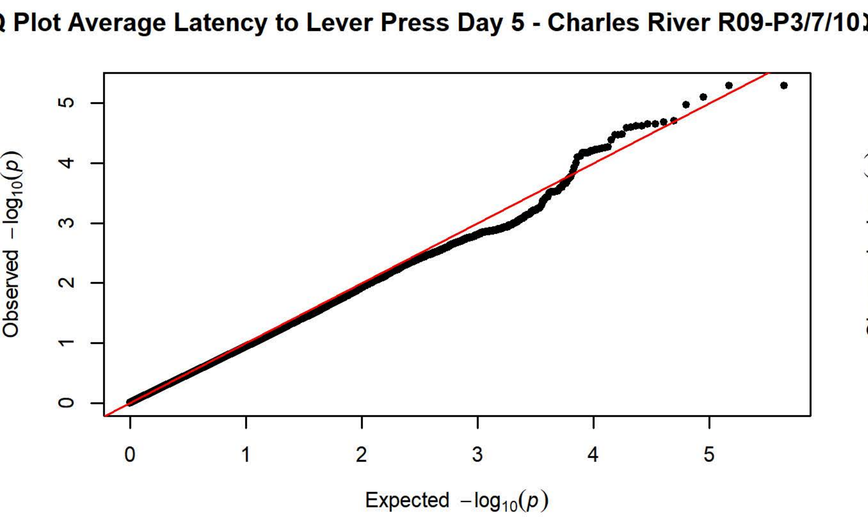
Q-Q Plot Average Latency to Lever Press Day 5 - Charles River R09-P3/7/10
Observed −log10(p)
Expected −log10(p)

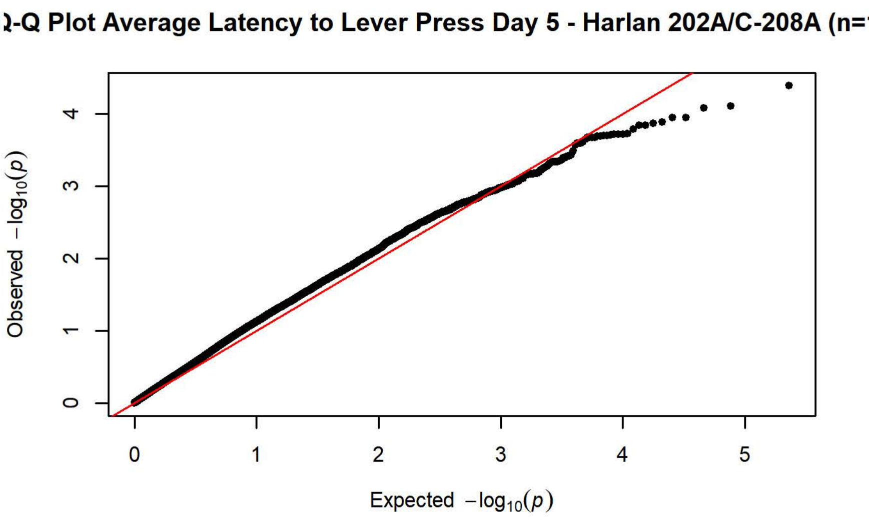
Q-Q Plot Average Latency to Lever Press Day 5 - Harlan 202A/C-208A (n=
Observed −log10(p)
Expected −log10(p)

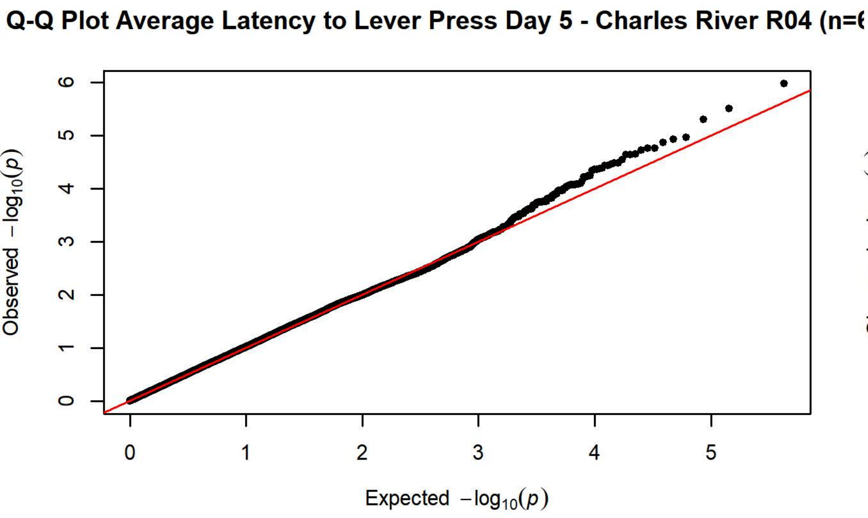
Q-Q Plot Average Latency to Lever Press Day 5 - Charles River R04 (n=
Observed −log10(p)
Expected −log10(p)

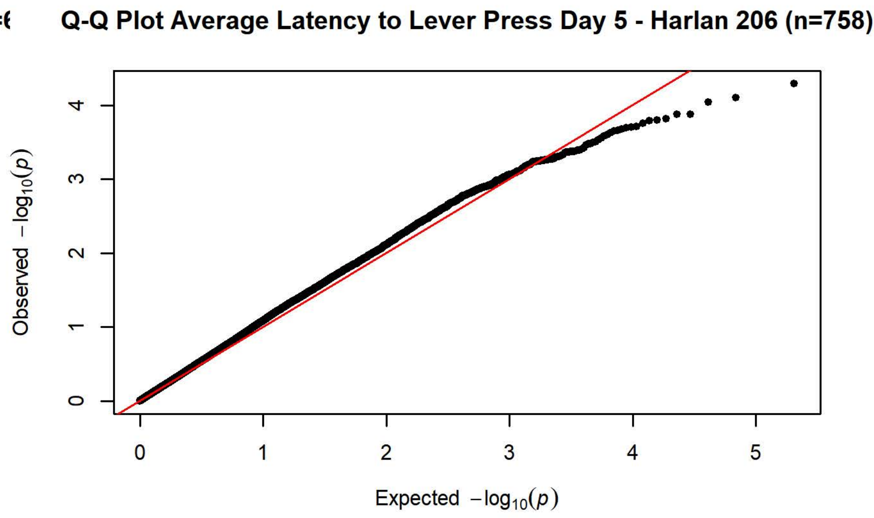
Q-Q Plot Average Latency to Lever Press Day 5 - Harlan 206 (n=758)
Observed −log10(p)
Expected −log10(p)

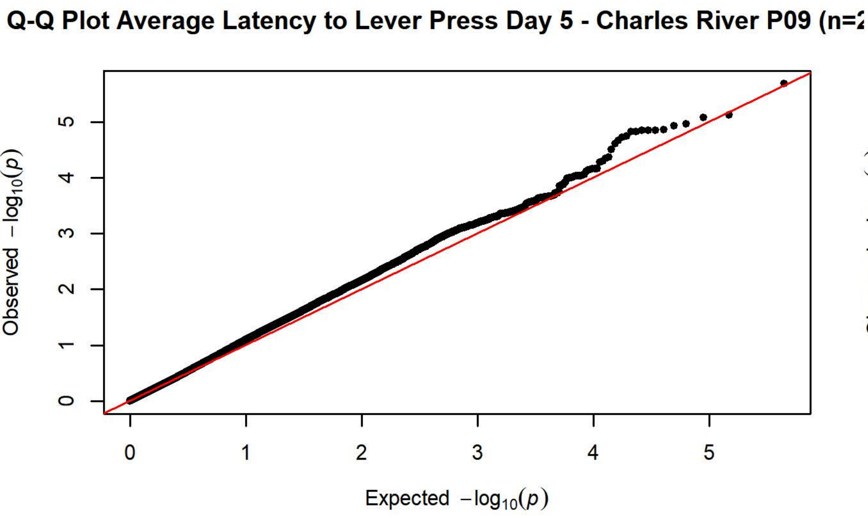
Q-Q Plot Average Latency to Lever Press Day 5 - Charles River P09 (n=
Observed −log10(p)
Expected −log10(p)

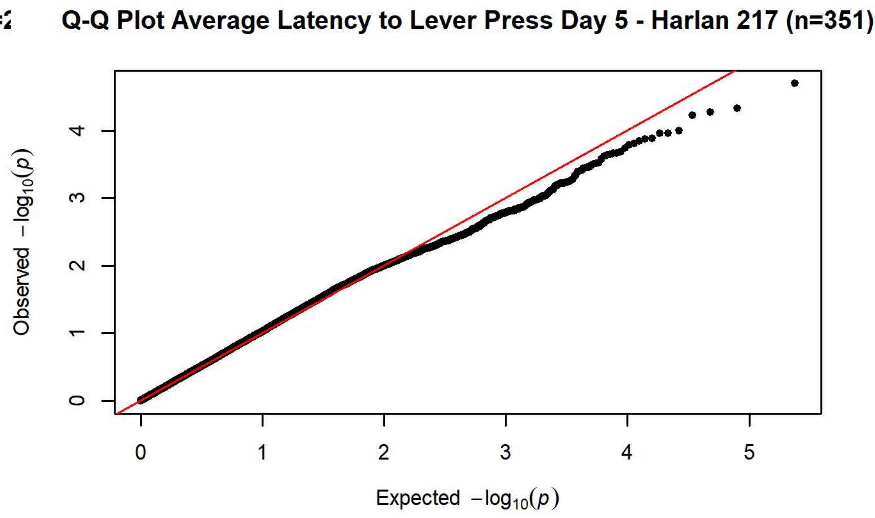
Q-Q Plot Average Latency to Lever Press Day 5 - Harlan 217 (n=351)
Observed −log10(p)
Expected −log10(p)

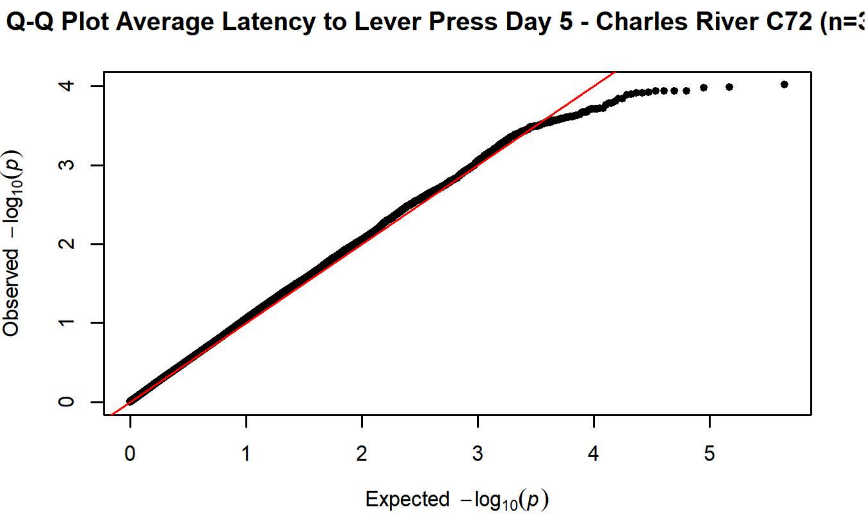
Q-Q Plot Average Latency to Lever Press Day 5 - Charles River C72 (n=
Observed −log10(p)
Expected −log10(p)

Plot Average Latency to Magazine Entry Day 1 - Charles River R09-P3/7/1

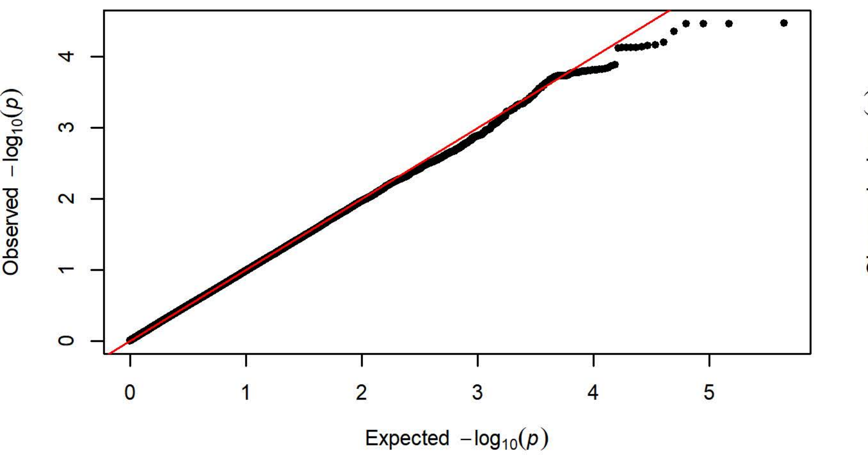


Q Plot Average Latency to Magazine Entry Day 1 - Harlan 202A/C-208A (n

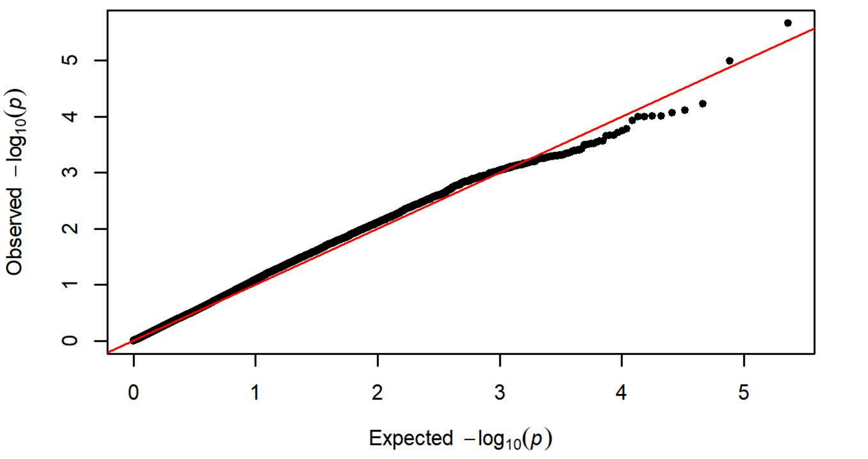


-Q Plot Average Latency to Magazine Entry Day 1 - Charles River R04 (n

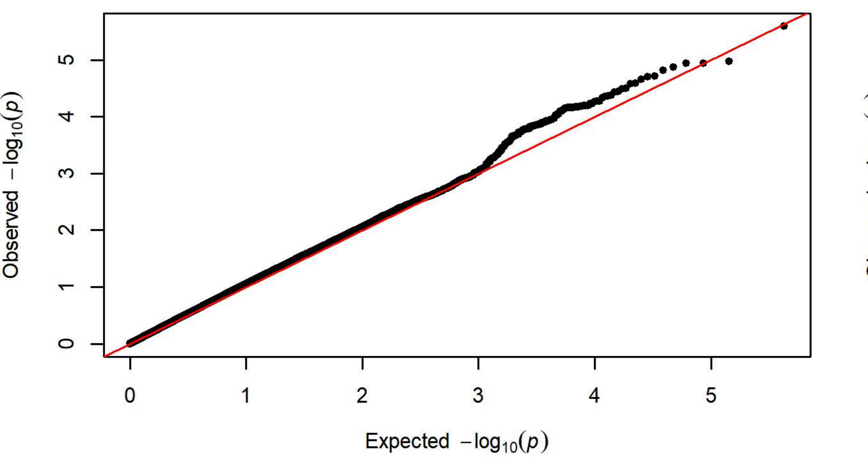


Q-Q Plot Average Latency to Magazine Entry Day 1 - Harlan 206 (n=75

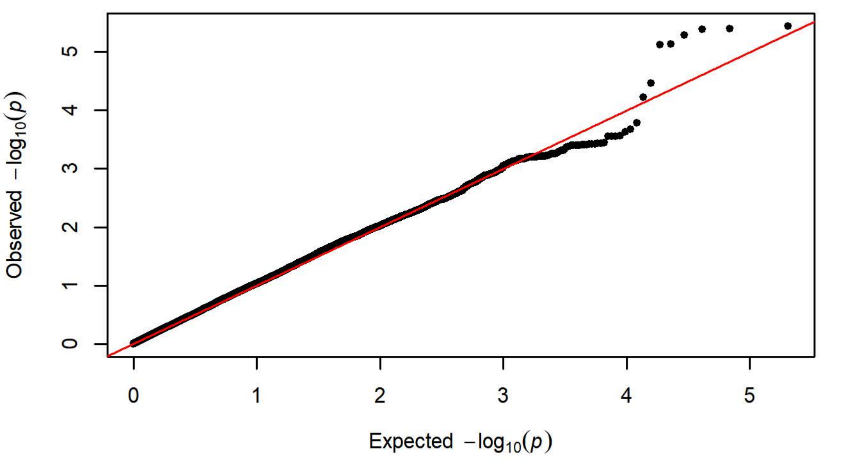


-Q Plot Average Latency to Magazine Entry Day 1 - Charles River P09 (n

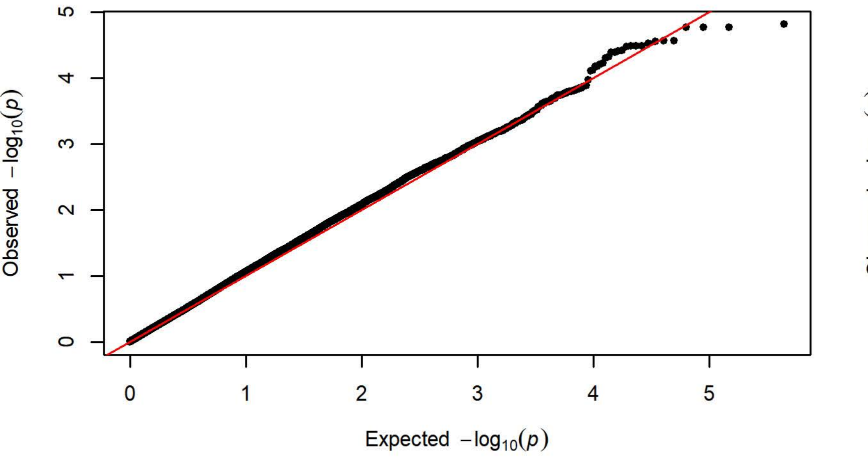


Q-Q Plot Average Latency to Magazine Entry Day 1 - Harlan 217 (n=35

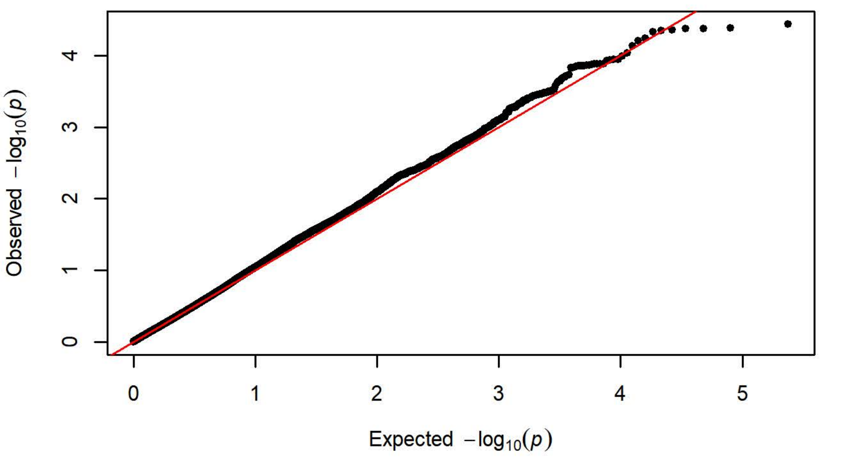


-Q Plot Average Latency to Magazine Entry Day 1 - Charles River C72 (n

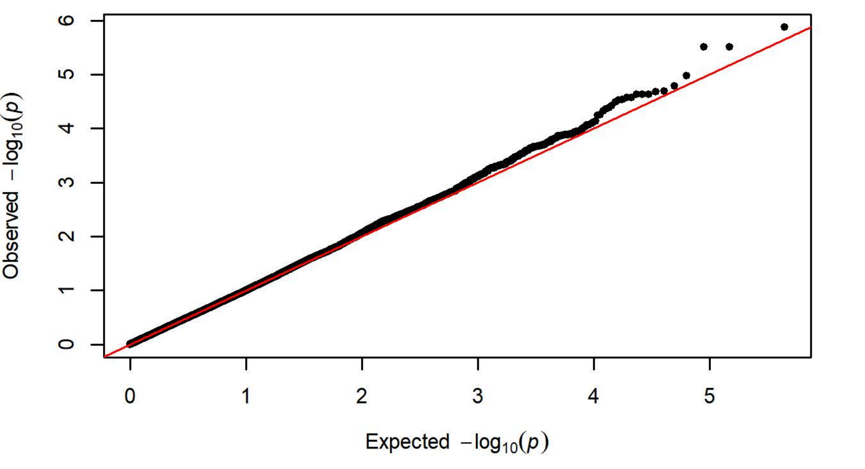

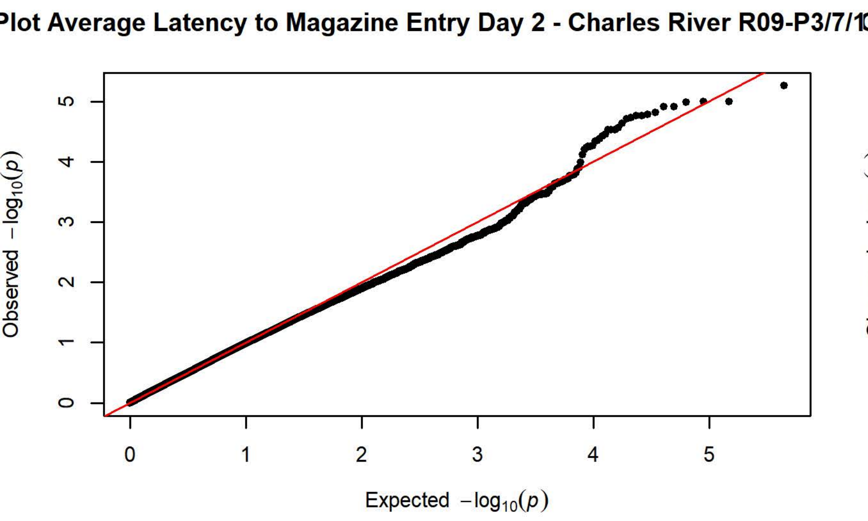
Plot Average Latency to Magazine Entry Day 2 - Charles River R09-P3/7/1
Observed $-\log_{10}(p)$
Expected $-\log_{10}(p)$

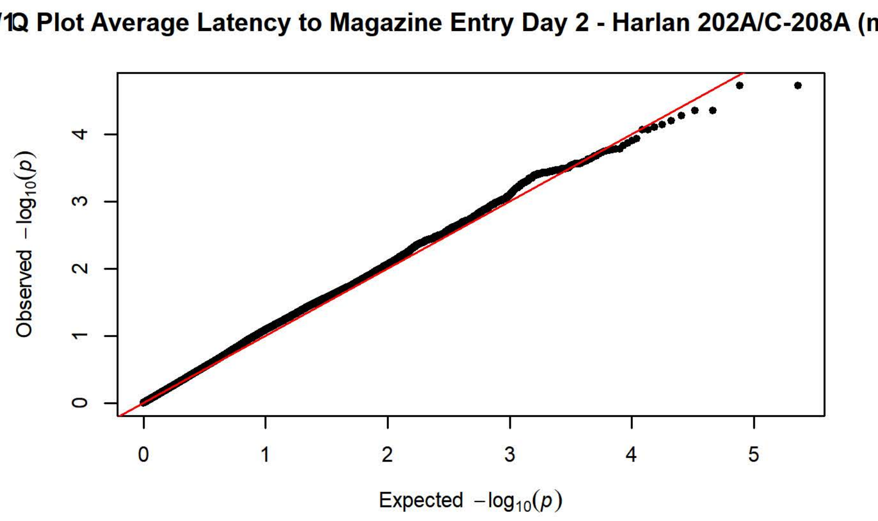
Q Plot Average Latency to Magazine Entry Day 2 - Harlan 202A/C-208A (n
Observed $-\log_{10}(p)$
Expected $-\log_{10}(p)$

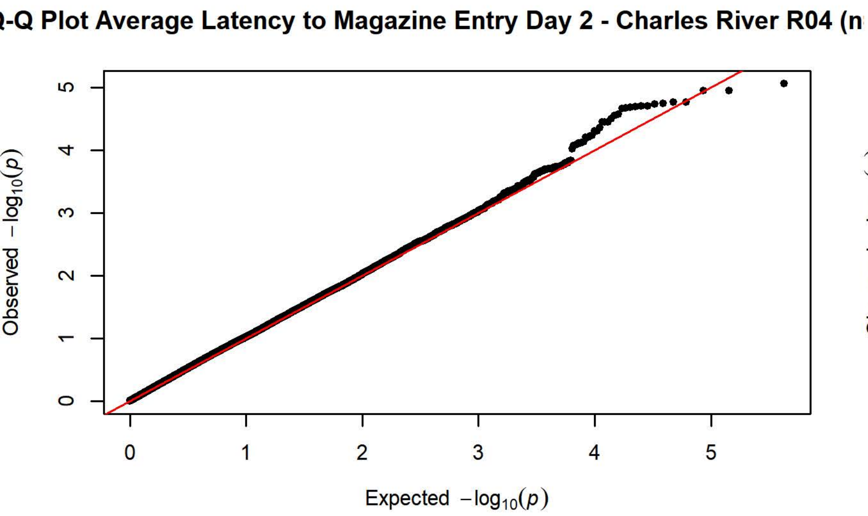
-Q Plot Average Latency to Magazine Entry Day 2 - Charles River R04 (n
Observed $-\log_{10}(p)$
Expected $-\log_{10}(p)$

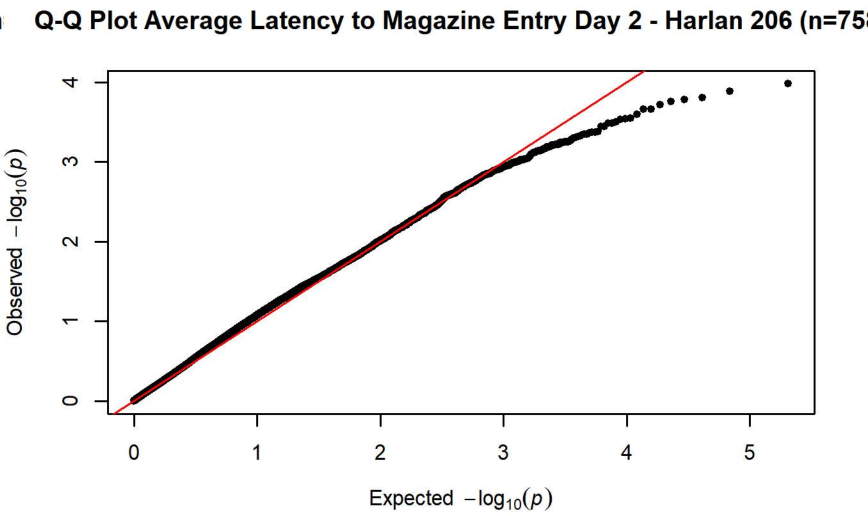
Q-Q Plot Average Latency to Magazine Entry Day 2 - Harlan 206 (n=75
Observed $-\log_{10}(p)$
Expected $-\log_{10}(p)$

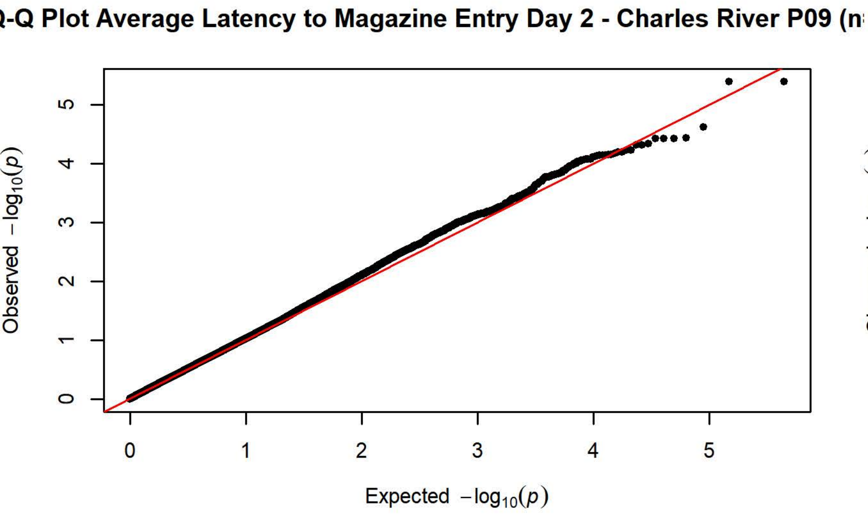
-Q Plot Average Latency to Magazine Entry Day 2 - Charles River P09 (n
Observed $-\log_{10}(p)$
Expected $-\log_{10}(p)$

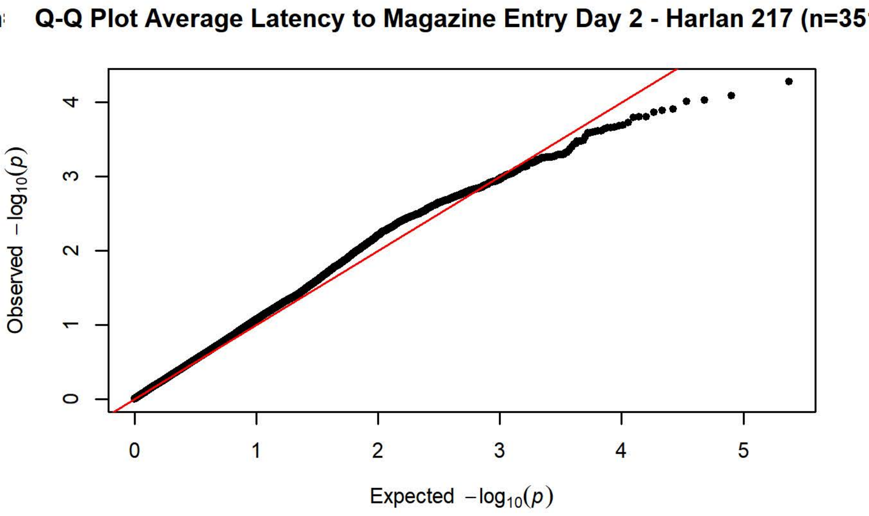
Q-Q Plot Average Latency to Magazine Entry Day 2 - Harlan 217 (n=35
Observed $-\log_{10}(p)$
Expected $-\log_{10}(p)$

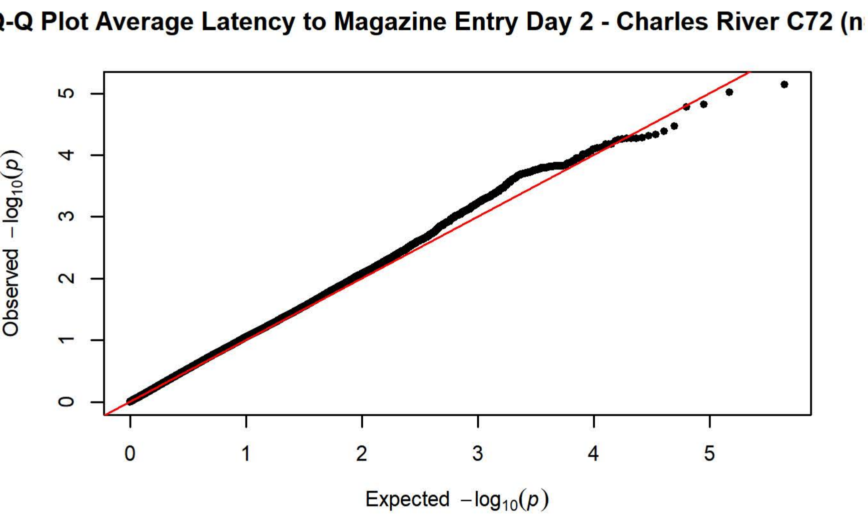
-Q Plot Average Latency to Magazine Entry Day 2 - Charles River C72 (n
Observed $-\log_{10}(p)$
Expected $-\log_{10}(p)$

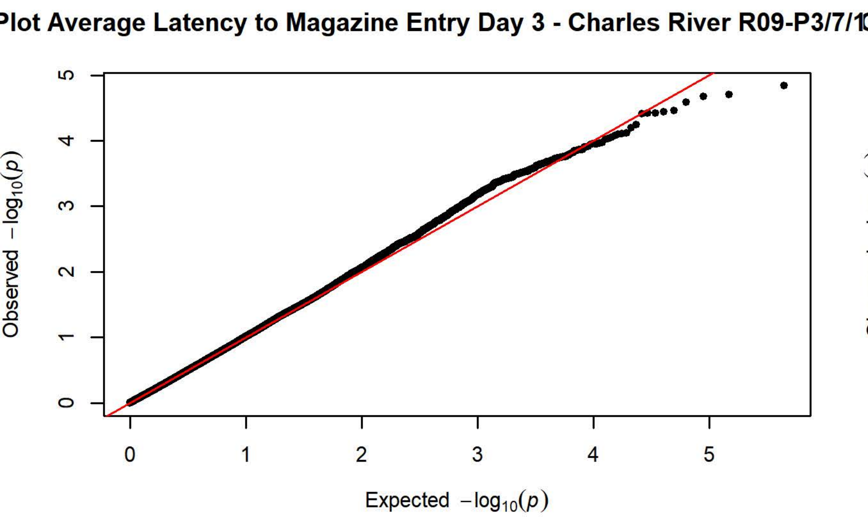
Plot Average Latency to Magazine Entry Day 3 - Charles River R09-P3/7/1
Observed −log10(p)
Expected −log10(p)

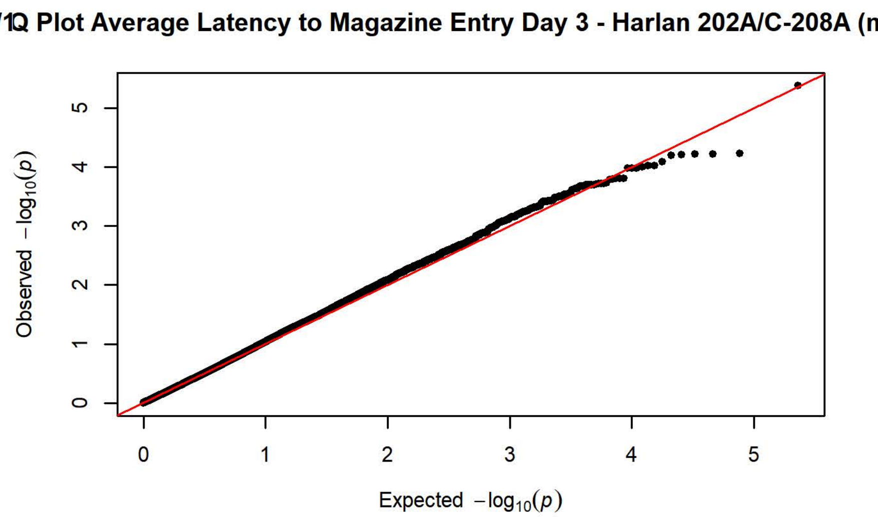
Q Plot Average Latency to Magazine Entry Day 3 - Harlan 202A/C-208A (n
Observed −log10(p)
Expected −log10(p)

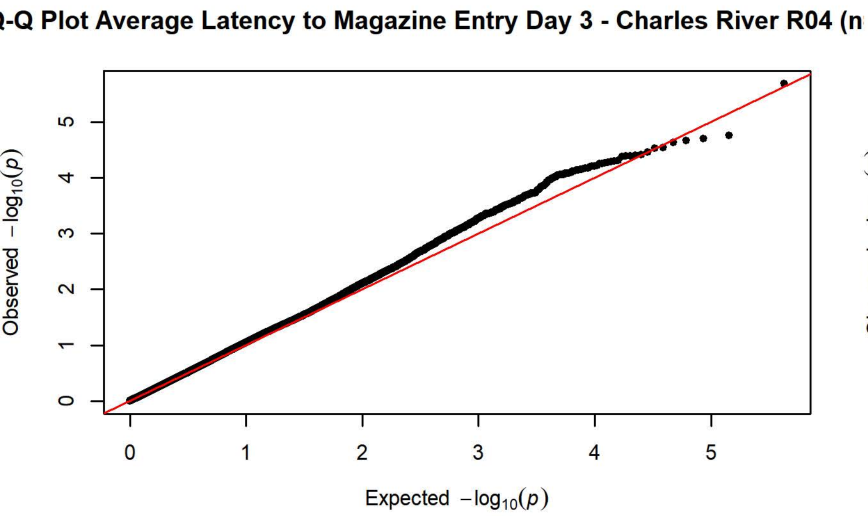
Q-Q Plot Average Latency to Magazine Entry Day 3 - Charles River R04 (n
Observed −log10(p)
Expected −log10(p)

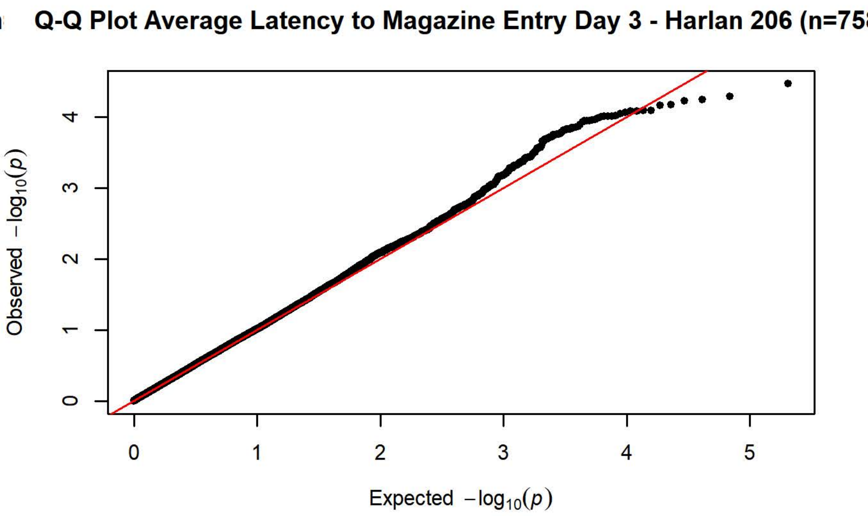
Q-Q Plot Average Latency to Magazine Entry Day 3 - Harlan 206 (n=75
Observed −log10(p)
Expected −log10(p)

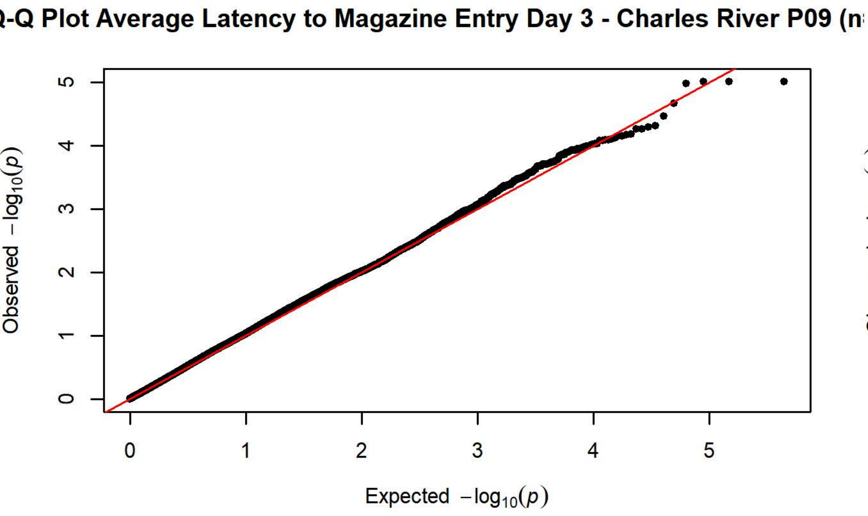
Q-Q Plot Average Latency to Magazine Entry Day 3 - Charles River P09 (n
Observed −log10(p)
Expected −log10(p)

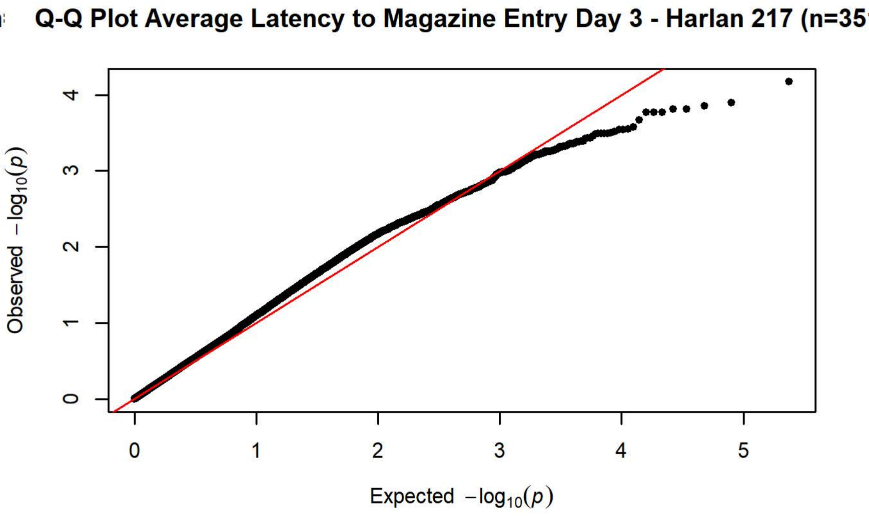
Q-Q Plot Average Latency to Magazine Entry Day 3 - Harlan 217 (n=35
Observed −log10(p)
Expected −log10(p)

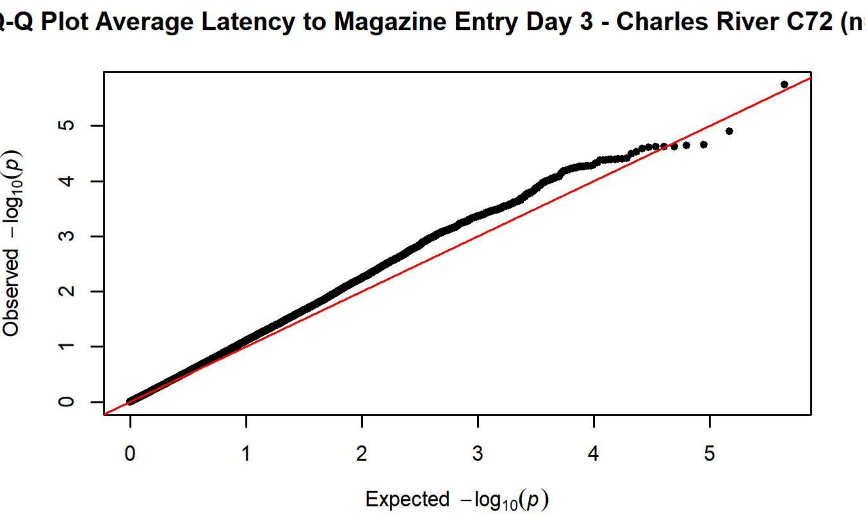
Q-Q Plot Average Latency to Magazine Entry Day 3 - Charles River C72 (n
Observed −log10(p)
Expected −log10(p)

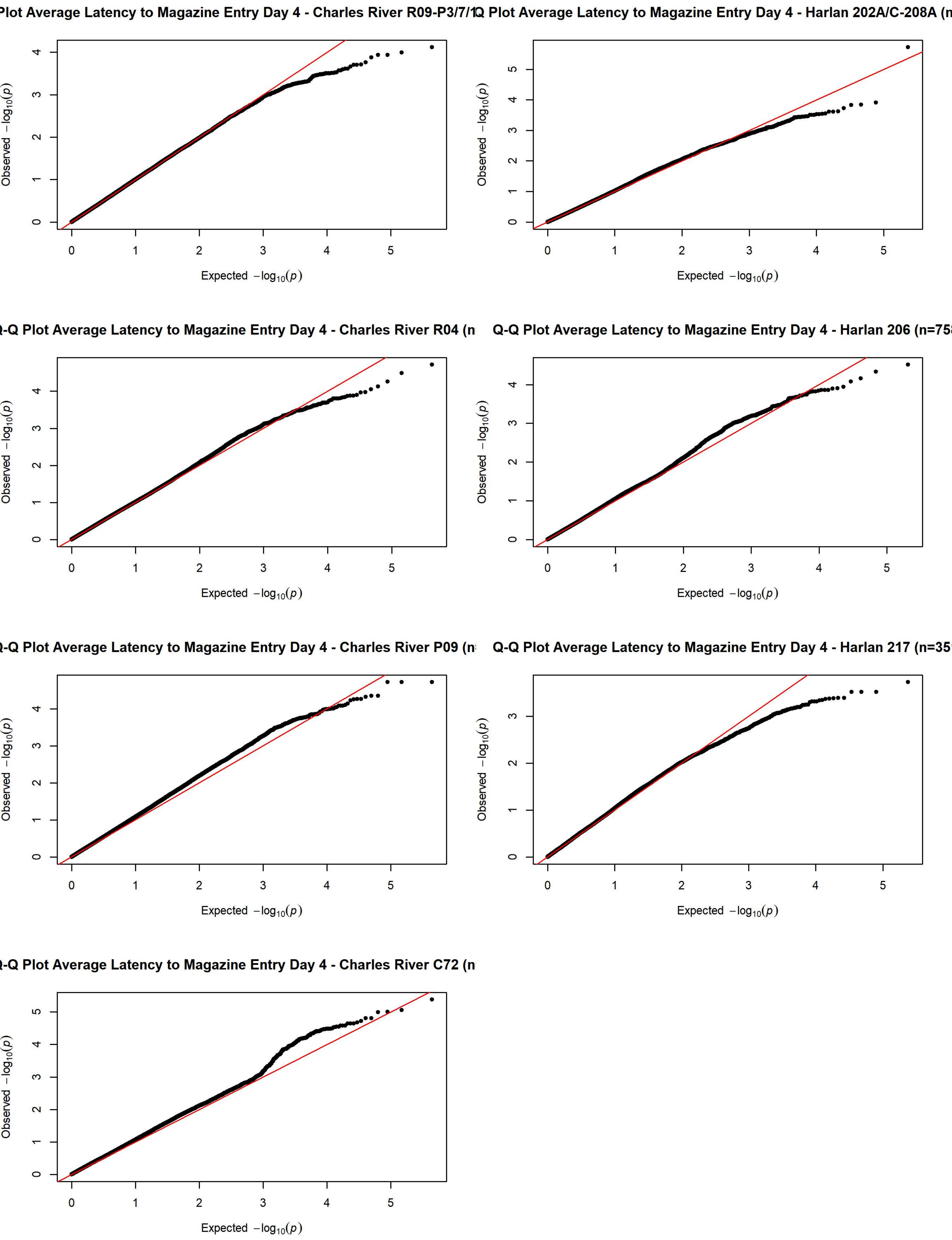

Plot Average Latency to Magazine Entry Day 4 - Charles River R09-P3/7/1
Q Plot Average Latency to Magazine Entry Day 4 - Harlan 202A/C-208A (n
-Q Plot Average Latency to Magazine Entry Day 4 - Charles River R04 (n
Q-Q Plot Average Latency to Magazine Entry Day 4 - Harlan 206 (n=75
-Q Plot Average Latency to Magazine Entry Day 4 - Charles River P09 (n
Q-Q Plot Average Latency to Magazine Entry Day 4 - Harlan 217 (n=35
-Q Plot Average Latency to Magazine Entry Day 4 - Charles River C72 (n
Expected $-\log_{10}(p)$
Observed $-\log_{10}(p)$

Plot Average Latency to Magazine Entry Day 5 - Charles River R09-P3/7/1

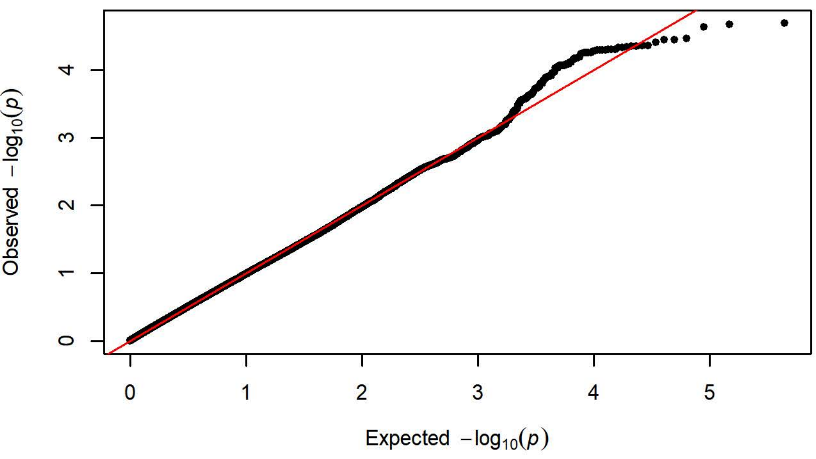


Q Plot Average Latency to Magazine Entry Day 5 - Harlan 202A/C-208A (n

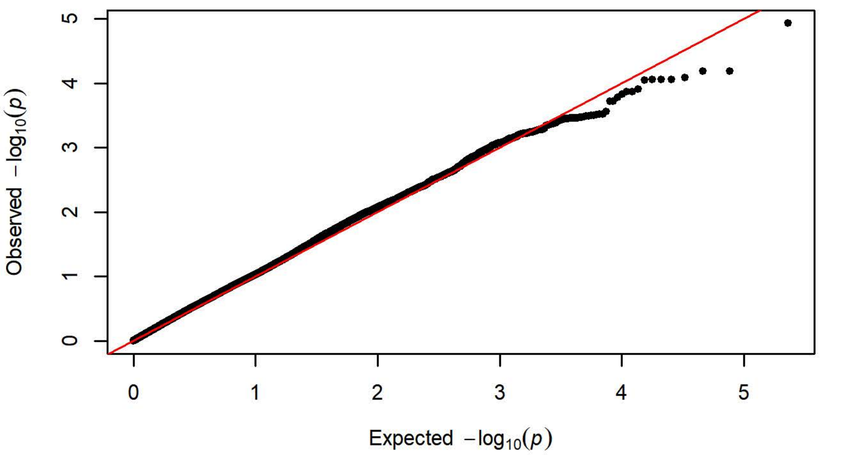


-Q Plot Average Latency to Magazine Entry Day 5 - Charles River R04 (n

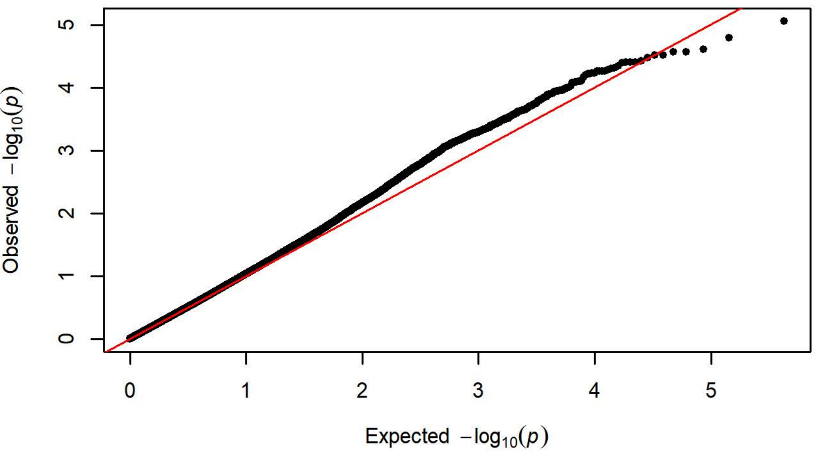


Q-Q Plot Average Latency to Magazine Entry Day 5 - Harlan 206 (n=75

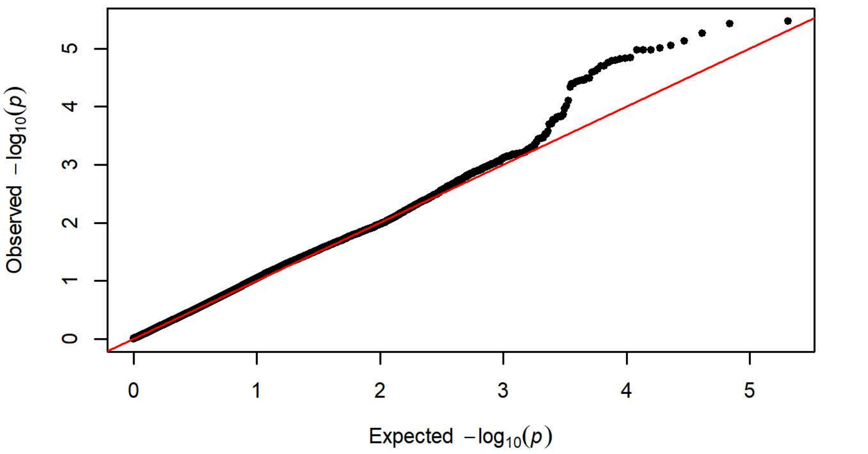


-Q Plot Average Latency to Magazine Entry Day 5 - Charles River P09 (n

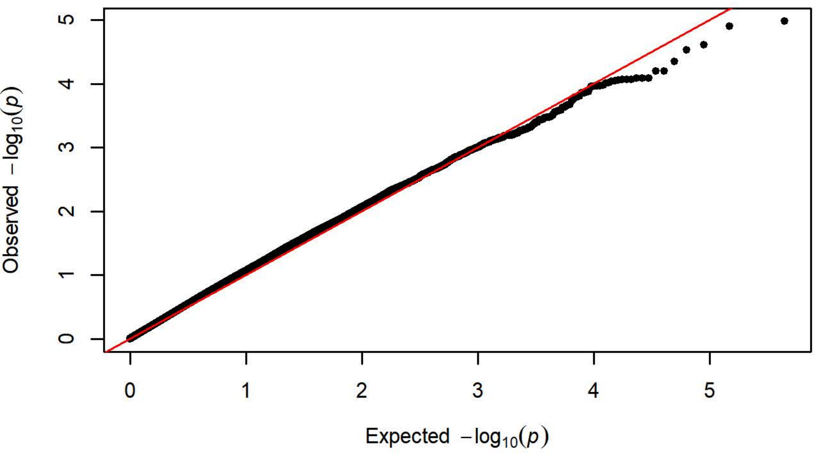


Q-Q Plot Average Latency to Magazine Entry Day 5 - Harlan 217 (n=35

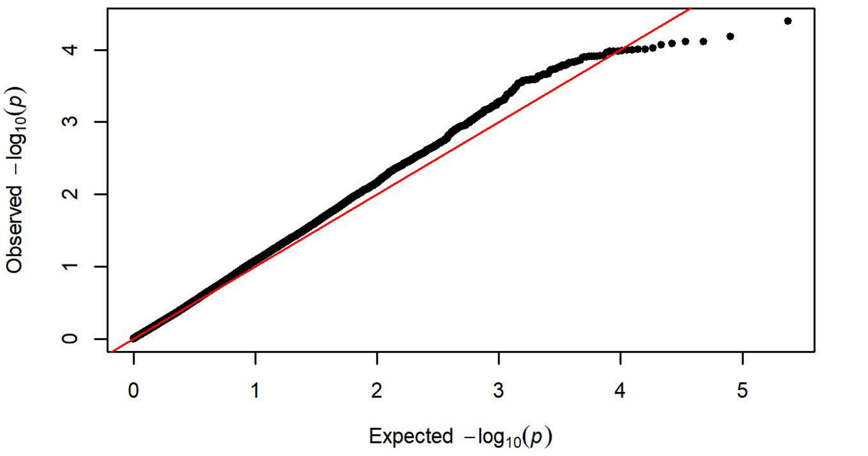


-Q Plot Average Latency to Magazine Entry Day 5 - Charles River C72 (n

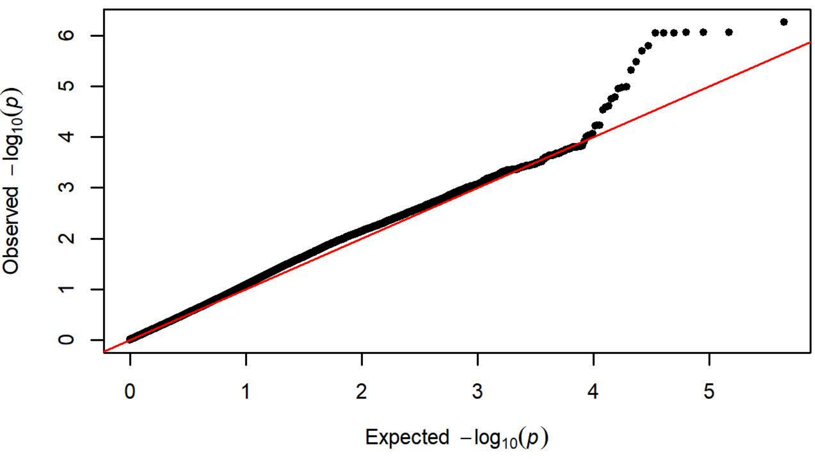

Q-Q Plot PavCA Index Score Day 1 - Charles River R09-P3/7/10 (n=422

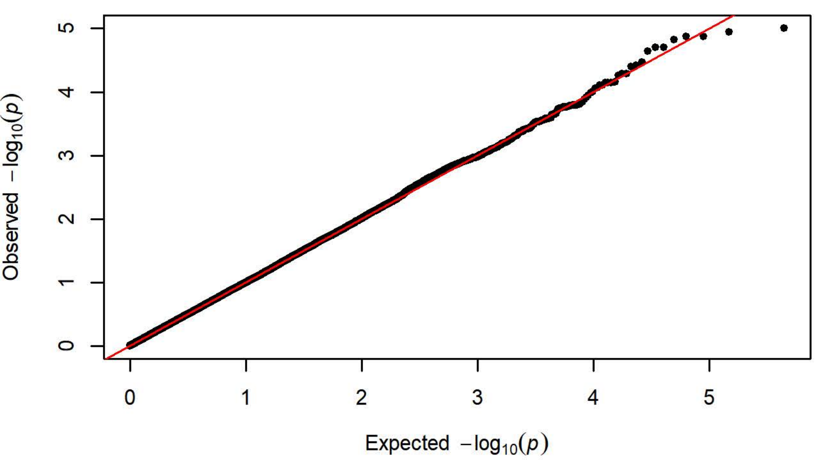


Q-Q Plot PavCA Index Score Day 1 - Harlan 202A/C-208A (n=1062)

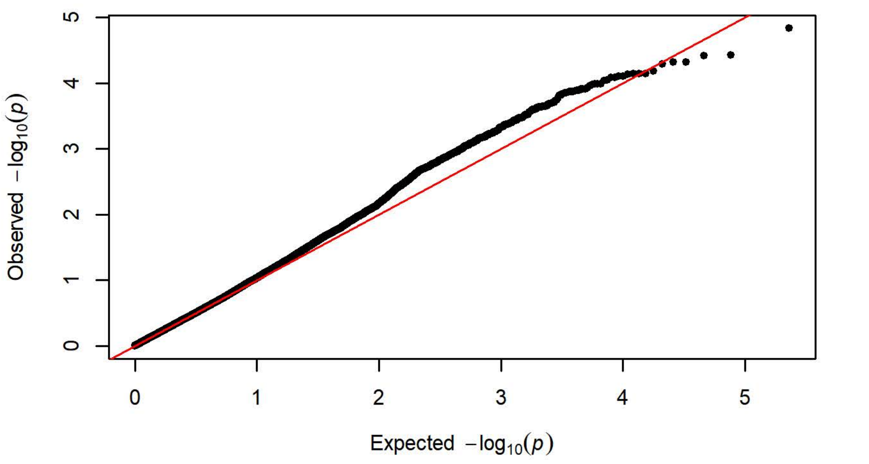


Q-Q Plot PavCA Index Score Day 1 - Charles River R04 (n=648)

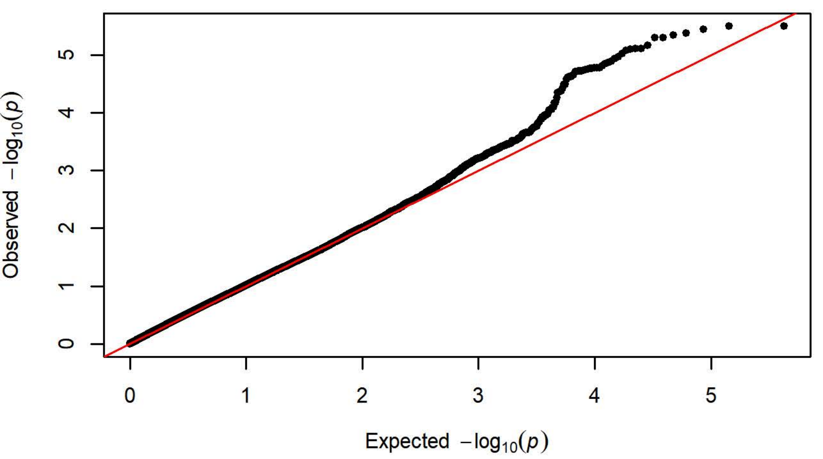


Q-Q Plot PavCA Index Score Day 1 - Harlan 206 (n=752)

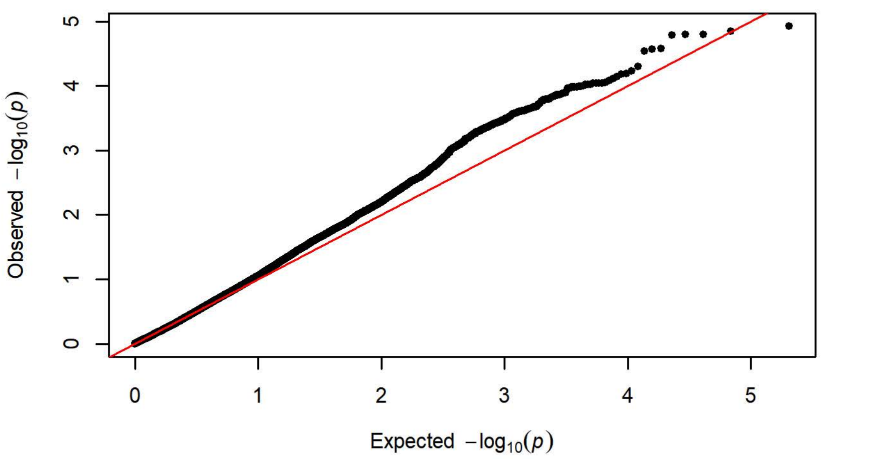


Q-Q Plot PavCA Index Score Day 1 - Charles River P09 (n=293)

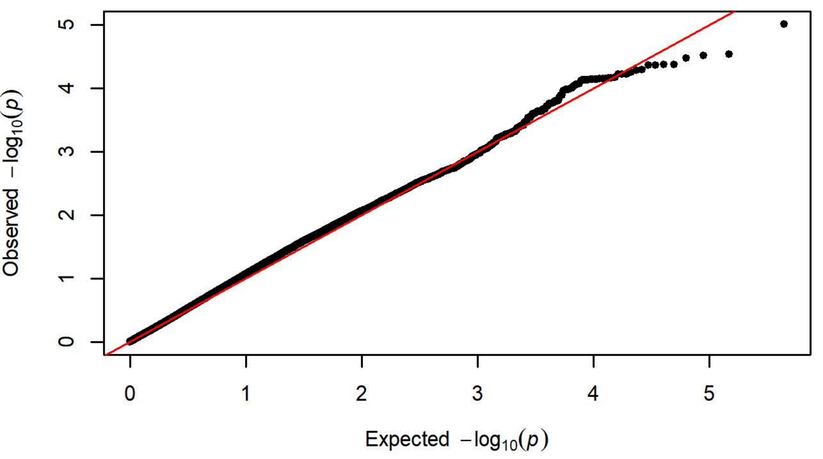


Q-Q Plot PavCA Index Score Day 1 - Harlan 217 (n=346)

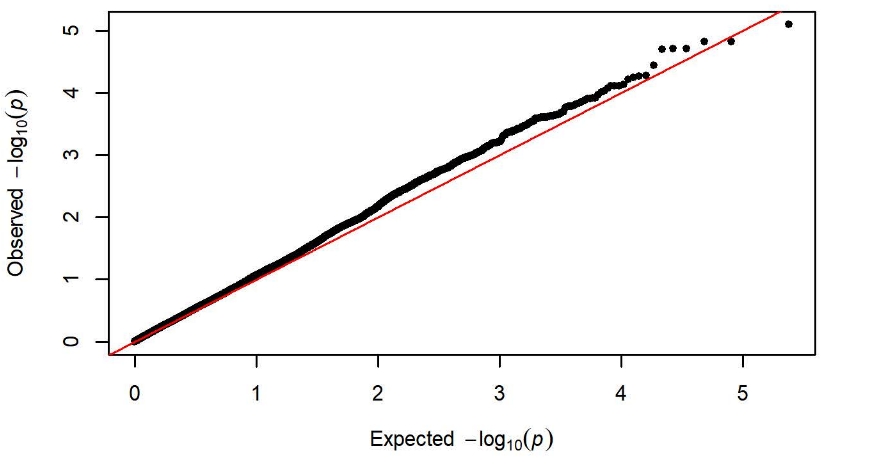


Q-Q Plot PavCA Index Score Day 1 - Charles River C72 (n=357)

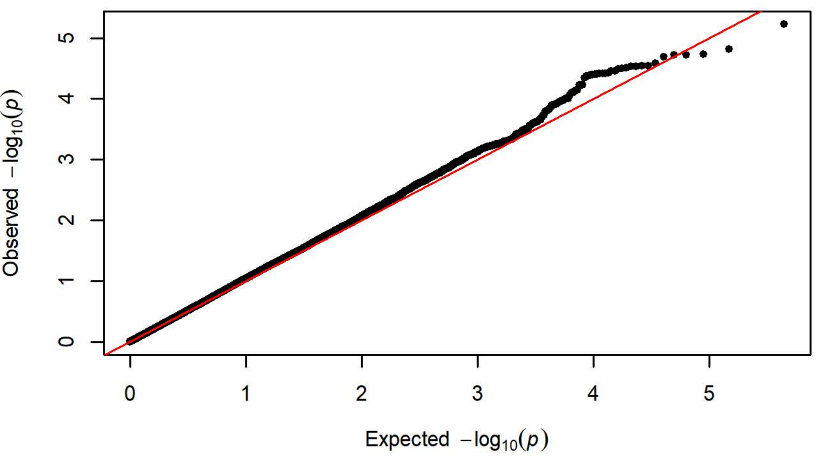

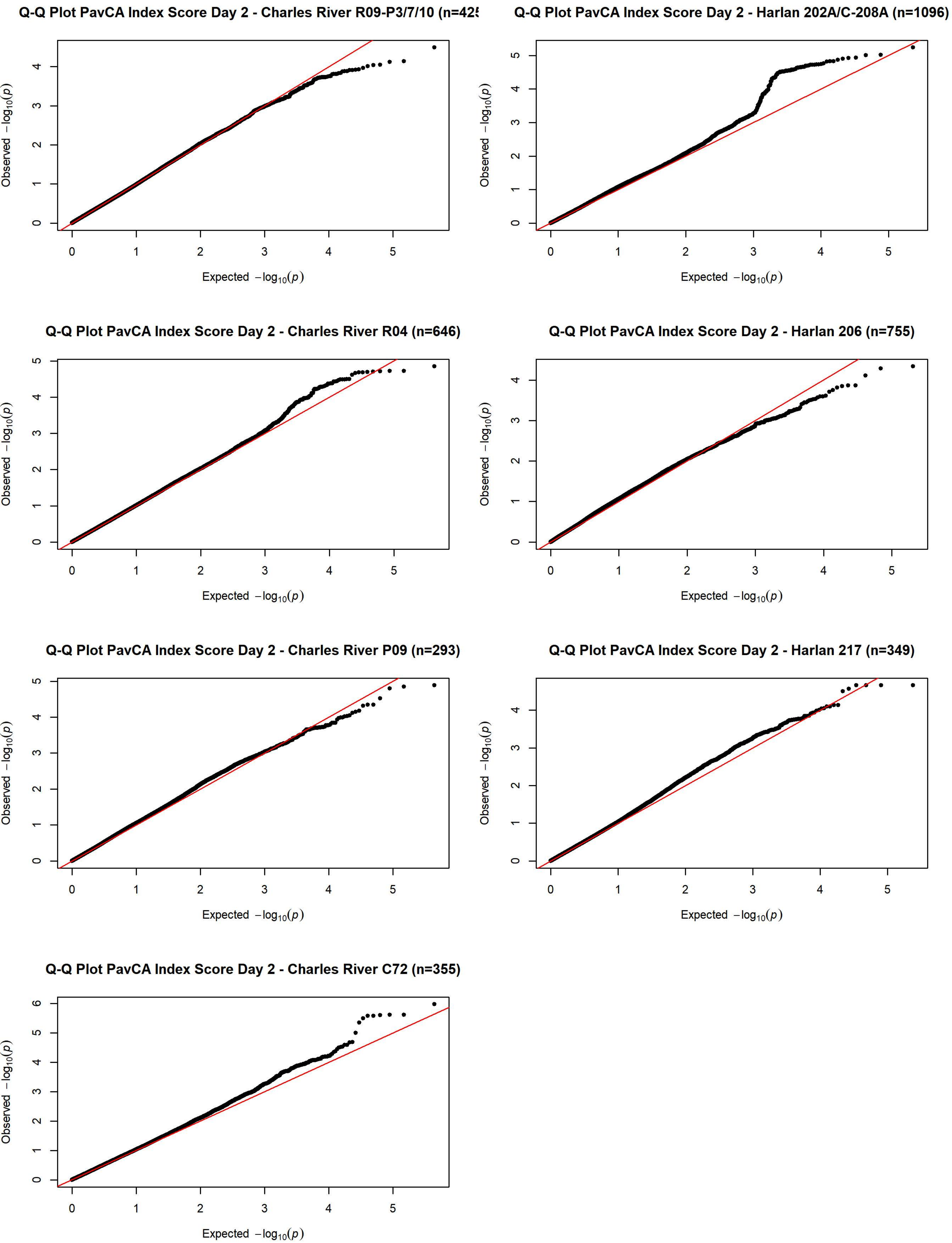
Q-Q Plot PavCA Index Score Day 2 - Charles River R09-P3/7/10 (n=42
Q-Q Plot PavCA Index Score Day 2 - Harlan 202A/C-208A (n=1096)
Q-Q Plot PavCA Index Score Day 2 - Charles River R04 (n=646)
Q-Q Plot PavCA Index Score Day 2 - Harlan 206 (n=755)
Q-Q Plot PavCA Index Score Day 2 - Charles River P09 (n=293)
Q-Q Plot PavCA Index Score Day 2 - Harlan 217 (n=349)
Q-Q Plot PavCA Index Score Day 2 - Charles River C72 (n=355)
Observed $-\log_{10}(p)$
Expected $-\log_{10}(p)$

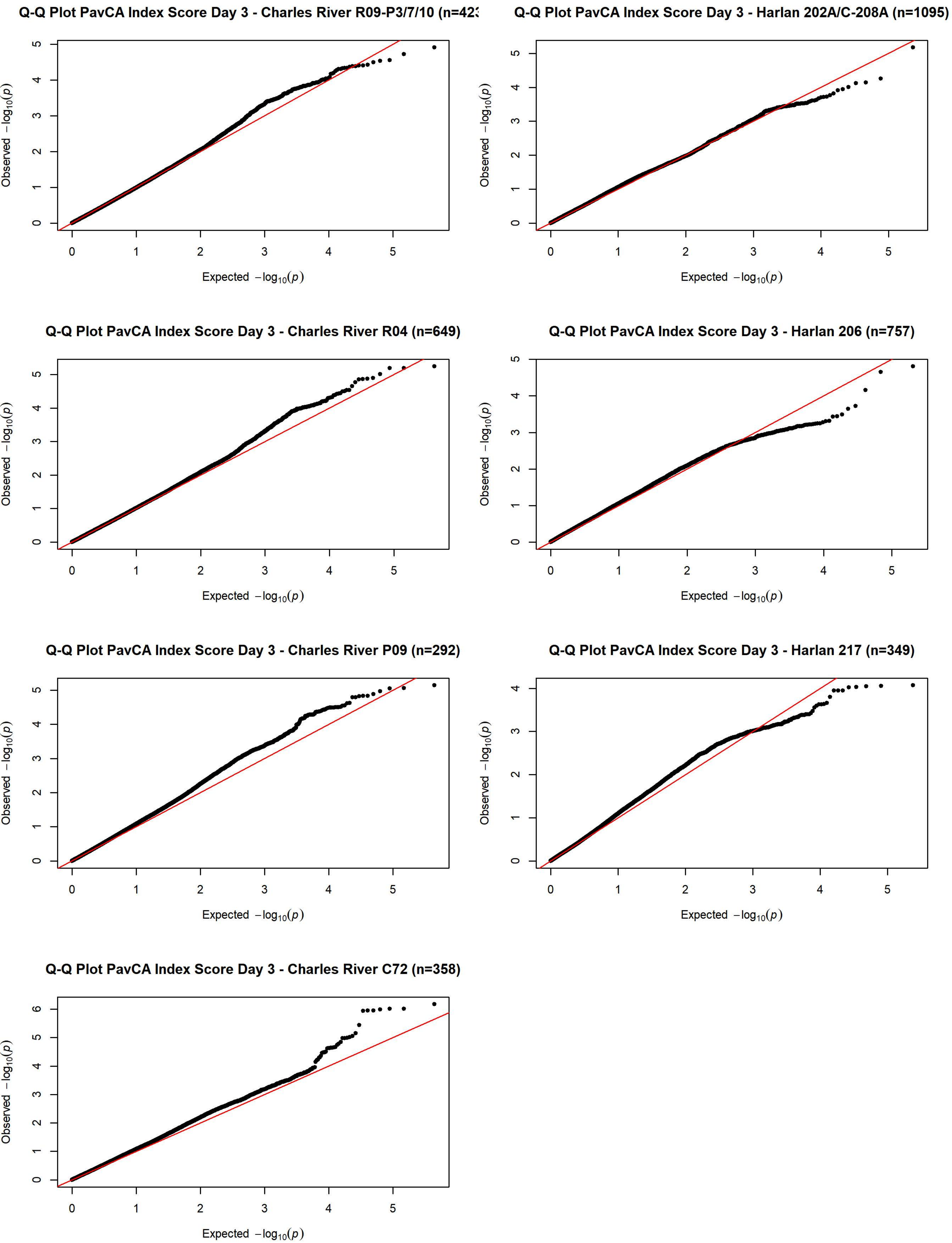
Q-Q Plot PavCA Index Score Day 3 - Charles River R09-P3/7/10 (n=423
Q-Q Plot PavCA Index Score Day 3 - Harlan 202A/C-208A (n=1095)
Q-Q Plot PavCA Index Score Day 3 - Charles River R04 (n=649)
Q-Q Plot PavCA Index Score Day 3 - Harlan 206 (n=757)
Q-Q Plot PavCA Index Score Day 3 - Charles River P09 (n=292)
Q-Q Plot PavCA Index Score Day 3 - Harlan 217 (n=349)
Q-Q Plot PavCA Index Score Day 3 - Charles River C72 (n=358)
Expected $-\log_{10}(p)$
Observed $-\log_{10}(p)$

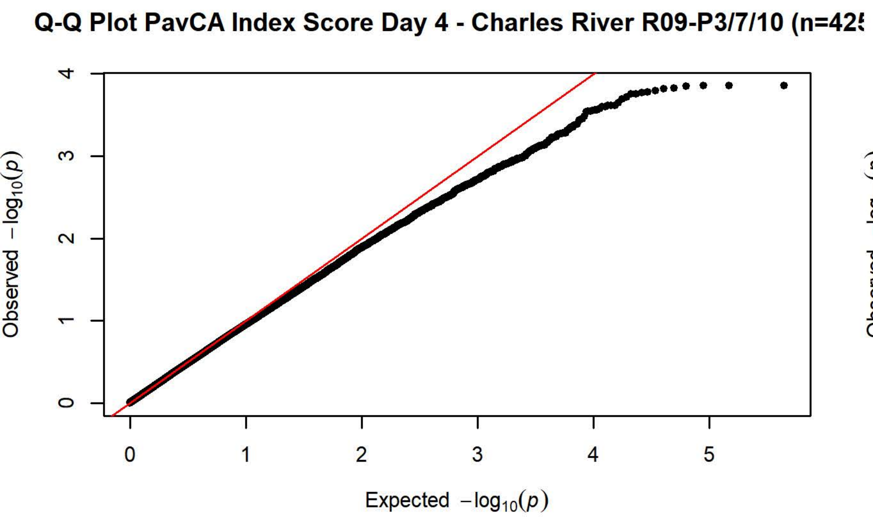
Q-Q Plot PavCA Index Score Day 4 - Charles River R09-P3/7/10 (n=42
Observed $-\log_{10}(p)$
Expected $-\log_{10}(p)$

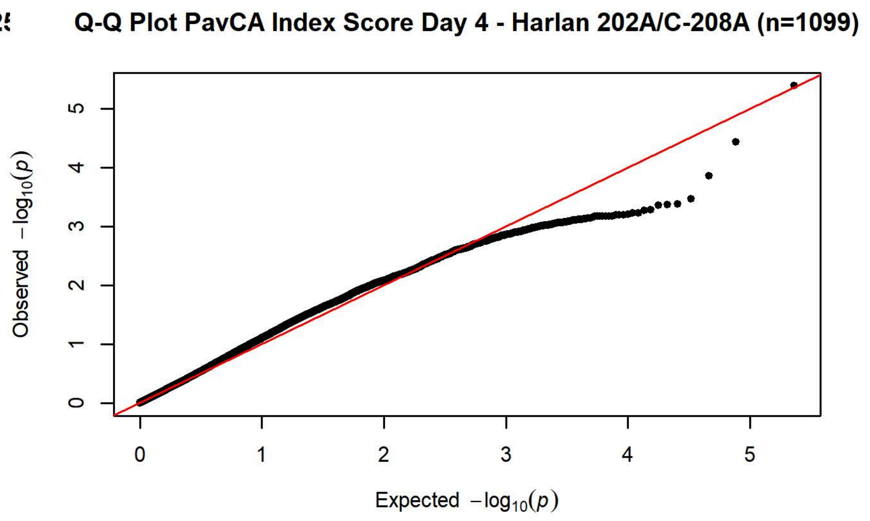
Q-Q Plot PavCA Index Score Day 4 - Harlan 202A/C-208A (n=1099)
Observed $-\log_{10}(p)$
Expected $-\log_{10}(p)$

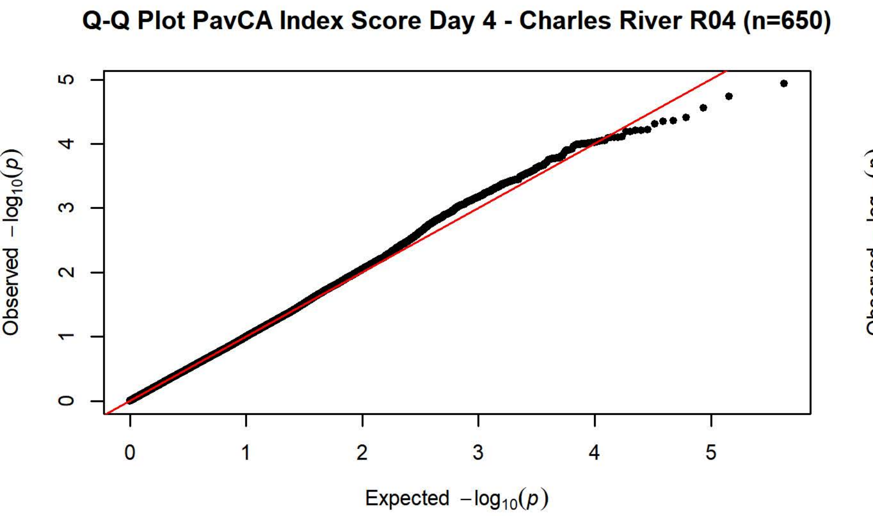
Q-Q Plot PavCA Index Score Day 4 - Charles River R04 (n=650)
Observed $-\log_{10}(p)$
Expected $-\log_{10}(p)$

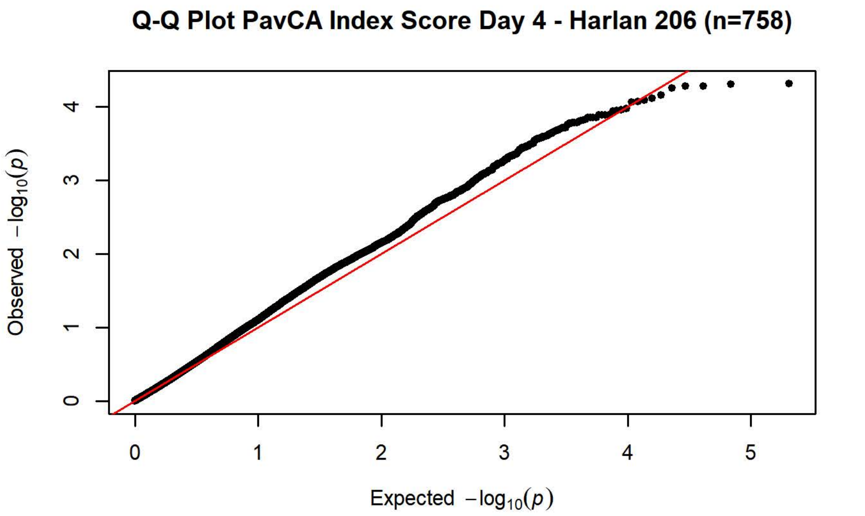
Q-Q Plot PavCA Index Score Day 4 - Harlan 206 (n=758)
Observed $-\log_{10}(p)$
Expected $-\log_{10}(p)$

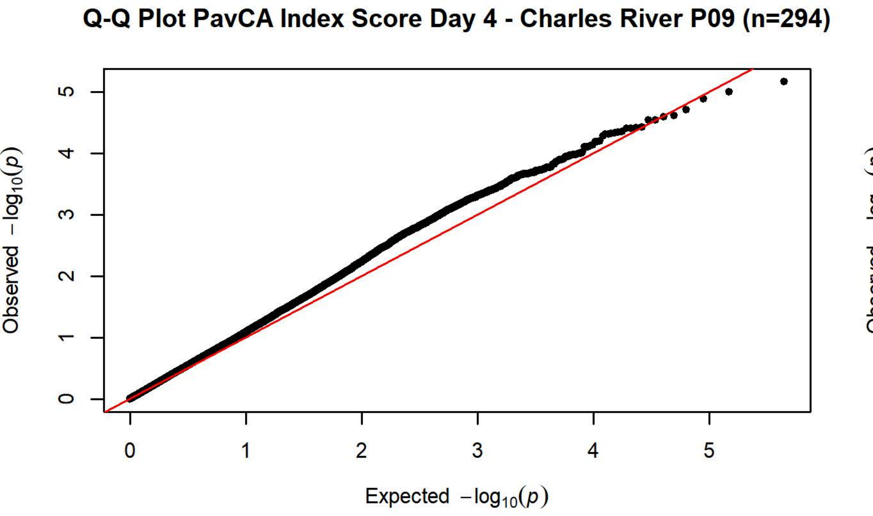
Q-Q Plot PavCA Index Score Day 4 - Charles River P09 (n=294)
Observed $-\log_{10}(p)$
Expected $-\log_{10}(p)$

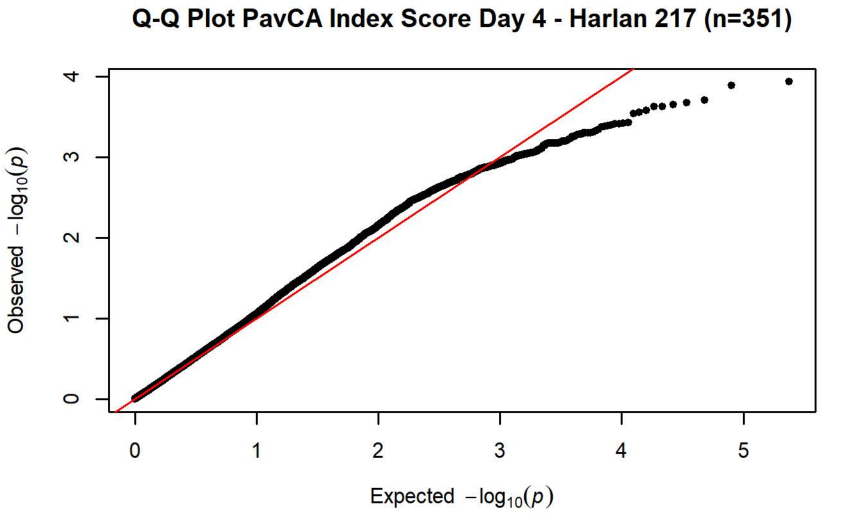
Q-Q Plot PavCA Index Score Day 4 - Harlan 217 (n=351)
Observed $-\log_{10}(p)$
Expected $-\log_{10}(p)$

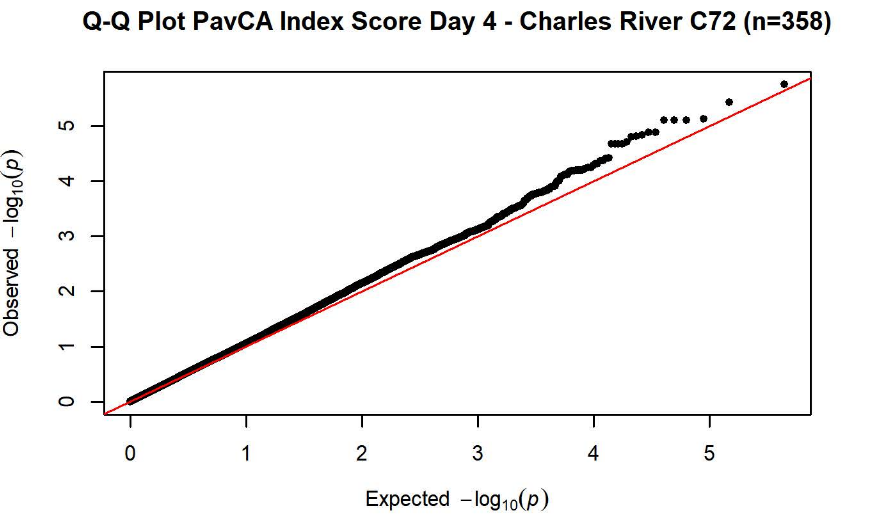
Q-Q Plot PavCA Index Score Day 4 - Charles River C72 (n=358)
Observed $-\log_{10}(p)$
Expected $-\log_{10}(p)$

**Q-Q Plot PavCA Index Score Day 5 - Charles River R09-P3/7/10 (n=42[illegible]**

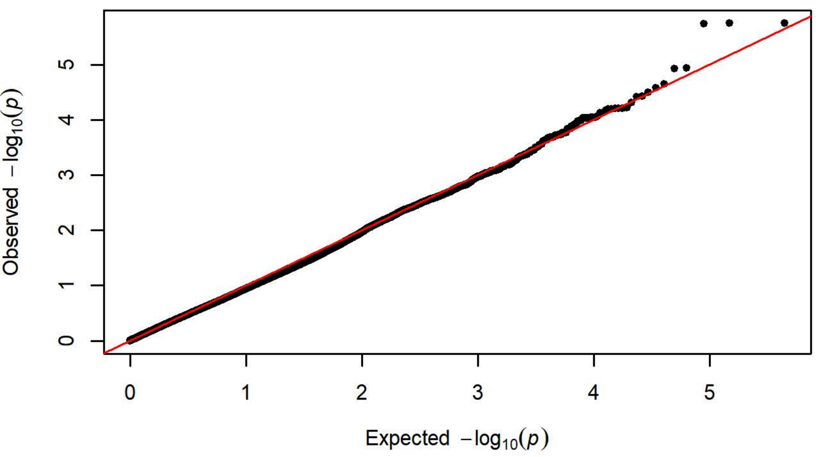


**Q-Q Plot PavCA Index Score Day 5 - Harlan 202A/C-208A (n=1099)**

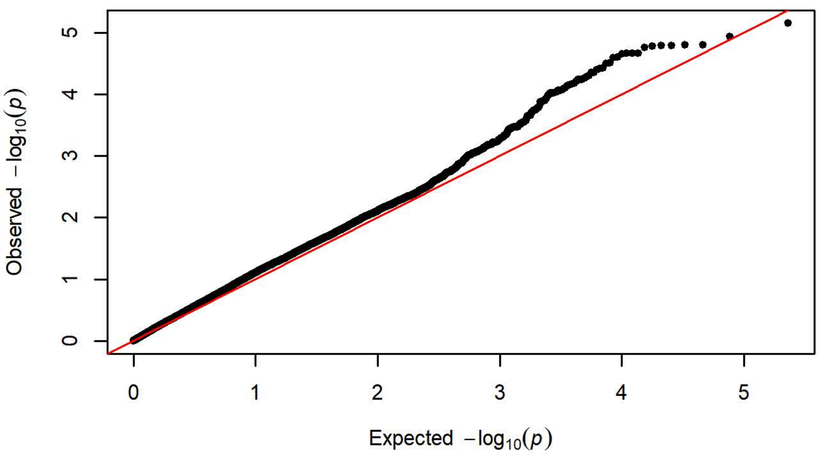


**Q-Q Plot PavCA Index Score Day 5 - Charles River R04 (n=650)**

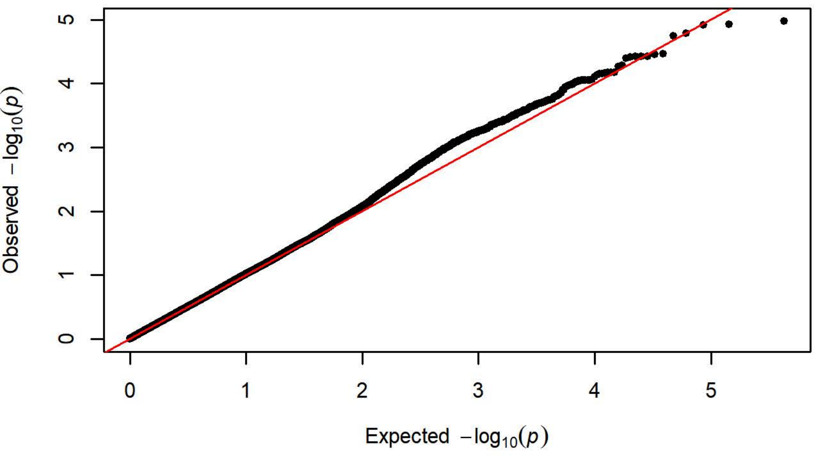


**Q-Q Plot PavCA Index Score Day 5 - Harlan 206 (n=757)**

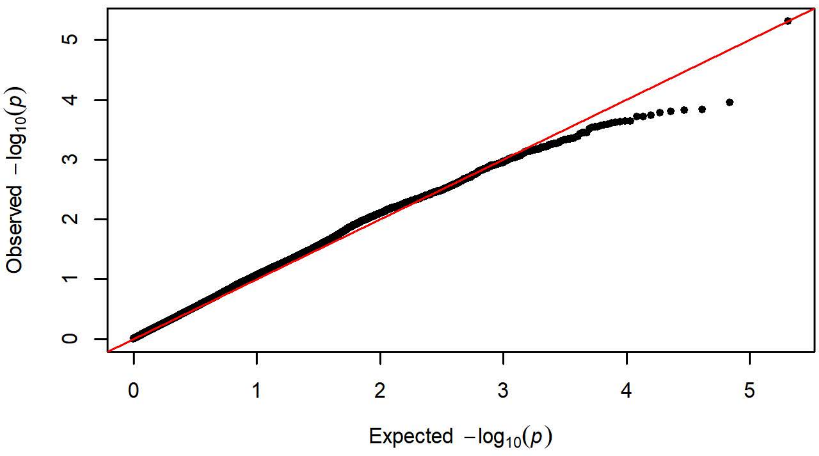


**Q-Q Plot PavCA Index Score Day 5 - Charles River P09 (n=293)**

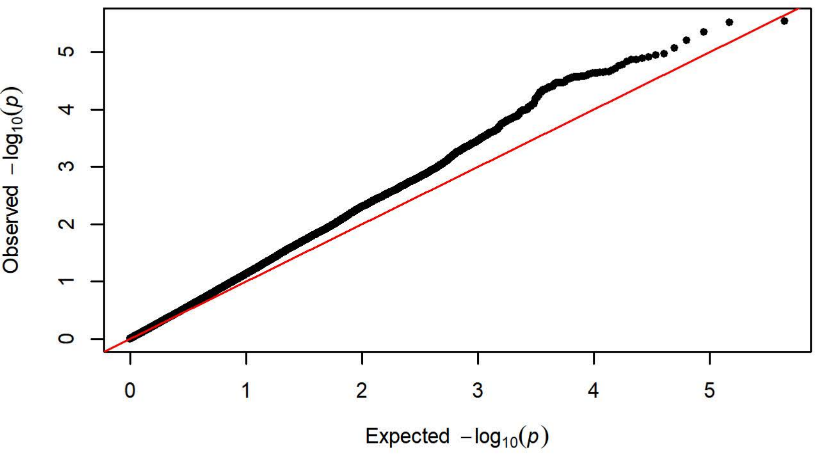


**Q-Q Plot PavCA Index Score Day 5 - Harlan 217 (n=351)**

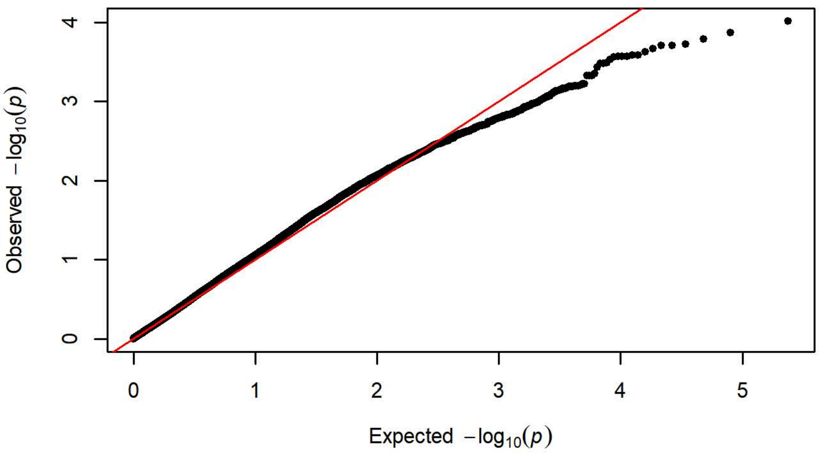


**Q-Q Plot PavCA Index Score Day 5 - Charles River C72 (n=358)**

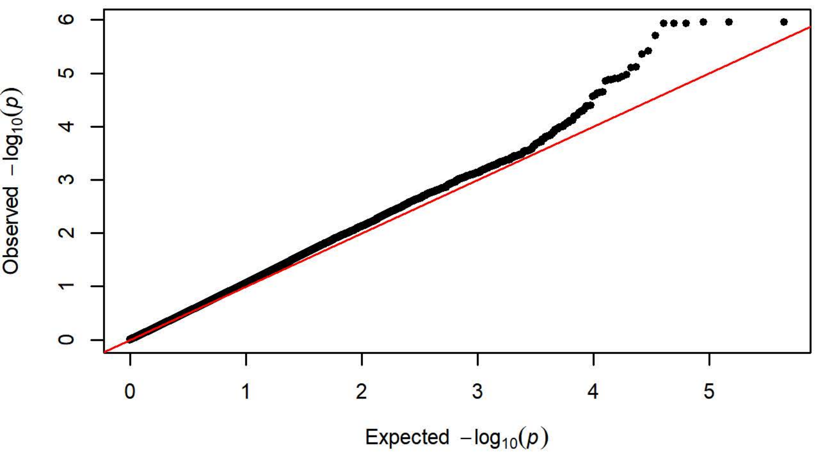

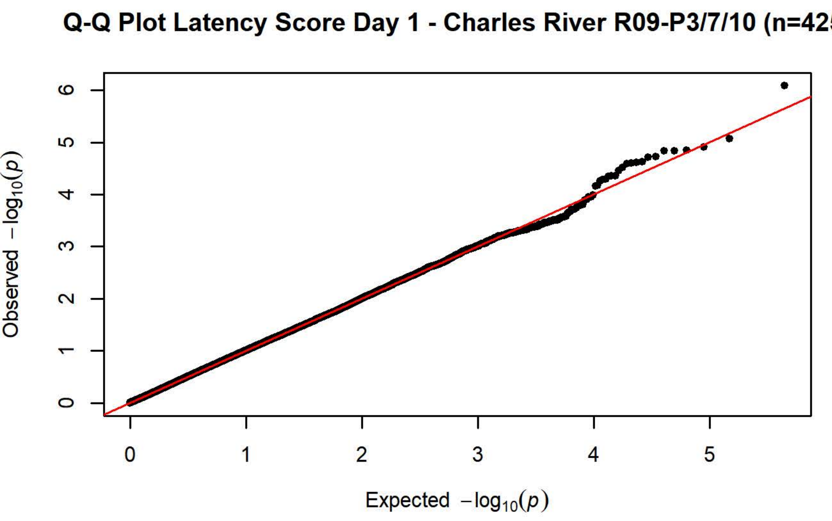
Q-Q Plot Latency Score Day 1 - Charles River R09-P3/7/10 (n=425)
Observed $-\log_{10}(p)$
Expected $-\log_{10}(p)$

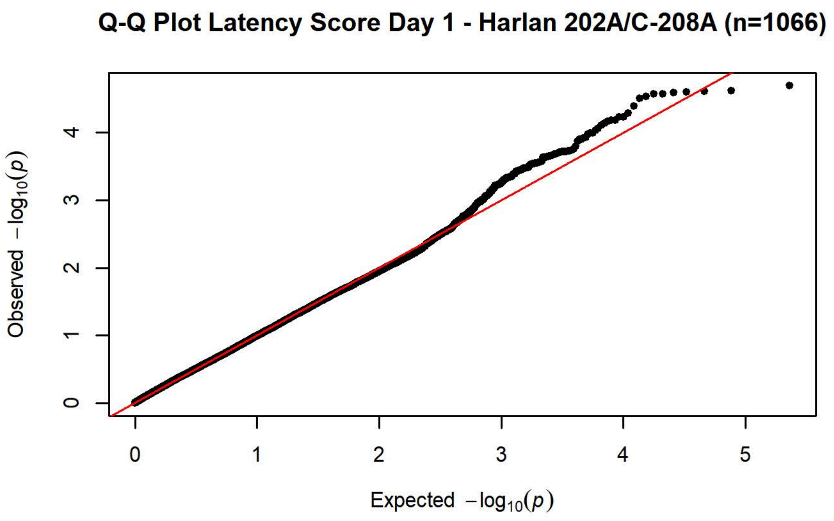
Q-Q Plot Latency Score Day 1 - Harlan 202A/C-208A (n=1066)
Observed $-\log_{10}(p)$
Expected $-\log_{10}(p)$

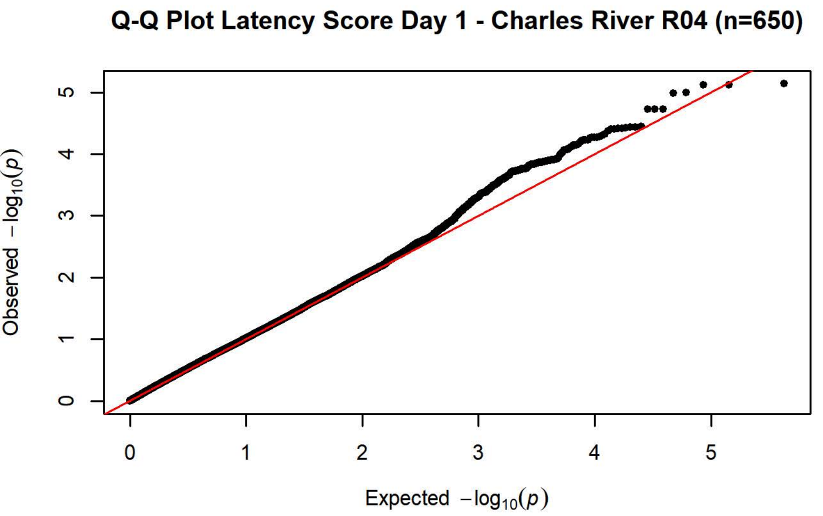
Q-Q Plot Latency Score Day 1 - Charles River R04 (n=650)
Observed $-\log_{10}(p)$
Expected $-\log_{10}(p)$

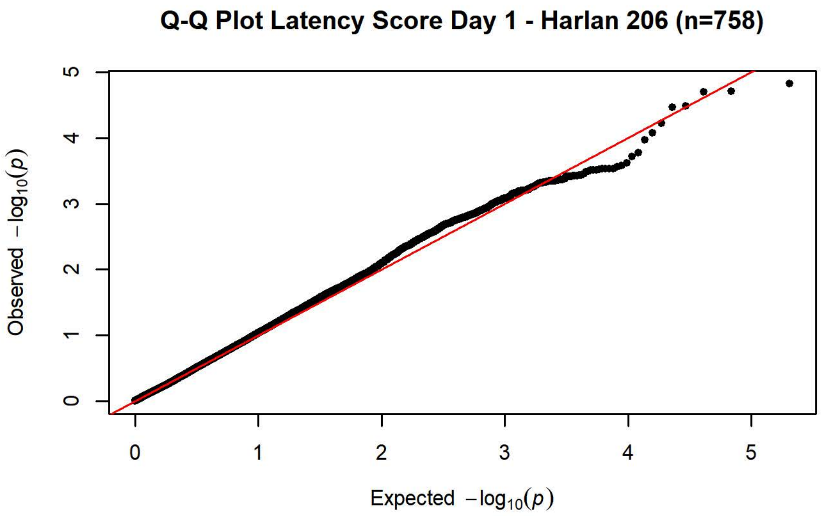
Q-Q Plot Latency Score Day 1 - Harlan 206 (n=758)
Observed $-\log_{10}(p)$
Expected $-\log_{10}(p)$

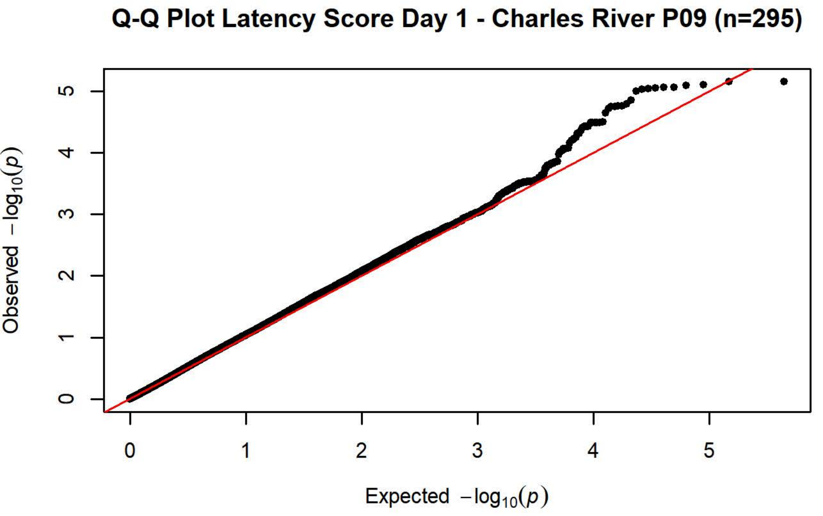
Q-Q Plot Latency Score Day 1 - Charles River P09 (n=295)
Observed $-\log_{10}(p)$
Expected $-\log_{10}(p)$

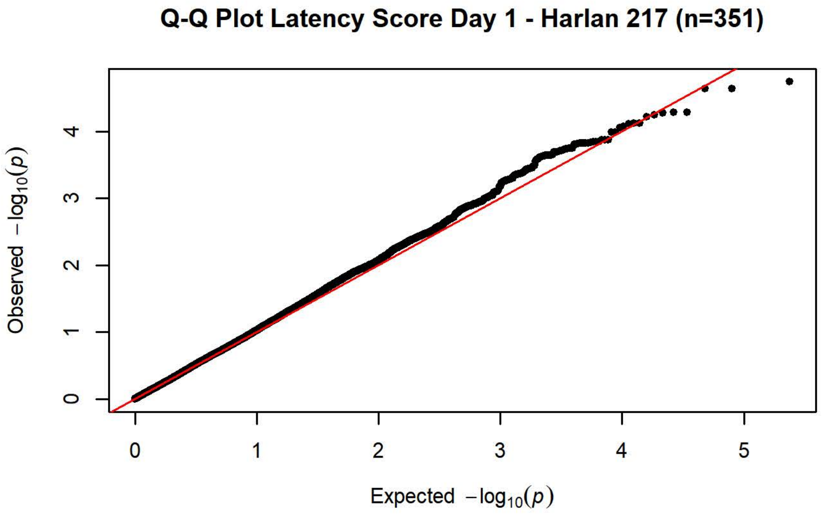
Q-Q Plot Latency Score Day 1 - Harlan 217 (n=351)
Observed $-\log_{10}(p)$
Expected $-\log_{10}(p)$

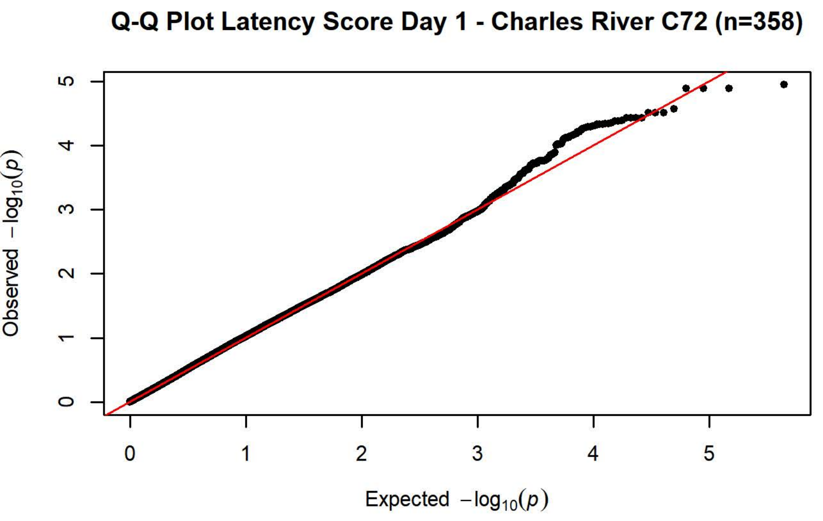
Q-Q Plot Latency Score Day 1 - Charles River C72 (n=358)
Observed $-\log_{10}(p)$
Expected $-\log_{10}(p)$

**Q-Q Plot Latency Score Day 2 - Charles River R09-P3/7/10 (n=425)**

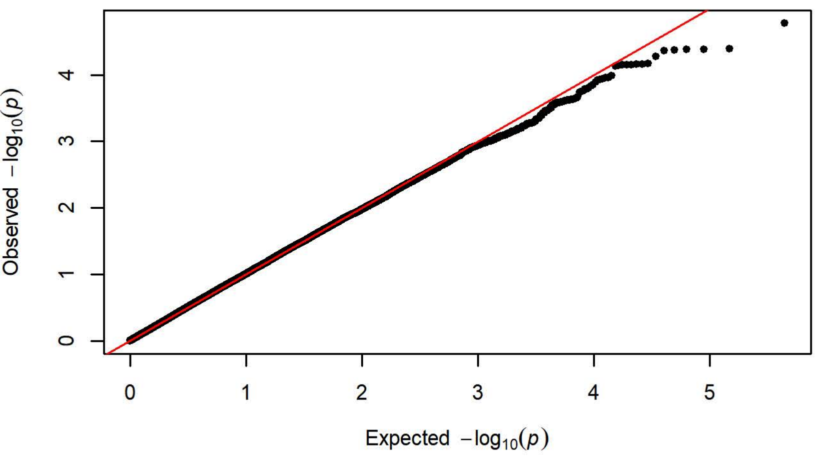


**Q-Q Plot Latency Score Day 2 - Harlan 202A/C-208A (n=1099)**

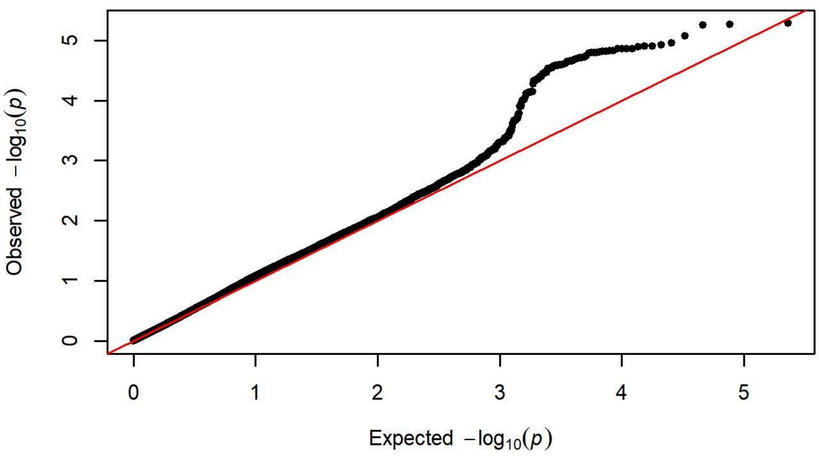


**Q-Q Plot Latency Score Day 2 - Charles River R04 (n=650)**

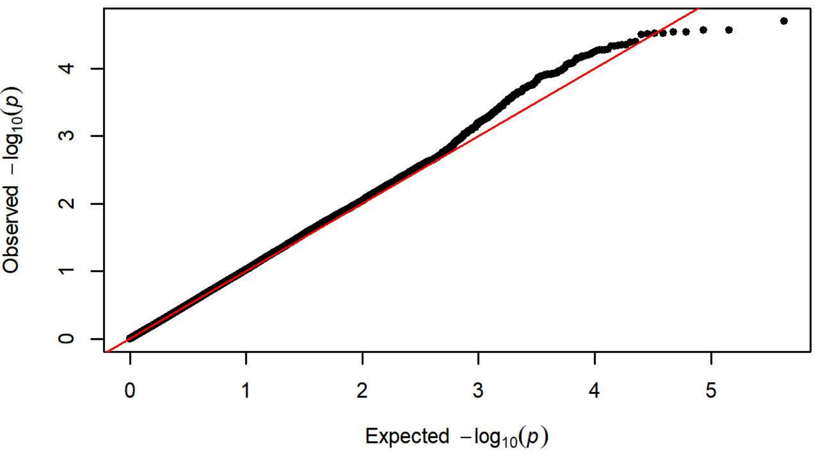


**Q-Q Plot Latency Score Day 2 - Harlan 206 (n=758)**

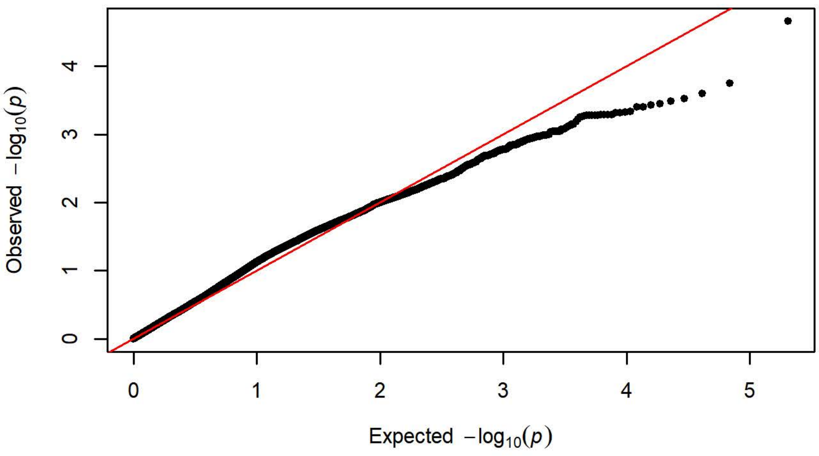


**Q-Q Plot Latency Score Day 2 - Charles River P09 (n=295)**

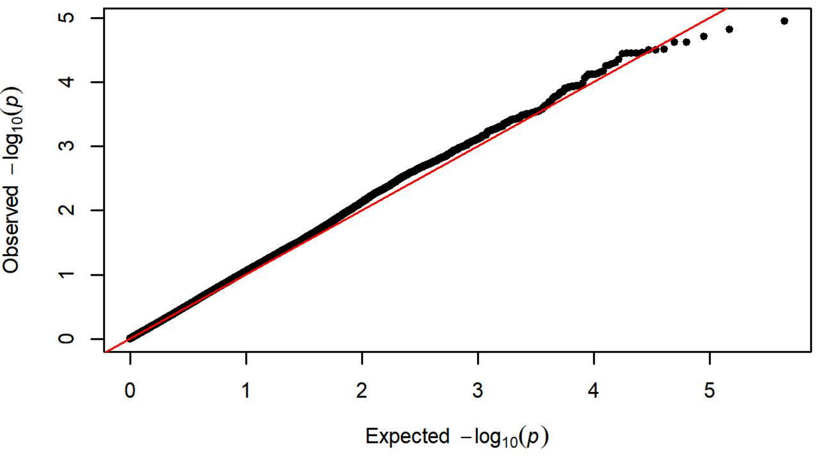


**Q-Q Plot Latency Score Day 2 - Harlan 217 (n=351)**

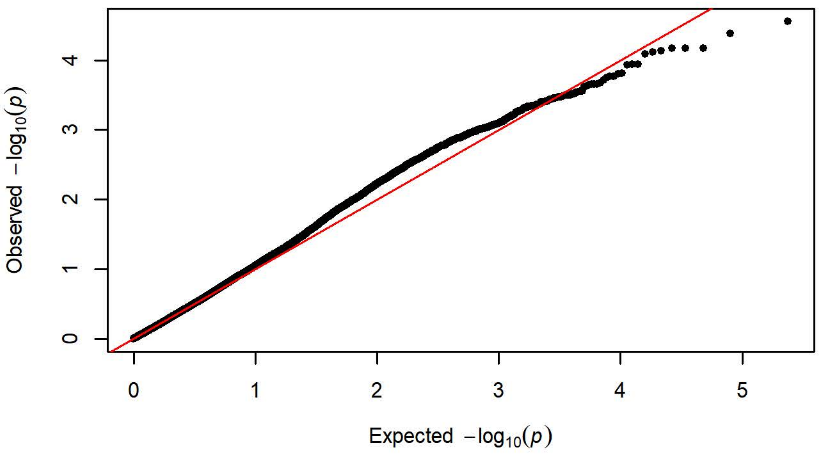


**Q-Q Plot Latency Score Day 2 - Charles River C72 (n=356)**

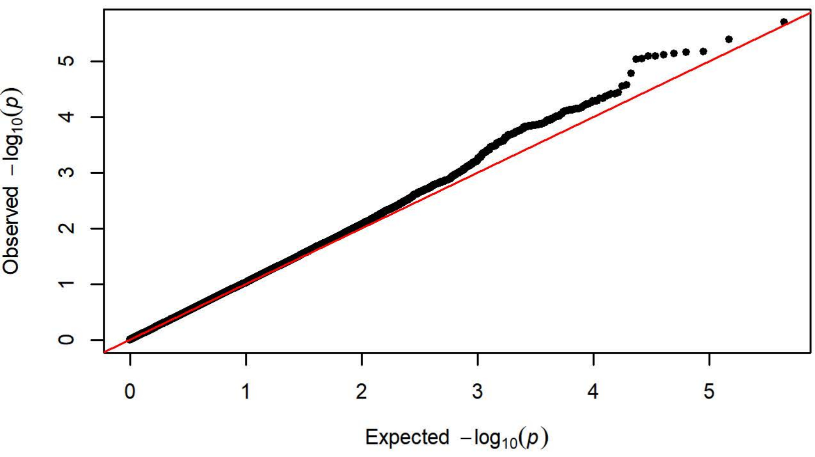

Q-Q Plot Latency Score Day 3 - Charles River R09-P3/7/10 (n=424)

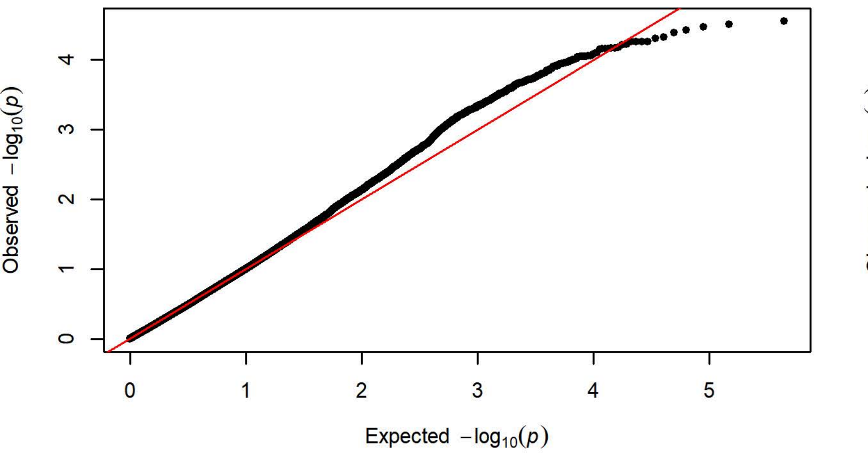


Q-Q Plot Latency Score Day 3 - Harlan 202A/C-208A (n=1096)

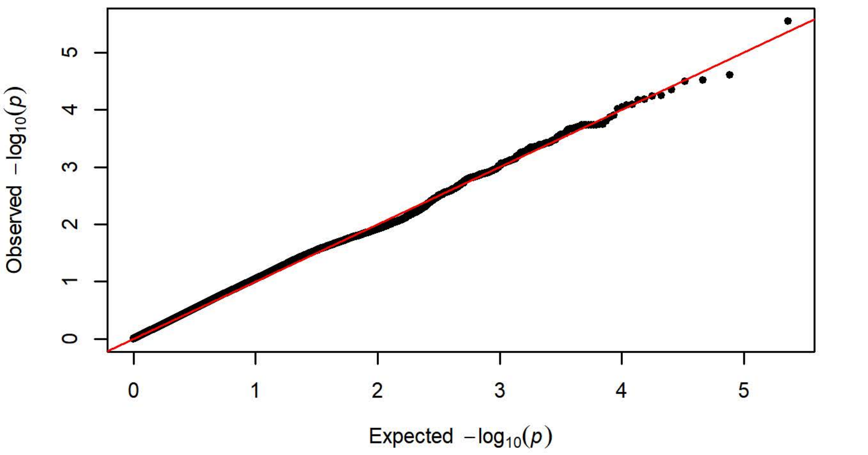


Q-Q Plot Latency Score Day 3 - Charles River R04 (n=650)

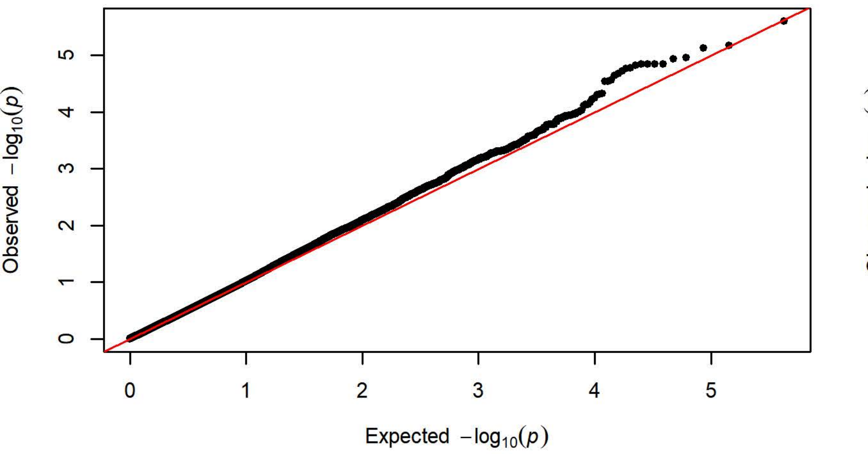


Q-Q Plot Latency Score Day 3 - Harlan 206 (n=758)

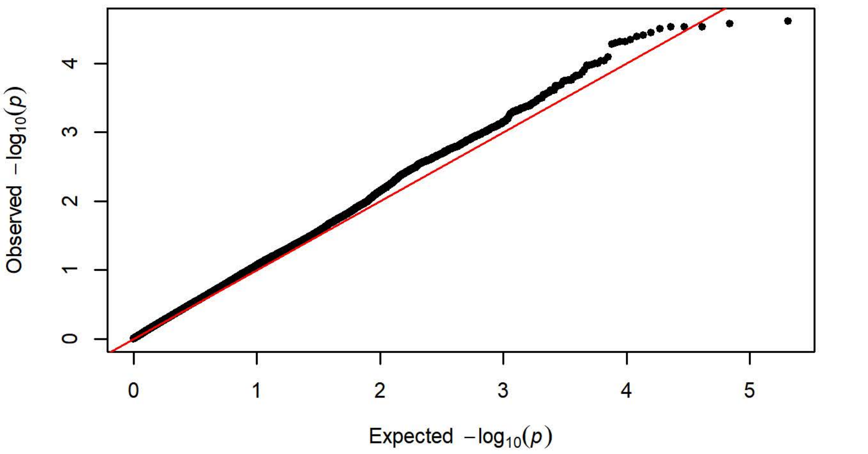


Q-Q Plot Latency Score Day 3 - Charles River P09 (n=295)

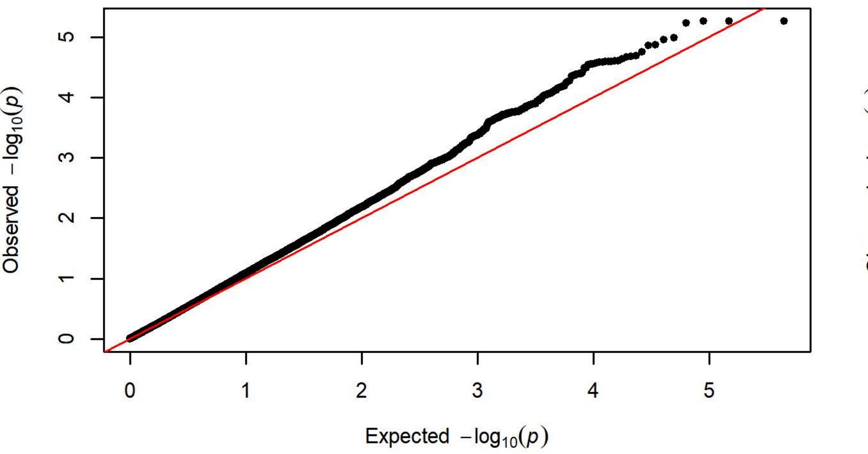


Q-Q Plot Latency Score Day 3 - Harlan 217 (n=351)

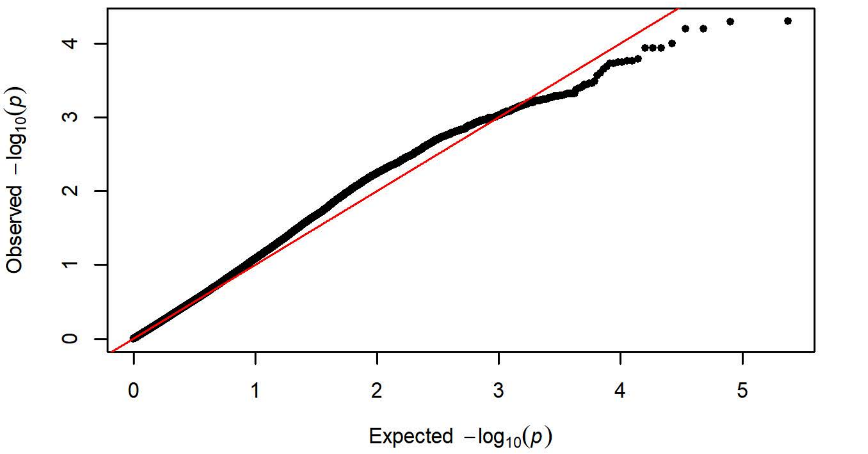


Q-Q Plot Latency Score Day 3 - Charles River C72 (n=358)

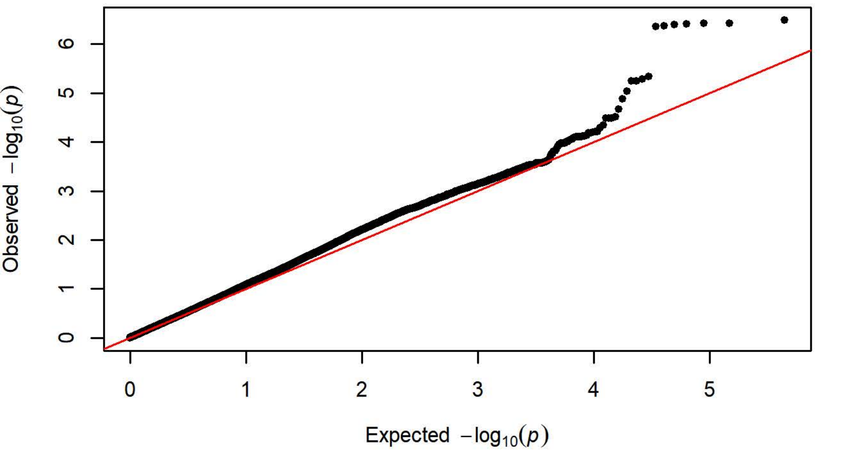

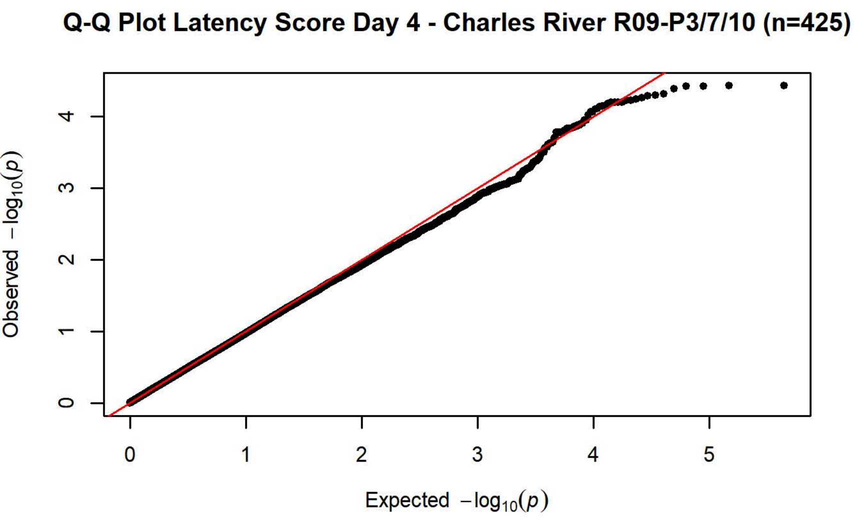
Q-Q Plot Latency Score Day 4 - Charles River R09-P3/7/10 (n=425)
Observed $-\log_{10}(p)$
Expected $-\log_{10}(p)$

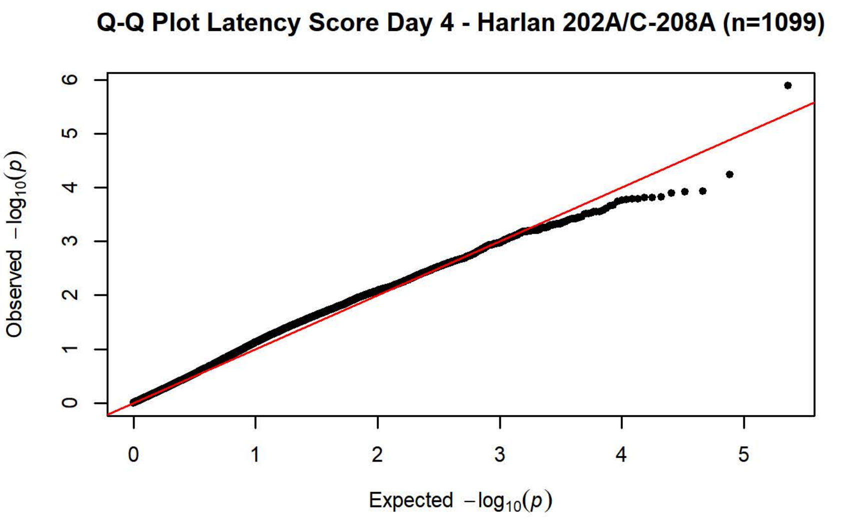
Q-Q Plot Latency Score Day 4 - Harlan 202A/C-208A (n=1099)
Observed $-\log_{10}(p)$
Expected $-\log_{10}(p)$

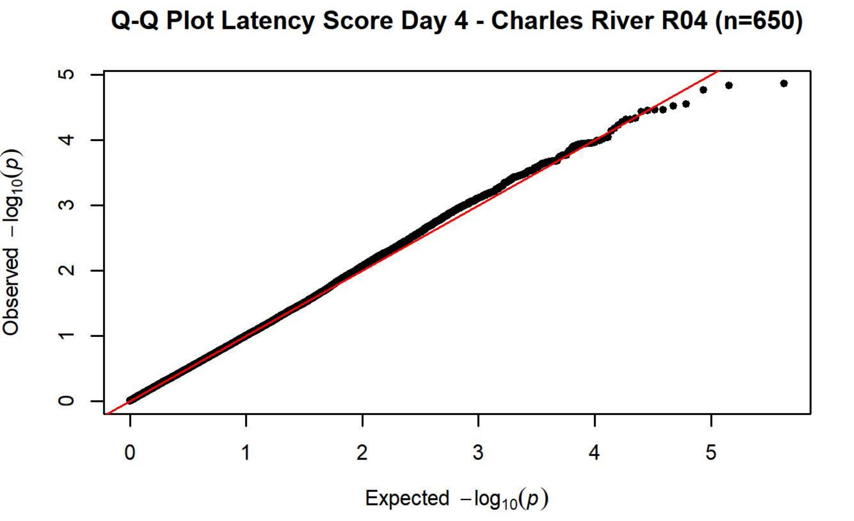
Q-Q Plot Latency Score Day 4 - Charles River R04 (n=650)
Observed $-\log_{10}(p)$
Expected $-\log_{10}(p)$

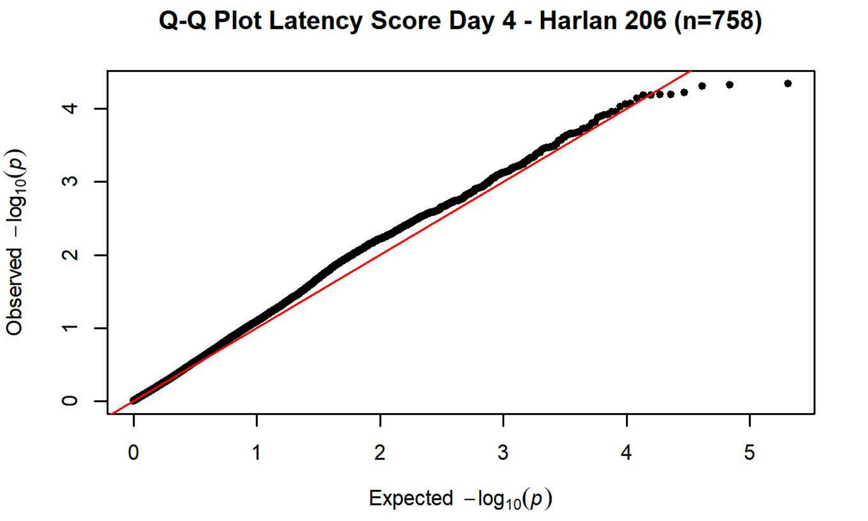
Q-Q Plot Latency Score Day 4 - Harlan 206 (n=758)
Observed $-\log_{10}(p)$
Expected $-\log_{10}(p)$

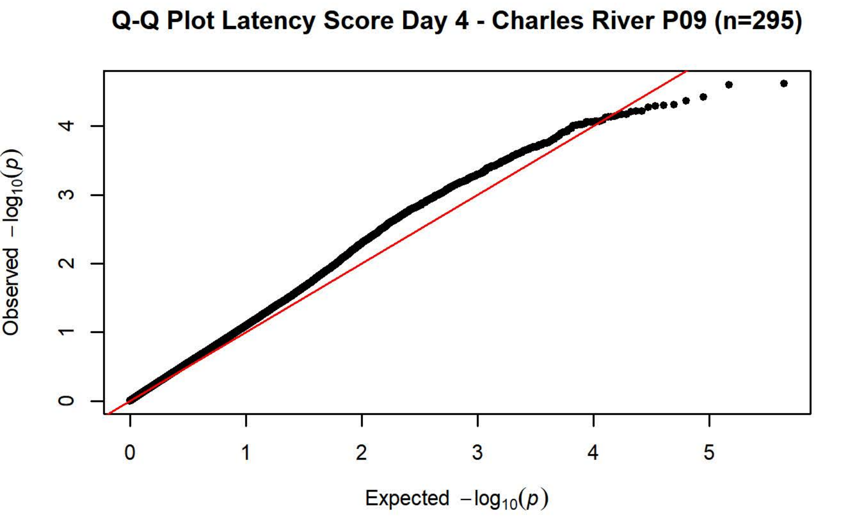
Q-Q Plot Latency Score Day 4 - Charles River P09 (n=295)
Observed $-\log_{10}(p)$
Expected $-\log_{10}(p)$

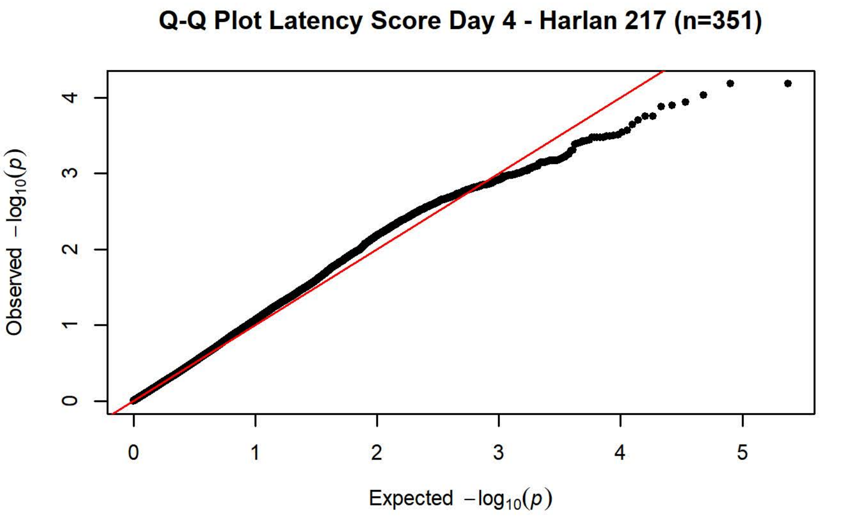
Q-Q Plot Latency Score Day 4 - Harlan 217 (n=351)
Observed $-\log_{10}(p)$
Expected $-\log_{10}(p)$

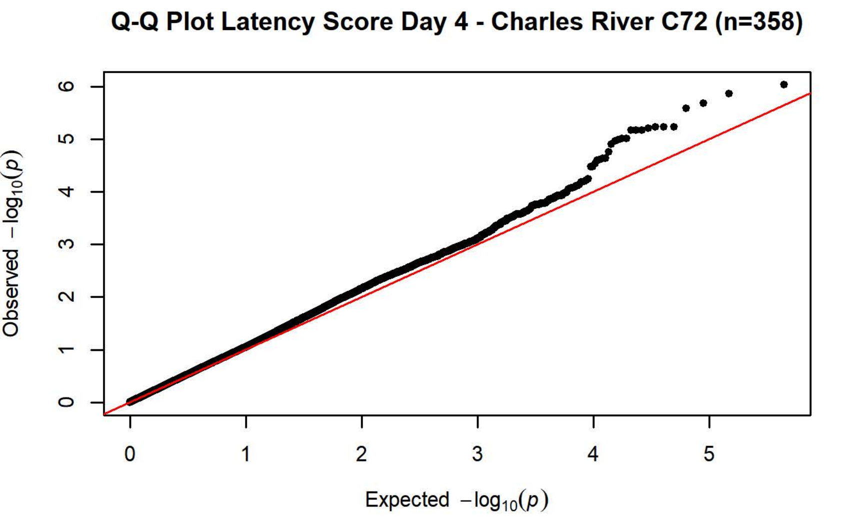
Q-Q Plot Latency Score Day 4 - Charles River C72 (n=358)
Observed $-\log_{10}(p)$
Expected $-\log_{10}(p)$

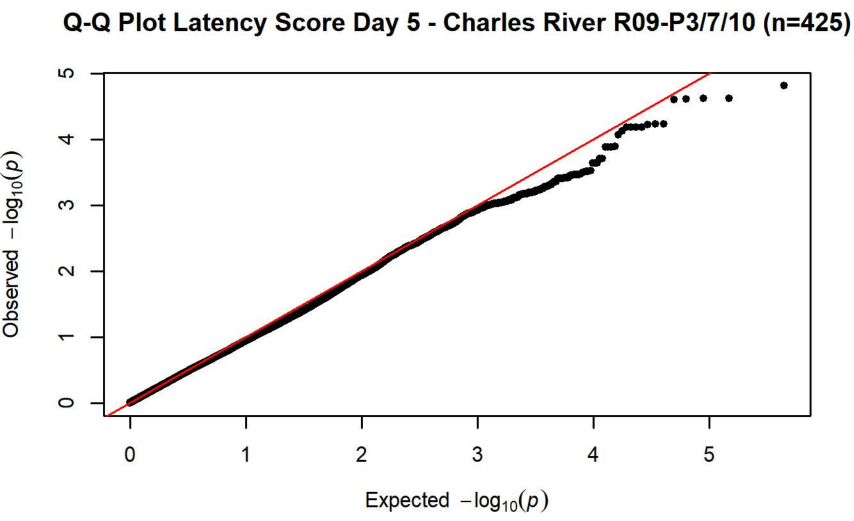
Q-Q Plot Latency Score Day 5 - Charles River R09-P3/7/10 (n=425)
Observed $-\log_{10}(p)$
Expected $-\log_{10}(p)$

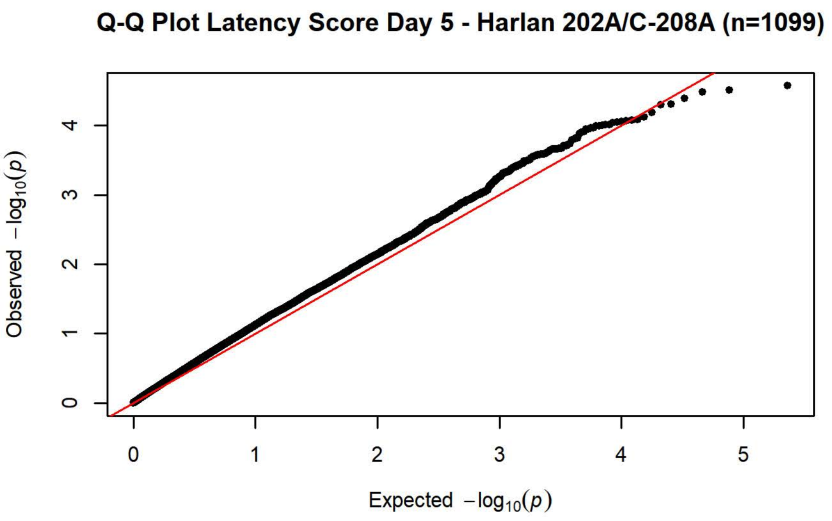
Q-Q Plot Latency Score Day 5 - Harlan 202A/C-208A (n=1099)
Observed $-\log_{10}(p)$
Expected $-\log_{10}(p)$

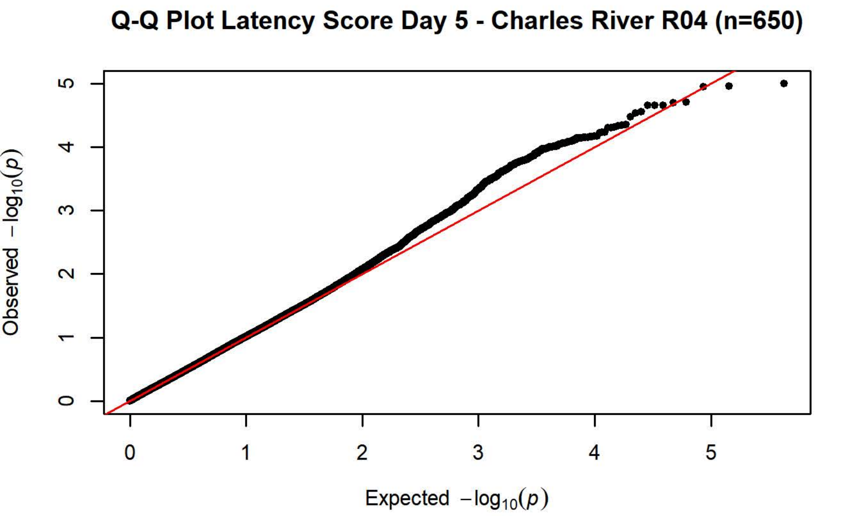
Q-Q Plot Latency Score Day 5 - Charles River R04 (n=650)
Observed $-\log_{10}(p)$
Expected $-\log_{10}(p)$

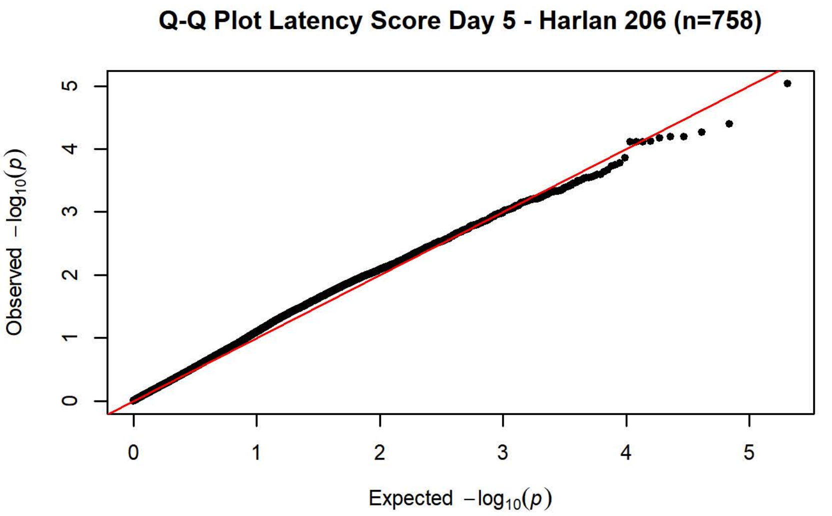
Q-Q Plot Latency Score Day 5 - Harlan 206 (n=758)
Observed $-\log_{10}(p)$
Expected $-\log_{10}(p)$

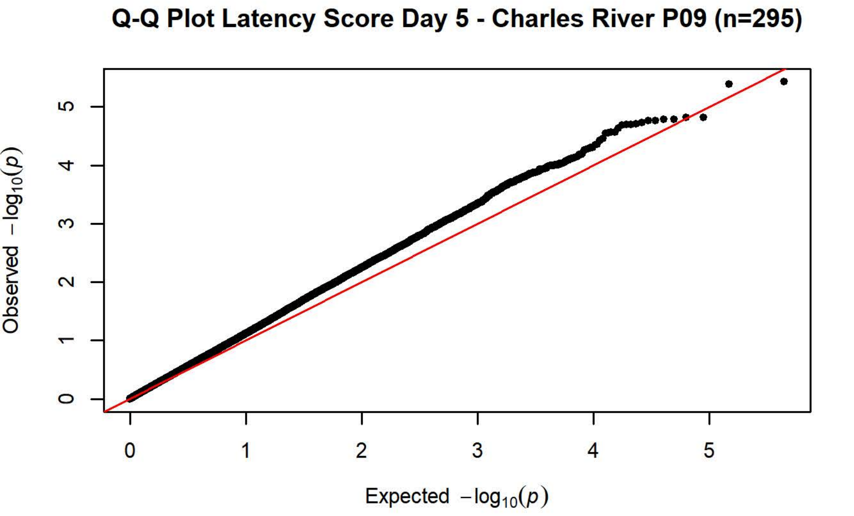
Q-Q Plot Latency Score Day 5 - Charles River P09 (n=295)
Observed $-\log_{10}(p)$
Expected $-\log_{10}(p)$

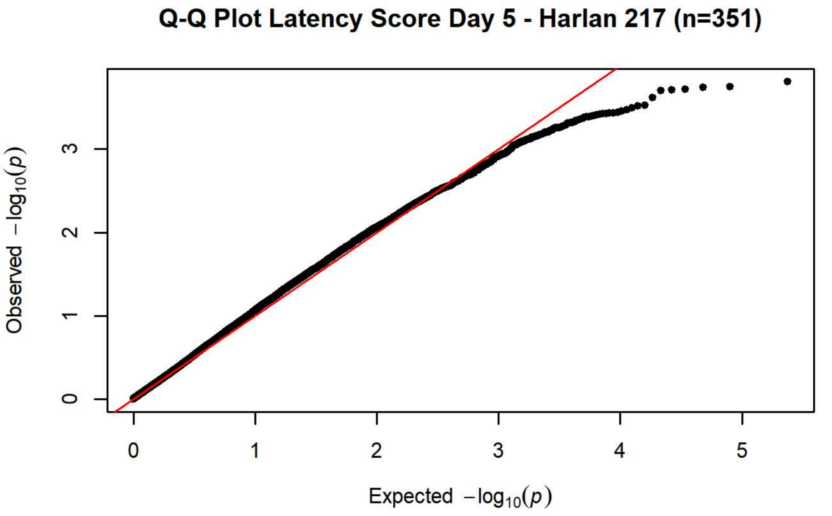
Q-Q Plot Latency Score Day 5 - Harlan 217 (n=351)
Observed $-\log_{10}(p)$
Expected $-\log_{10}(p)$

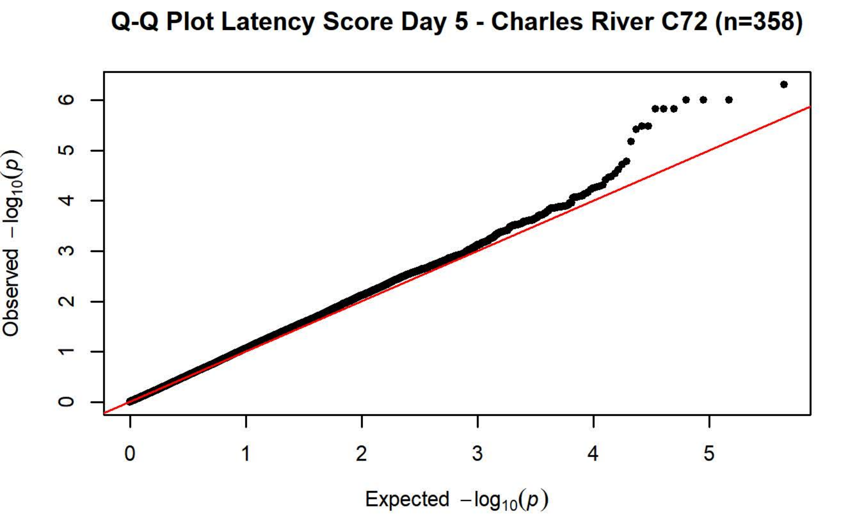
Q-Q Plot Latency Score Day 5 - Charles River C72 (n=358)
Observed $-\log_{10}(p)$
Expected $-\log_{10}(p)$

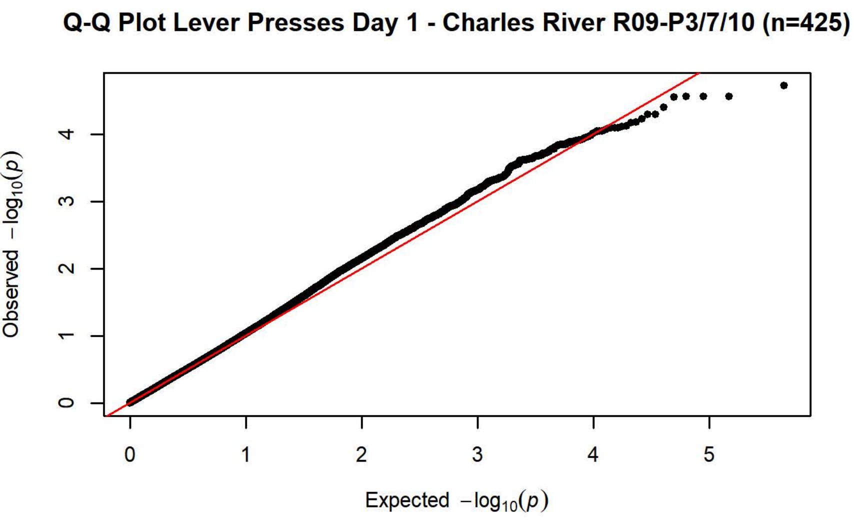
Q-Q Plot Lever Presses Day 1 - Charles River R09-P3/7/10 (n=425)
Observed $-\log_{10}(p)$
Expected $-\log_{10}(p)$

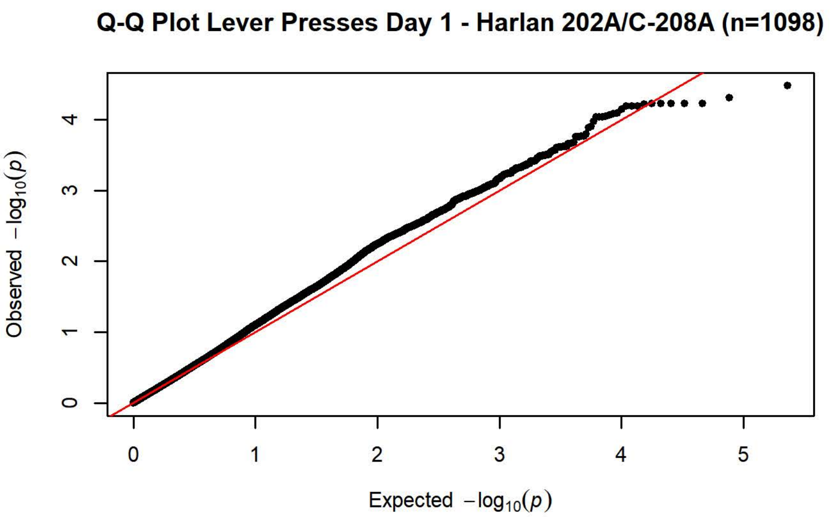
Q-Q Plot Lever Presses Day 1 - Harlan 202A/C-208A (n=1098)
Observed $-\log_{10}(p)$
Expected $-\log_{10}(p)$

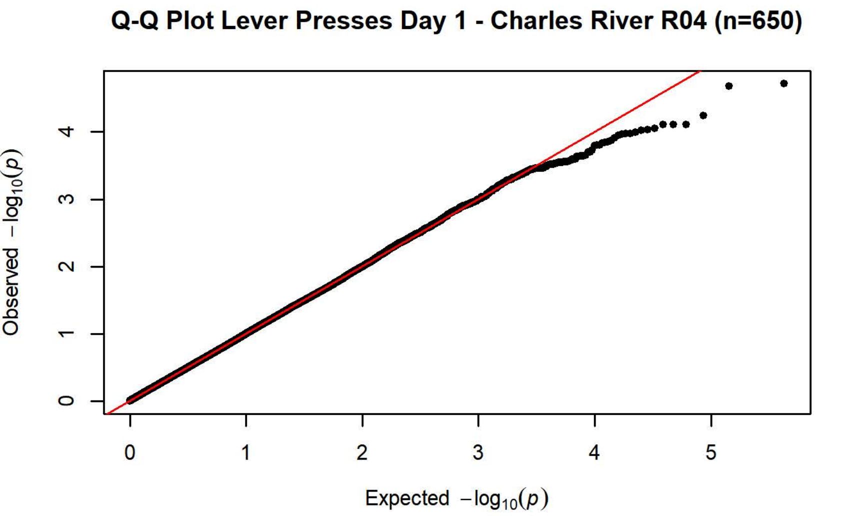
Q-Q Plot Lever Presses Day 1 - Charles River R04 (n=650)
Observed $-\log_{10}(p)$
Expected $-\log_{10}(p)$

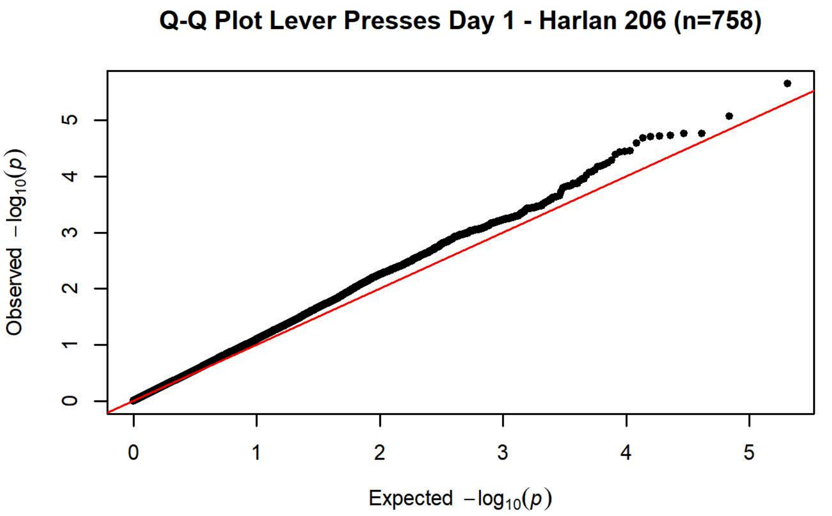
Q-Q Plot Lever Presses Day 1 - Harlan 206 (n=758)
Observed $-\log_{10}(p)$
Expected $-\log_{10}(p)$

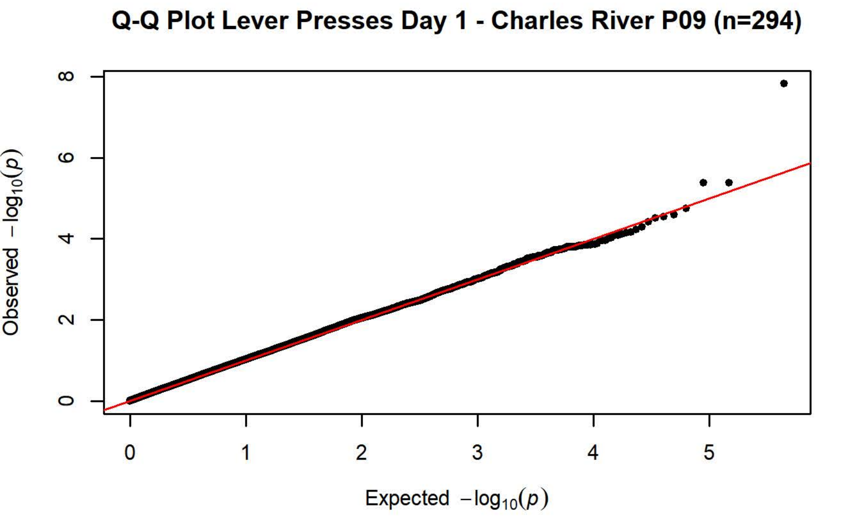
Q-Q Plot Lever Presses Day 1 - Charles River P09 (n=294)
Observed $-\log_{10}(p)$
Expected $-\log_{10}(p)$

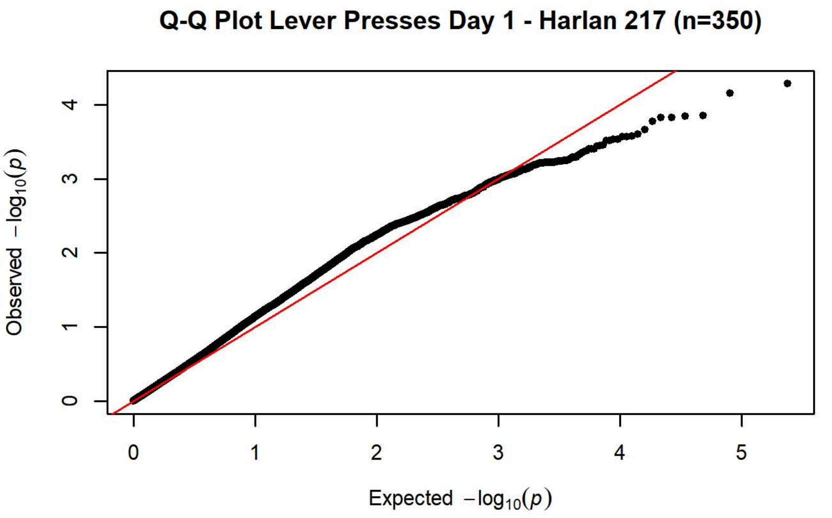
Q-Q Plot Lever Presses Day 1 - Harlan 217 (n=350)
Observed $-\log_{10}(p)$
Expected $-\log_{10}(p)$

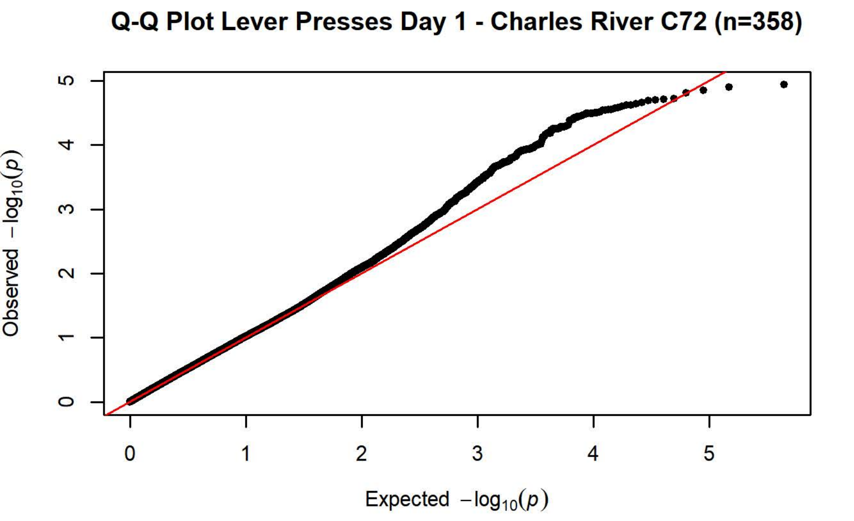
Q-Q Plot Lever Presses Day 1 - Charles River C72 (n=358)
Observed $-\log_{10}(p)$
Expected $-\log_{10}(p)$

**Q-Q Plot Lever Presses Day 2 - Charles River R09-P3/7/10 (n=425)**

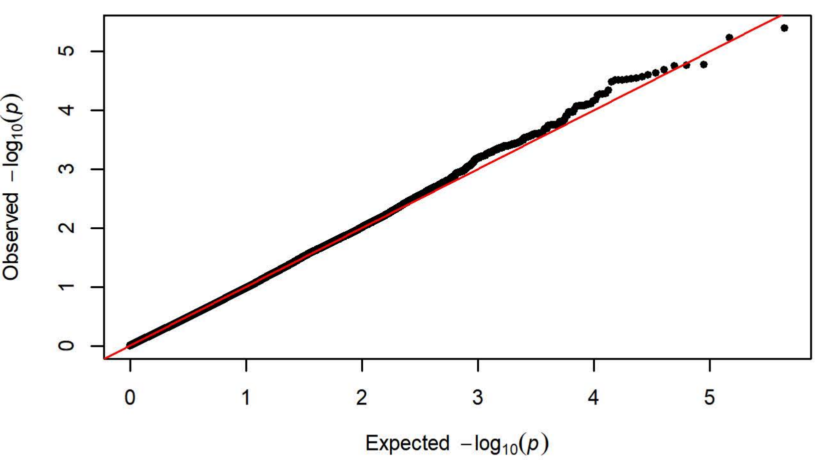


**Q-Q Plot Lever Presses Day 2 - Harlan 202A/C-208A (n=1098)**

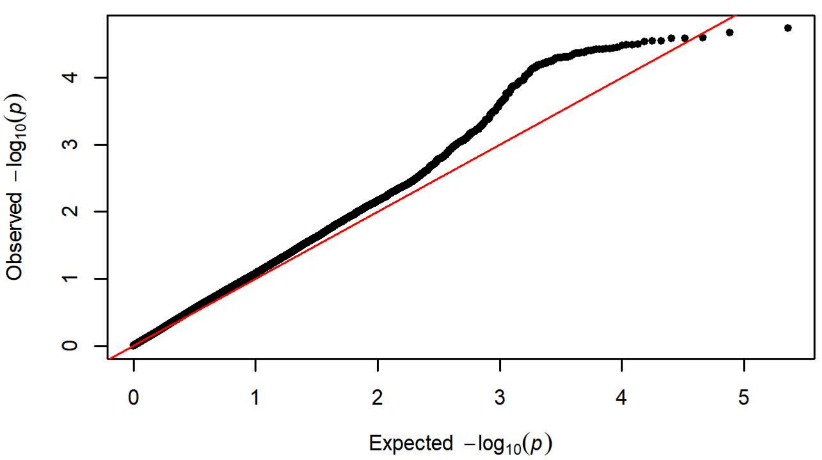


**Q-Q Plot Lever Presses Day 2 - Charles River R04 (n=650)**

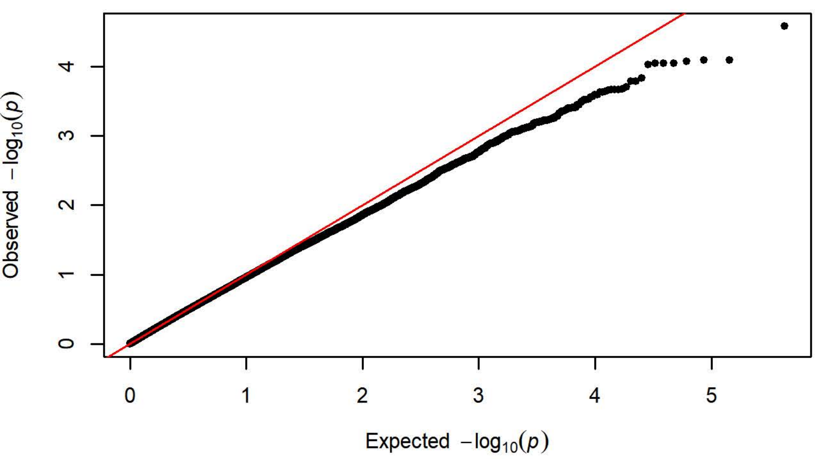


**Q-Q Plot Lever Presses Day 2 - Harlan 206 (n=757)**

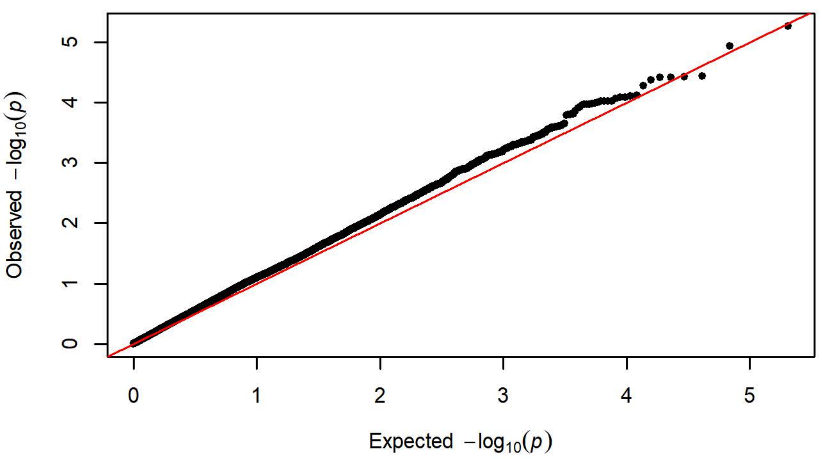


**Q-Q Plot Lever Presses Day 2 - Charles River P09 (n=295)**

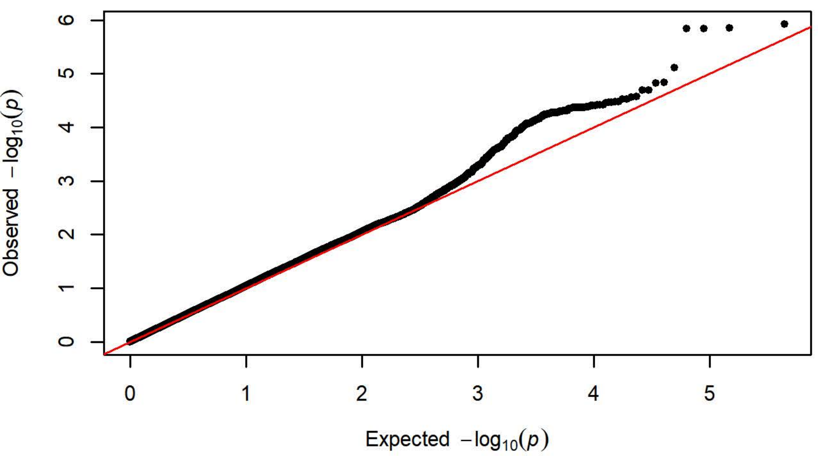


**Q-Q Plot Lever Presses Day 2 - Harlan 217 (n=350)**

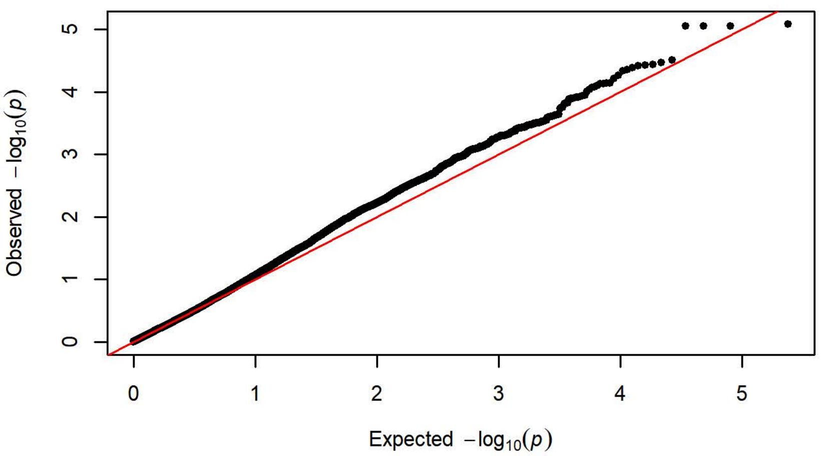


**Q-Q Plot Lever Presses Day 2 - Charles River C72 (n=356)**

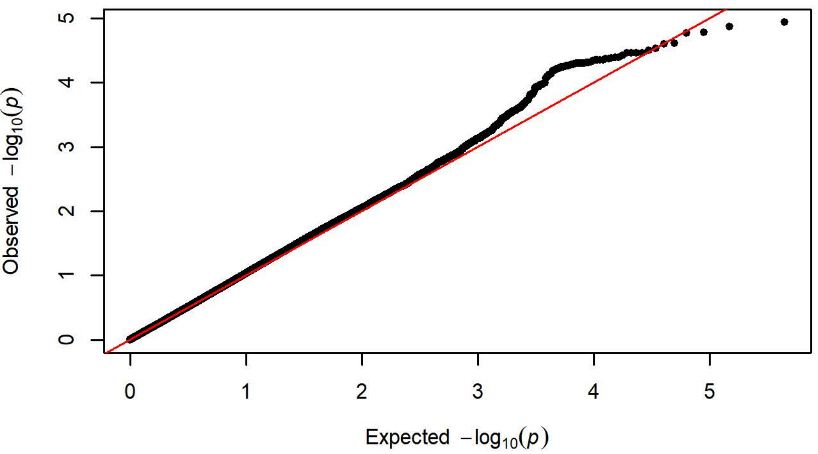

**Q-Q Plot Lever Presses Day 3 - Charles River R09-P3/7/10 (n=424)**

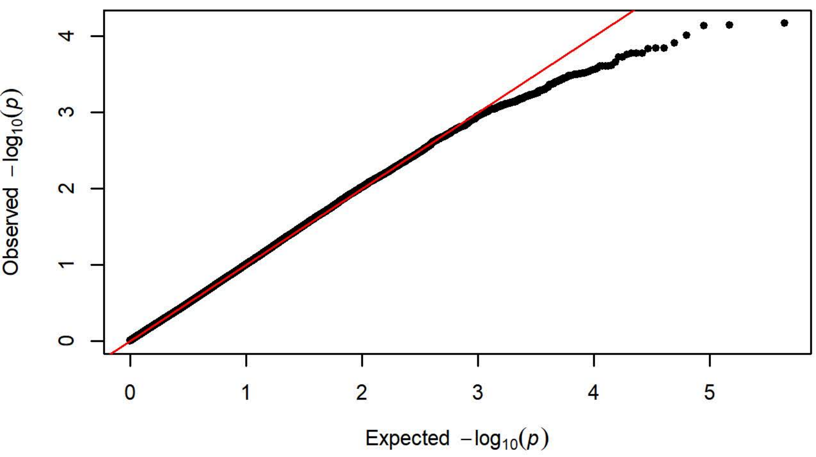


**Q-Q Plot Lever Presses Day 3 - Harlan 202A/C-208A (n=1095)**

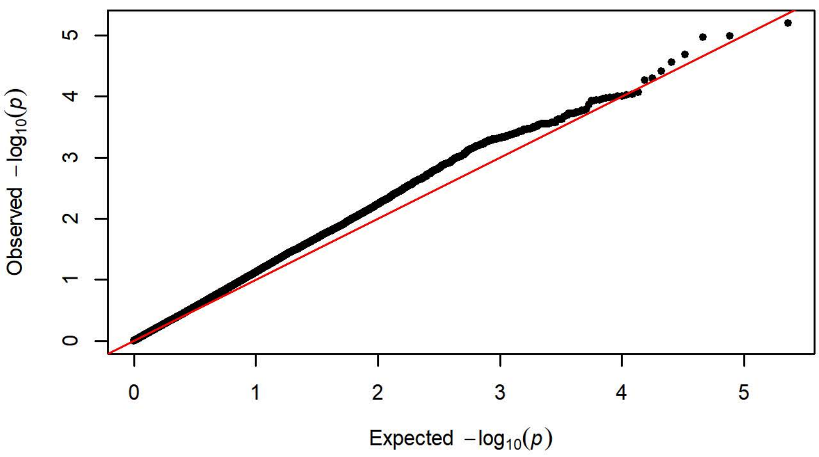


**Q-Q Plot Lever Presses Day 3 - Charles River R04 (n=650)**

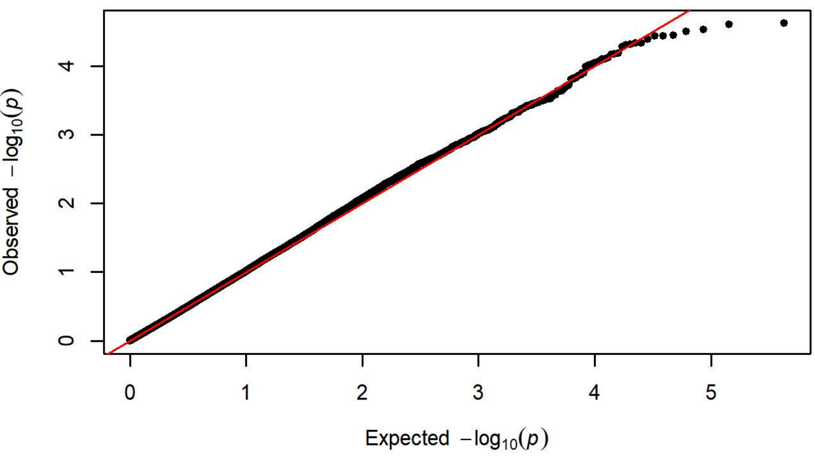


**Q-Q Plot Lever Presses Day 3 - Harlan 206 (n=758)**

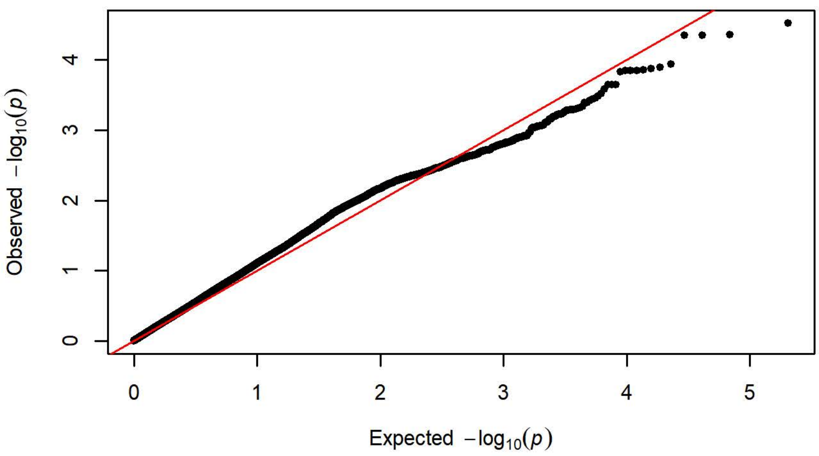


**Q-Q Plot Lever Presses Day 3 - Charles River P09 (n=295)**

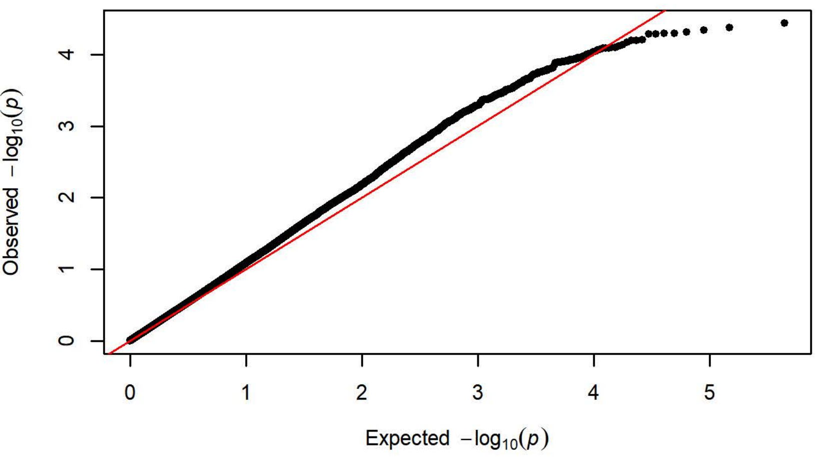


**Q-Q Plot Lever Presses Day 3 - Harlan 217 (n=351)**

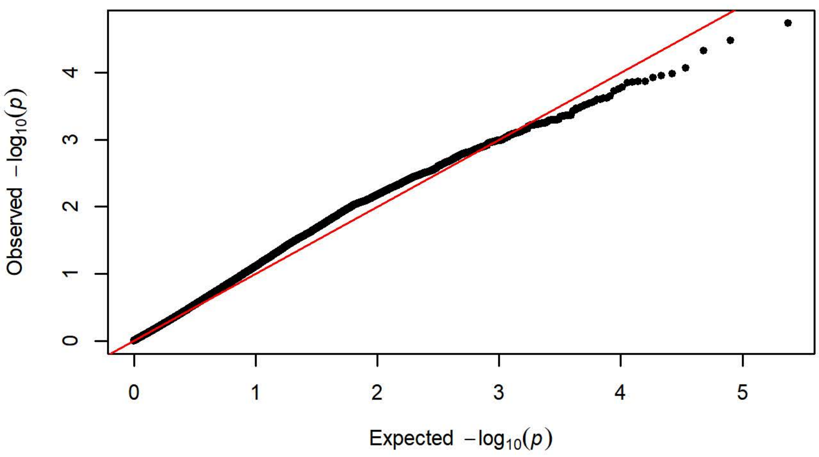


**Q-Q Plot Lever Presses Day 3 - Charles River C72 (n=358)**

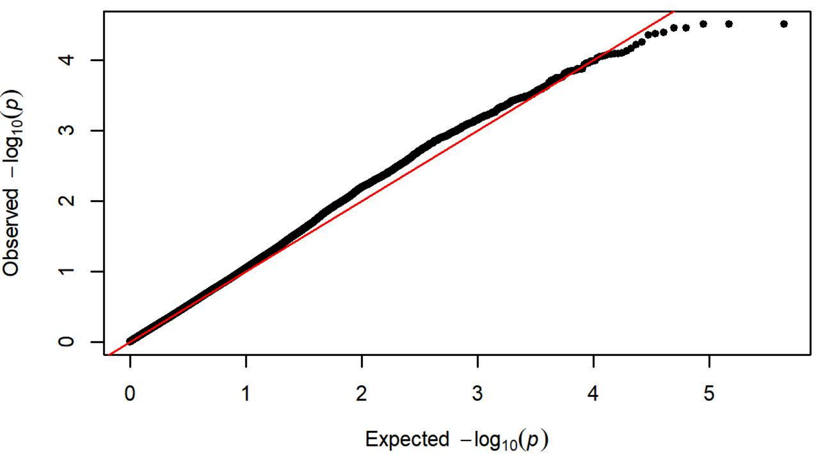

**Q-Q Plot Lever Presses Day 4 - Charles River R09-P3/7/10 (n=425)**

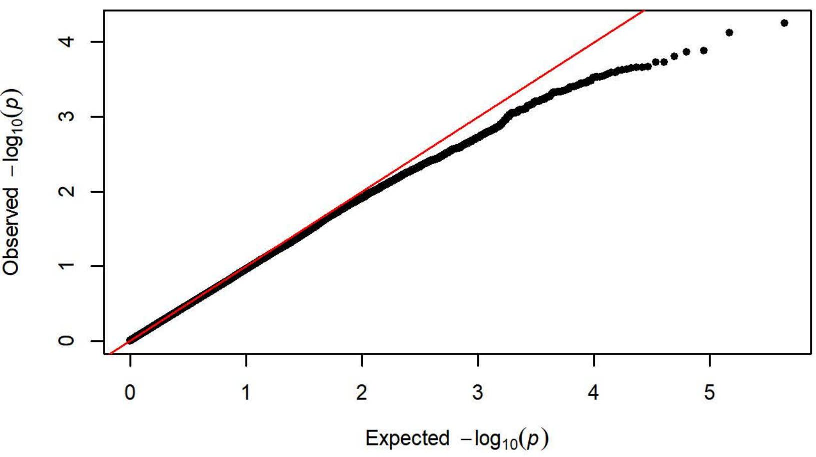


**Q-Q Plot Lever Presses Day 4 - Harlan 202A/C-208A (n=1099)**

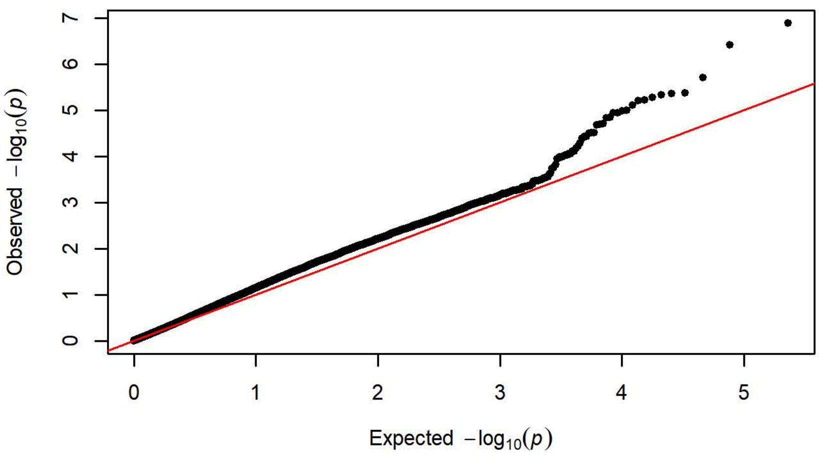


**Q-Q Plot Lever Presses Day 4 - Charles River R04 (n=650)**

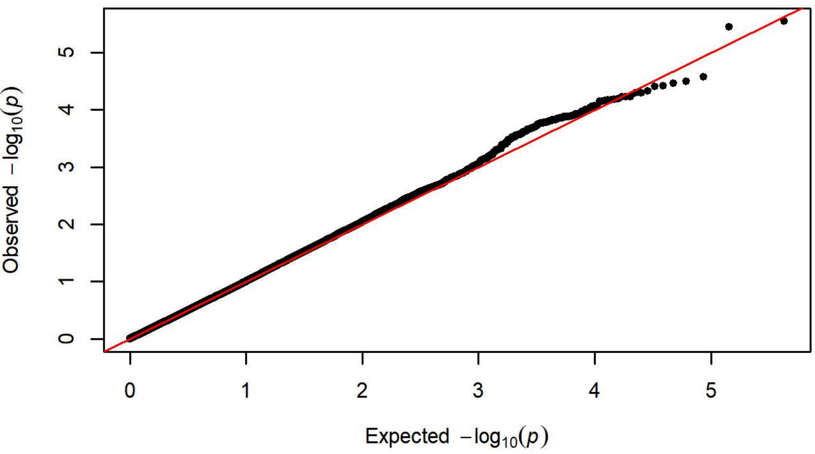


**Q-Q Plot Lever Presses Day 4 - Harlan 206 (n=758)**

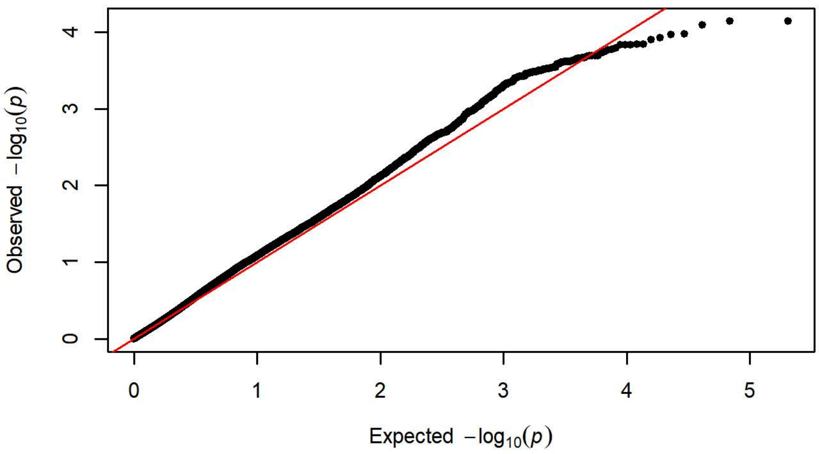


**Q-Q Plot Lever Presses Day 4 - Charles River P09 (n=295)**

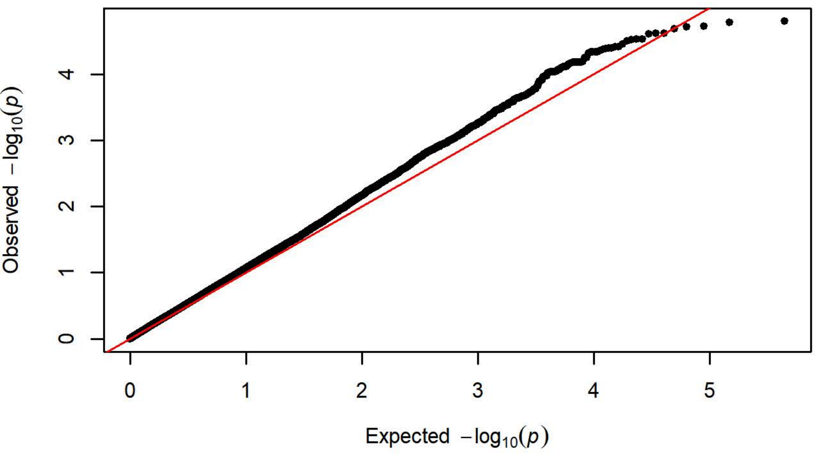


**Q-Q Plot Lever Presses Day 4 - Harlan 217 (n=351)**

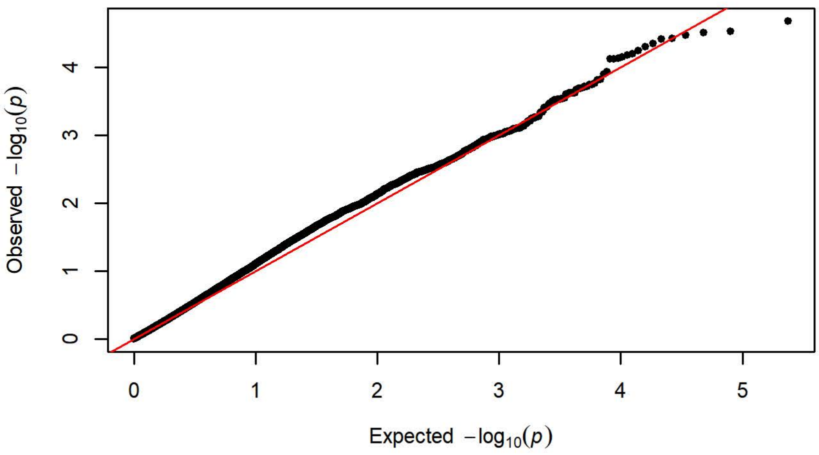


**Q-Q Plot Lever Presses Day 4 - Charles River C72 (n=358)**

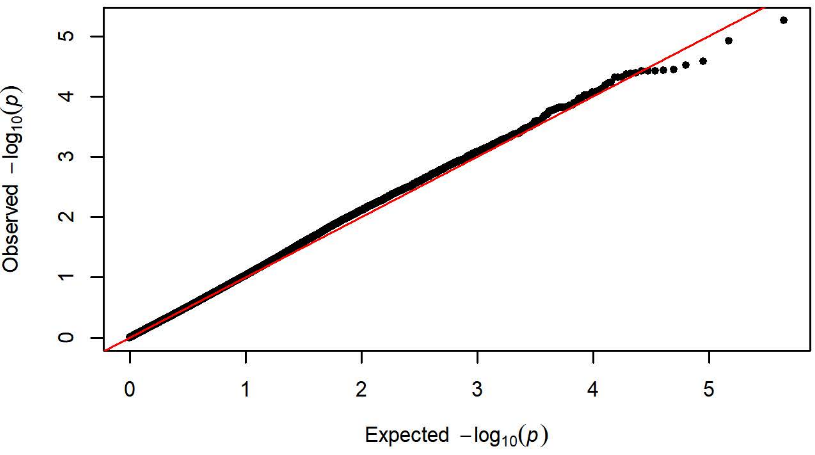

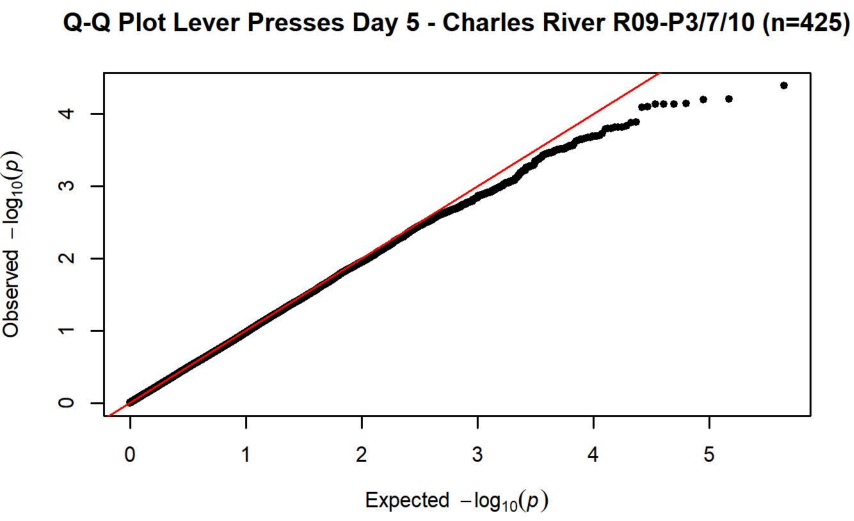
Q-Q Plot Lever Presses Day 5 - Charles River R09-P3/7/10 (n=425)
Observed −log10(p)
Expected −log10(p)

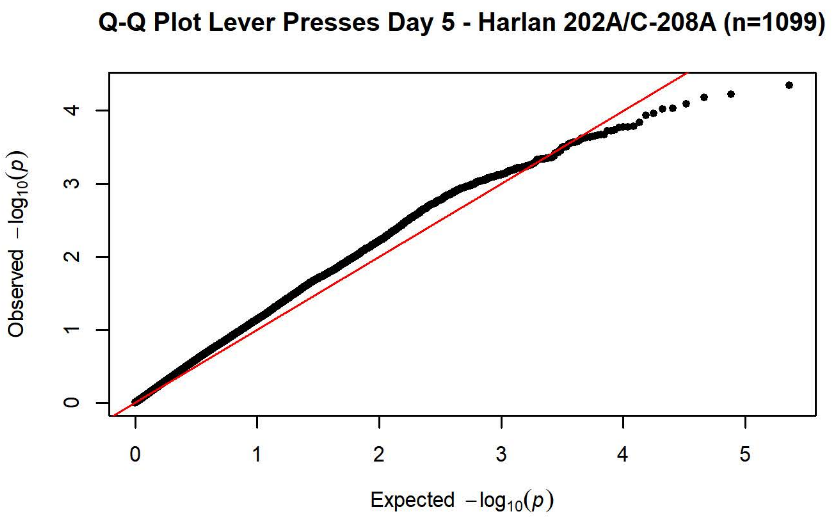
Q-Q Plot Lever Presses Day 5 - Harlan 202A/C-208A (n=1099)
Observed −log10(p)
Expected −log10(p)

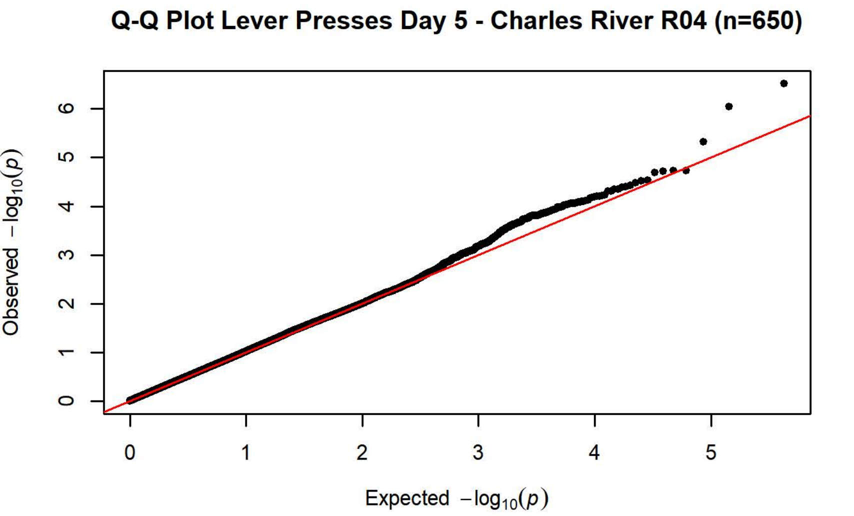
Q-Q Plot Lever Presses Day 5 - Charles River R04 (n=650)
Observed −log10(p)
Expected −log10(p)

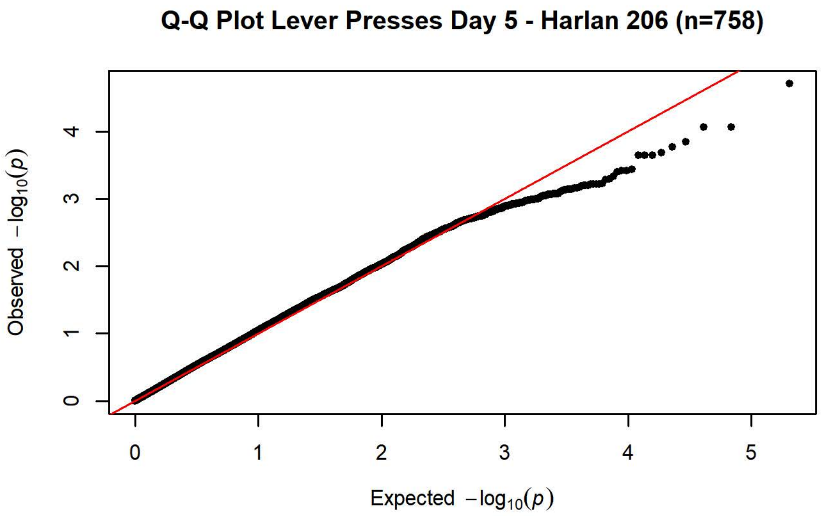
Q-Q Plot Lever Presses Day 5 - Harlan 206 (n=758)
Observed −log10(p)
Expected −log10(p)

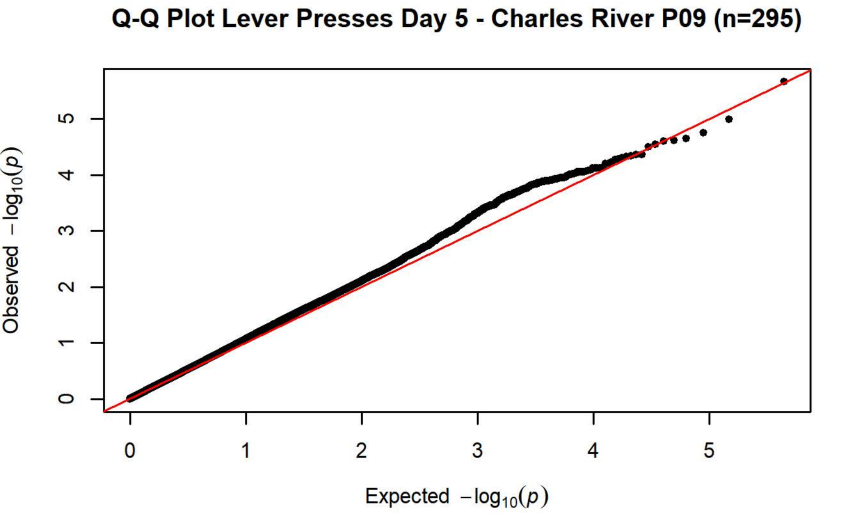
Q-Q Plot Lever Presses Day 5 - Charles River P09 (n=295)
Observed −log10(p)
Expected −log10(p)

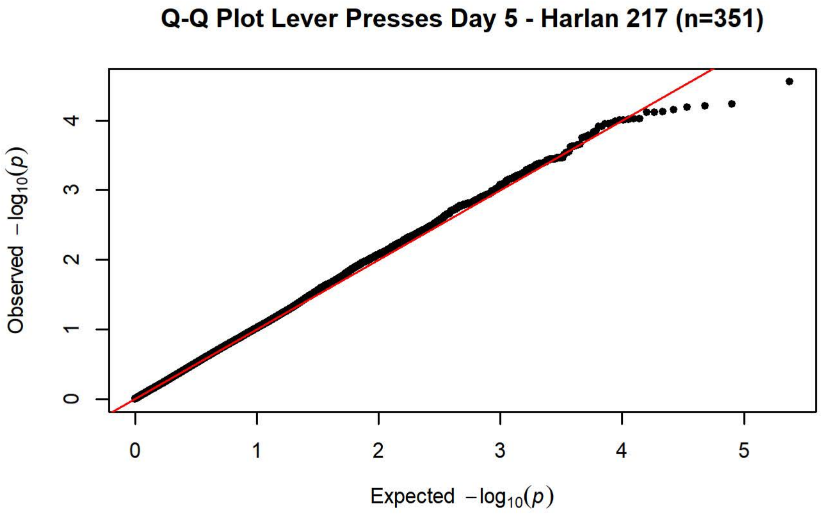
Q-Q Plot Lever Presses Day 5 - Harlan 217 (n=351)
Observed −log10(p)
Expected −log10(p)

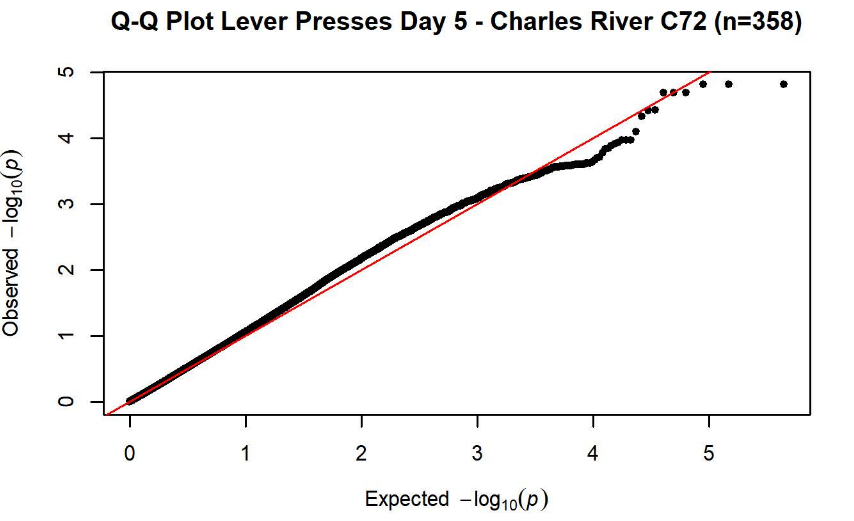
Q-Q Plot Lever Presses Day 5 - Charles River C72 (n=358)
Observed −log10(p)
Expected −log10(p)

**Q-Q Plot Magazaine Entries Day 1 - Charles River R09-P3/7/10 (n=425**

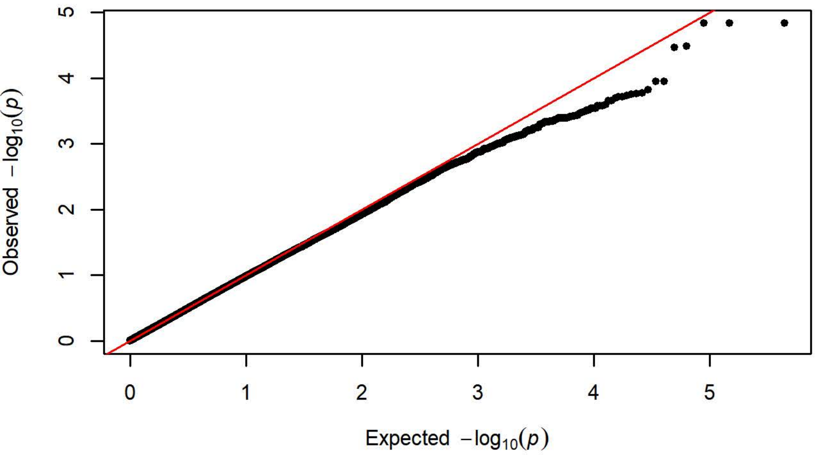


**Q-Q Plot Magazaine Entries Day 1 - Harlan 202A/C-208A (n=1099)**

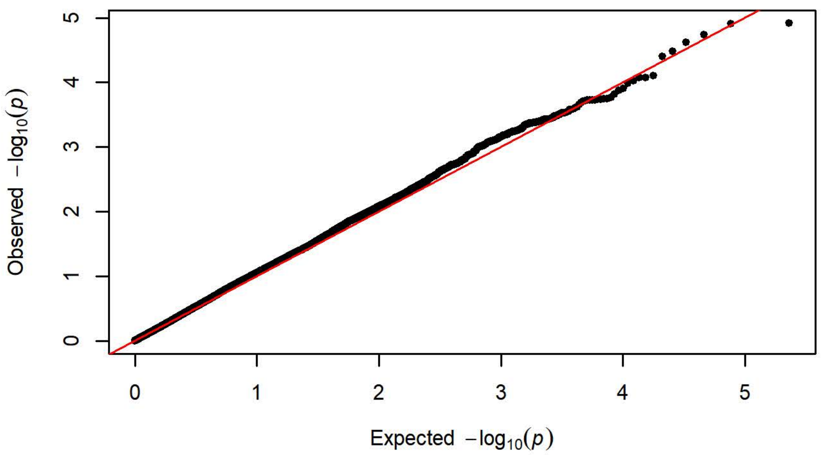


**Q-Q Plot Magazaine Entries Day 1 - Charles River R04 (n=650)**

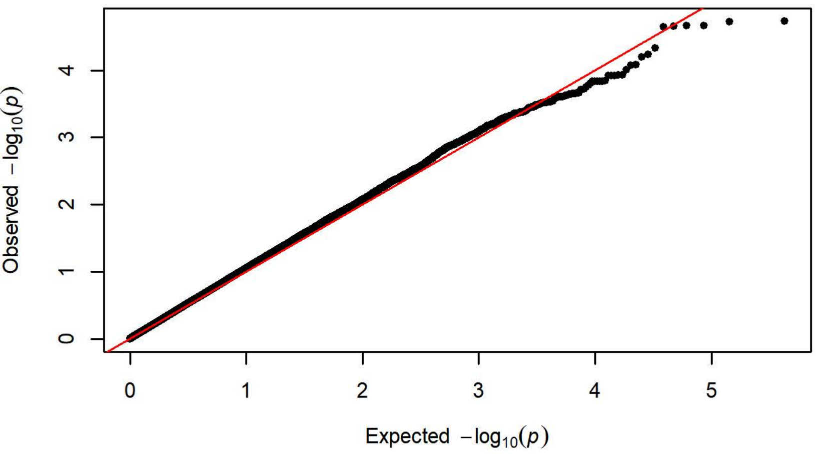


**Q-Q Plot Magazaine Entries Day 1 - Harlan 206 (n=757)**

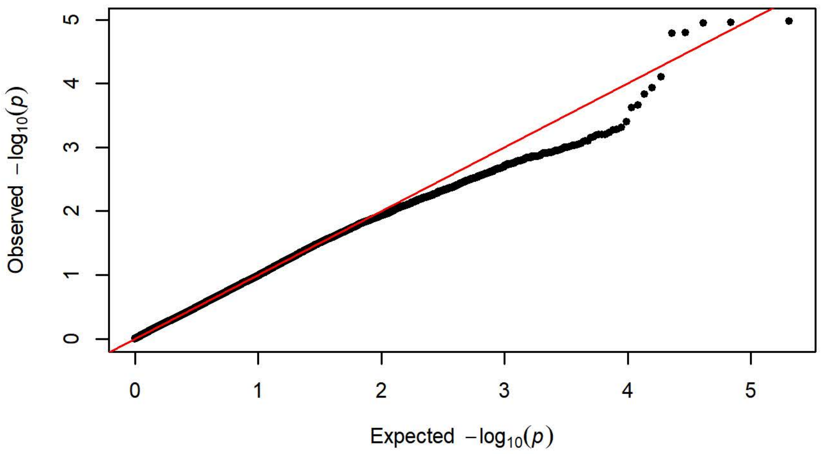


**Q-Q Plot Magazaine Entries Day 1 - Charles River P09 (n=295)**

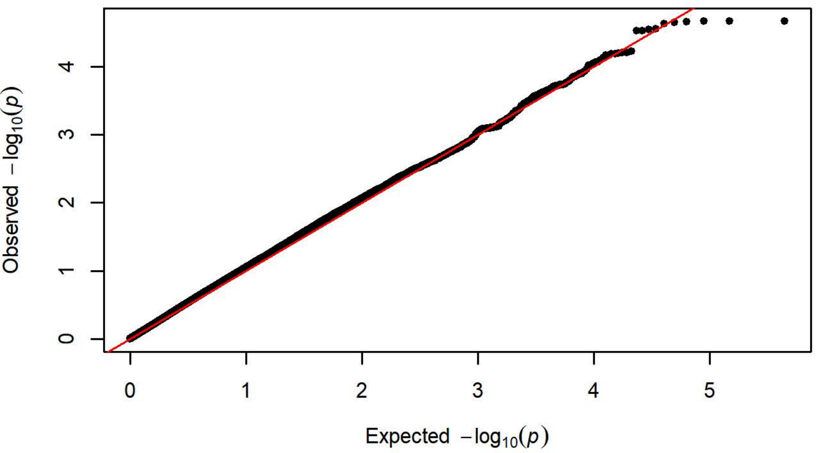


**Q-Q Plot Magazaine Entries Day 1 - Harlan 217 (n=349)**

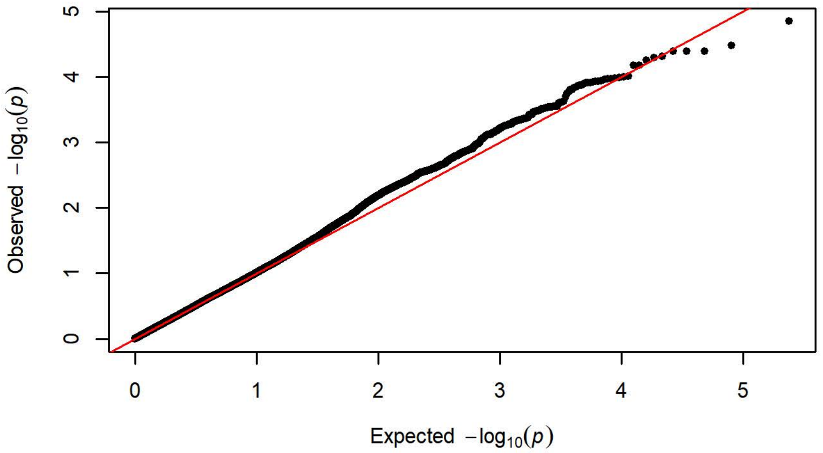


**Q-Q Plot Magazaine Entries Day 1 - Charles River C72 (n=358)**

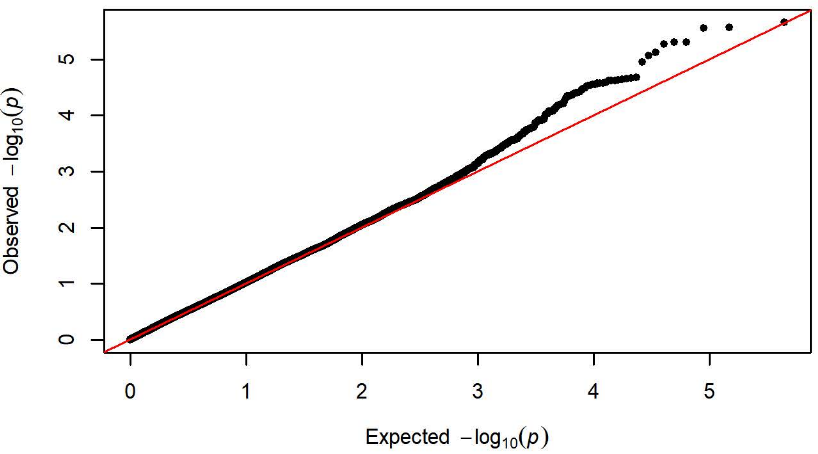

Q-Q Plot Magazaine Entries Day 2 - Charles River R09-P3/7/10 (n=425

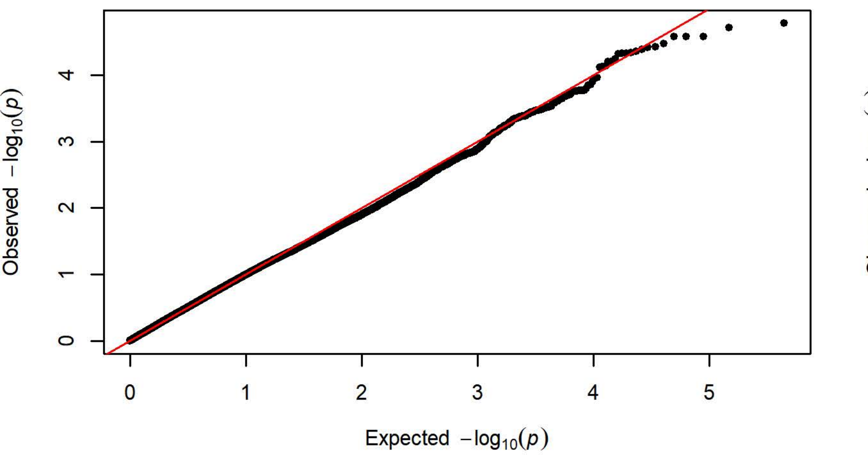


Q-Q Plot Magazaine Entries Day 2 - Harlan 202A/C-208A (n=1099)

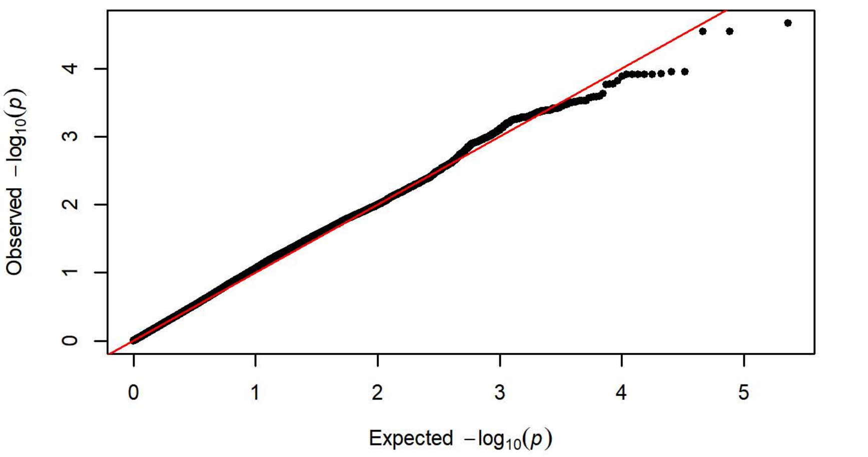


Q-Q Plot Magazaine Entries Day 2 - Charles River R04 (n=650)

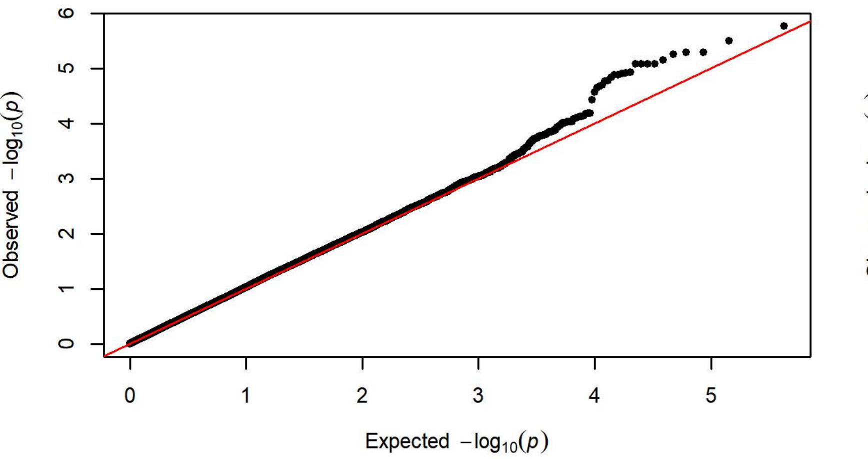


Q-Q Plot Magazaine Entries Day 2 - Harlan 206 (n=758)

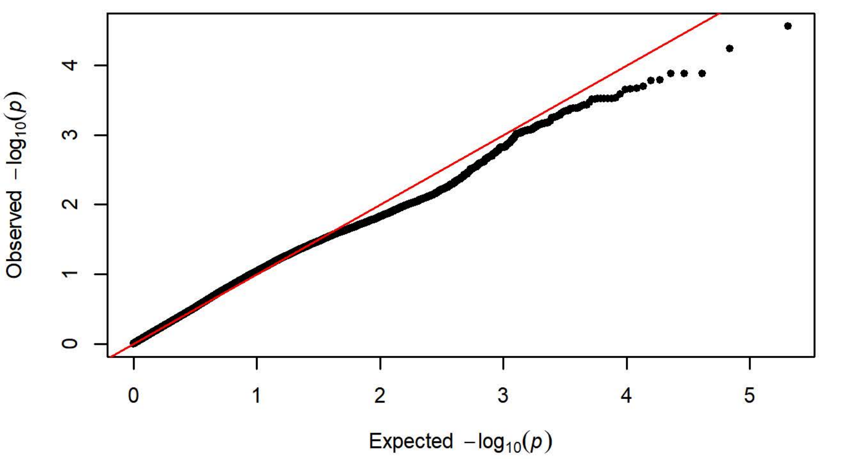


Q-Q Plot Magazaine Entries Day 2 - Charles River P09 (n=295)

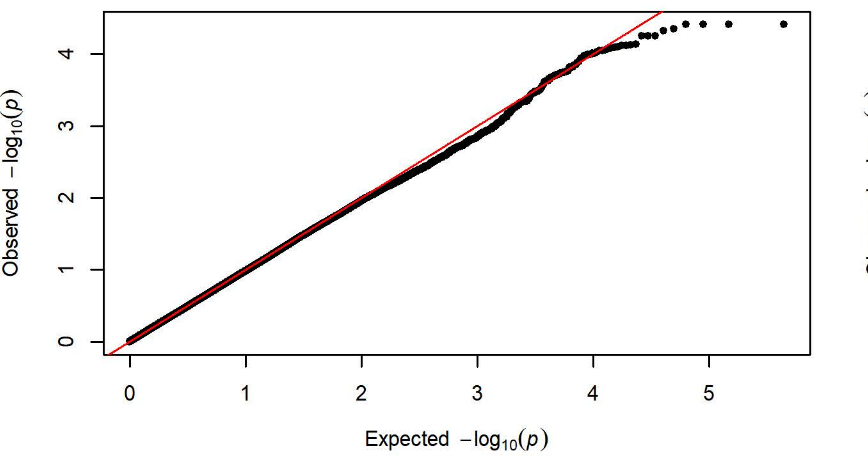


Q-Q Plot Magazaine Entries Day 2 - Harlan 217 (n=350)

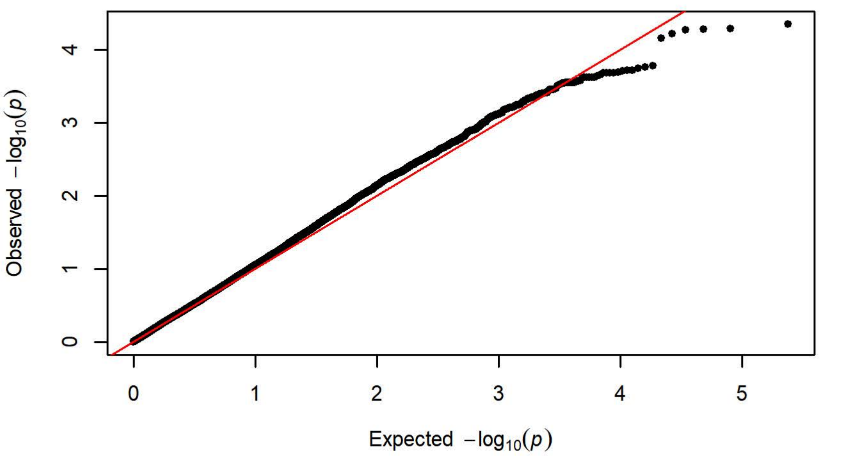


Q-Q Plot Magazaine Entries Day 2 - Charles River C72 (n=356)

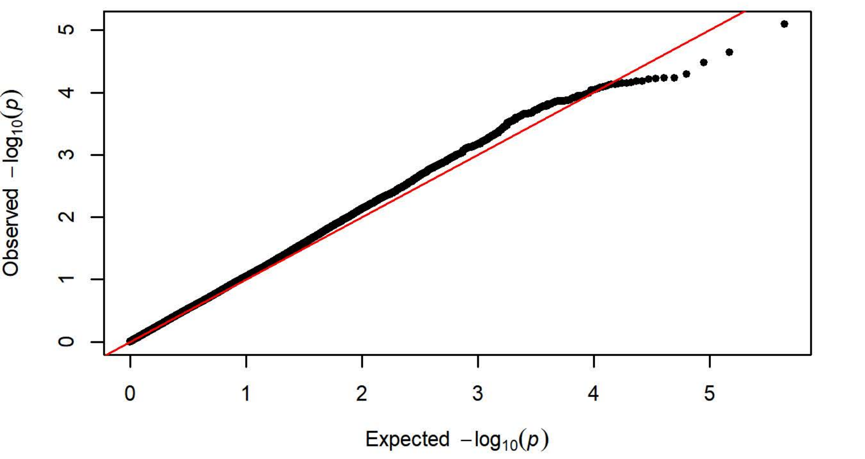

**Q-Q Plot Magazaine Entries Day 3 - Charles River R09-P3/7/10 (n=424**

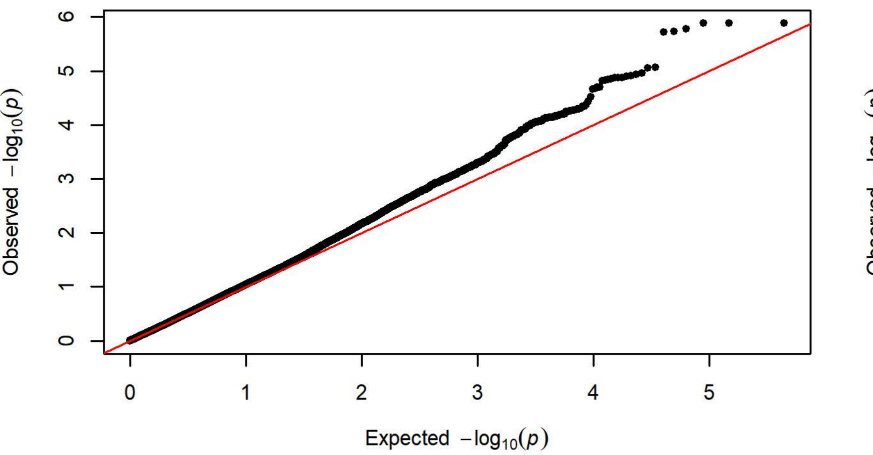


**Q-Q Plot Magazaine Entries Day 3 - Harlan 202A/C-208A (n=1096)**

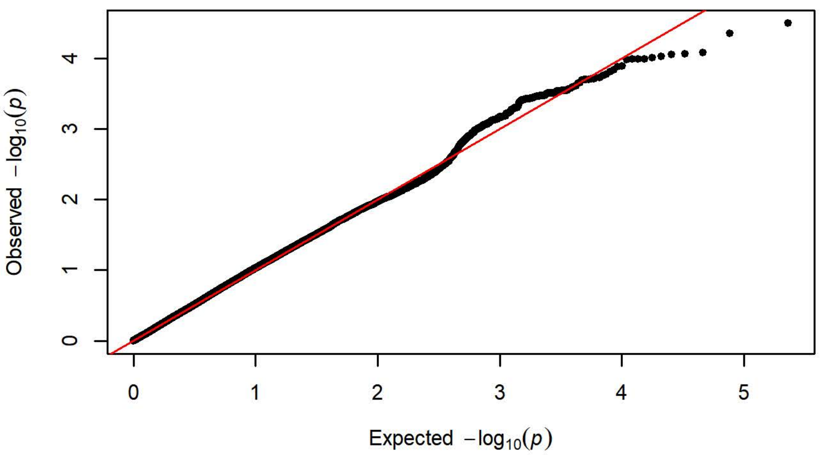


**Q-Q Plot Magazaine Entries Day 3 - Charles River R04 (n=650)**

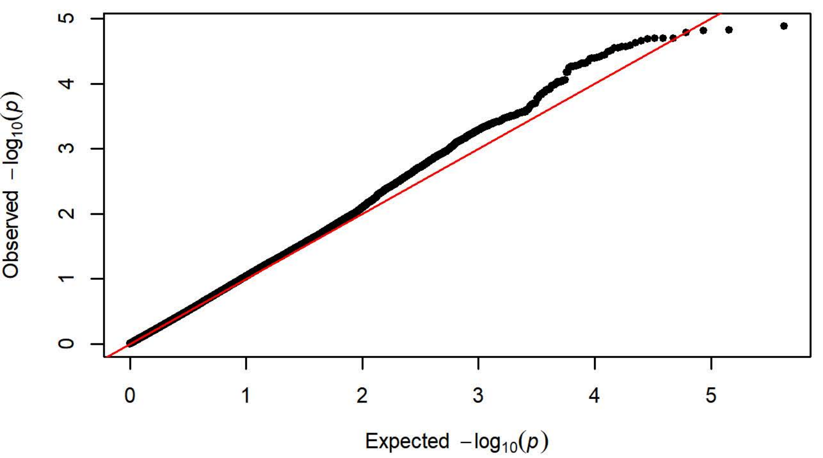


**Q-Q Plot Magazaine Entries Day 3 - Harlan 206 (n=758)**

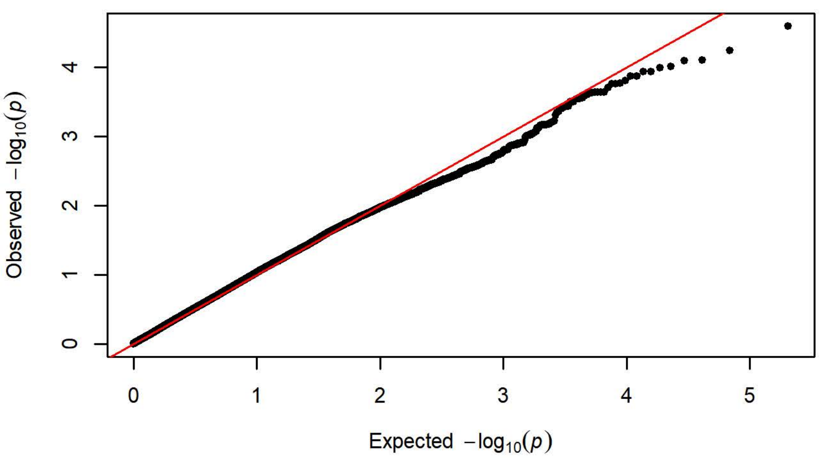


**Q-Q Plot Magazaine Entries Day 3 - Charles River P09 (n=295)**

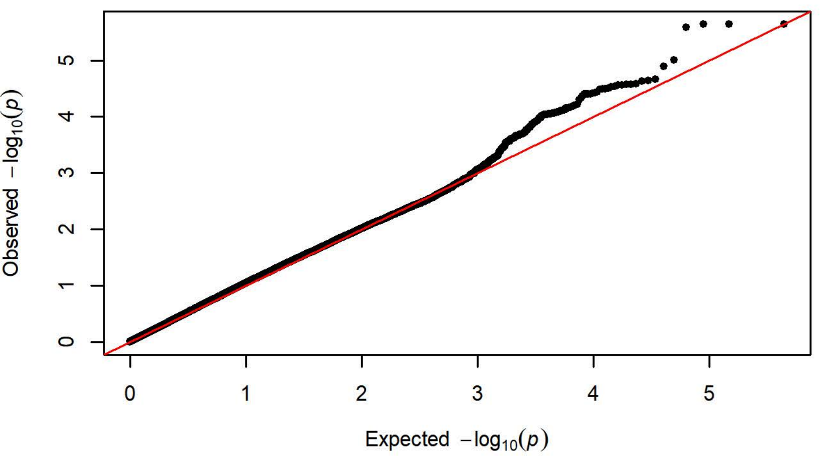


**Q-Q Plot Magazaine Entries Day 3 - Harlan 217 (n=350)**

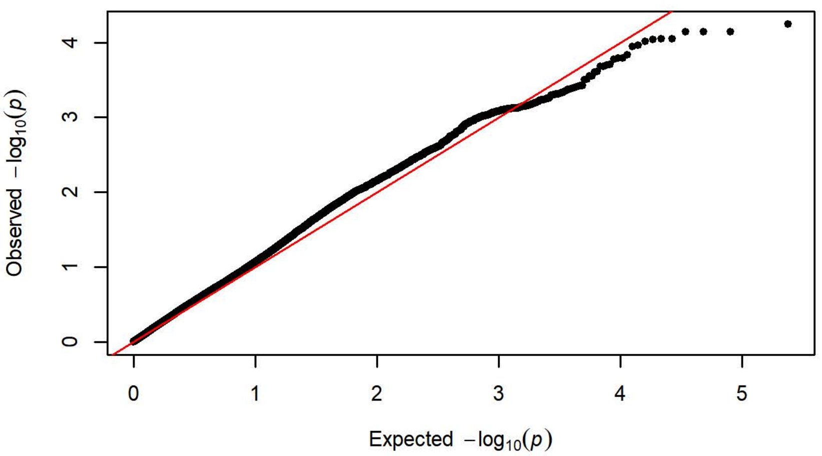


**Q-Q Plot Magazaine Entries Day 3 - Charles River C72 (n=358)**

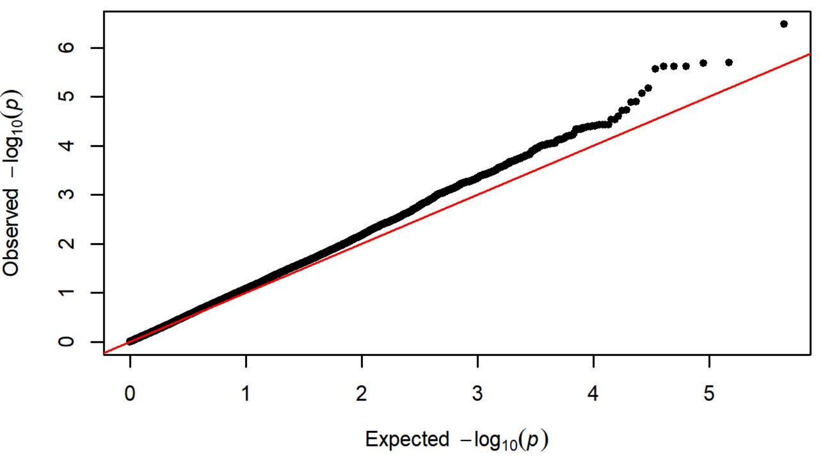

**Q-Q Plot Magazaine Entries Day 4 - Charles River R09-P3/7/10 (n=425**

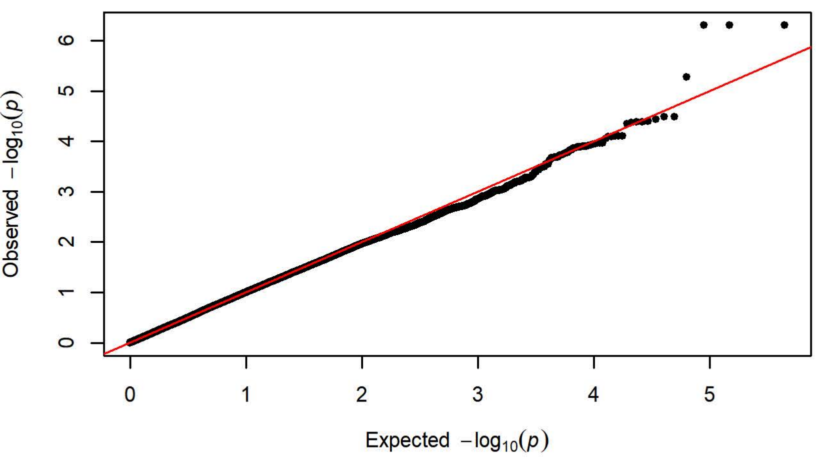


**Q-Q Plot Magazaine Entries Day 4 - Harlan 202A/C-208A (n=1099)**

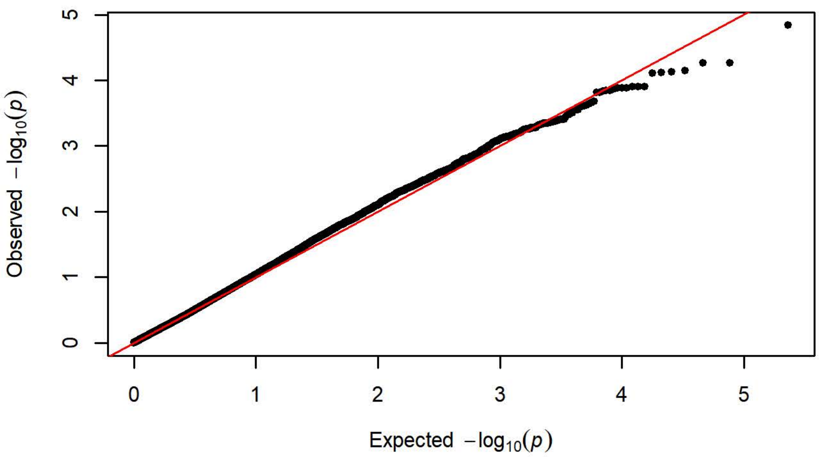


**Q-Q Plot Magazaine Entries Day 4 - Charles River R04 (n=650)**

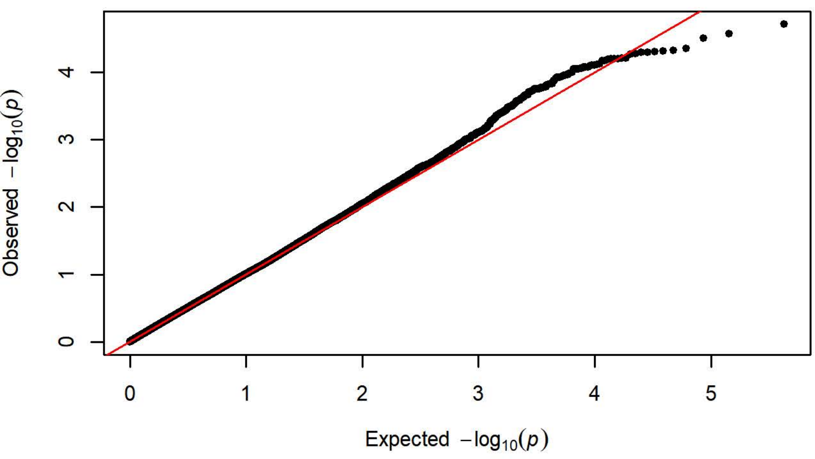


**Q-Q Plot Magazaine Entries Day 4 - Harlan 206 (n=758)**

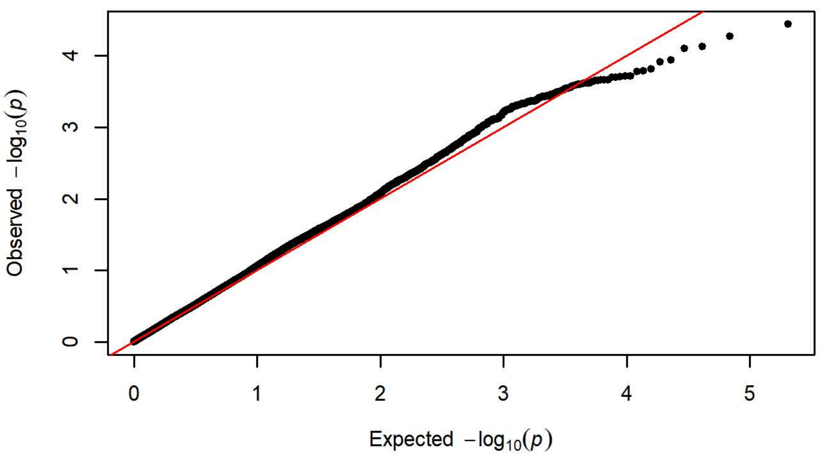


**Q-Q Plot Magazaine Entries Day 4 - Charles River P09 (n=295)**

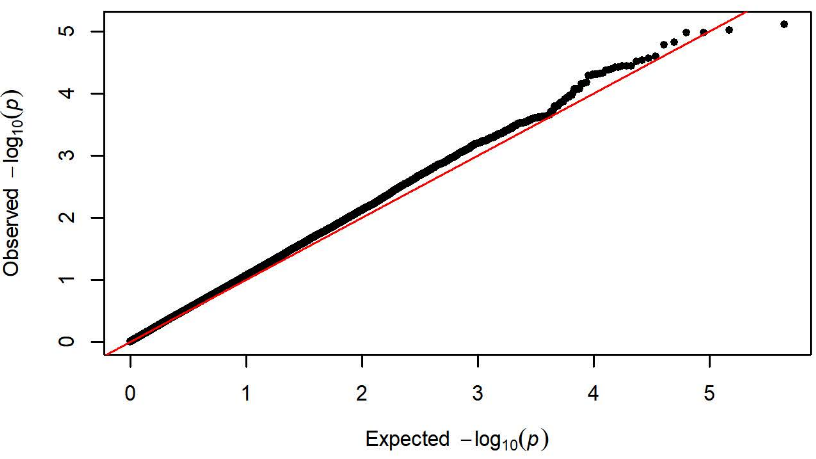


**Q-Q Plot Magazaine Entries Day 4 - Harlan 217 (n=351)**

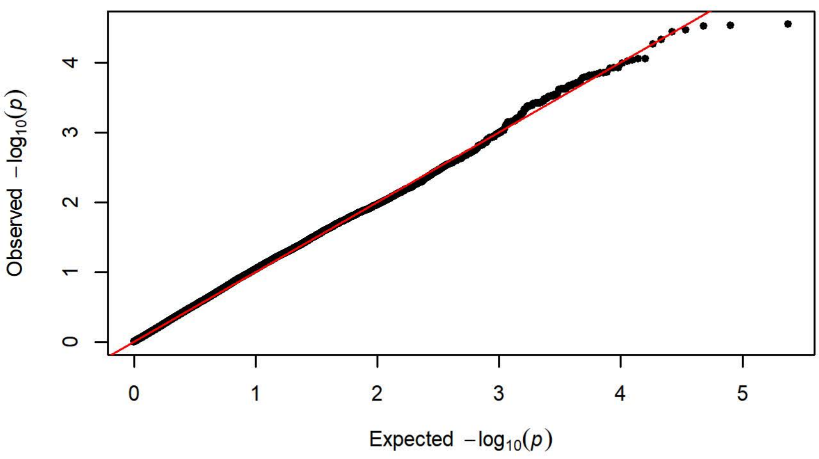


**Q-Q Plot Magazaine Entries Day 4 - Charles River C72 (n=358)**

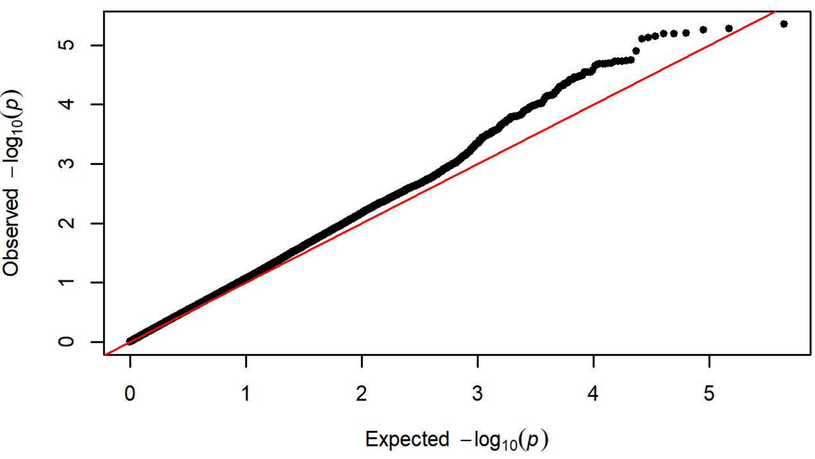

Q-Q Plot Magazaine Entries Day 5 - Charles River R09-P3/7/10 (n=425

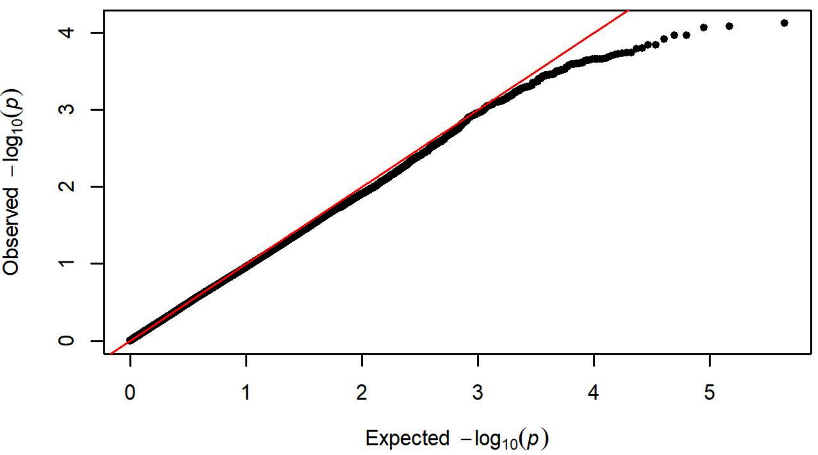


Q-Q Plot Magazaine Entries Day 5 - Harlan 202A/C-208A (n=1099)

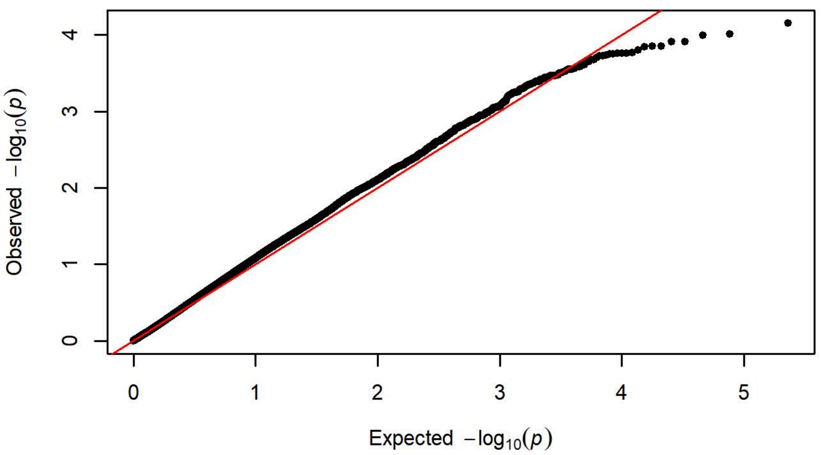


Q-Q Plot Magazaine Entries Day 5 - Charles River R04 (n=650)

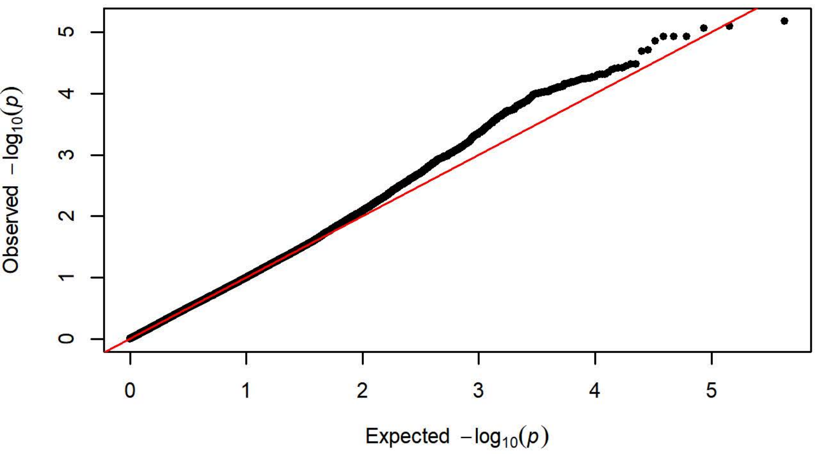


Q-Q Plot Magazaine Entries Day 5 - Harlan 206 (n=758)

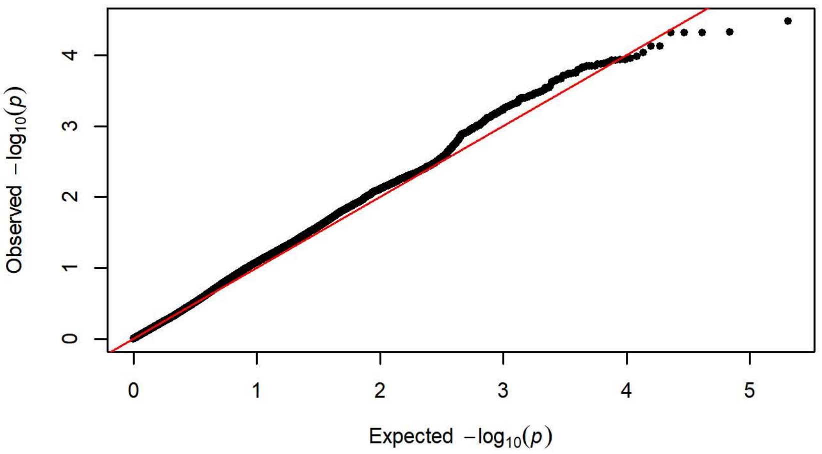


Q-Q Plot Magazaine Entries Day 5 - Charles River P09 (n=295)

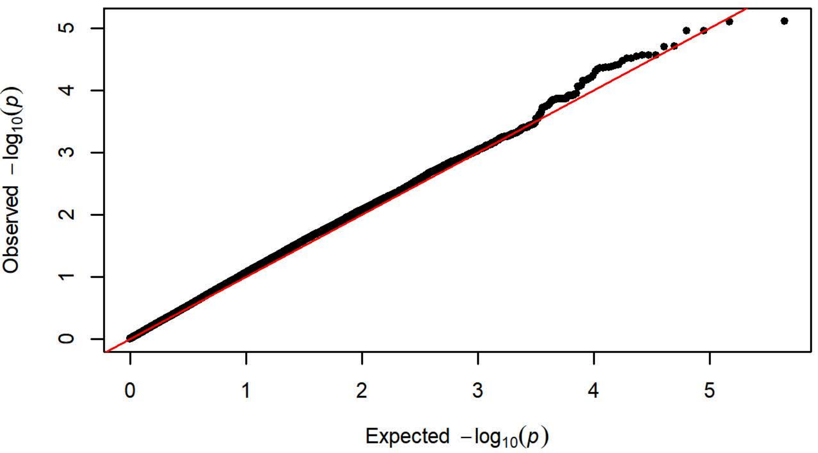


Q-Q Plot Magazaine Entries Day 5 - Harlan 217 (n=351)

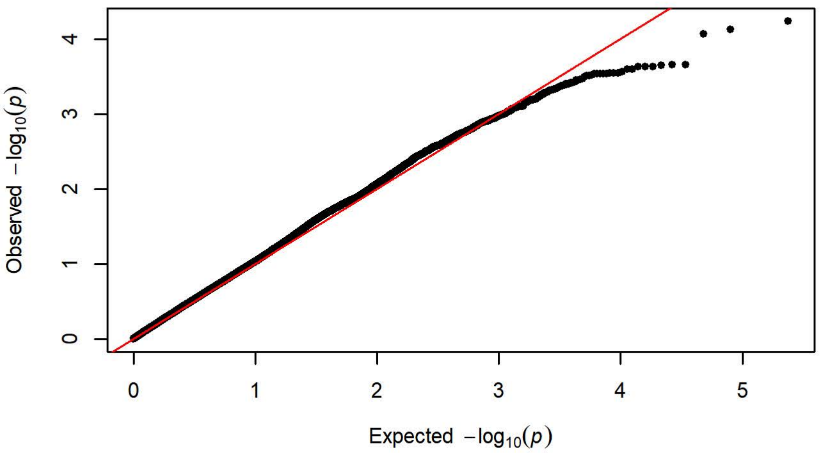


Q-Q Plot Magazaine Entries Day 5 - Charles River C72 (n=358)

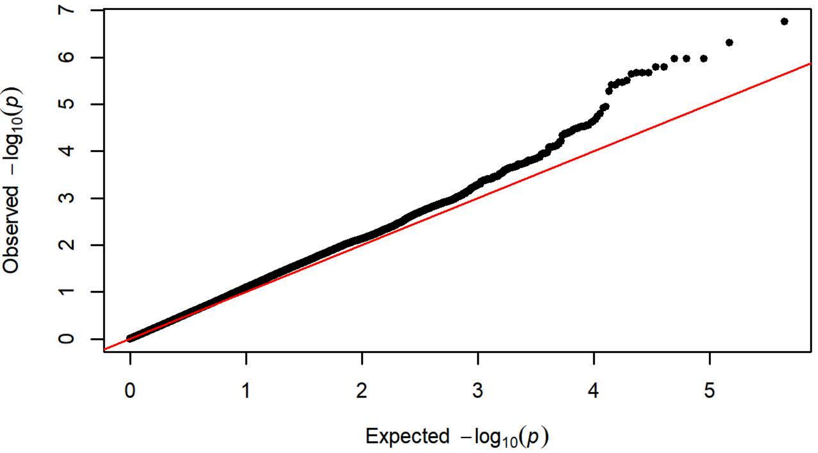

**Q-Q Plot Magazine Entries NCS Day 1 - Charles River R09-P3/7/10 (n=4[illegible]**

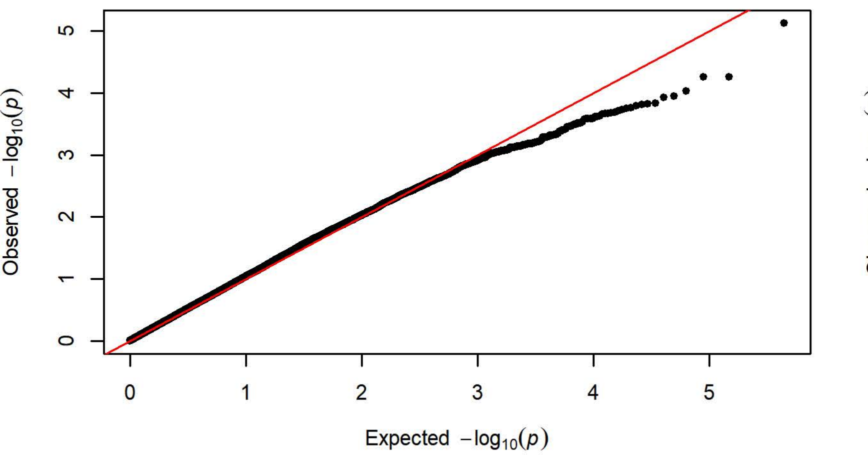


**Q-Q Plot Magazine Entries NCS Day 1 - Harlan 202A/C-208A (n=1099)**

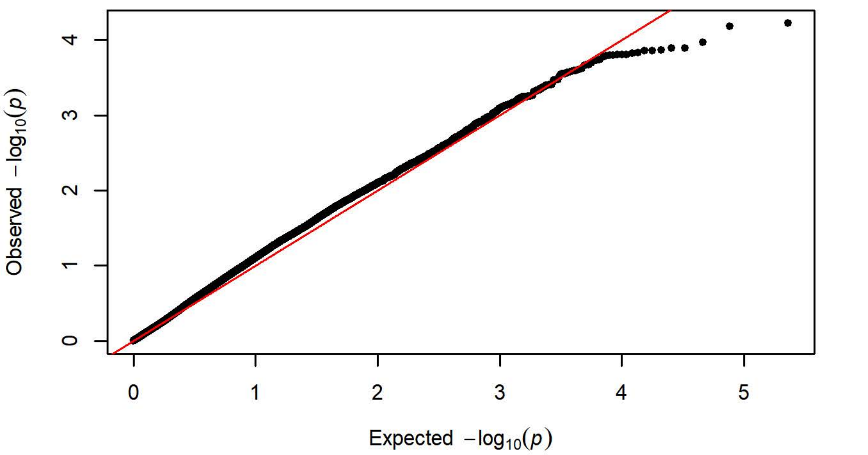


**Q-Q Plot Magazine Entries NCS Day 1 - Charles River R04 (n=650)**

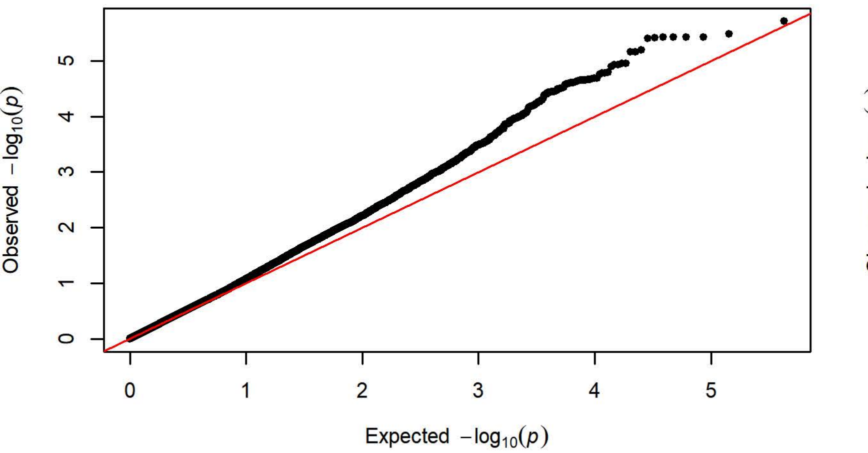


**Q-Q Plot Magazine Entries NCS Day 1 - Harlan 206 (n=757)**

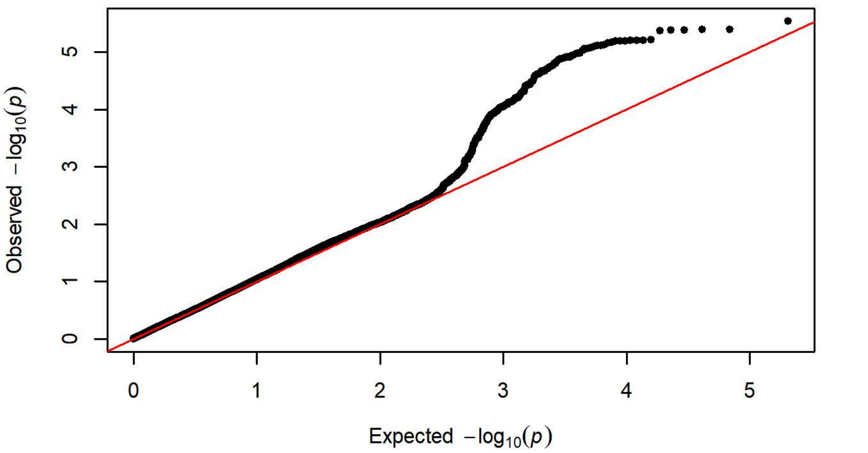


**Q-Q Plot Magazine Entries NCS Day 1 - Charles River P09 (n=295)**

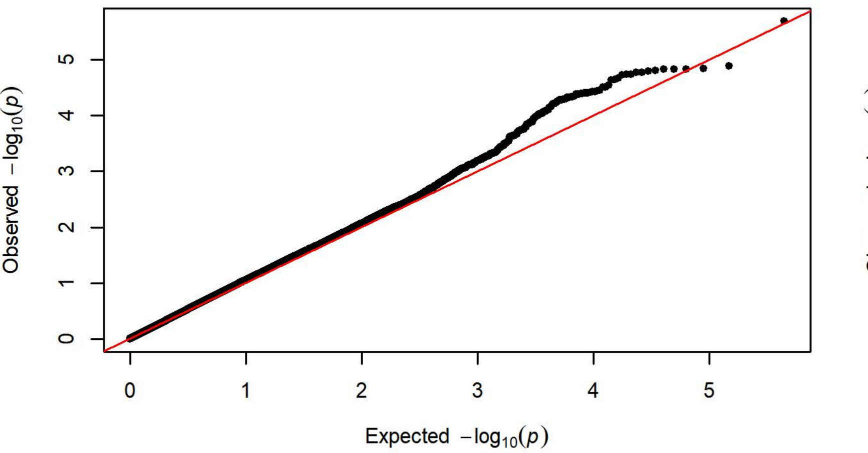


**Q-Q Plot Magazine Entries NCS Day 1 - Harlan 217 (n=349)**

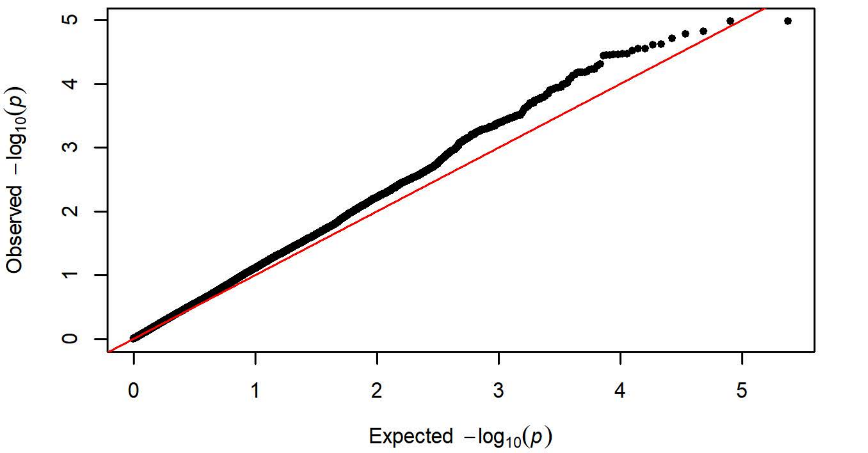


**Q-Q Plot Magazine Entries NCS Day 1 - Charles River C72 (n=358)**

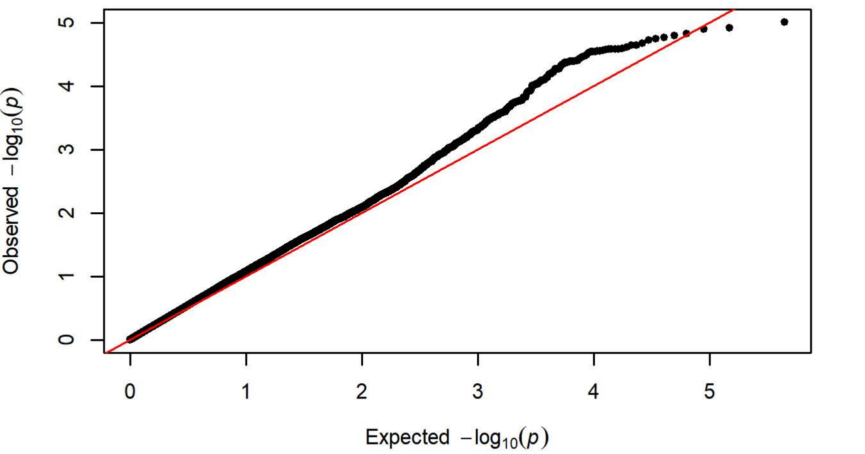

Q-Q Plot Magazine Entries NCS Day 2 - Charles River R09-P3/7/10 (n=4

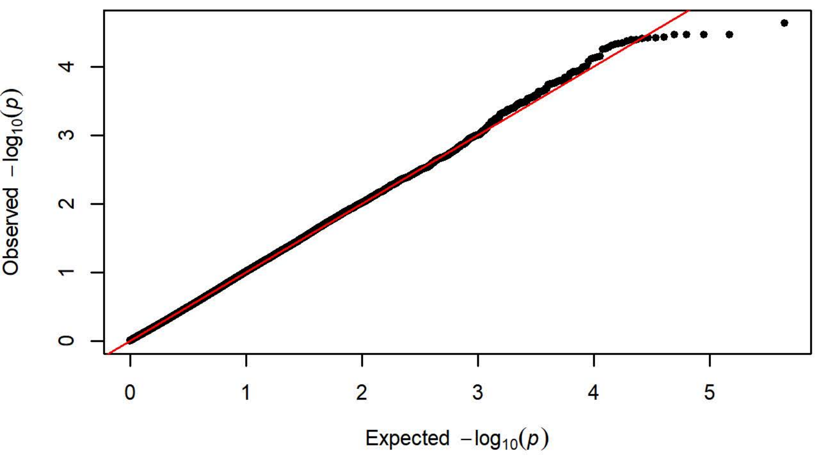


Q-Q Plot Magazine Entries NCS Day 2 - Harlan 202A/C-208A (n=1099)

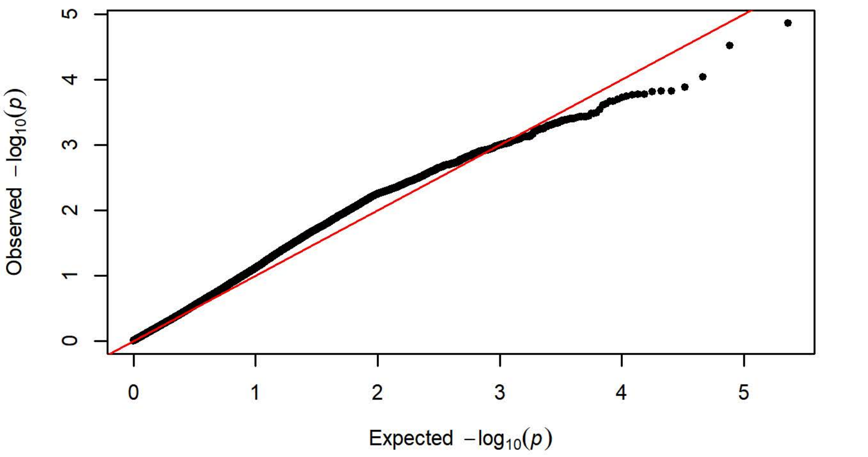


Q-Q Plot Magazine Entries NCS Day 2 - Charles River R04 (n=650)

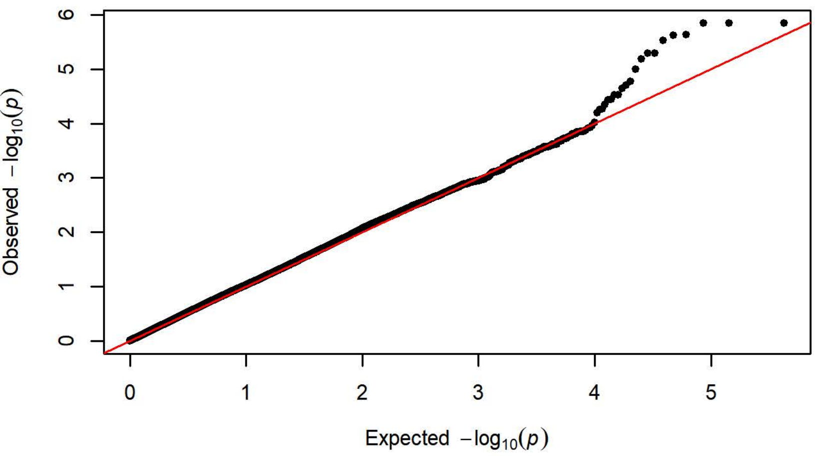


Q-Q Plot Magazine Entries NCS Day 2 - Harlan 206 (n=758)

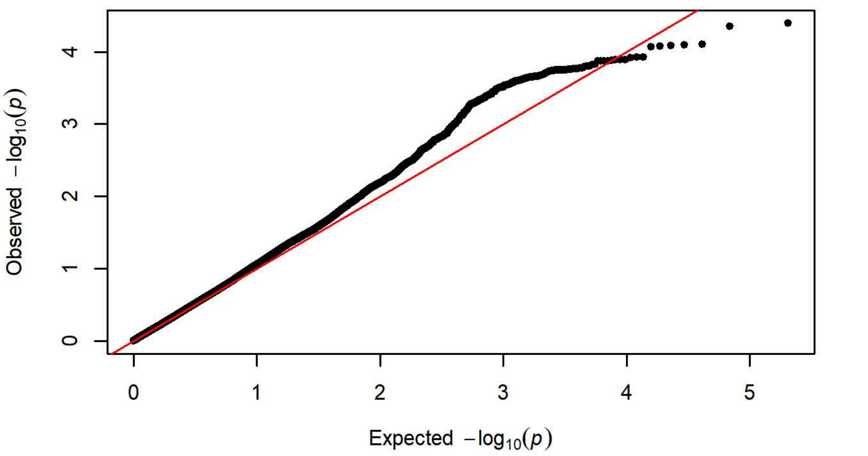


Q-Q Plot Magazine Entries NCS Day 2 - Charles River P09 (n=295)

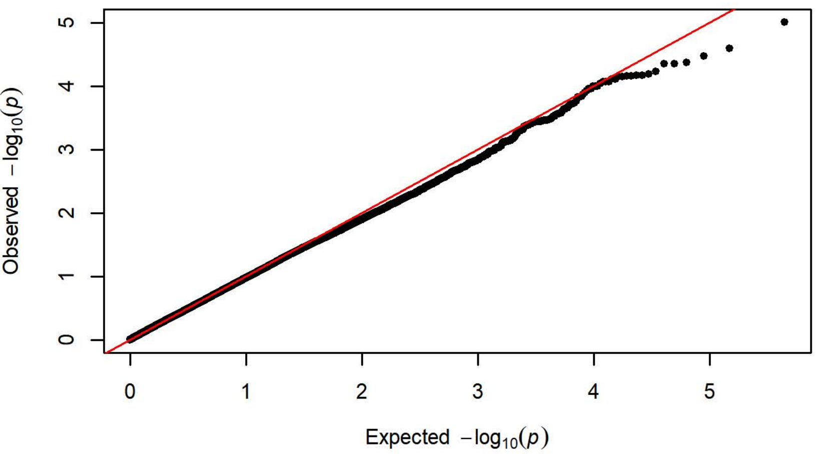


Q-Q Plot Magazine Entries NCS Day 2 - Harlan 217 (n=350)

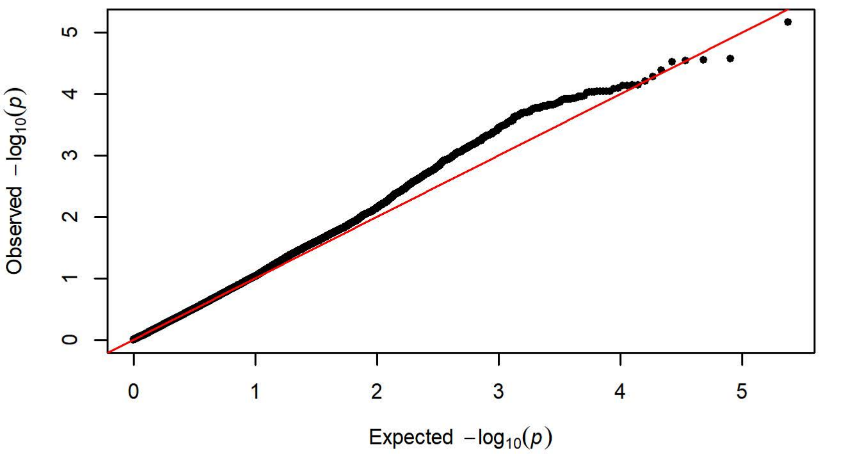


Q-Q Plot Magazine Entries NCS Day 2 - Charles River C72 (n=356)

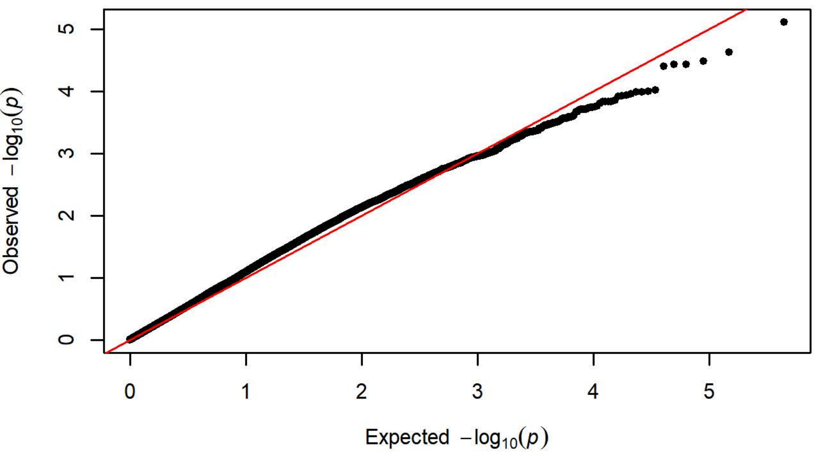

**Q-Q Plot Magazine Entries NCS Day 3 - Charles River R09-P3/7/10 (n=4**

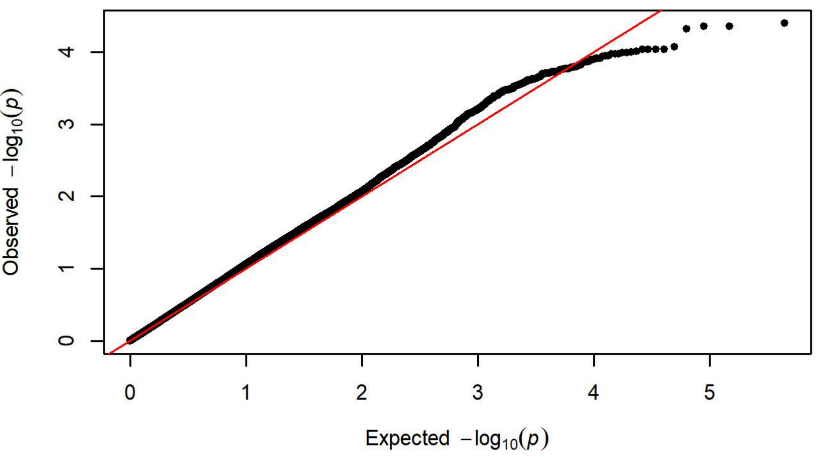


**Q-Q Plot Magazine Entries NCS Day 3 - Harlan 202A/C-208A (n=1096)**

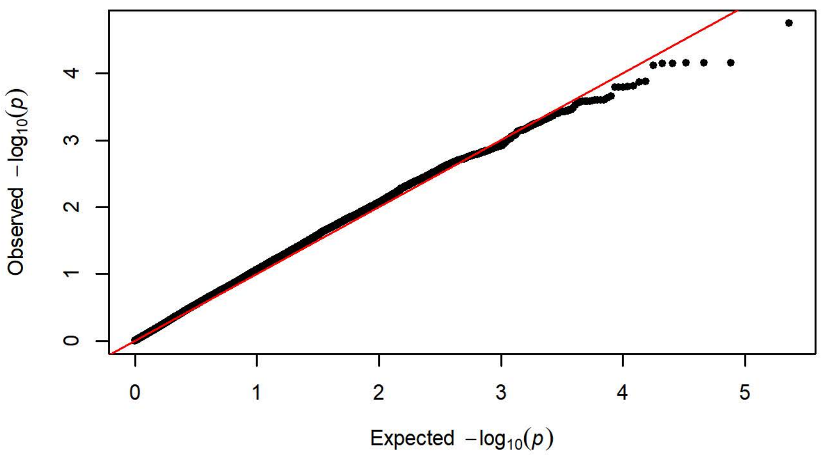


**Q-Q Plot Magazine Entries NCS Day 3 - Charles River R04 (n=650)**

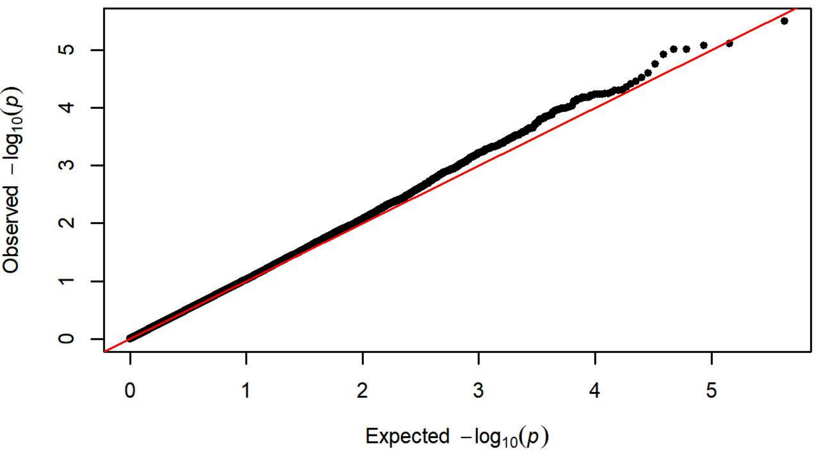


**Q-Q Plot Magazine Entries NCS Day 3 - Harlan 206 (n=758)**

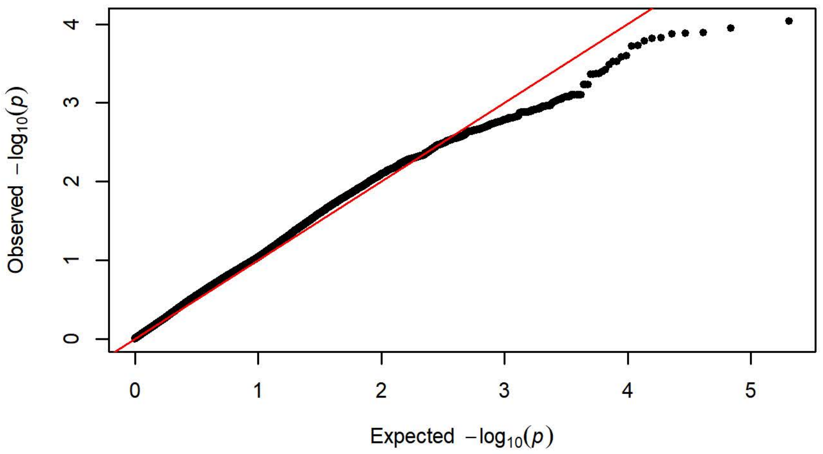


**Q-Q Plot Magazine Entries NCS Day 3 - Charles River P09 (n=295)**

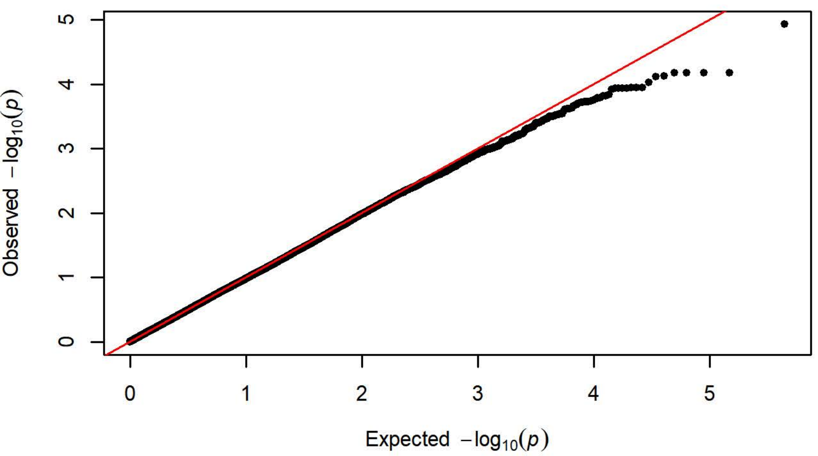


**Q-Q Plot Magazine Entries NCS Day 3 - Harlan 217 (n=350)**

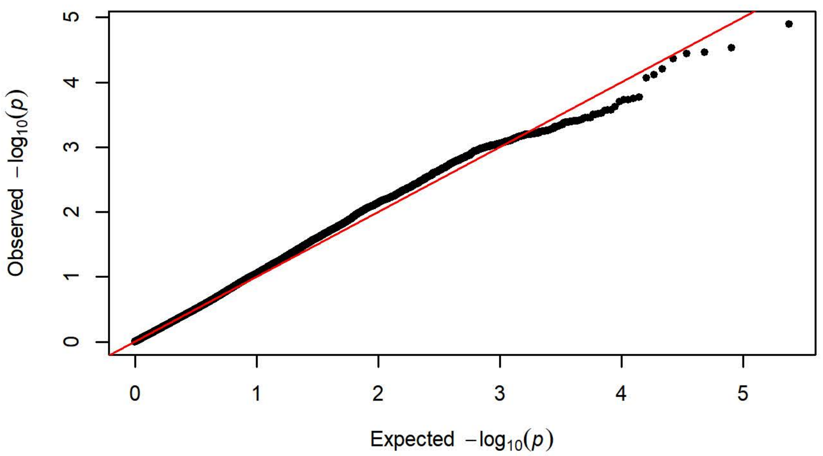


**Q-Q Plot Magazine Entries NCS Day 3 - Charles River C72 (n=358)**

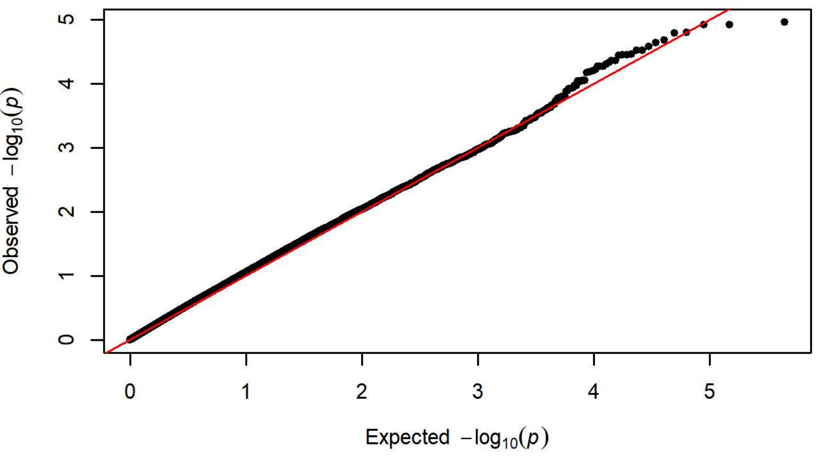

Q-Q Plot Magazine Entries NCS Day 4 - Charles River R09-P3/7/10 (n=4

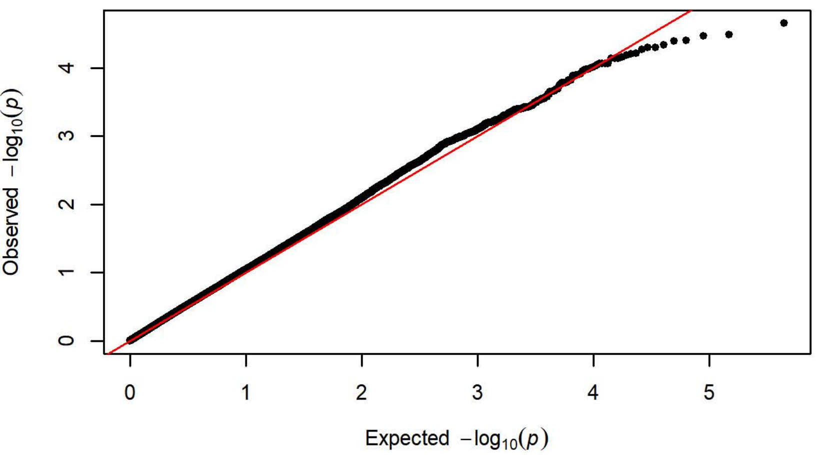


Q-Q Plot Magazine Entries NCS Day 4 - Harlan 202A/C-208A (n=1099)

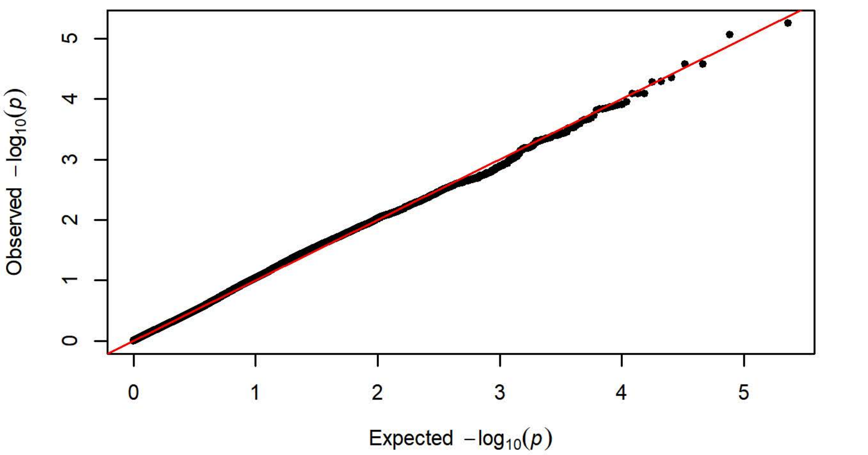


Q-Q Plot Magazine Entries NCS Day 4 - Charles River R04 (n=650)

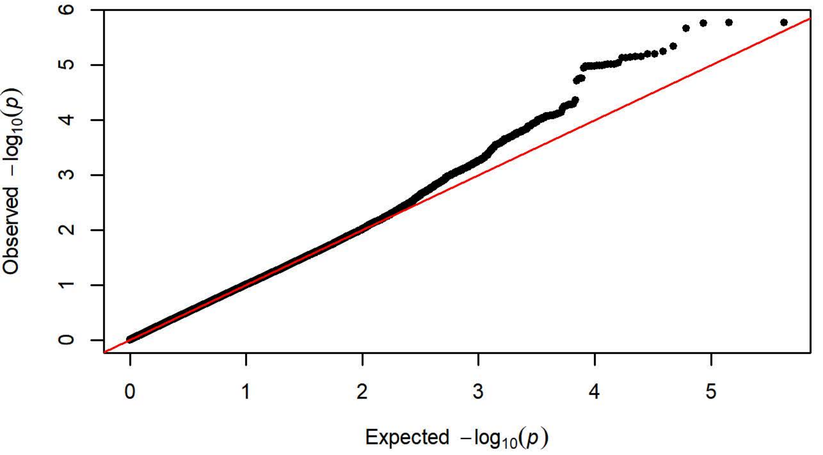


Q-Q Plot Magazine Entries NCS Day 4 - Harlan 206 (n=758)

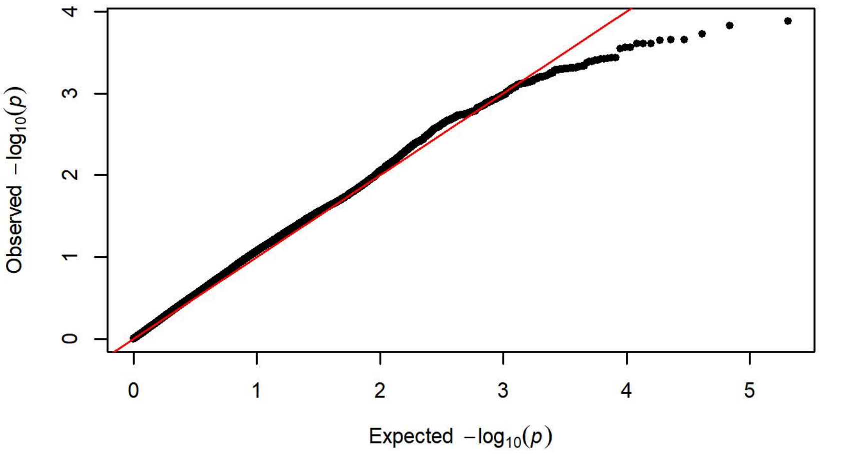


Q-Q Plot Magazine Entries NCS Day 4 - Charles River P09 (n=295)

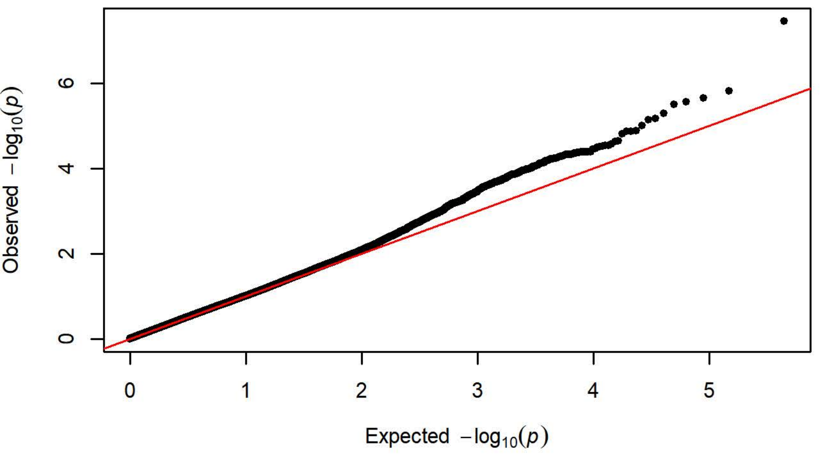


Q-Q Plot Magazine Entries NCS Day 4 - Harlan 217 (n=351)

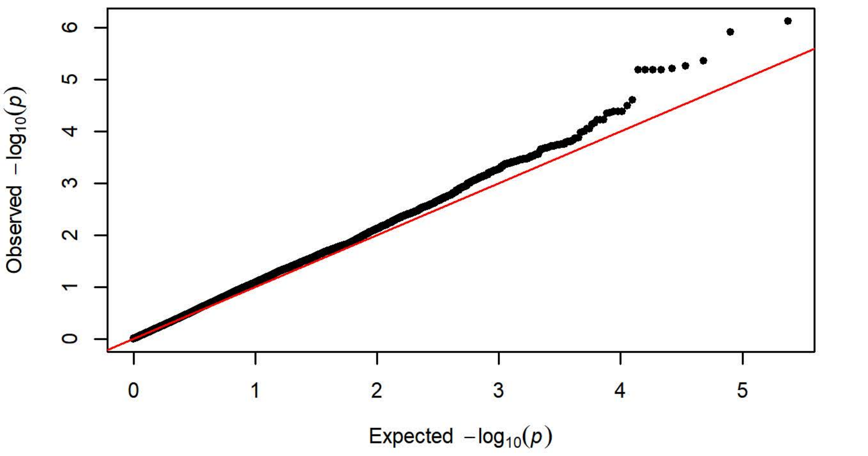


Q-Q Plot Magazine Entries NCS Day 4 - Charles River C72 (n=358)

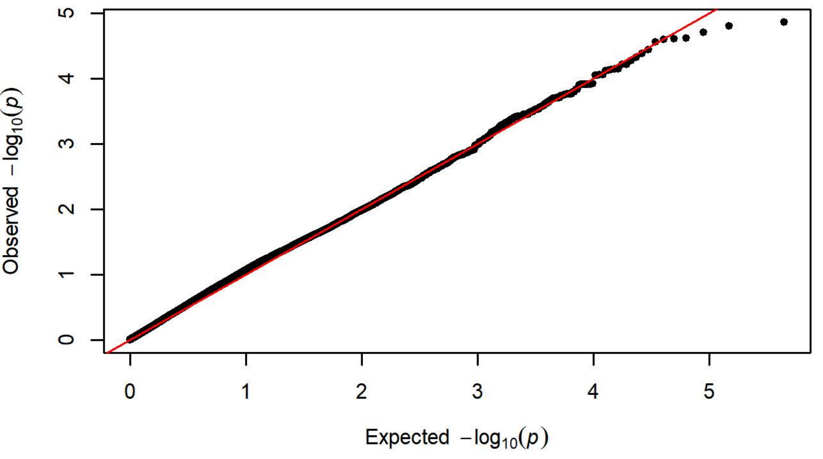

**Q-Q Plot Magazine Entries NCS Day 5 - Charles River R09-P3/7/10 (n=4**

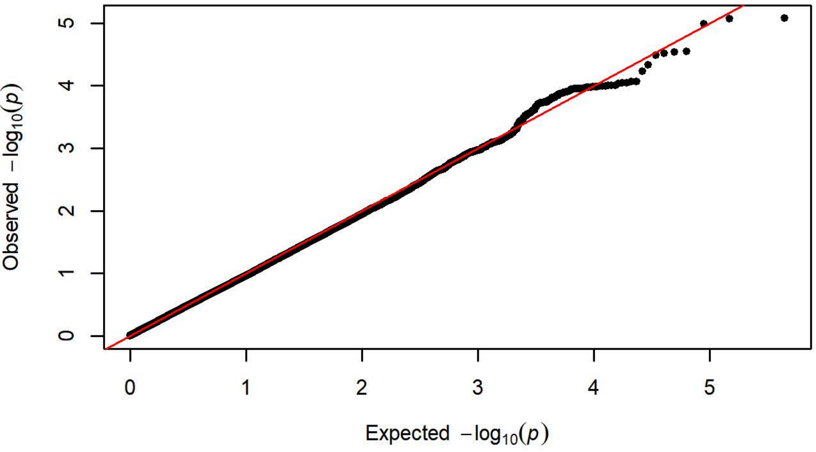


**Q-Q Plot Magazine Entries NCS Day 5 - Harlan 202A/C-208A (n=1099)**

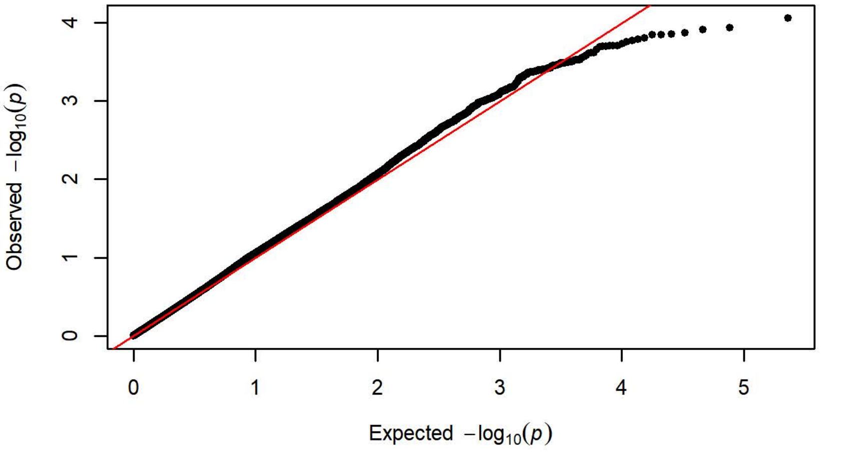


**Q-Q Plot Magazine Entries NCS Day 5 - Charles River R04 (n=650)**

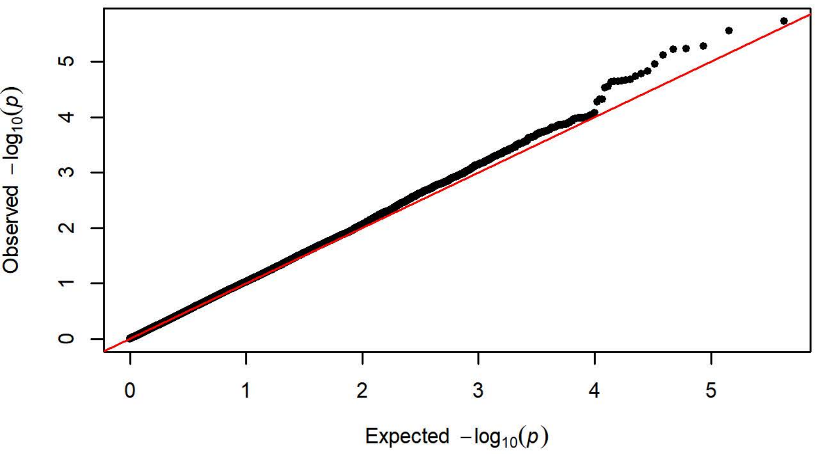


**Q-Q Plot Magazine Entries NCS Day 5 - Harlan 206 (n=758)**

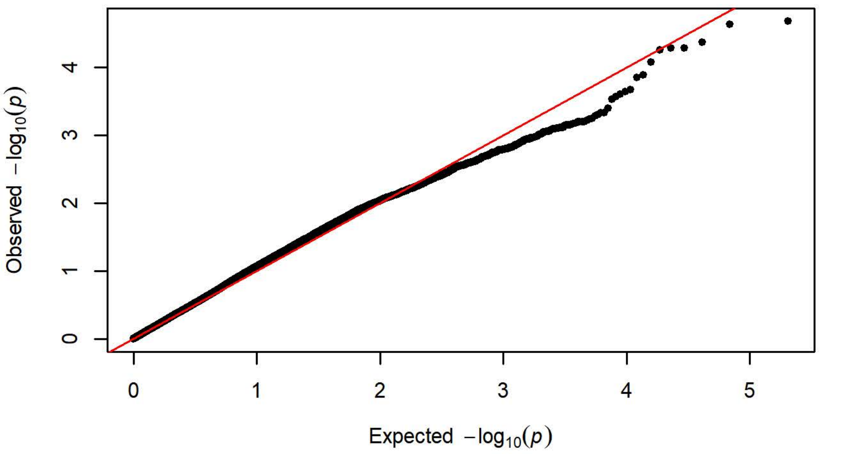


**Q-Q Plot Magazine Entries NCS Day 5 - Charles River P09 (n=295)**

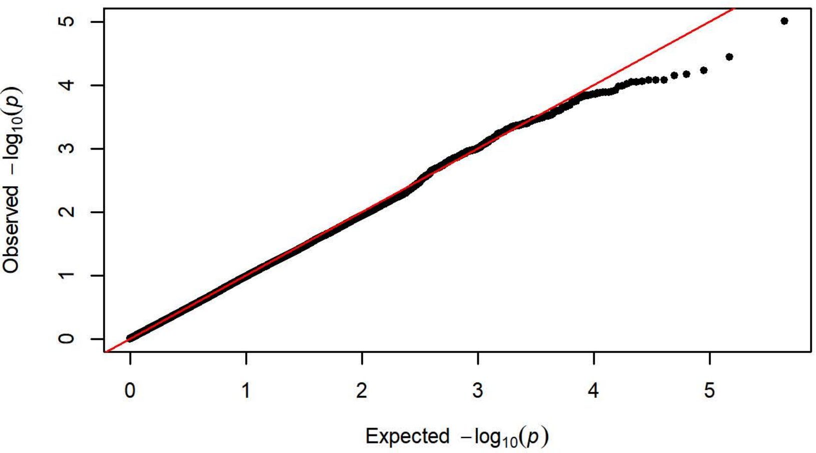


**Q-Q Plot Magazine Entries NCS Day 5 - Harlan 217 (n=351)**

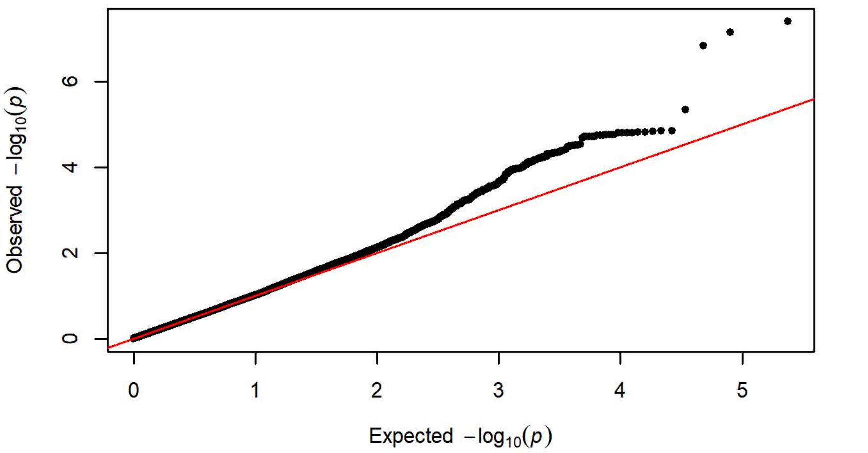


**Q-Q Plot Magazine Entries NCS Day 5 - Charles River C72 (n=358)**

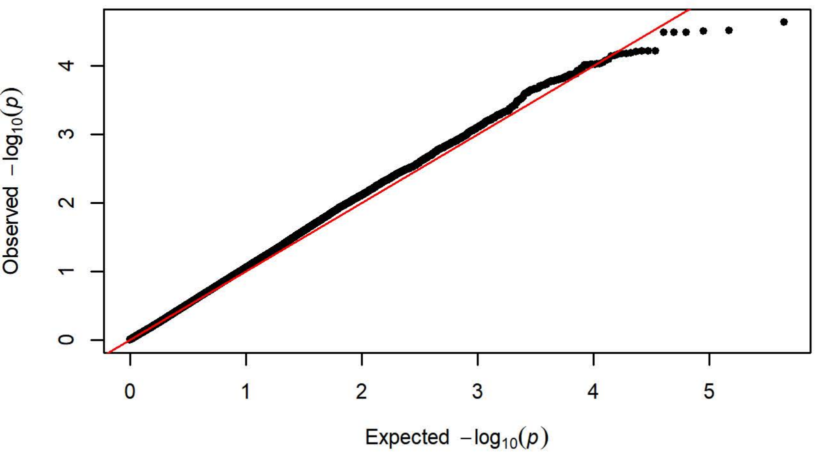

**Q-Q Plot Probability Difference Day 1 - Charles River R09-P3/7/10 (n=4[illegible]**

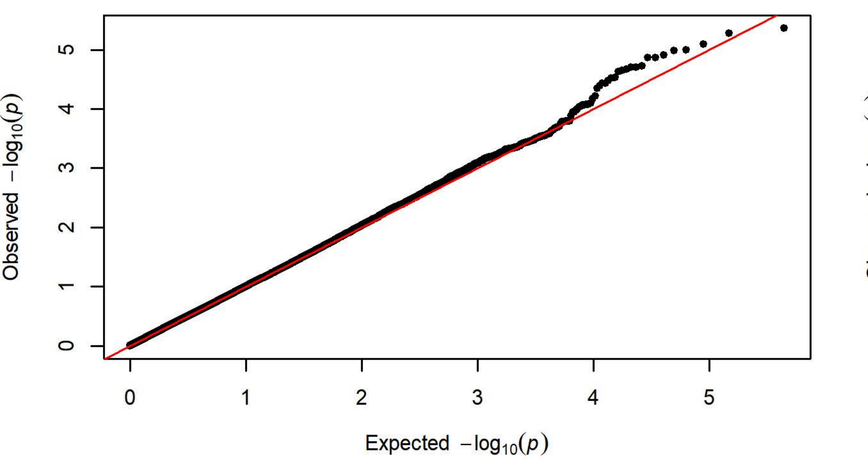


**Q-Q Plot Probability Difference Day 1 - Harlan 202A/C-208A (n=1066)**

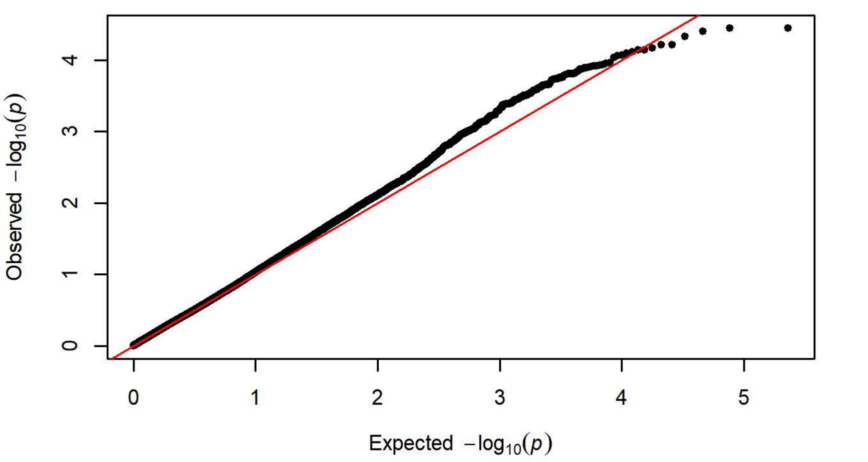


**Q-Q Plot Probability Difference Day 1 - Charles River R04 (n=650)**

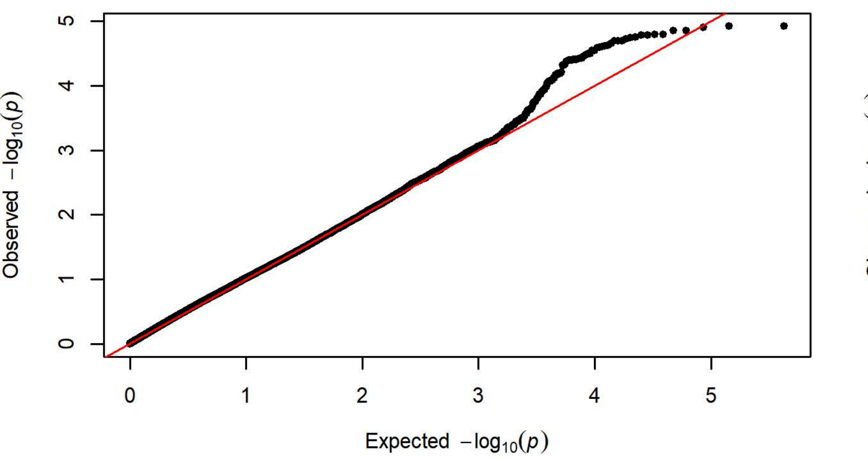


**Q-Q Plot Probability Difference Day 1 - Harlan 206 (n=758)**

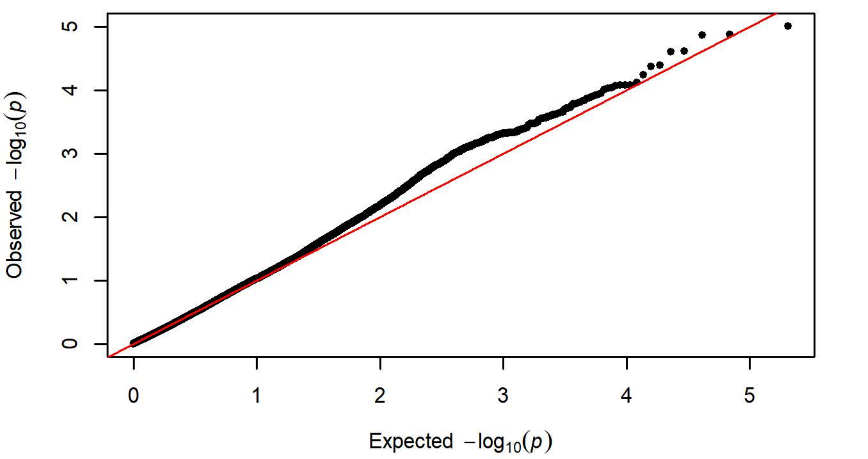


**Q-Q Plot Probability Difference Day 1 - Charles River P09 (n=295)**

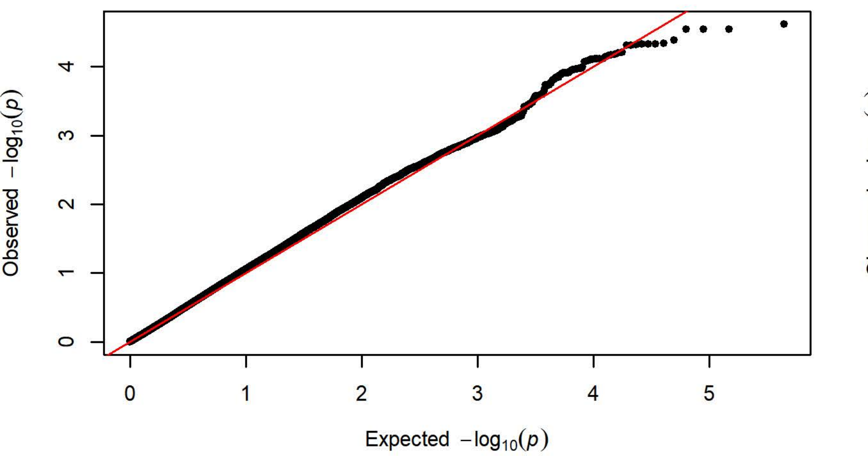


**Q-Q Plot Probability Difference Day 1 - Harlan 217 (n=351)**

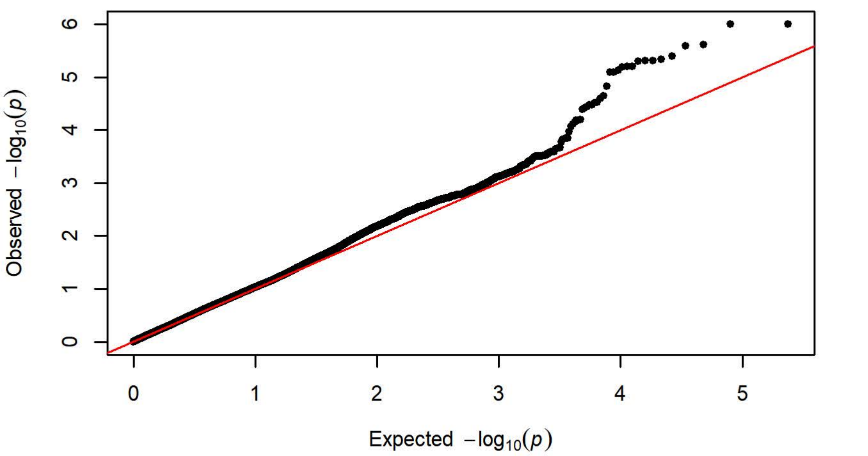


**Q-Q Plot Probability Difference Day 1 - Charles River C72 (n=358)**

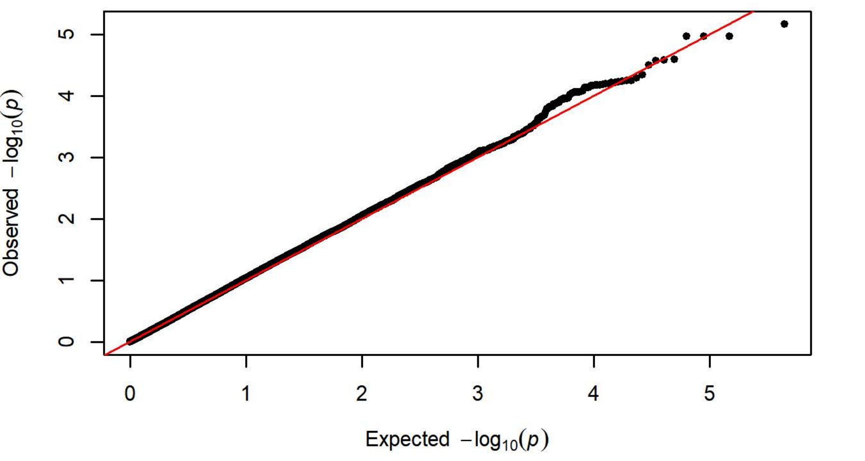

Q-Q Plot Probability Difference Day 2 - Charles River R09-P3/7/10 (n=4[illegible]

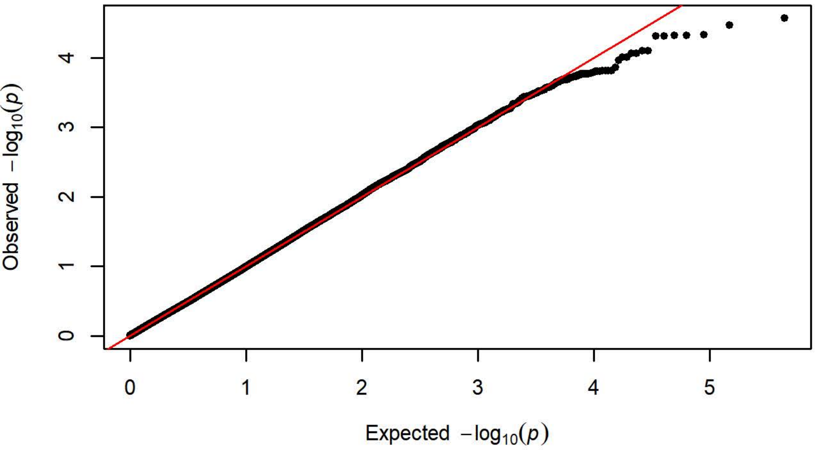


Q-Q Plot Probability Difference Day 2 - Harlan 202A/C-208A (n=1099)

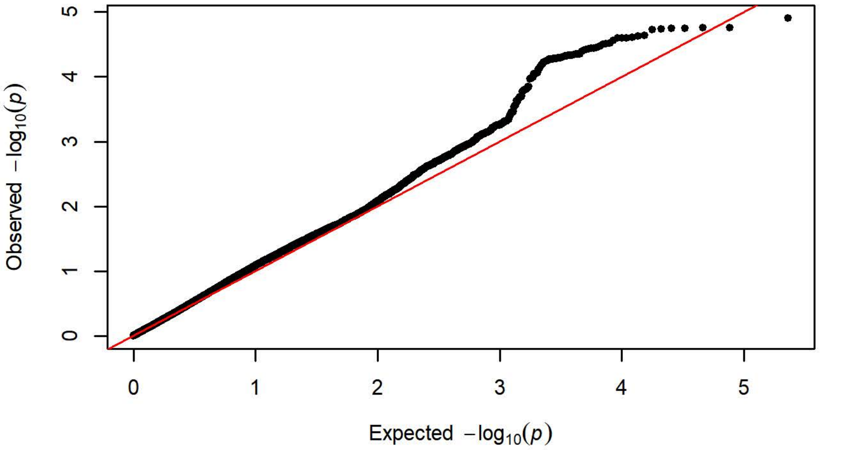


Q-Q Plot Probability Difference Day 2 - Charles River R04 (n=650)

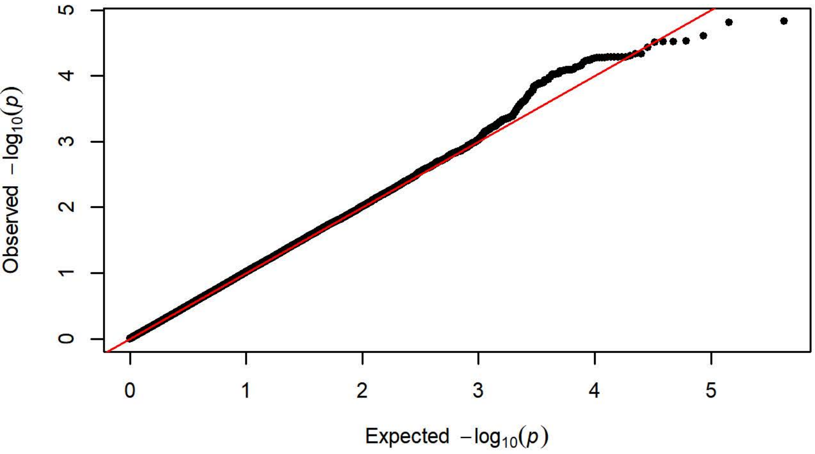


Q-Q Plot Probability Difference Day 2 - Harlan 206 (n=758)

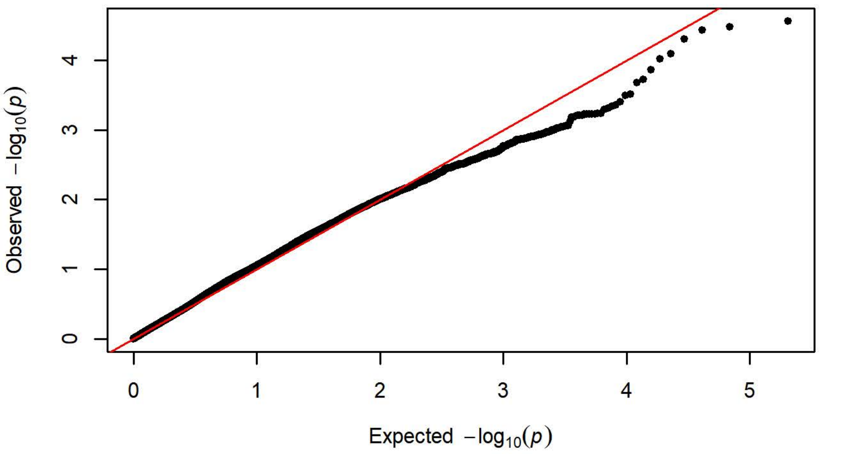


Q-Q Plot Probability Difference Day 2 - Charles River P09 (n=295)

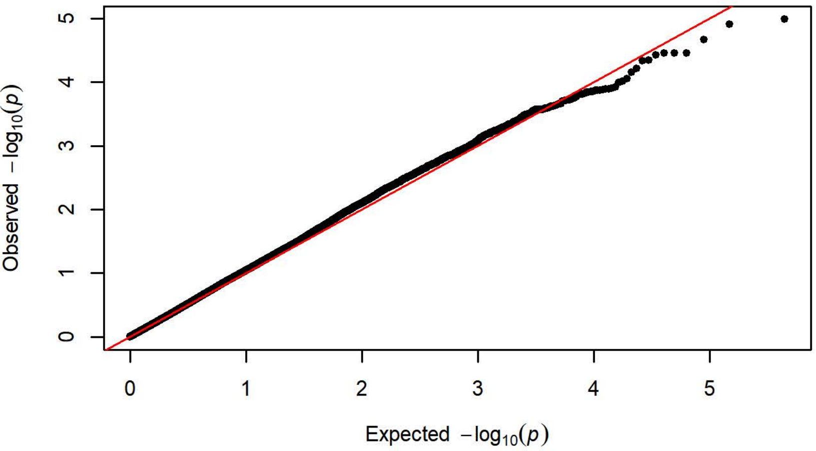


Q-Q Plot Probability Difference Day 2 - Harlan 217 (n=351)

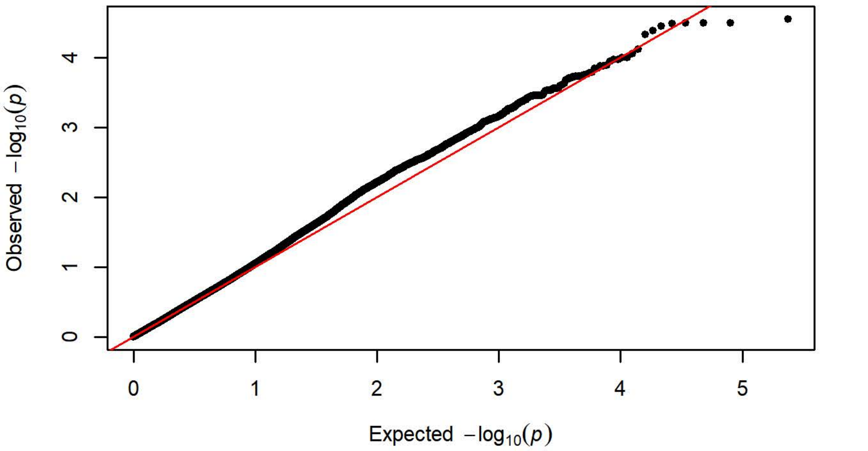


Q-Q Plot Probability Difference Day 2 - Charles River C72 (n=356)

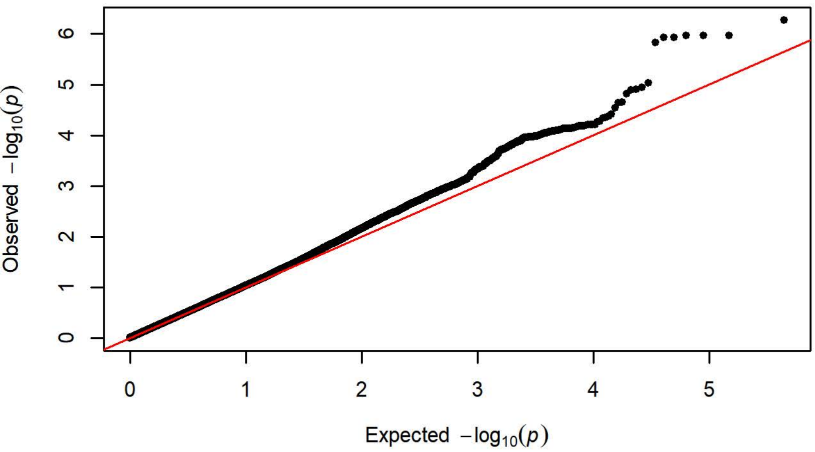

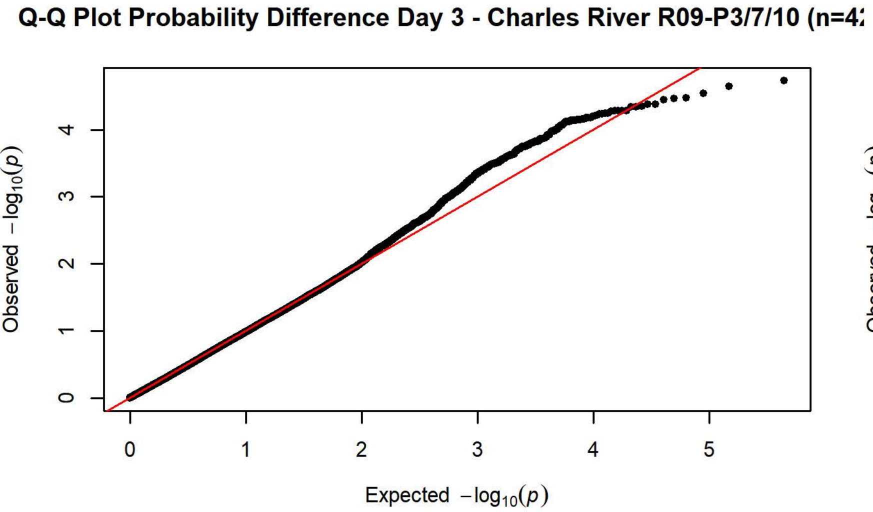
Q-Q Plot Probability Difference Day 3 - Charles River R09-P3/7/10 (n=4
Observed −log10(p)
Expected −log10(p)

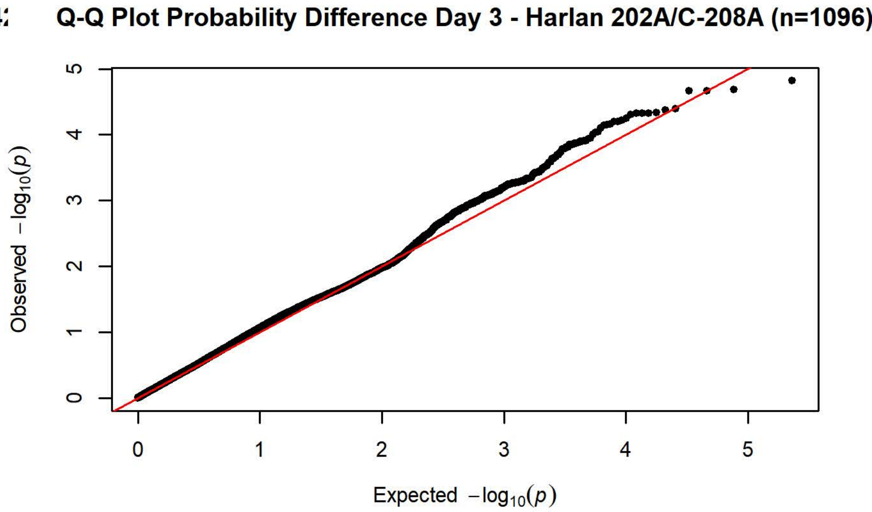
Q-Q Plot Probability Difference Day 3 - Harlan 202A/C-208A (n=1096)
Observed −log10(p)
Expected −log10(p)

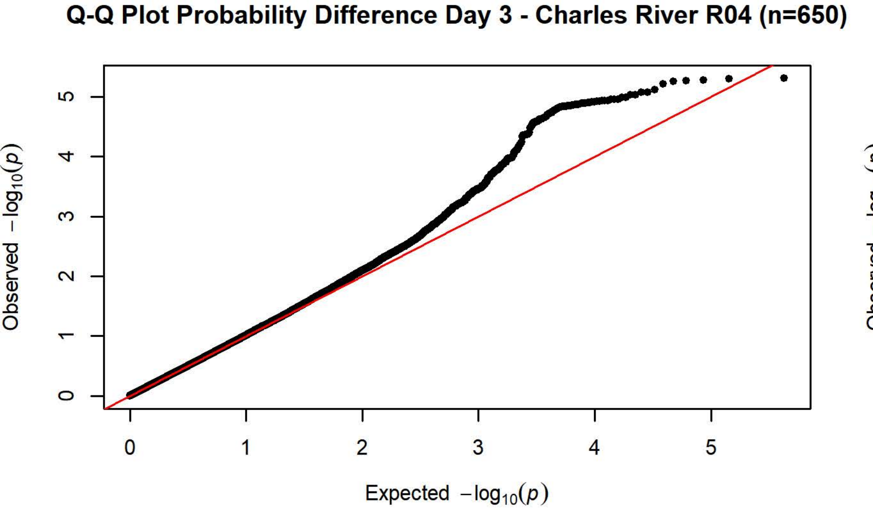
Q-Q Plot Probability Difference Day 3 - Charles River R04 (n=650)
Observed −log10(p)
Expected −log10(p)

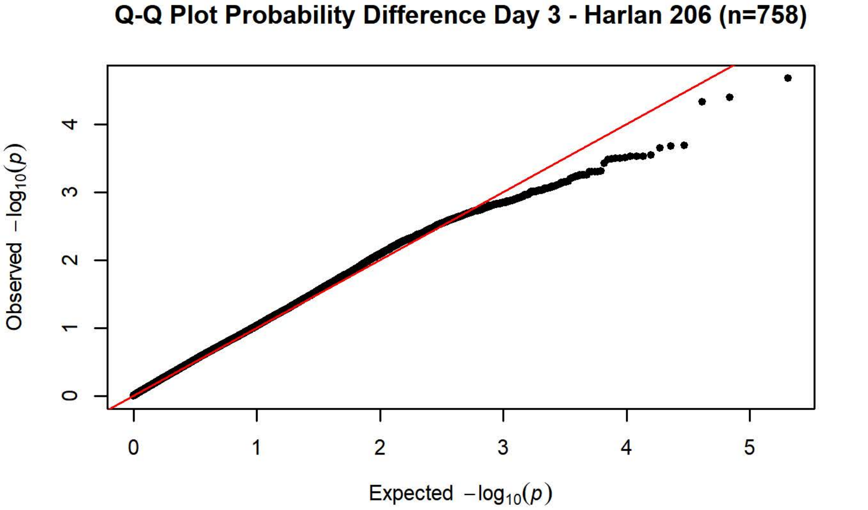
Q-Q Plot Probability Difference Day 3 - Harlan 206 (n=758)
Observed −log10(p)
Expected −log10(p)

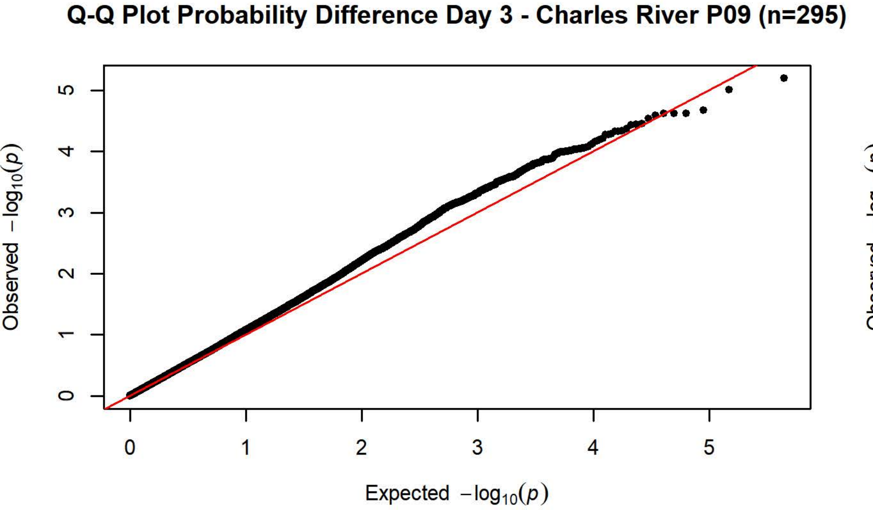
Q-Q Plot Probability Difference Day 3 - Charles River P09 (n=295)
Observed −log10(p)
Expected −log10(p)

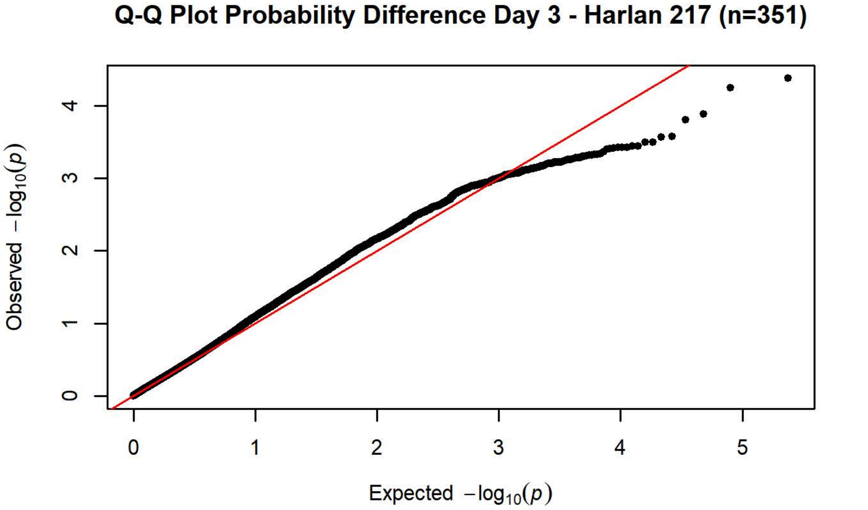
Q-Q Plot Probability Difference Day 3 - Harlan 217 (n=351)
Observed −log10(p)
Expected −log10(p)

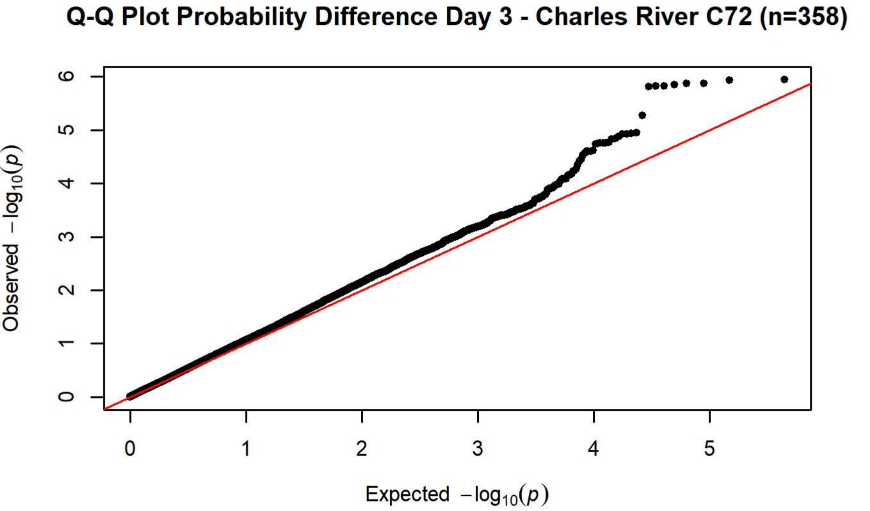
Q-Q Plot Probability Difference Day 3 - Charles River C72 (n=358)
Observed −log10(p)
Expected −log10(p)

**Q-Q Plot Probability Difference Day 4 - Charles River R09-P3/7/10 (n=4[illegible]**

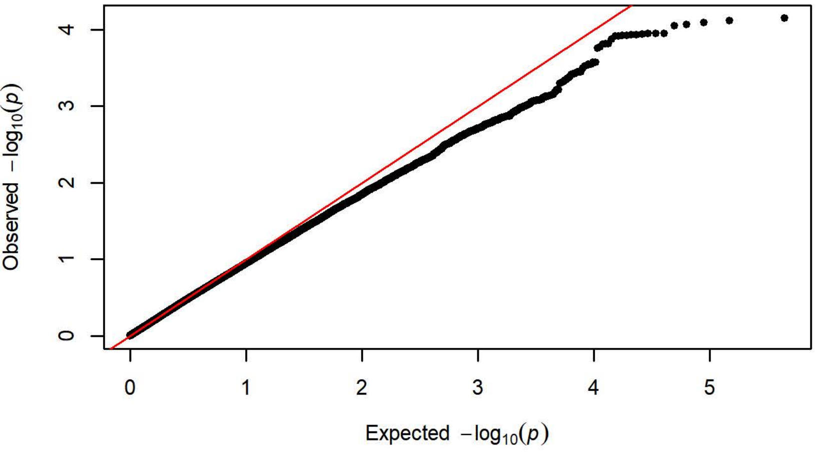


**Q-Q Plot Probability Difference Day 4 - Harlan 202A/C-208A (n=1099)**

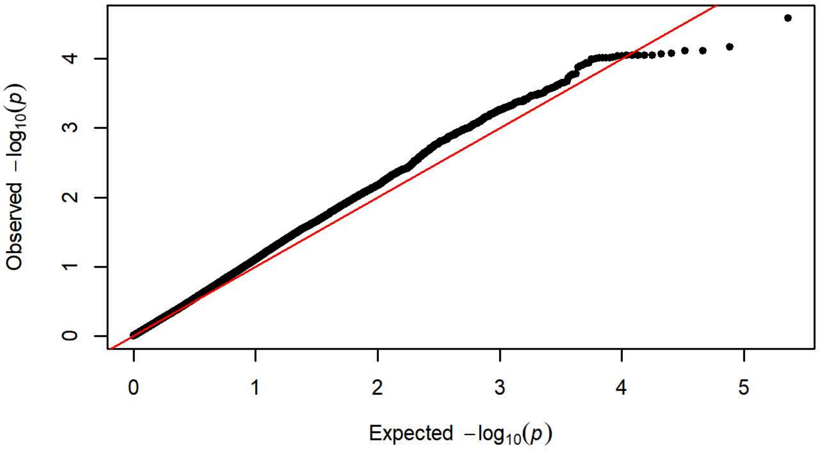


**Q-Q Plot Probability Difference Day 4 - Charles River R04 (n=650)**

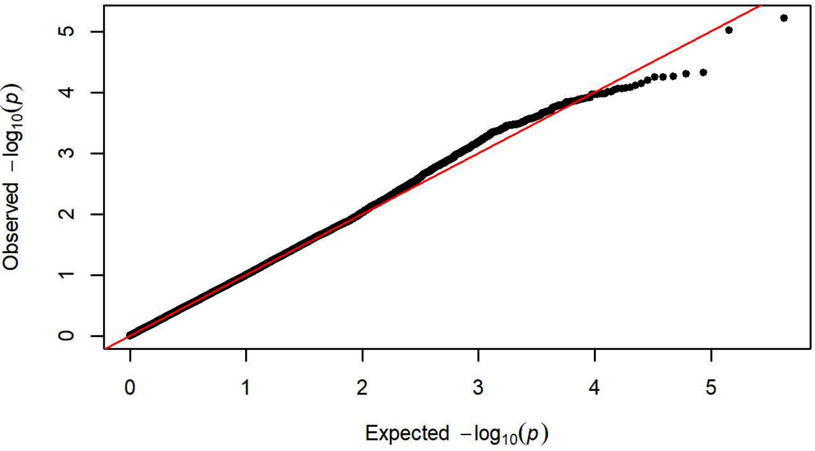


**Q-Q Plot Probability Difference Day 4 - Harlan 206 (n=758)**

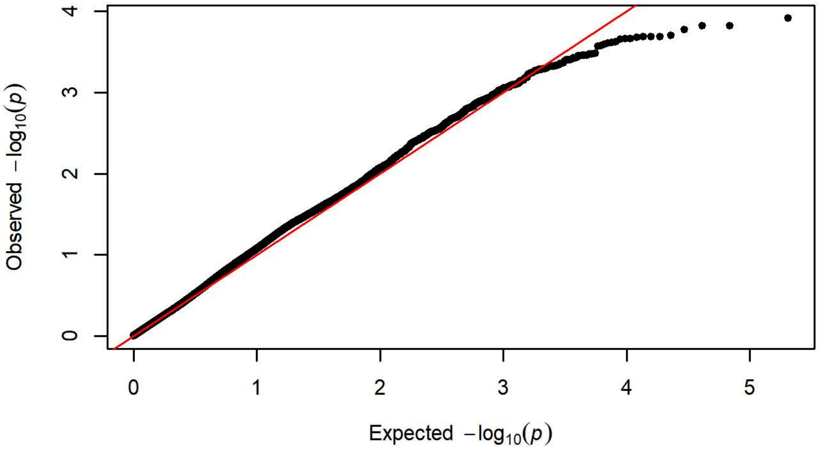


**Q-Q Plot Probability Difference Day 4 - Charles River P09 (n=295)**

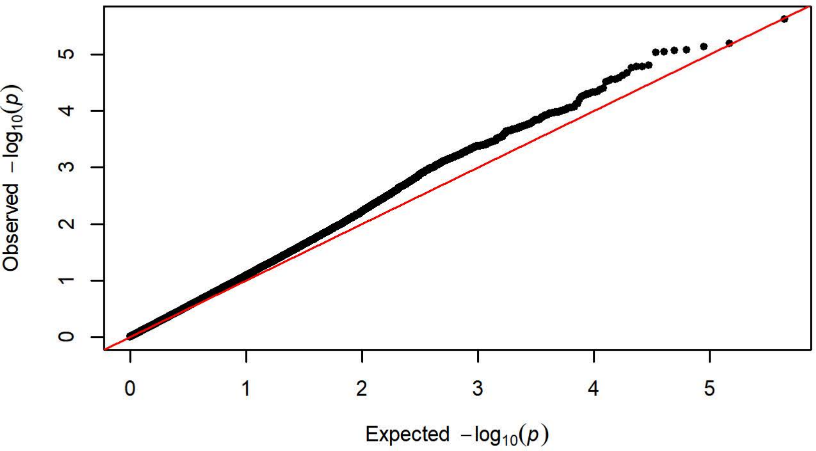


**Q-Q Plot Probability Difference Day 4 - Harlan 217 (n=351)**

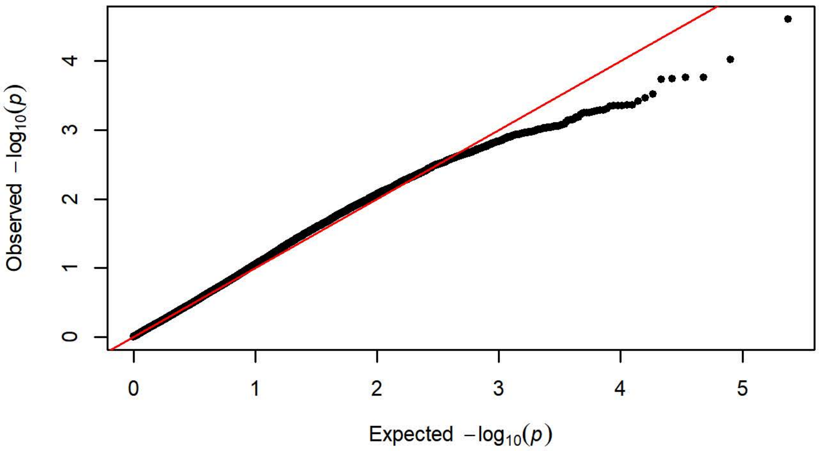


**Q-Q Plot Probability Difference Day 4 - Charles River C72 (n=358)**

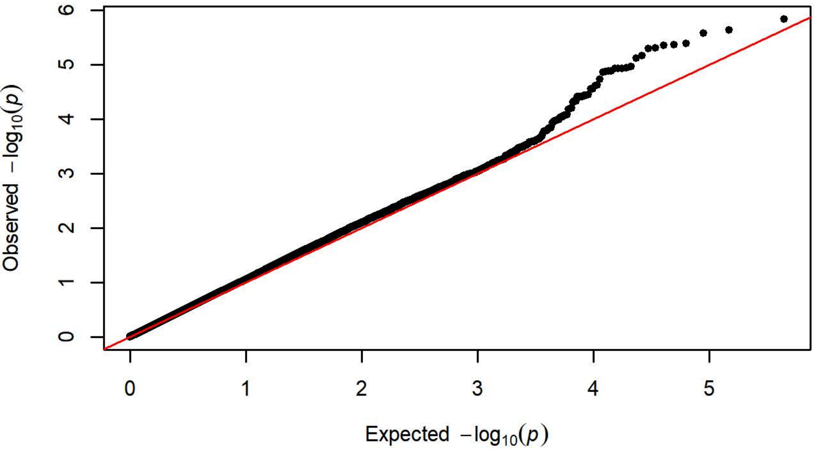

**Q-Q Plot Probability Difference Day 5 - Charles River R09-P3/7/10 (n=4[illegible]**

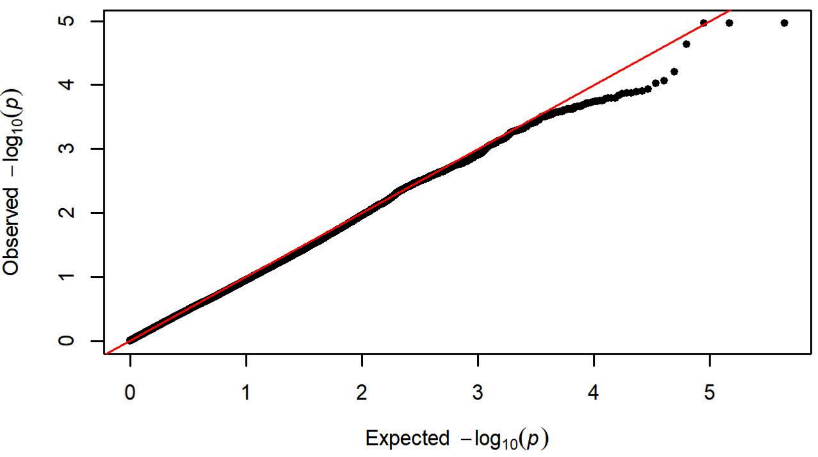


**Q-Q Plot Probability Difference Day 5 - Harlan 202A/C-208A (n=1099)**

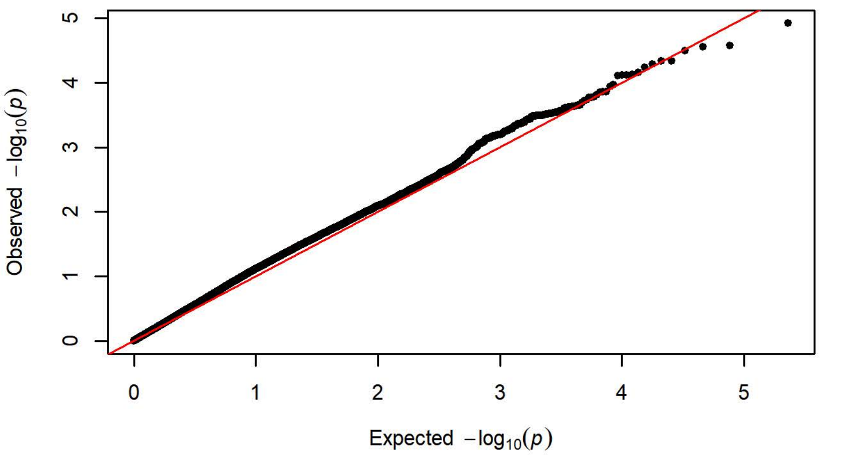


**Q-Q Plot Probability Difference Day 5 - Charles River R04 (n=650)**

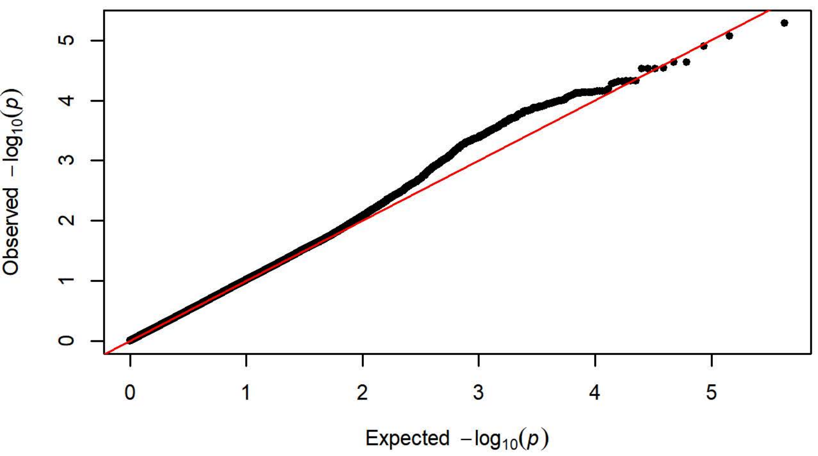


**Q-Q Plot Probability Difference Day 5 - Harlan 206 (n=758)**

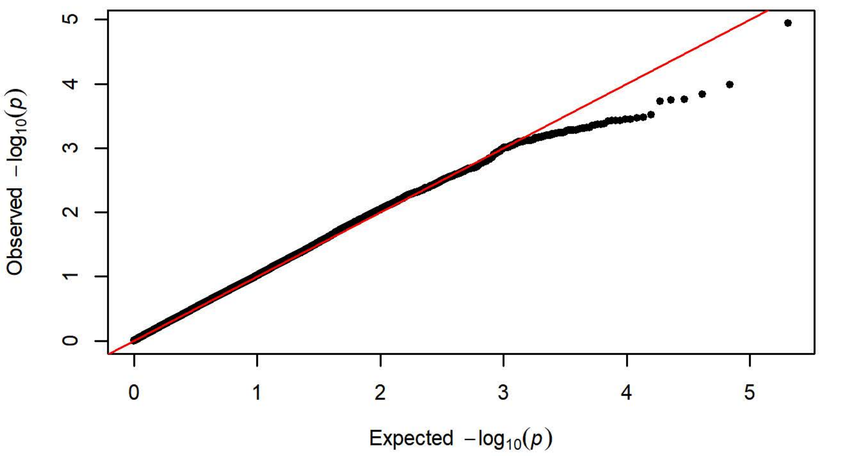


**Q-Q Plot Probability Difference Day 5 - Charles River P09 (n=295)**

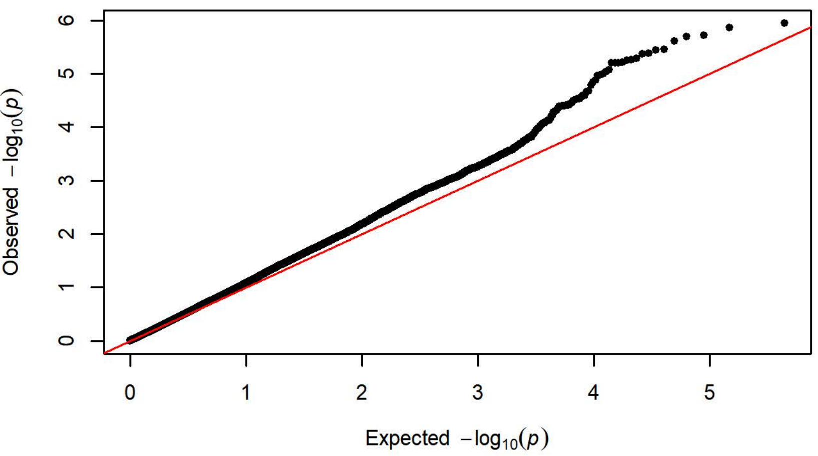


**Q-Q Plot Probability Difference Day 5 - Harlan 217 (n=351)**

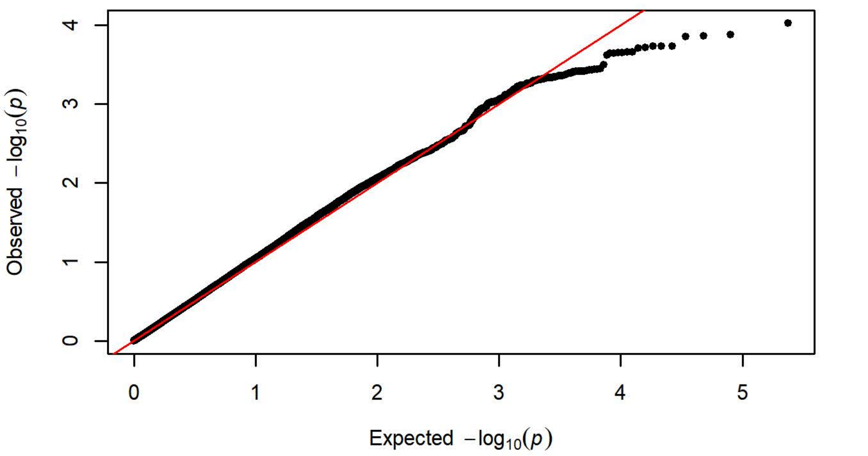


**Q-Q Plot Probability Difference Day 5 - Charles River C72 (n=358)**

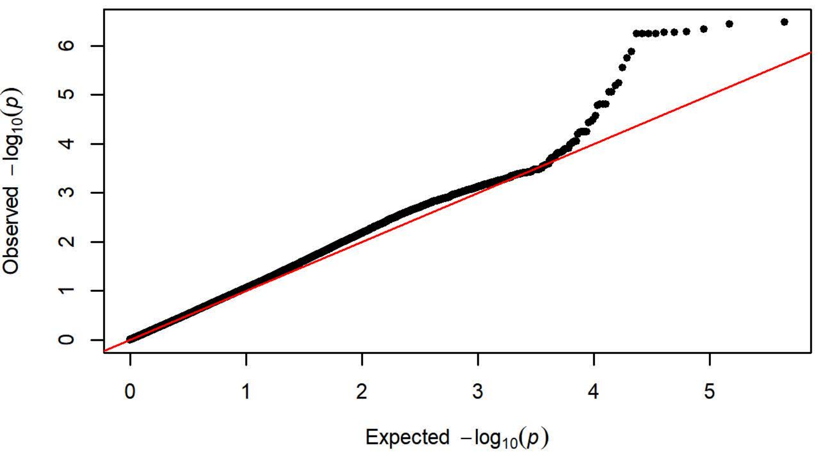

Q-Q Plot Probability of Lever Press Day 1 - Charles River R09-P3/7/10 (n=

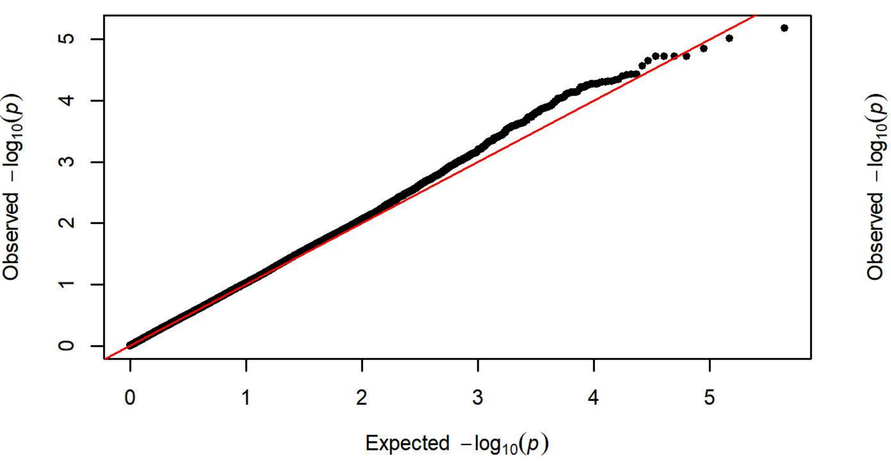


Q-Q Plot Probability of Lever Press Day 1 - Harlan 202A/C-208A (n=10

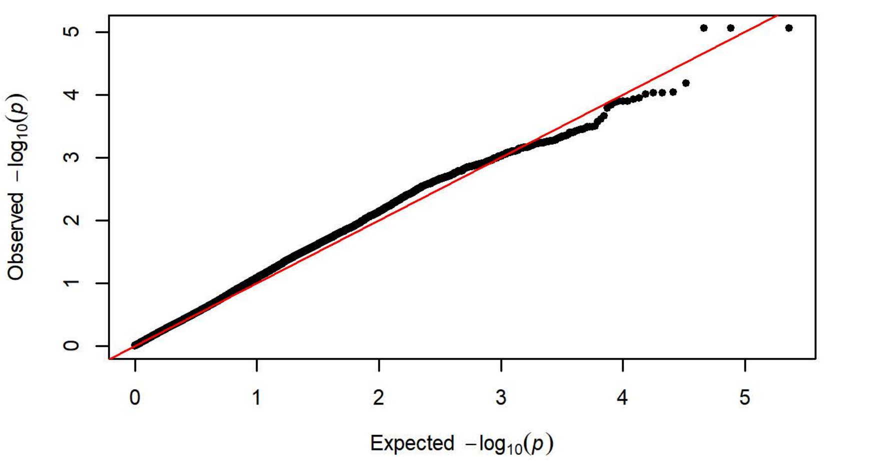


Q-Q Plot Probability of Lever Press Day 1 - Charles River R04 (n=650

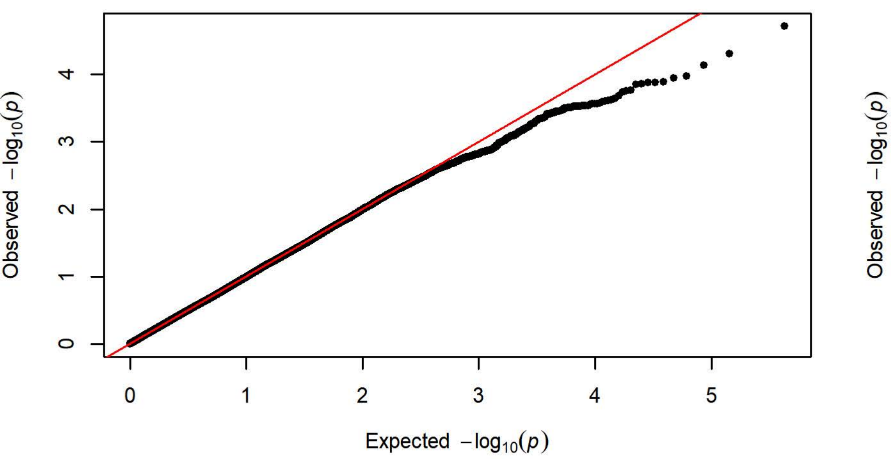


Q-Q Plot Probability of Lever Press Day 1 - Harlan 206 (n=758)

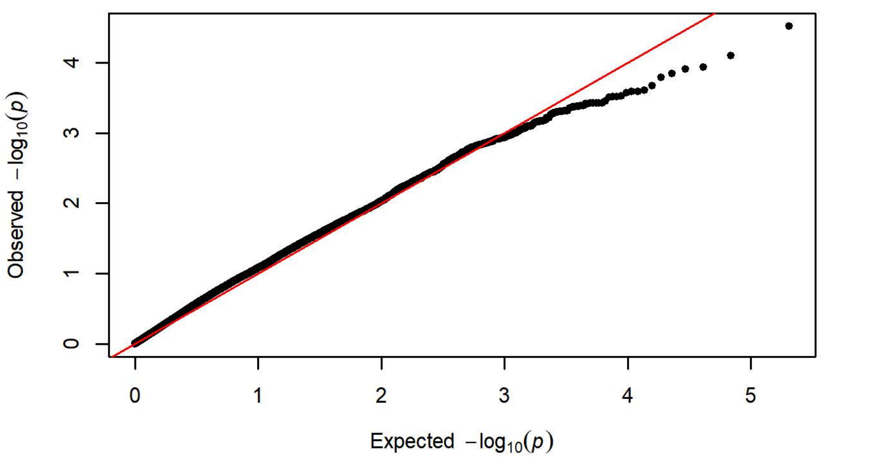


Q-Q Plot Probability of Lever Press Day 1 - Charles River P09 (n=295

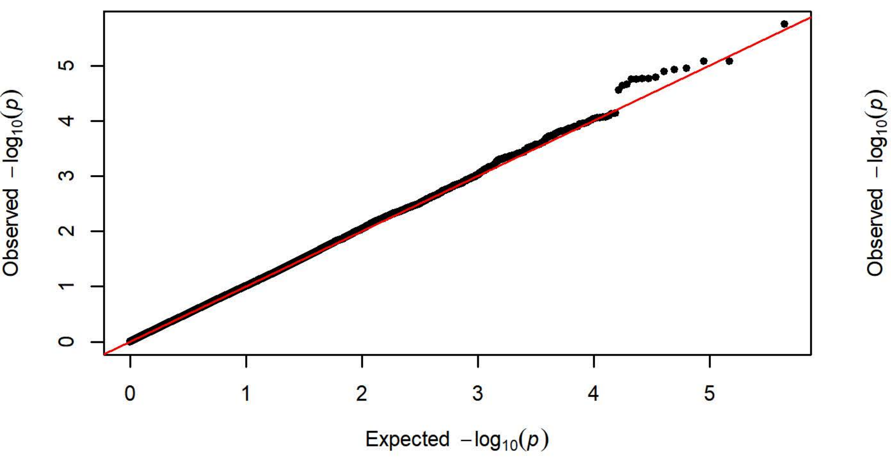


Q-Q Plot Probability of Lever Press Day 1 - Harlan 217 (n=351)

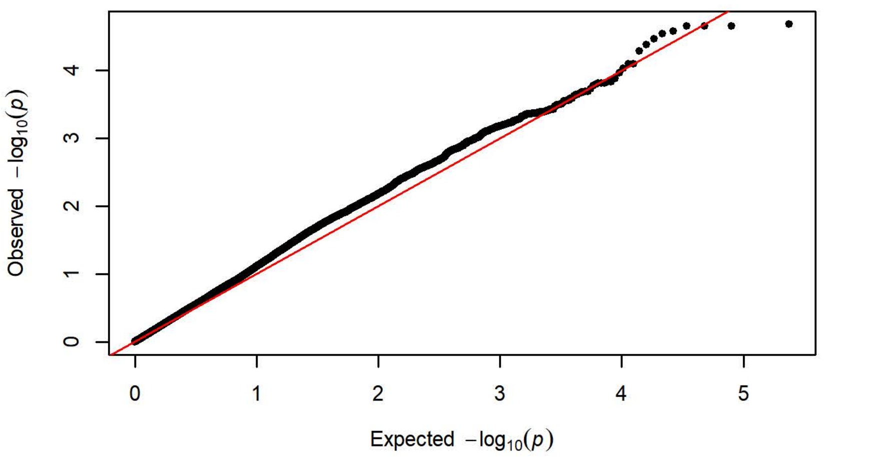


Q-Q Plot Probability of Lever Press Day 1 - Charles River C72 (n=358

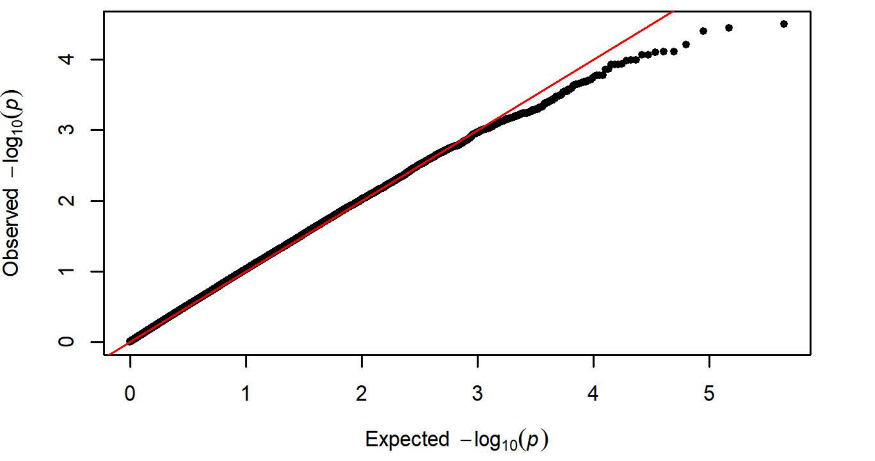

Q-Q Plot Probability of Lever Press Day 2 - Charles River R09-P3/7/10 (n=

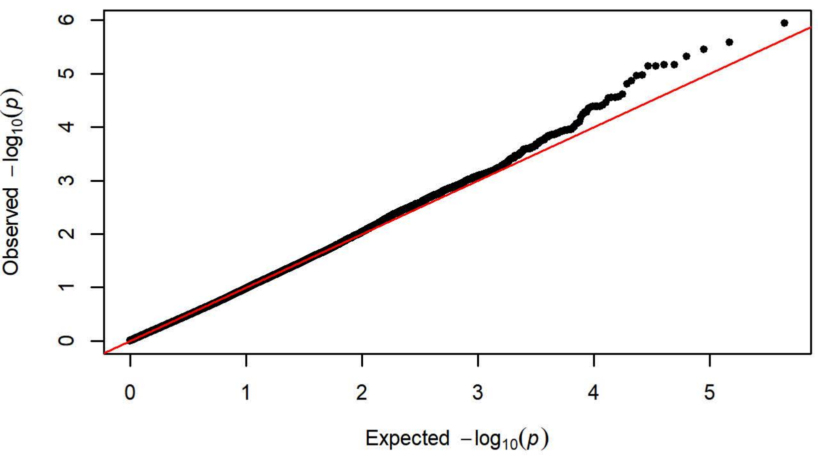


Q-Q Plot Probability of Lever Press Day 2 - Harlan 202A/C-208A (n=109

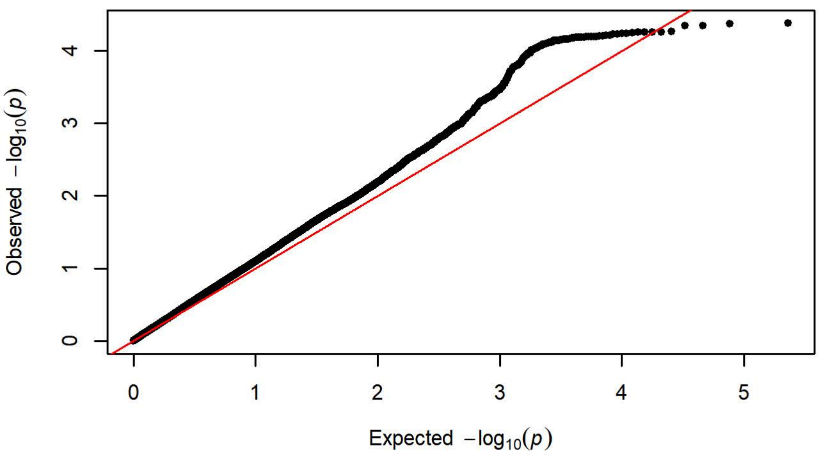


Q-Q Plot Probability of Lever Press Day 2 - Charles River R04 (n=650

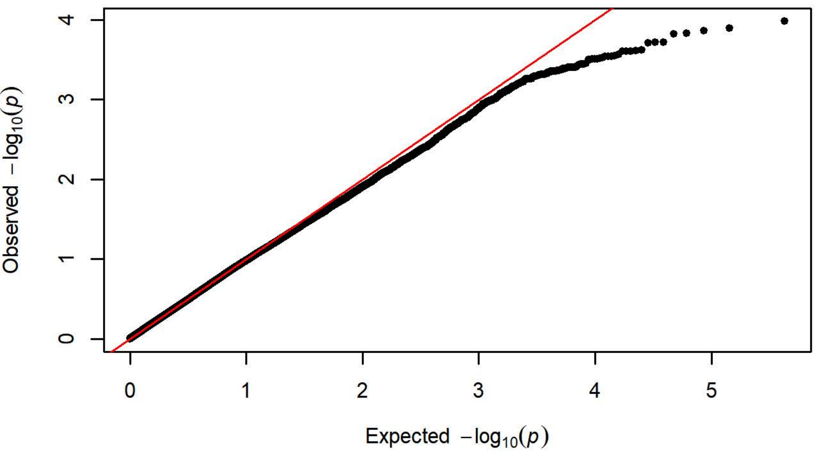


Q-Q Plot Probability of Lever Press Day 2 - Harlan 206 (n=758)

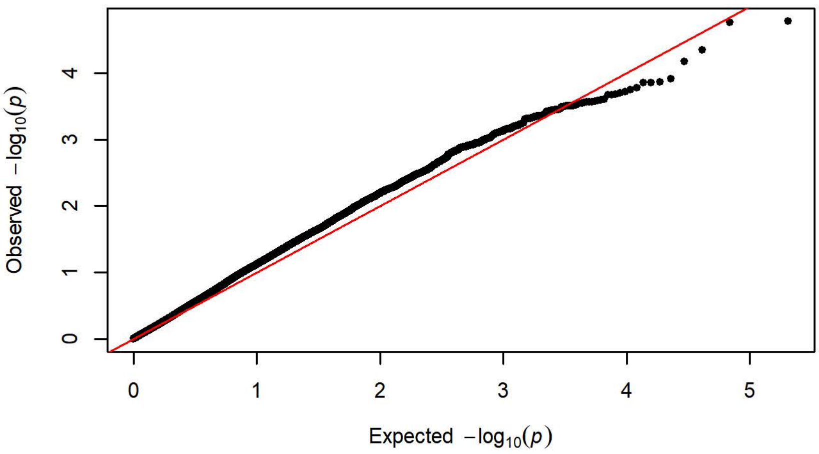


Q-Q Plot Probability of Lever Press Day 2 - Charles River P09 (n=295

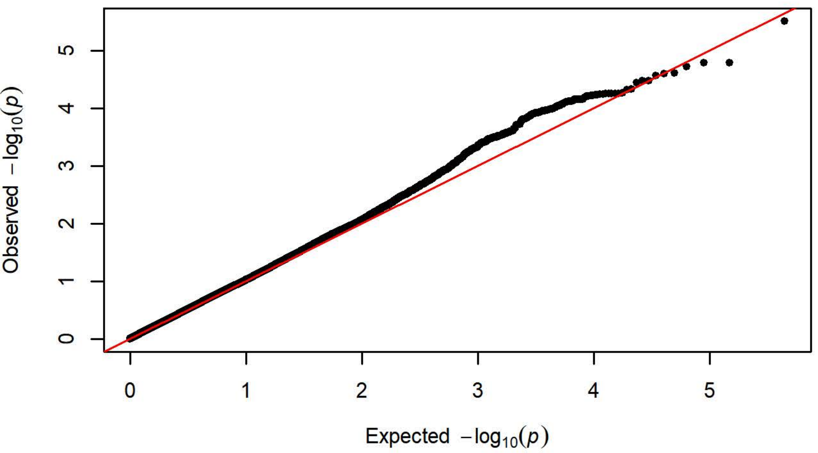


Q-Q Plot Probability of Lever Press Day 2 - Harlan 217 (n=351)

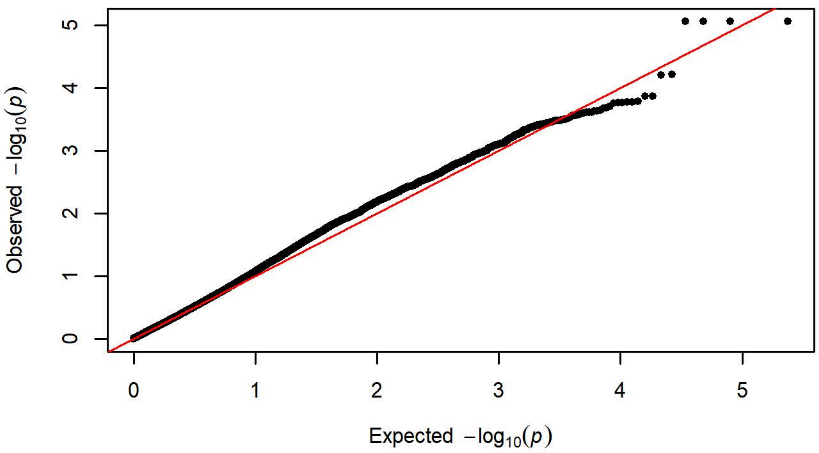


Q-Q Plot Probability of Lever Press Day 2 - Charles River C72 (n=356

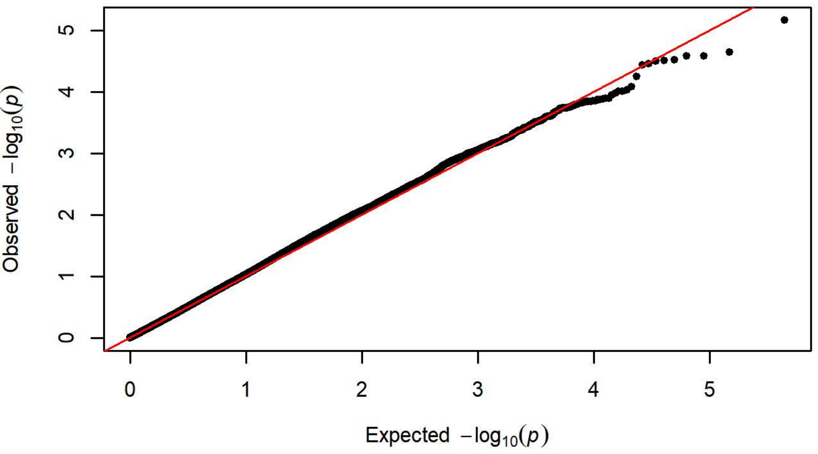

Q-Q Plot Probability of Lever Press Day 3 - Charles River R09-P3/7/10 (n=

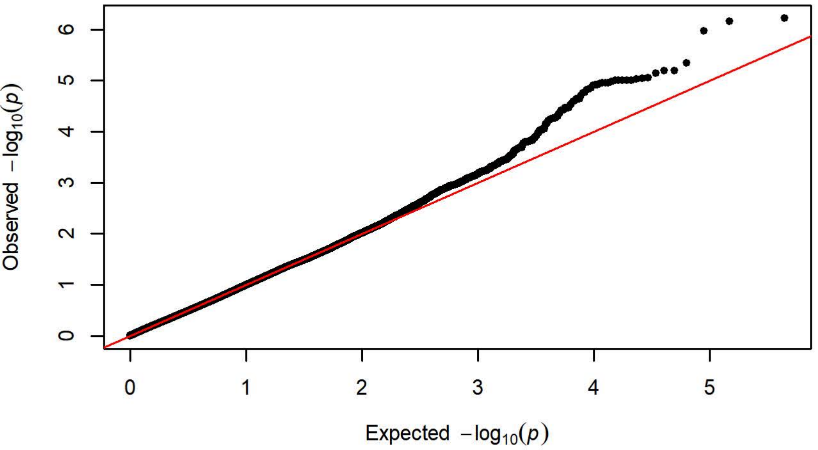


Q-Q Plot Probability of Lever Press Day 3 - Harlan 202A/C-208A (n=109

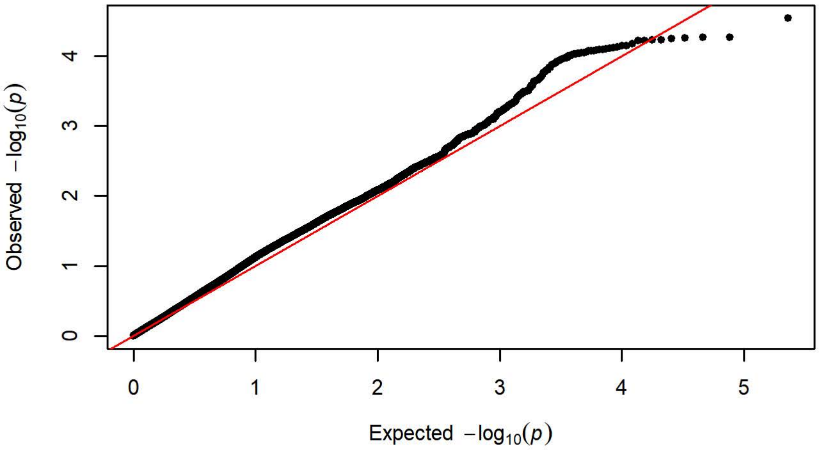


Q-Q Plot Probability of Lever Press Day 3 - Charles River R04 (n=650

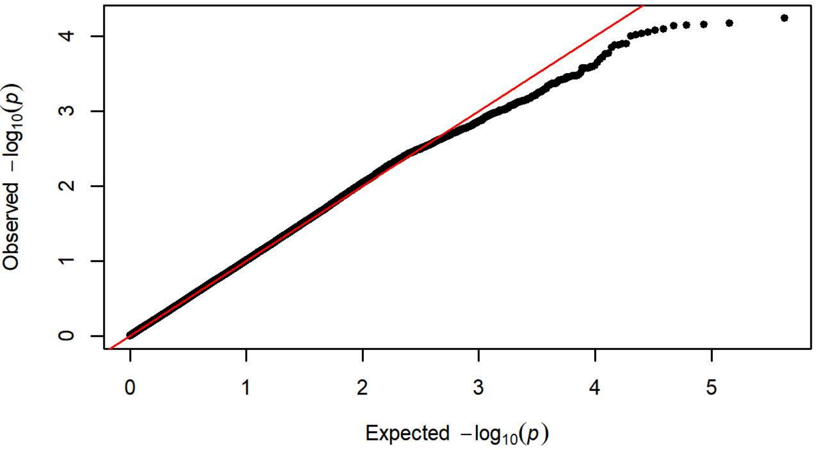


Q-Q Plot Probability of Lever Press Day 3 - Harlan 206 (n=758)

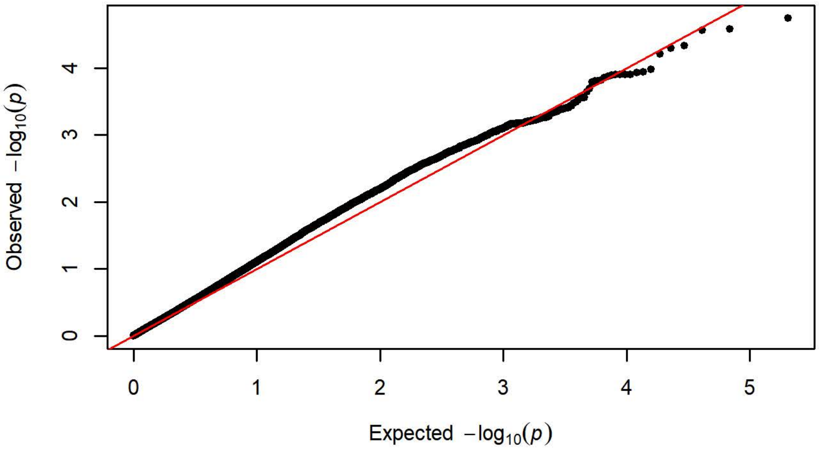


Q-Q Plot Probability of Lever Press Day 3 - Charles River P09 (n=295

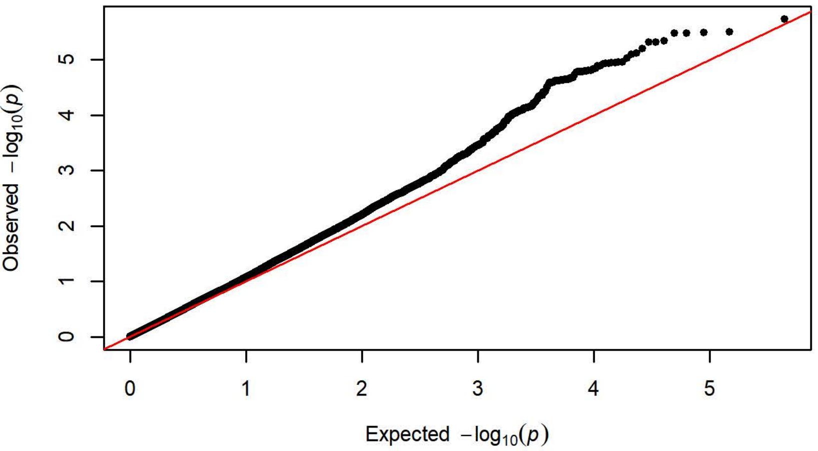


Q-Q Plot Probability of Lever Press Day 3 - Harlan 217 (n=351)

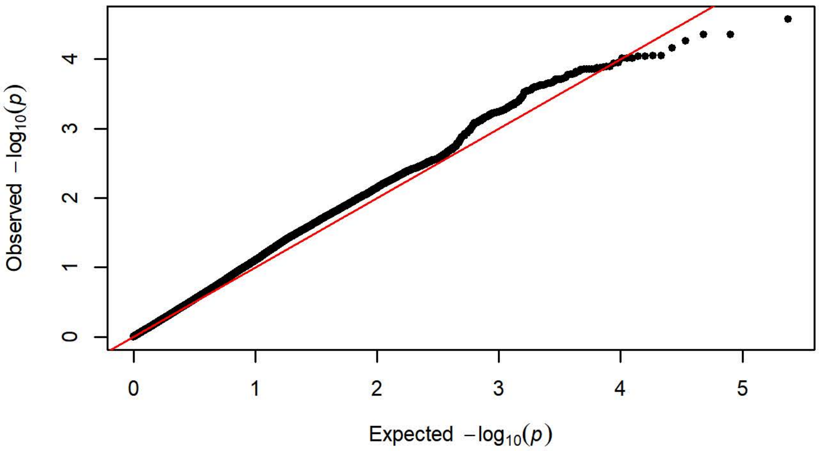


Q-Q Plot Probability of Lever Press Day 3 - Charles River C72 (n=358

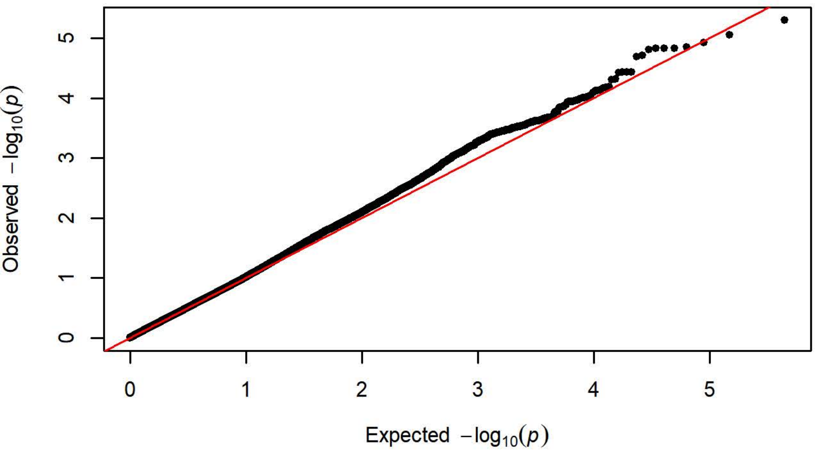

Q-Q Plot Probability of Lever Press Day 4 - Charles River R09-P3/7/10 (n=

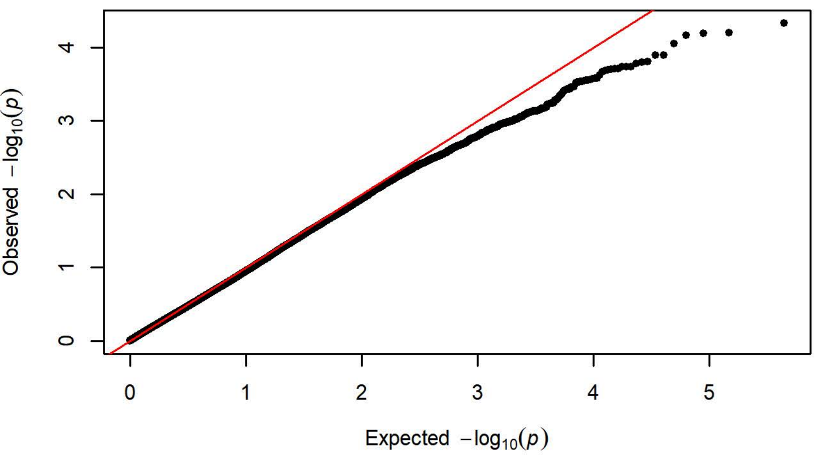


Q-Q Plot Probability of Lever Press Day 4 - Harlan 202A/C-208A (n=109

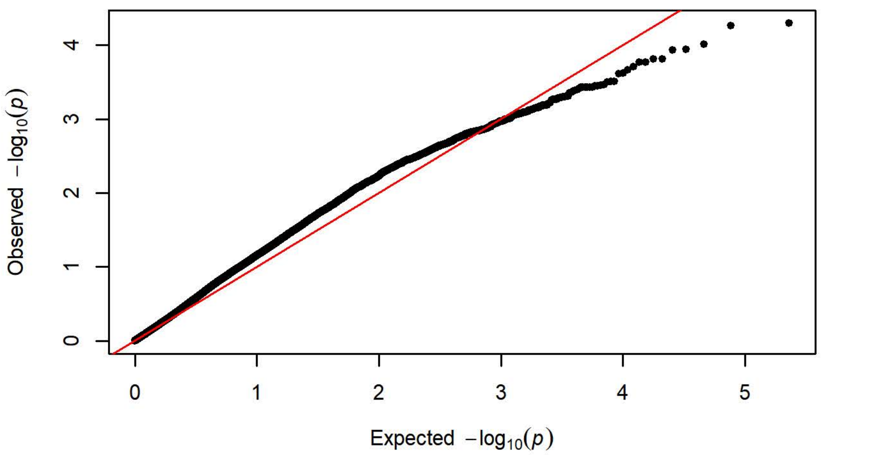


Q-Q Plot Probability of Lever Press Day 4 - Charles River R04 (n=650

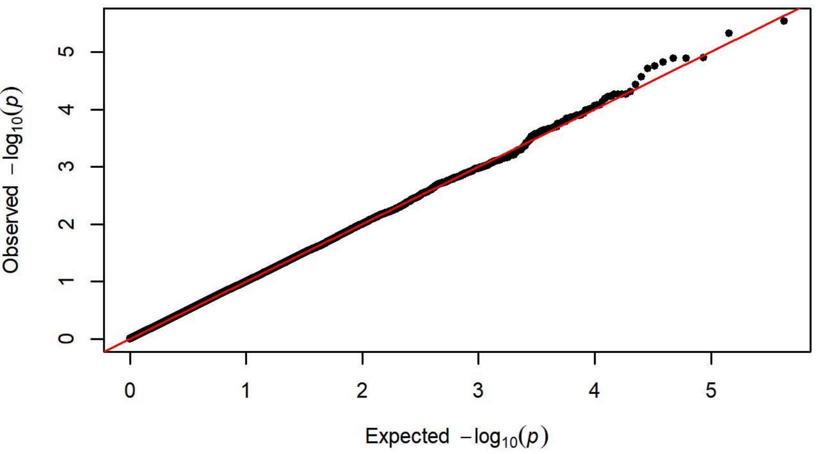


Q-Q Plot Probability of Lever Press Day 4 - Harlan 206 (n=758)

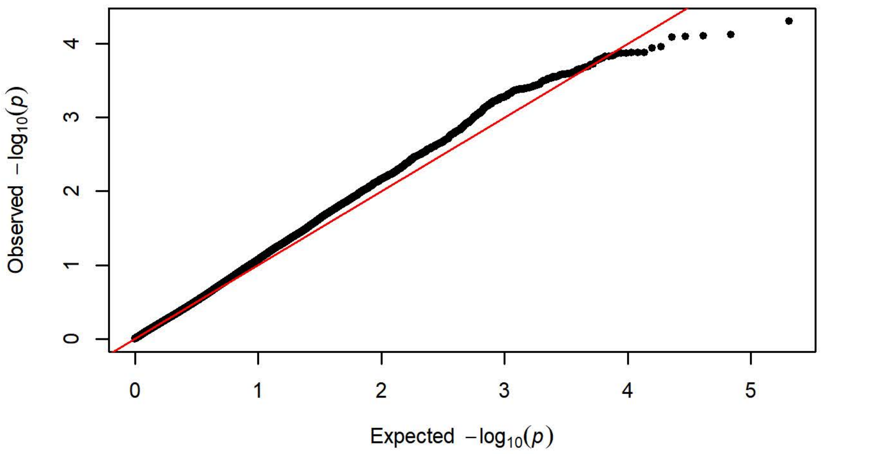


Q-Q Plot Probability of Lever Press Day 4 - Charles River P09 (n=295

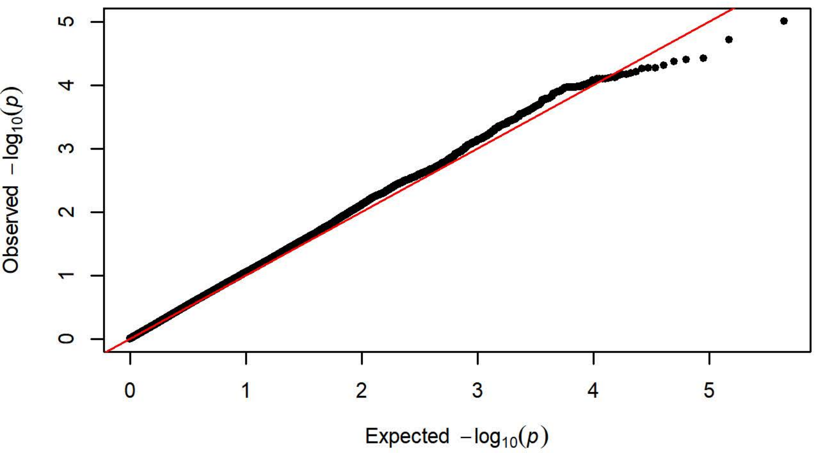


Q-Q Plot Probability of Lever Press Day 4 - Harlan 217 (n=351)

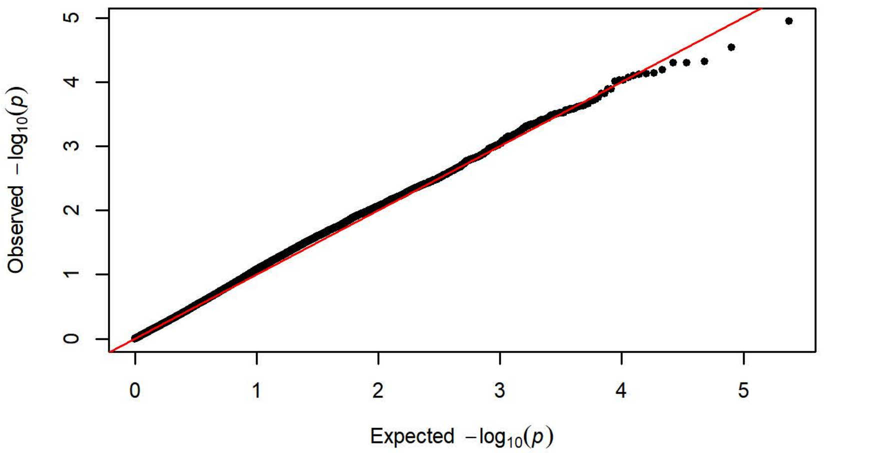


Q-Q Plot Probability of Lever Press Day 4 - Charles River C72 (n=358

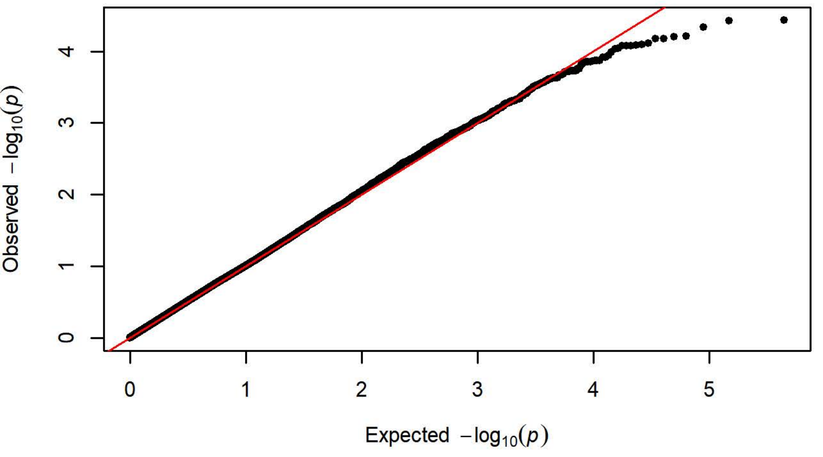

Q-Q Plot Probability of Lever Press Day 5 - Charles River R09-P3/7/10 (n=

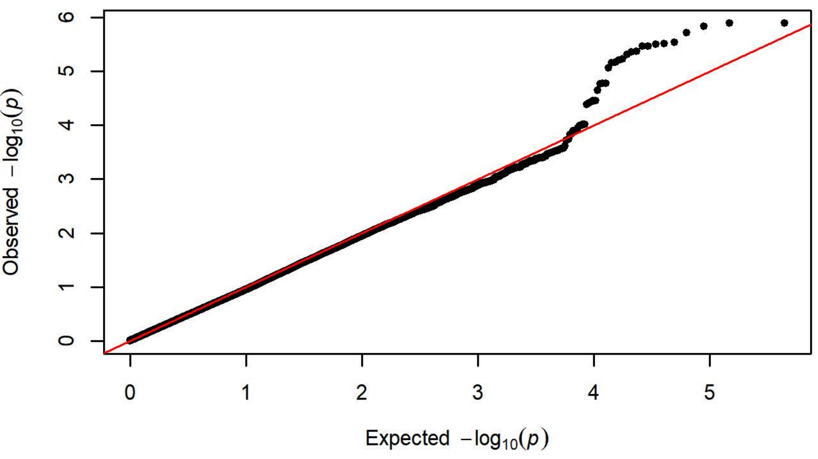


Q-Q Plot Probability of Lever Press Day 5 - Harlan 202A/C-208A (n=109

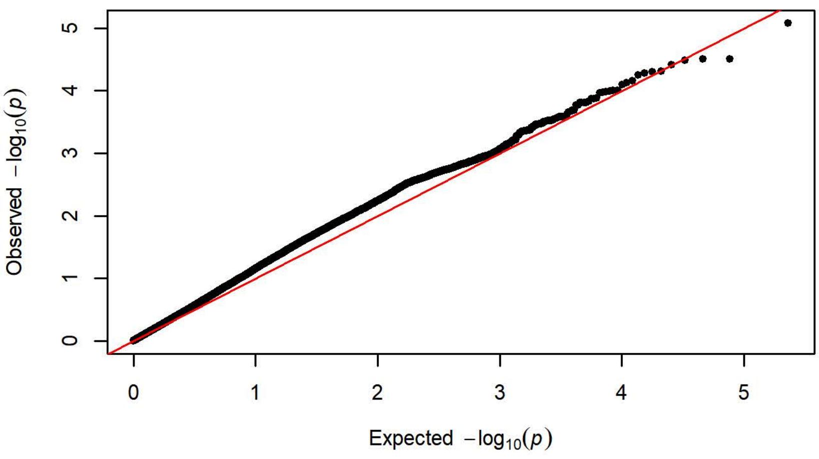


Q-Q Plot Probability of Lever Press Day 5 - Charles River R04 (n=650

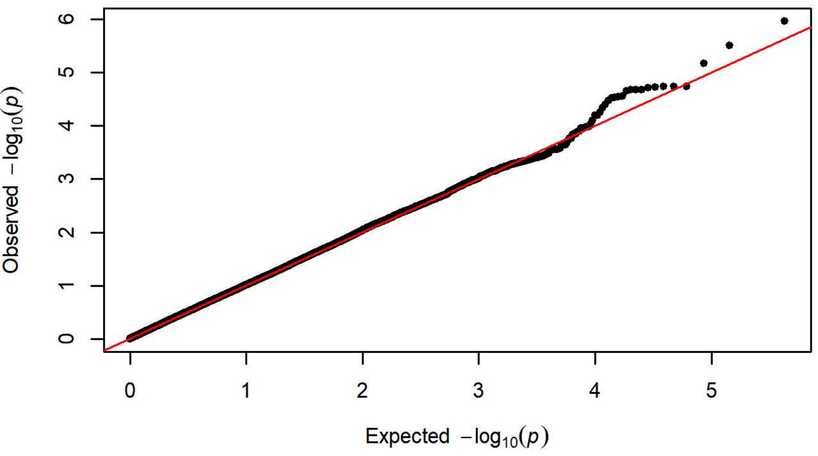


Q-Q Plot Probability of Lever Press Day 5 - Harlan 206 (n=758)

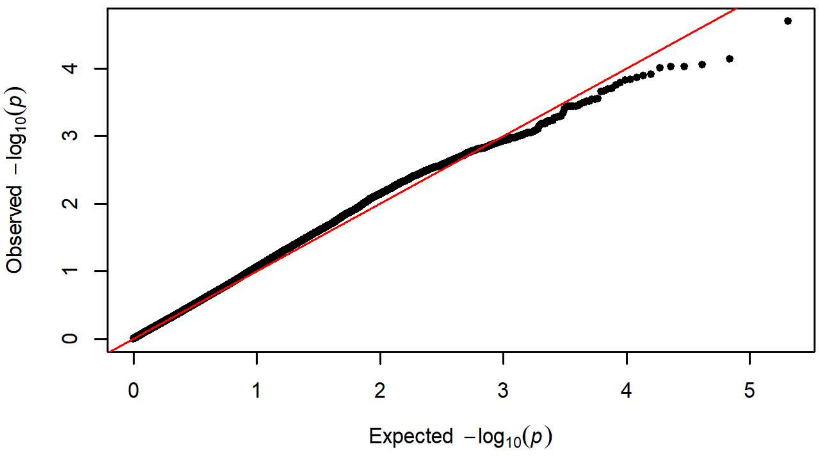


Q-Q Plot Probability of Lever Press Day 5 - Charles River P09 (n=295

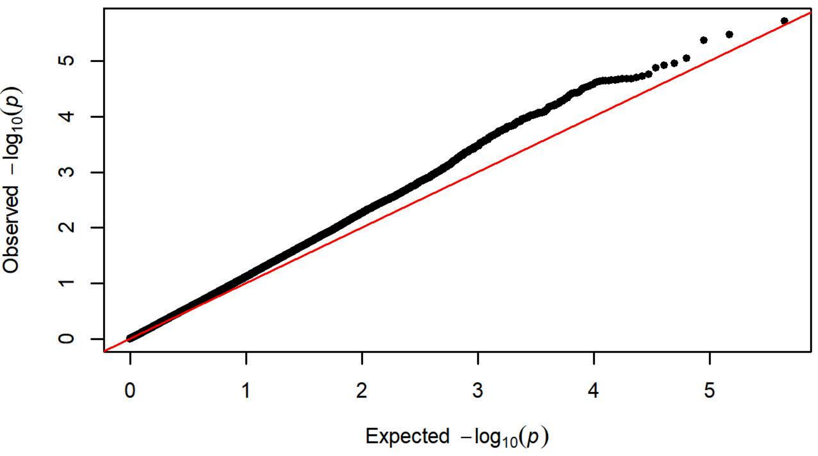


Q-Q Plot Probability of Lever Press Day 5 - Harlan 217 (n=351)

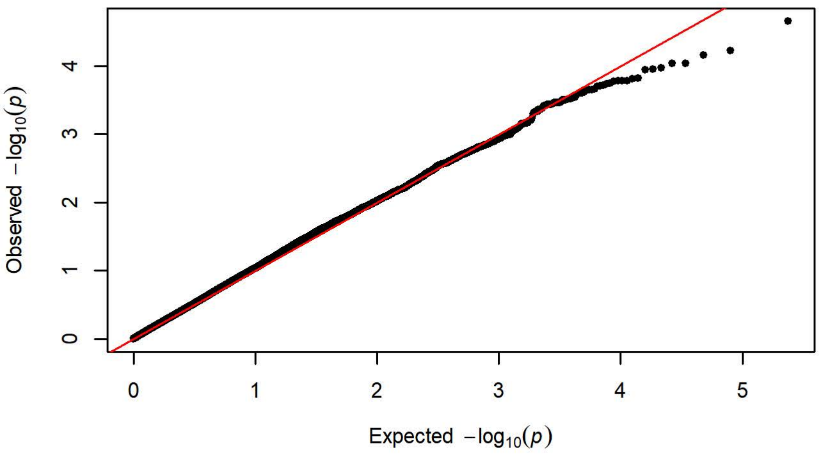


Q-Q Plot Probability of Lever Press Day 5 - Charles River C72 (n=358

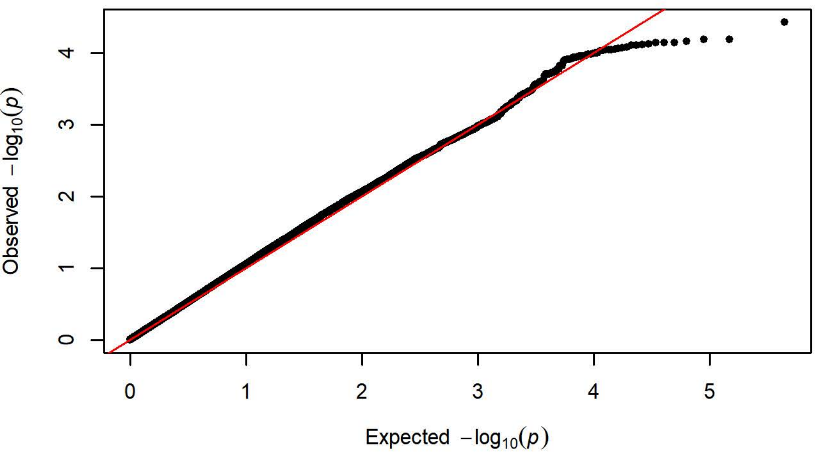

Q Plot Probability of Magazine Entry Day 1 - Charles River R09-P3/7/10 (

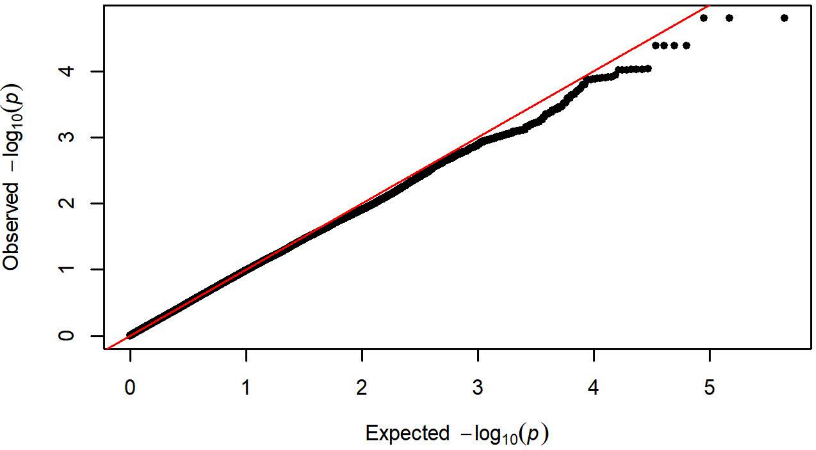


Q-Q Plot Probability of Magazine Entry Day 1 - Harlan 202A/C-208A (n=1

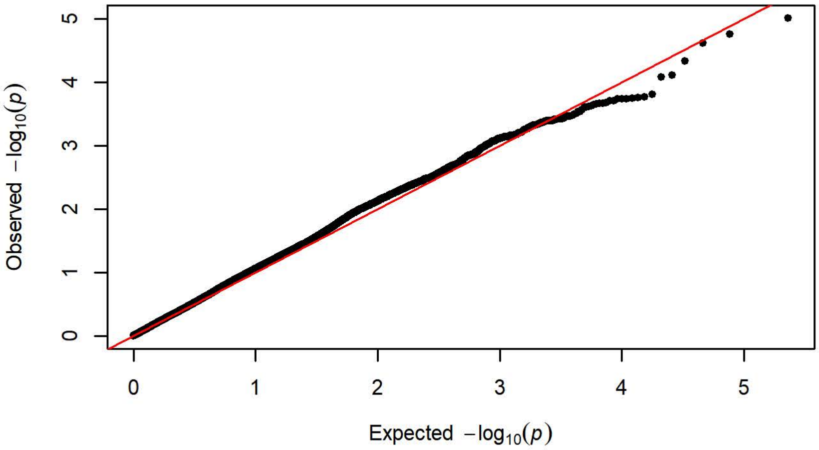


Q-Q Plot Probability of Magazine Entry Day 1 - Charles River R04 (n=6

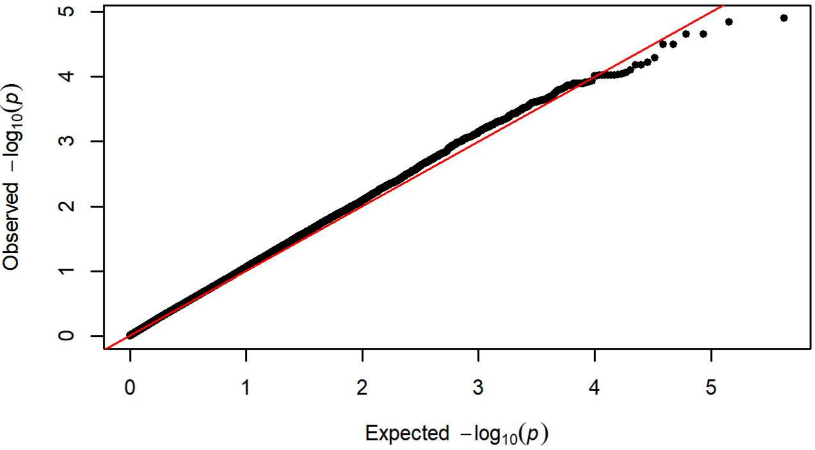


Q-Q Plot Probability of Magazine Entry Day 1 - Harlan 206 (n=758)

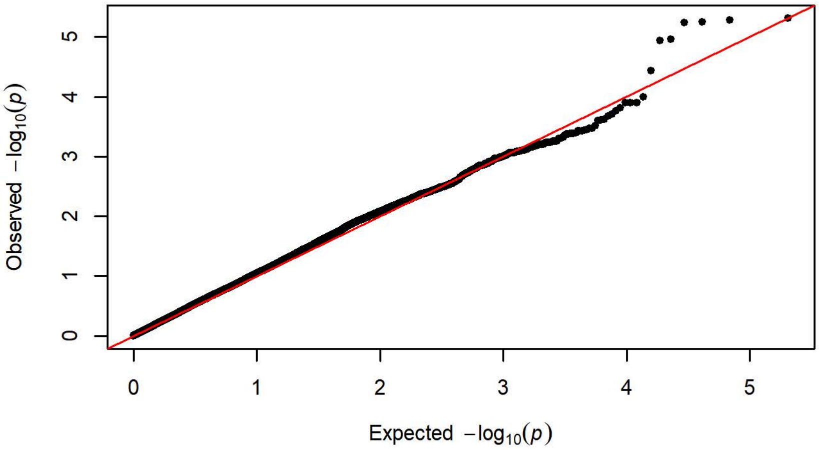


Q-Q Plot Probability of Magazine Entry Day 1 - Charles River P09 (n=2

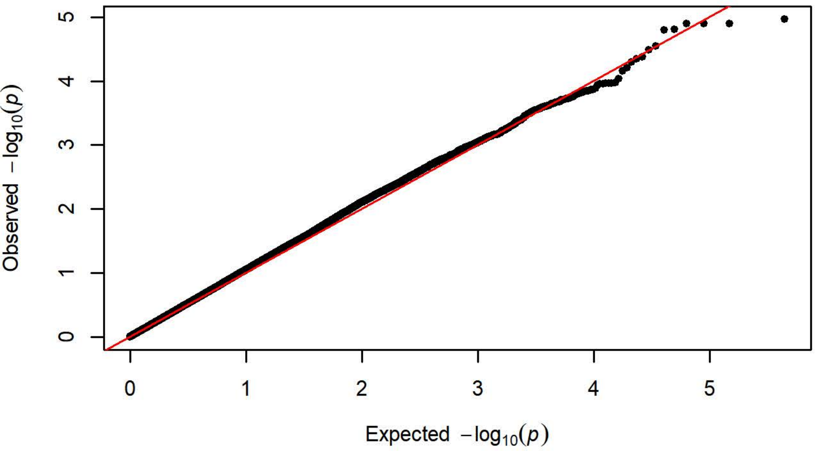


Q-Q Plot Probability of Magazine Entry Day 1 - Harlan 217 (n=351)

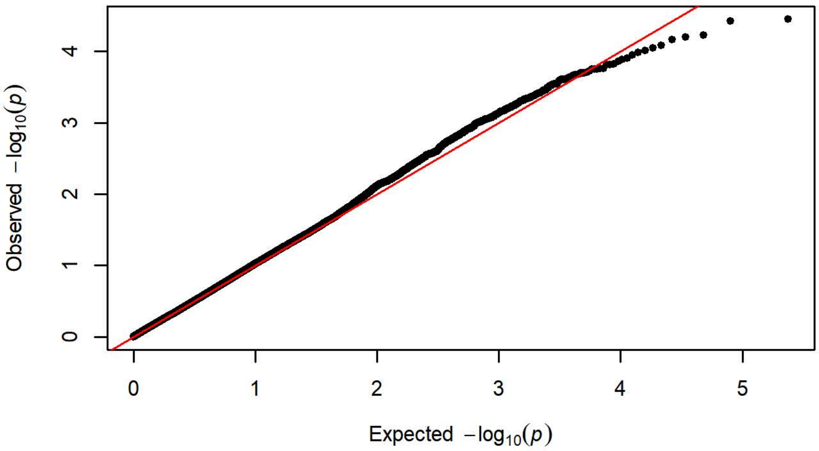


Q-Q Plot Probability of Magazine Entry Day 1 - Charles River C72 (n=3

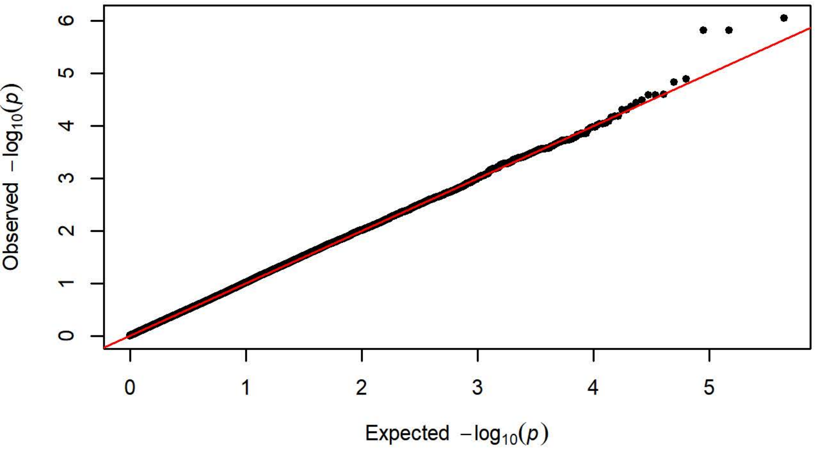

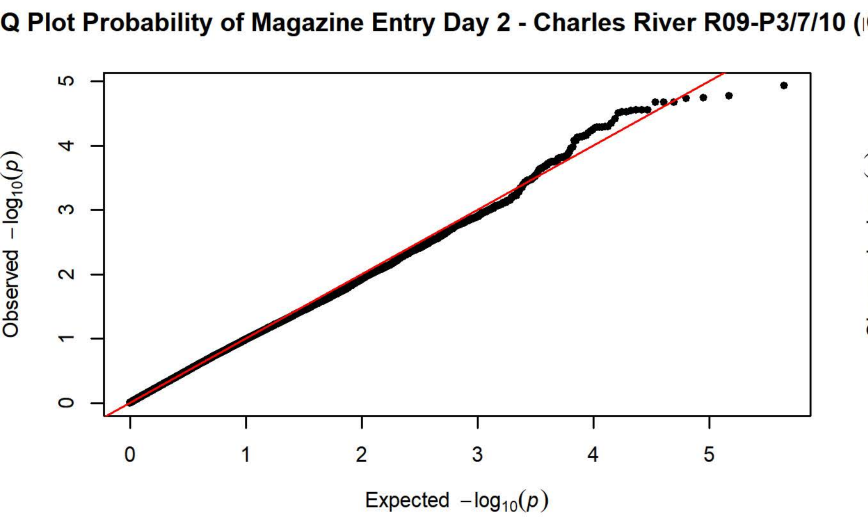
Q Plot Probability of Magazine Entry Day 2 - Charles River R09-P3/7/10 (
Observed $-\log_{10}(p)$
Expected $-\log_{10}(p)$

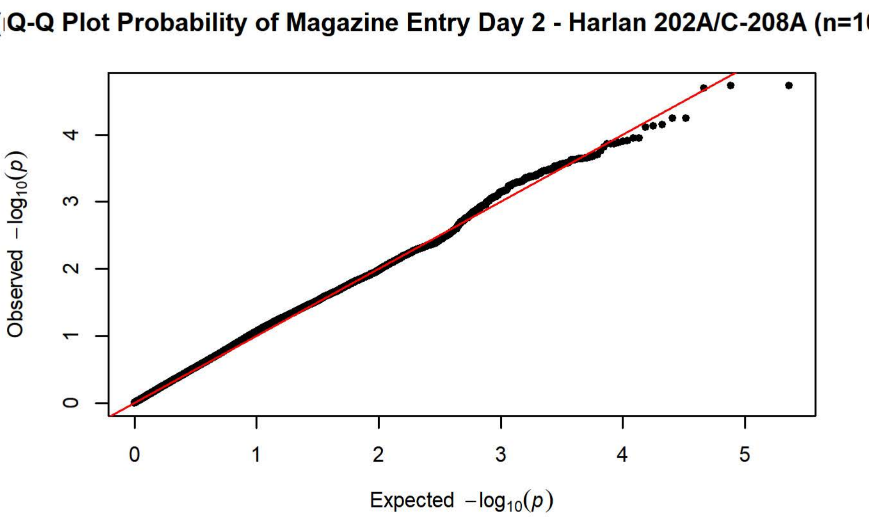
Q-Q Plot Probability of Magazine Entry Day 2 - Harlan 202A/C-208A (n=1
Observed $-\log_{10}(p)$
Expected $-\log_{10}(p)$

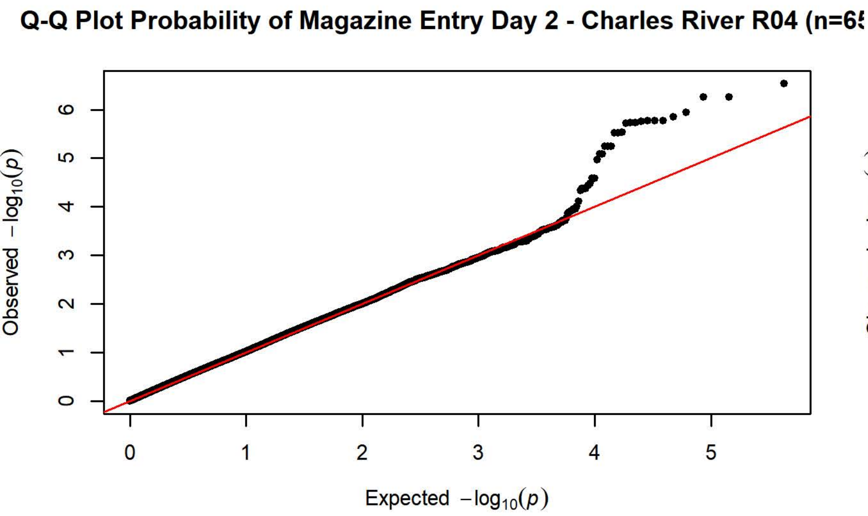
Q-Q Plot Probability of Magazine Entry Day 2 - Charles River R04 (n=6
Observed $-\log_{10}(p)$
Expected $-\log_{10}(p)$

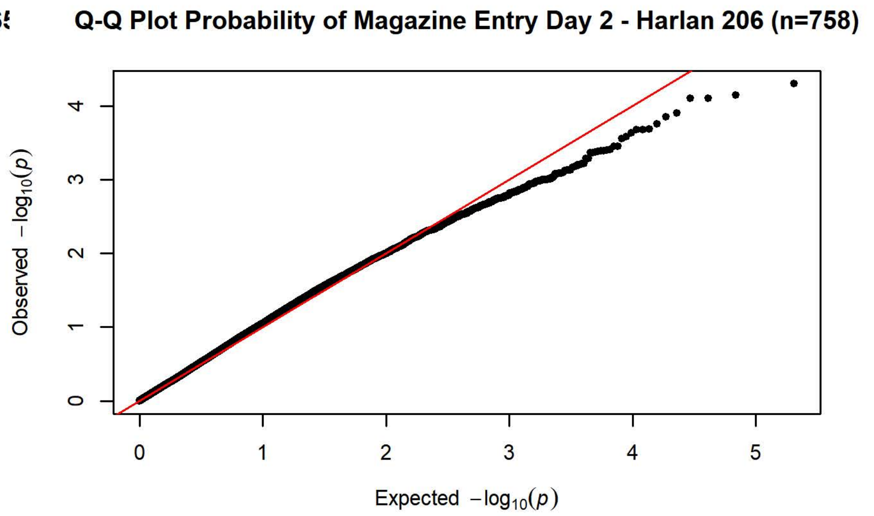
Q-Q Plot Probability of Magazine Entry Day 2 - Harlan 206 (n=758)
Observed $-\log_{10}(p)$
Expected $-\log_{10}(p)$

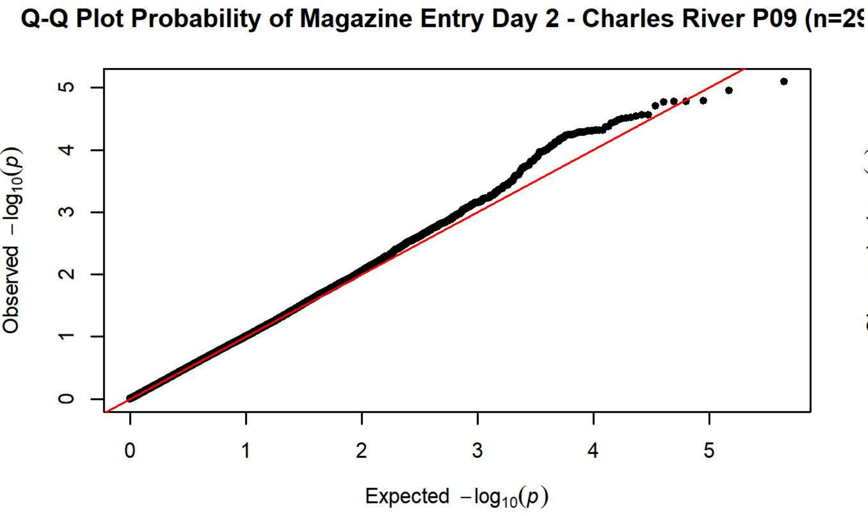
Q-Q Plot Probability of Magazine Entry Day 2 - Charles River P09 (n=2
Observed $-\log_{10}(p)$
Expected $-\log_{10}(p)$

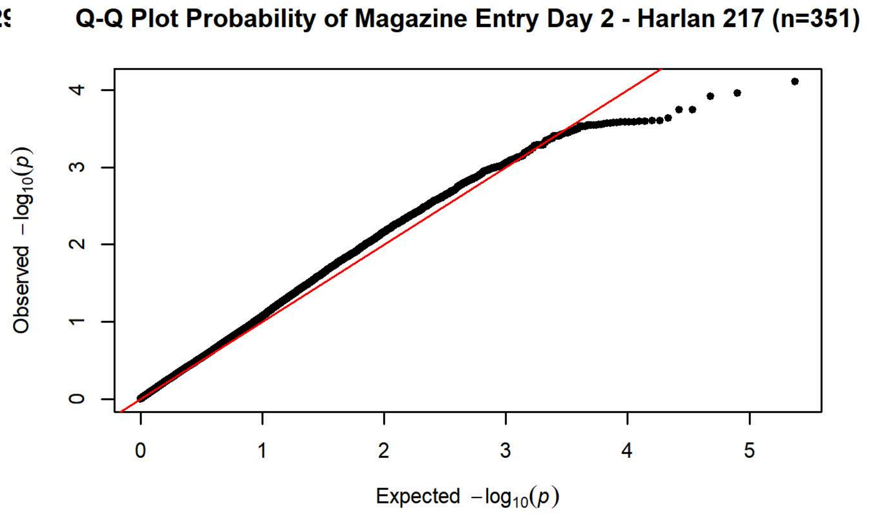
Q-Q Plot Probability of Magazine Entry Day 2 - Harlan 217 (n=351)
Observed $-\log_{10}(p)$
Expected $-\log_{10}(p)$

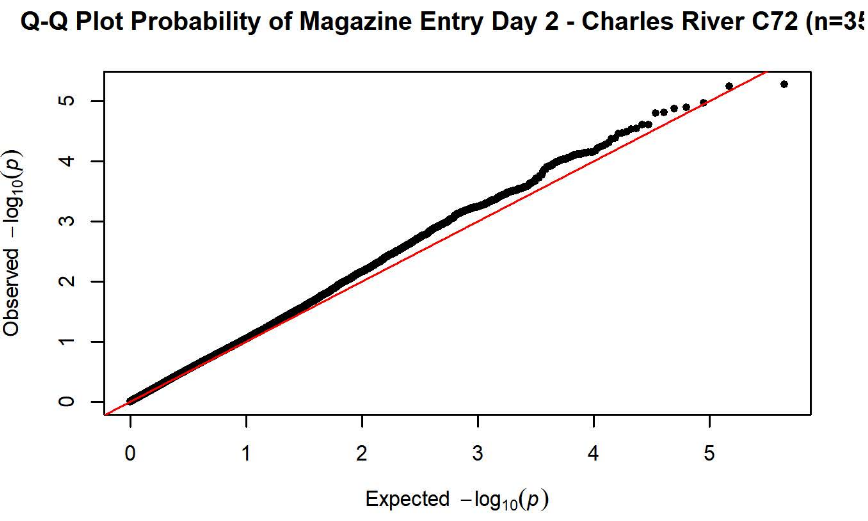
Q-Q Plot Probability of Magazine Entry Day 2 - Charles River C72 (n=3
Observed $-\log_{10}(p)$
Expected $-\log_{10}(p)$

Q Plot Probability of Magazine Entry Day 3 - Charles River R09-P3/7/10 (

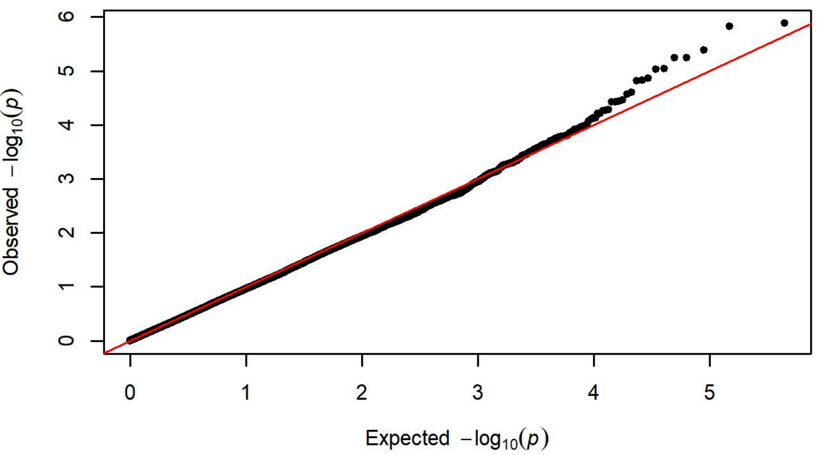


Q-Q Plot Probability of Magazine Entry Day 3 - Harlan 202A/C-208A (n=1

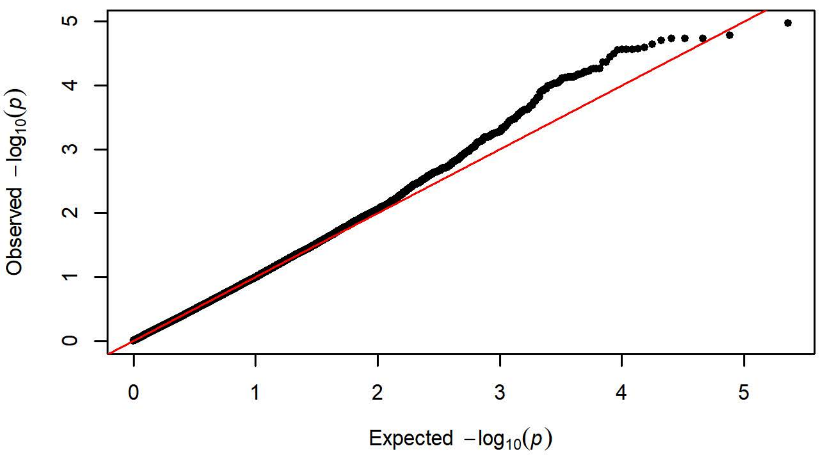


Q-Q Plot Probability of Magazine Entry Day 3 - Charles River R04 (n=6

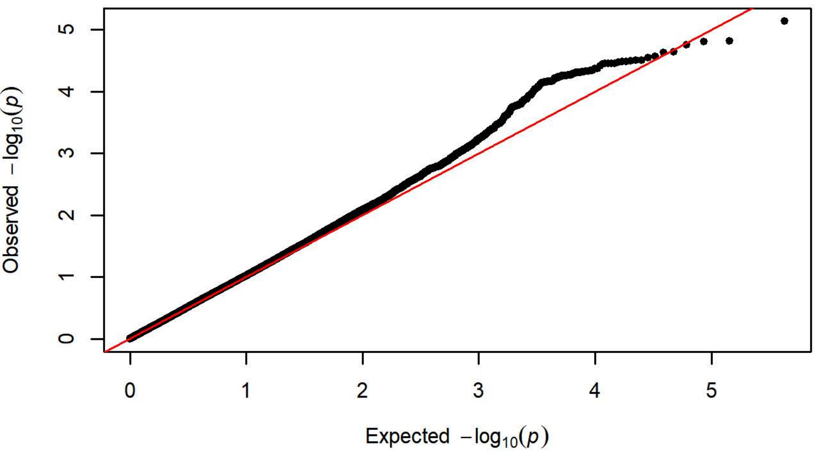


Q-Q Plot Probability of Magazine Entry Day 3 - Harlan 206 (n=758)

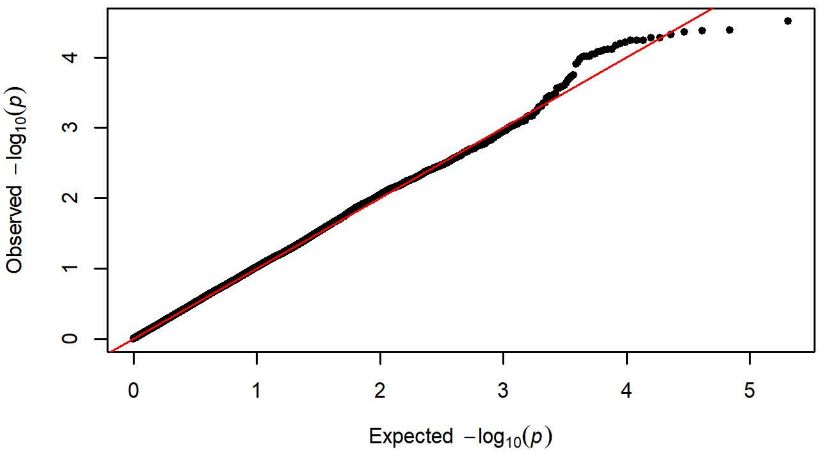


Q-Q Plot Probability of Magazine Entry Day 3 - Charles River P09 (n=2

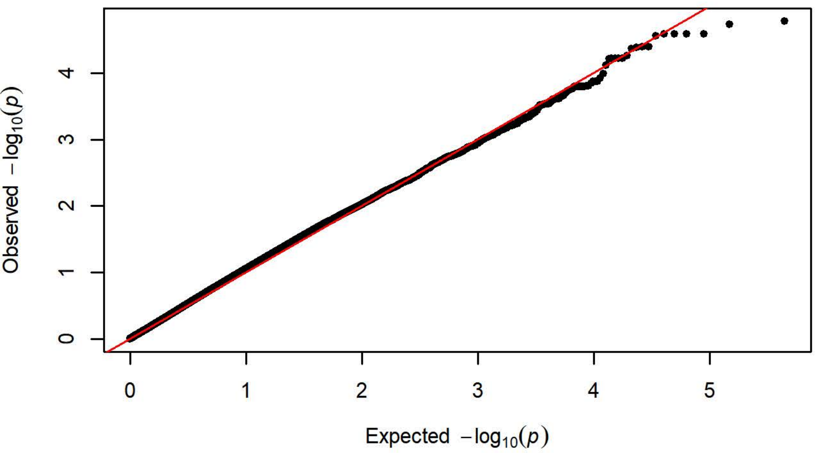


Q-Q Plot Probability of Magazine Entry Day 3 - Harlan 217 (n=351)

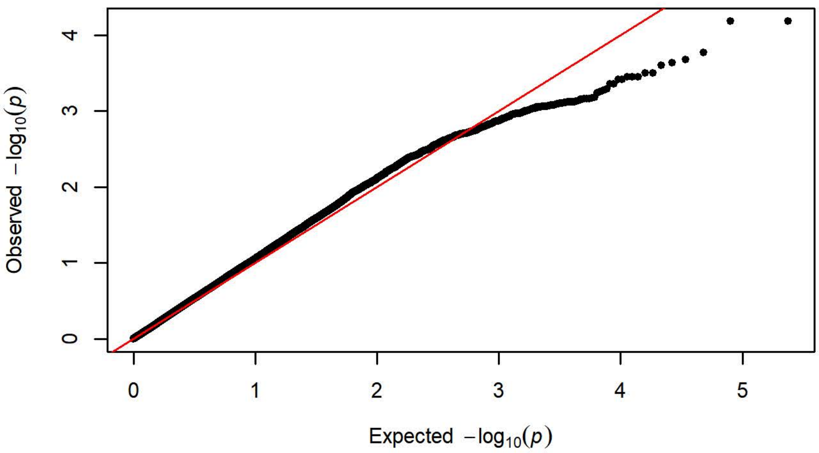


Q-Q Plot Probability of Magazine Entry Day 3 - Charles River C72 (n=3

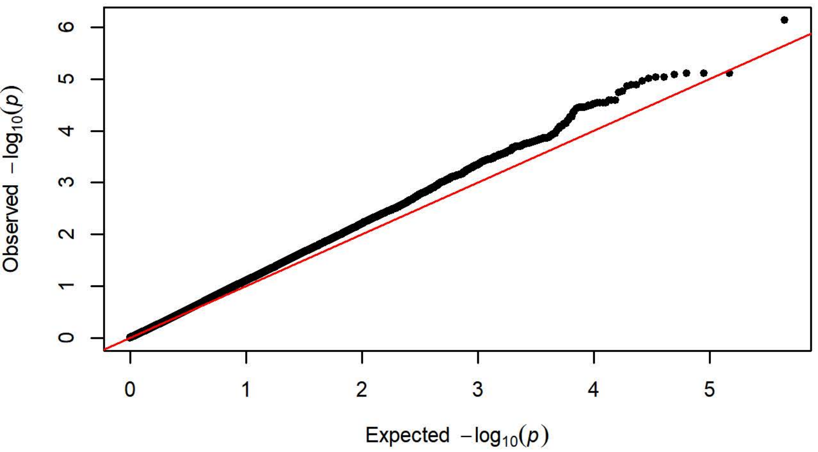

Q Plot Probability of Magazine Entry Day 4 - Charles River R09-P3/7/10 (

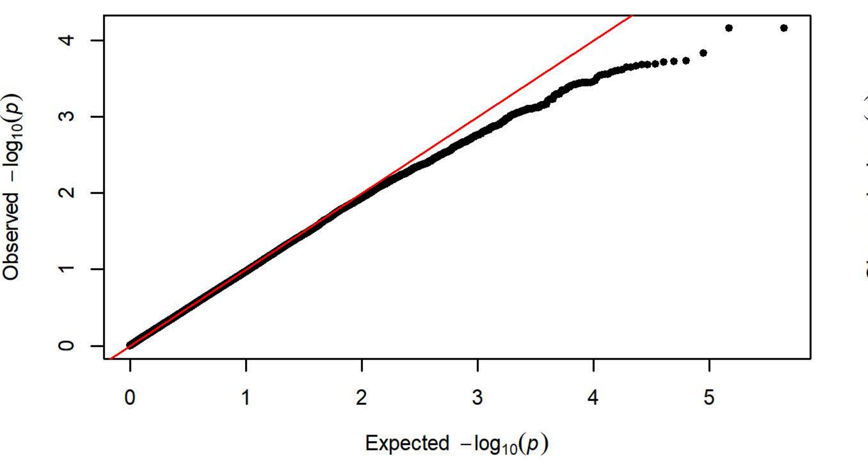


Q-Q Plot Probability of Magazine Entry Day 4 - Harlan 202A/C-208A (n=1

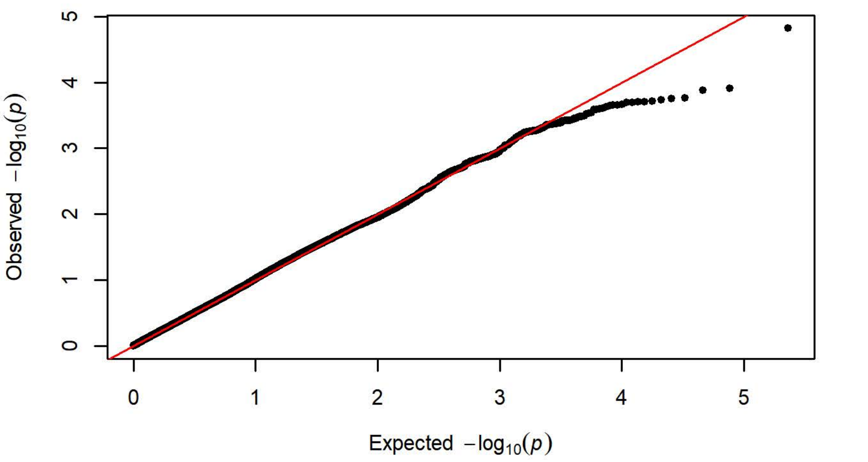


Q-Q Plot Probability of Magazine Entry Day 4 - Charles River R04 (n=6

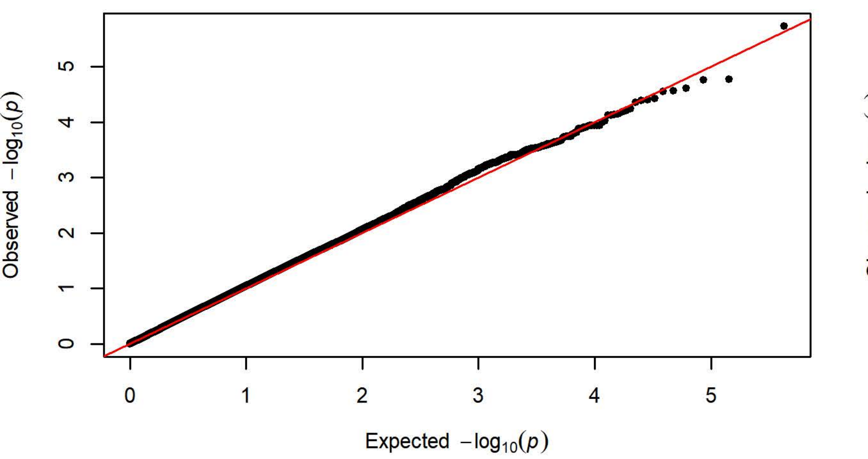


Q-Q Plot Probability of Magazine Entry Day 4 - Harlan 206 (n=758)

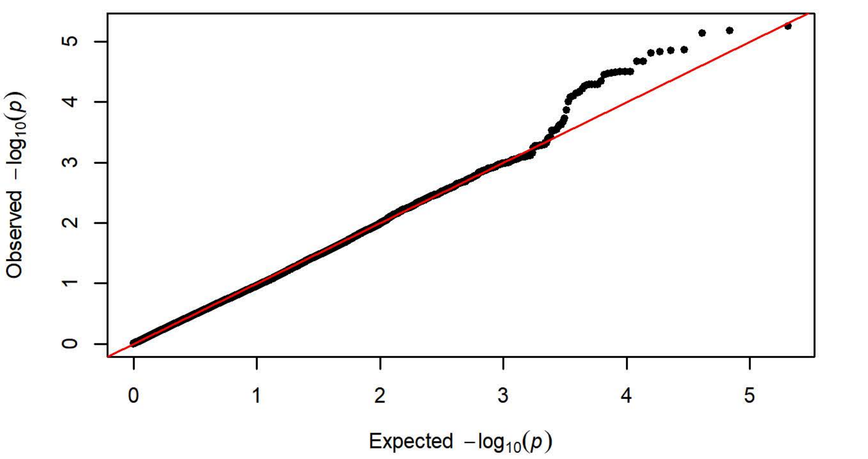


Q-Q Plot Probability of Magazine Entry Day 4 - Charles River P09 (n=2

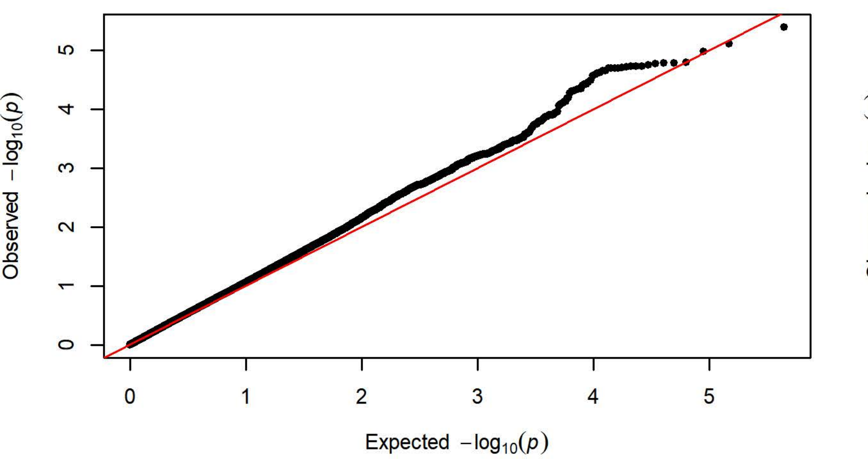


Q-Q Plot Probability of Magazine Entry Day 4 - Harlan 217 (n=351)

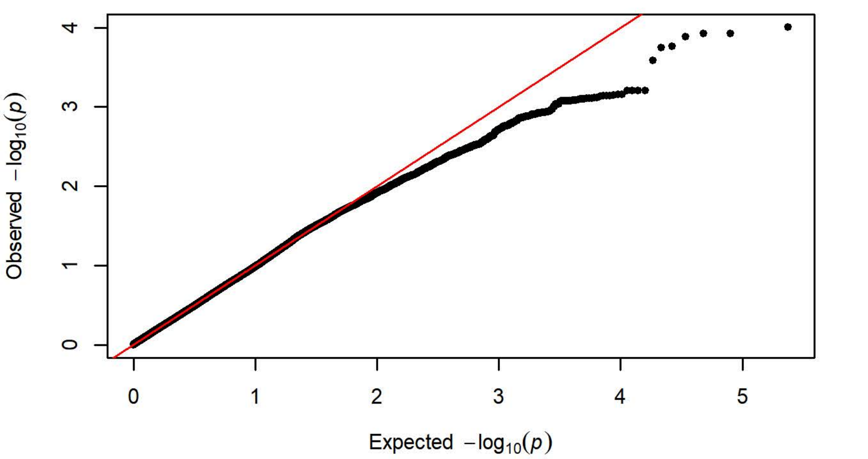


Q-Q Plot Probability of Magazine Entry Day 4 - Charles River C72 (n=3

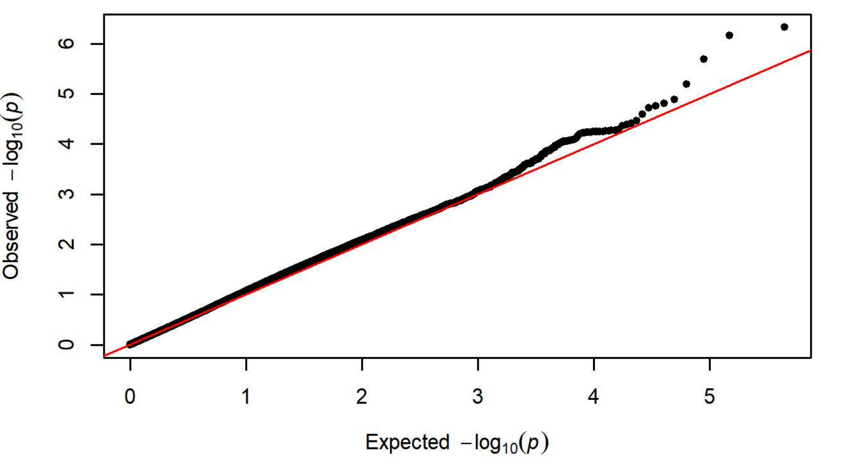

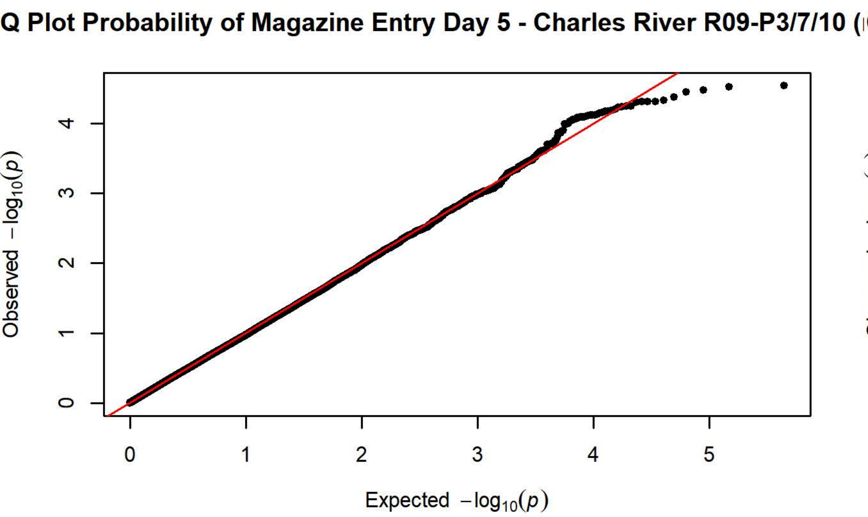
Q Plot Probability of Magazine Entry Day 5 - Charles River R09-P3/7/10 (
Observed $-\log_{10}(p)$
Expected $-\log_{10}(p)$

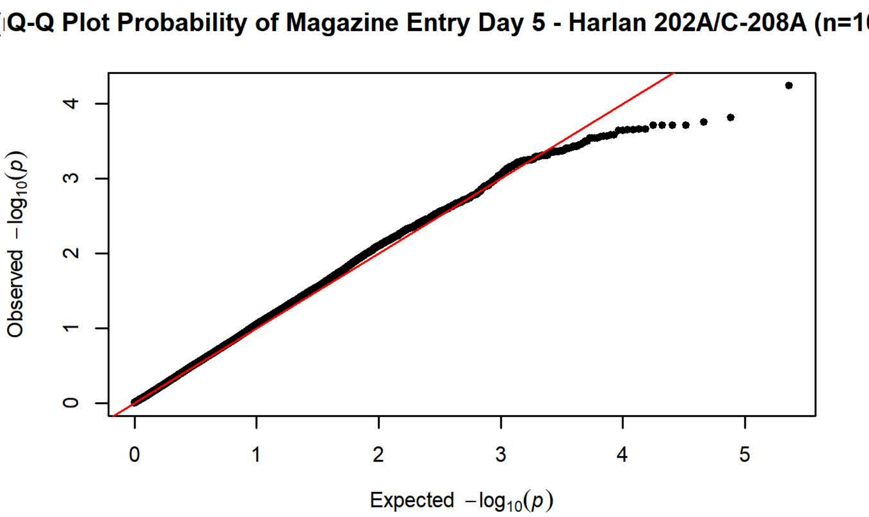
Q-Q Plot Probability of Magazine Entry Day 5 - Harlan 202A/C-208A (n=1
Observed $-\log_{10}(p)$
Expected $-\log_{10}(p)$

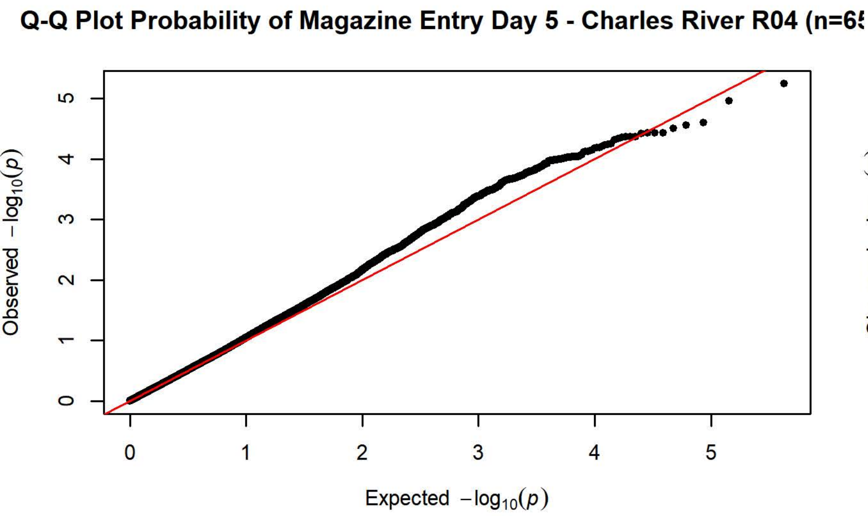
Q-Q Plot Probability of Magazine Entry Day 5 - Charles River R04 (n=6
Observed $-\log_{10}(p)$
Expected $-\log_{10}(p)$

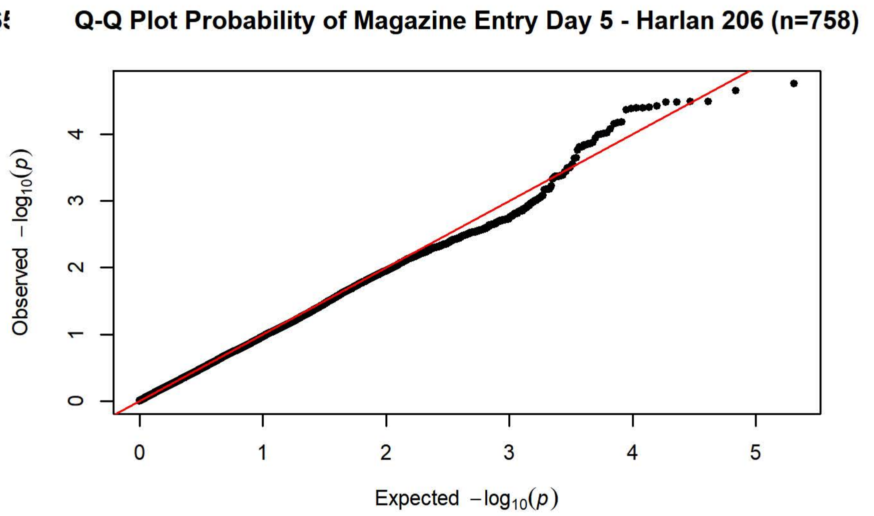
Q-Q Plot Probability of Magazine Entry Day 5 - Harlan 206 (n=758)
Observed $-\log_{10}(p)$
Expected $-\log_{10}(p)$

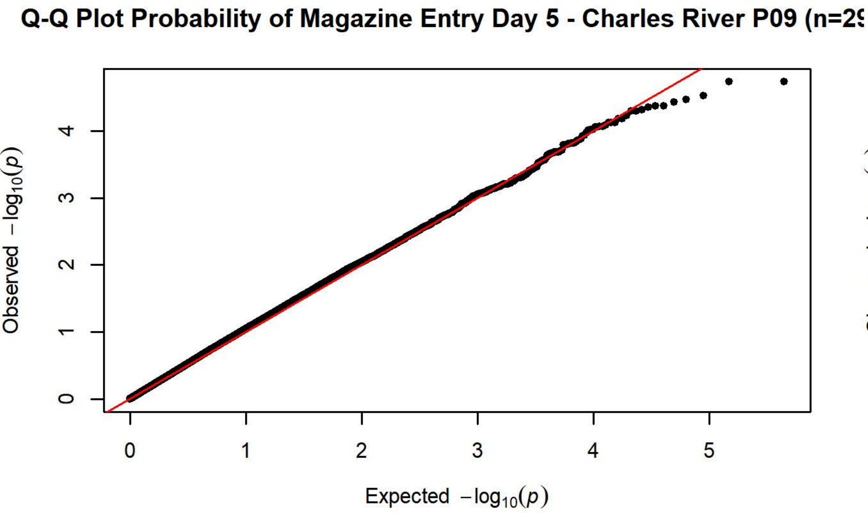
Q-Q Plot Probability of Magazine Entry Day 5 - Charles River P09 (n=2
Observed $-\log_{10}(p)$
Expected $-\log_{10}(p)$

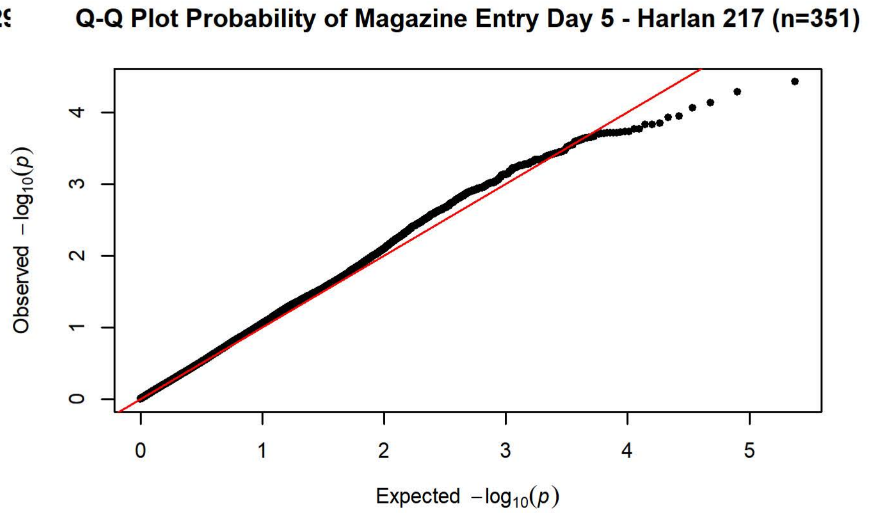
Q-Q Plot Probability of Magazine Entry Day 5 - Harlan 217 (n=351)
Observed $-\log_{10}(p)$
Expected $-\log_{10}(p)$

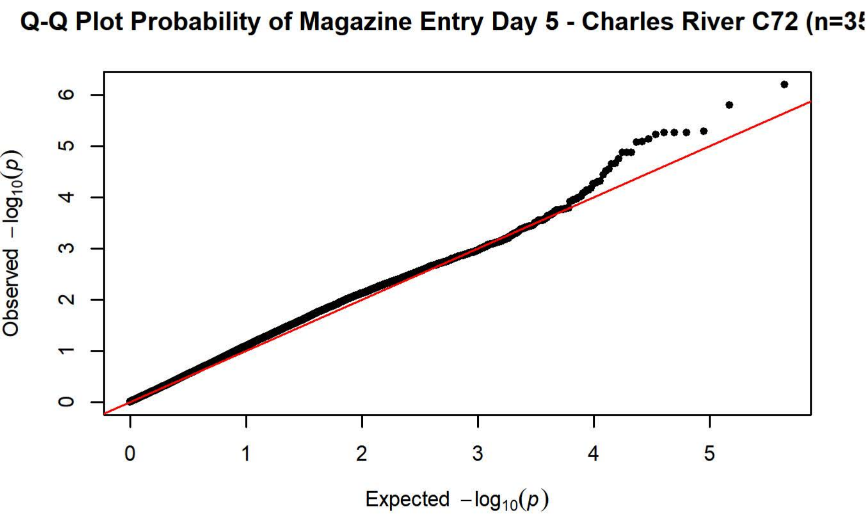
Q-Q Plot Probability of Magazine Entry Day 5 - Charles River C72 (n=3
Observed $-\log_{10}(p)$
Expected $-\log_{10}(p)$

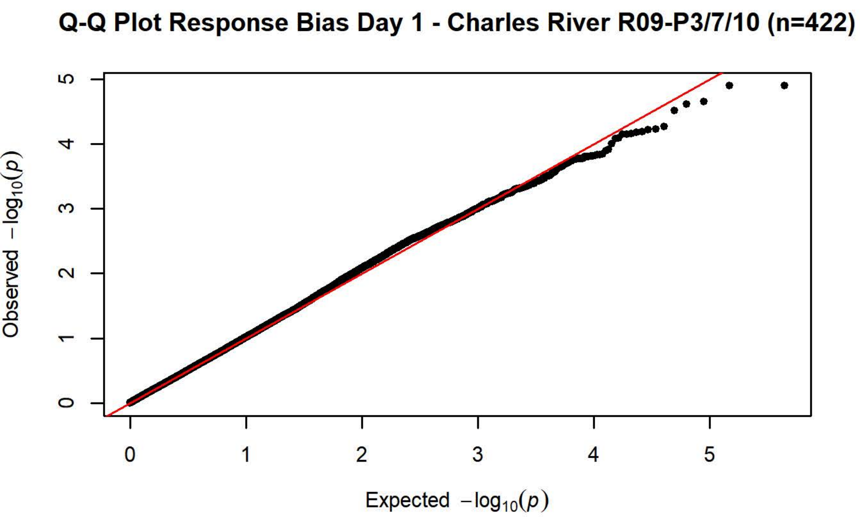
Q-Q Plot Response Bias Day 1 - Charles River R09-P3/7/10 (n=422)
Observed $-\log_{10}(p)$
Expected $-\log_{10}(p)$

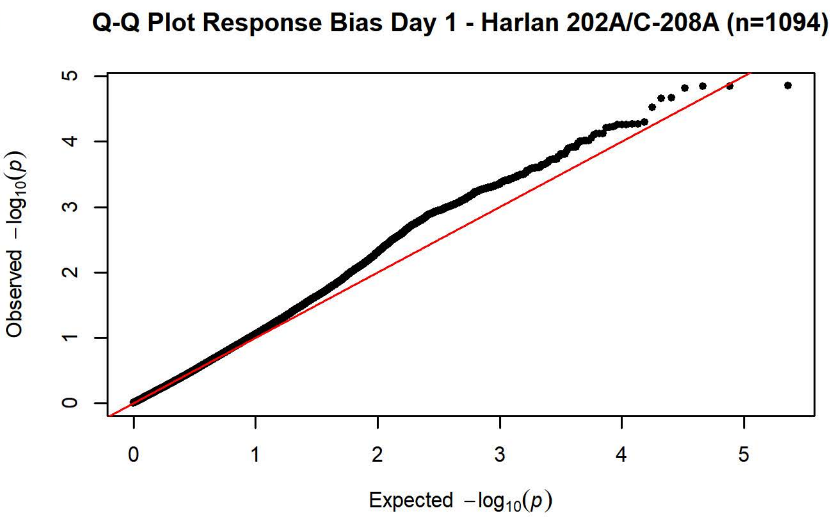
Q-Q Plot Response Bias Day 1 - Harlan 202A/C-208A (n=1094)
Observed $-\log_{10}(p)$
Expected $-\log_{10}(p)$

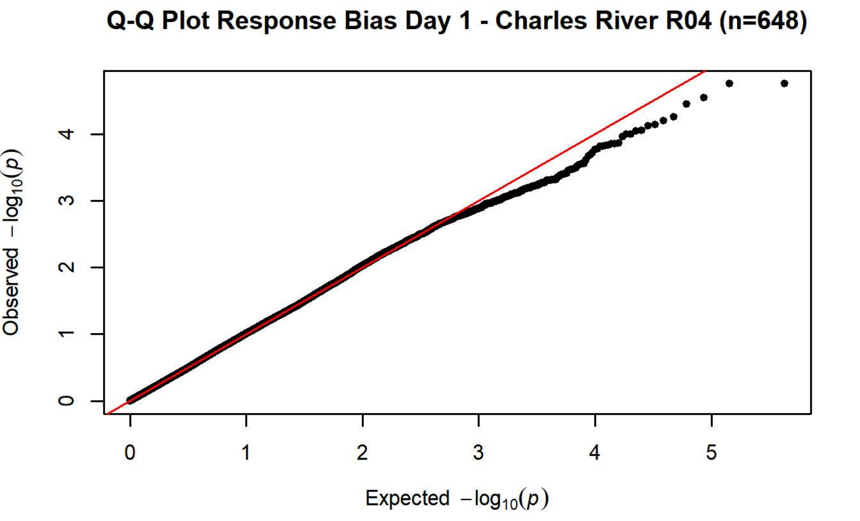
Q-Q Plot Response Bias Day 1 - Charles River R04 (n=648)
Observed $-\log_{10}(p)$
Expected $-\log_{10}(p)$

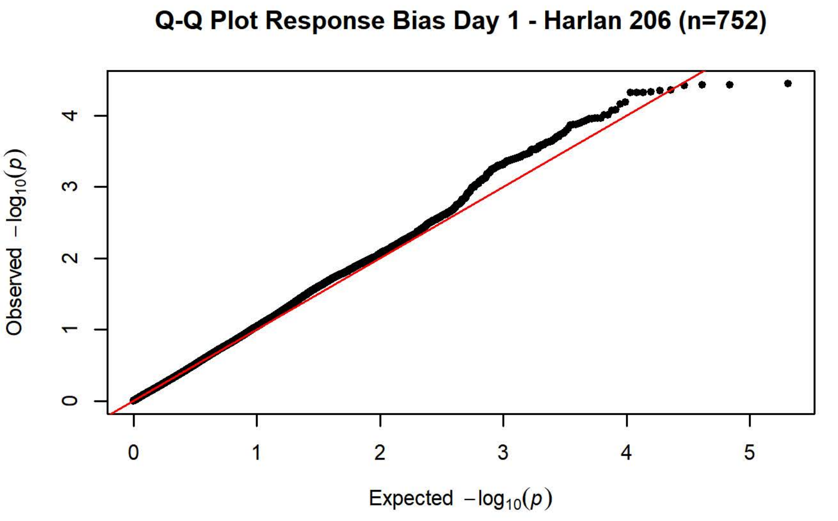
Q-Q Plot Response Bias Day 1 - Harlan 206 (n=752)
Observed $-\log_{10}(p)$
Expected $-\log_{10}(p)$

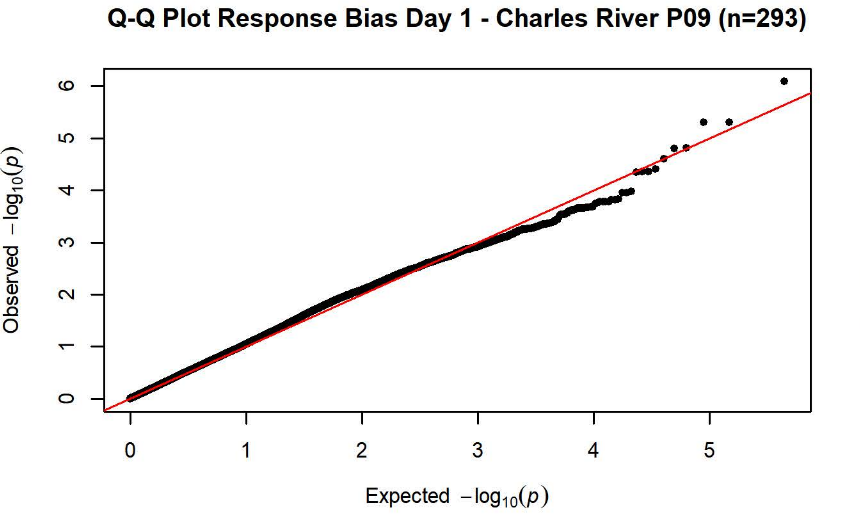
Q-Q Plot Response Bias Day 1 - Charles River P09 (n=293)
Observed $-\log_{10}(p)$
Expected $-\log_{10}(p)$

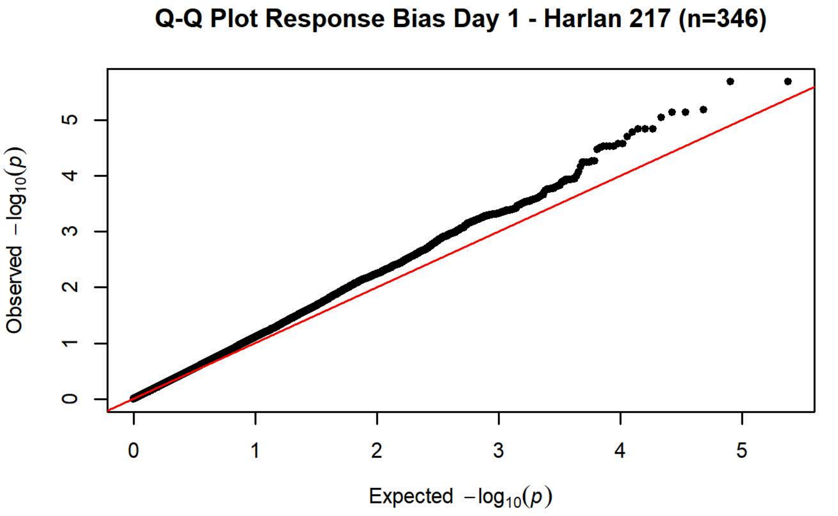
Q-Q Plot Response Bias Day 1 - Harlan 217 (n=346)
Observed $-\log_{10}(p)$
Expected $-\log_{10}(p)$

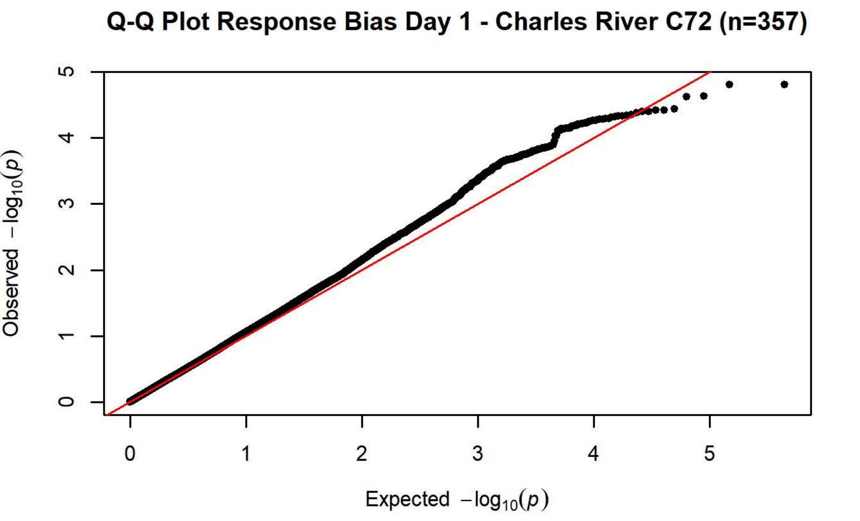
Q-Q Plot Response Bias Day 1 - Charles River C72 (n=357)
Observed $-\log_{10}(p)$
Expected $-\log_{10}(p)$

**Q-Q Plot Response Bias Day 2 - Charles River R09-P3/7/10 (n=425)**

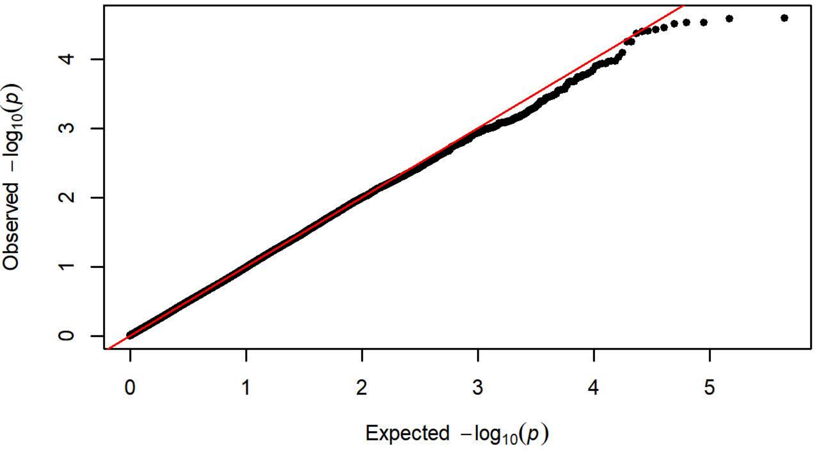


**Q-Q Plot Response Bias Day 2 - Harlan 202A/C-208A (n=1096)**

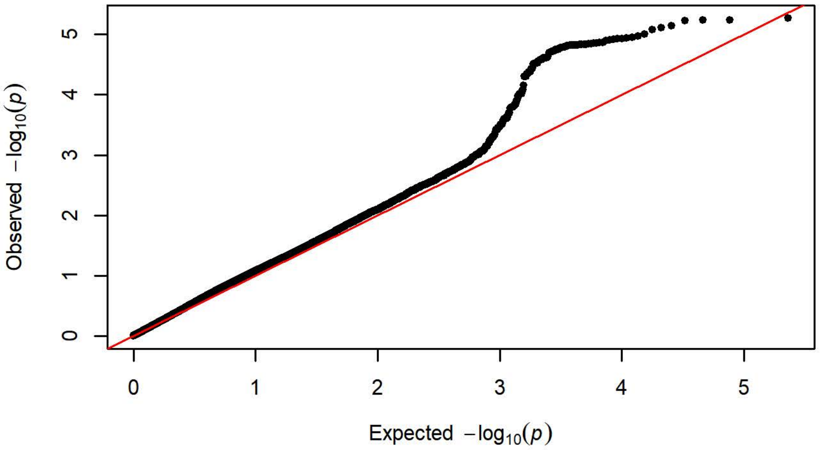


**Q-Q Plot Response Bias Day 2 - Charles River R04 (n=646)**

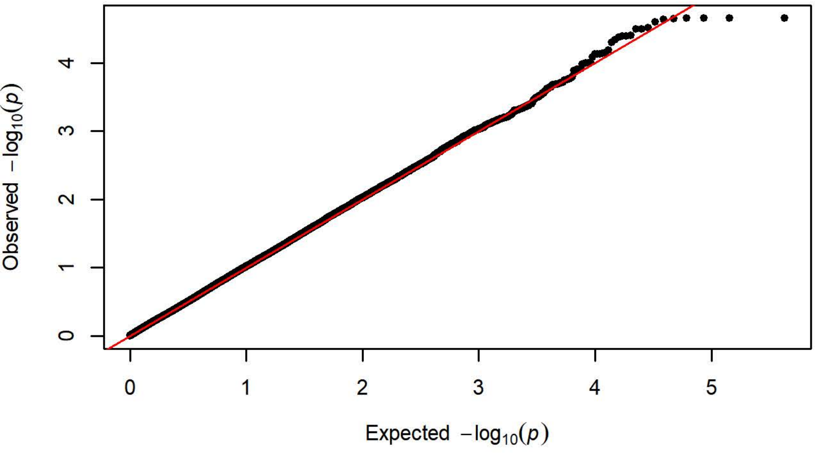


**Q-Q Plot Response Bias Day 2 - Harlan 206 (n=755)**

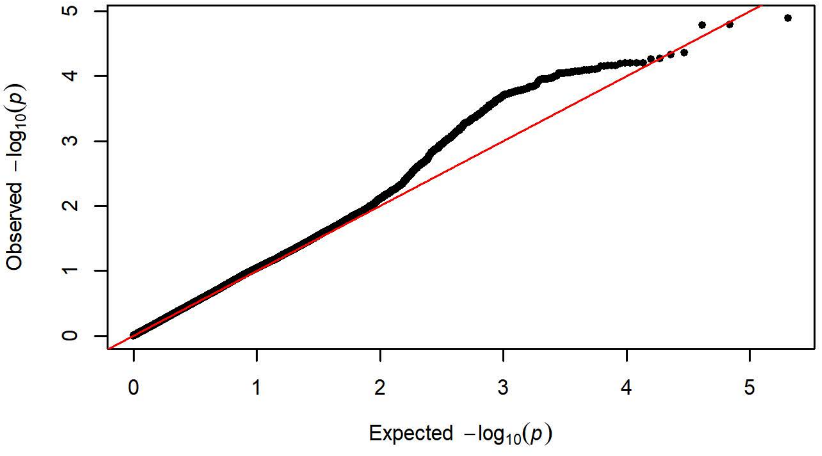


**Q-Q Plot Response Bias Day 2 - Charles River P09 (n=293)**

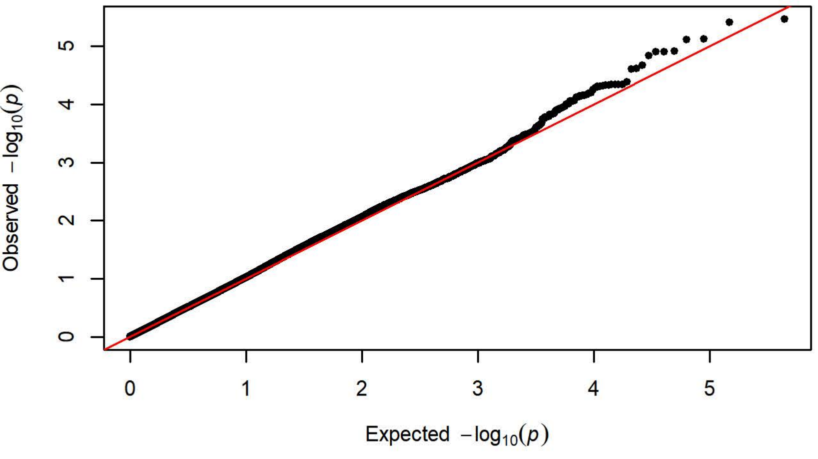


**Q-Q Plot Response Bias Day 2 - Harlan 217 (n=349)**

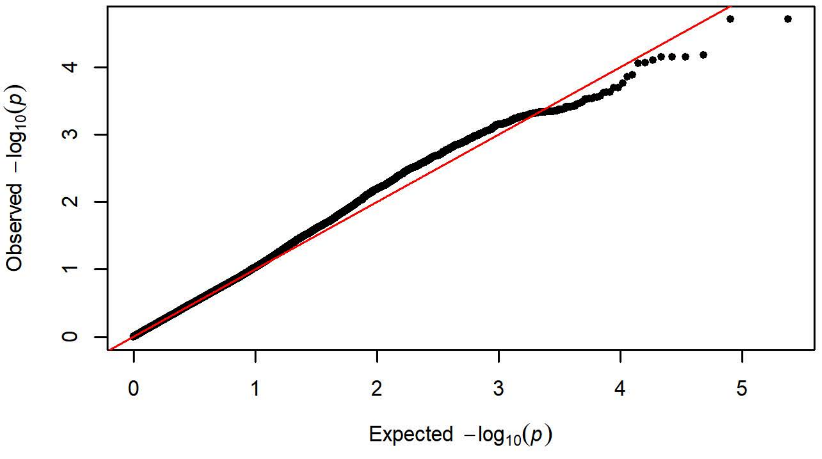


**Q-Q Plot Response Bias Day 2 - Charles River C72 (n=355)**

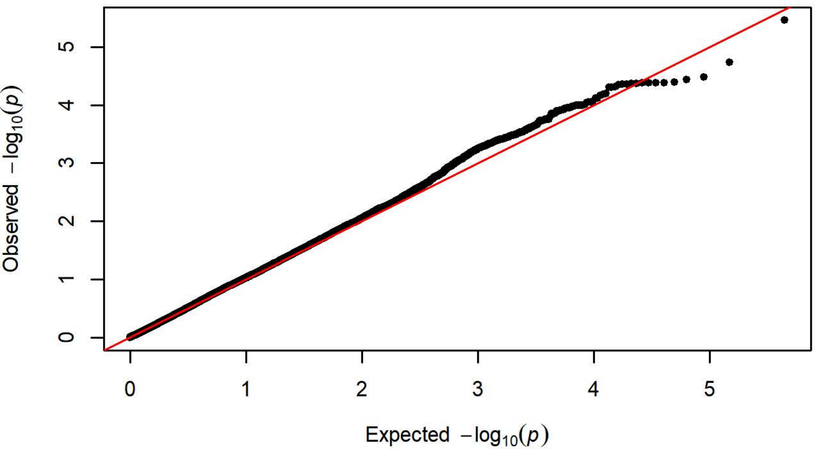

**Q-Q Plot Response Bias Day 3 - Charles River R09-P3/7/10 (n=423)**

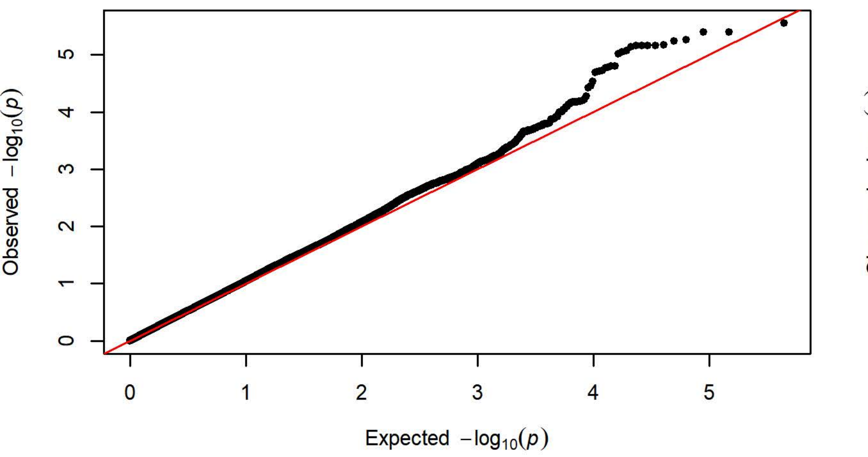


**Q-Q Plot Response Bias Day 3 - Harlan 202A/C-208A (n=1095)**

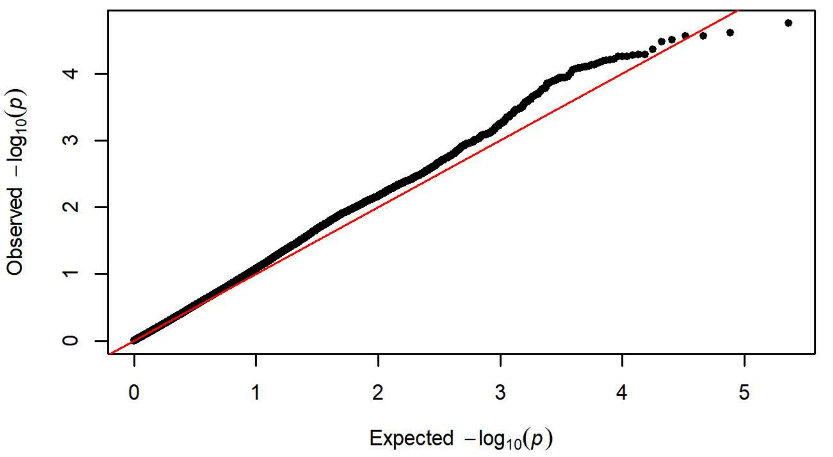


**Q-Q Plot Response Bias Day 3 - Charles River R04 (n=649)**

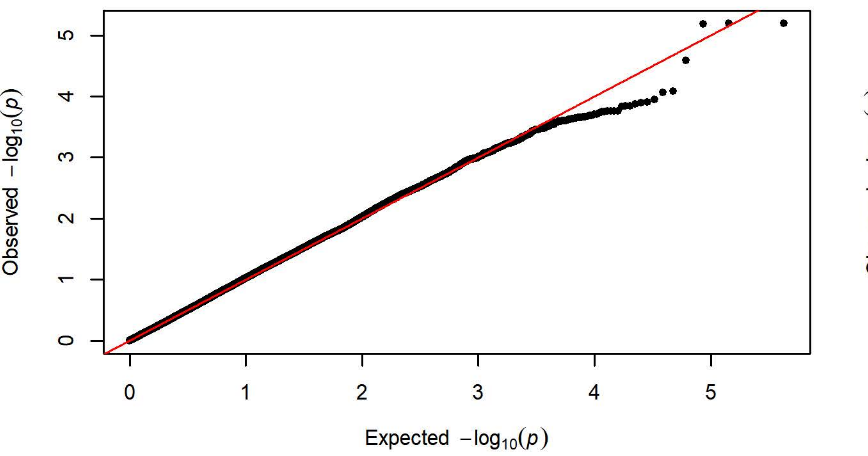


**Q-Q Plot Response Bias Day 3 - Harlan 206 (n=757)**

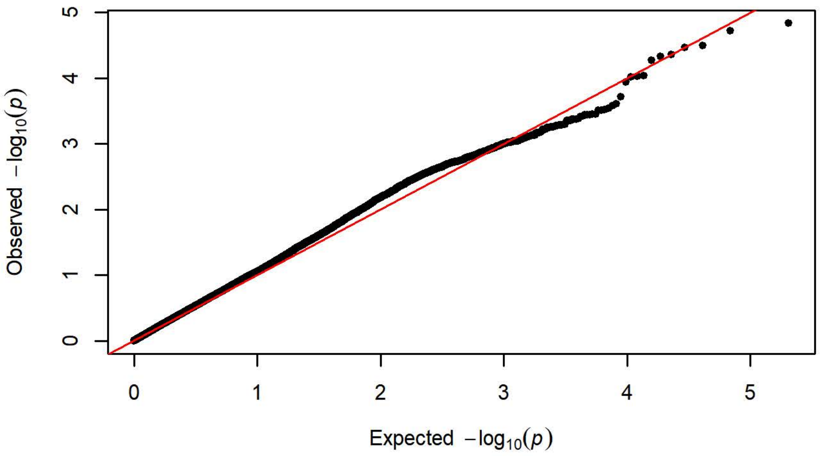


**Q-Q Plot Response Bias Day 3 - Charles River P09 (n=292)**

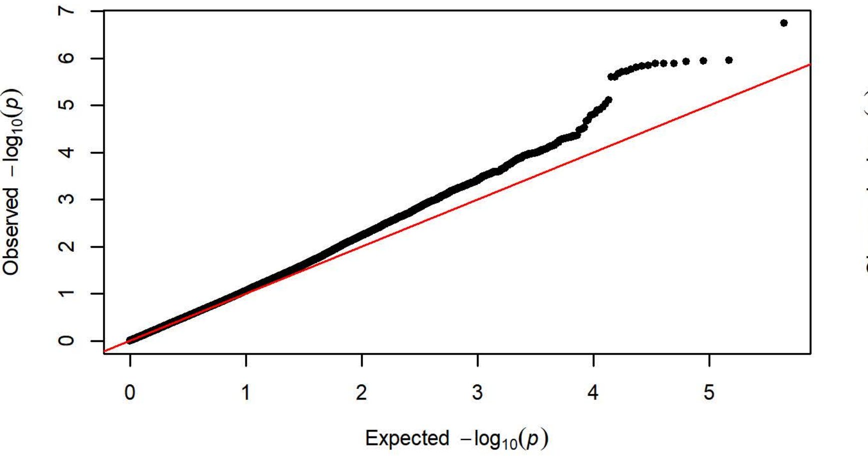


**Q-Q Plot Response Bias Day 3 - Harlan 217 (n=349)**

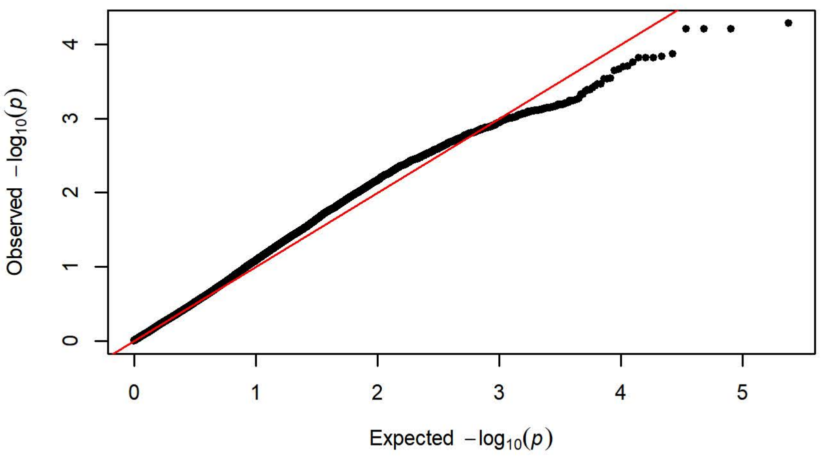


**Q-Q Plot Response Bias Day 3 - Charles River C72 (n=358)**

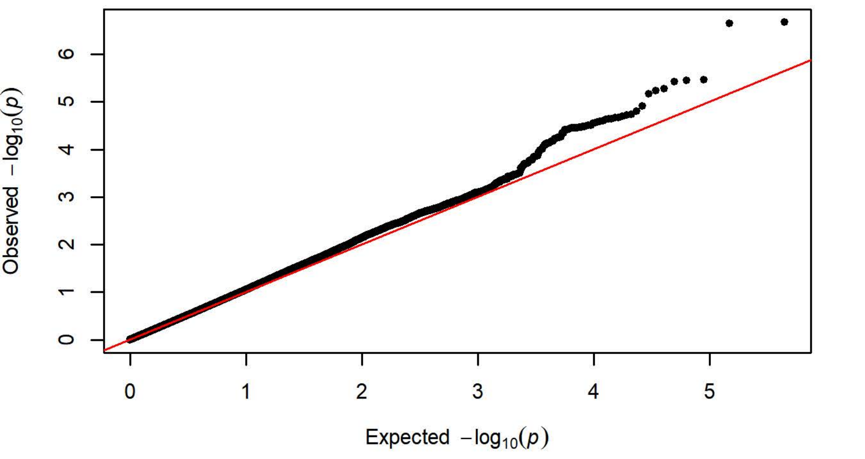

**Q-Q Plot Response Bias Day 4 - Charles River R09-P3/7/10 (n=425)**

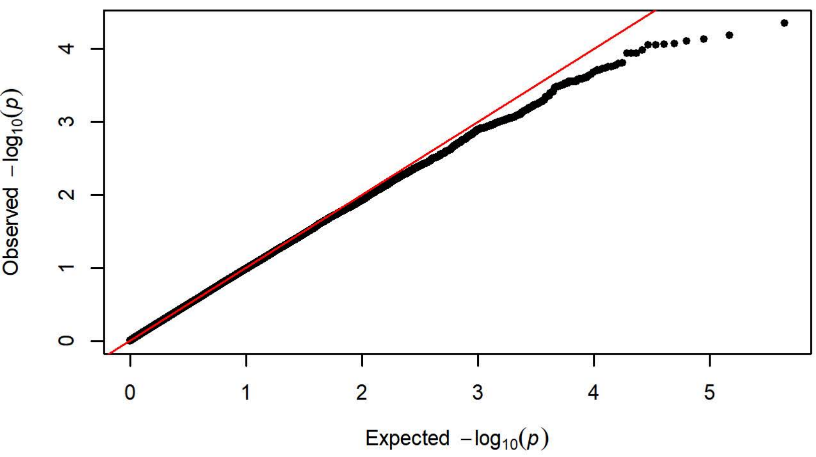


**Q-Q Plot Response Bias Day 4 - Harlan 202A/C-208A (n=1099)**

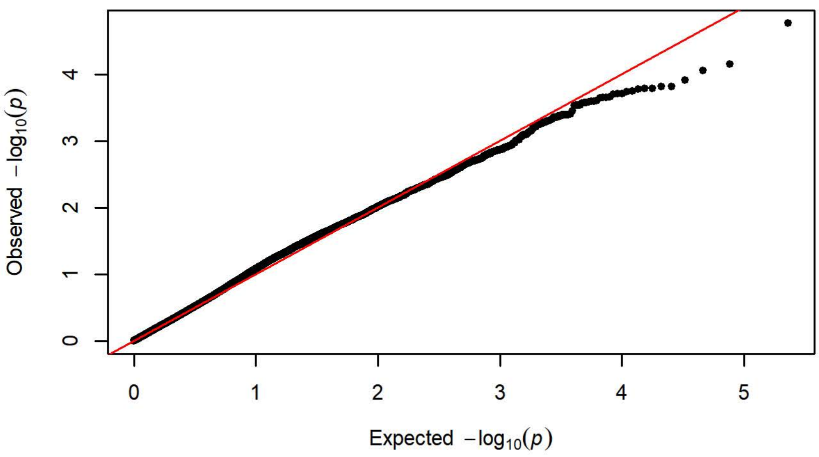


**Q-Q Plot Response Bias Day 4 - Charles River R04 (n=650)**

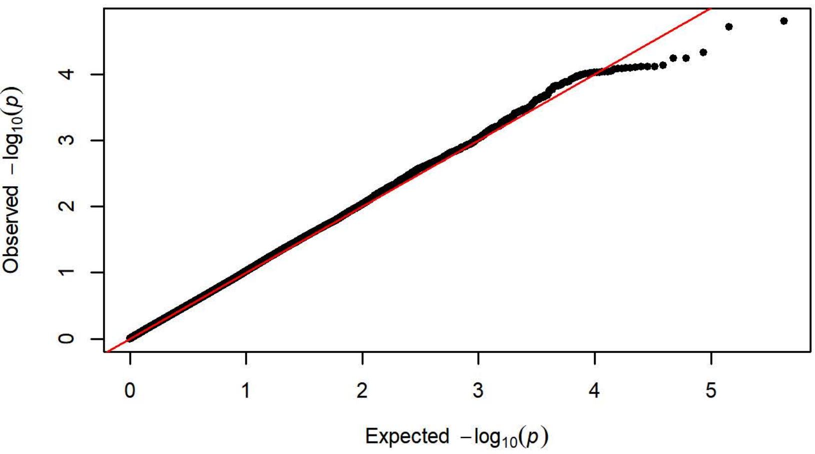


**Q-Q Plot Response Bias Day 4 - Harlan 206 (n=758)**

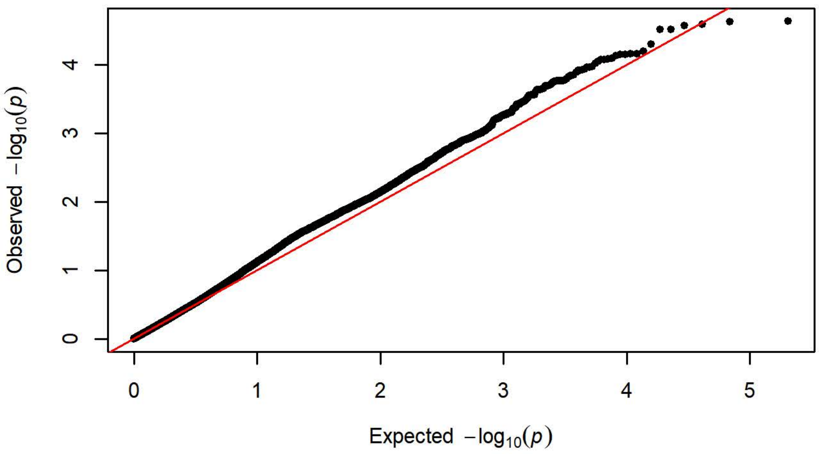


**Q-Q Plot Response Bias Day 4 - Charles River P09 (n=294)**

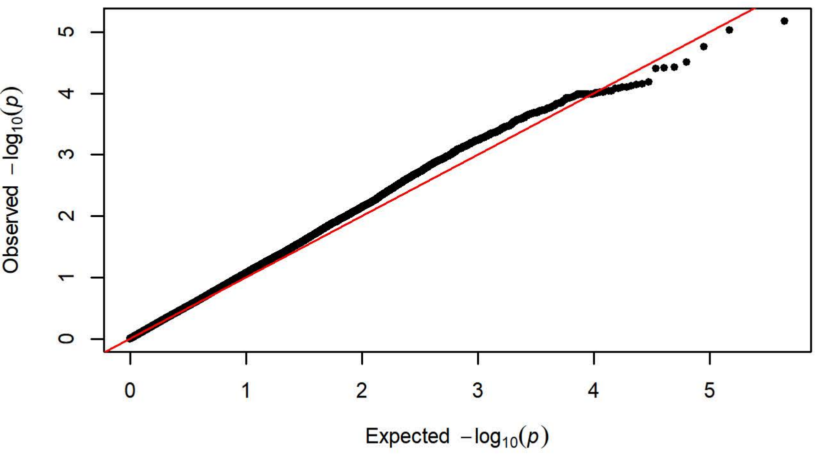


**Q-Q Plot Response Bias Day 4 - Harlan 217 (n=351)**

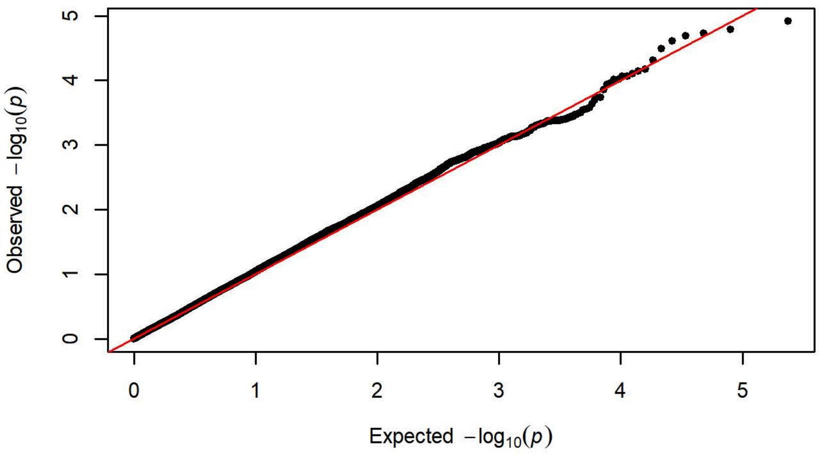


**Q-Q Plot Response Bias Day 4 - Charles River C72 (n=358)**

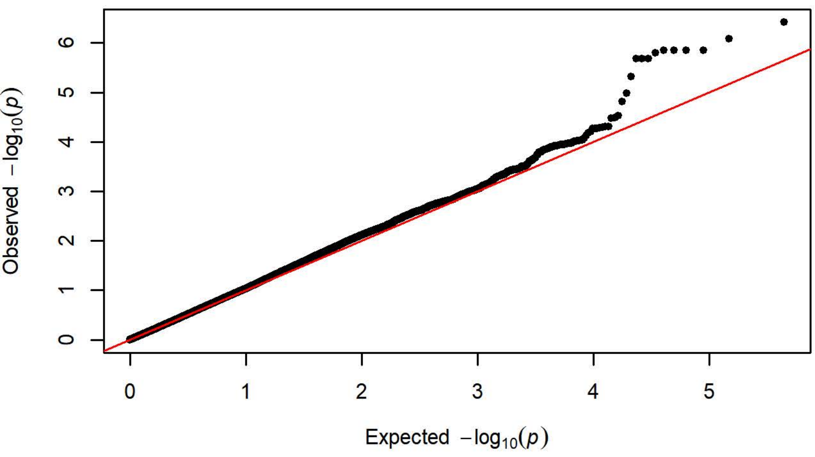

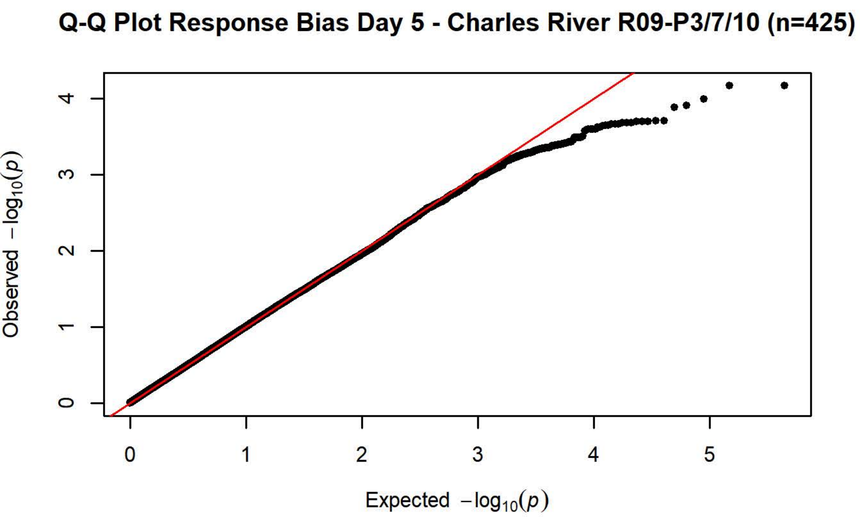
Q-Q Plot Response Bias Day 5 - Charles River R09-P3/7/10 (n=425)
Observed $-\log_{10}(p)$
Expected $-\log_{10}(p)$

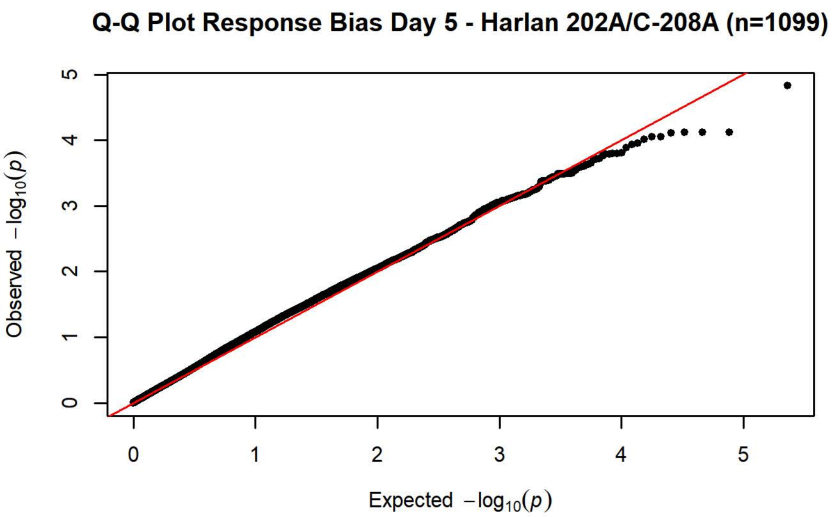
Q-Q Plot Response Bias Day 5 - Harlan 202A/C-208A (n=1099)
Observed $-\log_{10}(p)$
Expected $-\log_{10}(p)$

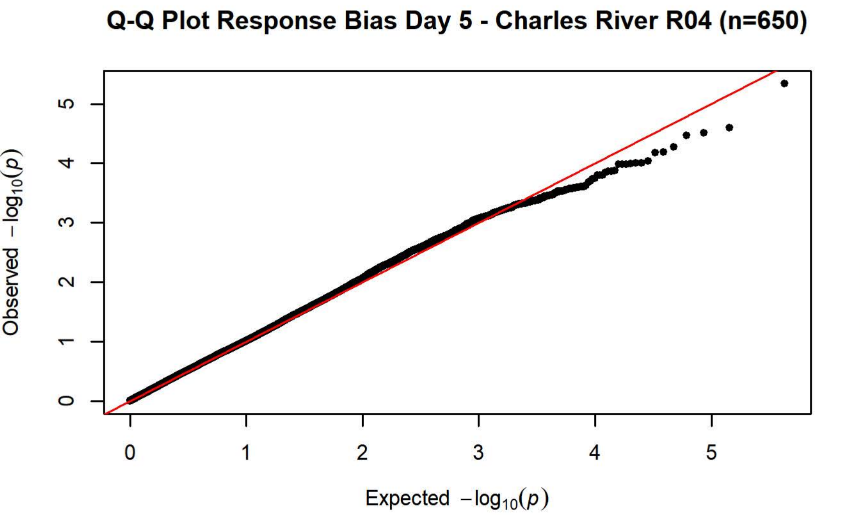
Q-Q Plot Response Bias Day 5 - Charles River R04 (n=650)
Observed $-\log_{10}(p)$
Expected $-\log_{10}(p)$

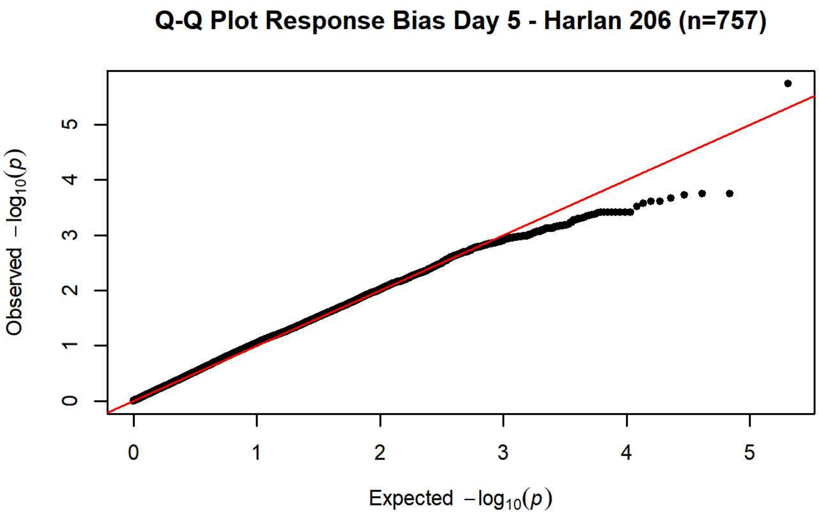
Q-Q Plot Response Bias Day 5 - Harlan 206 (n=757)
Observed $-\log_{10}(p)$
Expected $-\log_{10}(p)$

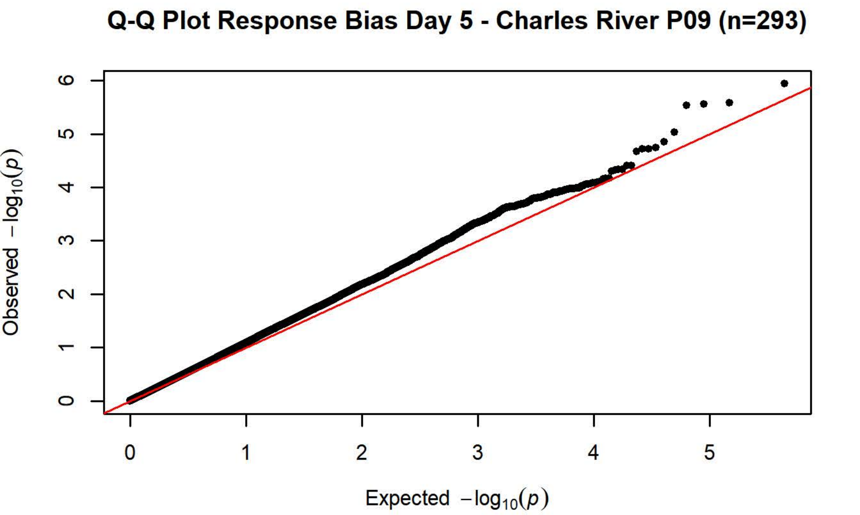
Q-Q Plot Response Bias Day 5 - Charles River P09 (n=293)
Observed $-\log_{10}(p)$
Expected $-\log_{10}(p)$

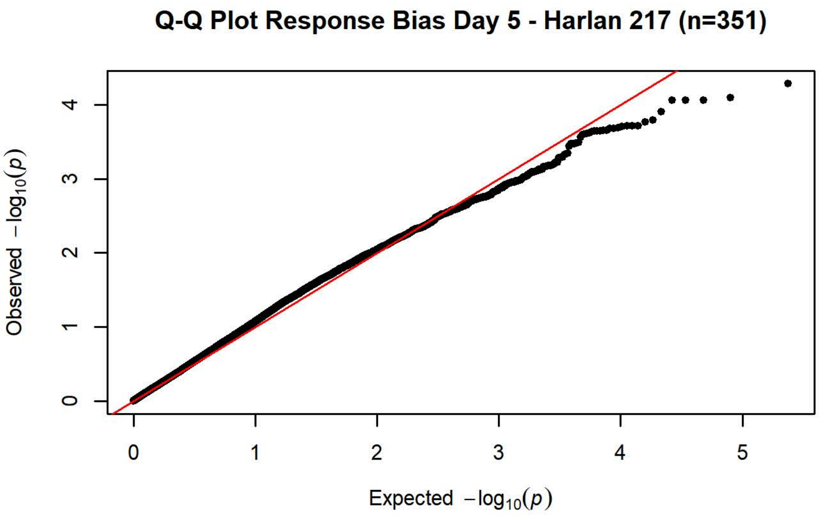
Q-Q Plot Response Bias Day 5 - Harlan 217 (n=351)
Observed $-\log_{10}(p)$
Expected $-\log_{10}(p)$

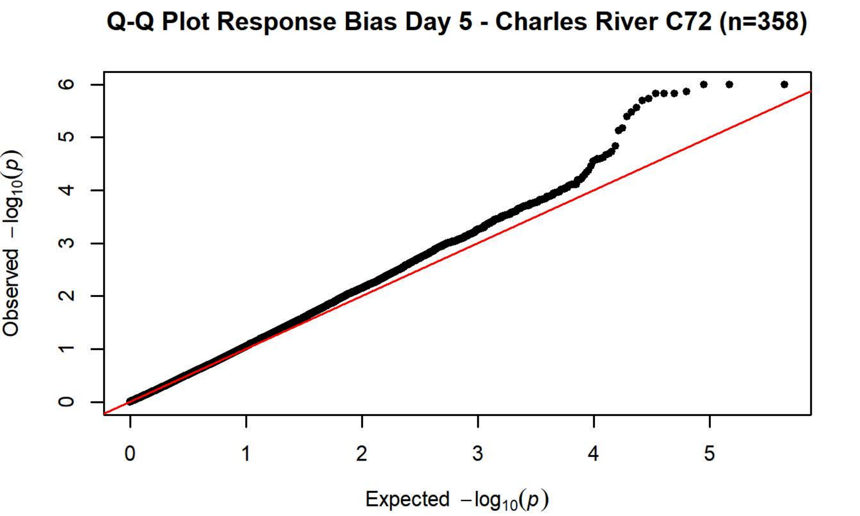
Q-Q Plot Response Bias Day 5 - Charles River C72 (n=358)
Observed $-\log_{10}(p)$
Expected $-\log_{10}(p)$
