## Supplementary material for "Genetic characterization of outbred Sprague Dawley rats and utility for genome-wide association studies": S4_File - PavCA GWAS Plots Meta-Analyses Days 1-5

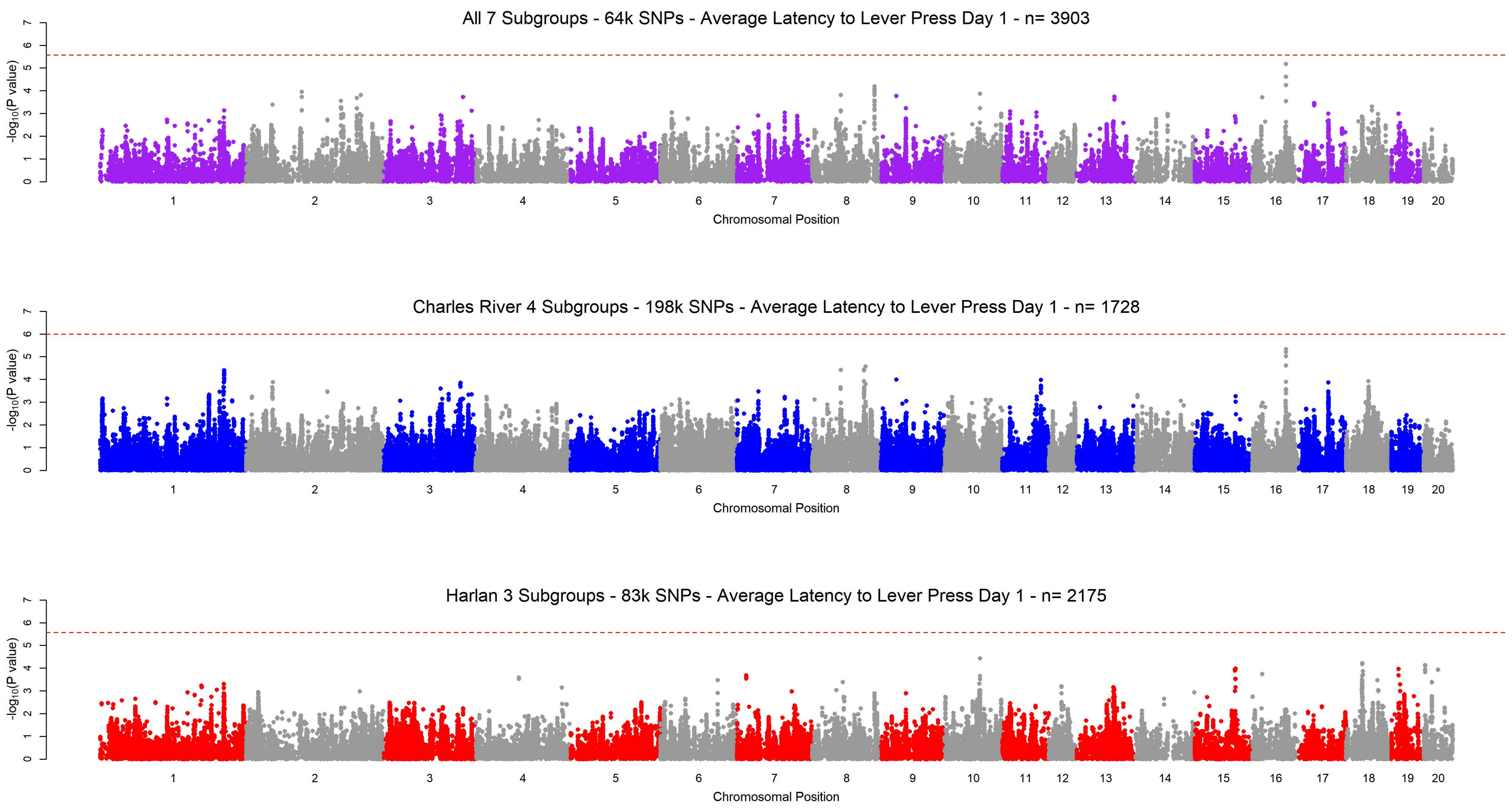
All 7 Subgroups - 64k SNPs - Average Latency to Lever Press Day 1 - n= 3903
Charles River 4 Subgroups - 198k SNPs - Average Latency to Lever Press Day 1 - n= 1728
Harlan 3 Subgroups - 83k SNPs - Average Latency to Lever Press Day 1 - n= 2175
$-\log_{10}$(P value)
0 1 2 3 4 5 6 7
1 2 3 4 5 6 7 8 9 10 11 12 13 14 15 16 17 18 19 20
Chromosomal Position

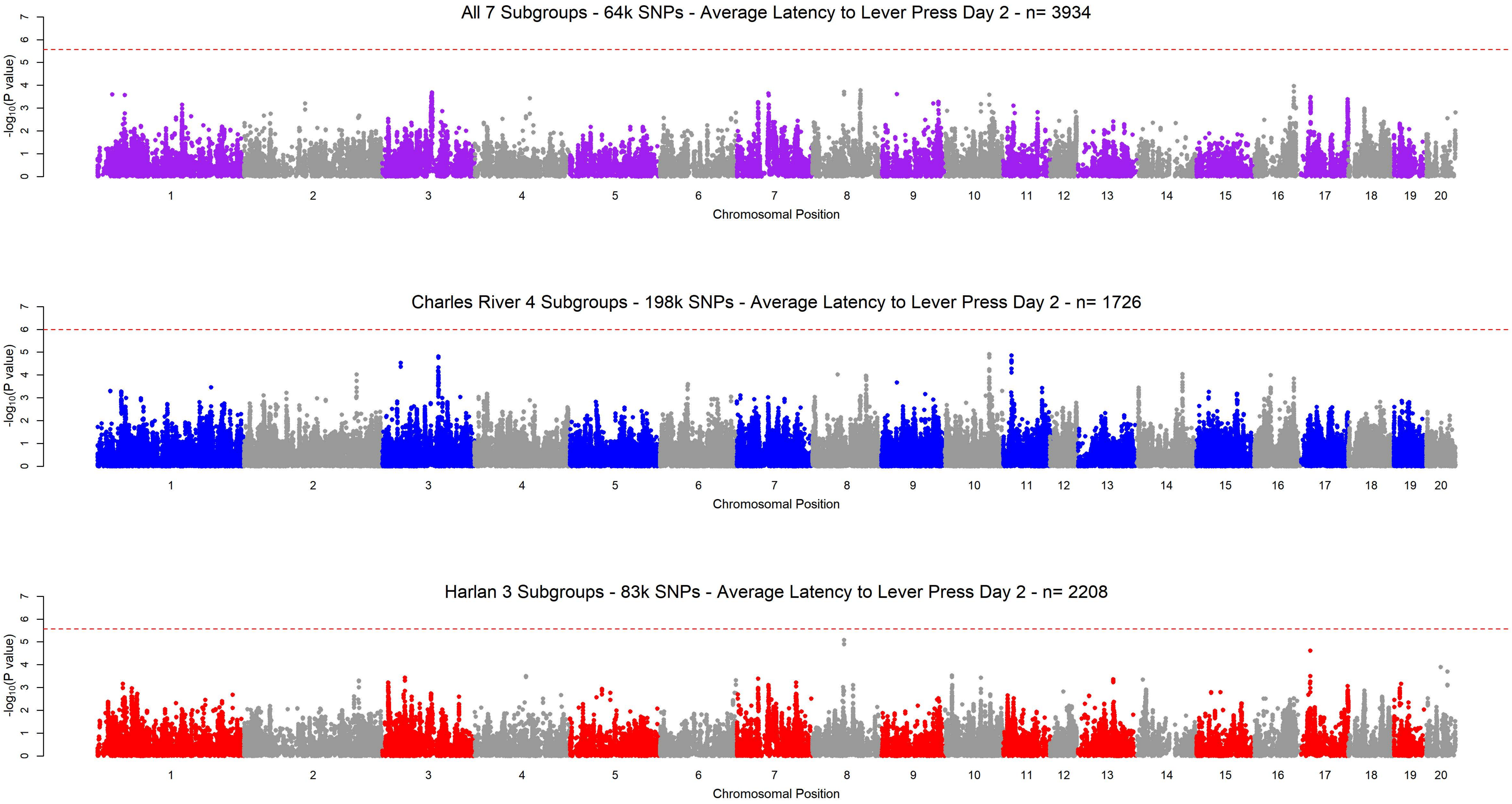

All 7 Subgroups - 64k SNPs - Average Latency to Lever Press Day 2 - n= 3934
Charles River 4 Subgroups - 198k SNPs - Average Latency to Lever Press Day 2 - n= 1726
Harlan 3 Subgroups - 83k SNPs - Average Latency to Lever Press Day 2 - n= 2208
$-\log_{10}$(P value)
0 1 2 3 4 5 6 7
1 2 3 4 5 6 7 8 9 10 11 12 13 14 15 16 17 18 19 20
Chromosomal Position

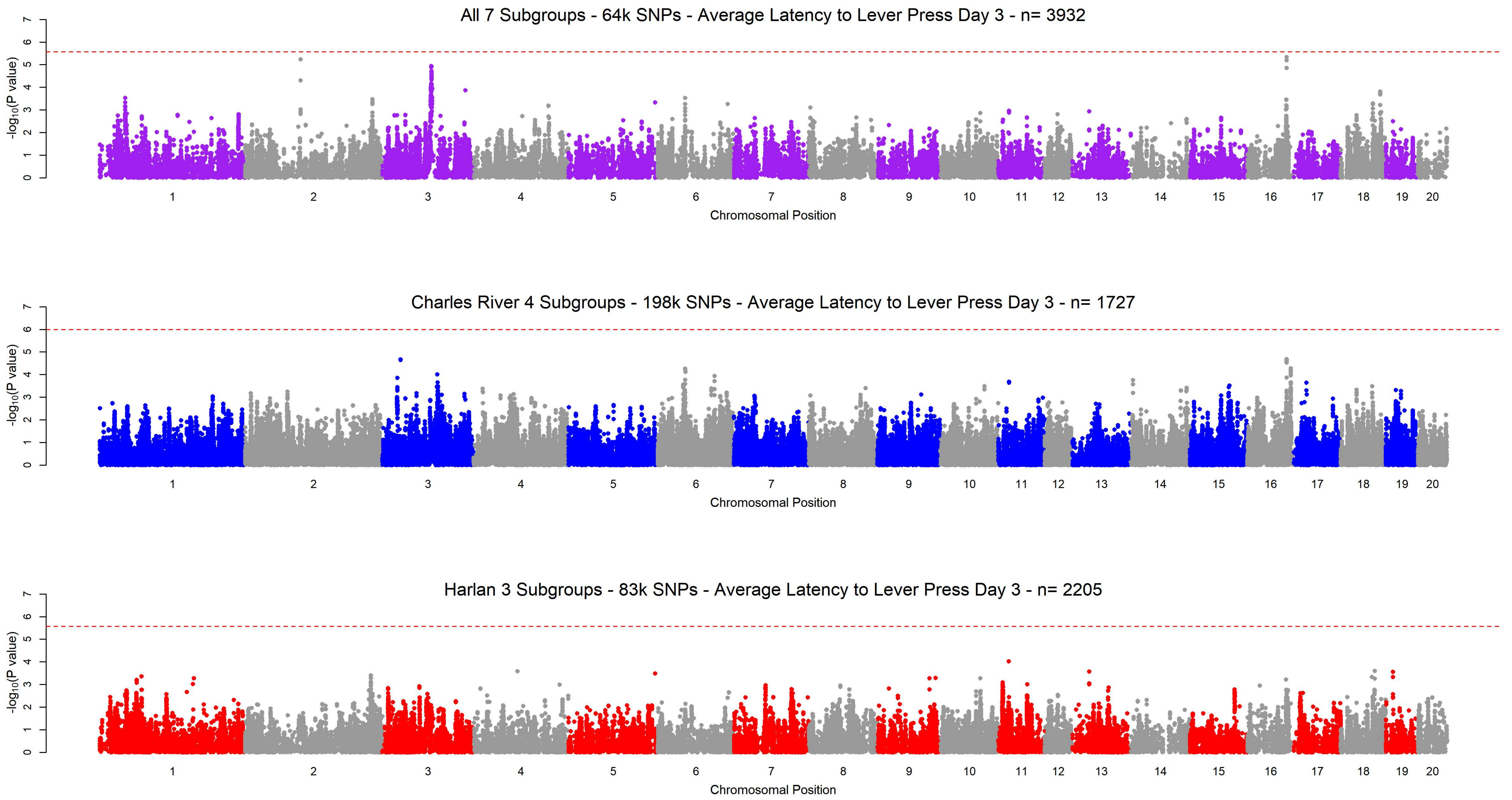
All 7 Subgroups - 64k SNPs - Average Latency to Lever Press Day 3 - n= 3932
-log10(P value)
0 1 2 3 4 5 6 7
1 2 3 4 5 6 7 8 9 10 11 12 13 14 15 16 17 18 19 20
Chromosomal Position
Charles River 4 Subgroups - 198k SNPs - Average Latency to Lever Press Day 3 - n= 1727
-log10(P value)
0 1 2 3 4 5 6 7
1 2 3 4 5 6 7 8 9 10 11 12 13 14 15 16 17 18 19 20
Chromosomal Position
Harlan 3 Subgroups - 83k SNPs - Average Latency to Lever Press Day 3 - n= 2205
-log10(P value)
0 1 2 3 4 5 6 7
1 2 3 4 5 6 7 8 9 10 11 12 13 14 15 16 17 18 19 20
Chromosomal Position

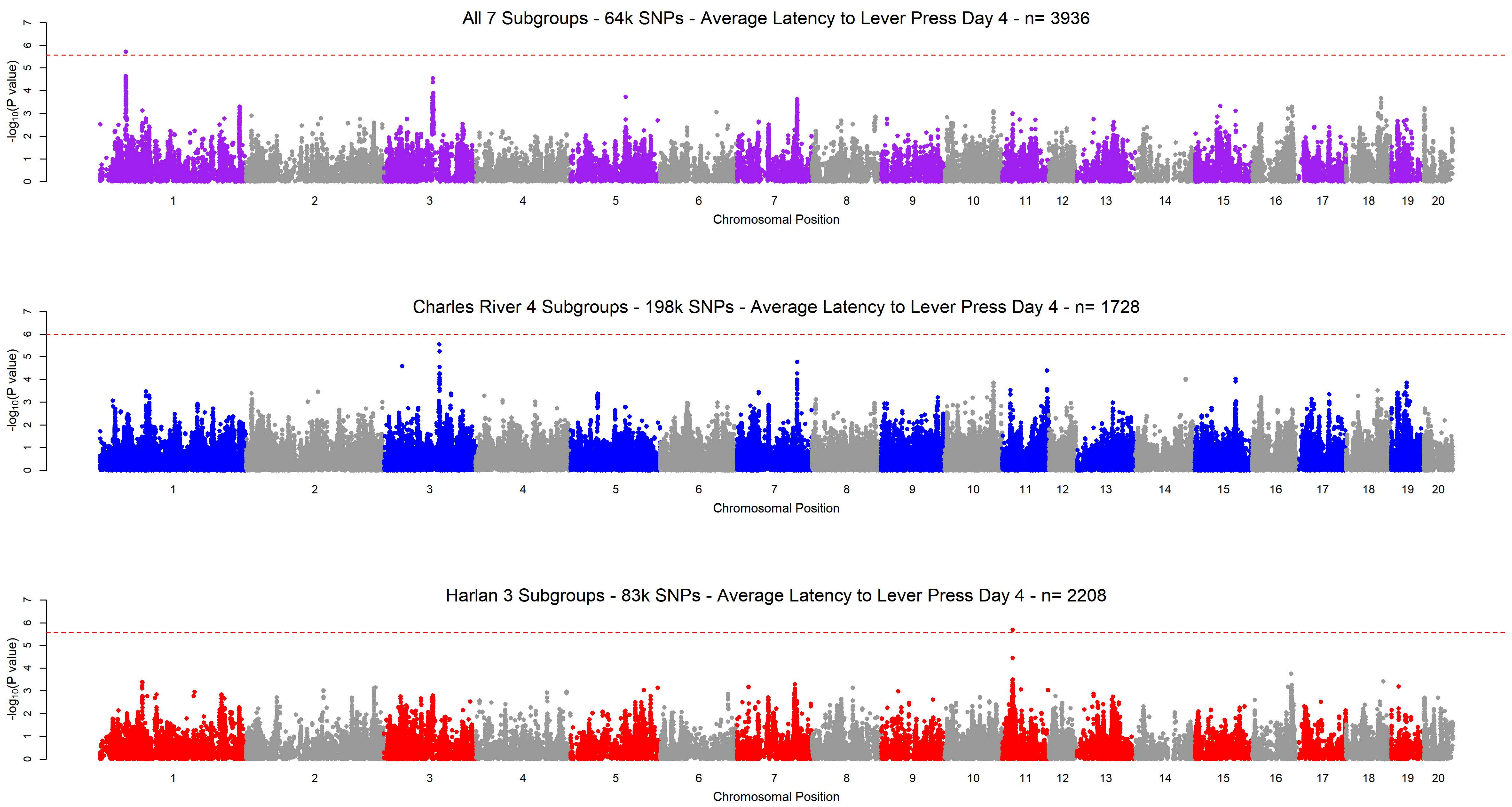
All 7 Subgroups - 64k SNPs - Average Latency to Lever Press Day 4 - n= 3936
-log10(P value)
0 1 2 3 4 5 6 7
1 2 3 4 5 6 7 8 9 10 11 12 13 14 15 16 17 18 19 20
Chromosomal Position
Charles River 4 Subgroups - 198k SNPs - Average Latency to Lever Press Day 4 - n= 1728
-log10(P value)
0 1 2 3 4 5 6 7
1 2 3 4 5 6 7 8 9 10 11 12 13 14 15 16 17 18 19 20
Chromosomal Position
Harlan 3 Subgroups - 83k SNPs - Average Latency to Lever Press Day 4 - n= 2208
-log10(P value)
0 1 2 3 4 5 6 7
1 2 3 4 5 6 7 8 9 10 11 12 13 14 15 16 17 18 19 20
Chromosomal Position

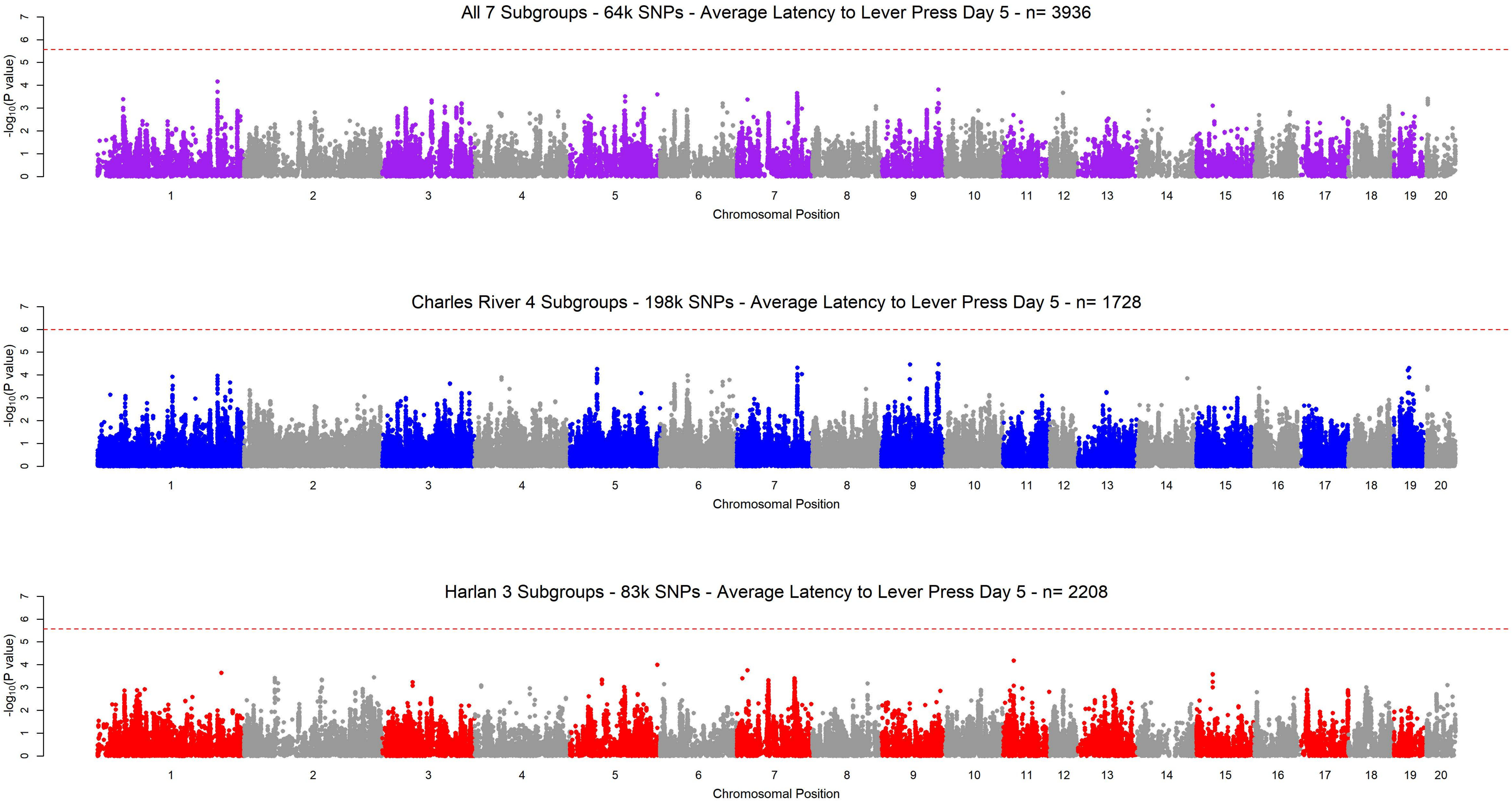
All 7 Subgroups - 64k SNPs - Average Latency to Lever Press Day 5 - n= 3936
-log10(P value)
0 1 2 3 4 5 6 7
1 2 3 4 5 6 7 8 9 10 11 12 13 14 15 16 17 18 19 20
Chromosomal Position
Charles River 4 Subgroups - 198k SNPs - Average Latency to Lever Press Day 5 - n= 1728
-log10(P value)
0 1 2 3 4 5 6 7
1 2 3 4 5 6 7 8 9 10 11 12 13 14 15 16 17 18 19 20
Chromosomal Position
Harlan 3 Subgroups - 83k SNPs - Average Latency to Lever Press Day 5 - n= 2208
-log10(P value)
0 1 2 3 4 5 6 7
1 2 3 4 5 6 7 8 9 10 11 12 13 14 15 16 17 18 19 20
Chromosomal Position

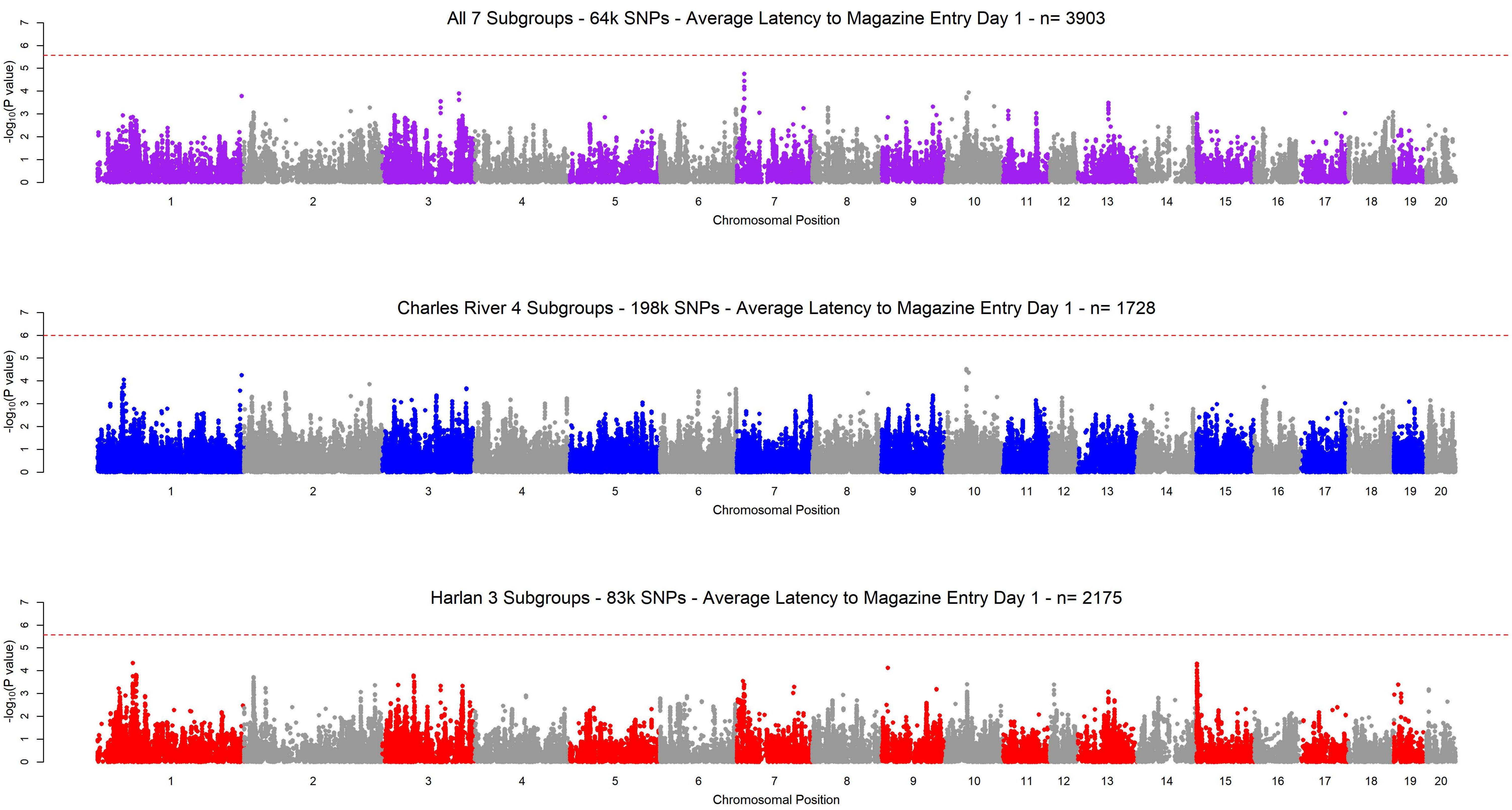
All 7 Subgroups - 64k SNPs - Average Latency to Magazine Entry Day 1 - n= 3903
Charles River 4 Subgroups - 198k SNPs - Average Latency to Magazine Entry Day 1 - n= 1728
Harlan 3 Subgroups - 83k SNPs - Average Latency to Magazine Entry Day 1 - n= 2175
-log10(P value)
Chromosomal Position
0 1 2 3 4 5 6 7
1 2 3 4 5 6 7 8 9 10 11 12 13 14 15 16 17 18 19 20

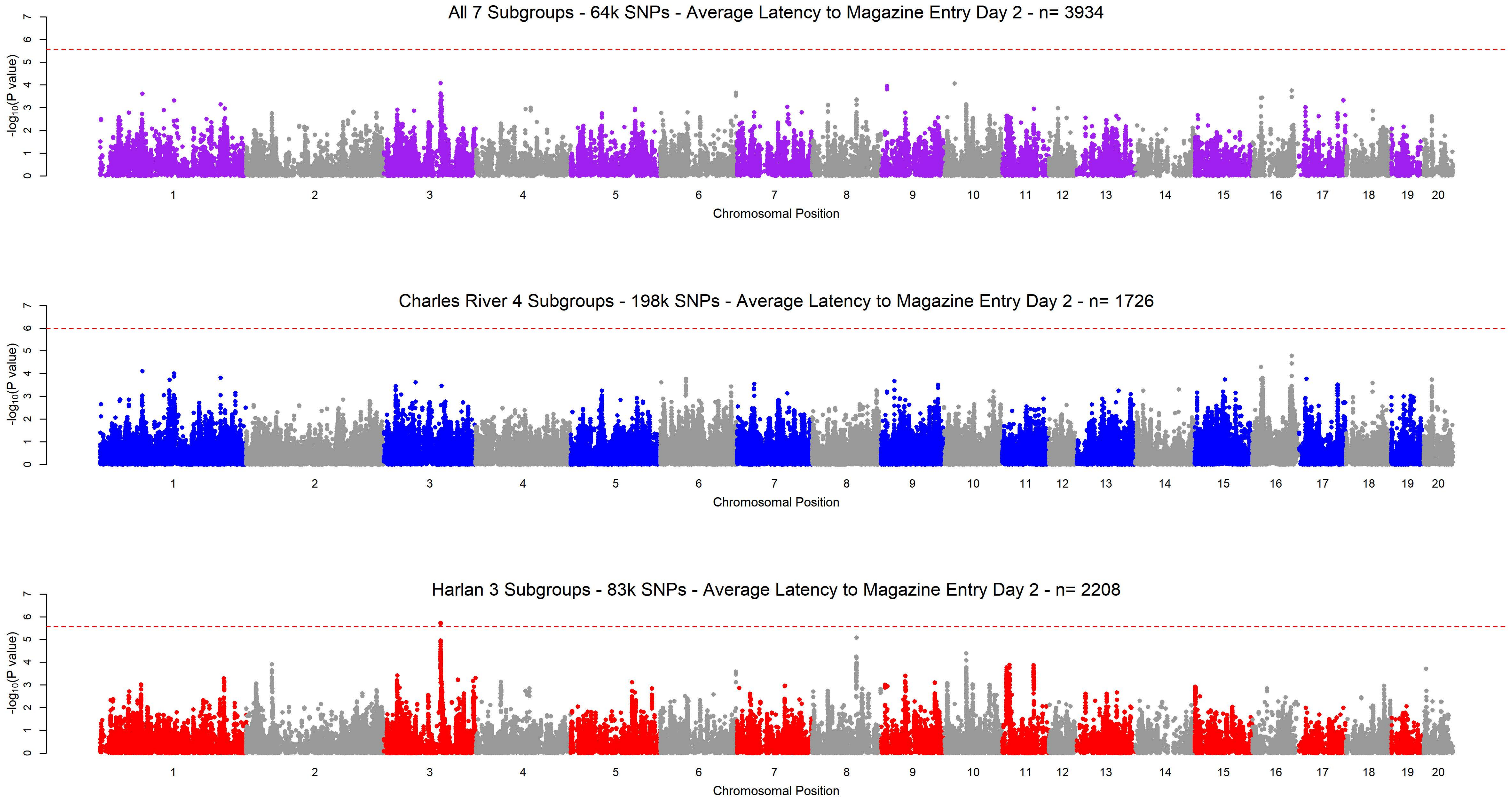
All 7 Subgroups - 64k SNPs - Average Latency to Magazine Entry Day 2 - n= 3934
Charles River 4 Subgroups - 198k SNPs - Average Latency to Magazine Entry Day 2 - n= 1726
Harlan 3 Subgroups - 83k SNPs - Average Latency to Magazine Entry Day 2 - n= 2208
$-\log_{10}$(P value)
Chromosomal Position
1 2 3 4 5 6 7 8 9 10 11 12 13 14 15 16 17 18 19 20
0 1 2 3 4 5 6 7

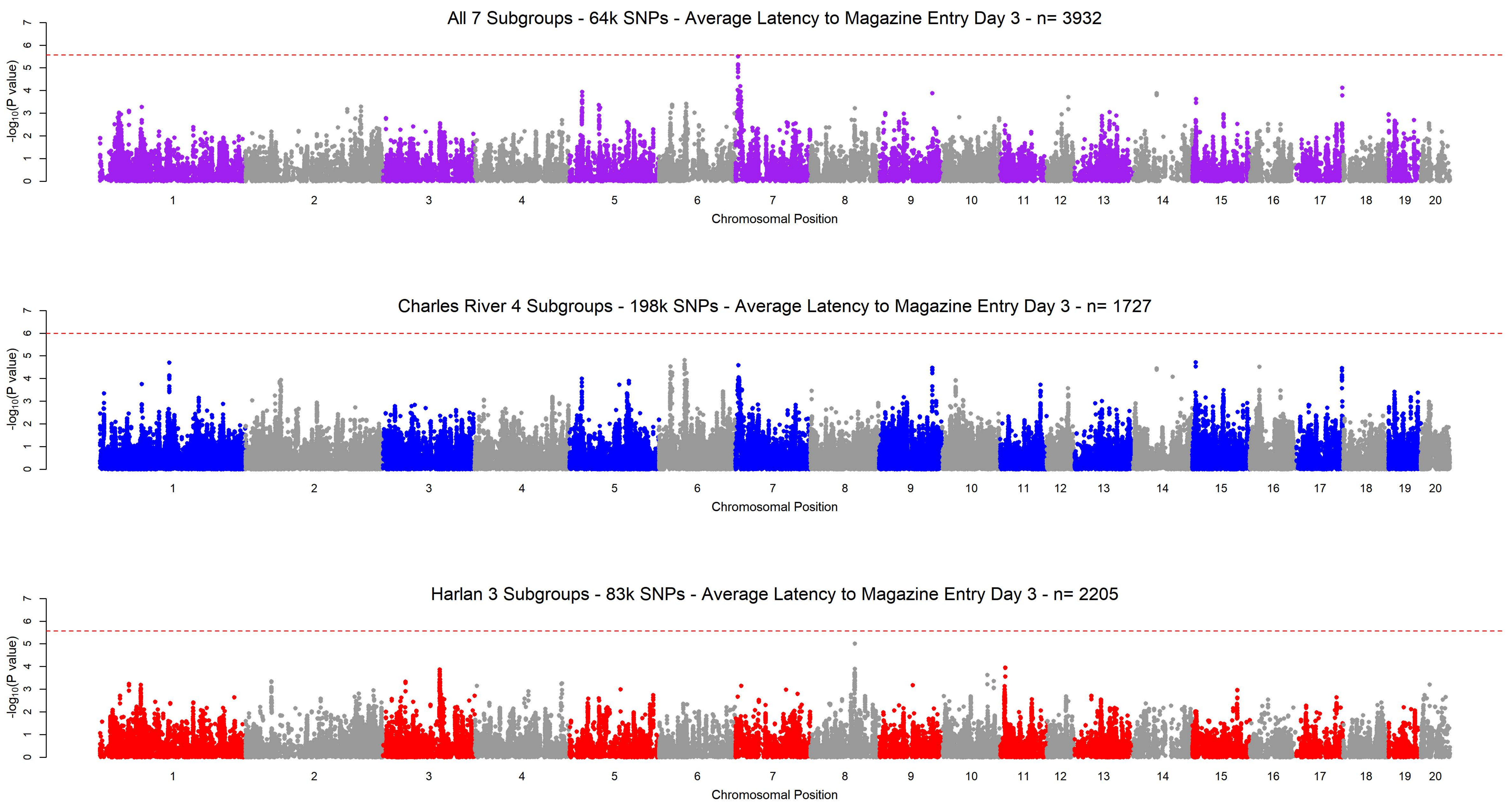
All 7 Subgroups - 64k SNPs - Average Latency to Magazine Entry Day 3 - n= 3932
$-\log_{10}$(P value)
0 1 2 3 4 5 6 7
1 2 3 4 5 6 7 8 9 10 11 12 13 14 15 16 17 18 19 20
Chromosomal Position
Charles River 4 Subgroups - 198k SNPs - Average Latency to Magazine Entry Day 3 - n= 1727
$-\log_{10}$(P value)
0 1 2 3 4 5 6 7
1 2 3 4 5 6 7 8 9 10 11 12 13 14 15 16 17 18 19 20
Chromosomal Position
Harlan 3 Subgroups - 83k SNPs - Average Latency to Magazine Entry Day 3 - n= 2205
$-\log_{10}$(P value)
0 1 2 3 4 5 6 7
1 2 3 4 5 6 7 8 9 10 11 12 13 14 15 16 17 18 19 20
Chromosomal Position

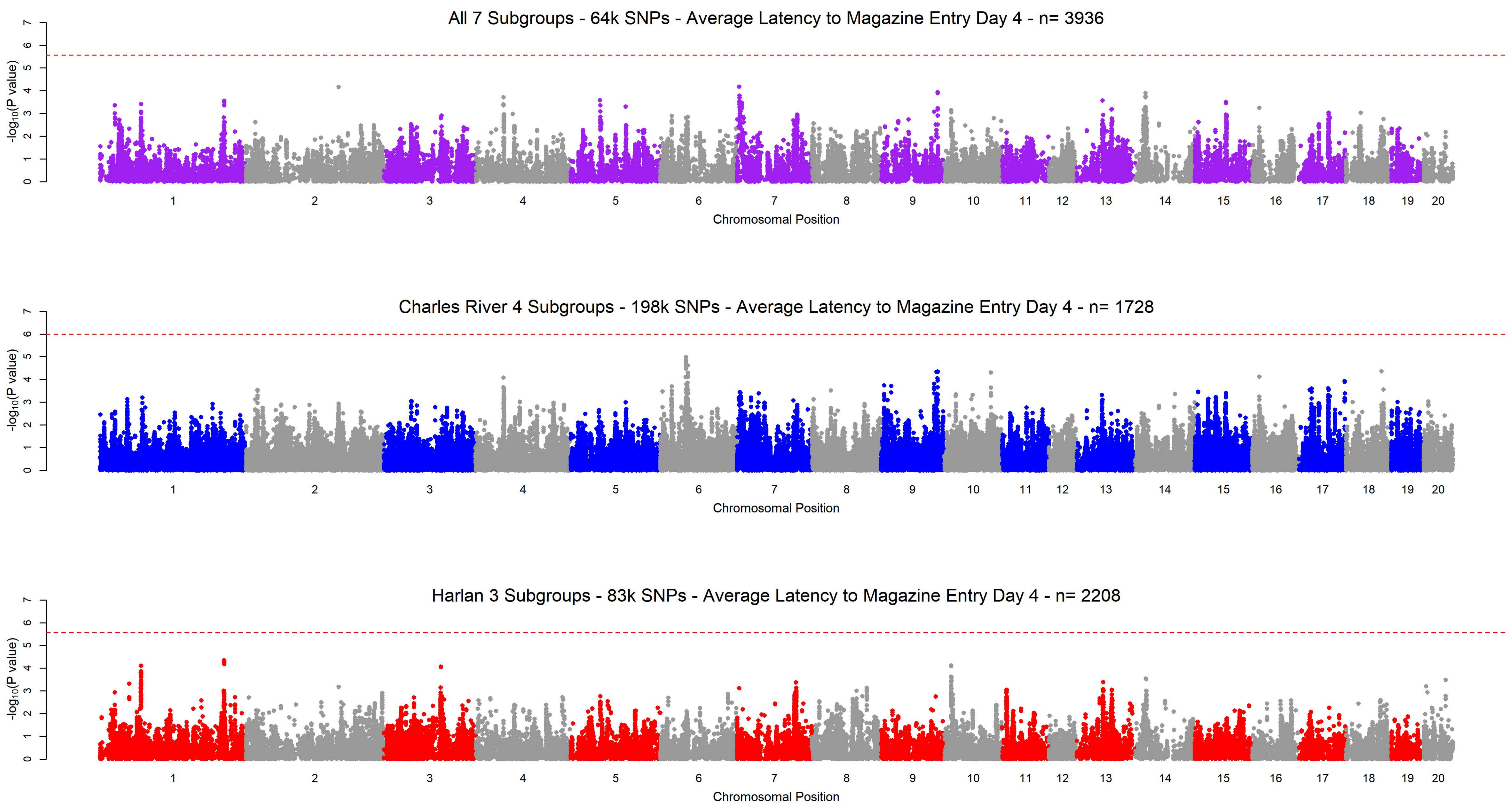
All 7 Subgroups - 64k SNPs - Average Latency to Magazine Entry Day 4 - n= 3936
Charles River 4 Subgroups - 198k SNPs - Average Latency to Magazine Entry Day 4 - n= 1728
Harlan 3 Subgroups - 83k SNPs - Average Latency to Magazine Entry Day 4 - n= 2208
-log10(P value)
Chromosomal Position
0 1 2 3 4 5 6 7
1 2 3 4 5 6 7 8 9 10 11 12 13 14 15 16 17 18 19 20

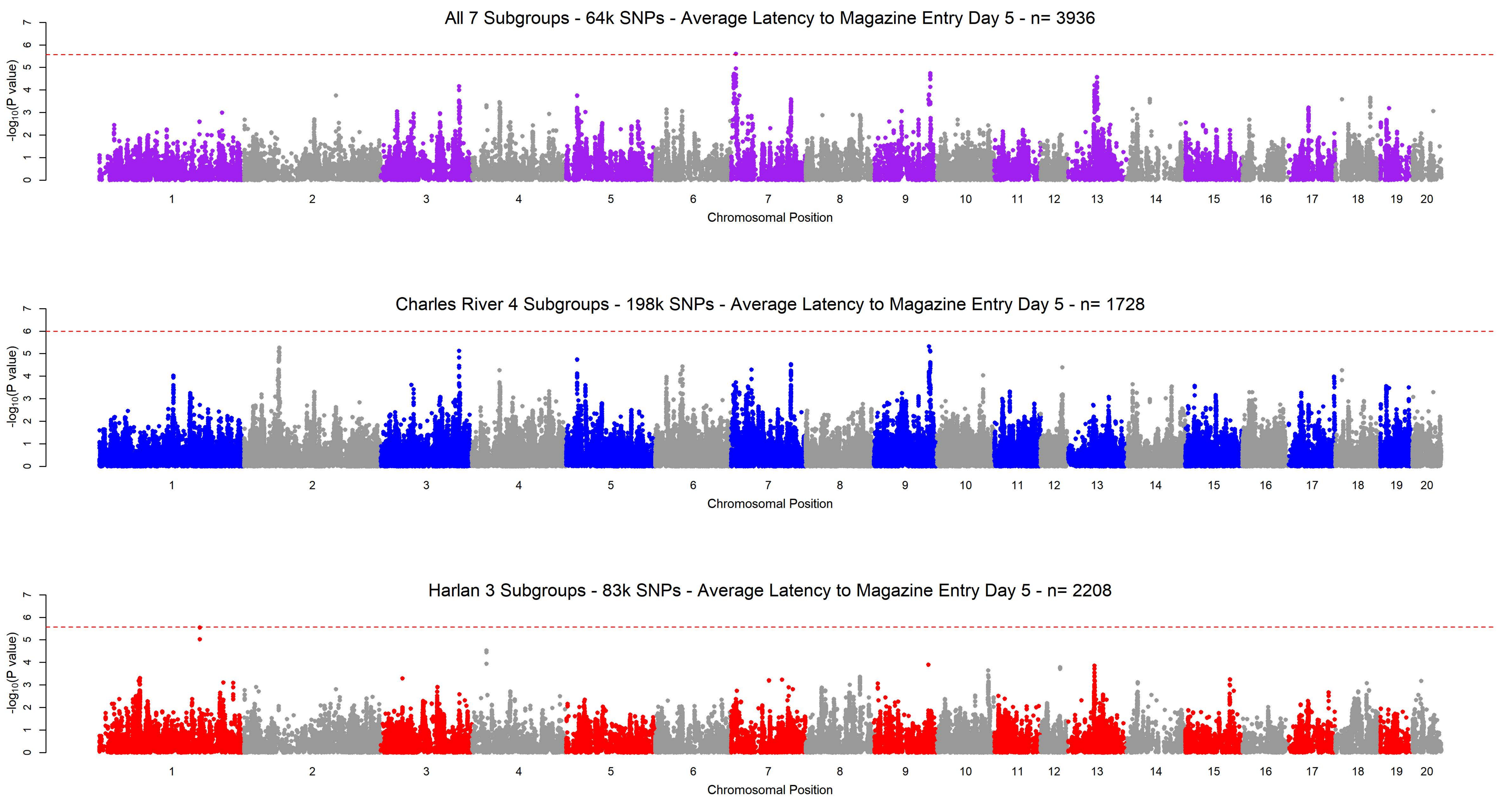
All 7 Subgroups - 64k SNPs - Average Latency to Magazine Entry Day 5 - n= 3936
Charles River 4 Subgroups - 198k SNPs - Average Latency to Magazine Entry Day 5 - n= 1728
Harlan 3 Subgroups - 83k SNPs - Average Latency to Magazine Entry Day 5 - n= 2208
-log10(P value)
Chromosomal Position
1 2 3 4 5 6 7 8 9 10 11 12 13 14 15 16 17 18 19 20
0 1 2 3 4 5 6 7

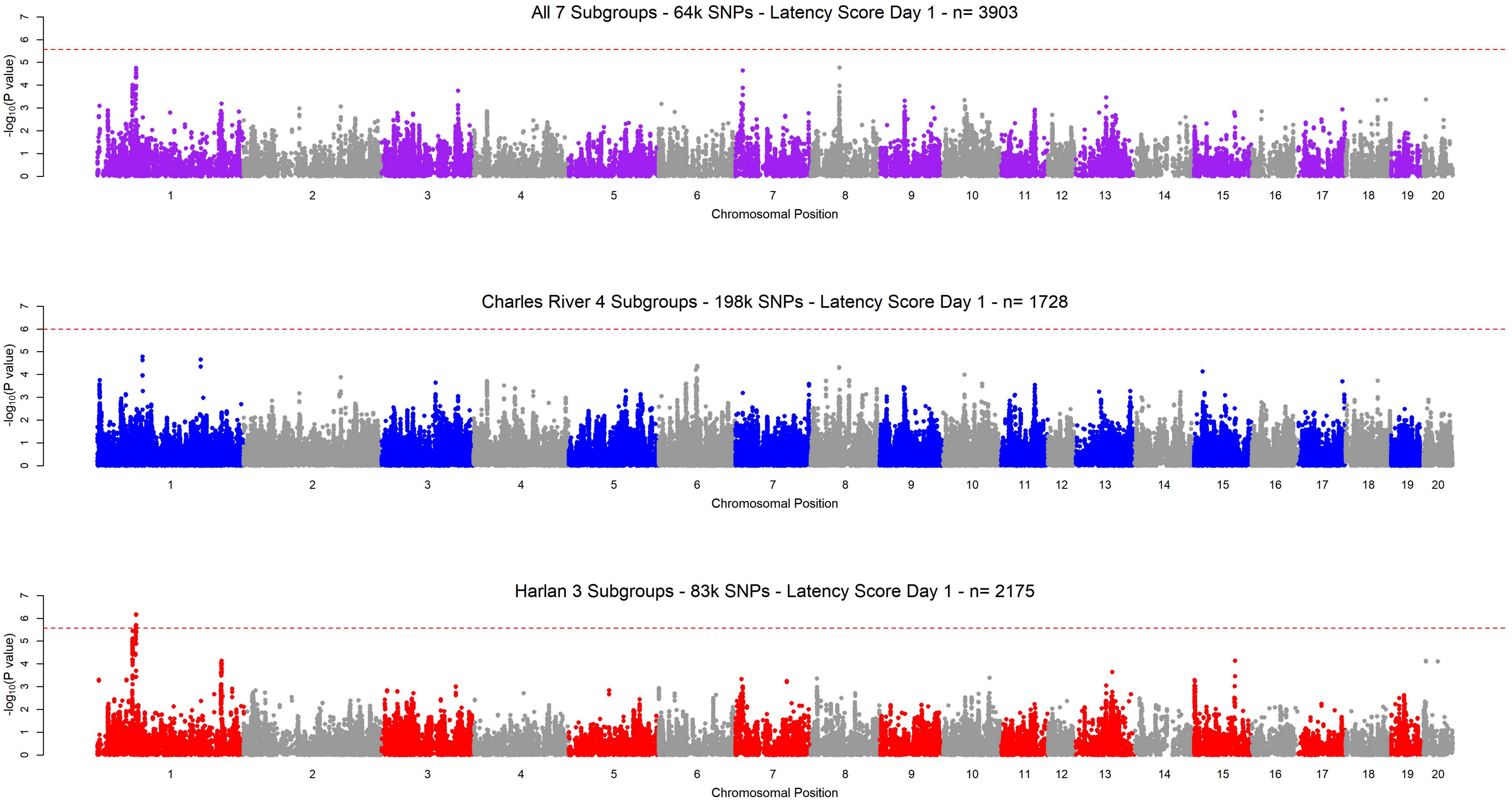
All 7 Subgroups - 64k SNPs - Latency Score Day 1 - n= 3903
Charles River 4 Subgroups - 198k SNPs - Latency Score Day 1 - n= 1728
Harlan 3 Subgroups - 83k SNPs - Latency Score Day 1 - n= 2175
-log10(P value)
Chromosomal Position
1 2 3 4 5 6 7 8 9 10 11 12 13 14 15 16 17 18 19 20
0 1 2 3 4 5 6 7

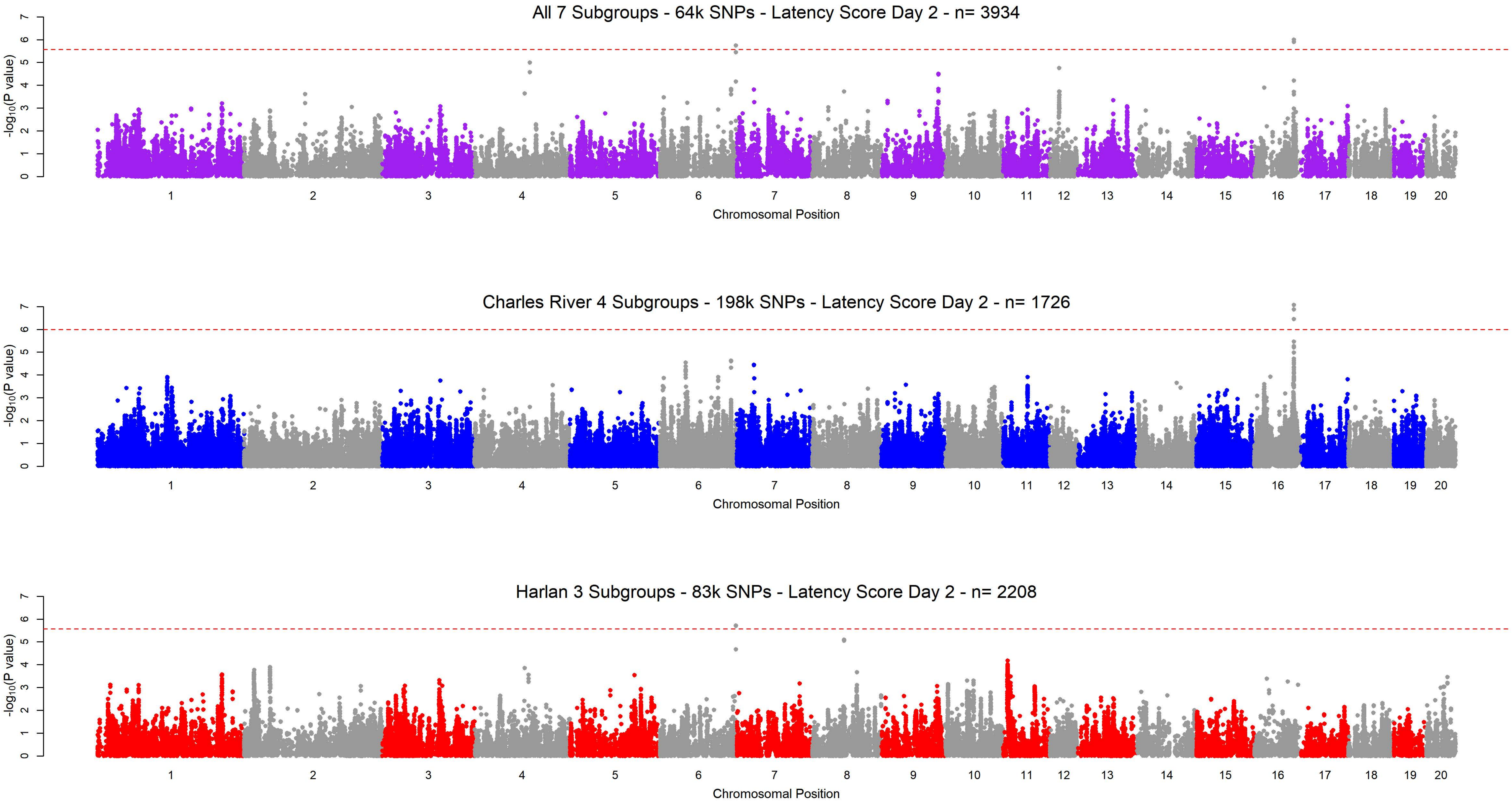

All 7 Subgroups - 64k SNPs - Latency Score Day 2 - n= 3934
Charles River 4 Subgroups - 198k SNPs - Latency Score Day 2 - n= 1726
Harlan 3 Subgroups - 83k SNPs - Latency Score Day 2 - n= 2208
-log10(P value)
Chromosomal Position
0 1 2 3 4 5 6 7
1 2 3 4 5 6 7 8 9 10 11 12 13 14 15 16 17 18 19 20

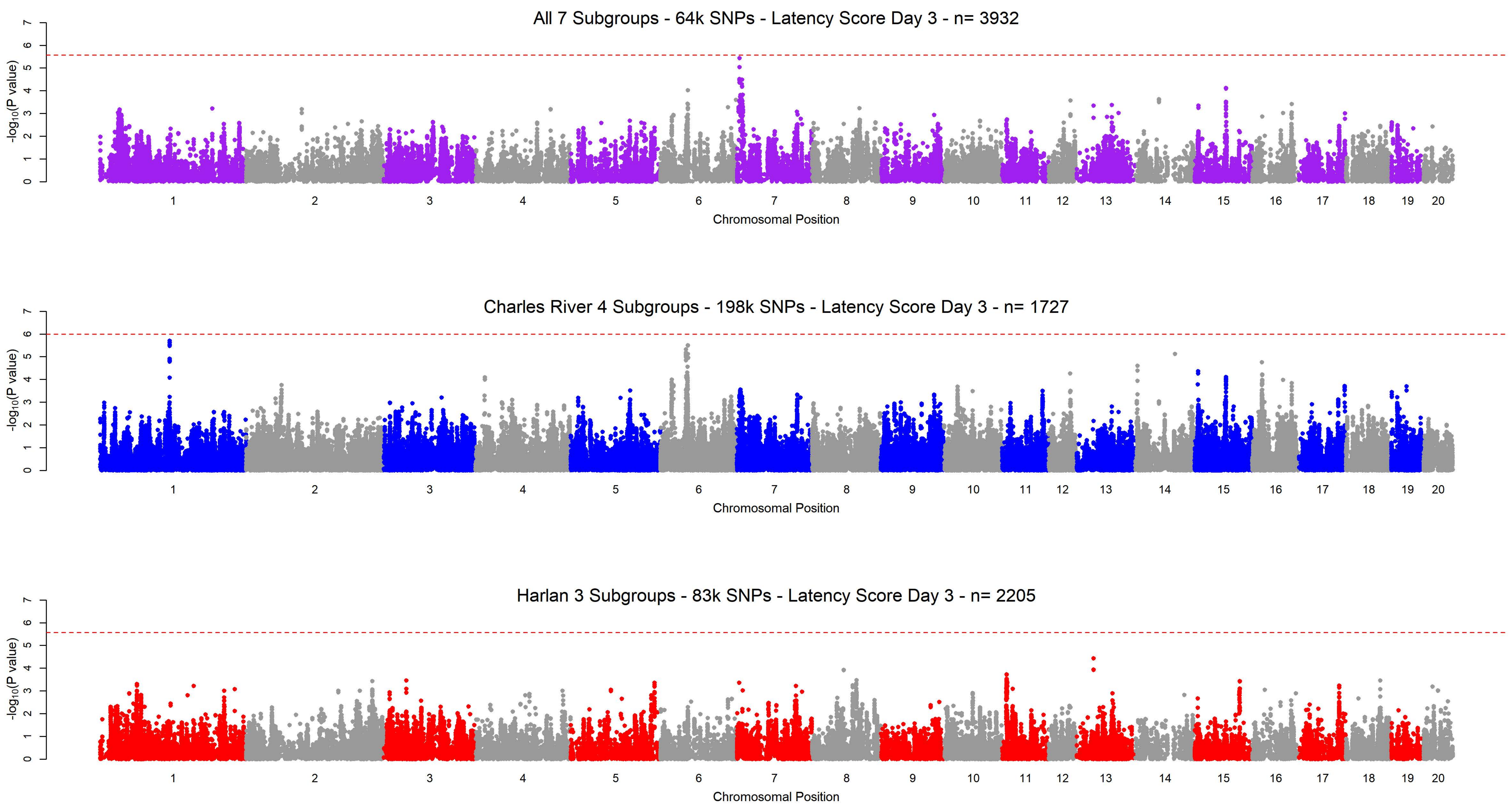

All 7 Subgroups - 64k SNPs - Latency Score Day 3 - n= 3932
Charles River 4 Subgroups - 198k SNPs - Latency Score Day 3 - n= 1727
Harlan 3 Subgroups - 83k SNPs - Latency Score Day 3 - n= 2205
-log10(P value)
Chromosomal Position
1 2 3 4 5 6 7 8 9 10 11 12 13 14 15 16 17 18 19 20
0 1 2 3 4 5 6 7

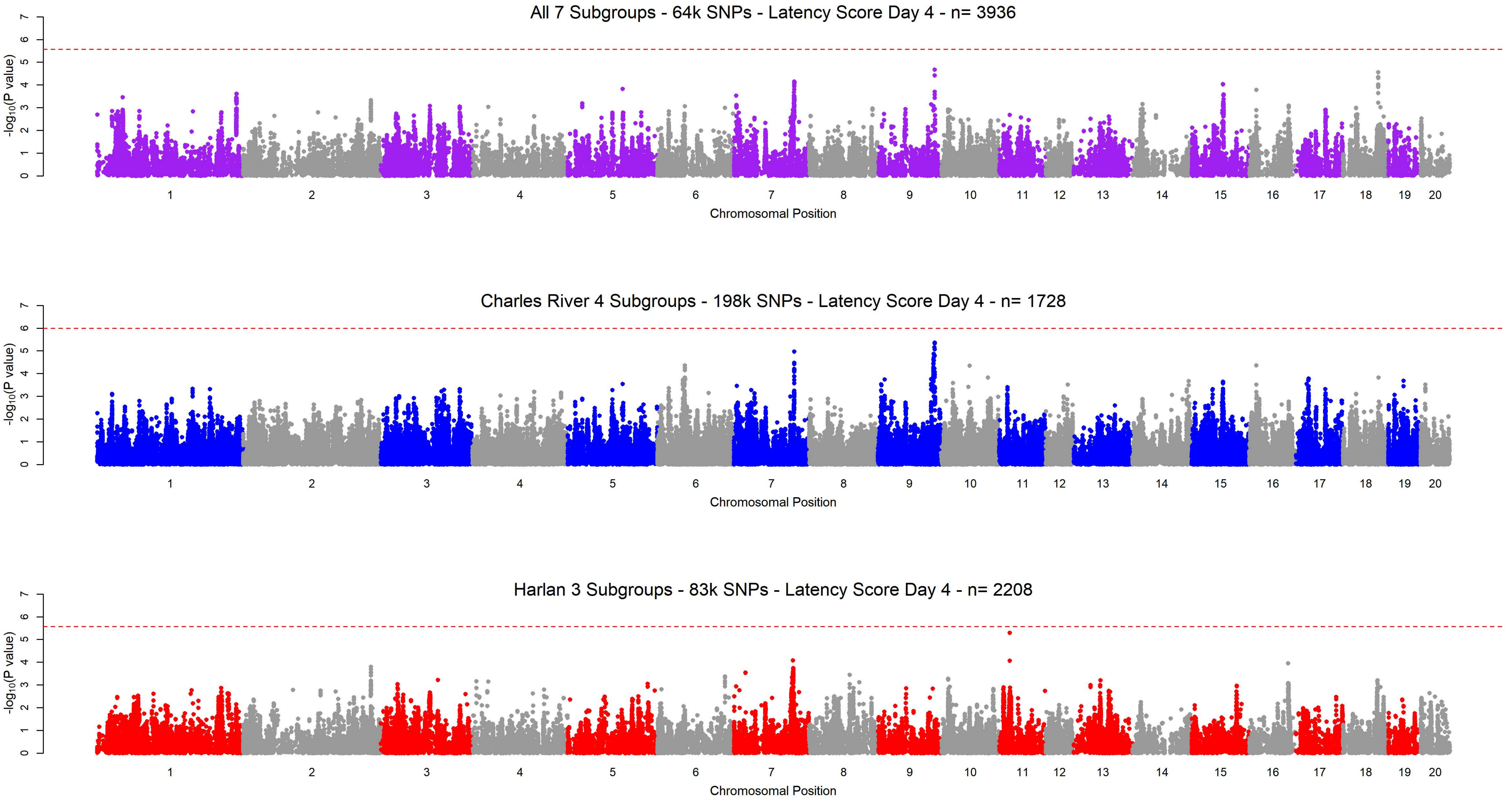
All 7 Subgroups - 64k SNPs - Latency Score Day 4 - n= 3936
Charles River 4 Subgroups - 198k SNPs - Latency Score Day 4 - n= 1728
Harlan 3 Subgroups - 83k SNPs - Latency Score Day 4 - n= 2208
-log10(P value)
0 1 2 3 4 5 6 7
1 2 3 4 5 6 7 8 9 10 11 12 13 14 15 16 17 18 19 20
Chromosomal Position

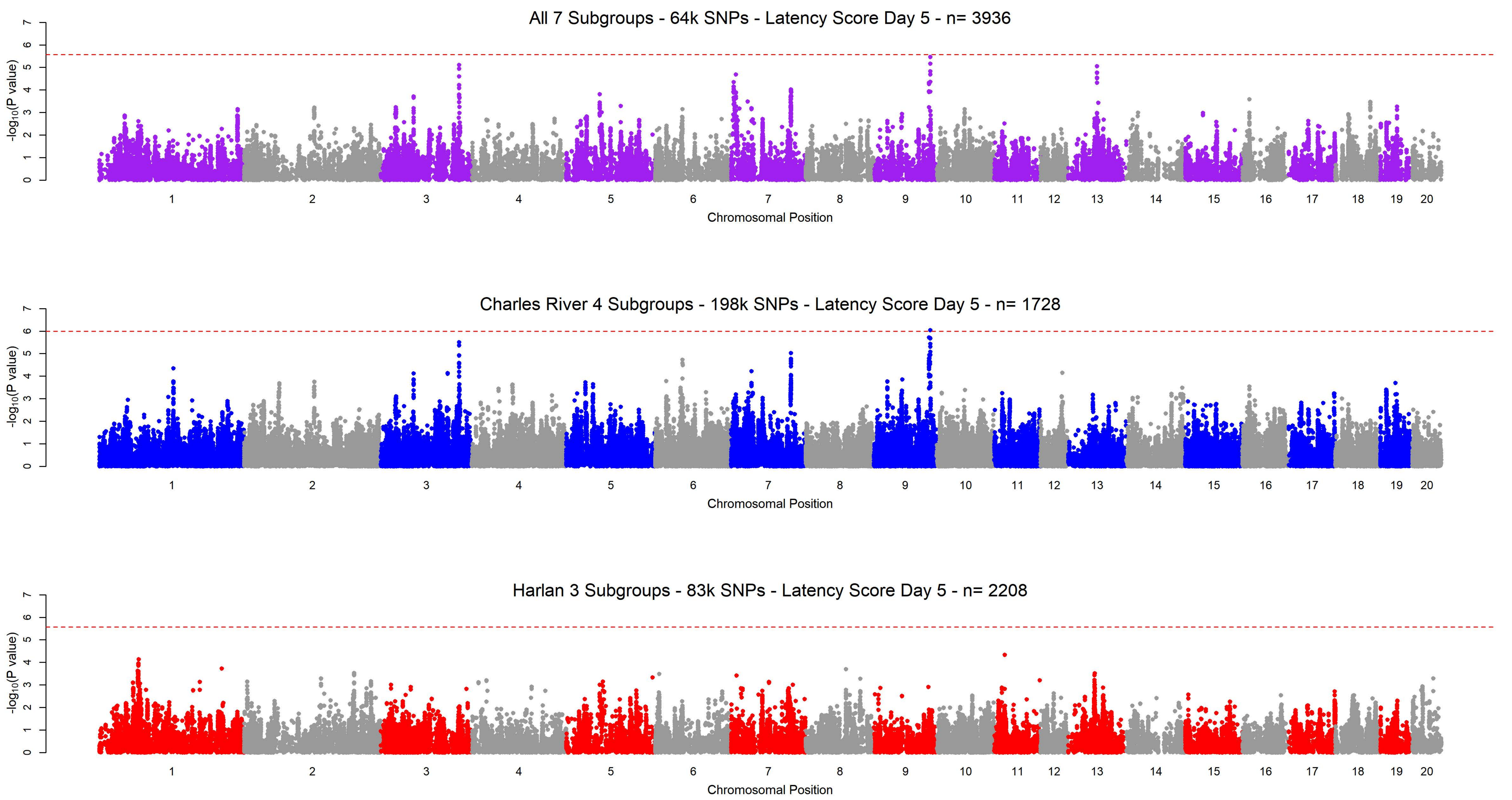

All 7 Subgroups - 64k SNPs - Latency Score Day 5 - n= 3936
Charles River 4 Subgroups - 198k SNPs - Latency Score Day 5 - n= 1728
Harlan 3 Subgroups - 83k SNPs - Latency Score Day 5 - n= 2208
$-\log_{10}$(P value)
Chromosomal Position
0 1 2 3 4 5 6 7
1 2 3 4 5 6 7 8 9 10 11 12 13 14 15 16 17 18 19 20

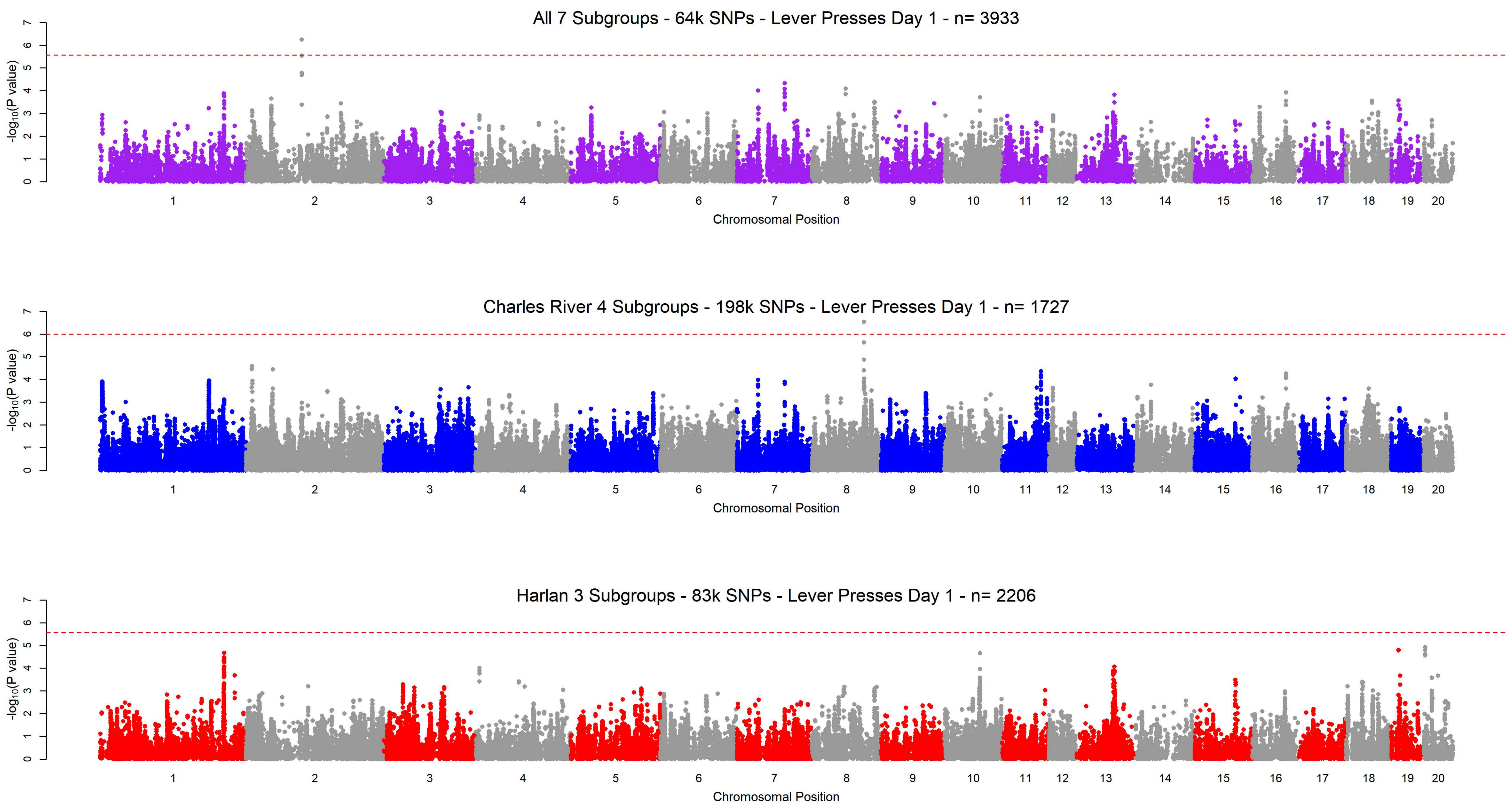
All 7 Subgroups - 64k SNPs - Lever Presses Day 1 - n= 3933
Charles River 4 Subgroups - 198k SNPs - Lever Presses Day 1 - n= 1727
Harlan 3 Subgroups - 83k SNPs - Lever Presses Day 1 - n= 2206
-log10(P value)
Chromosomal Position
0
1
2
3
4
5
6
7

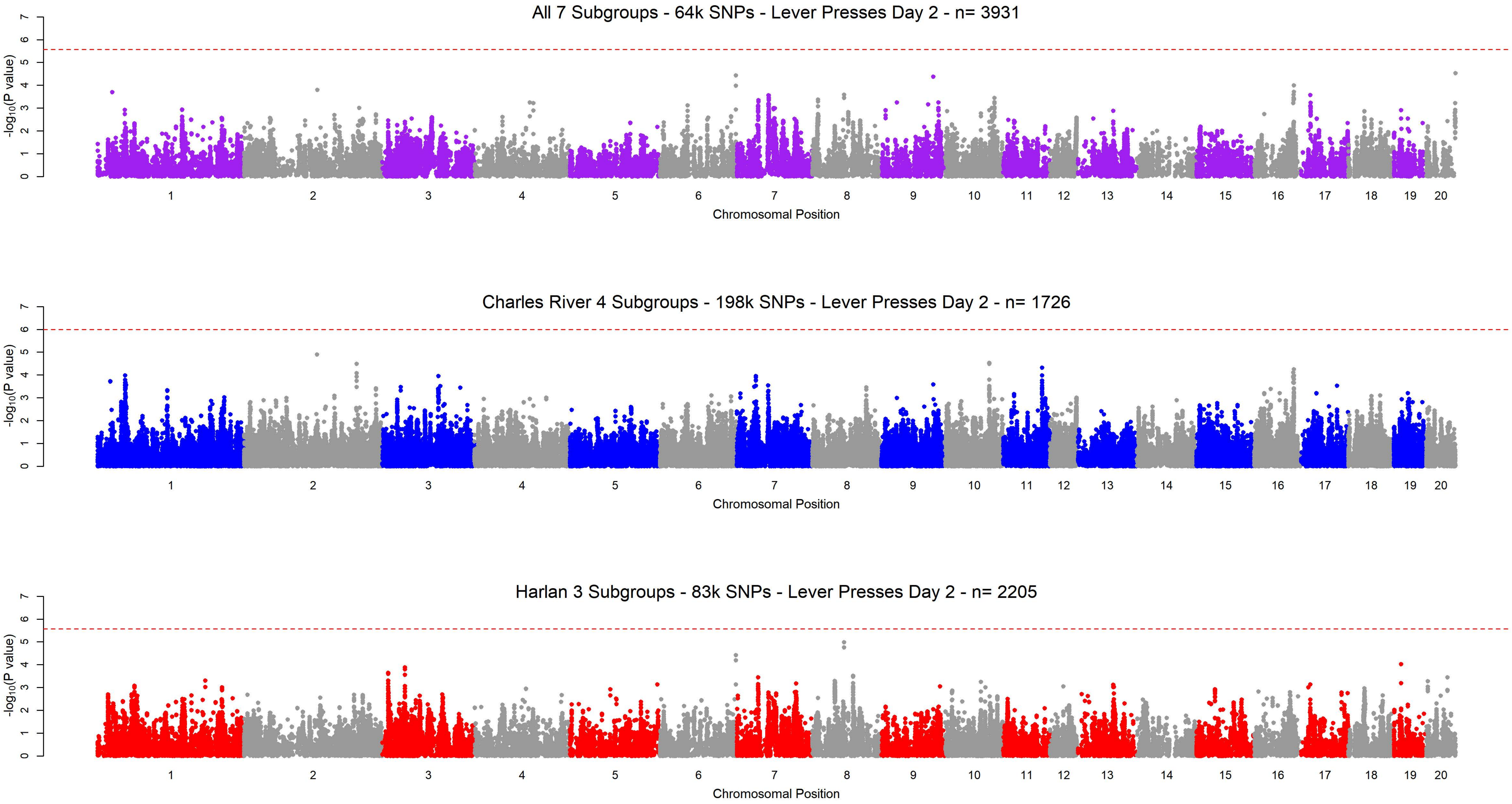

All 7 Subgroups - 64k SNPs - Lever Presses Day 2 - n= 3931
-log10(P value)
Chromosomal Position
Charles River 4 Subgroups - 198k SNPs - Lever Presses Day 2 - n= 1726
-log10(P value)
Chromosomal Position
Harlan 3 Subgroups - 83k SNPs - Lever Presses Day 2 - n= 2205
-log10(P value)
Chromosomal Position

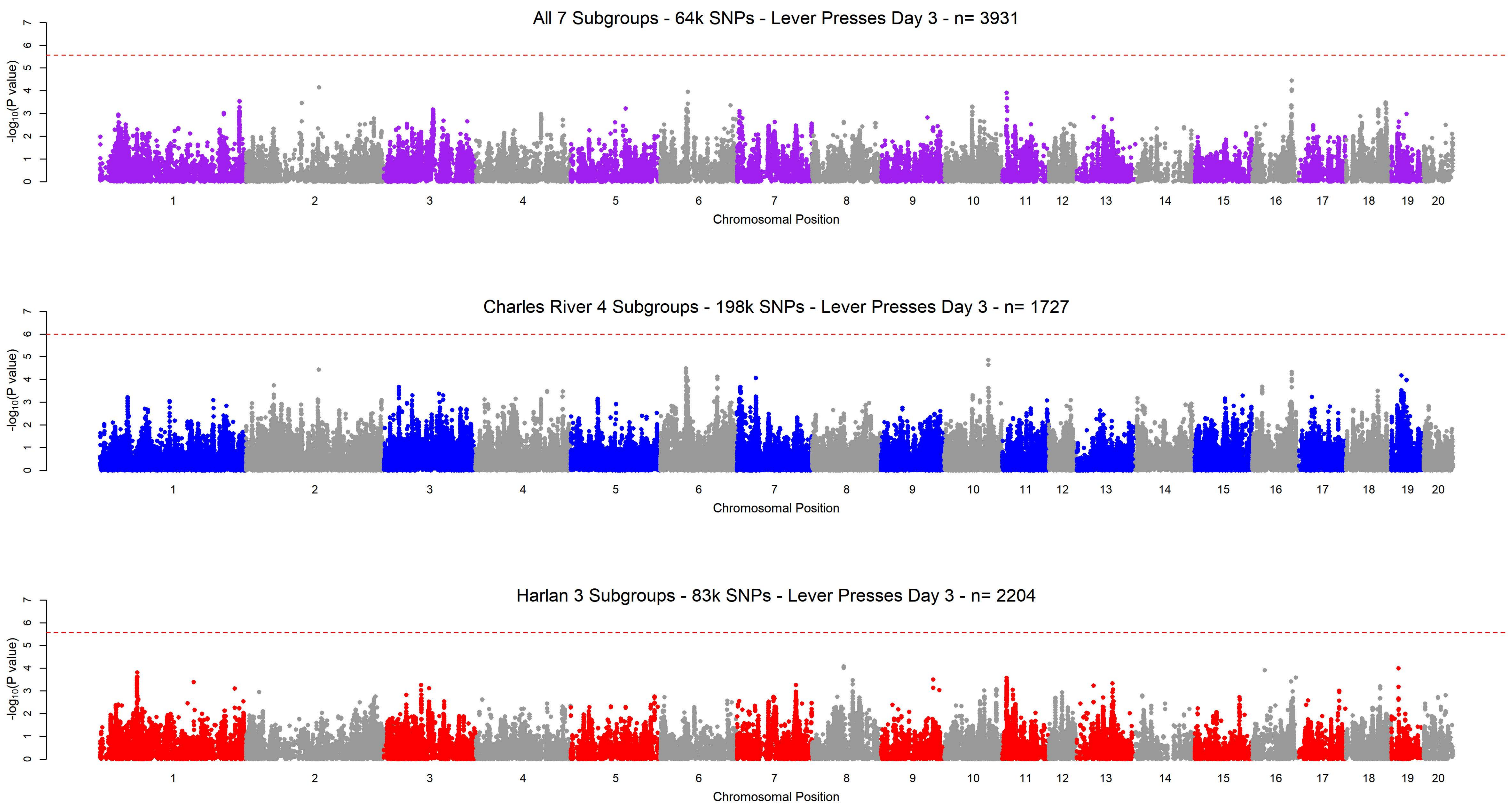
All 7 Subgroups - 64k SNPs - Lever Presses Day 3 - n= 3931
Charles River 4 Subgroups - 198k SNPs - Lever Presses Day 3 - n= 1727
Harlan 3 Subgroups - 83k SNPs - Lever Presses Day 3 - n= 2204
-log10(P value)
Chromosomal Position
1 2 3 4 5 6 7 8 9 10 11 12 13 14 15 16 17 18 19 20

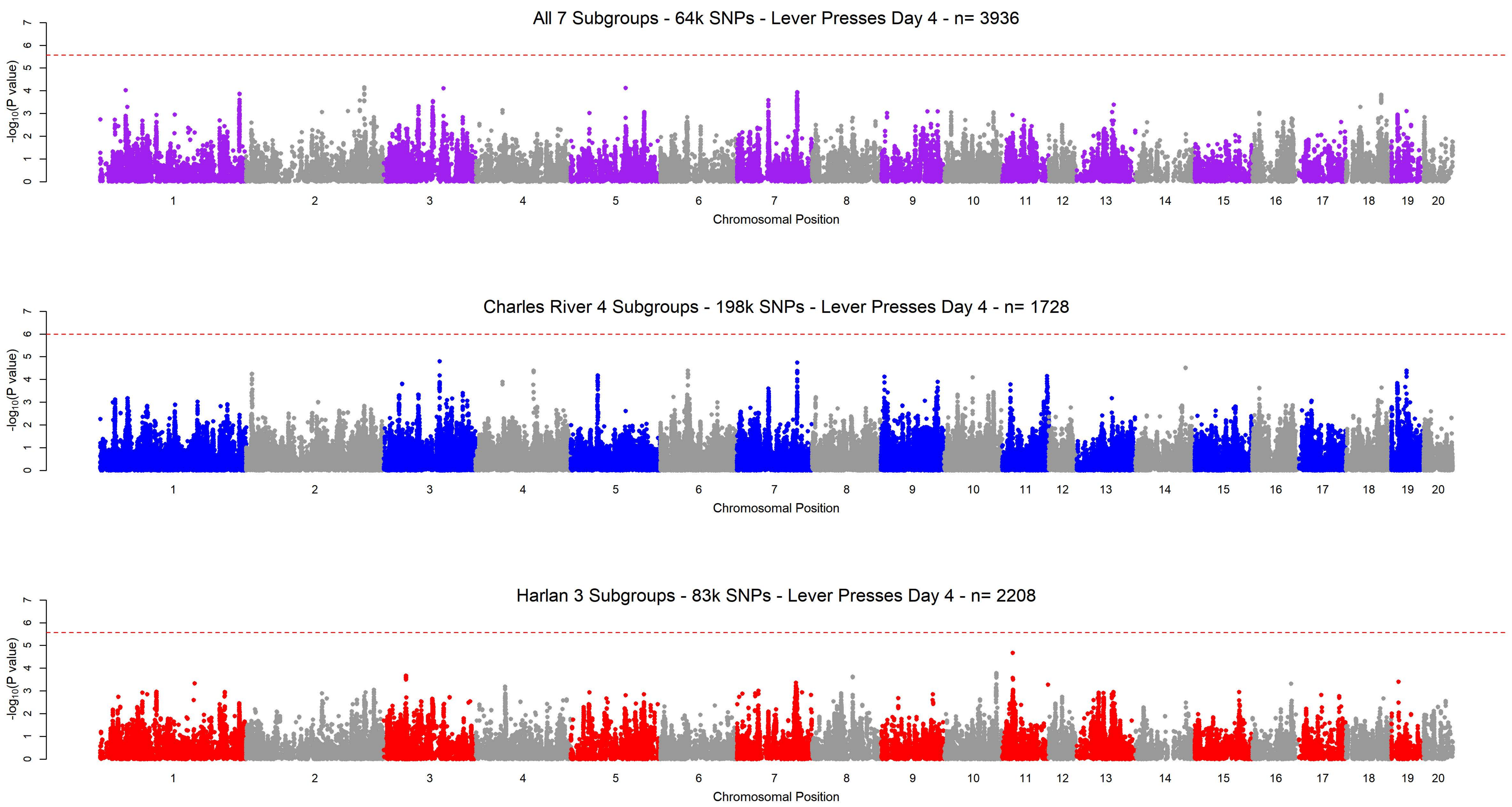

All 7 Subgroups - 64k SNPs - Lever Presses Day 4 - n= 3936
-log10(P value)
0 1 2 3 4 5 6 7
1 2 3 4 5 6 7 8 9 10 11 12 13 14 15 16 17 18 19 20
Chromosomal Position
Charles River 4 Subgroups - 198k SNPs - Lever Presses Day 4 - n= 1728
-log10(P value)
0 1 2 3 4 5 6 7
1 2 3 4 5 6 7 8 9 10 11 12 13 14 15 16 17 18 19 20
Chromosomal Position
Harlan 3 Subgroups - 83k SNPs - Lever Presses Day 4 - n= 2208
-log10(P value)
0 1 2 3 4 5 6 7
1 2 3 4 5 6 7 8 9 10 11 12 13 14 15 16 17 18 19 20
Chromosomal Position

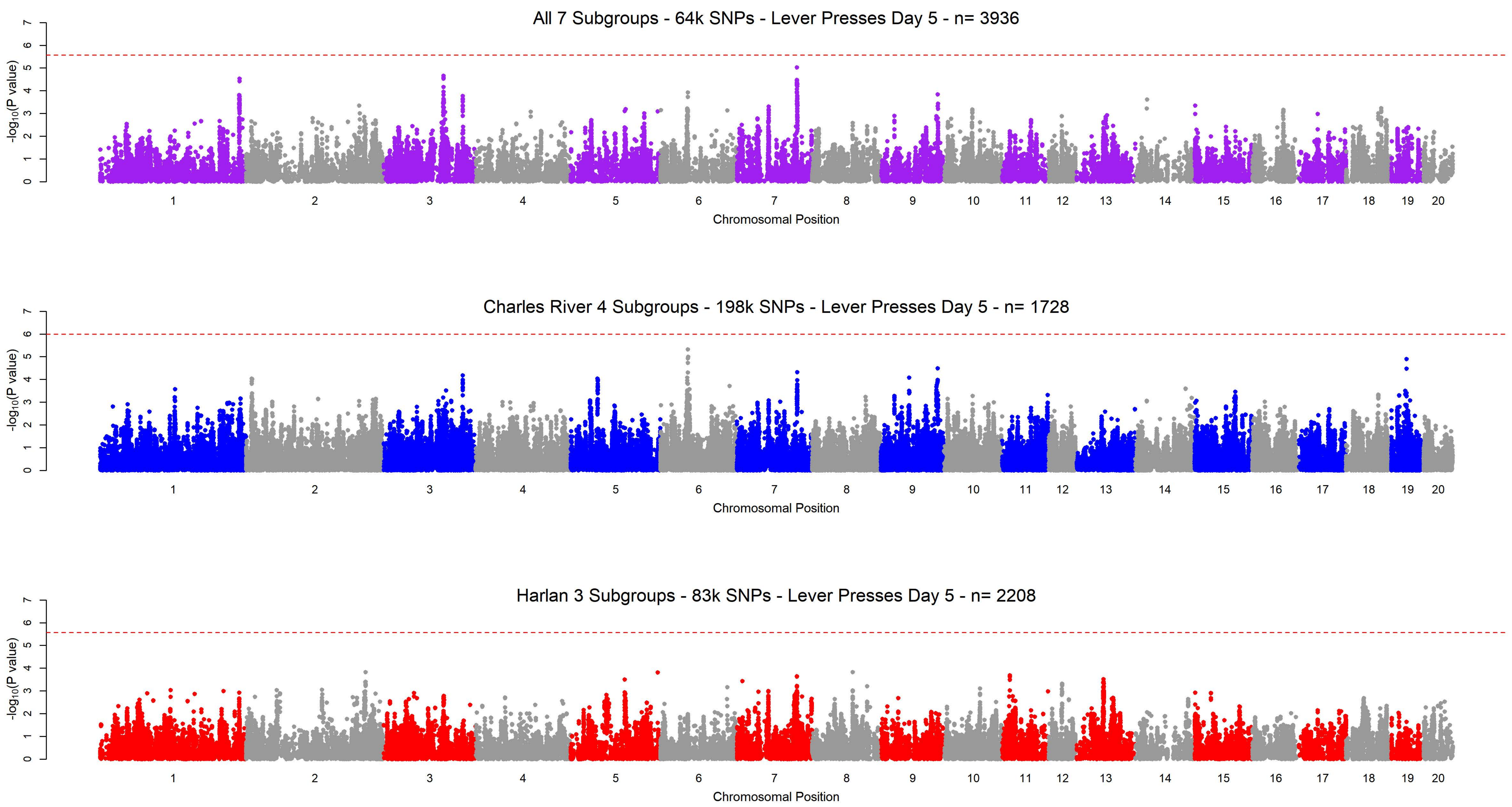
All 7 Subgroups - 64k SNPs - Lever Presses Day 5 - n= 3936
Charles River 4 Subgroups - 198k SNPs - Lever Presses Day 5 - n= 1728
Harlan 3 Subgroups - 83k SNPs - Lever Presses Day 5 - n= 2208
-log10(P value)
Chromosomal Position
1 2 3 4 5 6 7 8 9 10 11 12 13 14 15 16 17 18 19 20
0 1 2 3 4 5 6 7

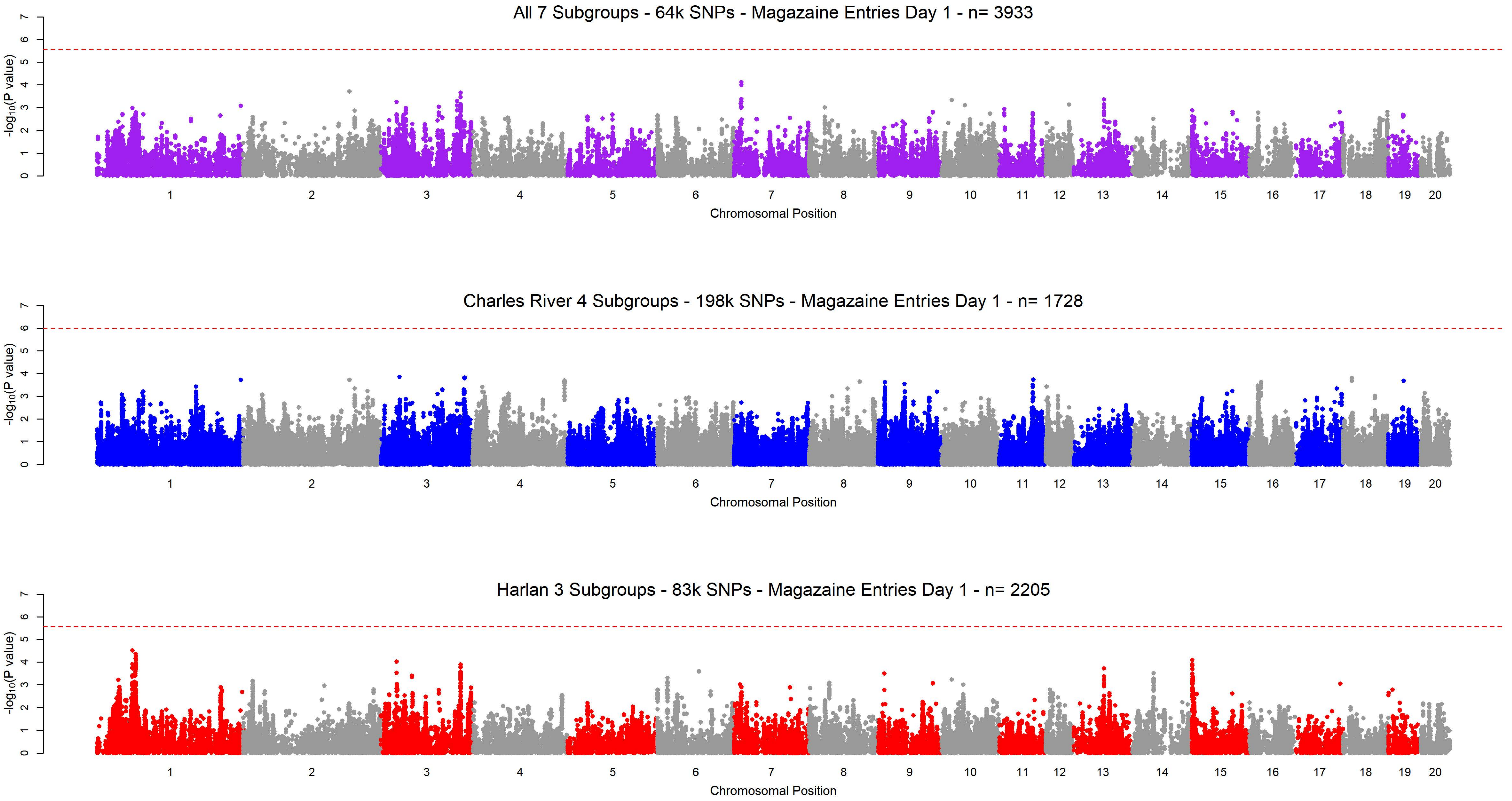
All 7 Subgroups - 64k SNPs - Magazaine Entries Day 1 - n= 3933
Charles River 4 Subgroups - 198k SNPs - Magazaine Entries Day 1 - n= 1728
Harlan 3 Subgroups - 83k SNPs - Magazaine Entries Day 1 - n= 2205
$-\log_{10}$(P value)
Chromosomal Position
1 2 3 4 5 6 7 8 9 10 11 12 13 14 15 16 17 18 19 20

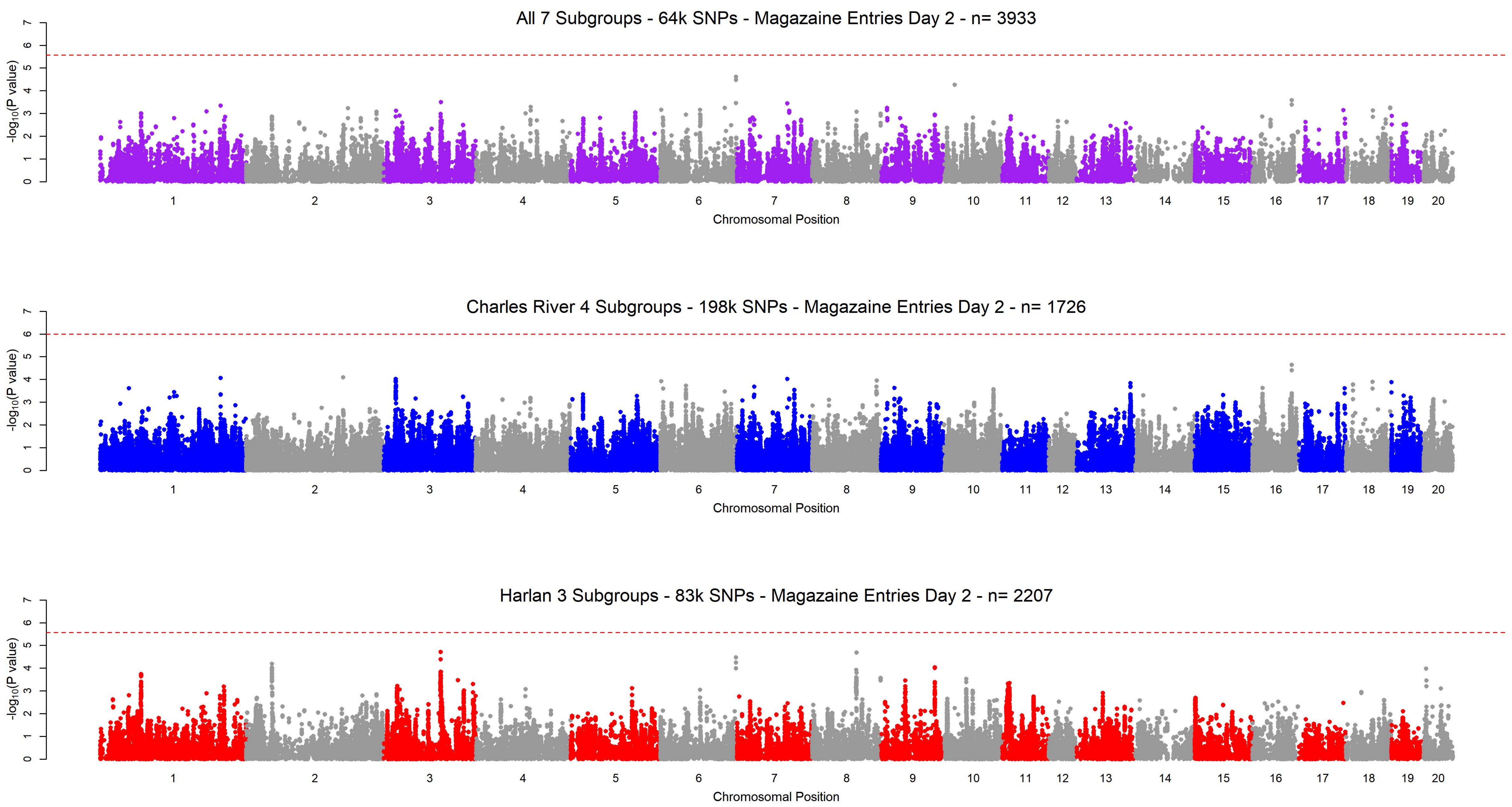

All 7 Subgroups - 64k SNPs - Magazaine Entries Day 2 - n= 3933
-log10(P value)
0
1
2
3
4
5
6
7
Chromosomal Position
Charles River 4 Subgroups - 198k SNPs - Magazaine Entries Day 2 - n= 1726
-log10(P value)
Chromosomal Position
Harlan 3 Subgroups - 83k SNPs - Magazaine Entries Day 2 - n= 2207
-log10(P value)
Chromosomal Position

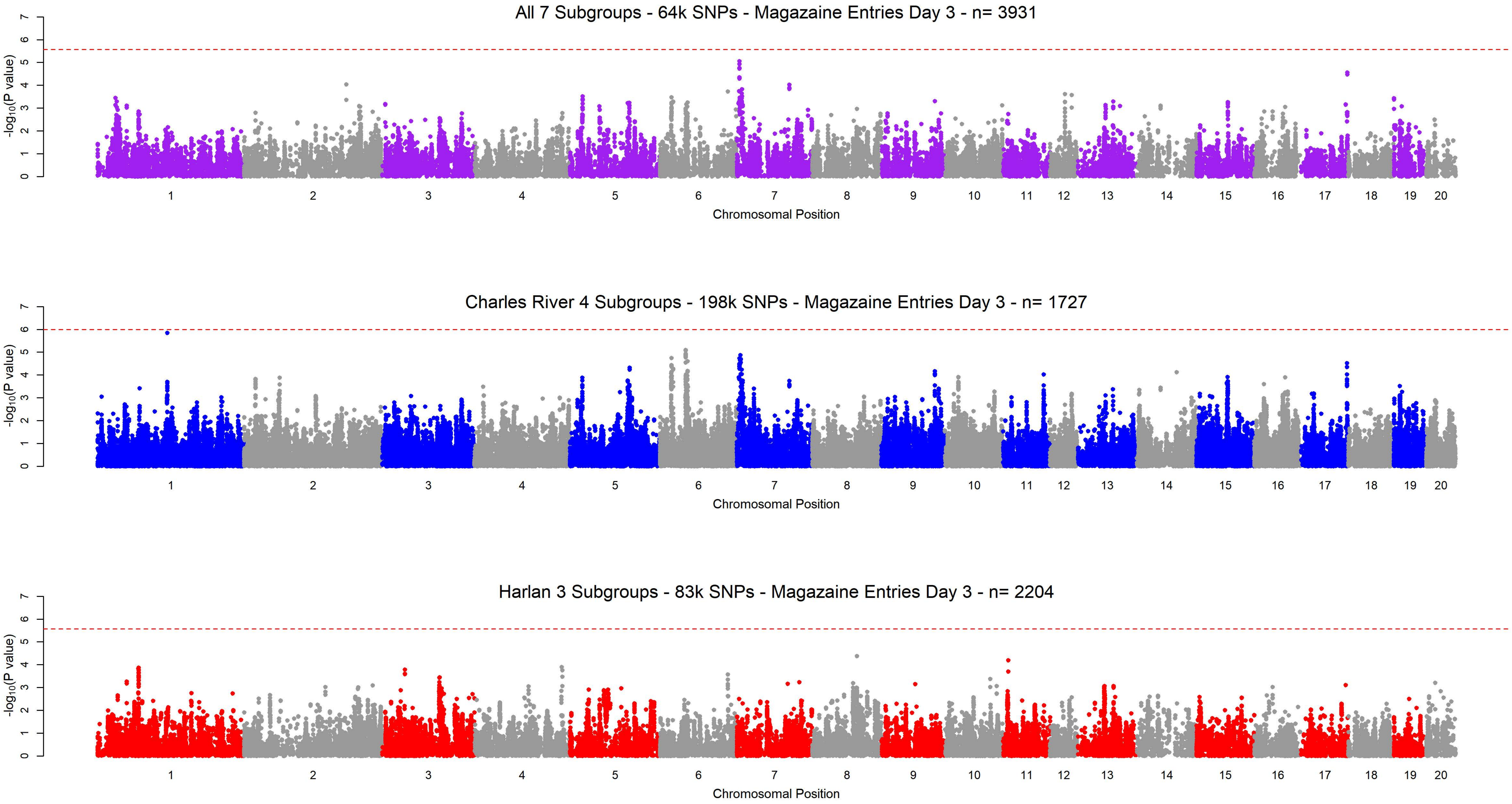

All 7 Subgroups - 64k SNPs - Magazaine Entries Day 3 - n= 3931
Charles River 4 Subgroups - 198k SNPs - Magazaine Entries Day 3 - n= 1727
Harlan 3 Subgroups - 83k SNPs - Magazaine Entries Day 3 - n= 2204
-log10(P value)
0 1 2 3 4 5 6 7
1 2 3 4 5 6 7 8 9 10 11 12 13 14 15 16 17 18 19 20
Chromosomal Position

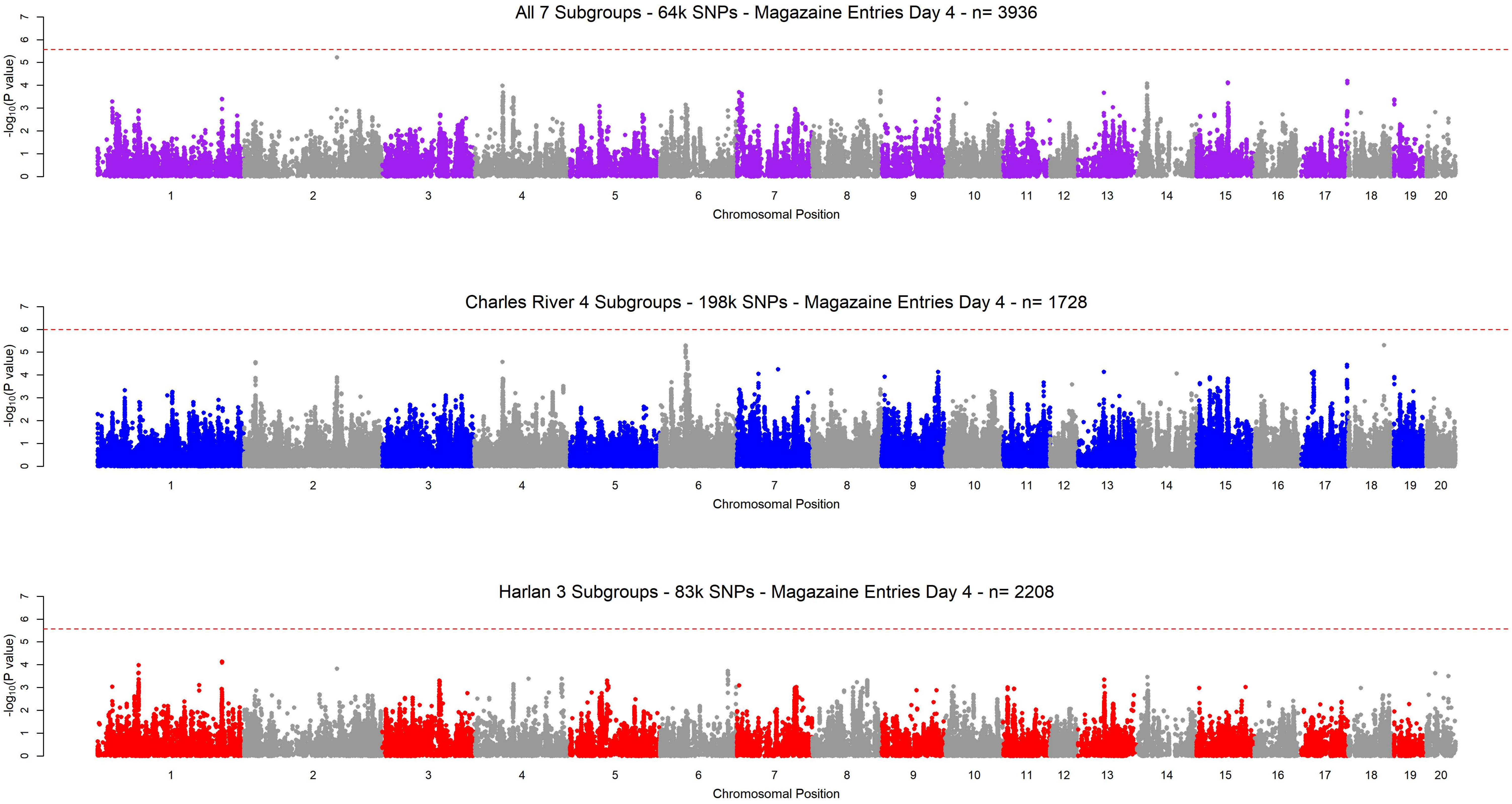
All 7 Subgroups - 64k SNPs - Magazaine Entries Day 4 - n= 3936
Charles River 4 Subgroups - 198k SNPs - Magazaine Entries Day 4 - n= 1728
Harlan 3 Subgroups - 83k SNPs - Magazaine Entries Day 4 - n= 2208
-log10(P value)
0 1 2 3 4 5 6 7
1 2 3 4 5 6 7 8 9 10 11 12 13 14 15 16 17 18 19 20
Chromosomal Position

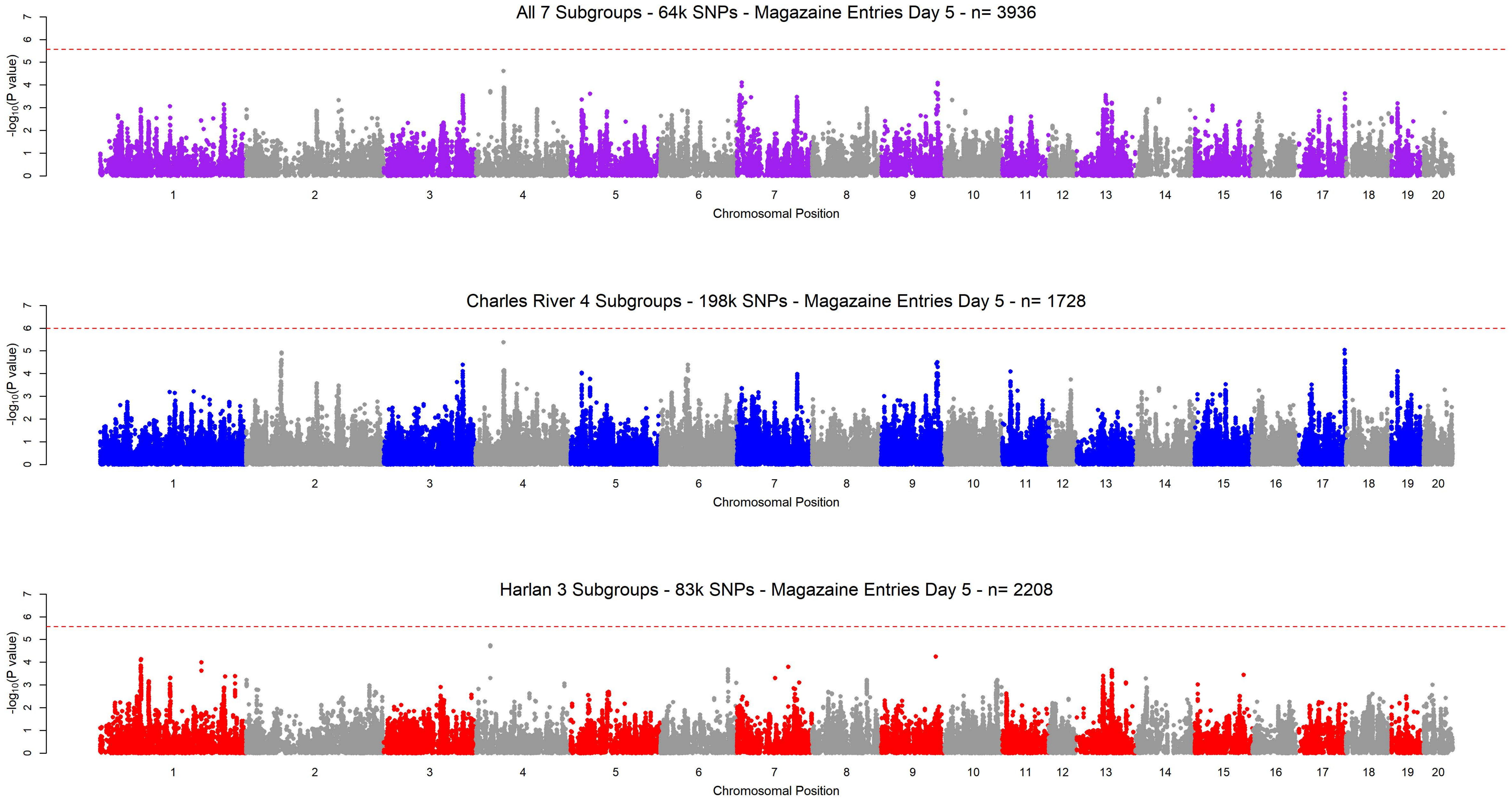
All 7 Subgroups - 64k SNPs - Magazaine Entries Day 5 - n= 3936
Charles River 4 Subgroups - 198k SNPs - Magazaine Entries Day 5 - n= 1728
Harlan 3 Subgroups - 83k SNPs - Magazaine Entries Day 5 - n= 2208
-log10(P value)
Chromosomal Position

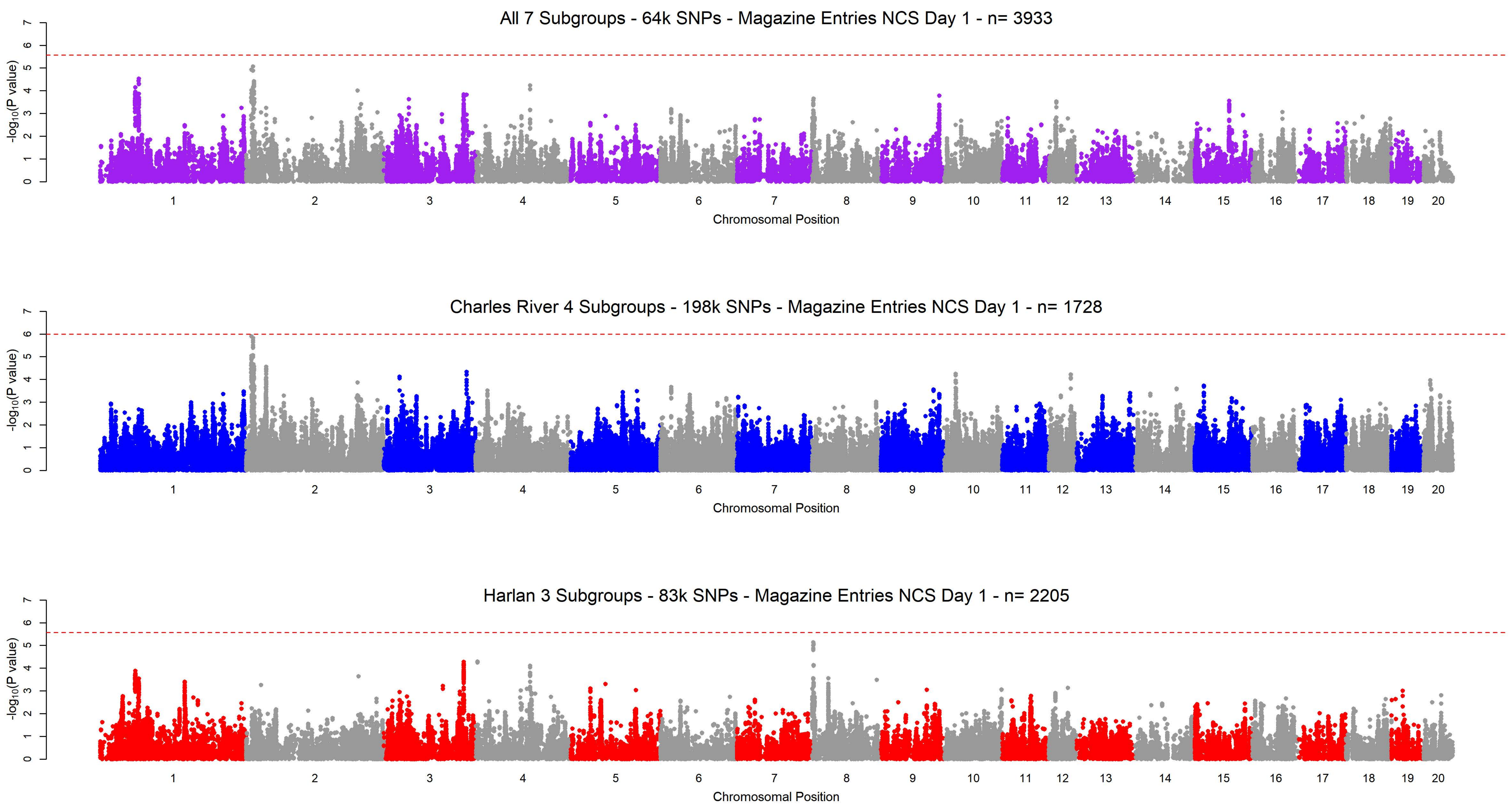

All 7 Subgroups - 64k SNPs - Magazine Entries NCS Day 1 - n= 3933
Charles River 4 Subgroups - 198k SNPs - Magazine Entries NCS Day 1 - n= 1728
Harlan 3 Subgroups - 83k SNPs - Magazine Entries NCS Day 1 - n= 2205
$-\log_{10}$(P value)
Chromosomal Position
0 1 2 3 4 5 6 7
1 2 3 4 5 6 7 8 9 10 11 12 13 14 15 16 17 18 19 20

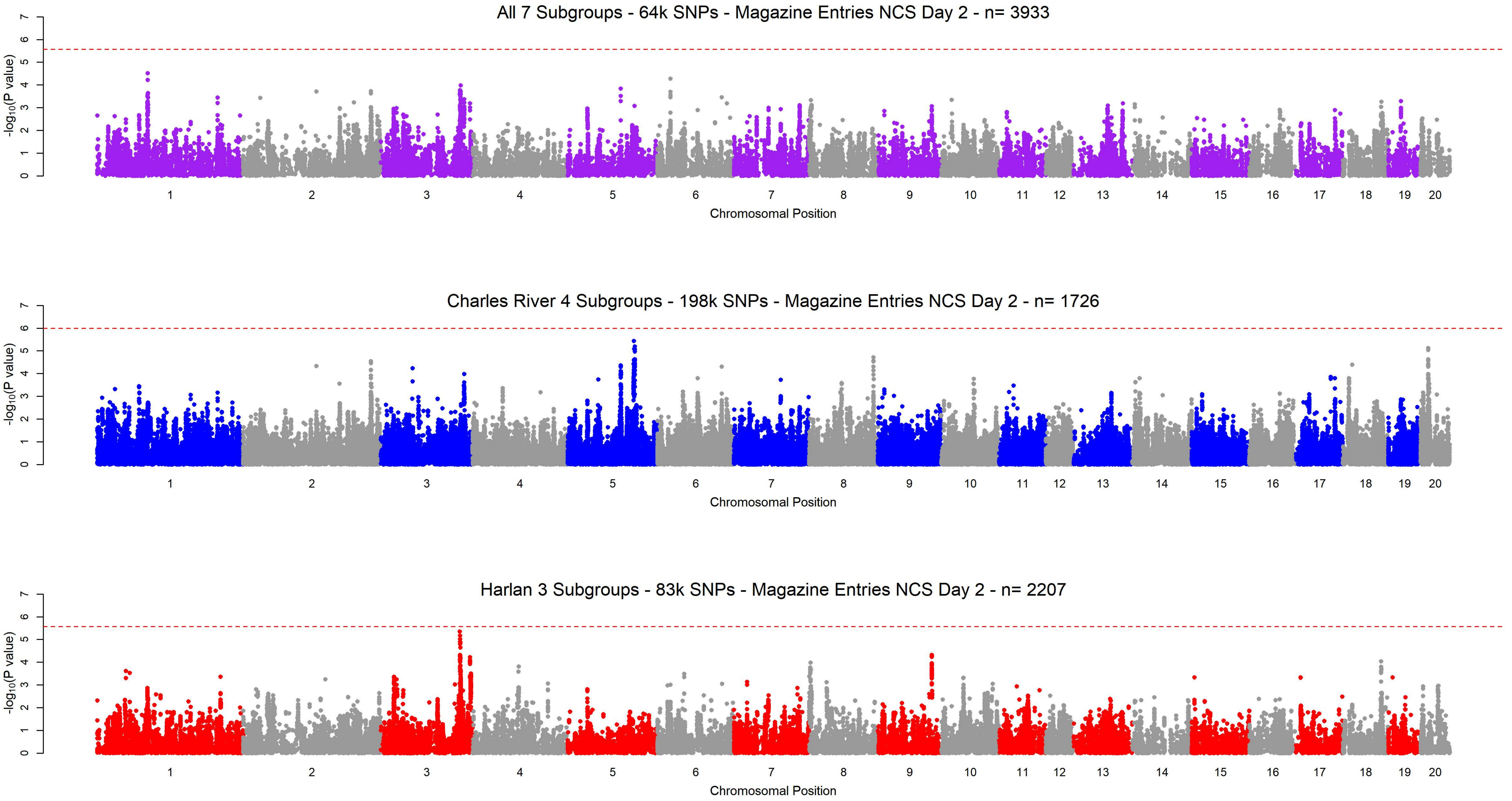
All 7 Subgroups - 64k SNPs - Magazine Entries NCS Day 2 - n= 3933
Charles River 4 Subgroups - 198k SNPs - Magazine Entries NCS Day 2 - n= 1726
Harlan 3 Subgroups - 83k SNPs - Magazine Entries NCS Day 2 - n= 2207
-log10(P value)
Chromosomal Position
0 1 2 3 4 5 6 7
1 2 3 4 5 6 7 8 9 10 11 12 13 14 15 16 17 18 19 20

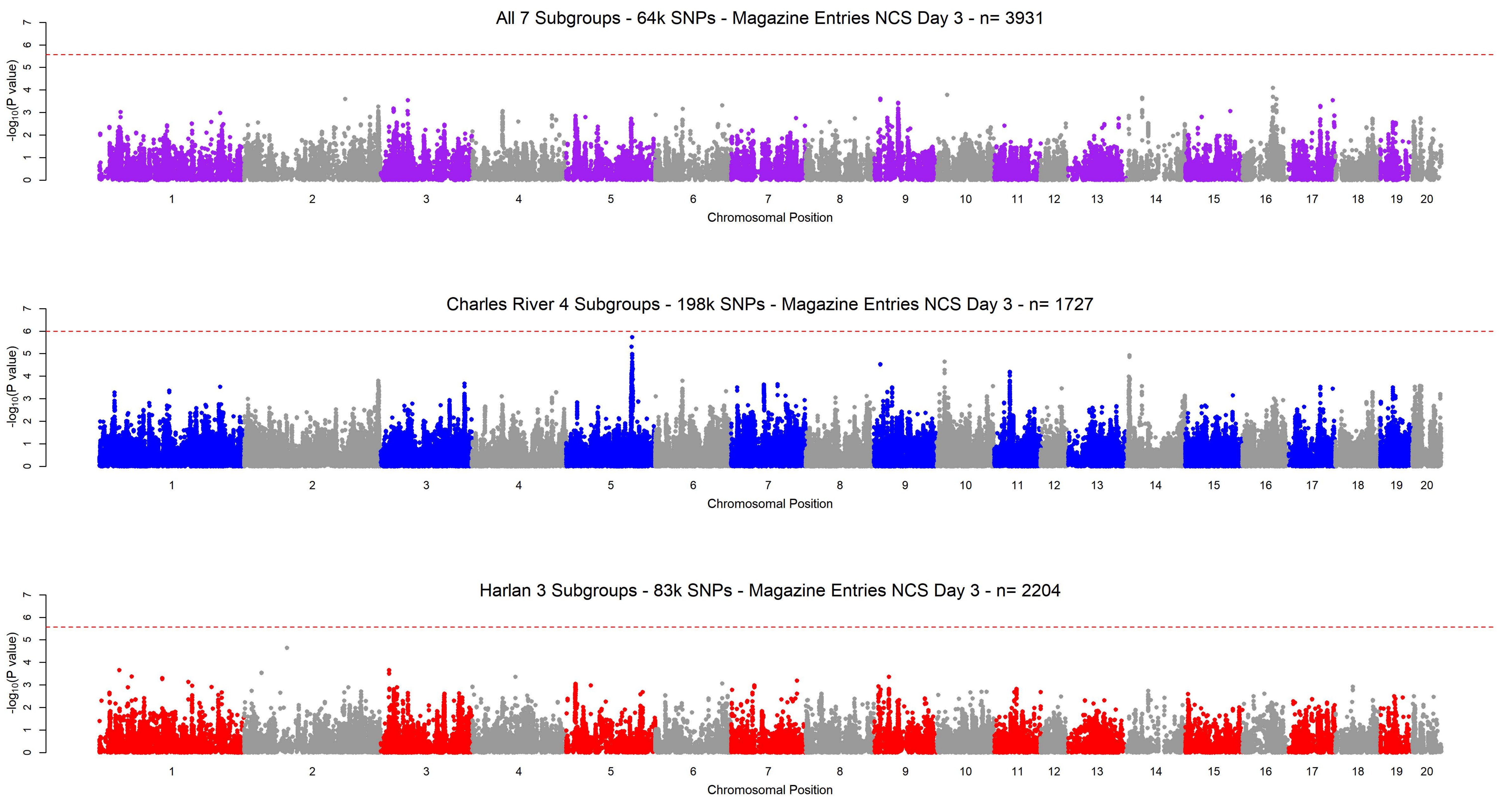
All 7 Subgroups - 64k SNPs - Magazine Entries NCS Day 3 - n= 3931
Charles River 4 Subgroups - 198k SNPs - Magazine Entries NCS Day 3 - n= 1727
Harlan 3 Subgroups - 83k SNPs - Magazine Entries NCS Day 3 - n= 2204
-log10(P value)
Chromosomal Position
1 2 3 4 5 6 7 8 9 10 11 12 13 14 15 16 17 18 19 20

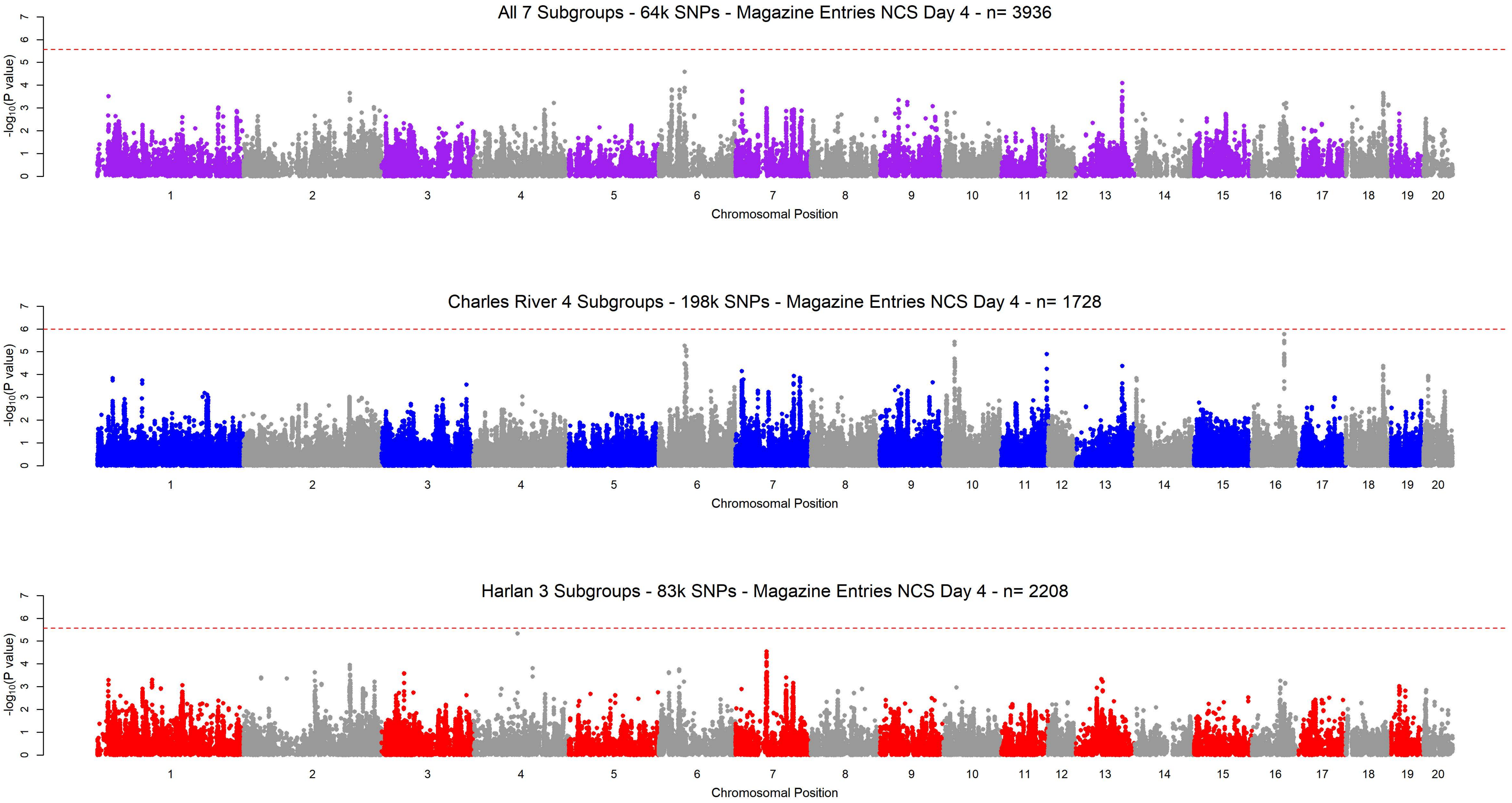
All 7 Subgroups - 64k SNPs - Magazine Entries NCS Day 4 - n= 3936
Charles River 4 Subgroups - 198k SNPs - Magazine Entries NCS Day 4 - n= 1728
Harlan 3 Subgroups - 83k SNPs - Magazine Entries NCS Day 4 - n= 2208
$-\log_{10}$(P value)
Chromosomal Position
0
1
2
3
4
5
6
7

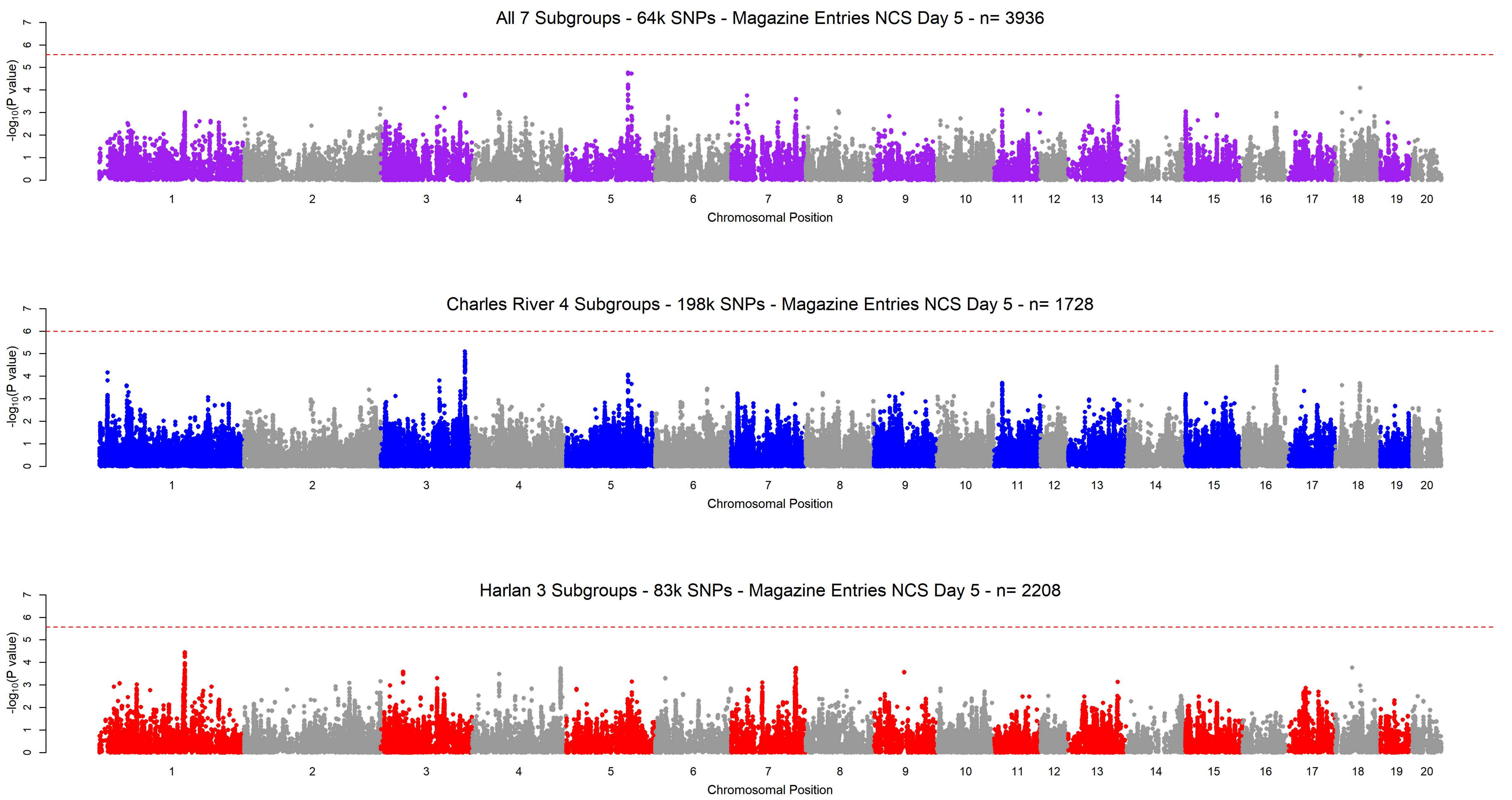
All 7 Subgroups - 64k SNPs - Magazine Entries NCS Day 5 - n= 3936
Charles River 4 Subgroups - 198k SNPs - Magazine Entries NCS Day 5 - n= 1728
Harlan 3 Subgroups - 83k SNPs - Magazine Entries NCS Day 5 - n= 2208
-log10(P value)
Chromosomal Position
0 1 2 3 4 5 6 7
1 2 3 4 5 6 7 8 9 10 11 12 13 14 15 16 17 18 19 20

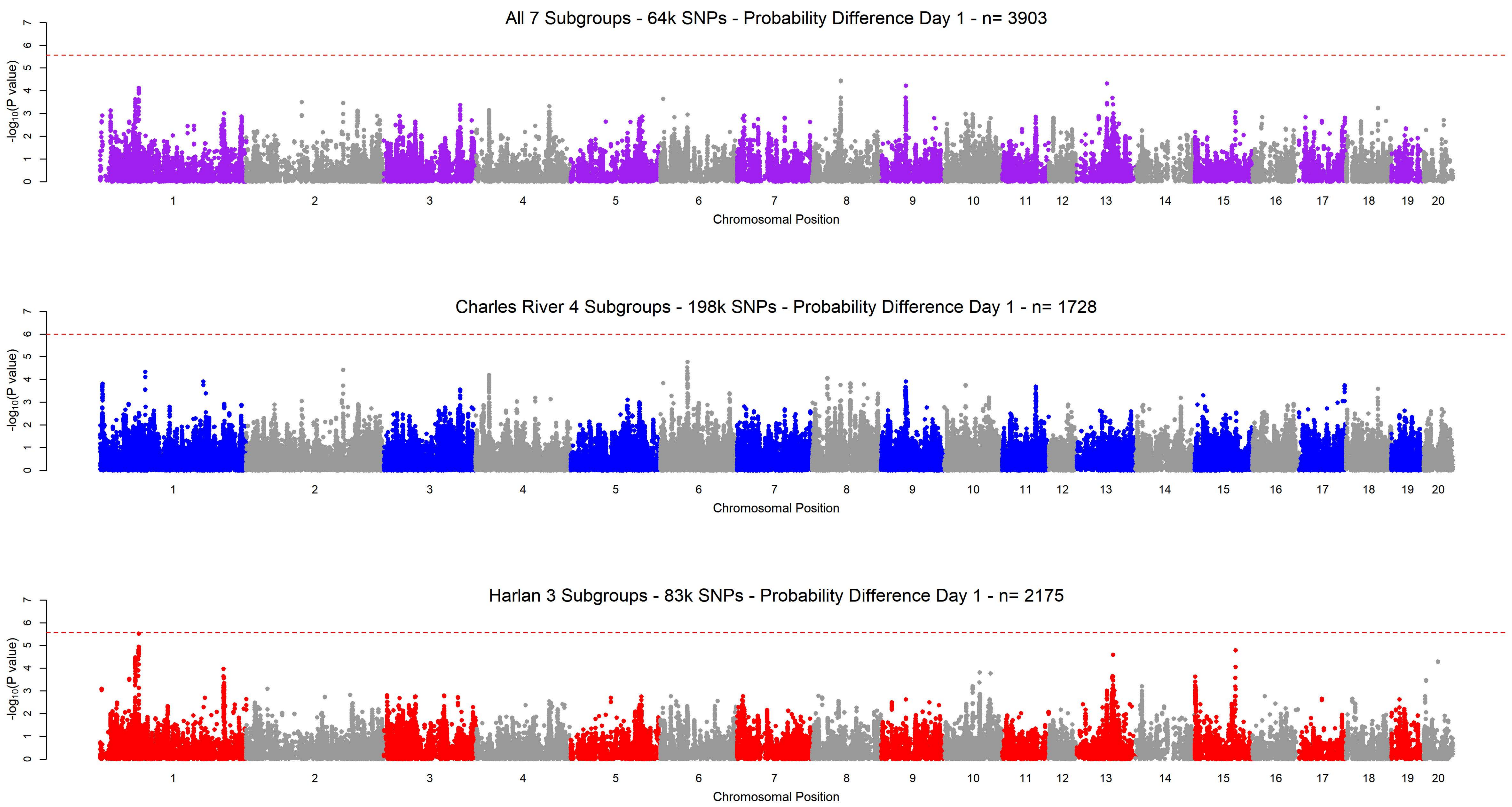
All 7 Subgroups - 64k SNPs - Probability Difference Day 1 - n= 3903
Charles River 4 Subgroups - 198k SNPs - Probability Difference Day 1 - n= 1728
Harlan 3 Subgroups - 83k SNPs - Probability Difference Day 1 - n= 2175
$-\log_{10}$(P value)
Chromosomal Position
1 2 3 4 5 6 7 8 9 10 11 12 13 14 15 16 17 18 19 20
0 1 2 3 4 5 6 7

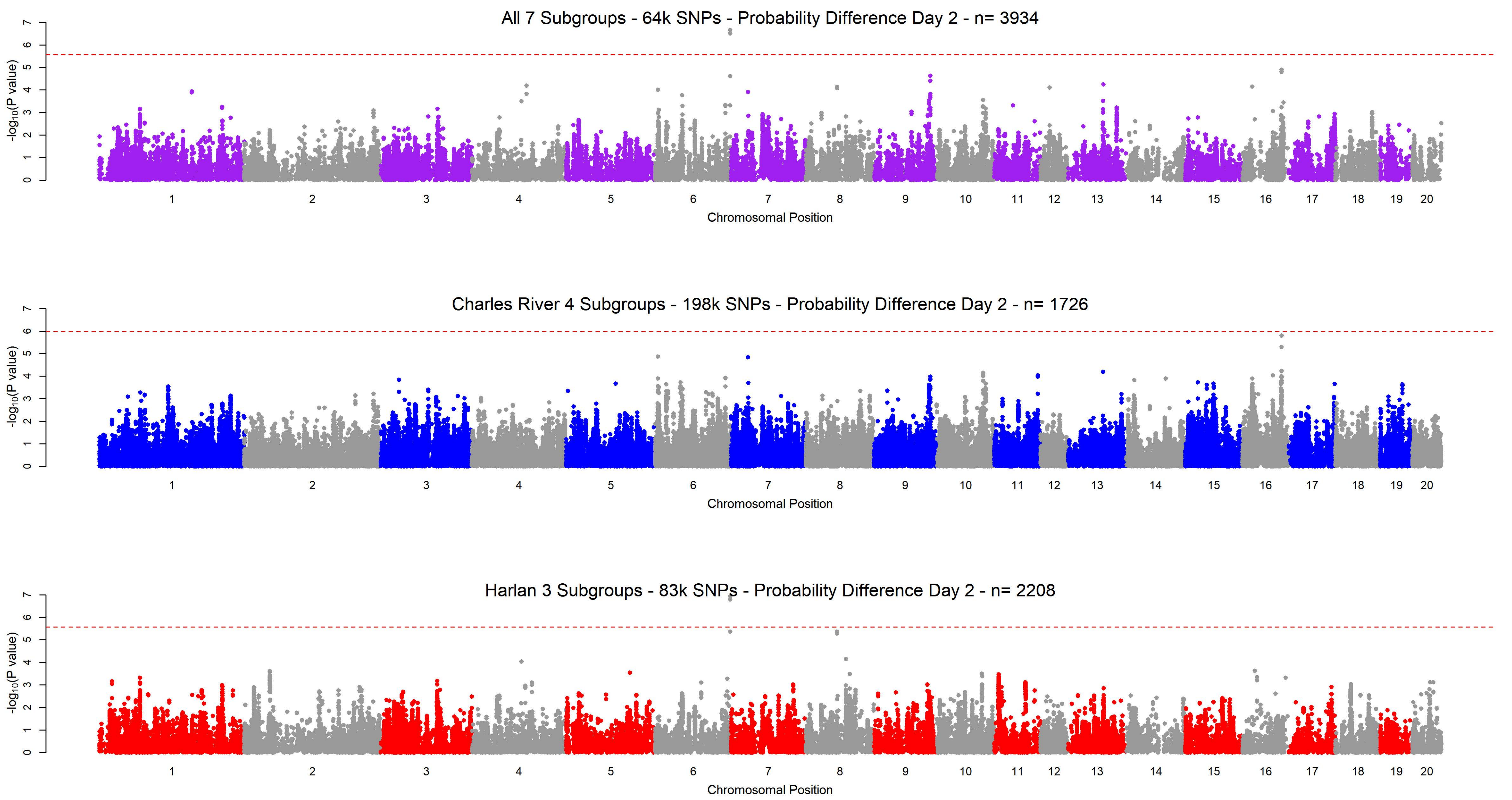

All 7 Subgroups - 64k SNPs - Probability Difference Day 2 - n= 3934
-log10(P value)
Chromosomal Position
Charles River 4 Subgroups - 198k SNPs - Probability Difference Day 2 - n= 1726
-log10(P value)
Chromosomal Position
Harlan 3 Subgroups - 83k SNPs - Probability Difference Day 2 - n= 2208
-log10(P value)
Chromosomal Position

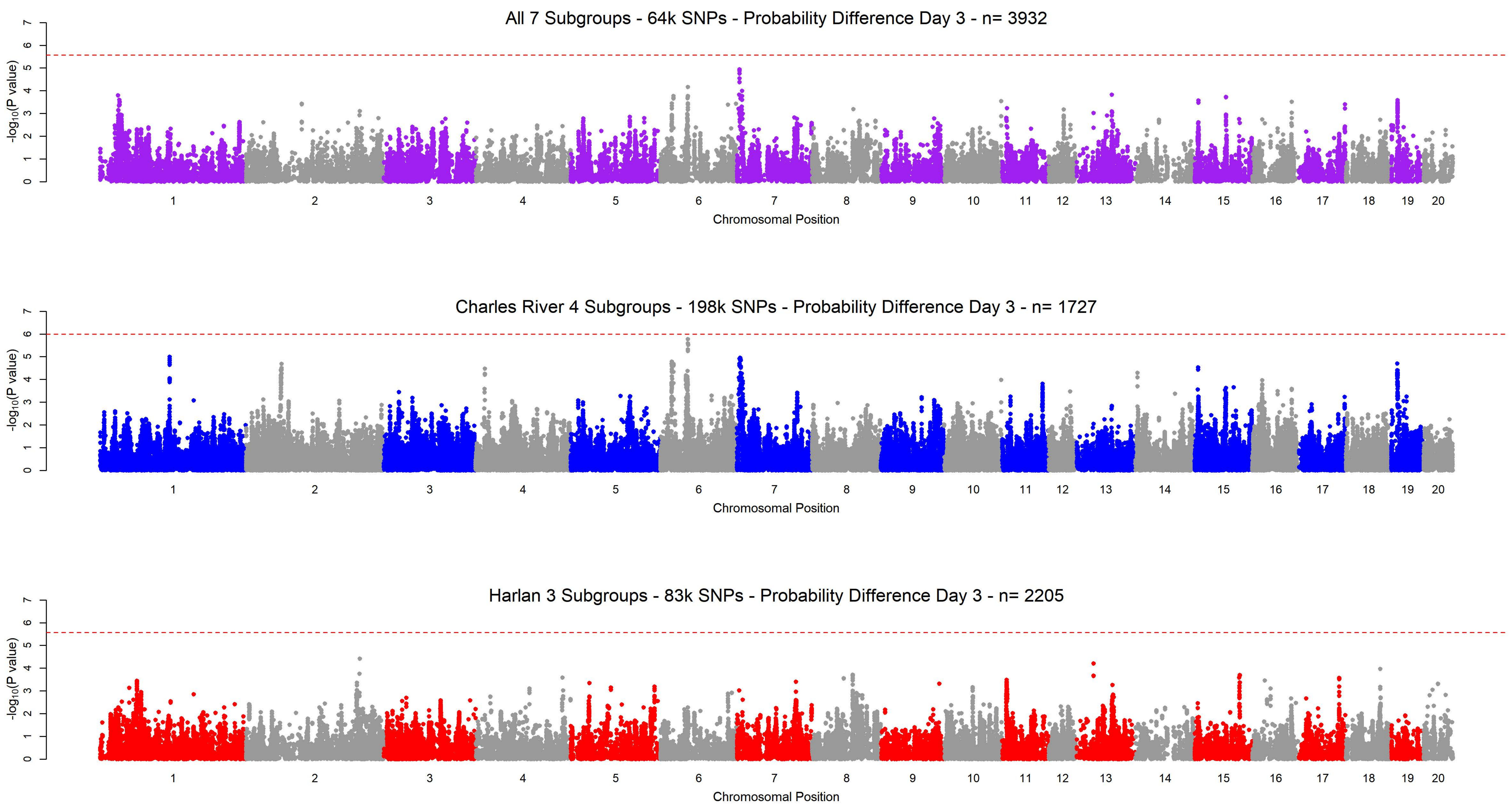

All 7 Subgroups - 64k SNPs - Probability Difference Day 3 - n= 3932
Charles River 4 Subgroups - 198k SNPs - Probability Difference Day 3 - n= 1727
Harlan 3 Subgroups - 83k SNPs - Probability Difference Day 3 - n= 2205
-log10(P value)
Chromosomal Position
1 2 3 4 5 6 7 8 9 10 11 12 13 14 15 16 17 18 19 20
0 1 2 3 4 5 6 7

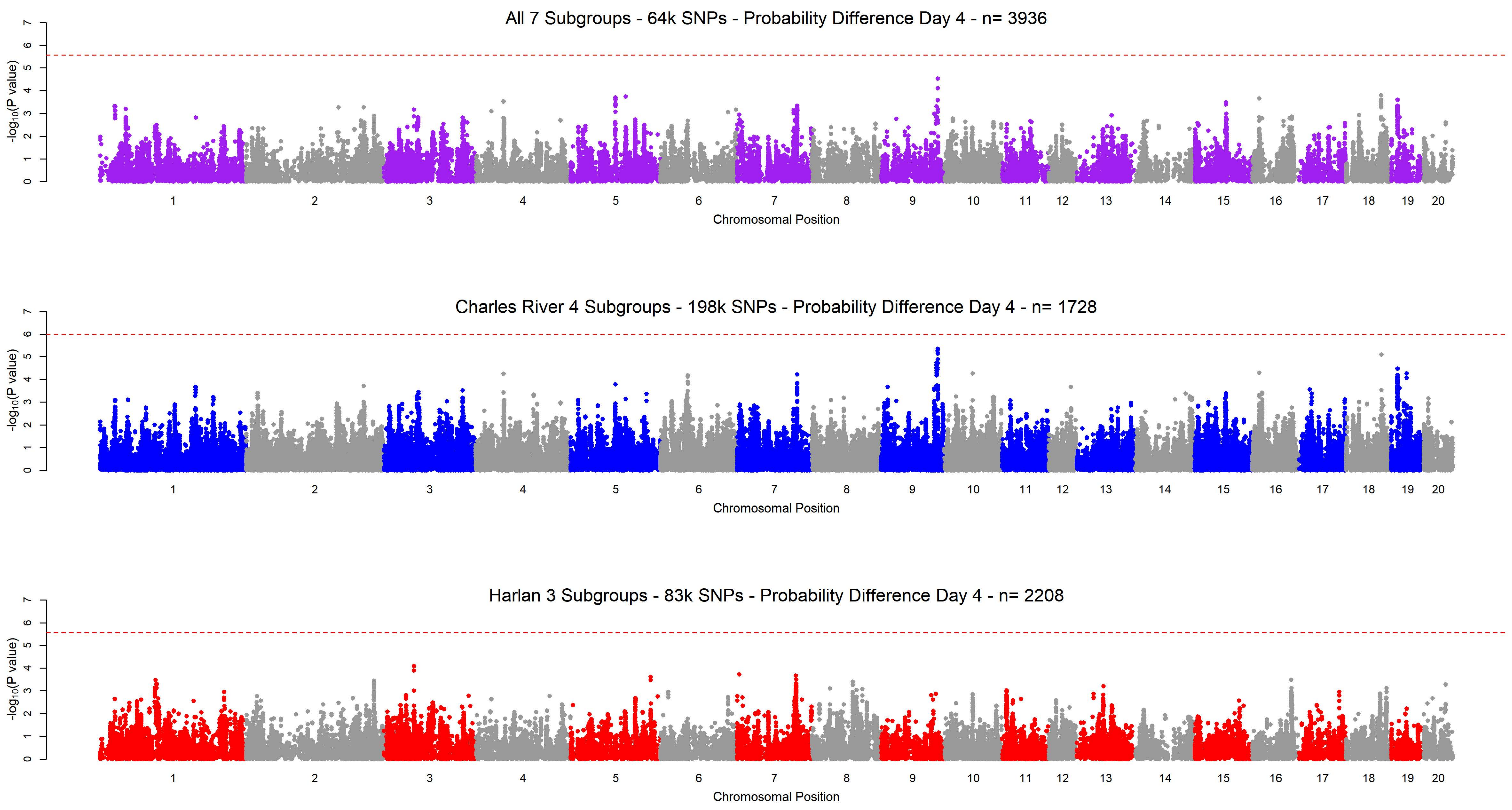

All 7 Subgroups - 64k SNPs - Probability Difference Day 4 - n= 3936
Charles River 4 Subgroups - 198k SNPs - Probability Difference Day 4 - n= 1728
Harlan 3 Subgroups - 83k SNPs - Probability Difference Day 4 - n= 2208
-log10(P value)
Chromosomal Position
0 1 2 3 4 5 6 7
1 2 3 4 5 6 7 8 9 10 11 12 13 14 15 16 17 18 19 20

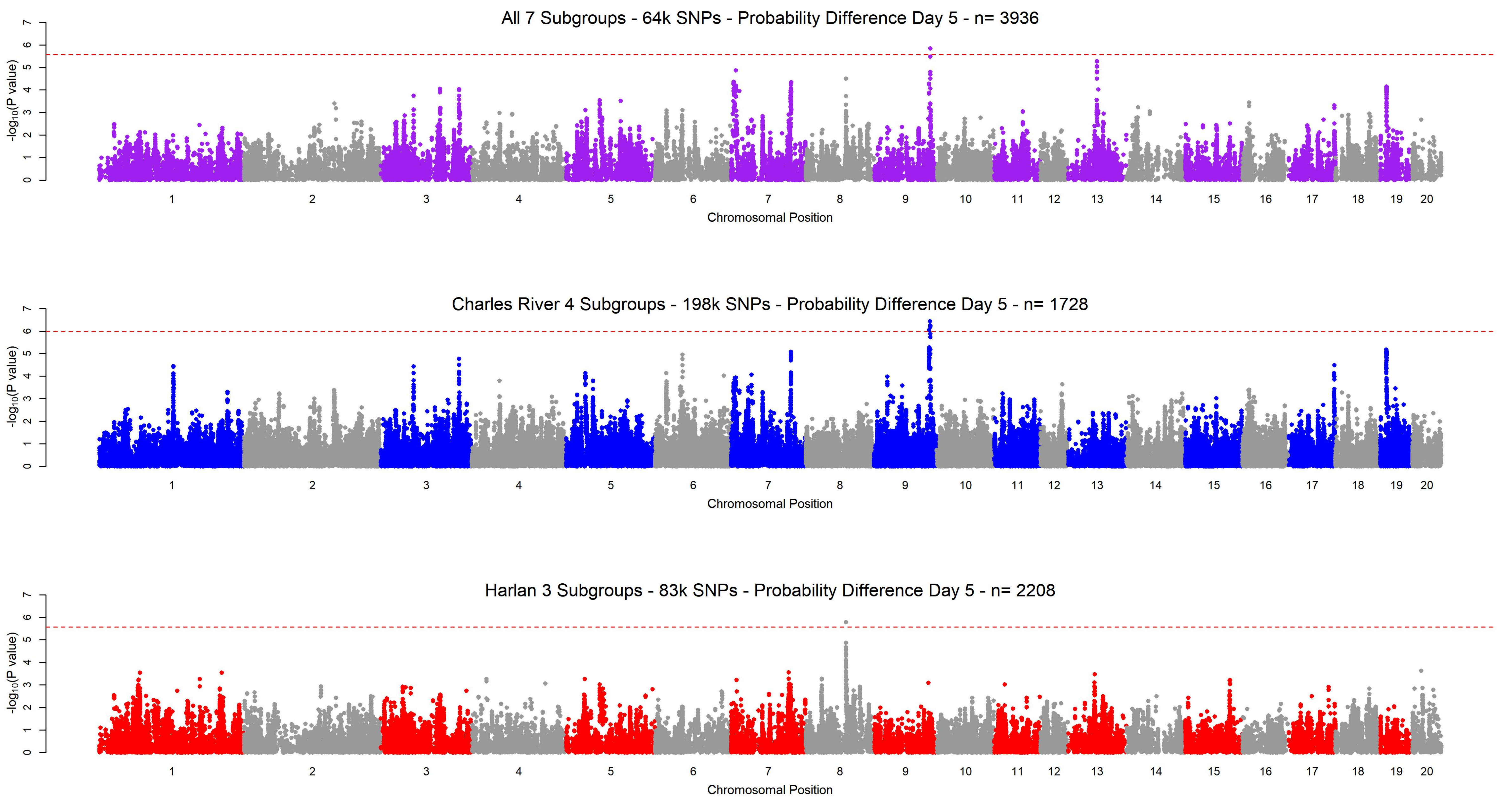
All 7 Subgroups - 64k SNPs - Probability Difference Day 5 - n= 3936
Charles River 4 Subgroups - 198k SNPs - Probability Difference Day 5 - n= 1728
Harlan 3 Subgroups - 83k SNPs - Probability Difference Day 5 - n= 2208
-log10(P value)
Chromosomal Position
1 2 3 4 5 6 7 8 9 10 11 12 13 14 15 16 17 18 19 20

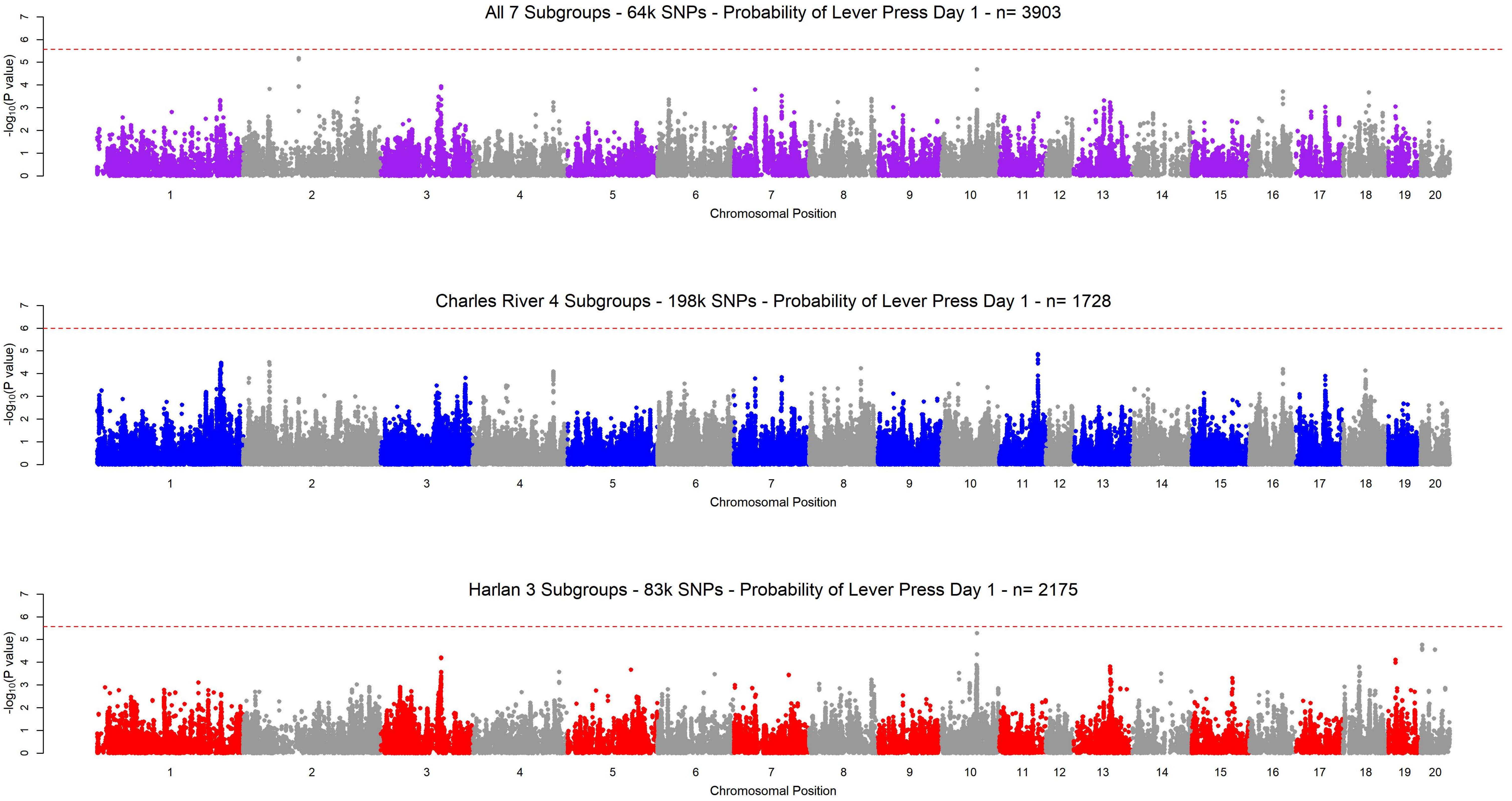
All 7 Subgroups - 64k SNPs - Probability of Lever Press Day 1 - n= 3903
Charles River 4 Subgroups - 198k SNPs - Probability of Lever Press Day 1 - n= 1728
Harlan 3 Subgroups - 83k SNPs - Probability of Lever Press Day 1 - n= 2175
-log10(P value)
0 1 2 3 4 5 6 7
1 2 3 4 5 6 7 8 9 10 11 12 13 14 15 16 17 18 19 20
Chromosomal Position

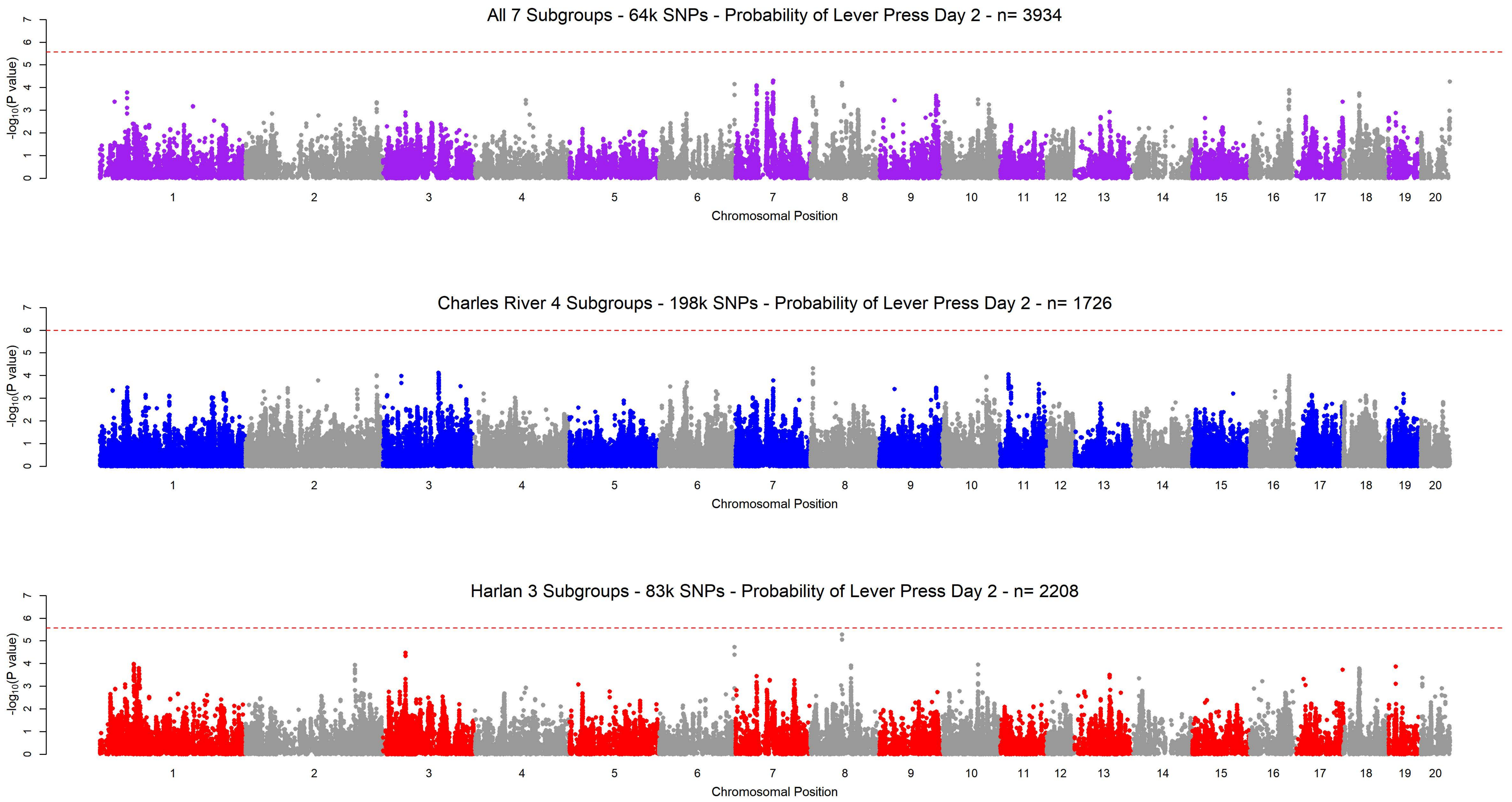
All 7 Subgroups - 64k SNPs - Probability of Lever Press Day 2 - n= 3934
Charles River 4 Subgroups - 198k SNPs - Probability of Lever Press Day 2 - n= 1726
Harlan 3 Subgroups - 83k SNPs - Probability of Lever Press Day 2 - n= 2208
-log10(P value)
Chromosomal Position

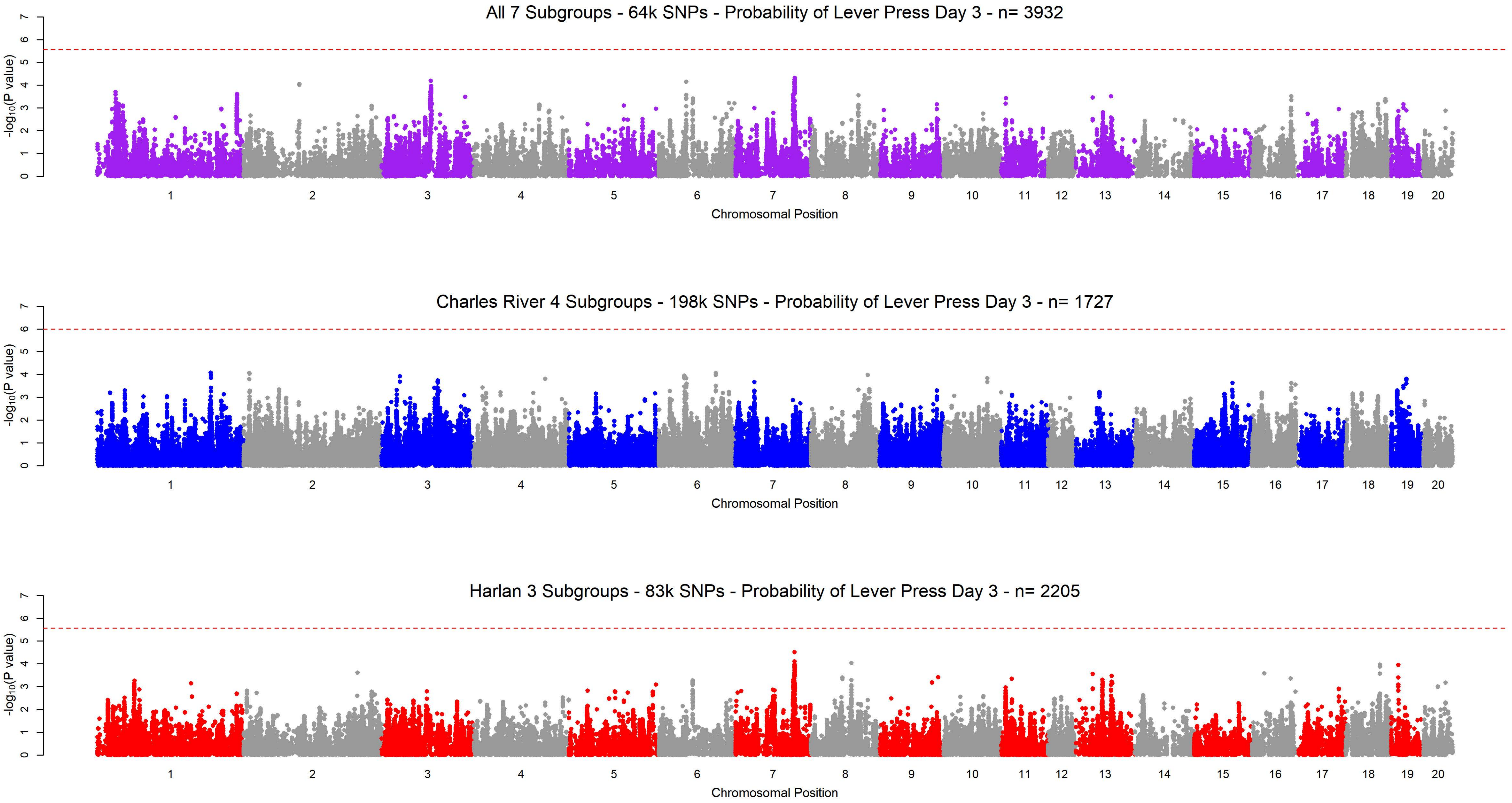

All 7 Subgroups - 64k SNPs - Probability of Lever Press Day 3 - n= 3932
Charles River 4 Subgroups - 198k SNPs - Probability of Lever Press Day 3 - n= 1727
Harlan 3 Subgroups - 83k SNPs - Probability of Lever Press Day 3 - n= 2205
-log10(P value)
Chromosomal Position
0
1
2
3
4
5
6
7

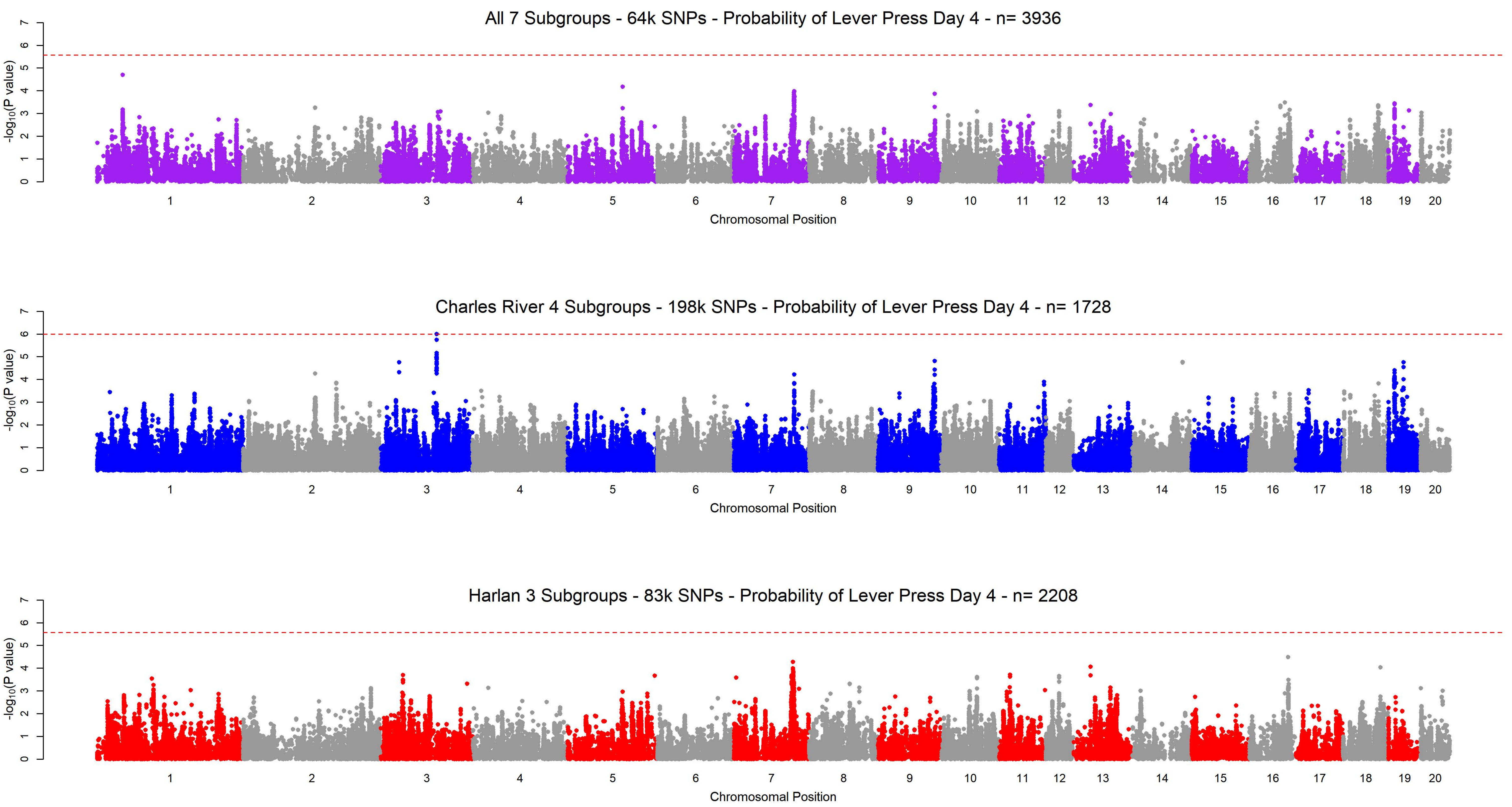
All 7 Subgroups - 64k SNPs - Probability of Lever Press Day 4 - n= 3936
Charles River 4 Subgroups - 198k SNPs - Probability of Lever Press Day 4 - n= 1728
Harlan 3 Subgroups - 83k SNPs - Probability of Lever Press Day 4 - n= 2208
-log10(P value)
0 1 2 3 4 5 6 7
1 2 3 4 5 6 7 8 9 10 11 12 13 14 15 16 17 18 19 20
Chromosomal Position

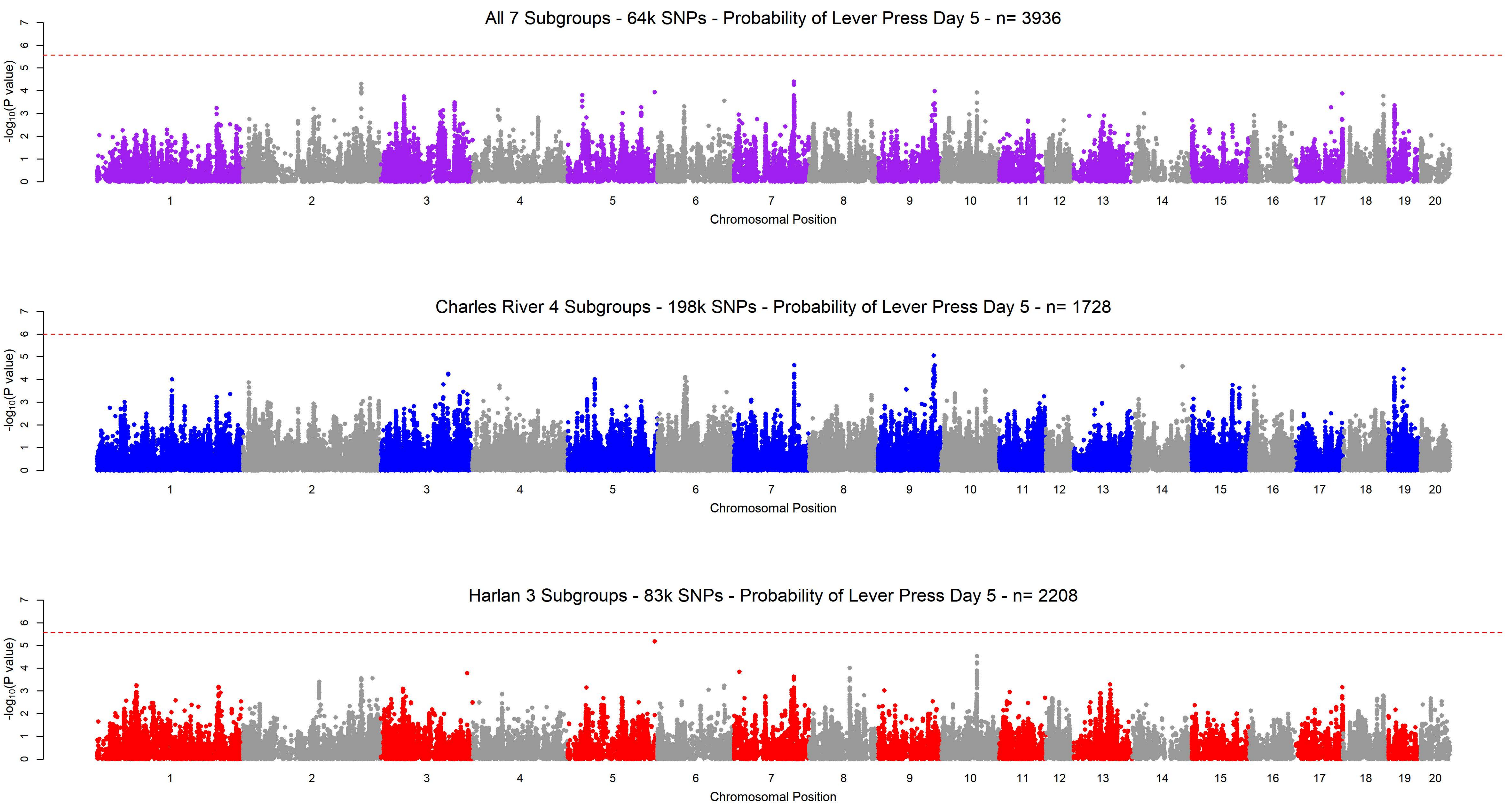
All 7 Subgroups - 64k SNPs - Probability of Lever Press Day 5 - n= 3936
-log10(P value)
Chromosomal Position
Charles River 4 Subgroups - 198k SNPs - Probability of Lever Press Day 5 - n= 1728
-log10(P value)
Chromosomal Position
Harlan 3 Subgroups - 83k SNPs - Probability of Lever Press Day 5 - n= 2208
-log10(P value)
Chromosomal Position

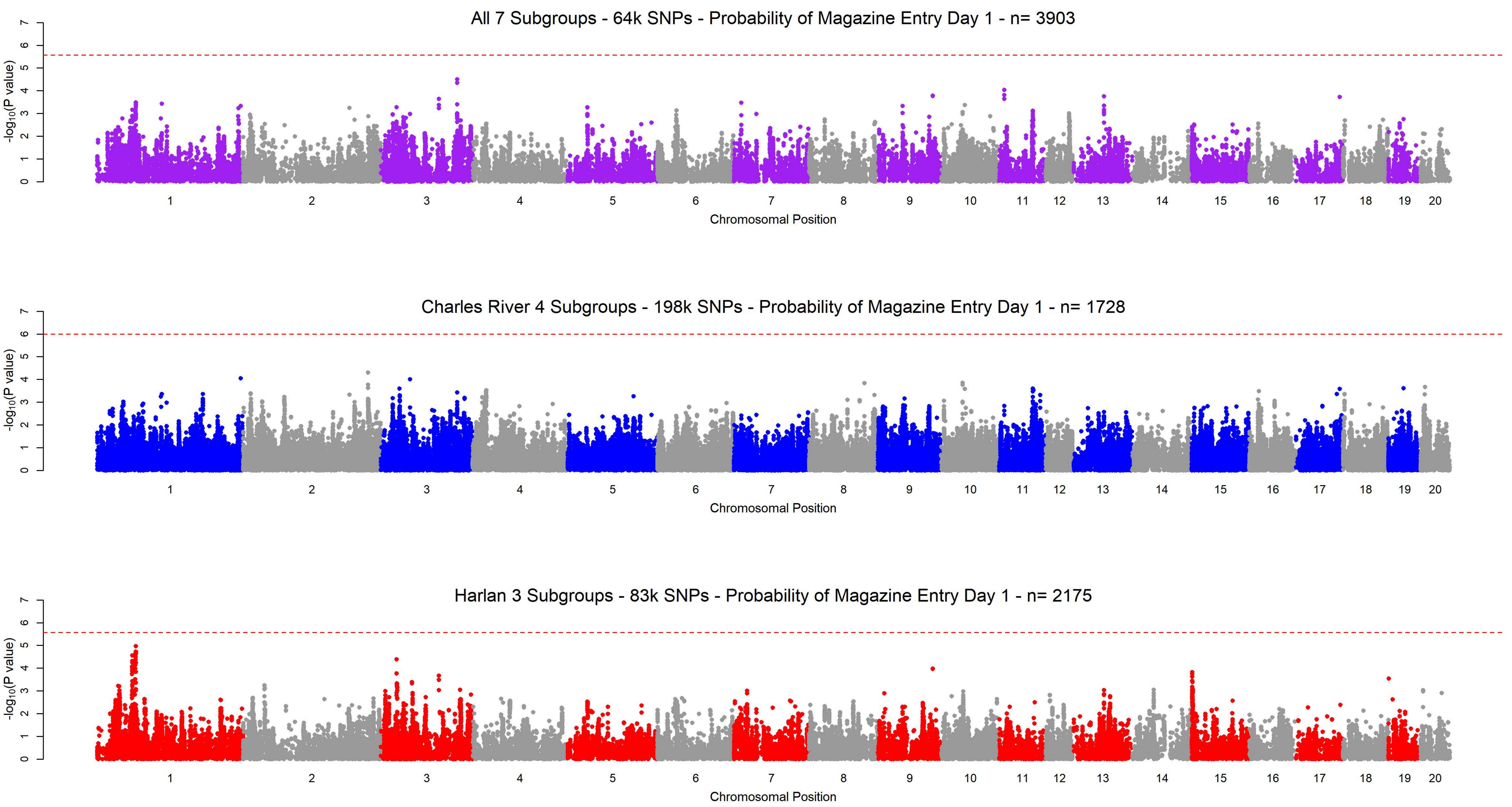
All 7 Subgroups - 64k SNPs - Probability of Magazine Entry Day 1 - n= 3903
-log10(P value)
0 1 2 3 4 5 6 7
1 2 3 4 5 6 7 8 9 10 11 12 13 14 15 16 17 18 19 20
Chromosomal Position
Charles River 4 Subgroups - 198k SNPs - Probability of Magazine Entry Day 1 - n= 1728
-log10(P value)
0 1 2 3 4 5 6 7
1 2 3 4 5 6 7 8 9 10 11 12 13 14 15 16 17 18 19 20
Chromosomal Position
Harlan 3 Subgroups - 83k SNPs - Probability of Magazine Entry Day 1 - n= 2175
-log10(P value)
0 1 2 3 4 5 6 7
1 2 3 4 5 6 7 8 9 10 11 12 13 14 15 16 17 18 19 20
Chromosomal Position

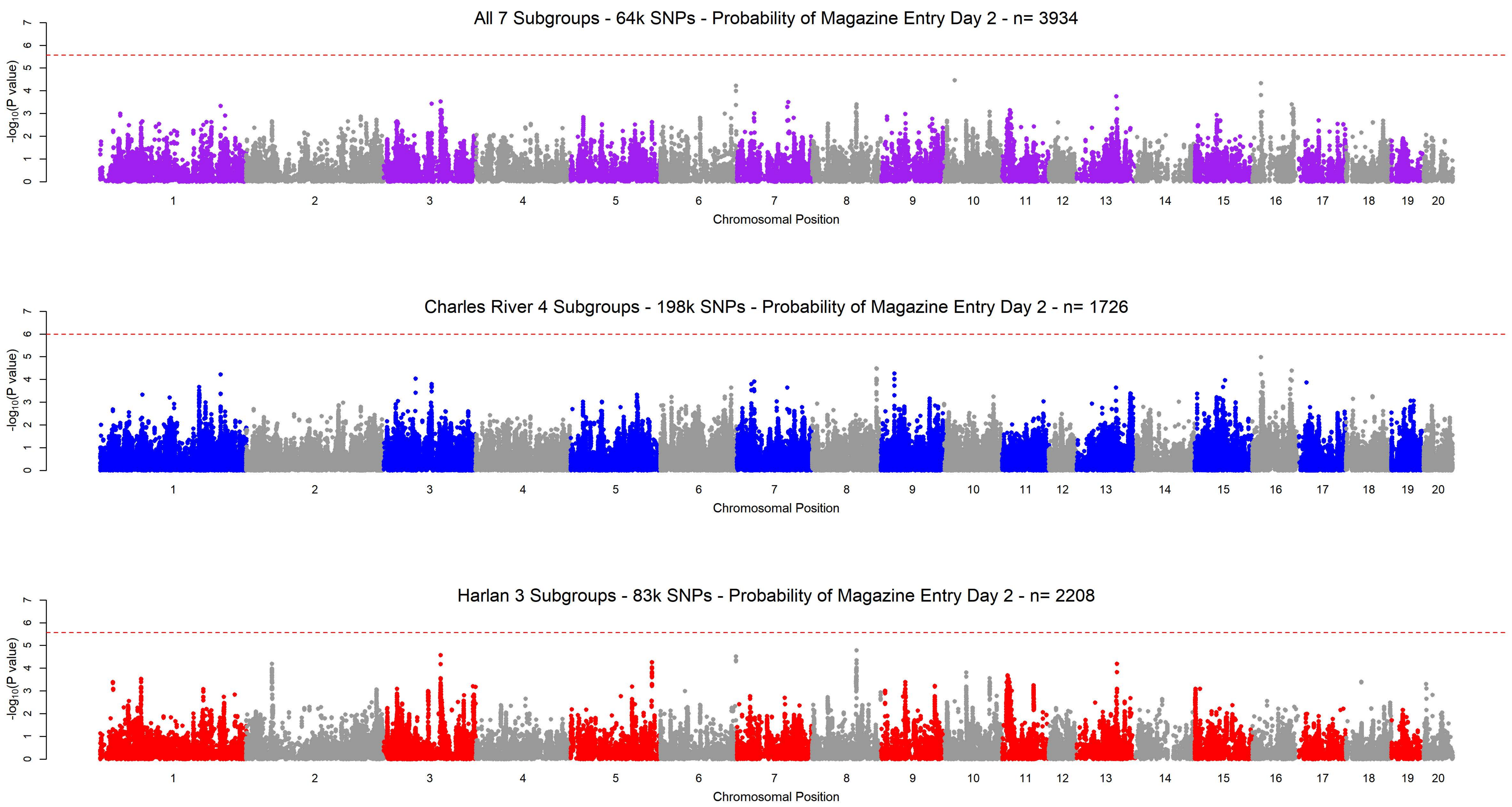

All 7 Subgroups - 64k SNPs - Probability of Magazine Entry Day 2 - n= 3934
Charles River 4 Subgroups - 198k SNPs - Probability of Magazine Entry Day 2 - n= 1726
Harlan 3 Subgroups - 83k SNPs - Probability of Magazine Entry Day 2 - n= 2208
$-\log_{10}$(P value)
Chromosomal Position
1 2 3 4 5 6 7 8 9 10 11 12 13 14 15 16 17 18 19 20
0 1 2 3 4 5 6 7

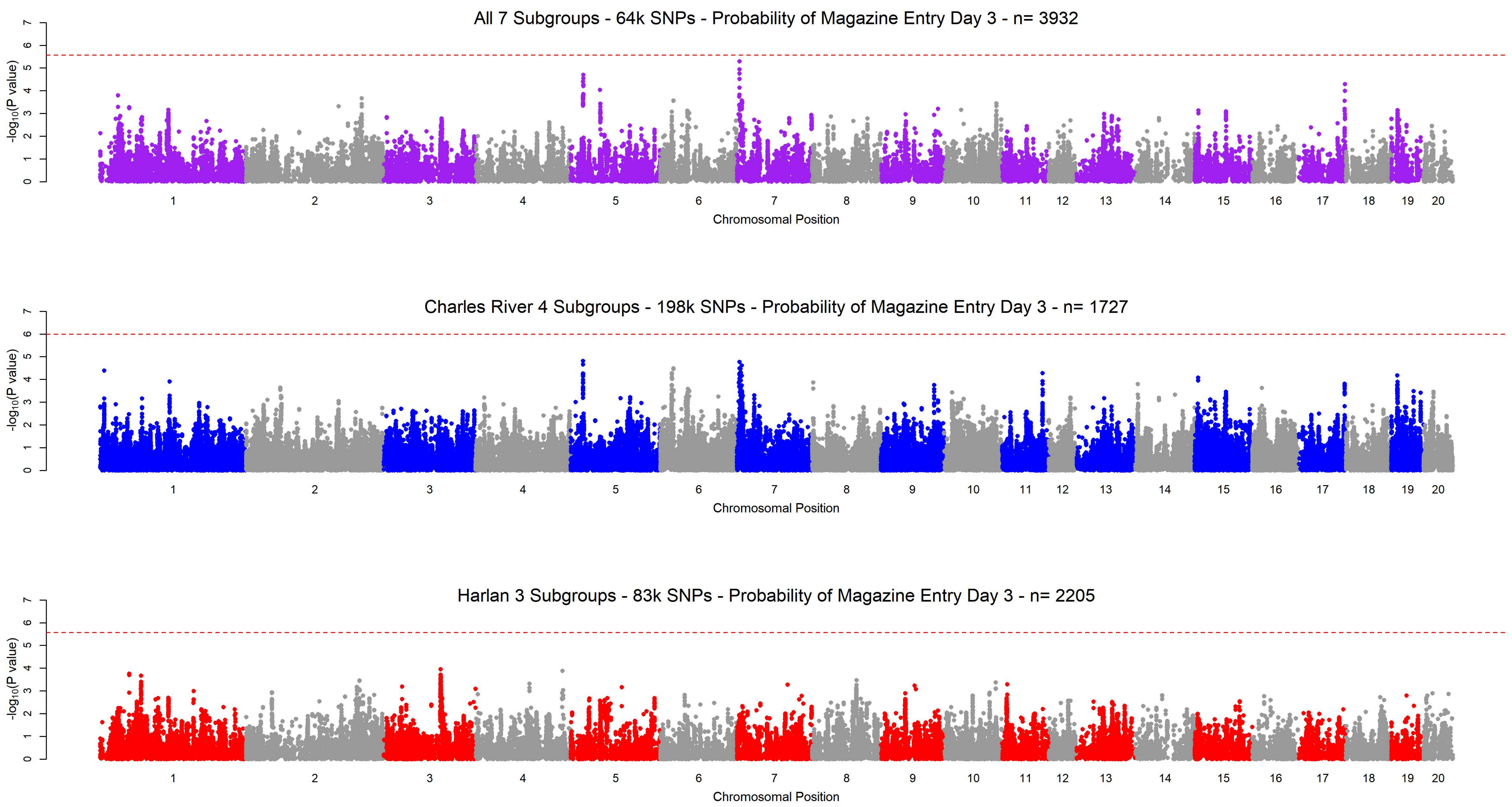

All 7 Subgroups - 64k SNPs - Probability of Magazine Entry Day 3 - n= 3932
-log10(P value)
Chromosomal Position
1 2 3 4 5 6 7 8 9 10 11 12 13 14 15 16 17 18 19 20
Charles River 4 Subgroups - 198k SNPs - Probability of Magazine Entry Day 3 - n= 1727
-log10(P value)
Chromosomal Position
1 2 3 4 5 6 7 8 9 10 11 12 13 14 15 16 17 18 19 20
Harlan 3 Subgroups - 83k SNPs - Probability of Magazine Entry Day 3 - n= 2205
-log10(P value)
Chromosomal Position
1 2 3 4 5 6 7 8 9 10 11 12 13 14 15 16 17 18 19 20

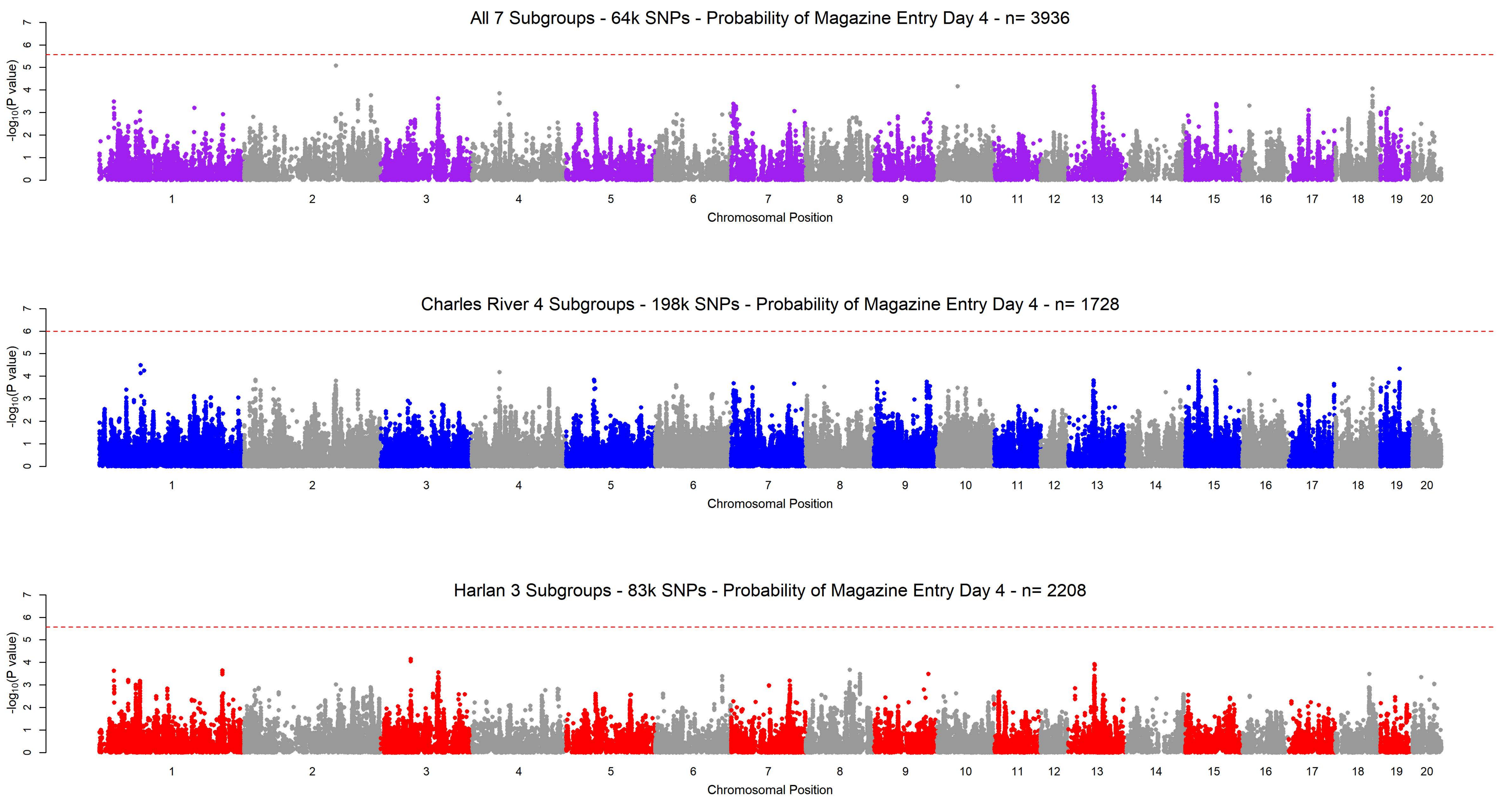
All 7 Subgroups - 64k SNPs - Probability of Magazine Entry Day 4 - n= 3936
-log10(P value)
Chromosomal Position
Charles River 4 Subgroups - 198k SNPs - Probability of Magazine Entry Day 4 - n= 1728
-log10(P value)
Chromosomal Position
Harlan 3 Subgroups - 83k SNPs - Probability of Magazine Entry Day 4 - n= 2208
-log10(P value)
Chromosomal Position

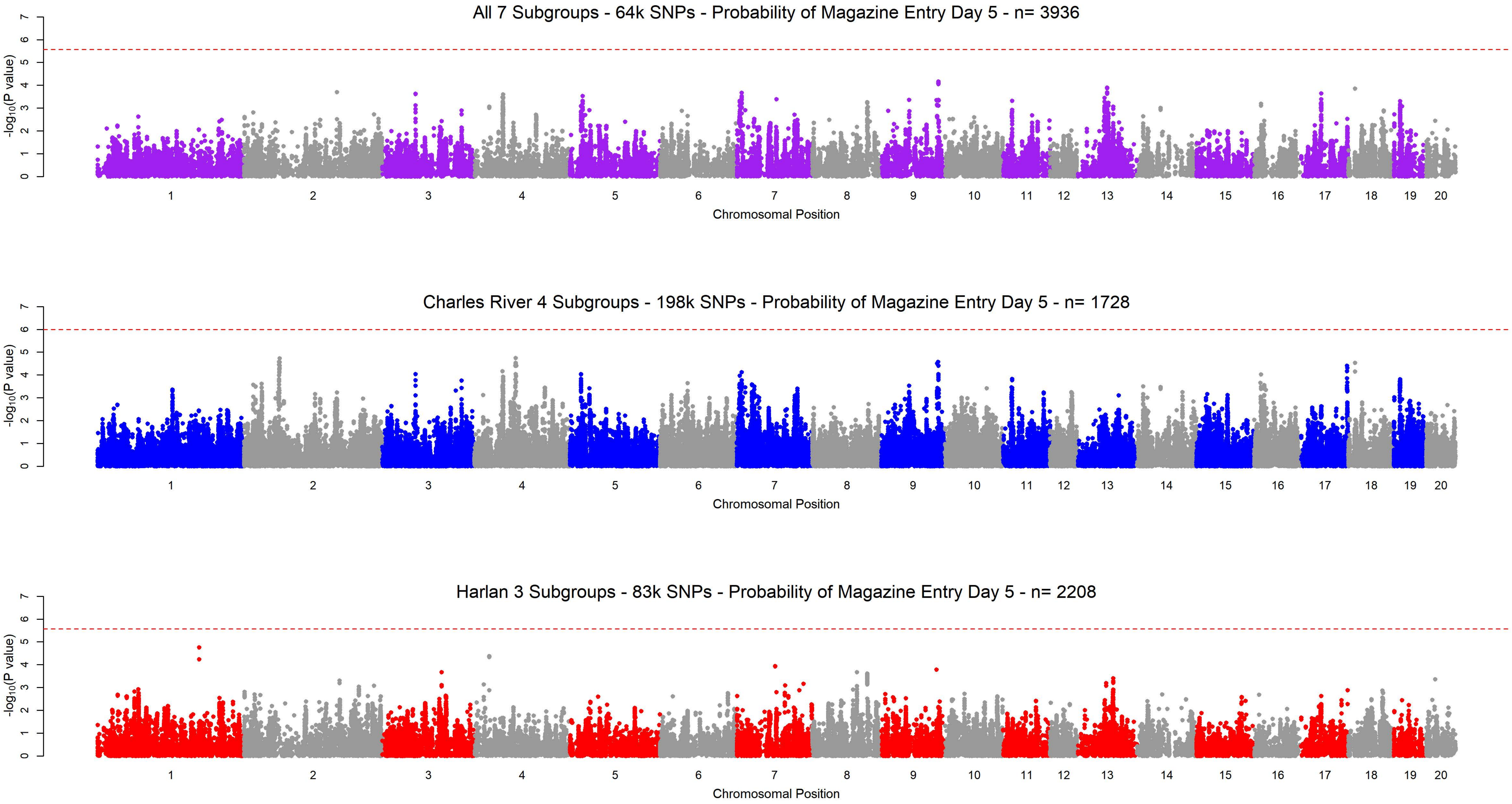
All 7 Subgroups - 64k SNPs - Probability of Magazine Entry Day 5 - n= 3936
Charles River 4 Subgroups - 198k SNPs - Probability of Magazine Entry Day 5 - n= 1728
Harlan 3 Subgroups - 83k SNPs - Probability of Magazine Entry Day 5 - n= 2208
-log10(P value)
0 1 2 3 4 5 6 7
1 2 3 4 5 6 7 8 9 10 11 12 13 14 15 16 17 18 19 20
Chromosomal Position

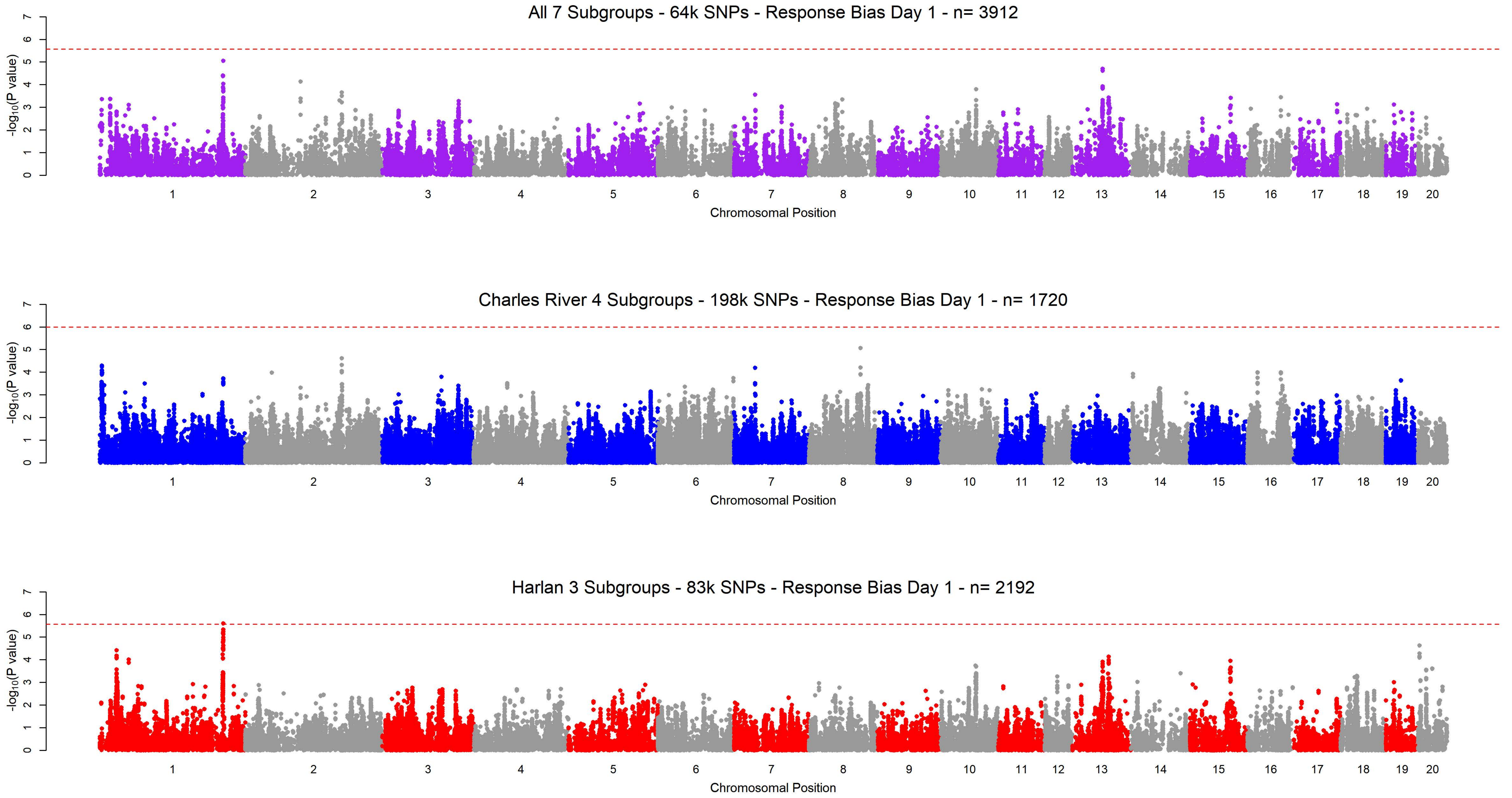
All 7 Subgroups - 64k SNPs - Response Bias Day 1 - n= 3912
Charles River 4 Subgroups - 198k SNPs - Response Bias Day 1 - n= 1720
Harlan 3 Subgroups - 83k SNPs - Response Bias Day 1 - n= 2192
-log10(P value)
Chromosomal Position
0 1 2 3 4 5 6 7
1 2 3 4 5 6 7 8 9 10 11 12 13 14 15 16 17 18 19 20

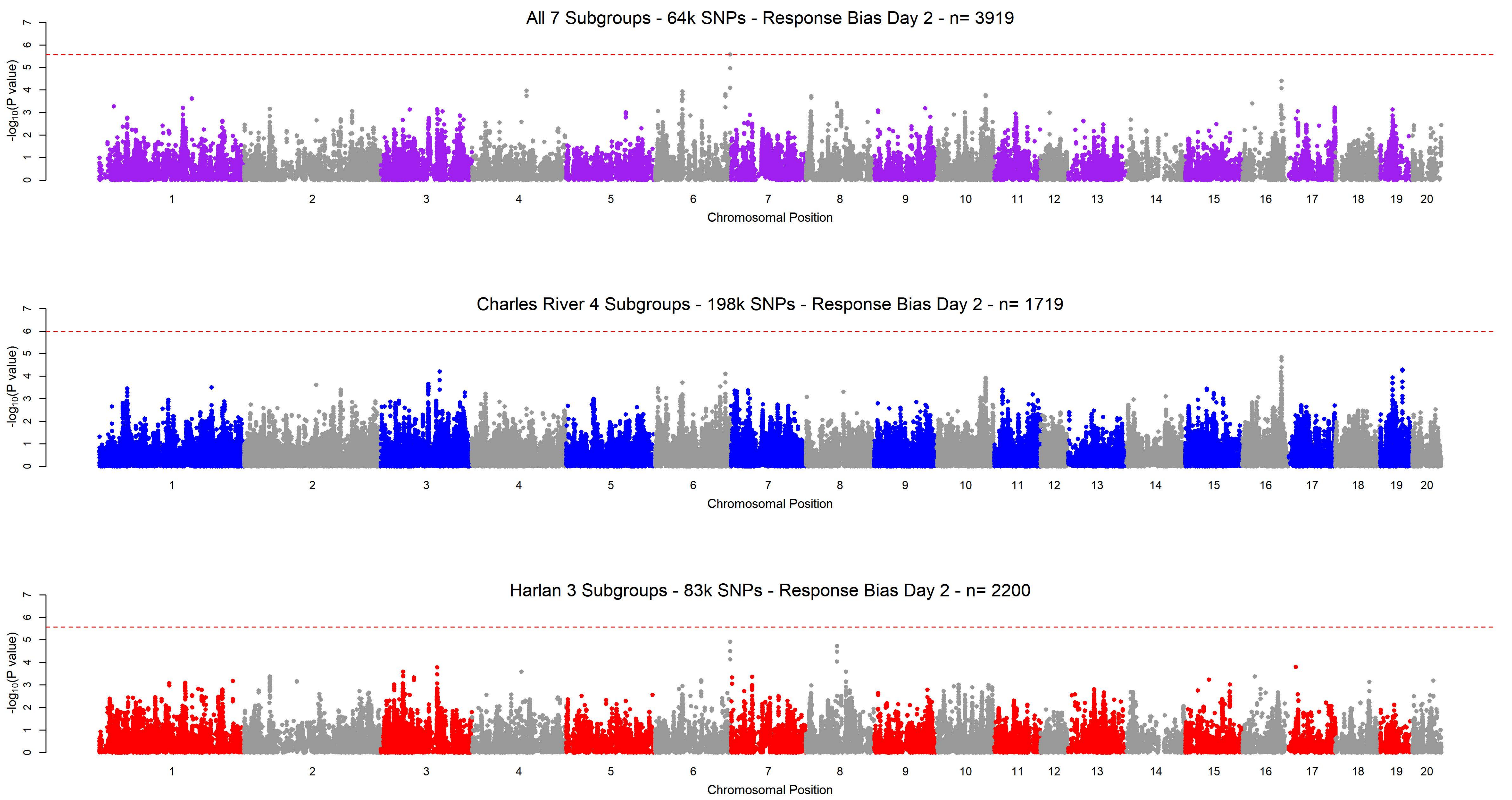
All 7 Subgroups - 64k SNPs - Response Bias Day 2 - n= 3919
Charles River 4 Subgroups - 198k SNPs - Response Bias Day 2 - n= 1719
Harlan 3 Subgroups - 83k SNPs - Response Bias Day 2 - n= 2200
$-\log_{10}$(P value)
Chromosomal Position
1 2 3 4 5 6 7 8 9 10 11 12 13 14 15 16 17 18 19 20
0 1 2 3 4 5 6 7

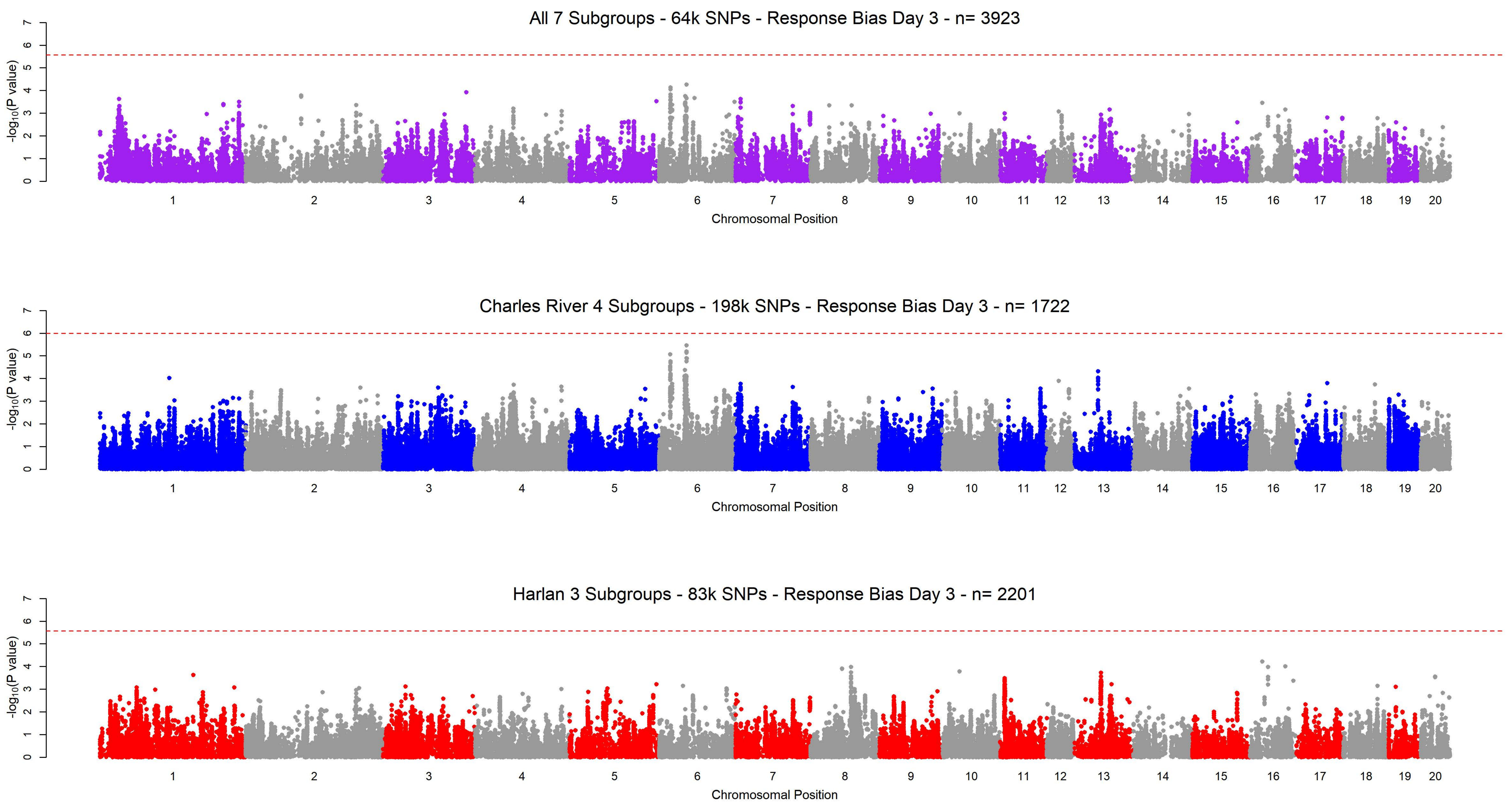
All 7 Subgroups - 64k SNPs - Response Bias Day 3 - n= 3923
$-\log_{10}$(P value)
Chromosomal Position
Charles River 4 Subgroups - 198k SNPs - Response Bias Day 3 - n= 1722
$-\log_{10}$(P value)
Chromosomal Position
Harlan 3 Subgroups - 83k SNPs - Response Bias Day 3 - n= 2201
$-\log_{10}$(P value)
Chromosomal Position

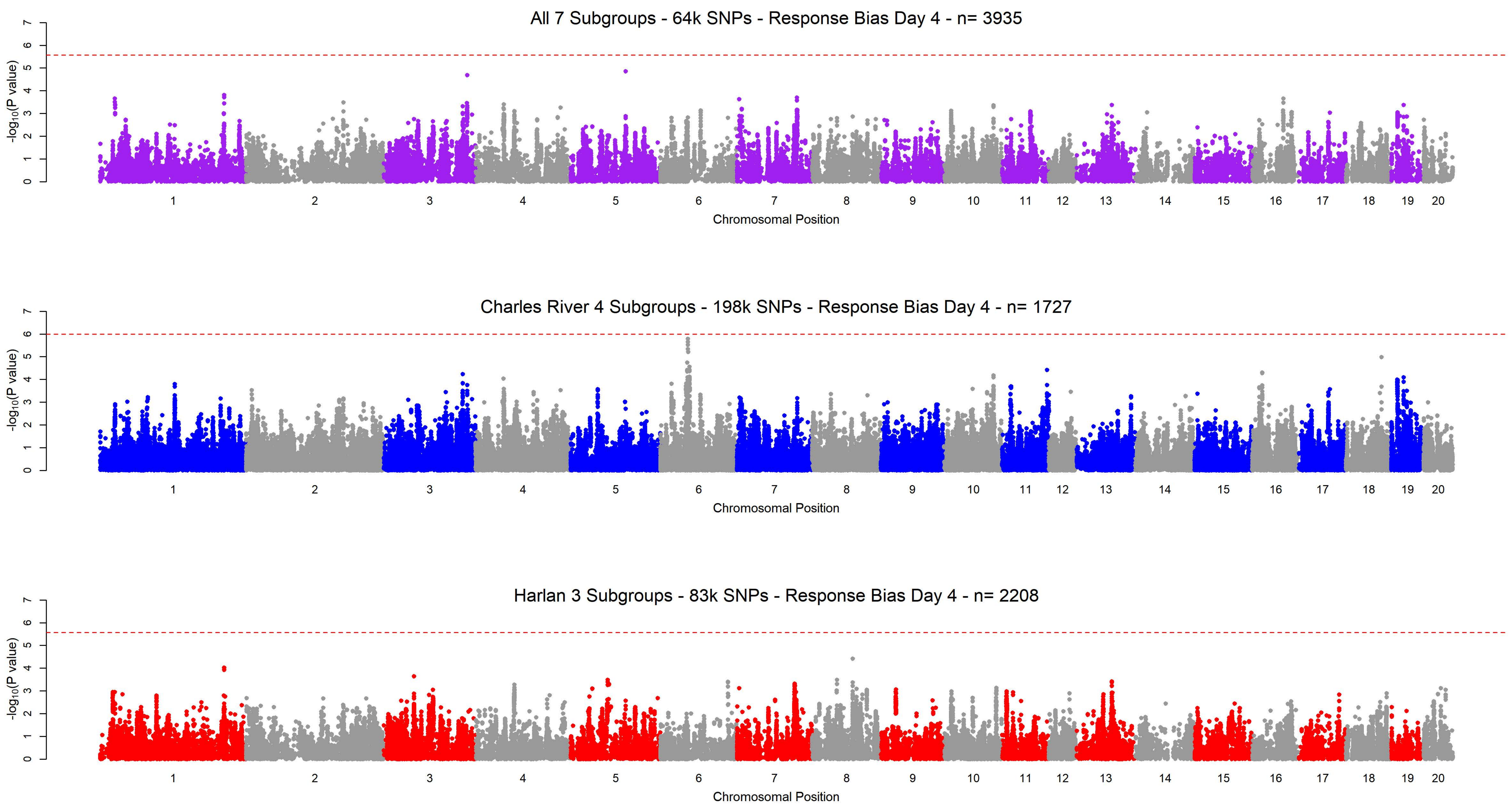
All 7 Subgroups - 64k SNPs - Response Bias Day 4 - n= 3935
-log10(P value)
Chromosomal Position
Charles River 4 Subgroups - 198k SNPs - Response Bias Day 4 - n= 1727
-log10(P value)
Chromosomal Position
Harlan 3 Subgroups - 83k SNPs - Response Bias Day 4 - n= 2208
-log10(P value)
Chromosomal Position

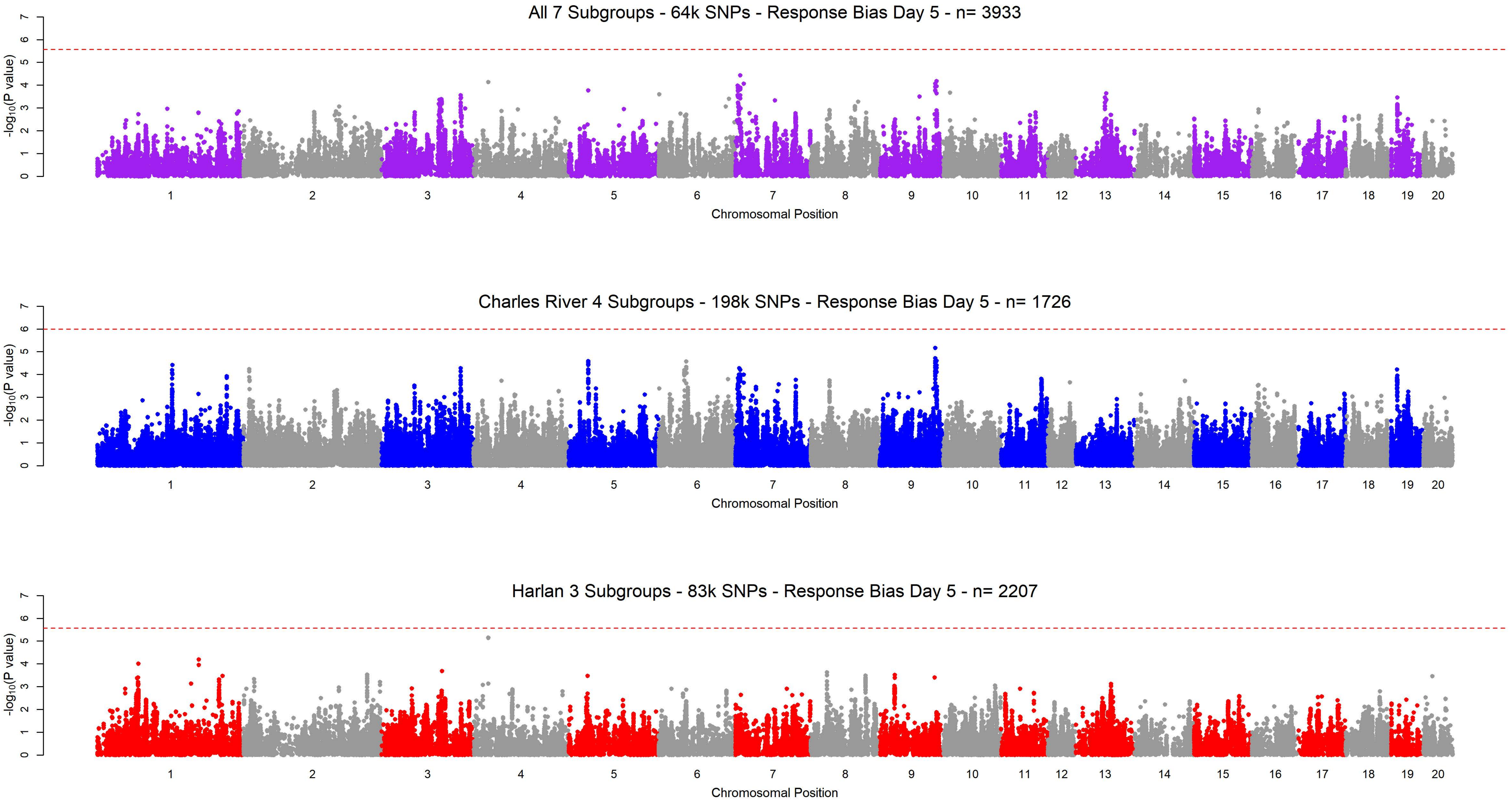

All 7 Subgroups - 64k SNPs - Response Bias Day 5 - n= 3933
Charles River 4 Subgroups - 198k SNPs - Response Bias Day 5 - n= 1726
Harlan 3 Subgroups - 83k SNPs - Response Bias Day 5 - n= 2207
-log10(P value)
Chromosomal Position

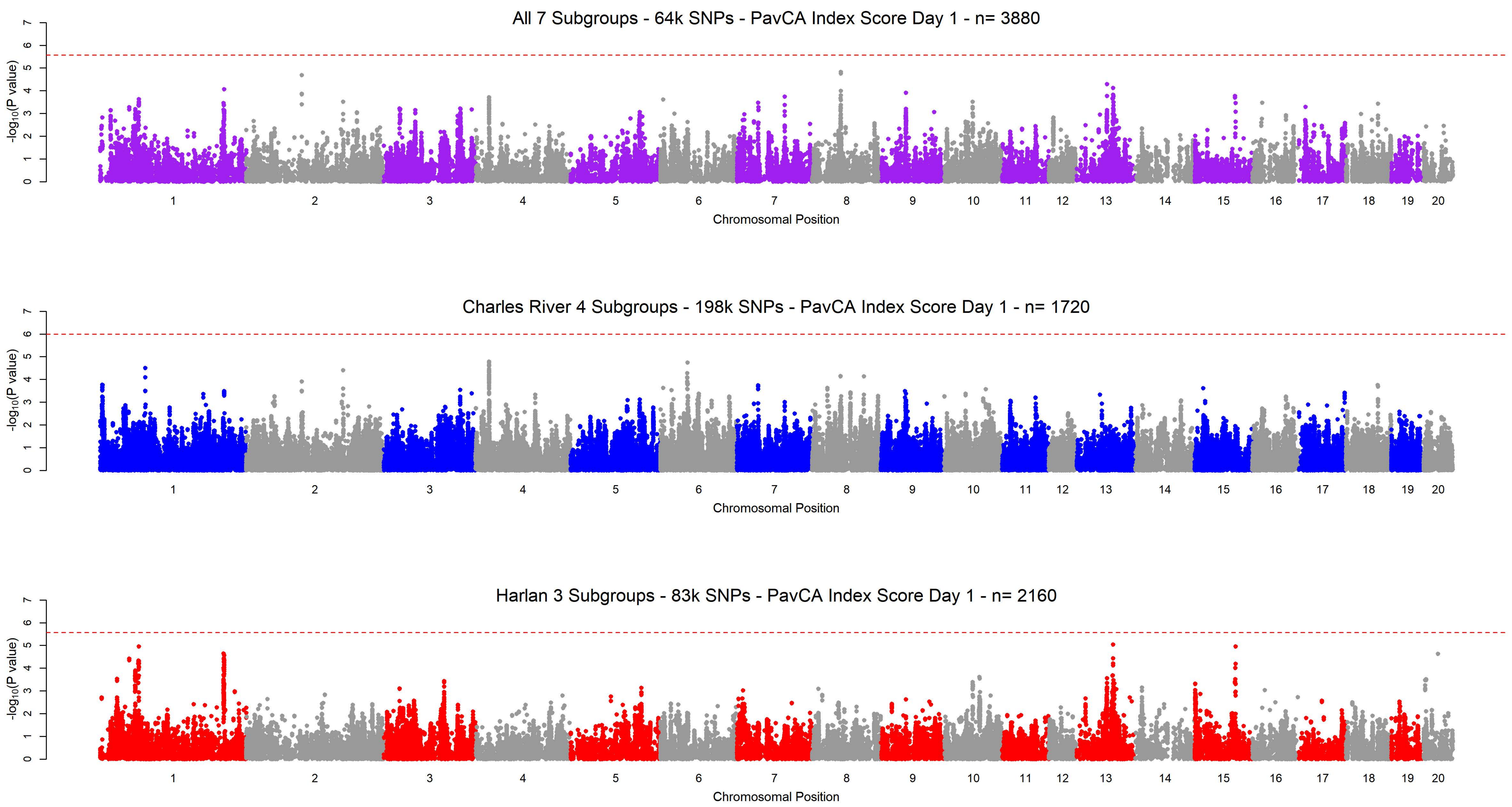
All 7 Subgroups - 64k SNPs - PavCA Index Score Day 1 - n= 3880
Charles River 4 Subgroups - 198k SNPs - PavCA Index Score Day 1 - n= 1720
Harlan 3 Subgroups - 83k SNPs - PavCA Index Score Day 1 - n= 2160
-log10(P value)
0 1 2 3 4 5 6 7
1 2 3 4 5 6 7 8 9 10 11 12 13 14 15 16 17 18 19 20
Chromosomal Position

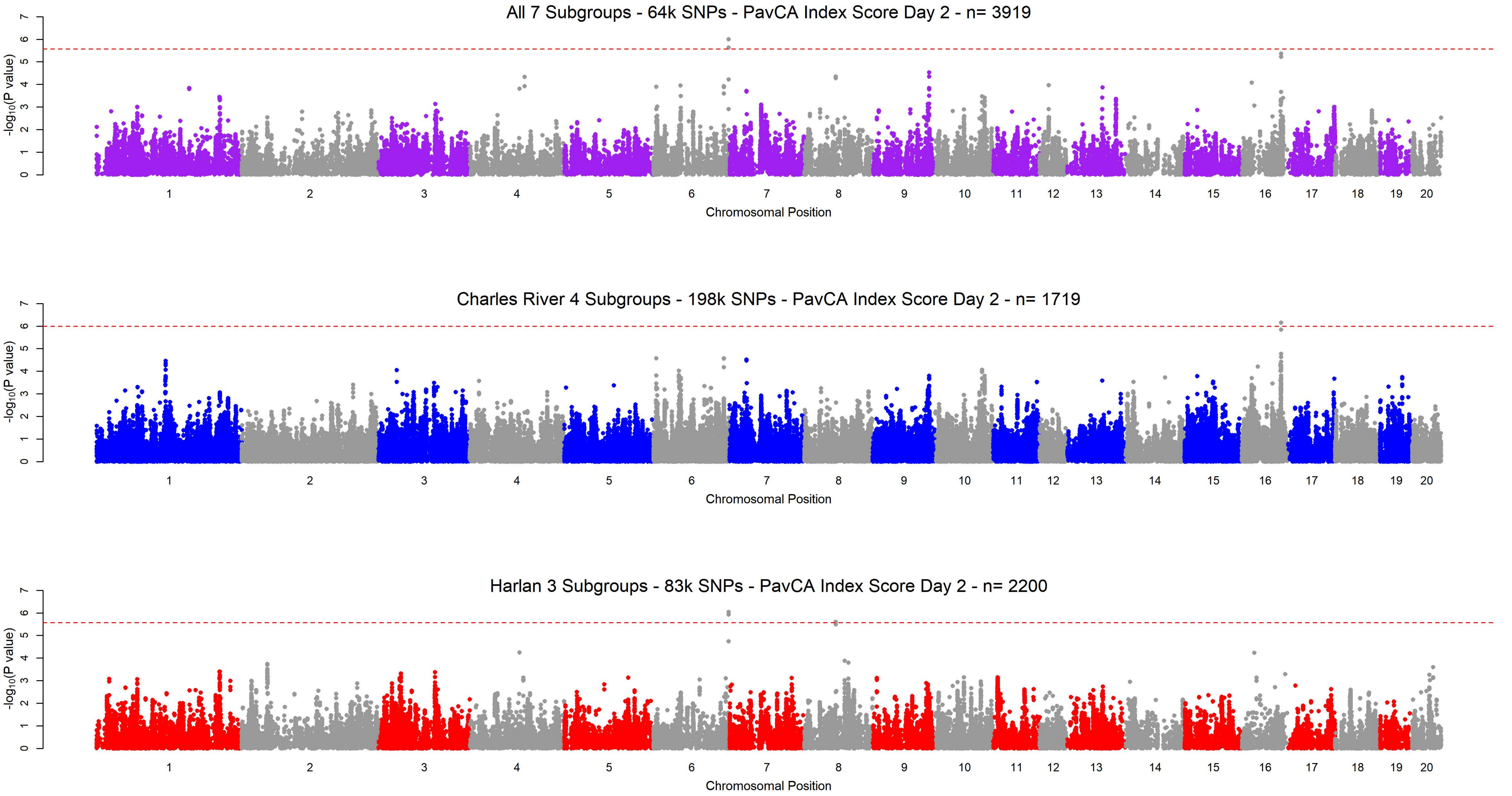

All 7 Subgroups - 64k SNPs - PavCA Index Score Day 2 - n= 3919
Charles River 4 Subgroups - 198k SNPs - PavCA Index Score Day 2 - n= 1719
Harlan 3 Subgroups - 83k SNPs - PavCA Index Score Day 2 - n= 2200
-log10(P value)
Chromosomal Position
1 2 3 4 5 6 7 8 9 10 11 12 13 14 15 16 17 18 19 20
0 1 2 3 4 5 6 7

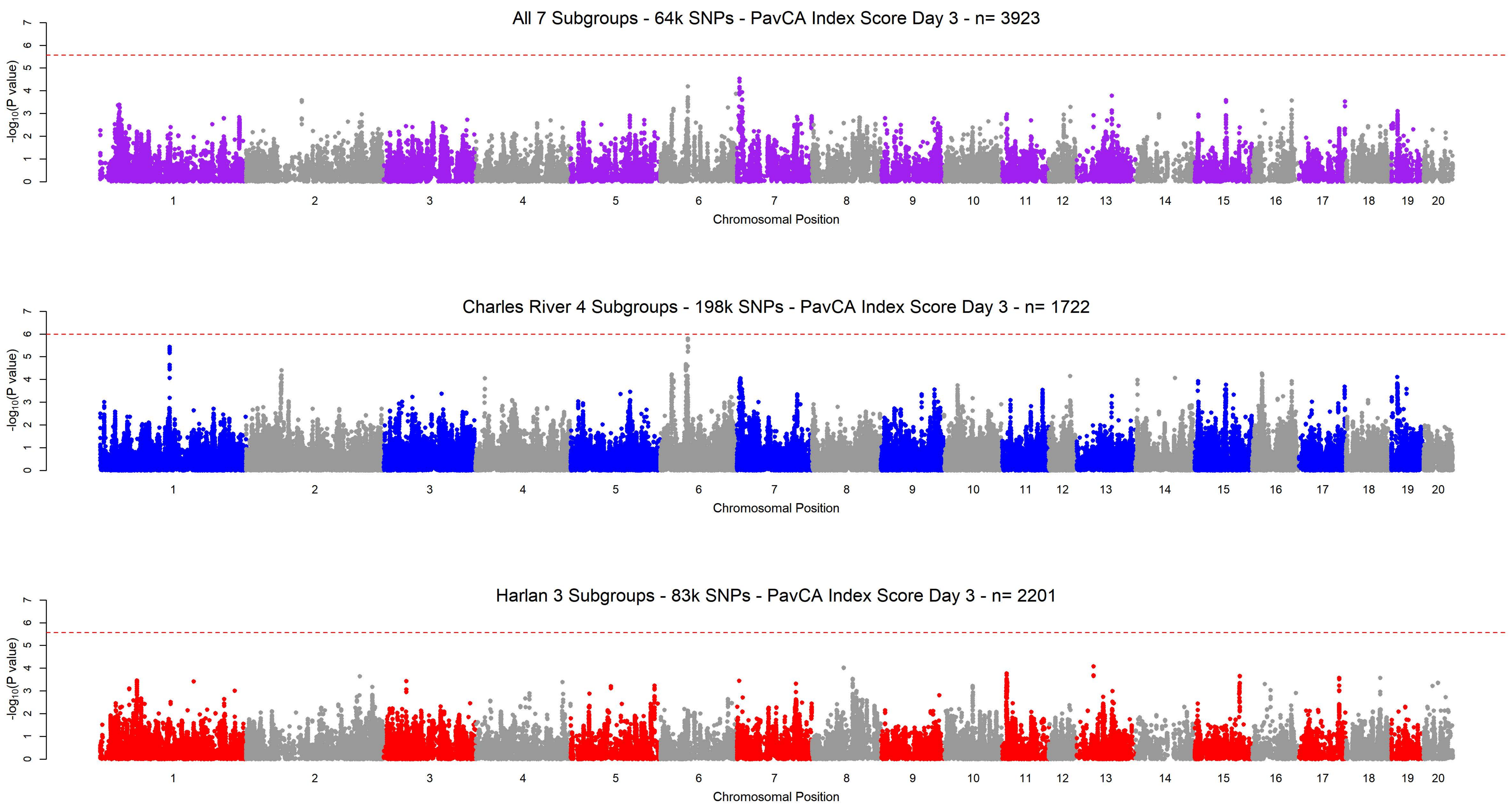
All 7 Subgroups - 64k SNPs - PavCA Index Score Day 3 - n= 3923
Charles River 4 Subgroups - 198k SNPs - PavCA Index Score Day 3 - n= 1722
Harlan 3 Subgroups - 83k SNPs - PavCA Index Score Day 3 - n= 2201
-log10(P value)
Chromosomal Position
1 2 3 4 5 6 7 8 9 10 11 12 13 14 15 16 17 18 19 20
0 1 2 3 4 5 6 7

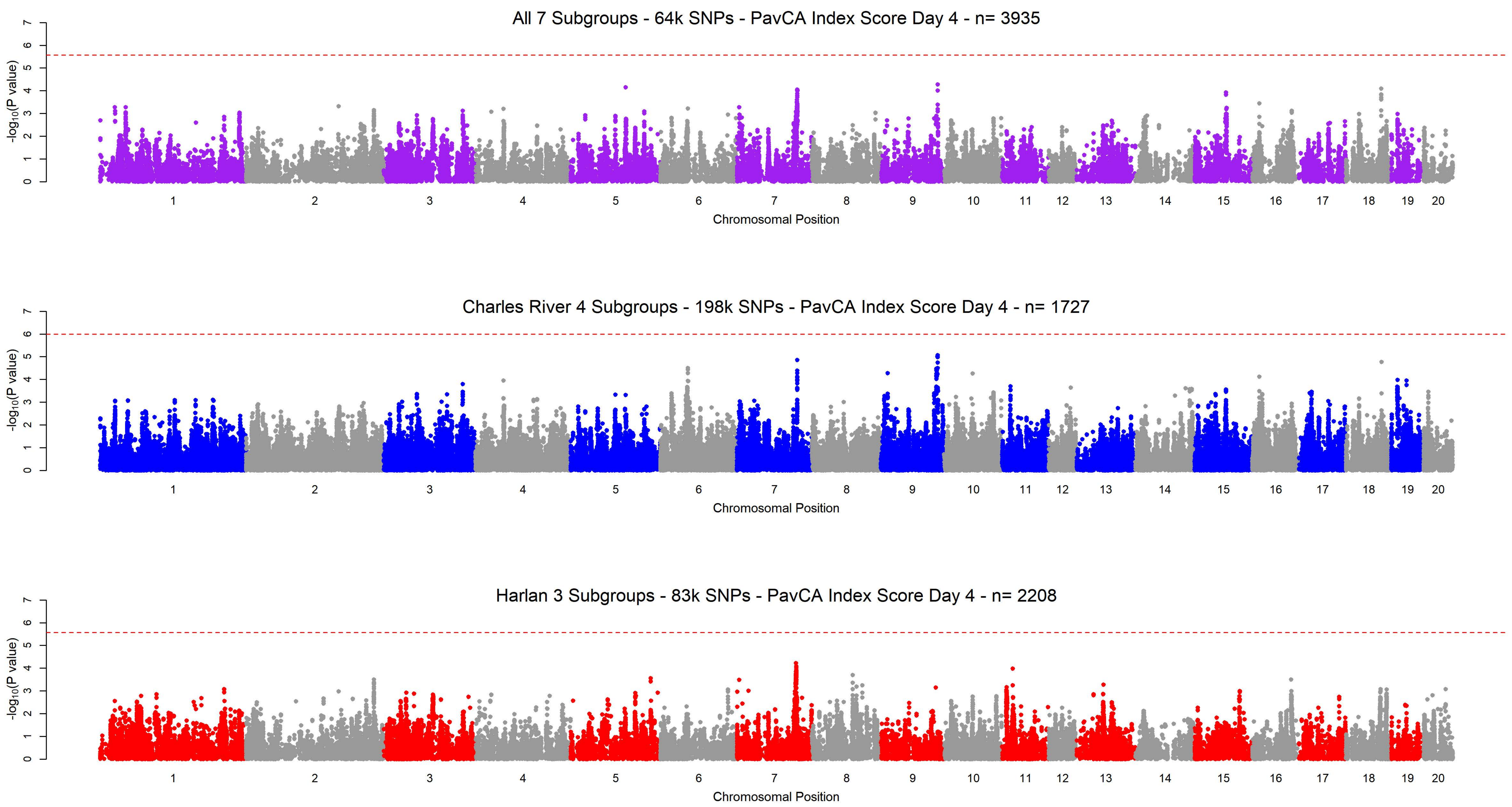

All 7 Subgroups - 64k SNPs - PavCA Index Score Day 4 - n= 3935
Charles River 4 Subgroups - 198k SNPs - PavCA Index Score Day 4 - n= 1727
Harlan 3 Subgroups - 83k SNPs - PavCA Index Score Day 4 - n= 2208
-log10(P value)
Chromosomal Position
1 2 3 4 5 6 7 8 9 10 11 12 13 14 15 16 17 18 19 20
0 1 2 3 4 5 6 7

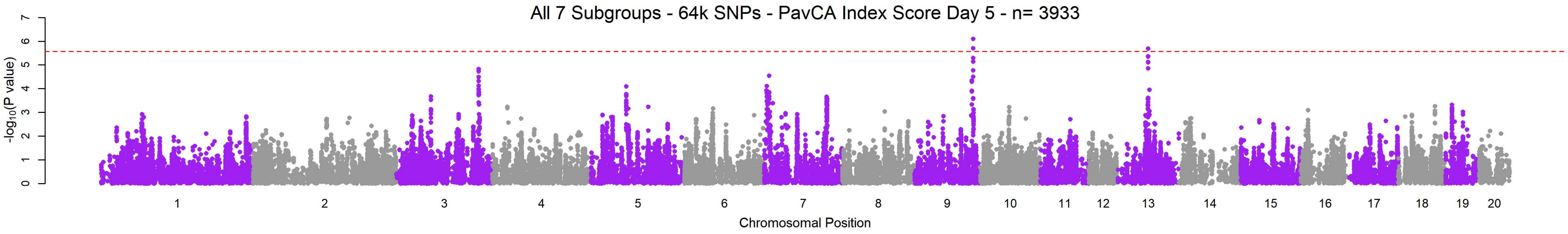
All 7 Subgroups - 64k SNPs - PavCA Index Score Day 5 - n= 3933
-log10(P value)
0
1
2
3
4
5
6
7
Chromosomal Position

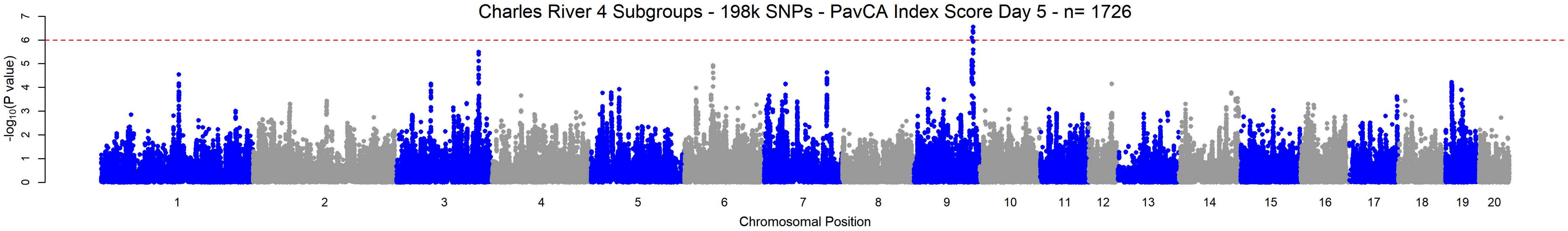
Charles River 4 Subgroups - 198k SNPs - PavCA Index Score Day 5 - n= 1726
-log10(P value)
0
1
2
3
4
5
6
7
Chromosomal Position

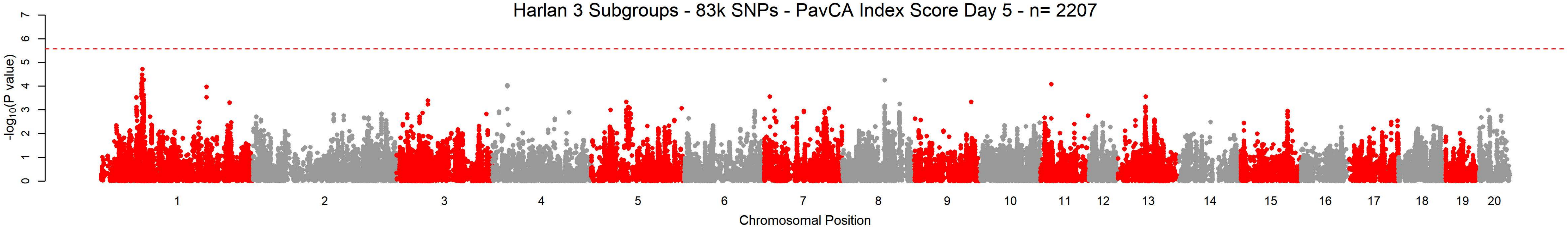
Harlan 3 Subgroups - 83k SNPs - PavCA Index Score Day 5 - n= 2207
-log10(P value)
0
1
2
3
4
5
6
7
Chromosomal Position
