## Supplementary material for "Genetic characterization of outbred Sprague Dawley rats and utility for genome-wide association studies": S5_File - Meta QQ-Plots

**Q-Q Plot Average Latency to Lever Press Day 1 - Meta-analysis of 7 Subgroups - 64k SNPs (n=3903)**

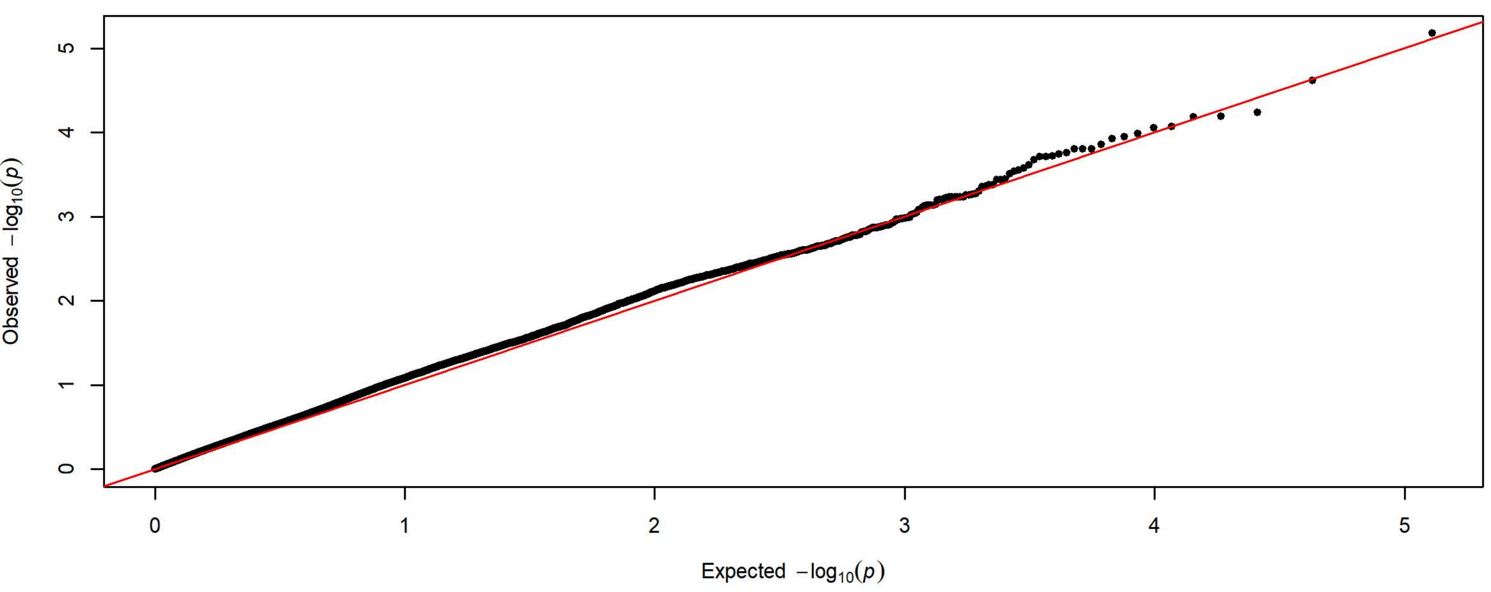


**Q-Q Plot Average Latency to Lever Press Day 1 - Charles River 4 Subgroups - 198k SNPs (n=1728)**

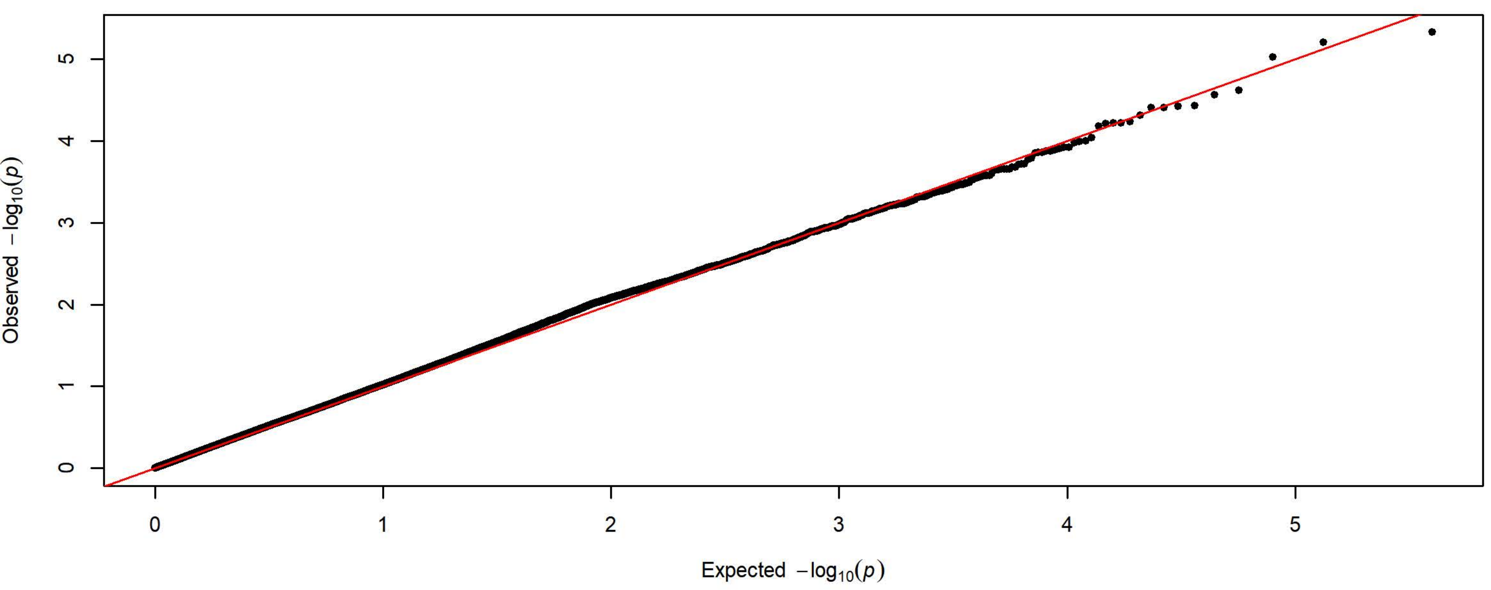


**Q-Q Plot Average Latency to Lever Press Day 1 - Harlan 3 Subgroups - 83k SNPs (n=2175)**

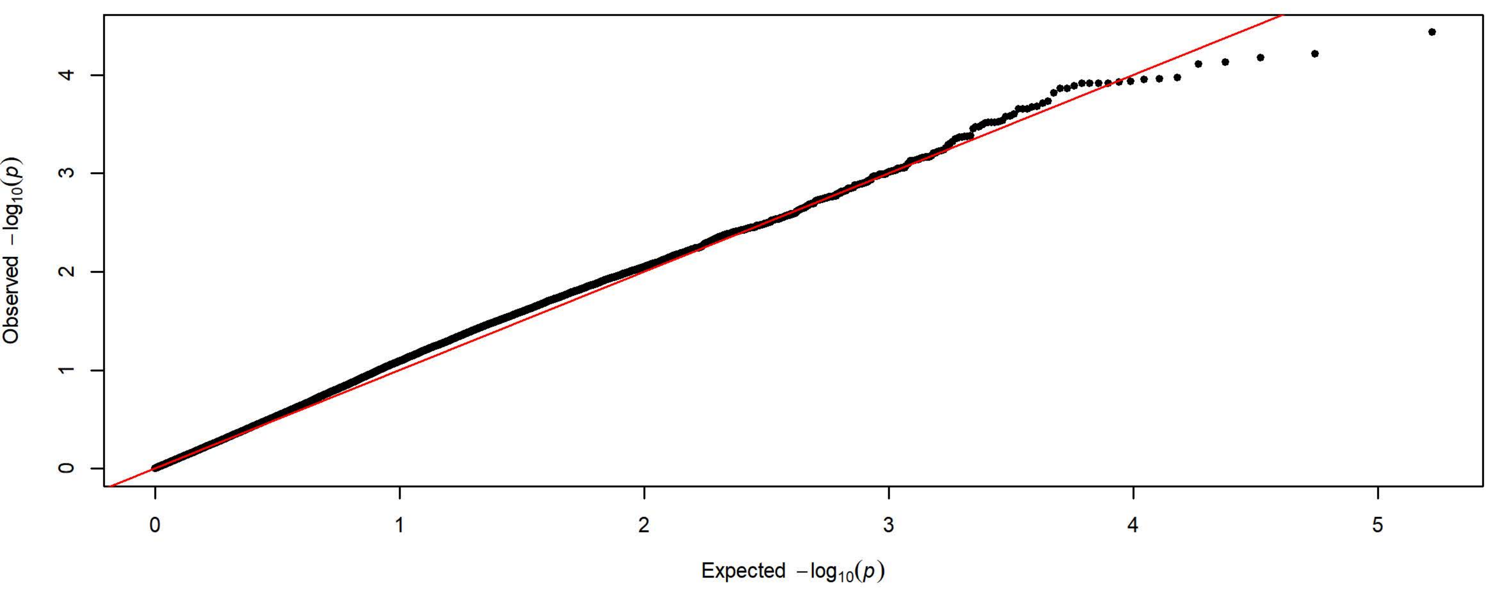

Q-Q Plot Average Latency to Lever Press Day 2 - Meta-analysis of 7 Subgroups - 64k SNPs (n=3934)

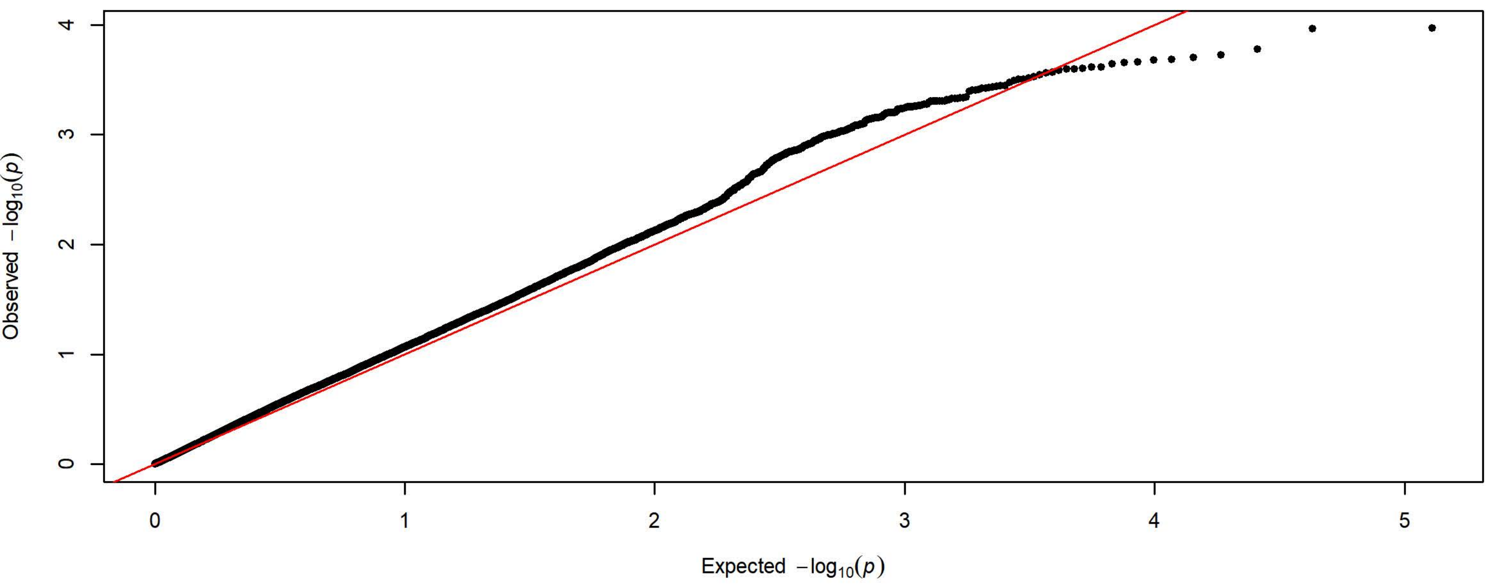


Q-Q Plot Average Latency to Lever Press Day 2 - Charles River 4 Subgroups - 198k SNPs (n=1726)

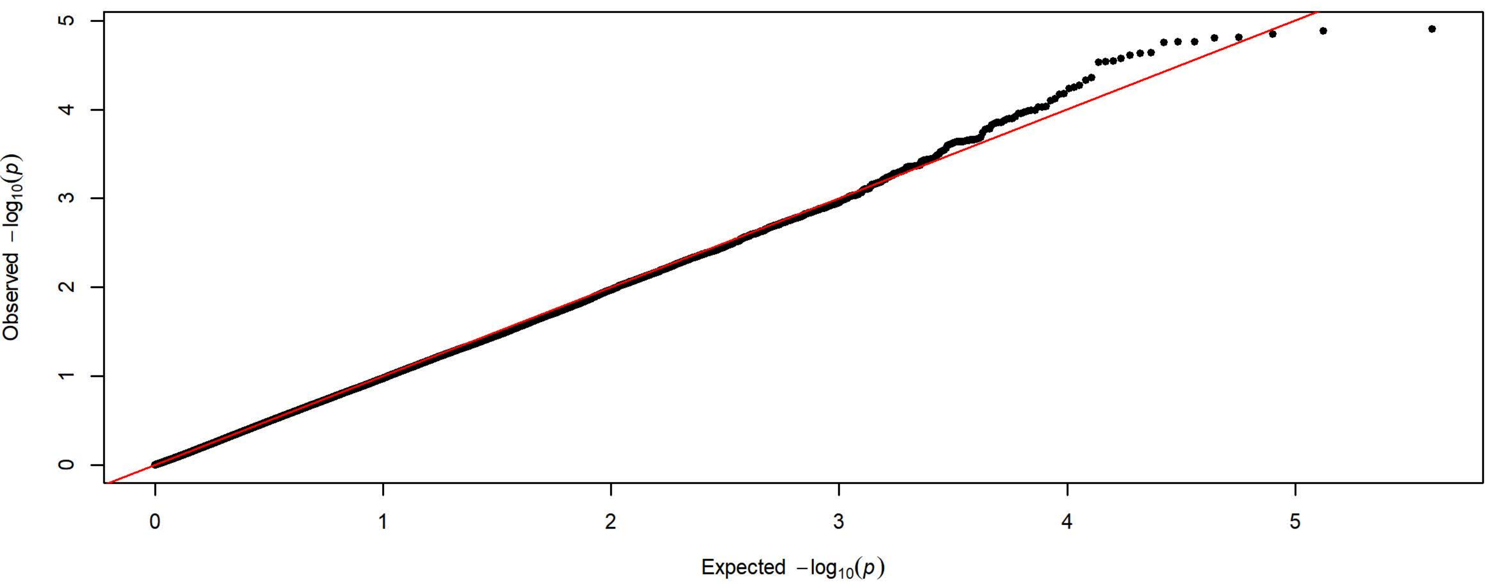


Q-Q Plot Average Latency to Lever Press Day 2 - Harlan 3 Subgroups - 83k SNPs (n=2208)

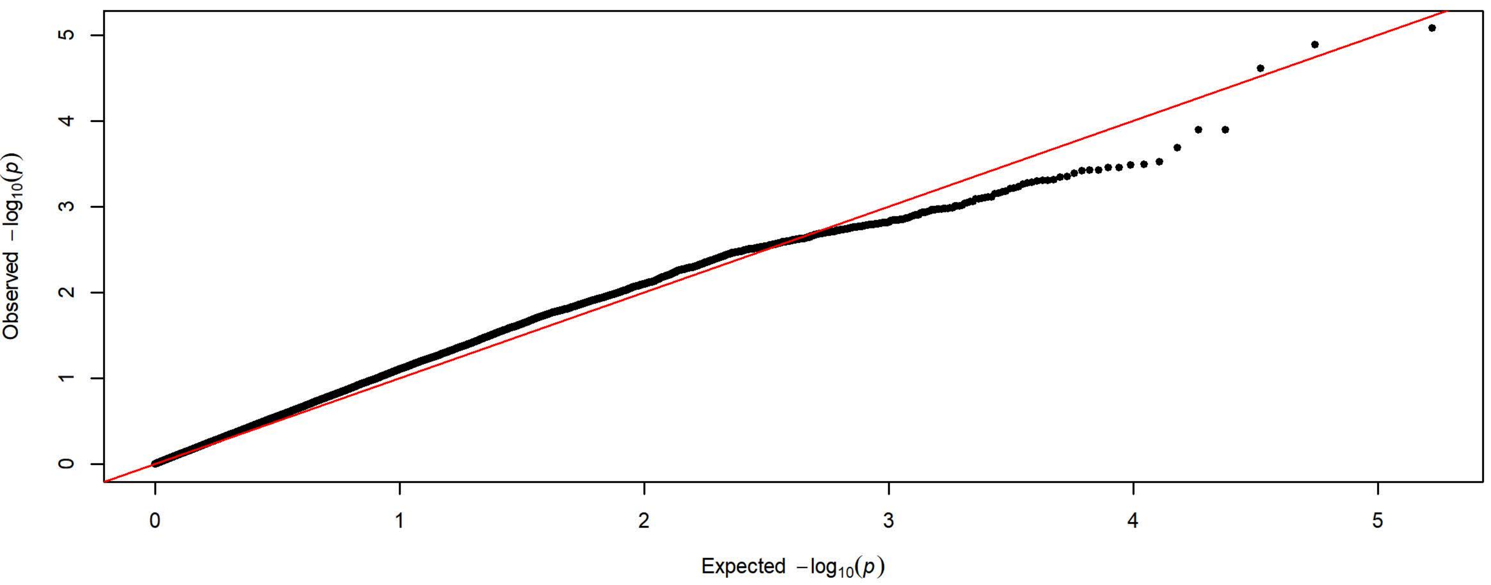

**Q-Q Plot Average Latency to Lever Press Day 3 - Meta-analysis of 7 Subgroups - 64k SNPs (n=3932)**

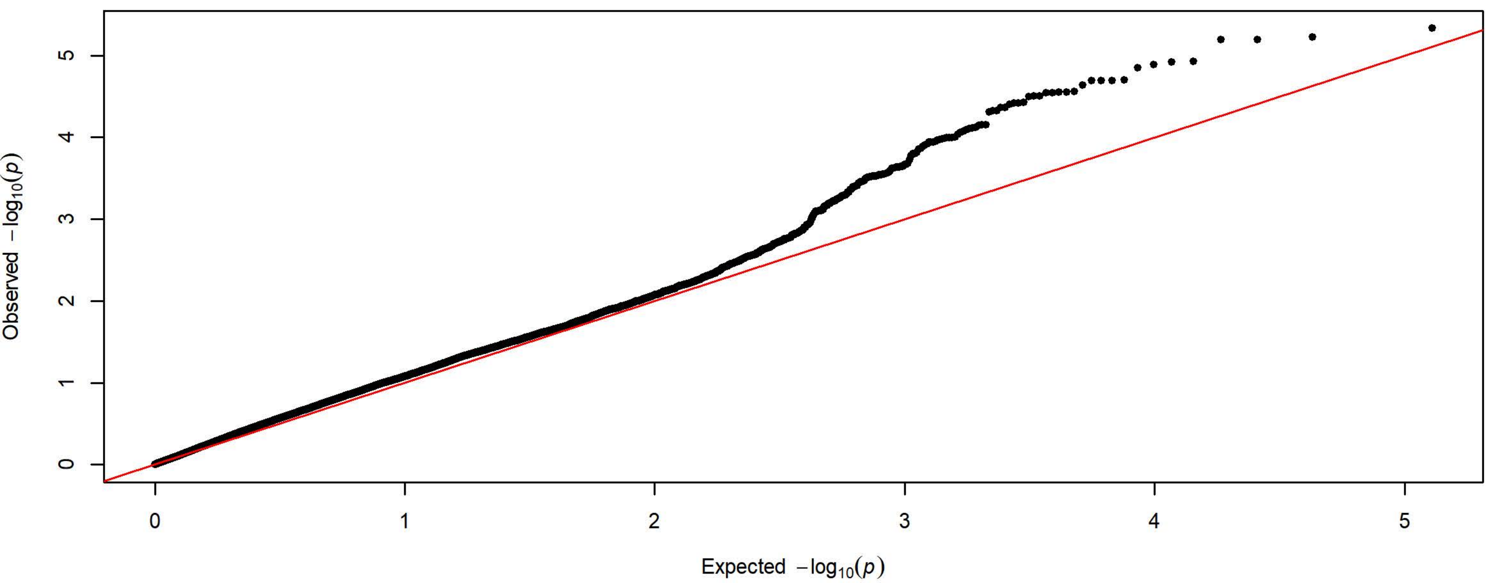


**Q-Q Plot Average Latency to Lever Press Day 3 - Charles River 4 Subgroups - 198k SNPs (n=1727)**

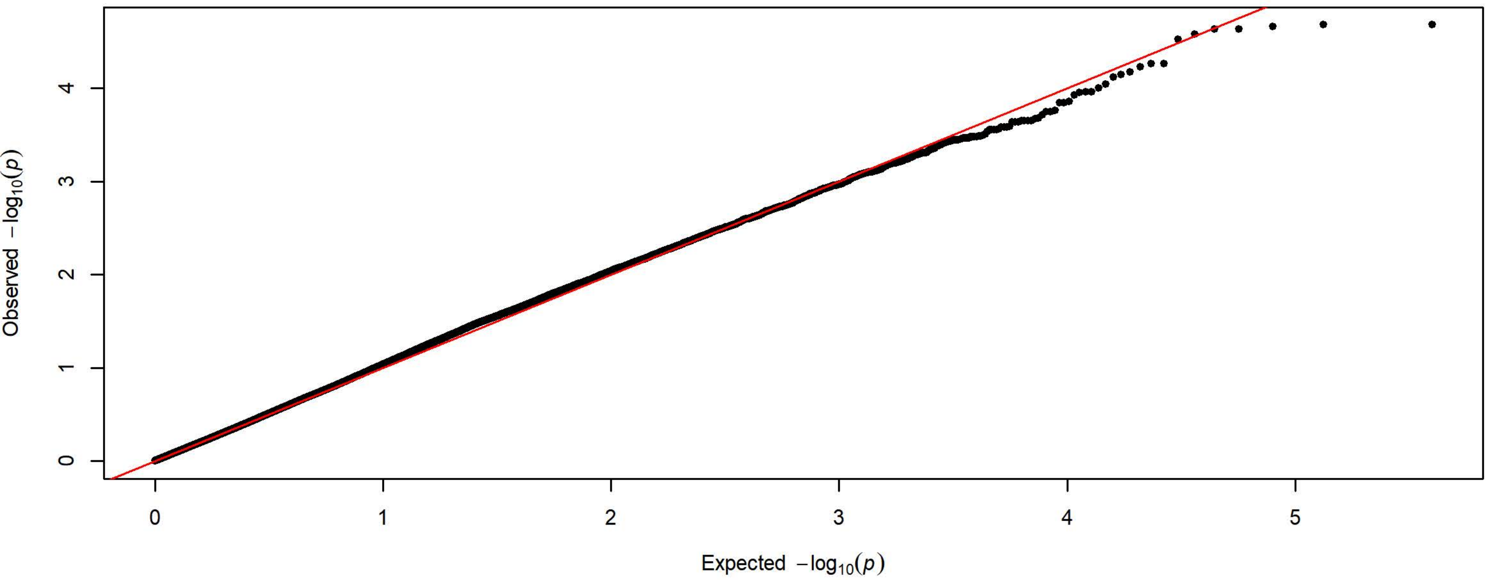


**Q-Q Plot Average Latency to Lever Press Day 3 - Harlan 3 Subgroups - 83k SNPs (n=2205)**

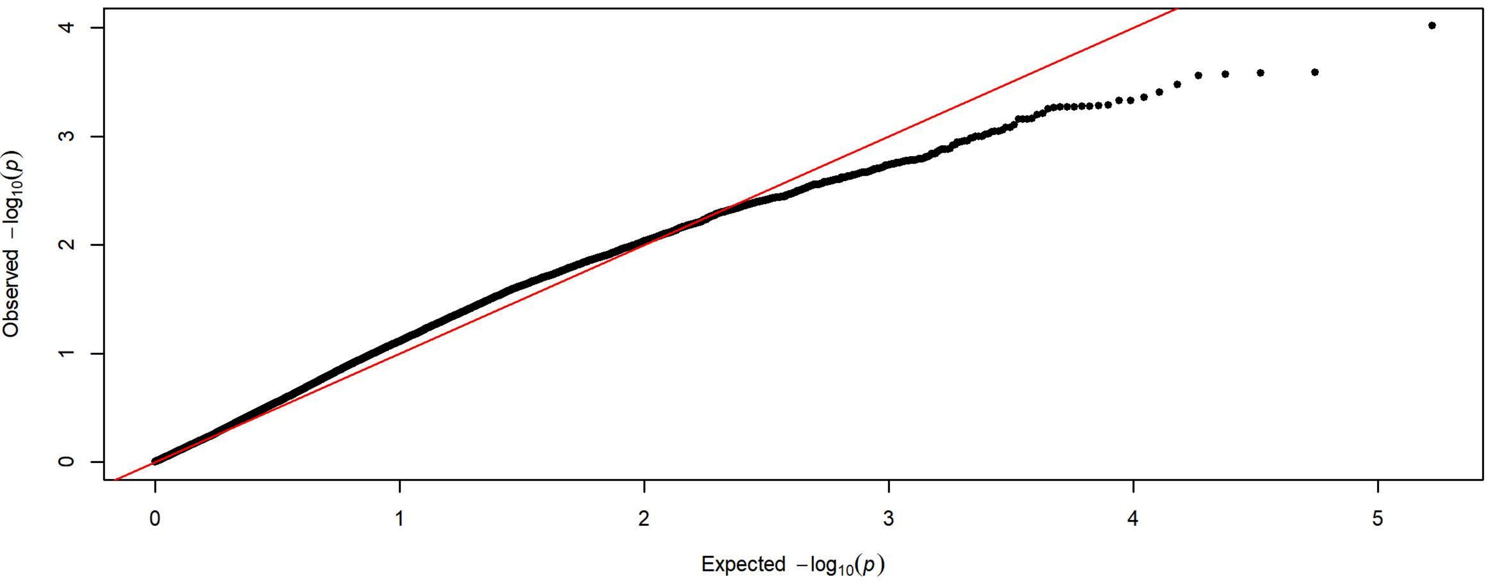

Q-Q Plot Average Latency to Lever Press Day 4 - Meta-analysis of 7 Subgroups - 64k SNPs (n=3936)

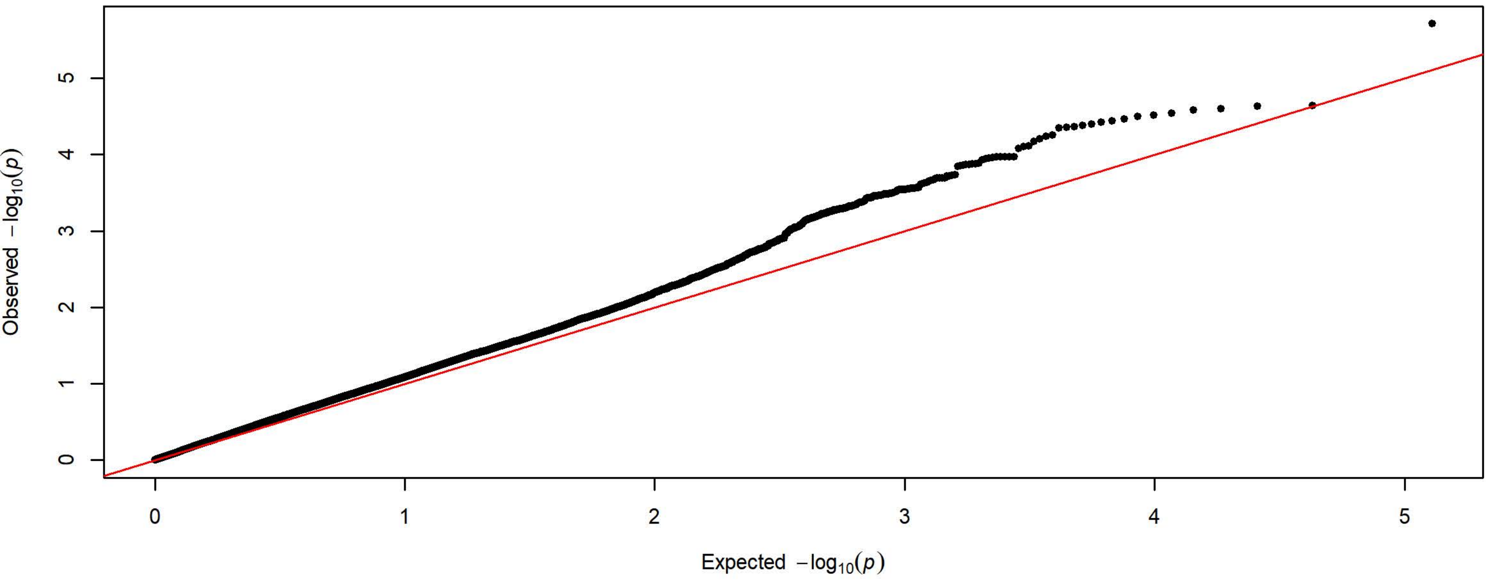


Q-Q Plot Average Latency to Lever Press Day 4 - Charles River 4 Subgroups - 198k SNPs (n=1728)

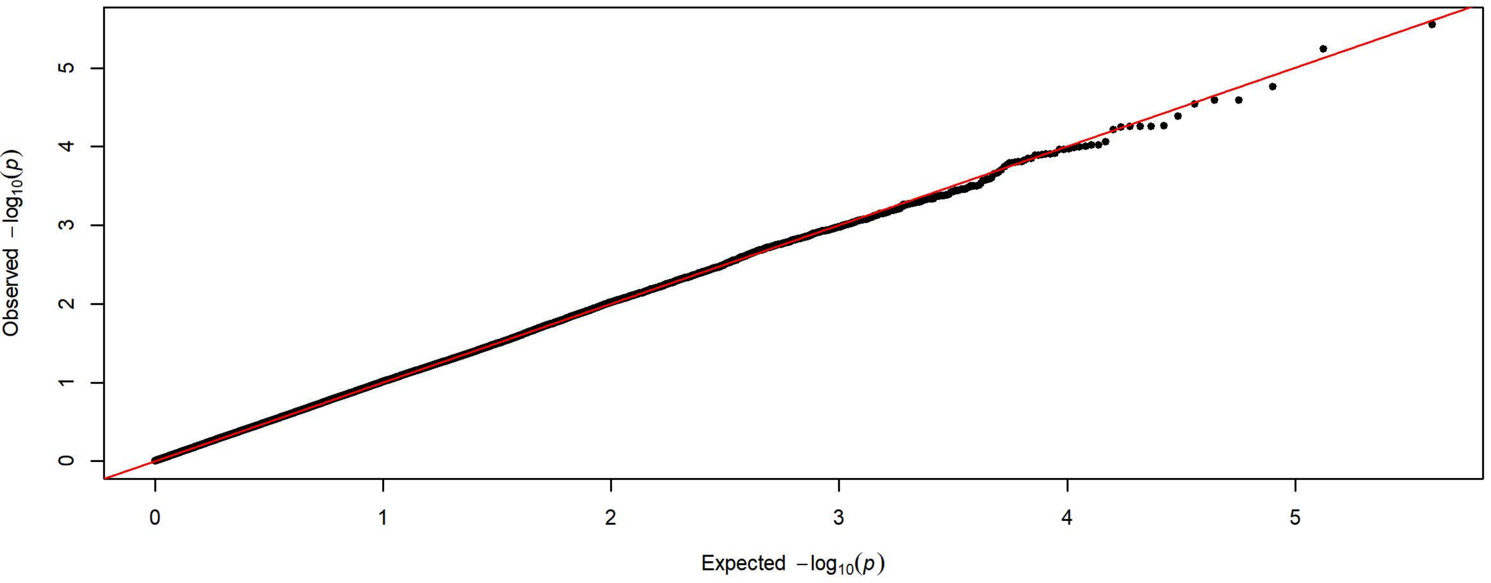


Q-Q Plot Average Latency to Lever Press Day 4 - Harlan 3 Subgroups - 83k SNPs (n=2208)

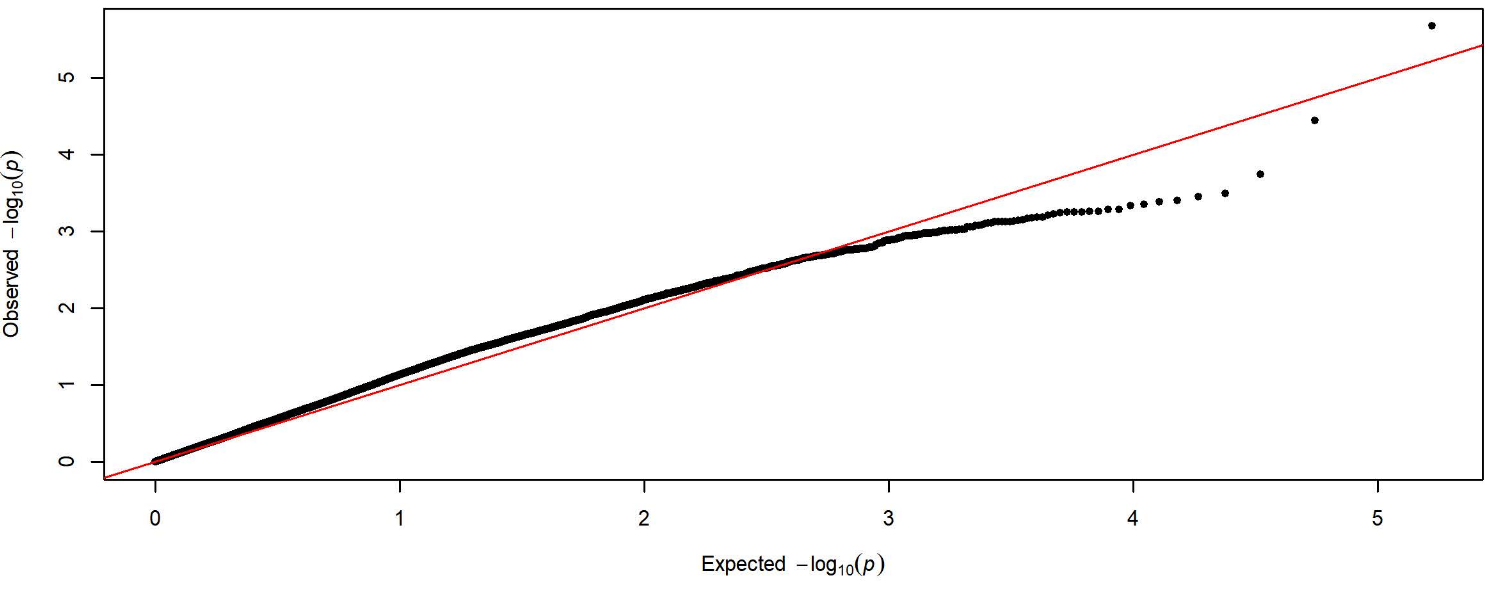

Q-Q Plot Average Latency to Lever Press Day 5 - Meta-analysis of 7 Subgroups - 64k SNPs (n=3936)

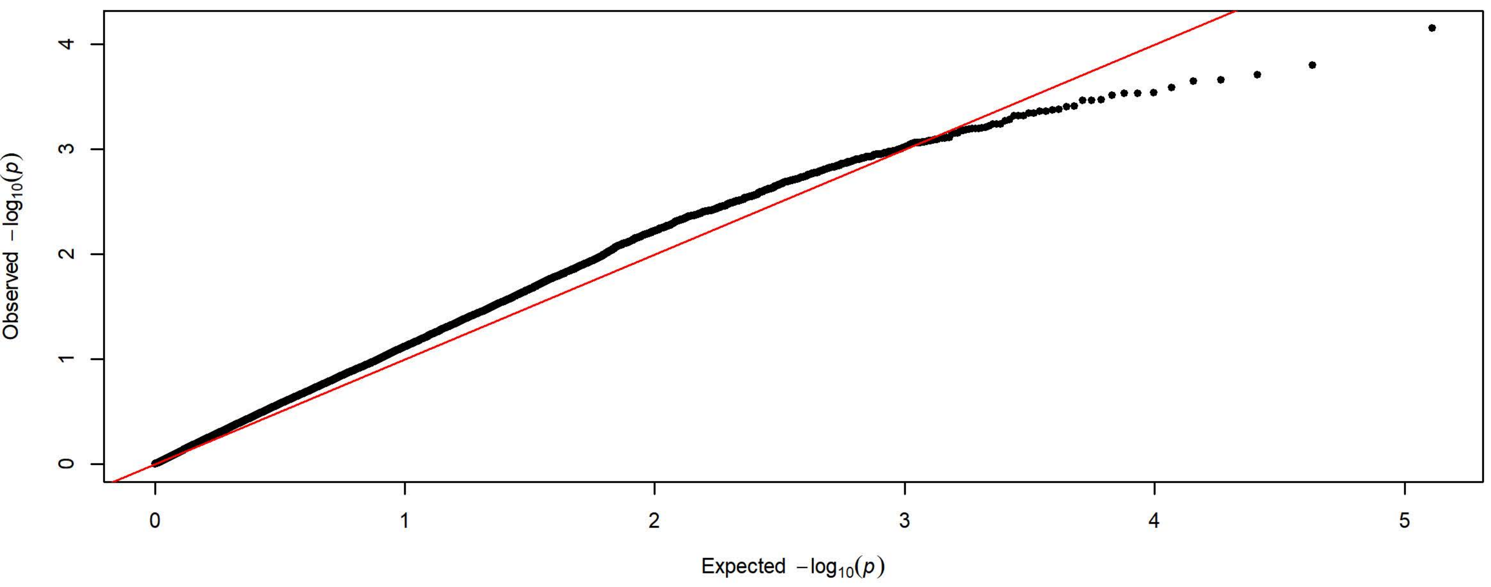


Q-Q Plot Average Latency to Lever Press Day 5 - Charles River 4 Subgroups - 198k SNPs (n=1728)

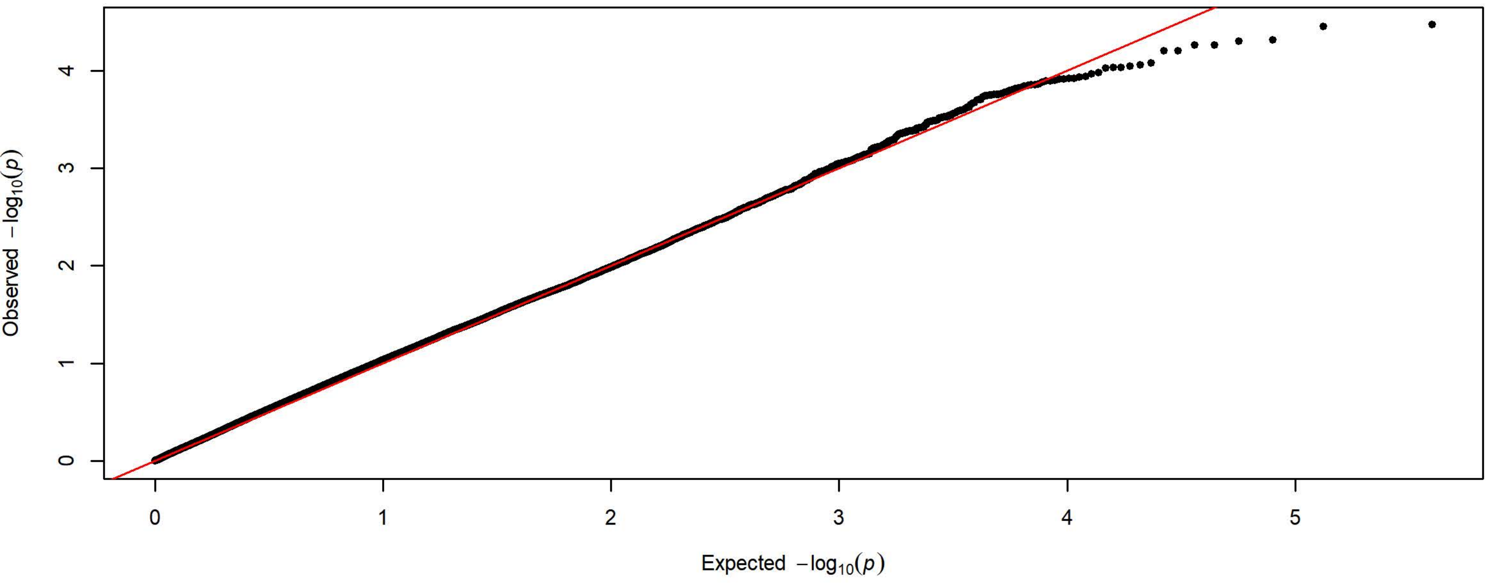


Q-Q Plot Average Latency to Lever Press Day 5 - Harlan 3 Subgroups - 83k SNPs (n=2208)

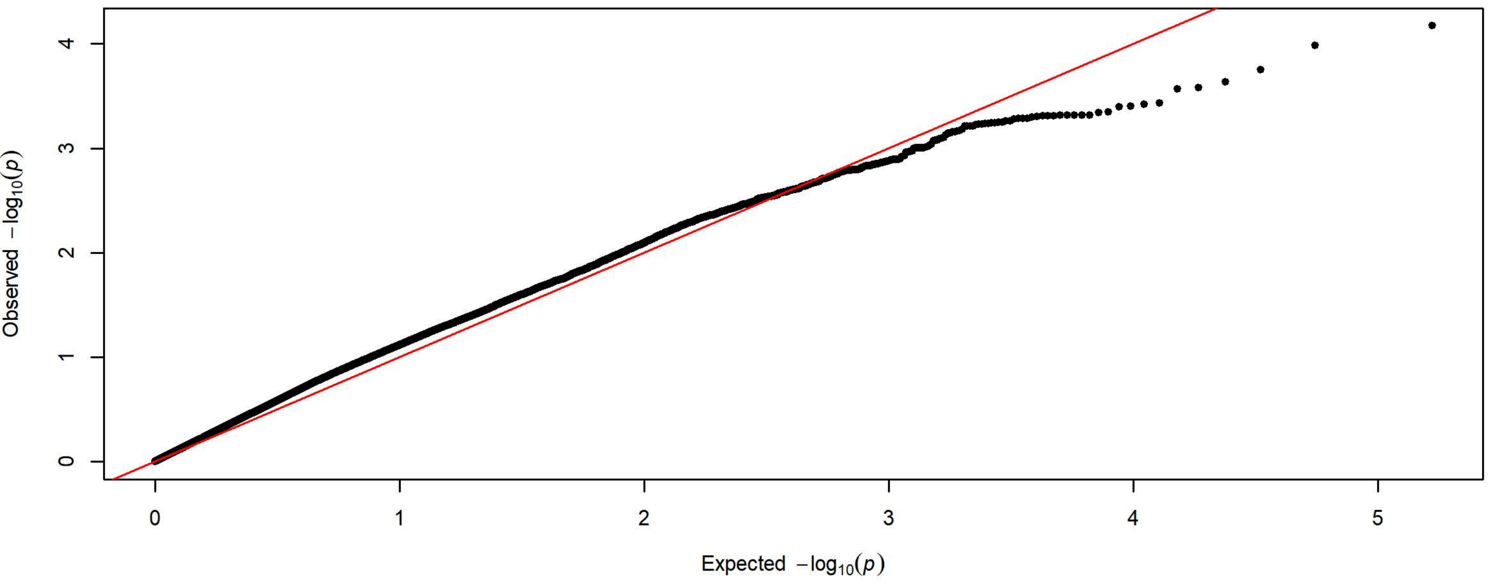

**Q-Q Plot Average Latency to Magazine Entry Day 1 - Meta-analysis of 7 Subgroups - 64k SNPs (n=3903)**

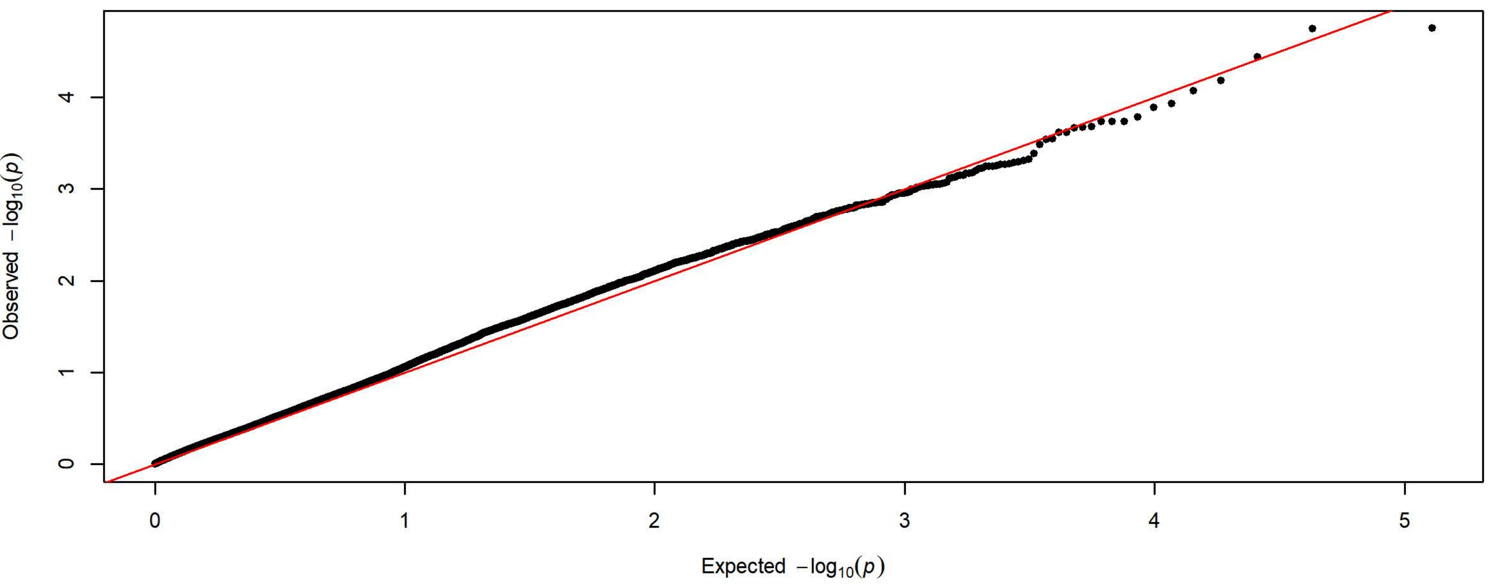


**Q-Q Plot Average Latency to Magazine Entry Day 1 - Charles River 4 Subgroups - 198k SNPs (n=1728)**

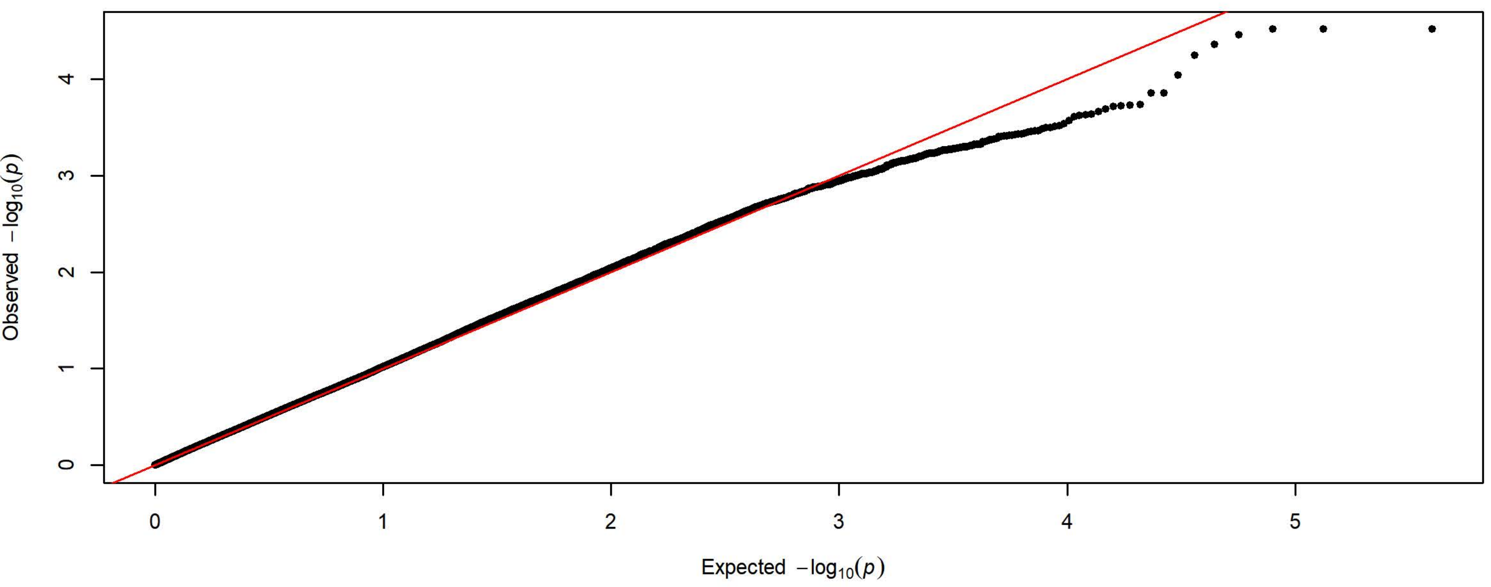


**Q-Q Plot Average Latency to Magazine Entry Day 1 - Harlan 3 Subgroups - 83k SNPs (n=2175)**

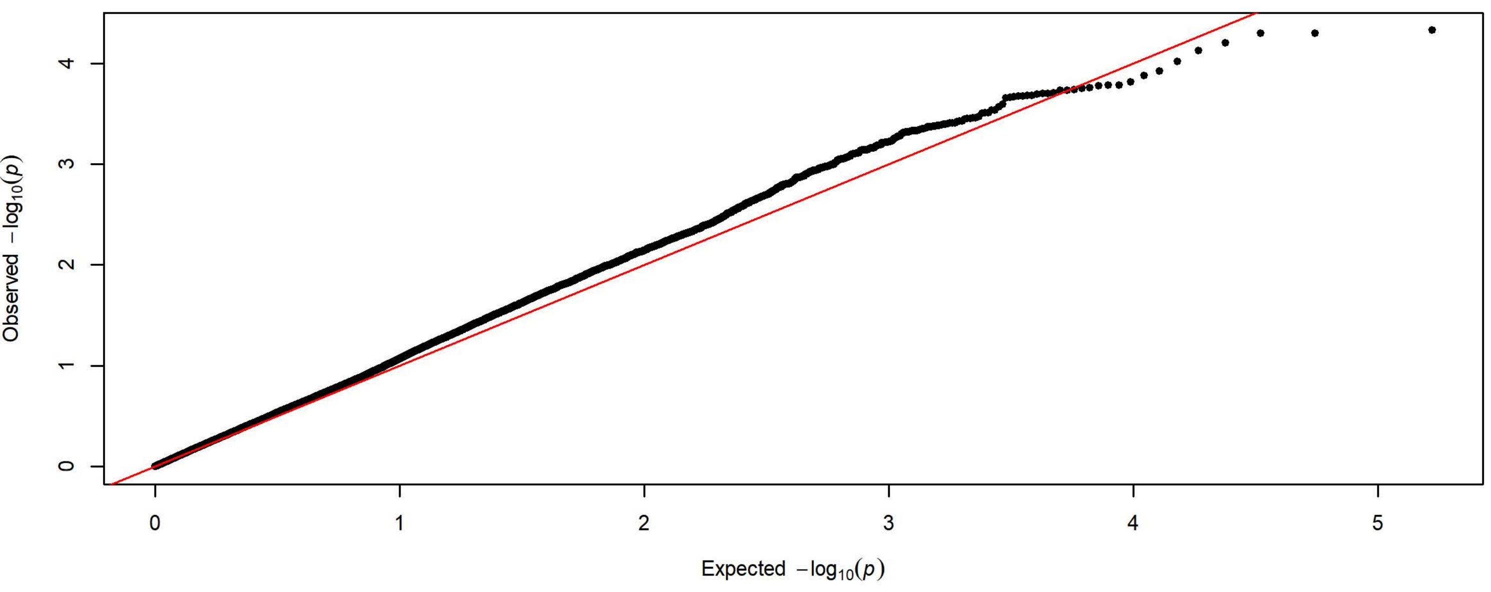

**Q-Q Plot Average Latency to Magazine Entry Day 2 - Meta-analysis of 7 Subgroups - 64k SNPs (n=3934)**

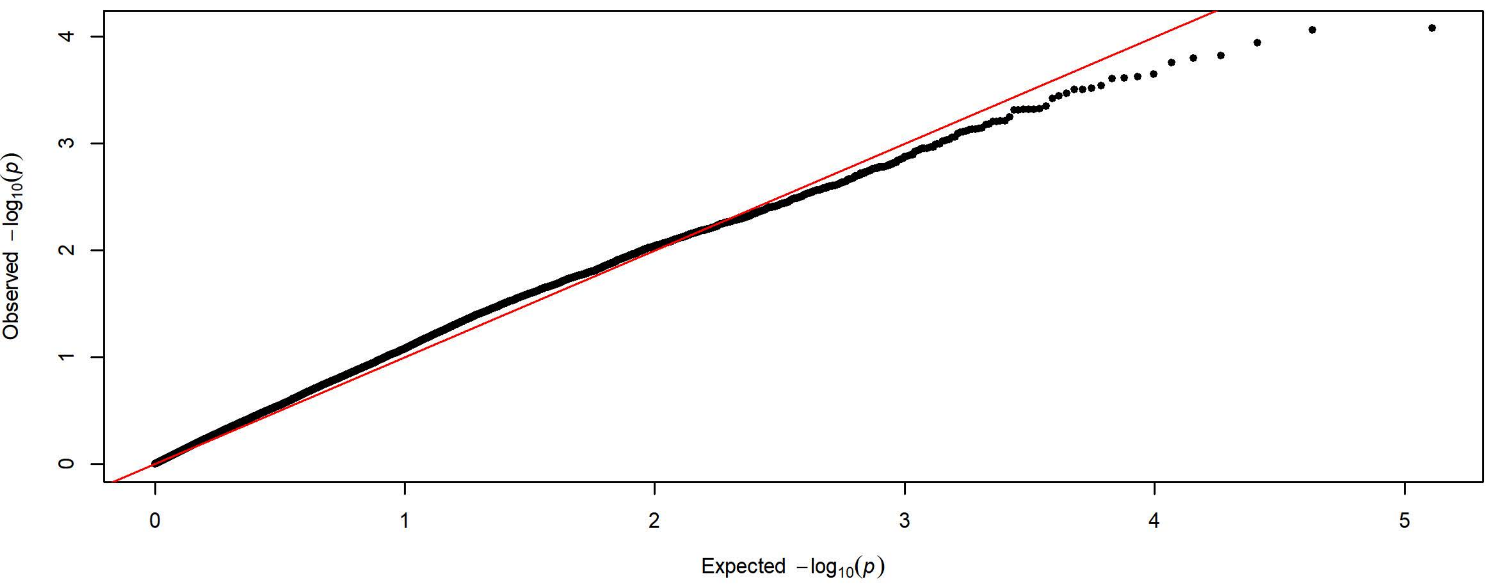


**Q-Q Plot Average Latency to Magazine Entry Day 2 - Charles River 4 Subgroups - 198k SNPs (n=1726)**

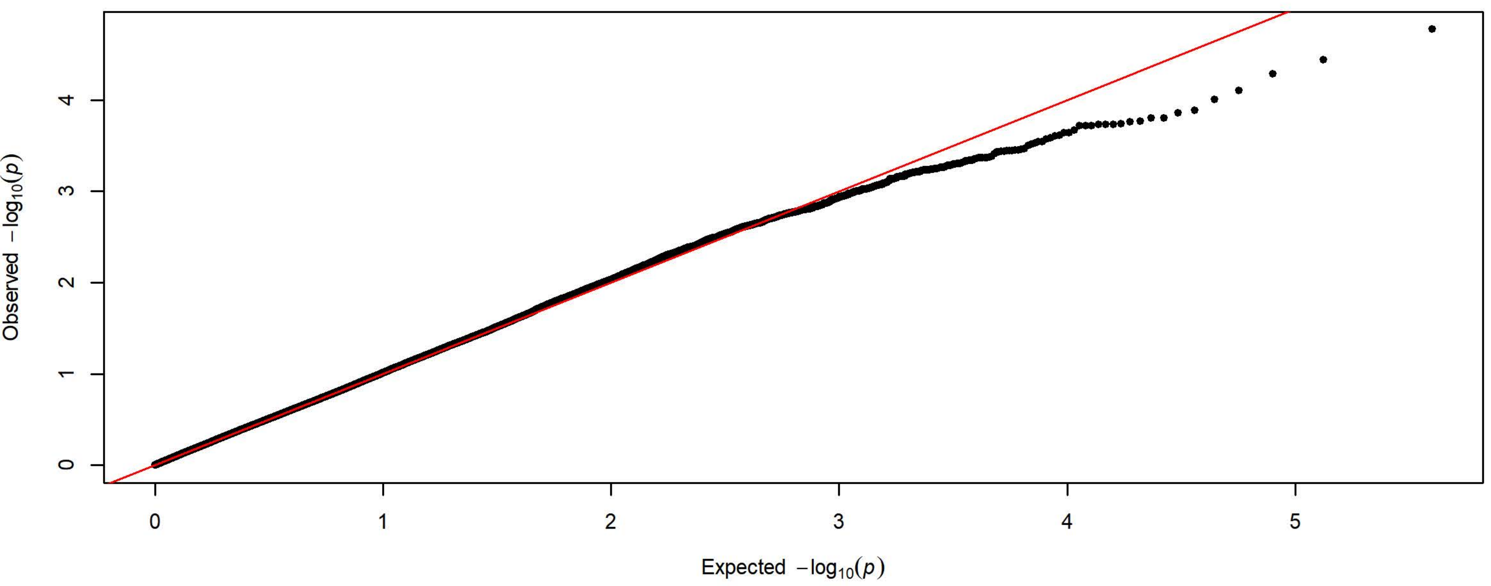


**Q-Q Plot Average Latency to Magazine Entry Day 2 - Harlan 3 Subgroups - 83k SNPs (n=2208)**

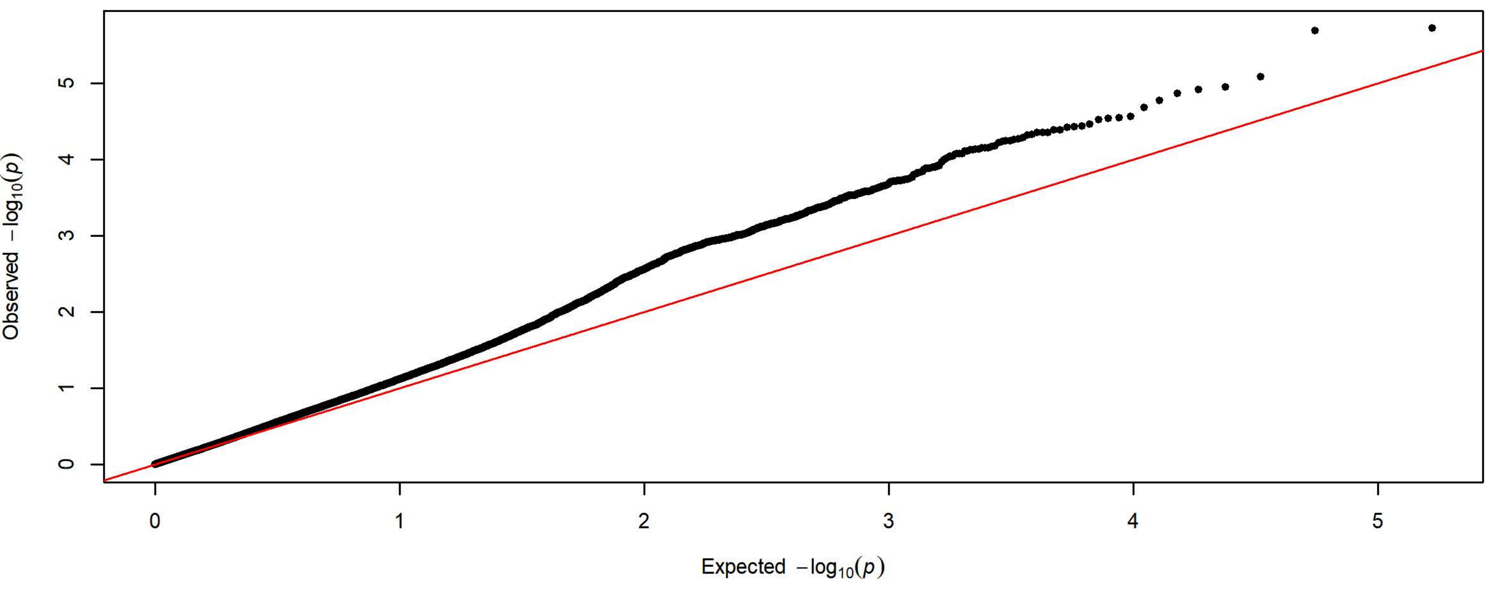

**Q-Q Plot Average Latency to Magazine Entry Day 3 - Meta-analysis of 7 Subgroups - 64k SNPs (n=3932)**

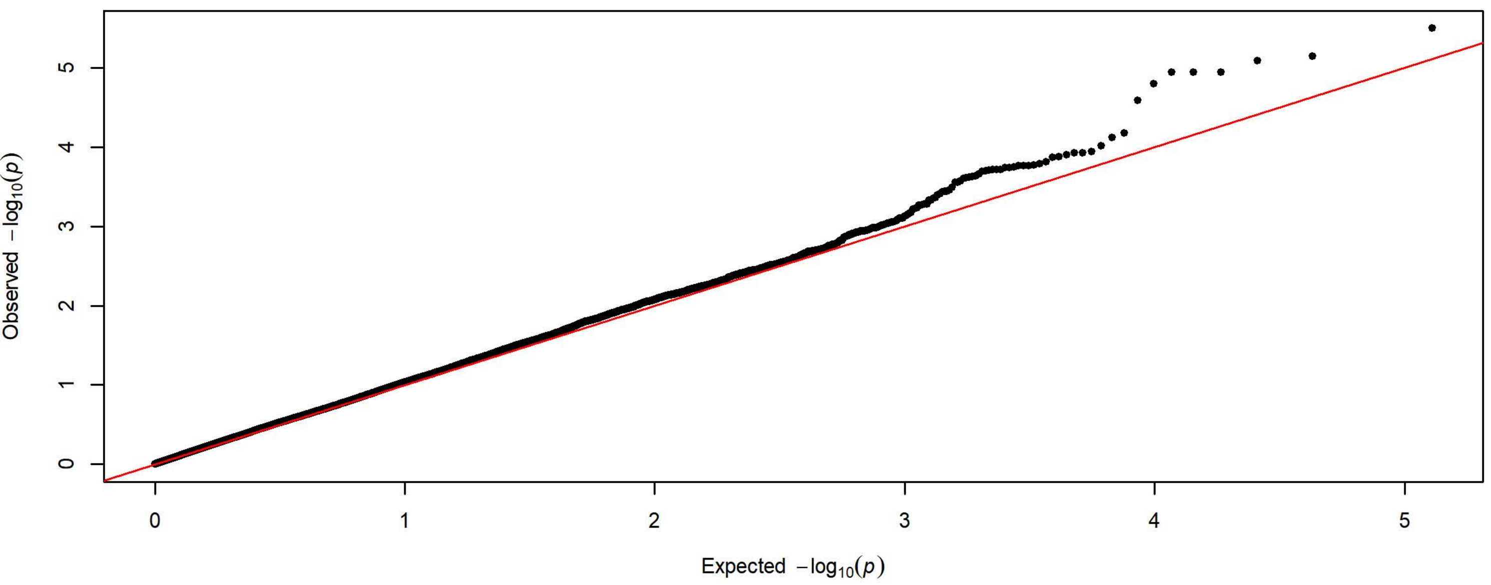


**Q-Q Plot Average Latency to Magazine Entry Day 3 - Charles River 4 Subgroups - 198k SNPs (n=1727)**

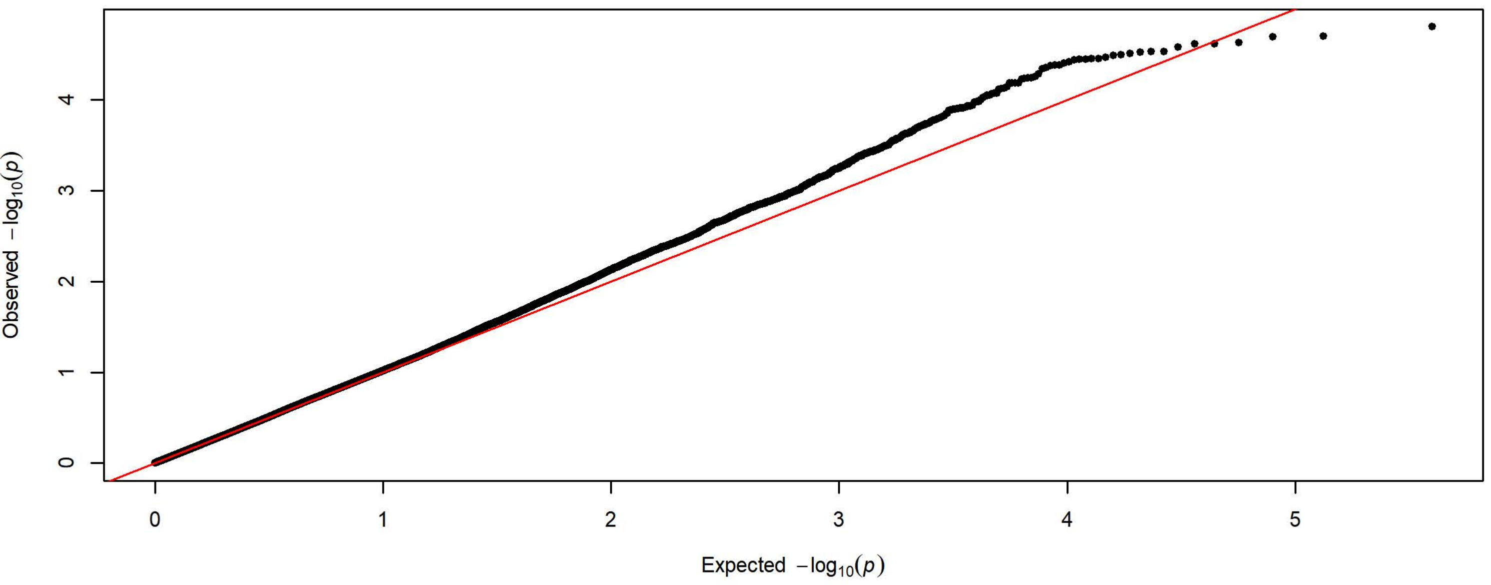


**Q-Q Plot Average Latency to Magazine Entry Day 3 - Harlan 3 Subgroups - 83k SNPs (n=2205)**

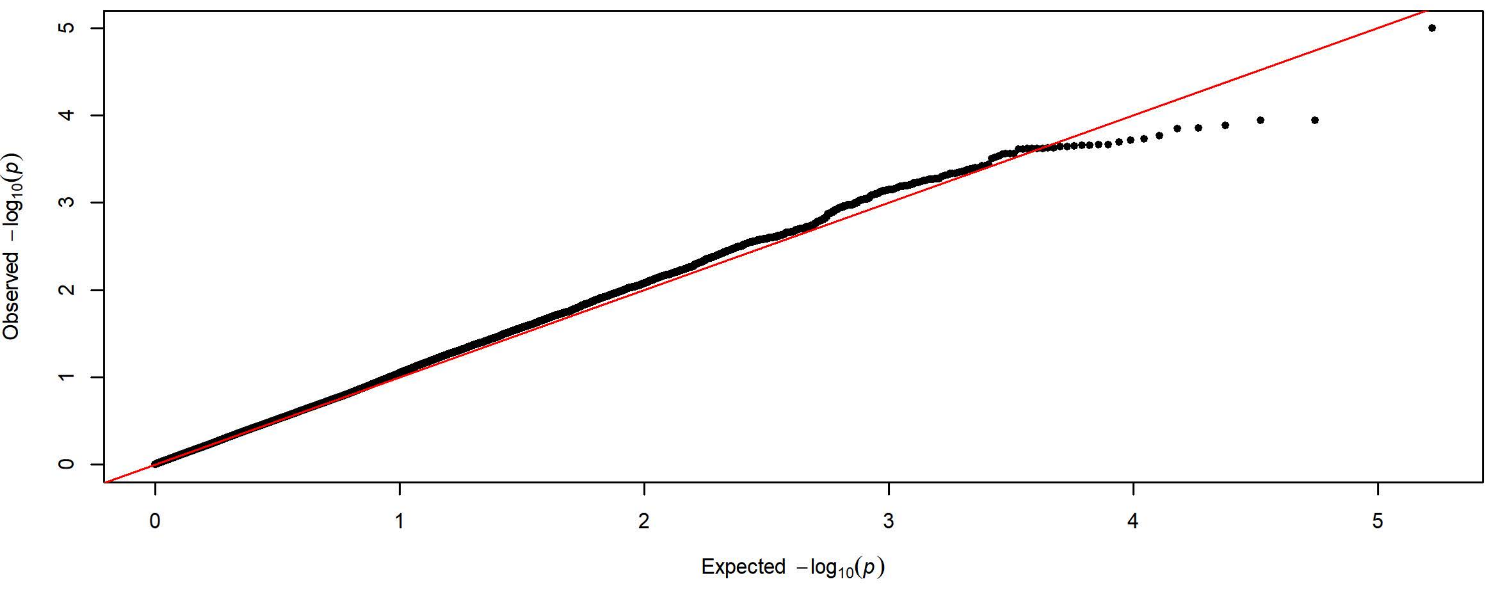

**Q-Q Plot Average Latency to Magazine Entry Day 4 - Meta-analysis of 7 Subgroups - 64k SNPs (n=3936)**

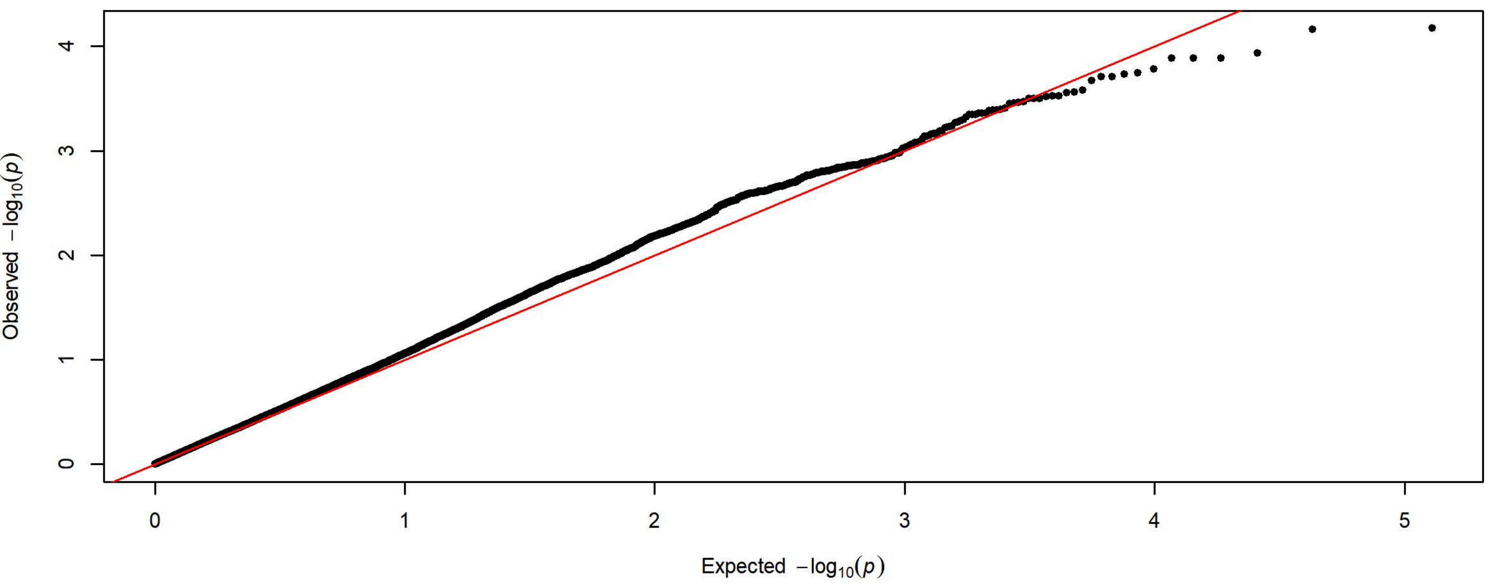


**Q-Q Plot Average Latency to Magazine Entry Day 4 - Charles River 4 Subgroups - 198k SNPs (n=1728)**

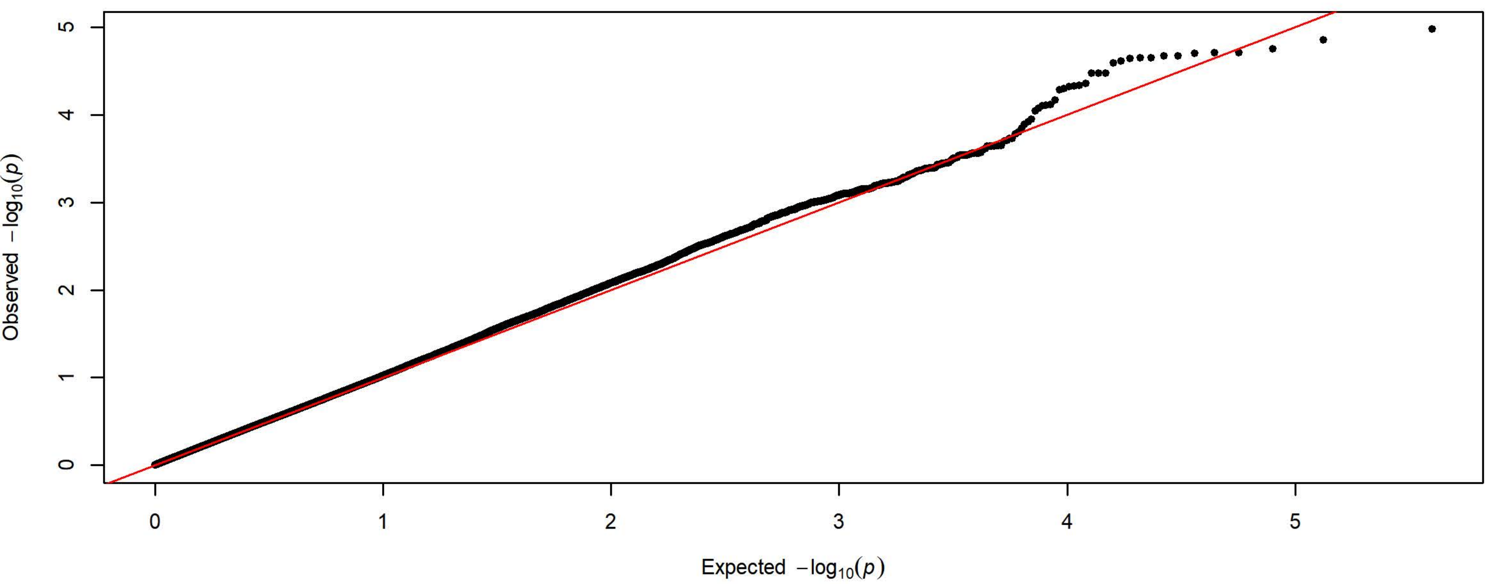


**Q-Q Plot Average Latency to Magazine Entry Day 4 - Harlan 3 Subgroups - 83k SNPs (n=2208)**

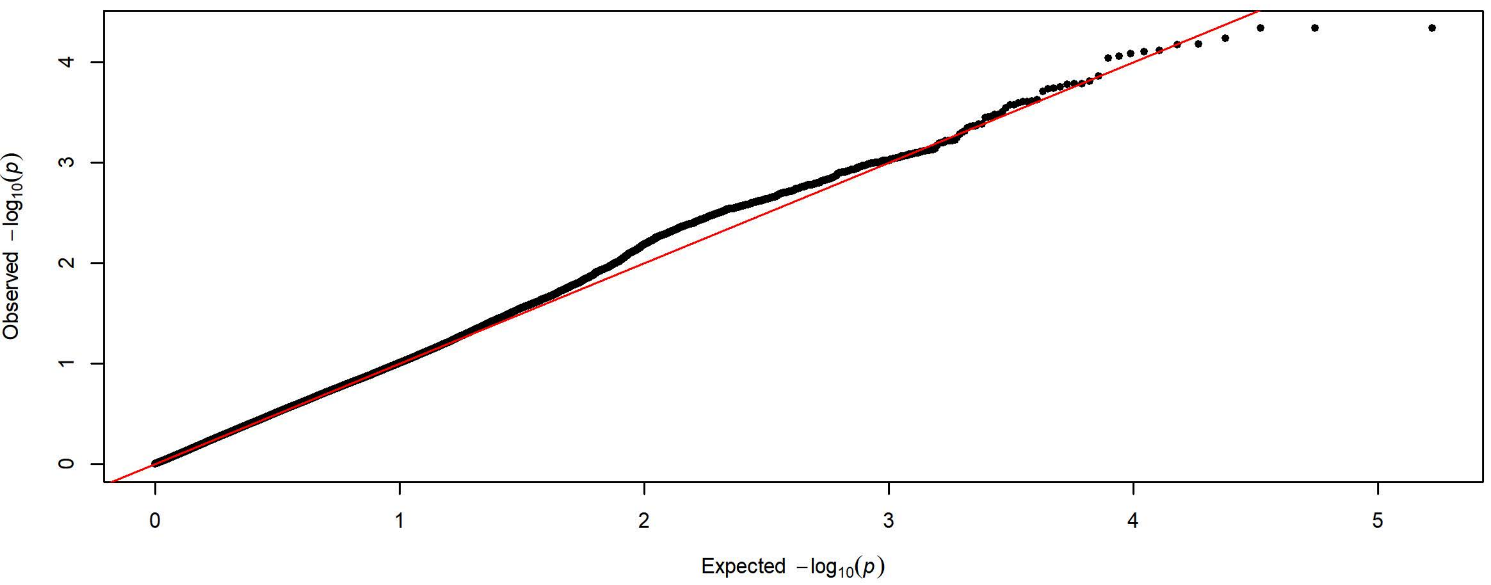

**Q-Q Plot Average Latency to Magazine Entry Day 5 - Meta-analysis of 7 Subgroups - 64k SNPs (n=3936)**

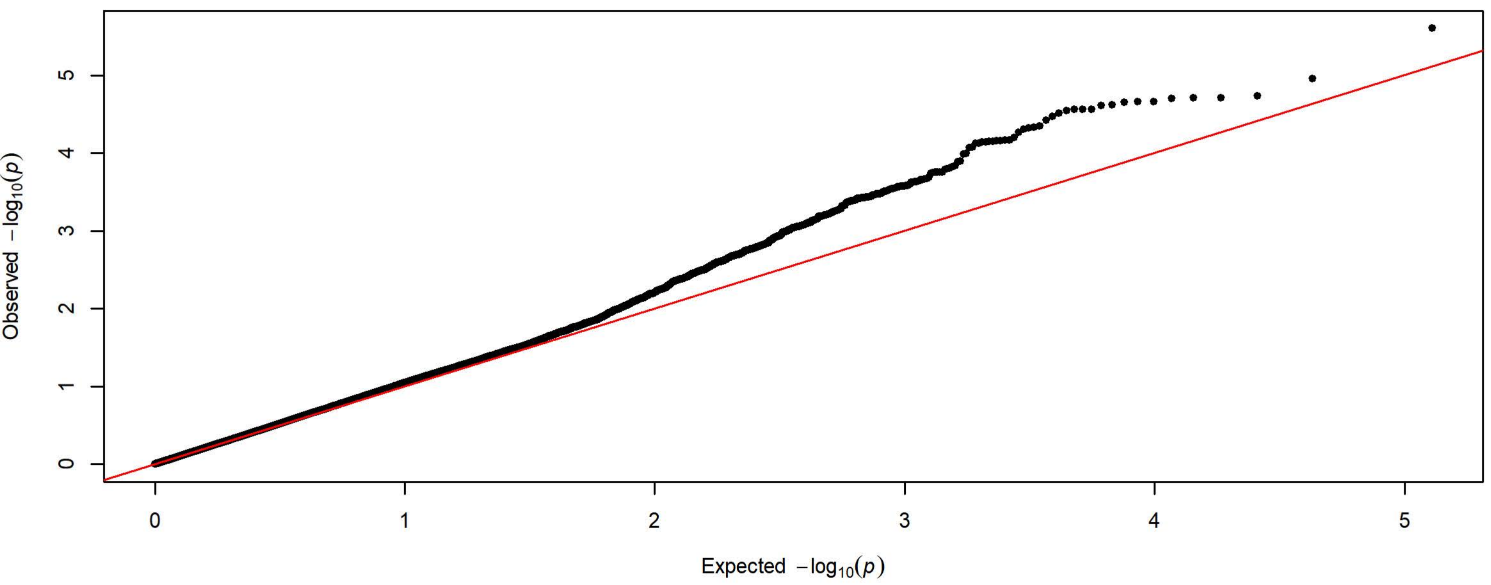


**Q-Q Plot Average Latency to Magazine Entry Day 5 - Charles River 4 Subgroups - 198k SNPs (n=1728)**

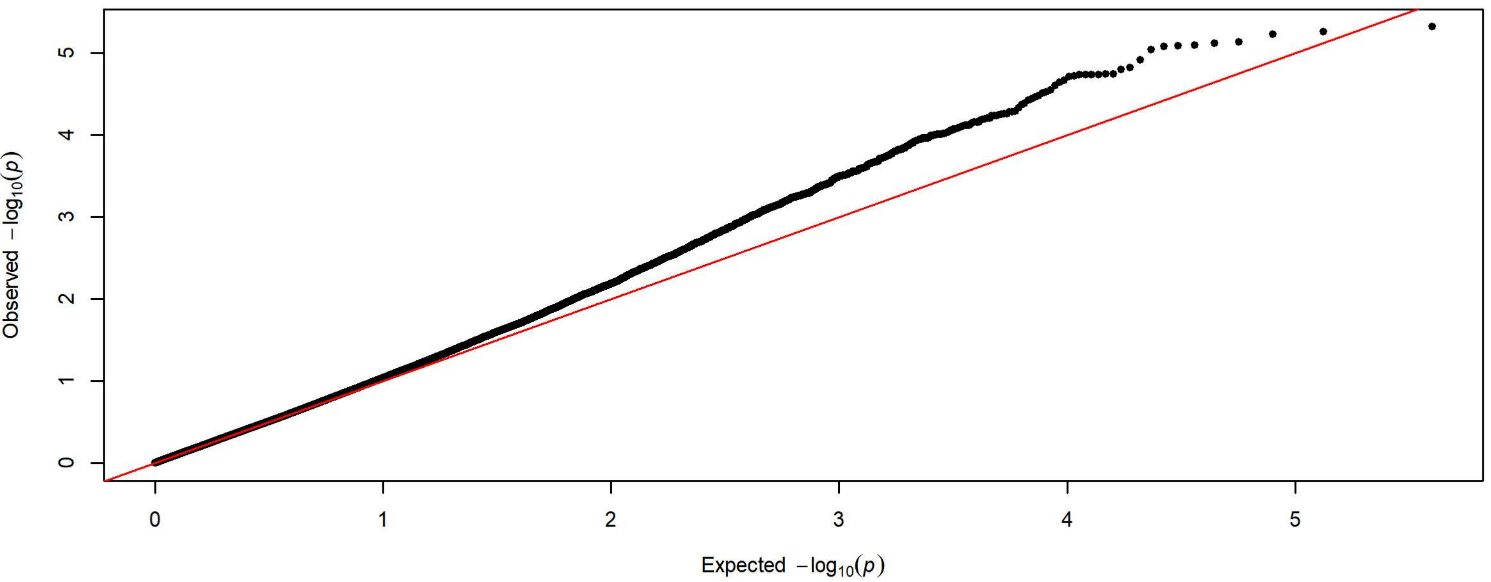


**Q-Q Plot Average Latency to Magazine Entry Day 5 - Harlan 3 Subgroups - 83k SNPs (n=2208)**

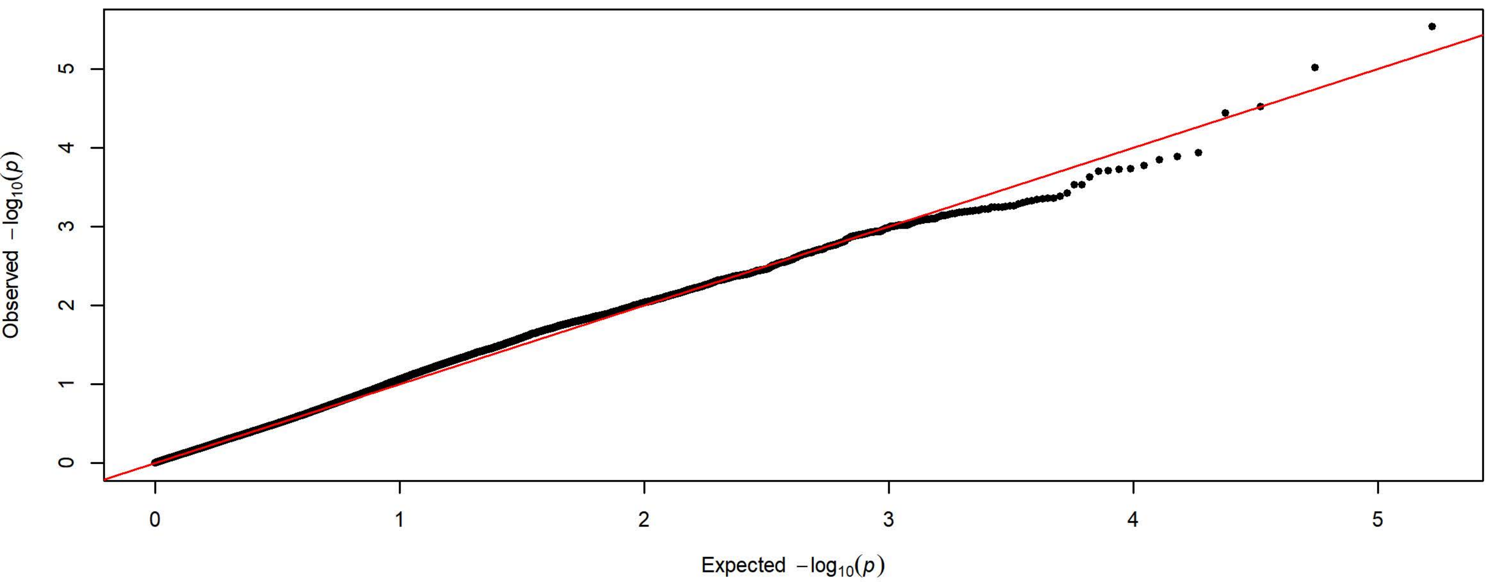

**Q-Q Plot PavCA Index Score Day 1 - Meta-analysis of 7 Subgroups - 64k SNPs (n=3880)**

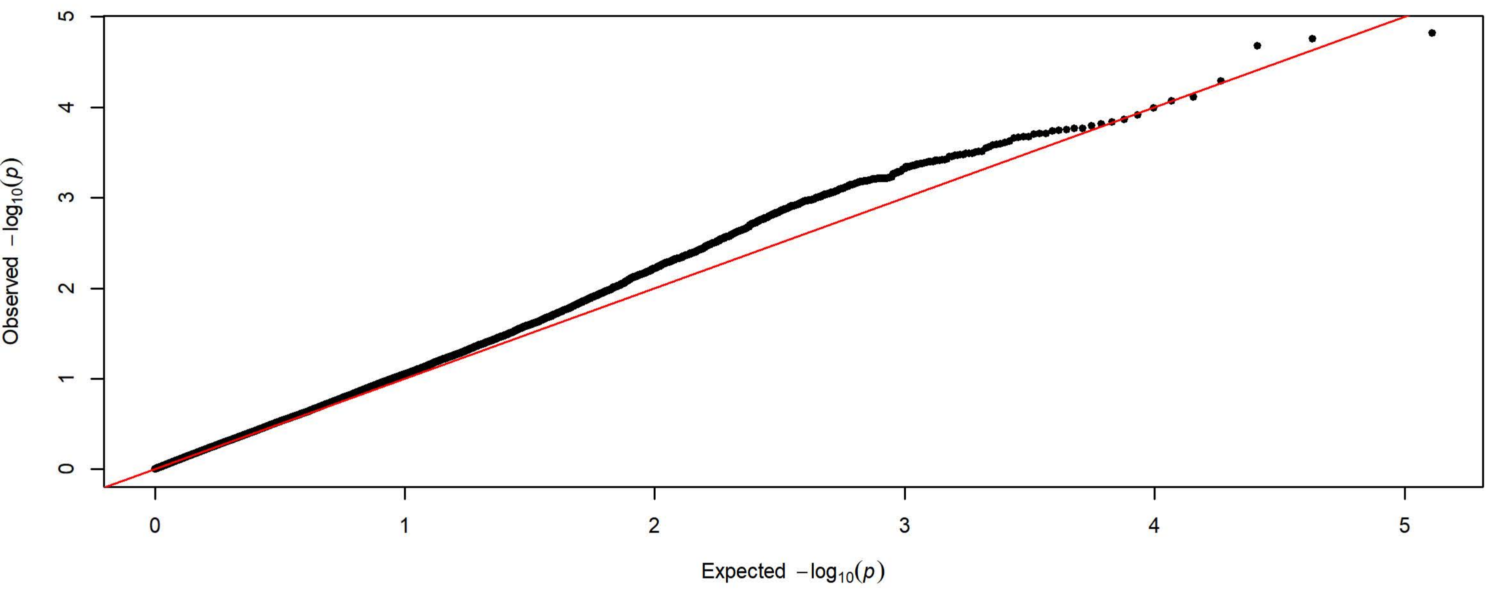


**Q-Q Plot PavCA Index Score Day 1 - Charles River 4 Subgroups - 198k SNPs (n=1720)**

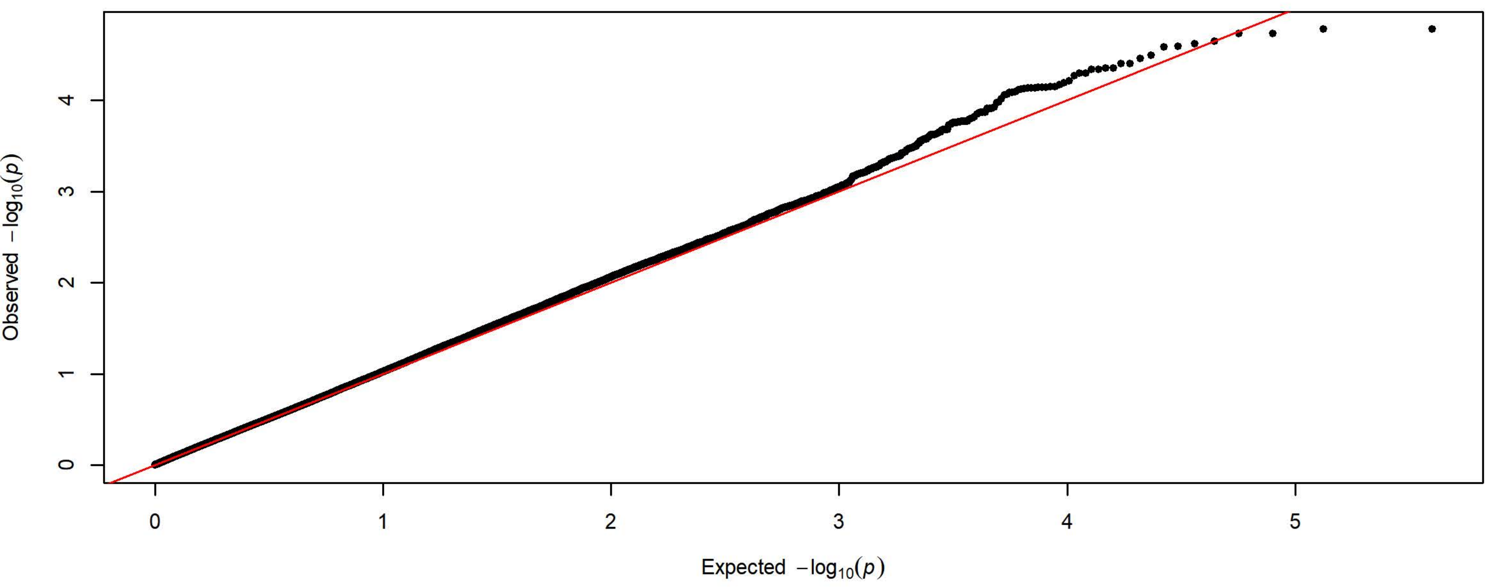


**Q-Q Plot PavCA Index Score Day 1 - Harlan 3 Subgroups - 83k SNPs (n=2160)**

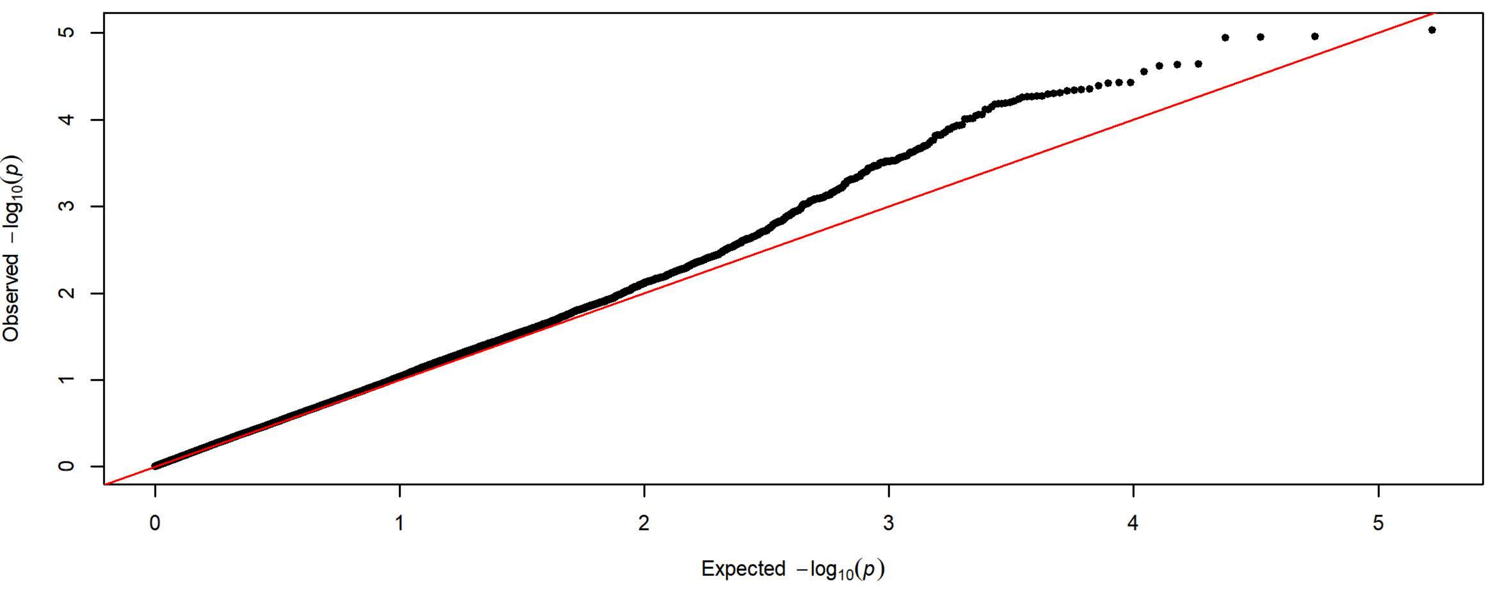

**Q-Q Plot PavCA Index Score Day 2 - Meta-analysis of 7 Subgroups - 64k SNPs (n=3919)**

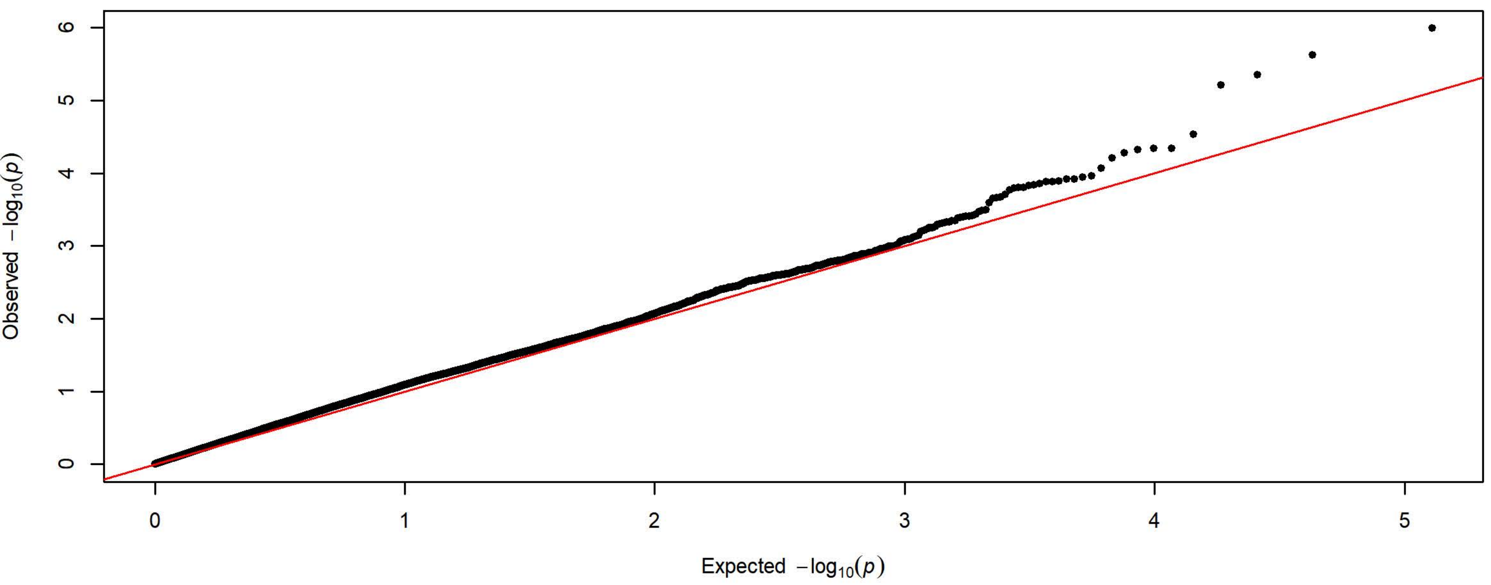


**Q-Q Plot PavCA Index Score Day 2 - Charles River 4 Subgroups - 198k SNPs (n=1719)**

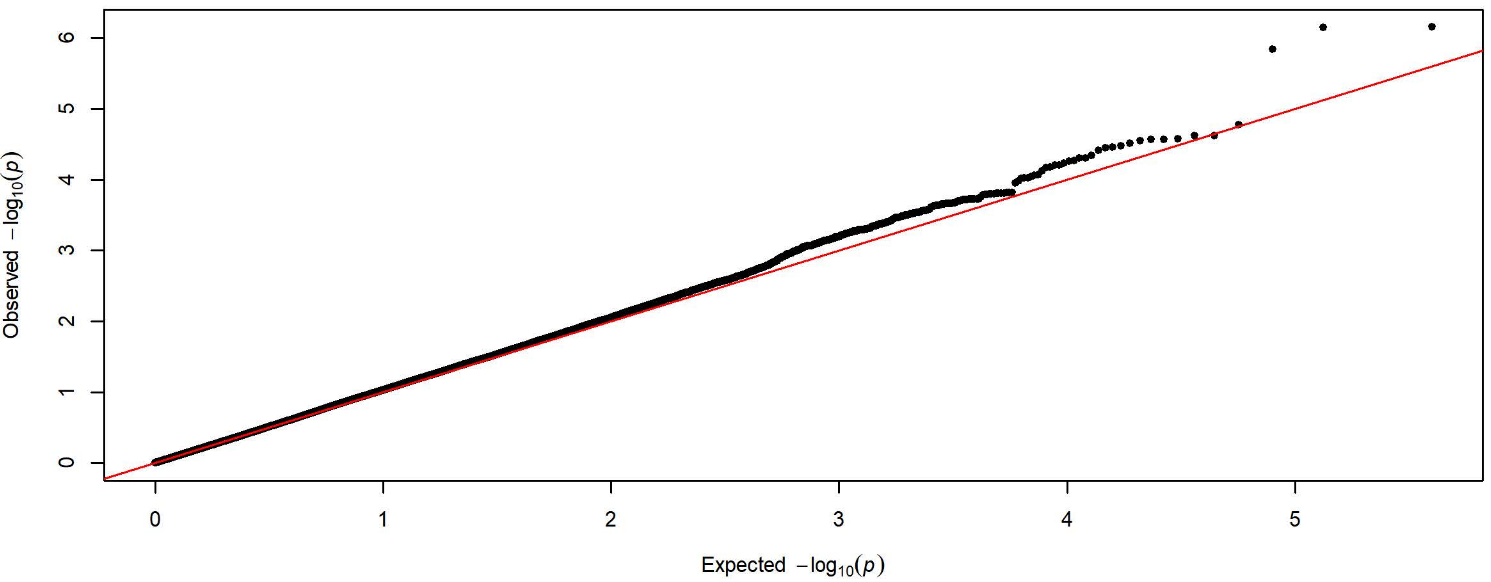


**Q-Q Plot PavCA Index Score Day 2 - Harlan 3 Subgroups - 83k SNPs (n=2200)**

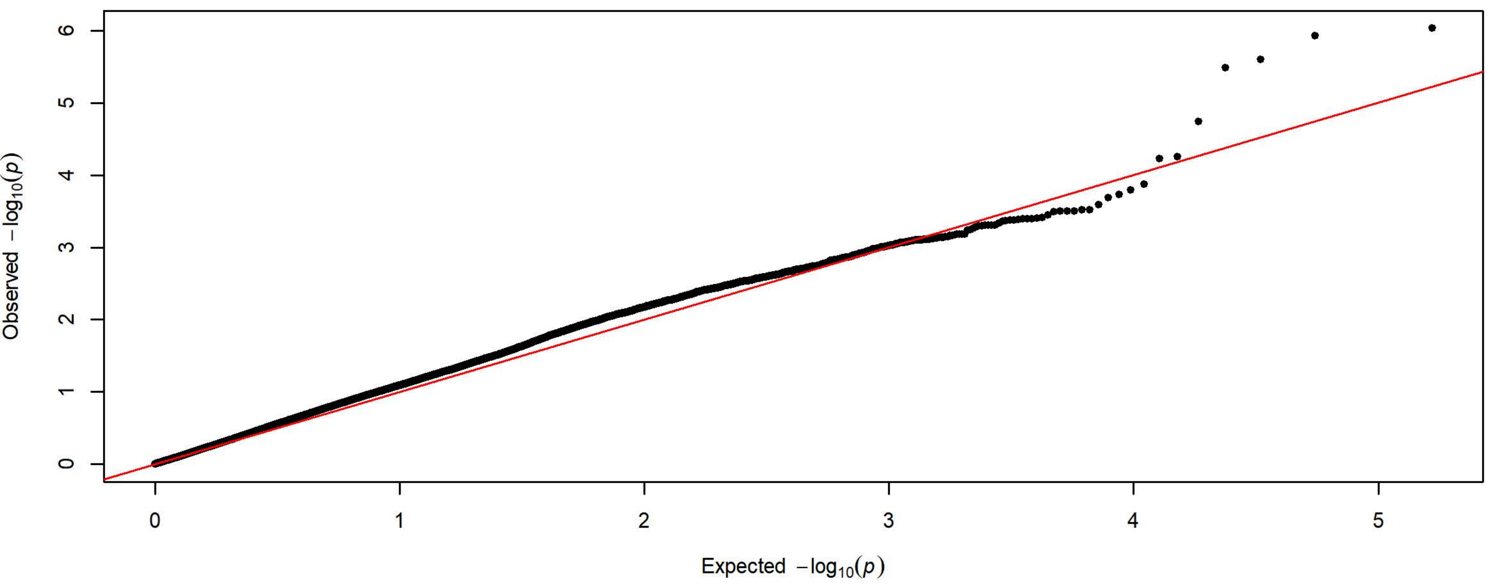

**Q-Q Plot PavCA Index Score Day 3 - Meta-analysis of 7 Subgroups - 64k SNPs (n=3923)**

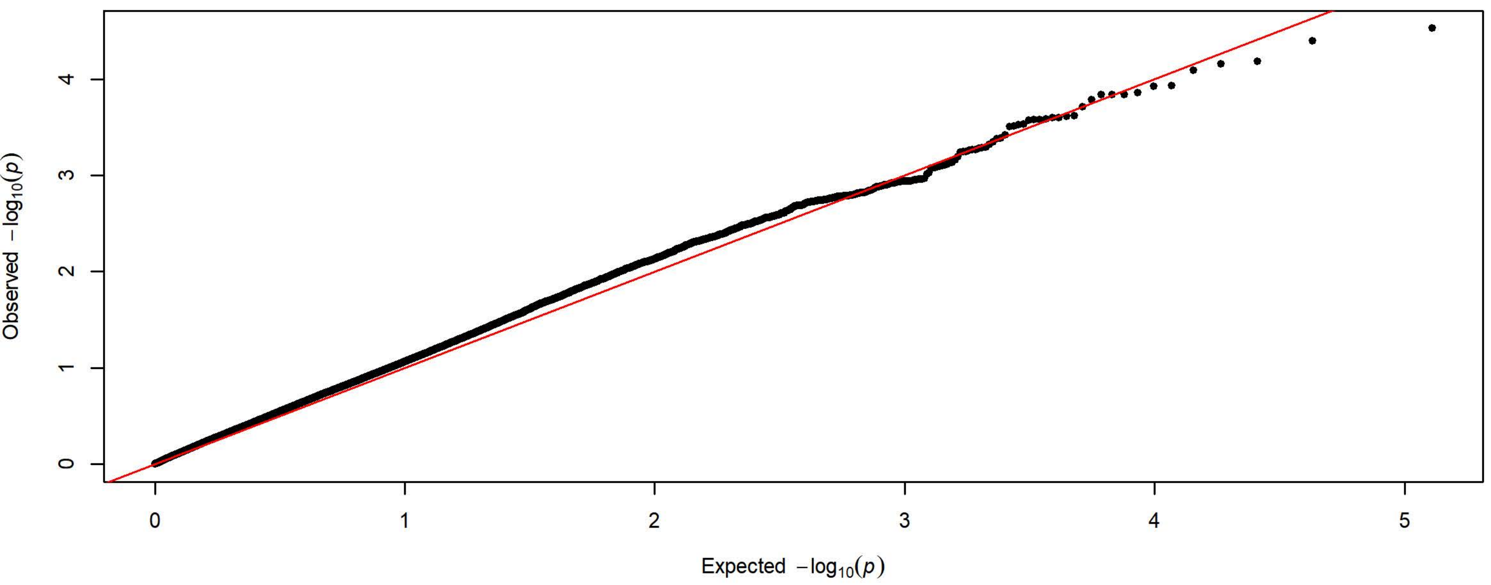


**Q-Q Plot PavCA Index Score Day 3 - Charles River 4 Subgroups - 198k SNPs (n=1722)**

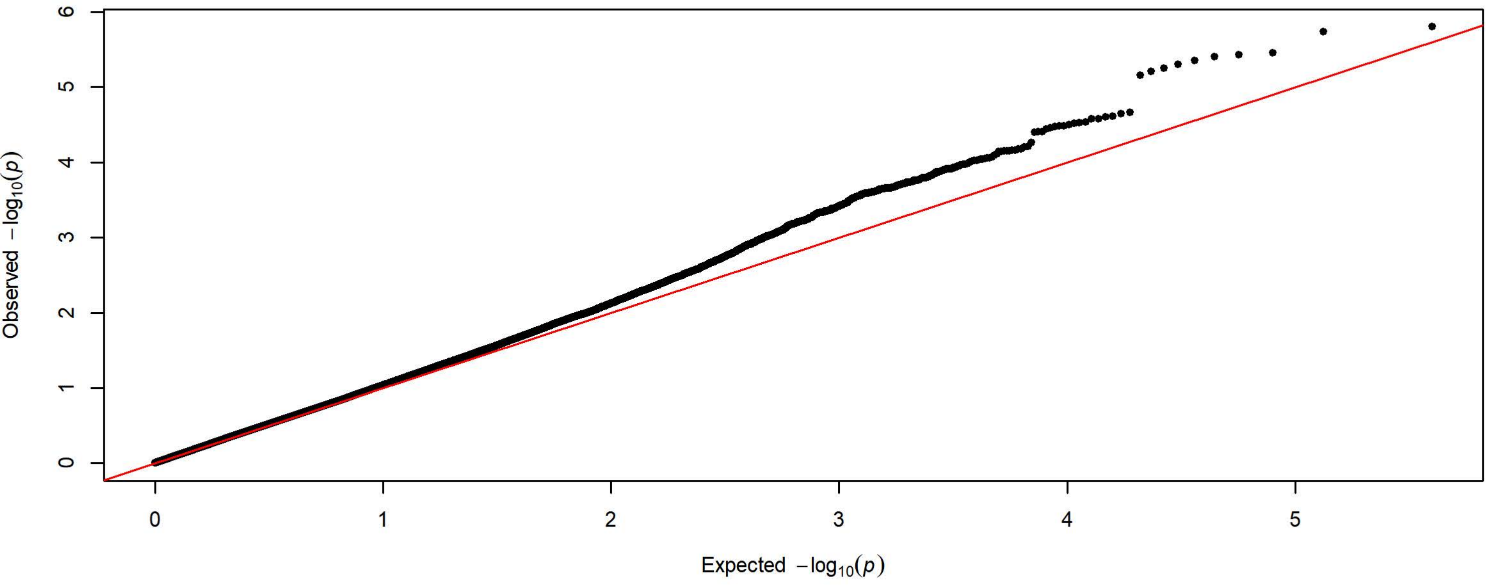


**Q-Q Plot PavCA Index Score Day 3 - Harlan 3 Subgroups - 83k SNPs (n=2201)**

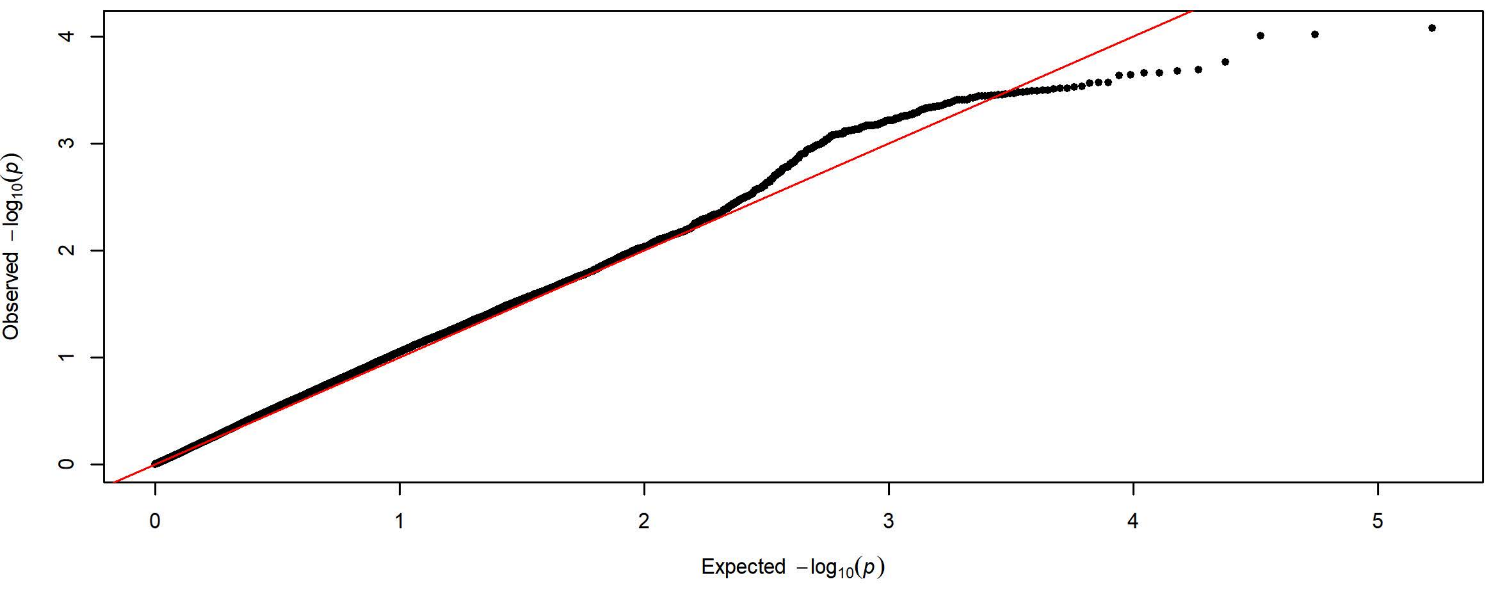

Q-Q Plot PavCA Index Score Day 4 - Meta-analysis of 7 Subgroups - 64k SNPs (n=3935)

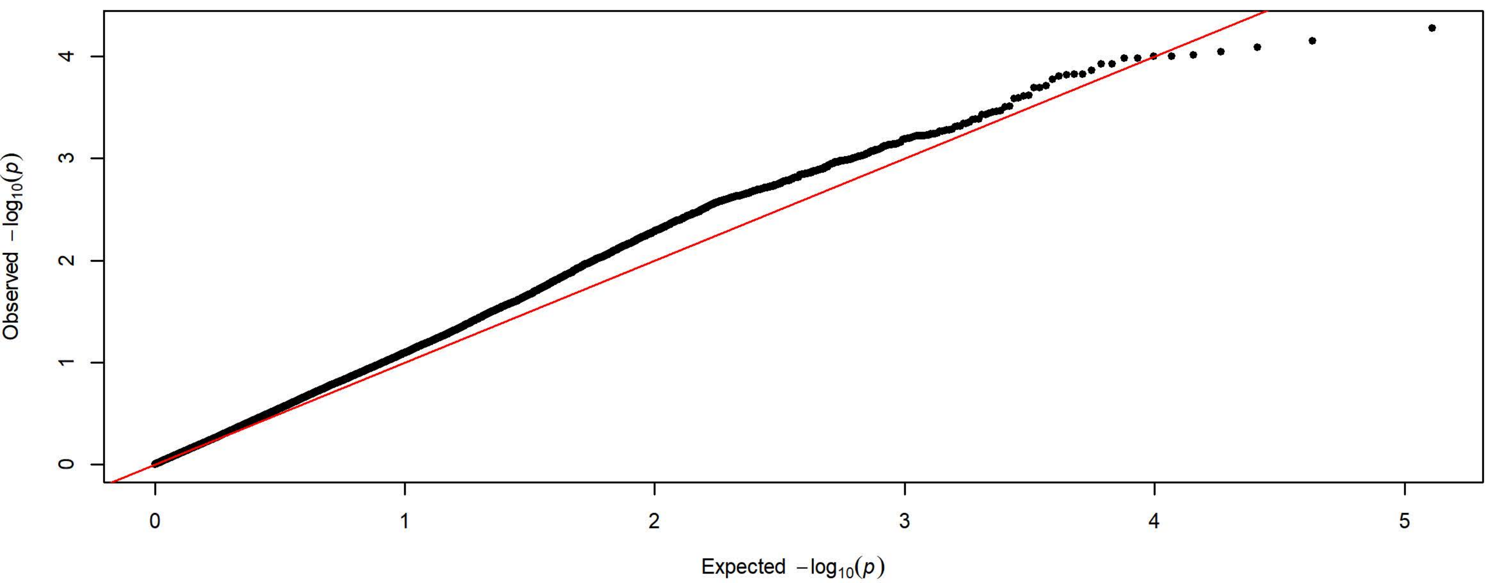


Q-Q Plot PavCA Index Score Day 4 - Charles River 4 Subgroups - 198k SNPs (n=1727)

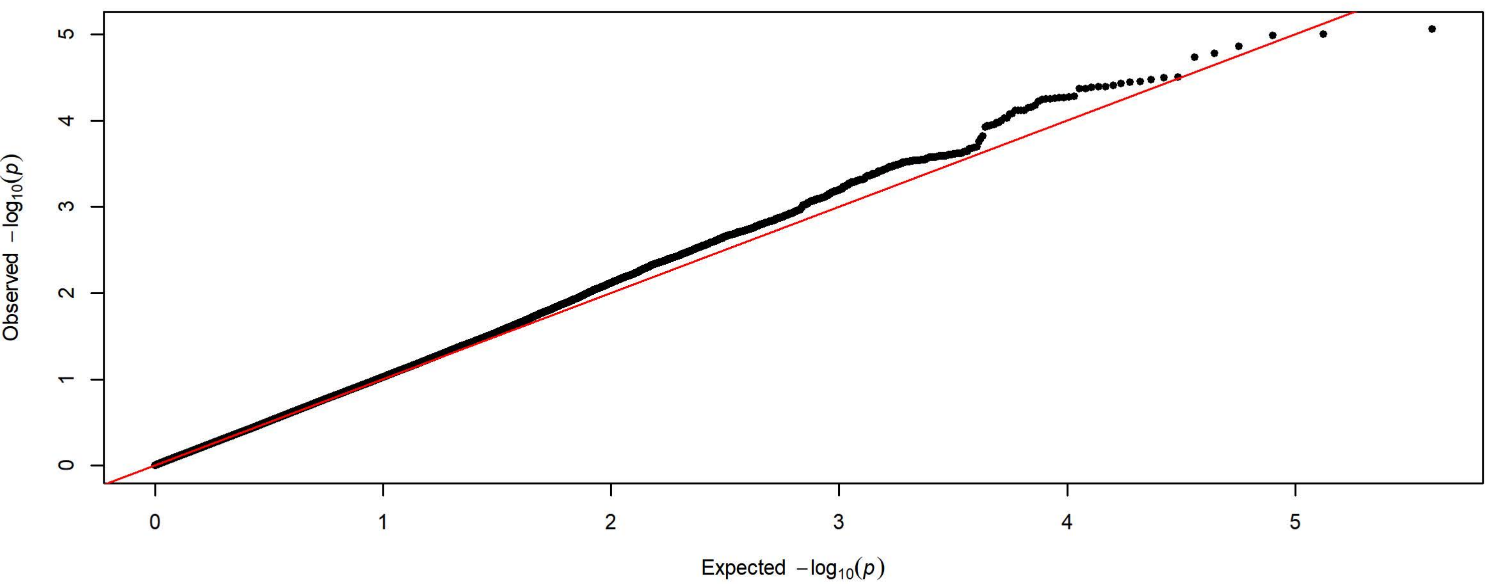


Q-Q Plot PavCA Index Score Day 4 - Harlan 3 Subgroups - 83k SNPs (n=2208)

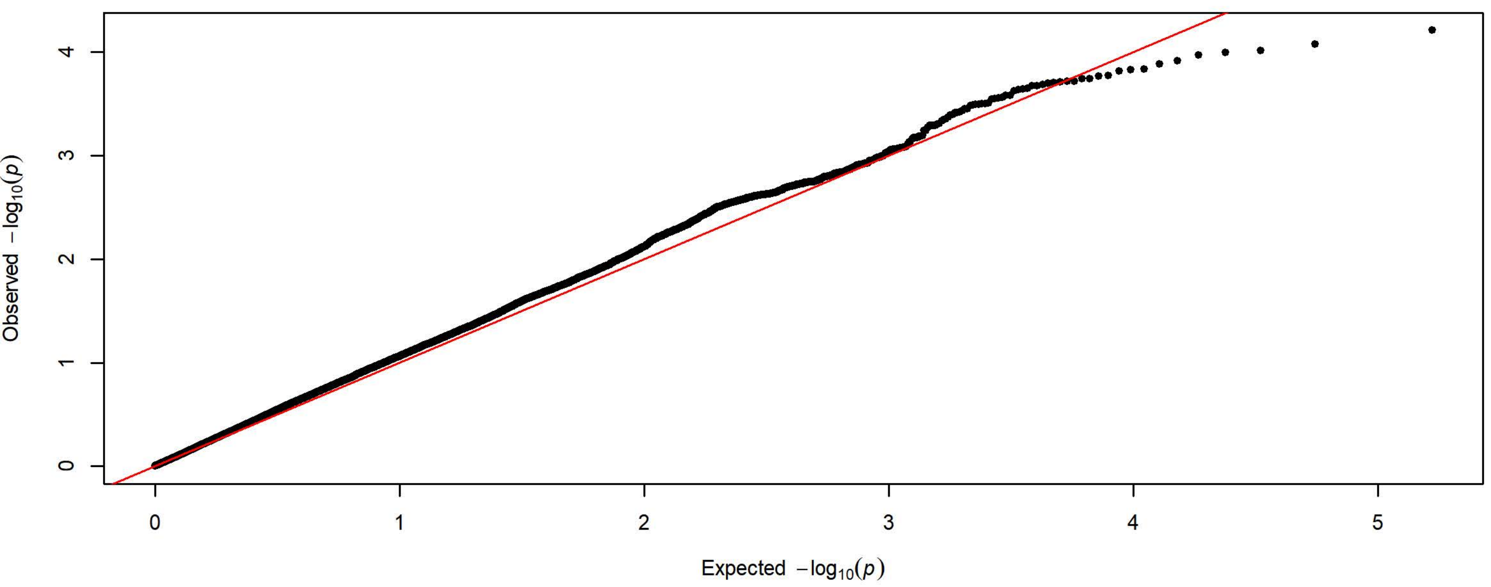

**Q-Q Plot PavCA Index Score Day 5 - Meta-analysis of 7 Subgroups - 64k SNPs (n=3933)**

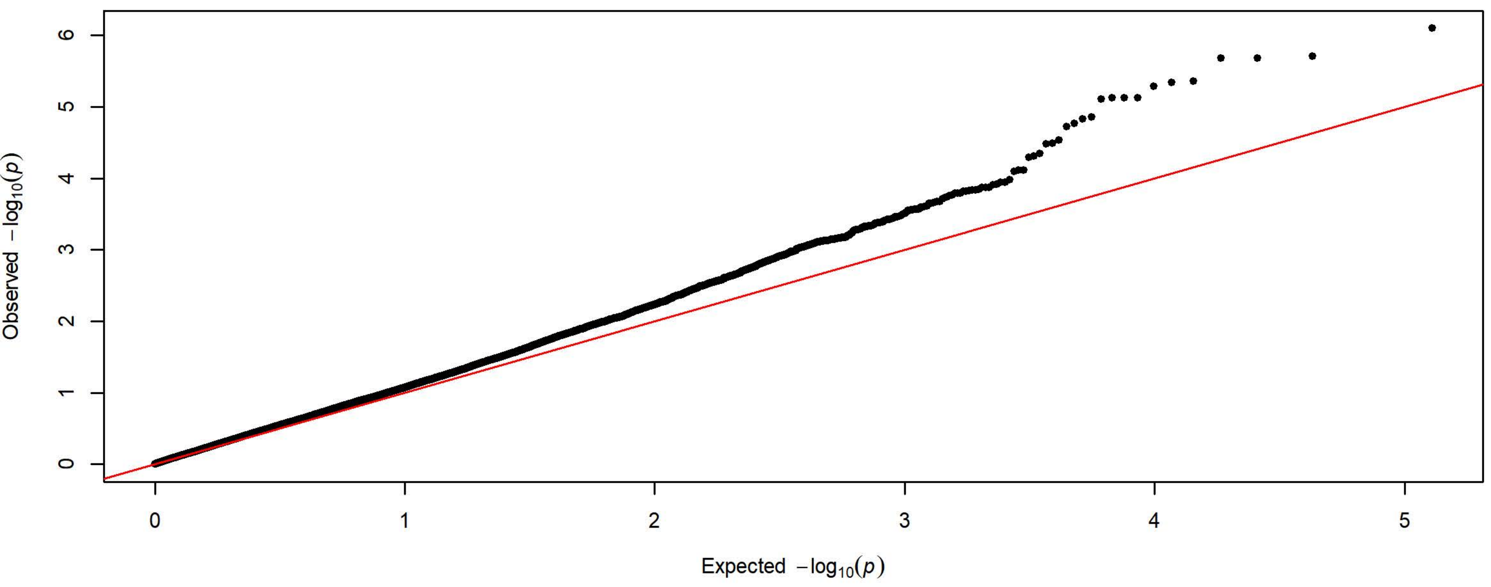


**Q-Q Plot PavCA Index Score Day 5 - Charles River 4 Subgroups - 198k SNPs (n=1726)**

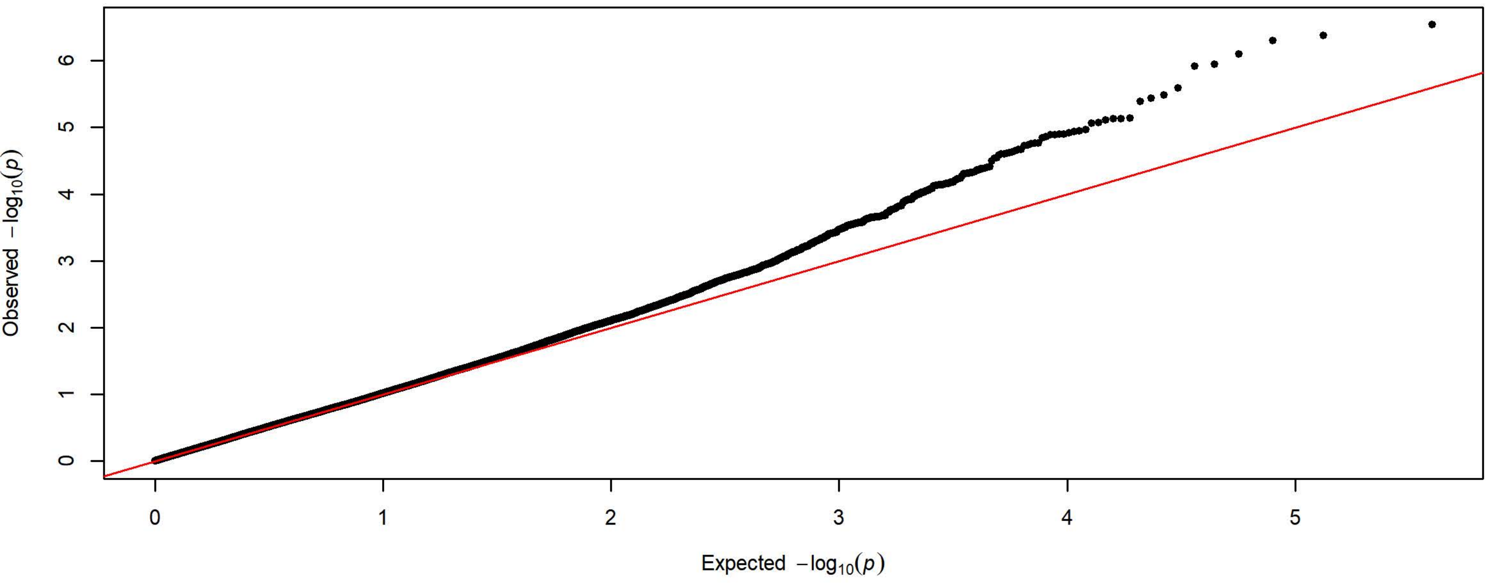


**Q-Q Plot PavCA Index Score Day 5 - Harlan 3 Subgroups - 83k SNPs (n=2207)**

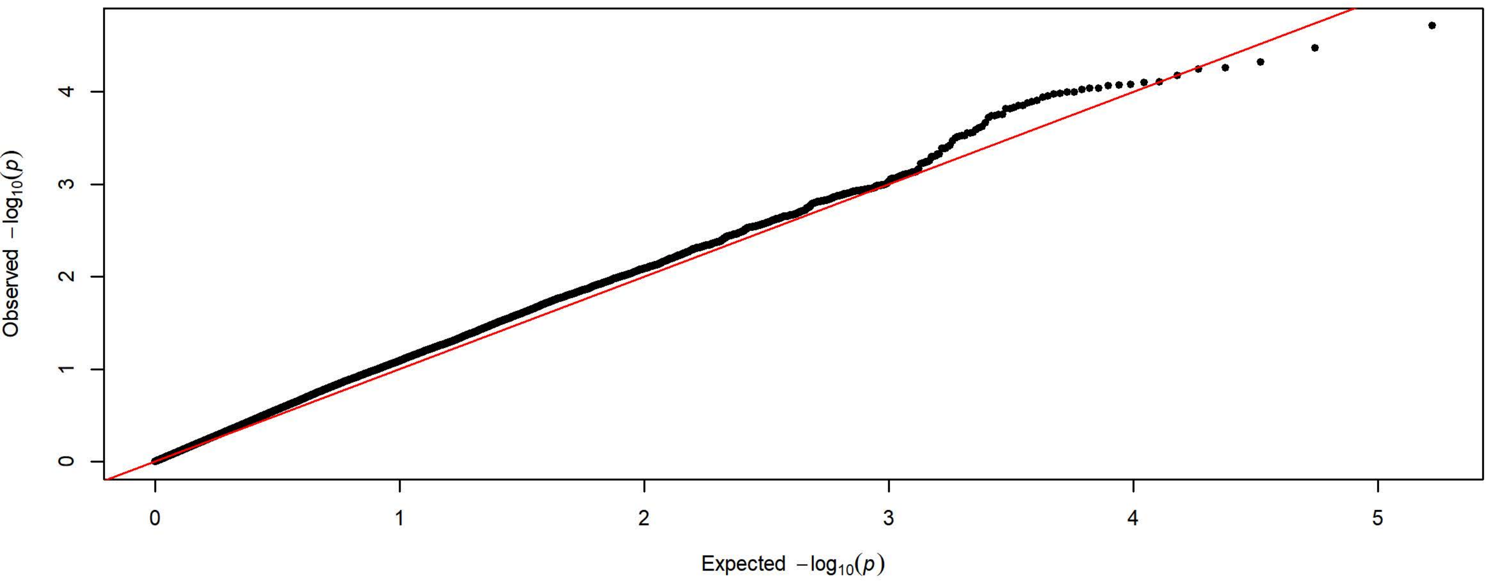

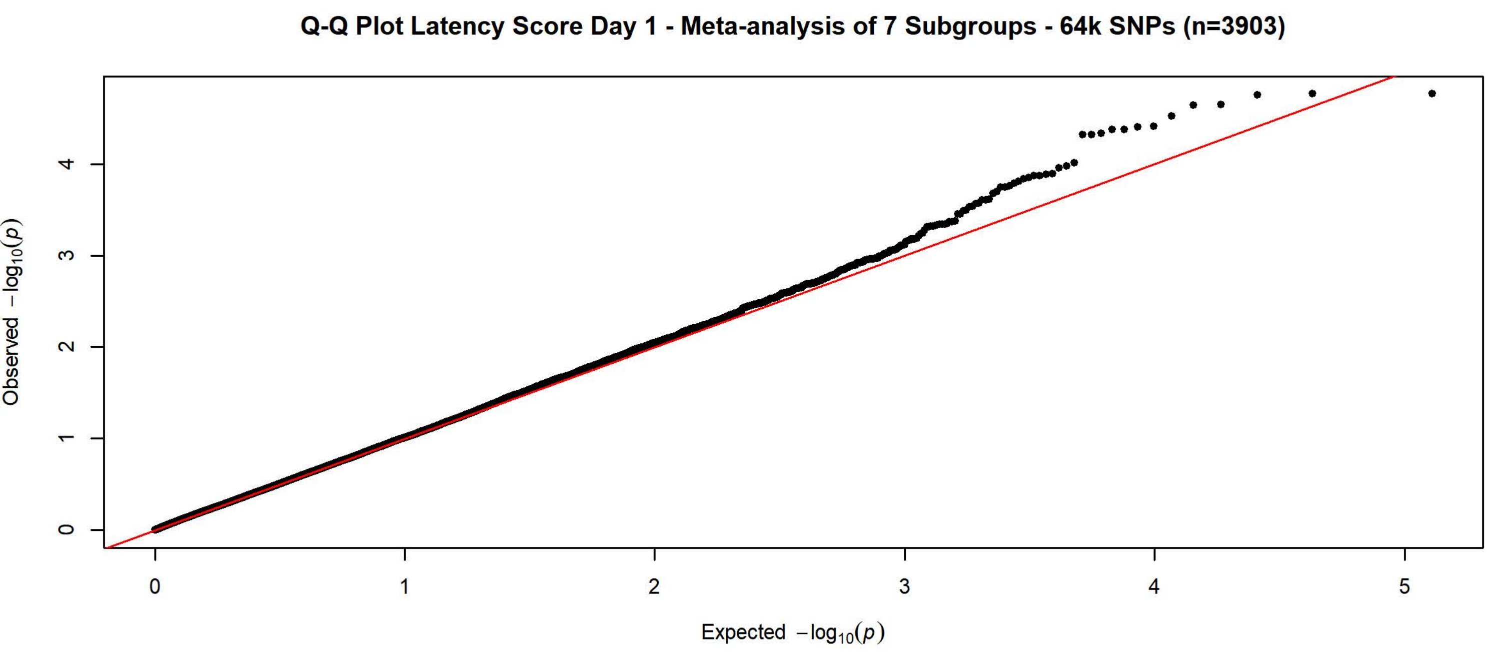

Q-Q Plot Latency Score Day 1 - Meta-analysis of 7 Subgroups - 64k SNPs (n=3903)
Observed $-\log_{10}(p)$
Expected $-\log_{10}(p)$


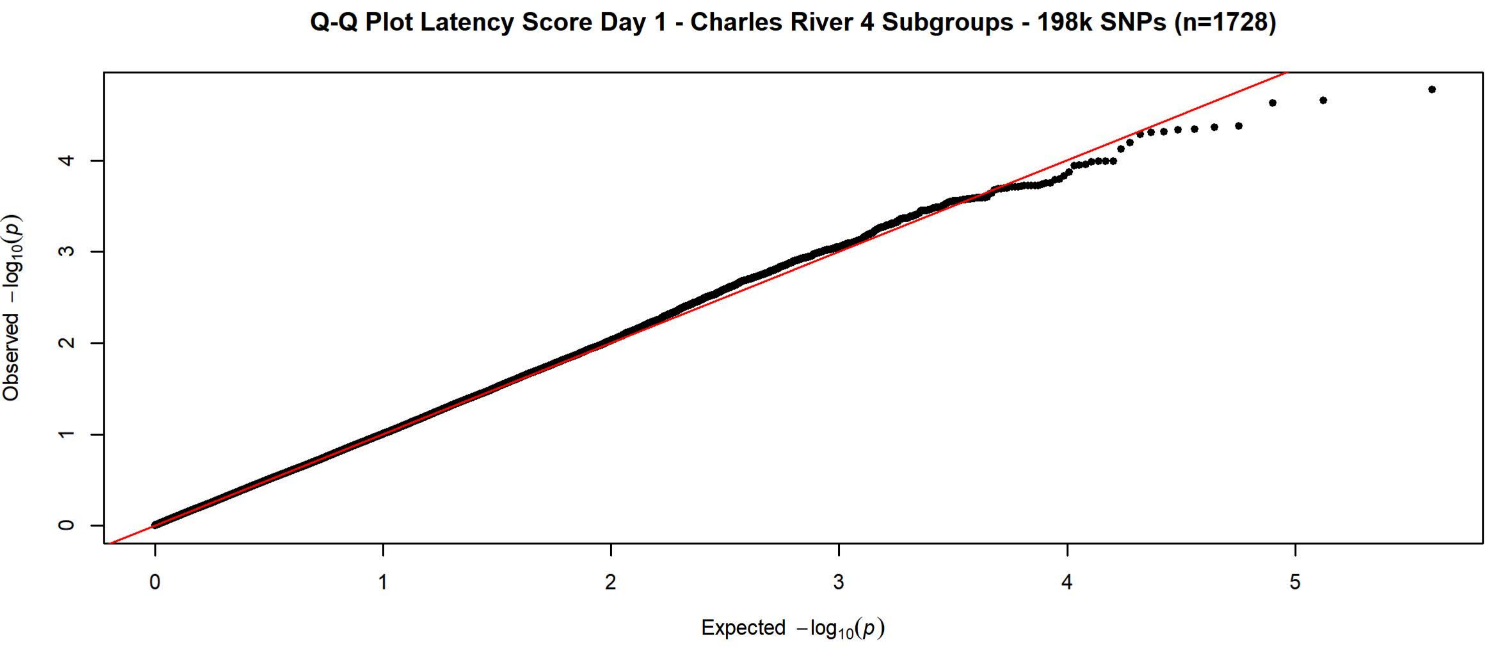

Q-Q Plot Latency Score Day 1 - Charles River 4 Subgroups - 198k SNPs (n=1728)
Observed $-\log_{10}(p)$
Expected $-\log_{10}(p)$


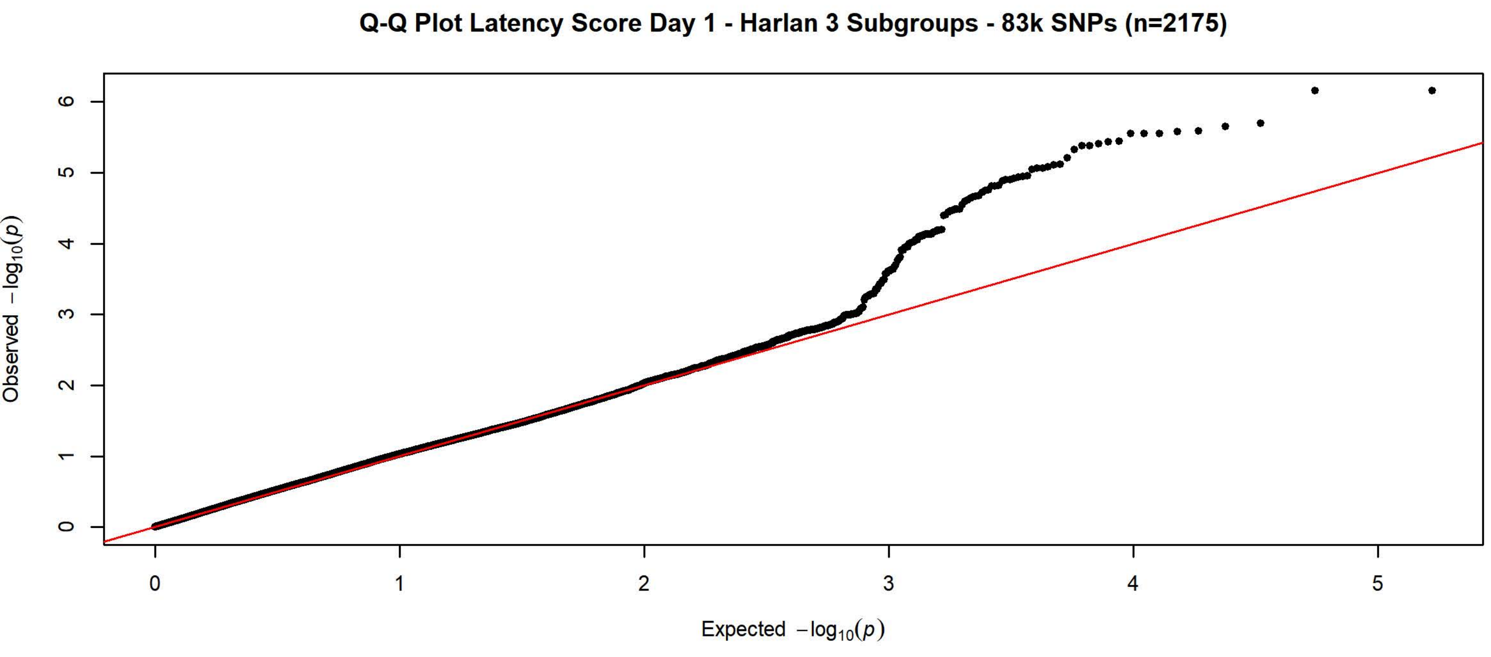

Q-Q Plot Latency Score Day 1 - Harlan 3 Subgroups - 83k SNPs (n=2175)
Observed $-\log_{10}(p)$
Expected $-\log_{10}(p)$

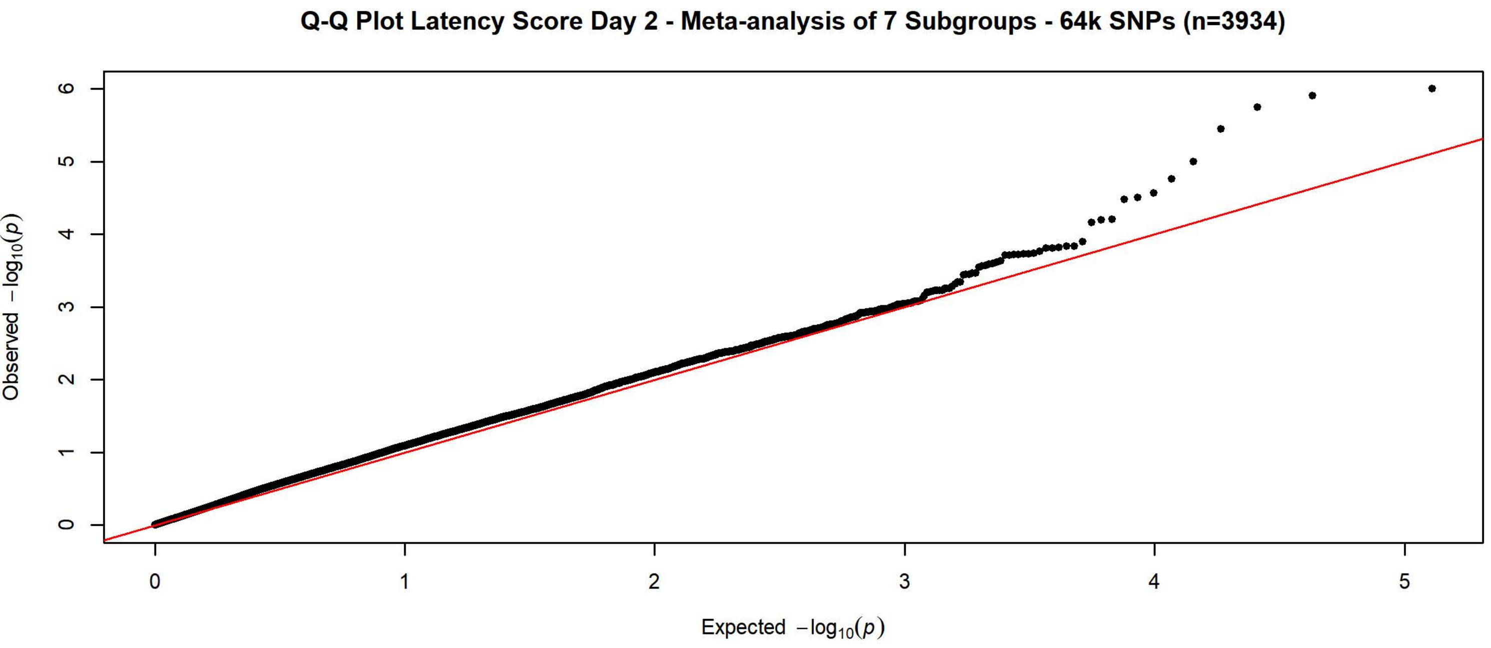
Q-Q Plot Latency Score Day 2 - Meta-analysis of 7 Subgroups - 64k SNPs (n=3934)
Observed $-\log_{10}(p)$
Expected $-\log_{10}(p)$

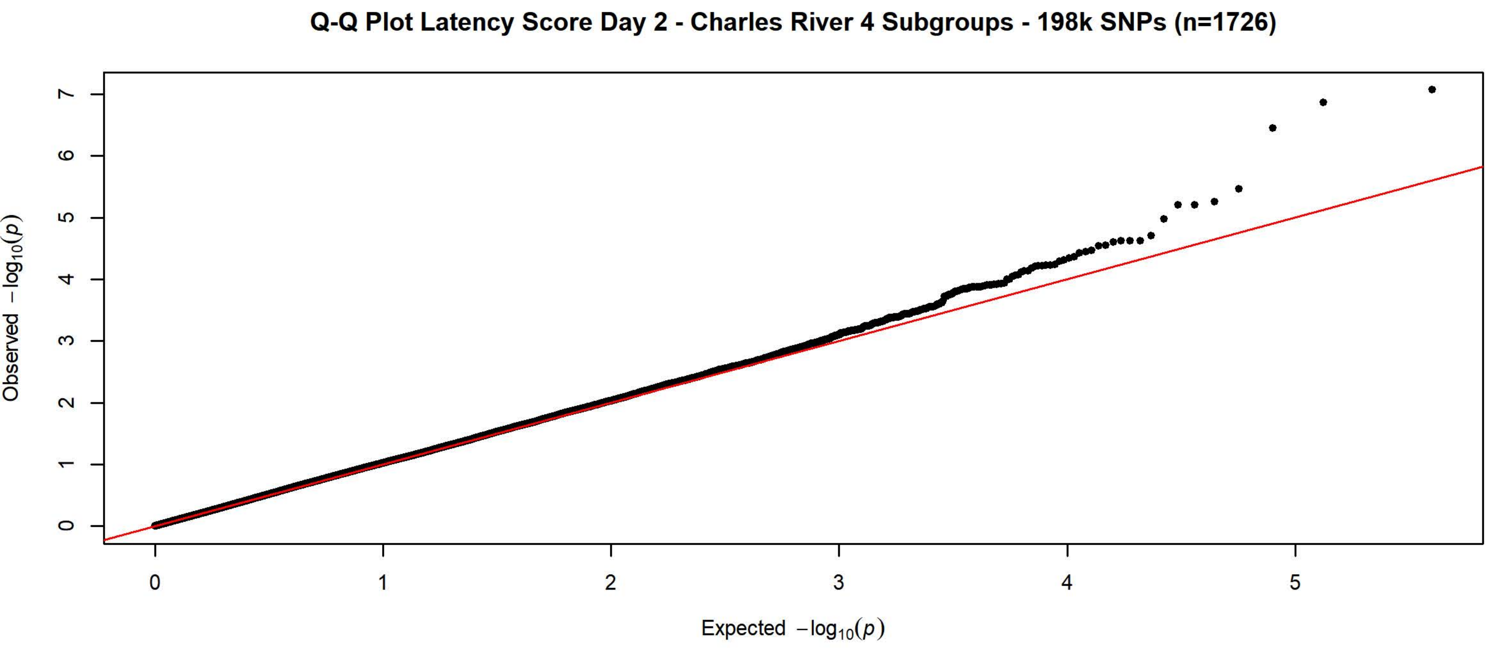
Q-Q Plot Latency Score Day 2 - Charles River 4 Subgroups - 198k SNPs (n=1726)
Observed $-\log_{10}(p)$
Expected $-\log_{10}(p)$

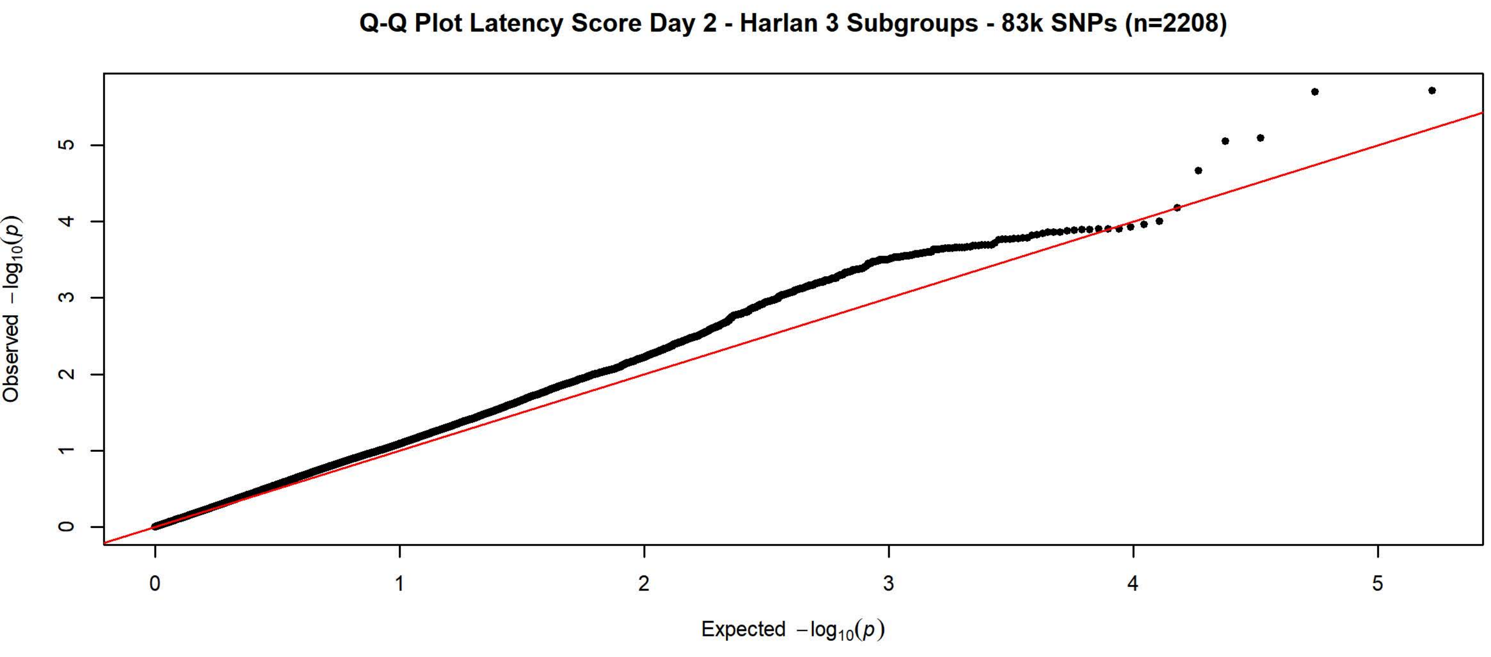
Q-Q Plot Latency Score Day 2 - Harlan 3 Subgroups - 83k SNPs (n=2208)
Observed $-\log_{10}(p)$
Expected $-\log_{10}(p)$

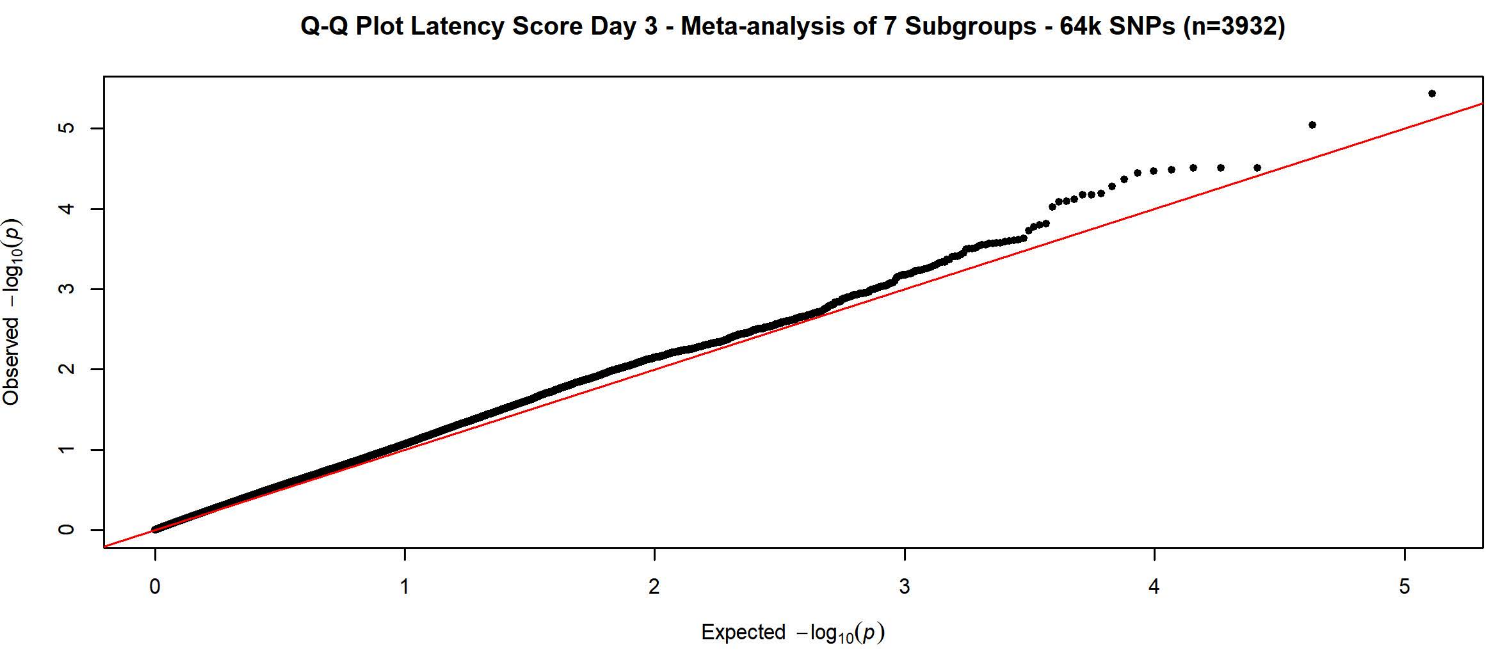
Q-Q Plot Latency Score Day 3 - Meta-analysis of 7 Subgroups - 64k SNPs (n=3932)
Observed $-\log_{10}(p)$
Expected $-\log_{10}(p)$

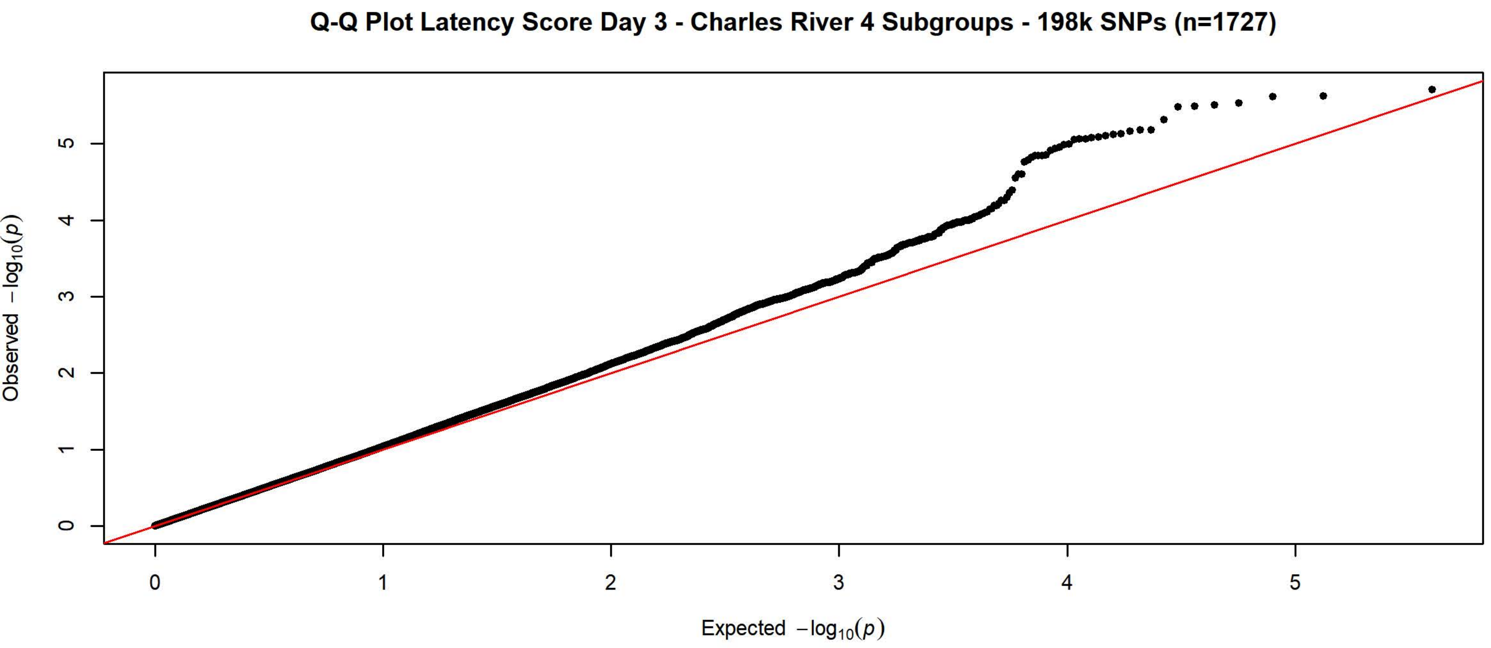
Q-Q Plot Latency Score Day 3 - Charles River 4 Subgroups - 198k SNPs (n=1727)
Observed $-\log_{10}(p)$
Expected $-\log_{10}(p)$

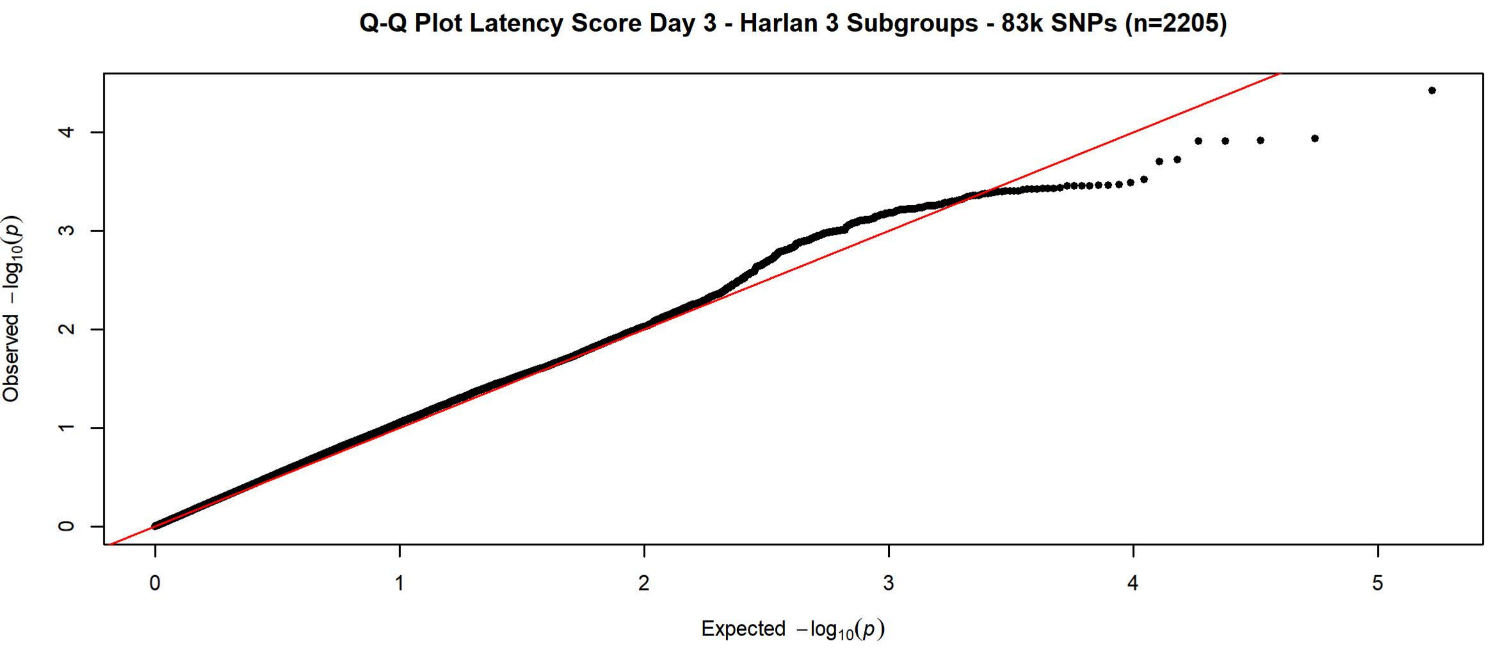
Q-Q Plot Latency Score Day 3 - Harlan 3 Subgroups - 83k SNPs (n=2205)
Observed $-\log_{10}(p)$
Expected $-\log_{10}(p)$

**Q-Q Plot Latency Score Day 4 - Meta-analysis of 7 Subgroups - 64k SNPs (n=3936)**

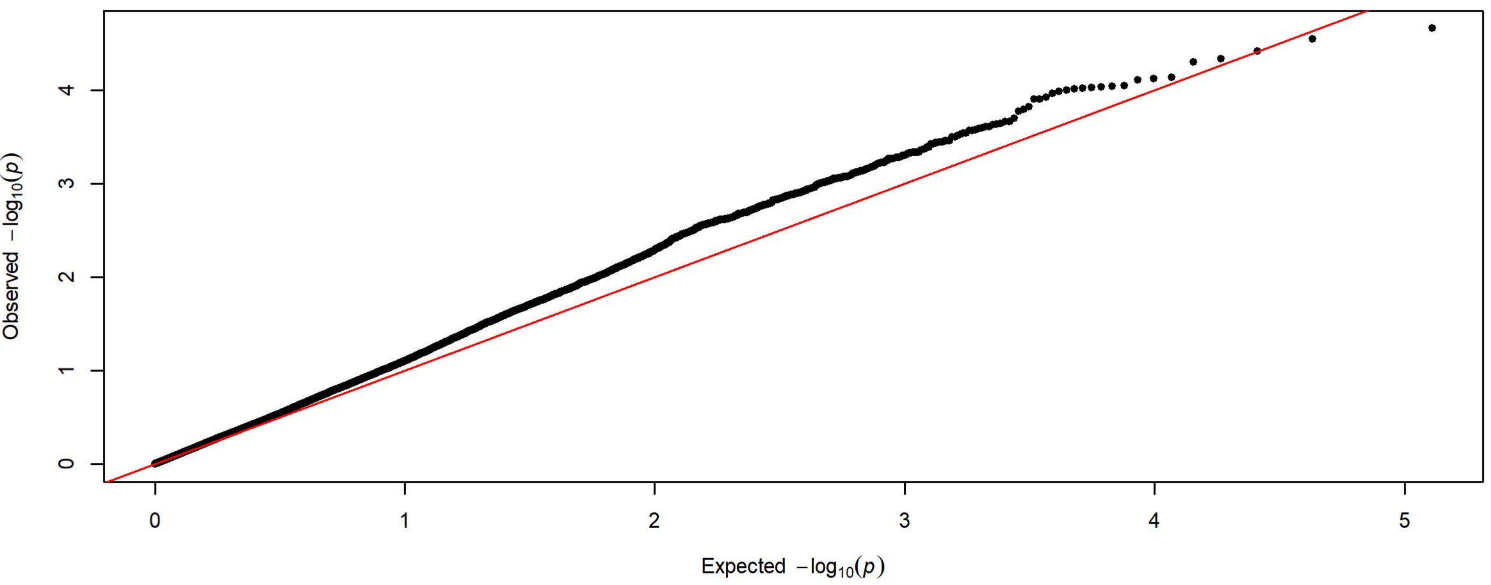


**Q-Q Plot Latency Score Day 4 - Charles River 4 Subgroups - 198k SNPs (n=1728)**

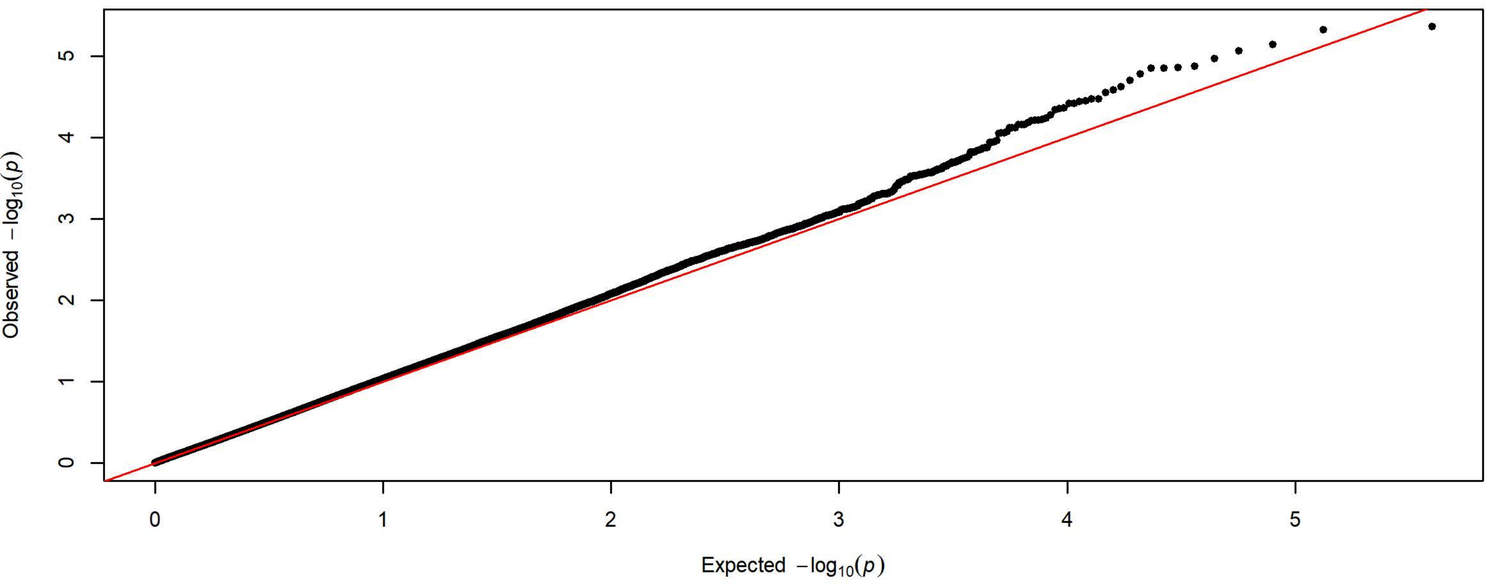


**Q-Q Plot Latency Score Day 4 - Harlan 3 Subgroups - 83k SNPs (n=2208)**

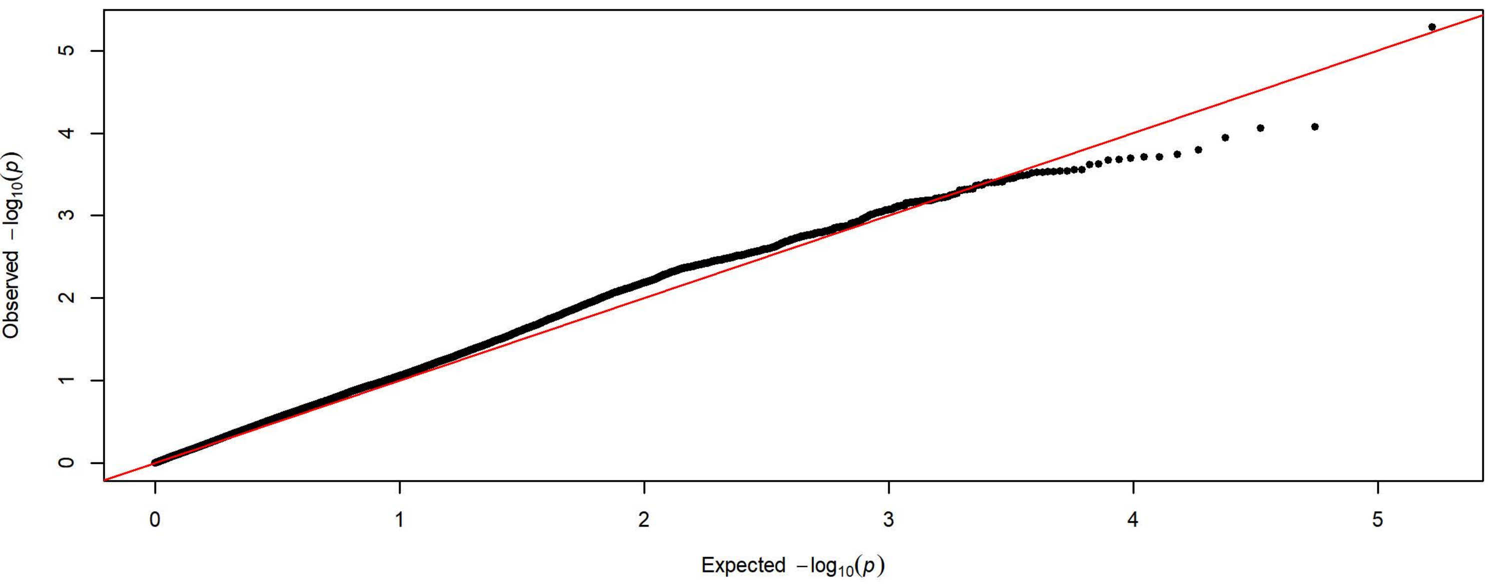

**Q-Q Plot Latency Score Day 5 - Meta-analysis of 7 Subgroups - 64k SNPs (n=3936)**

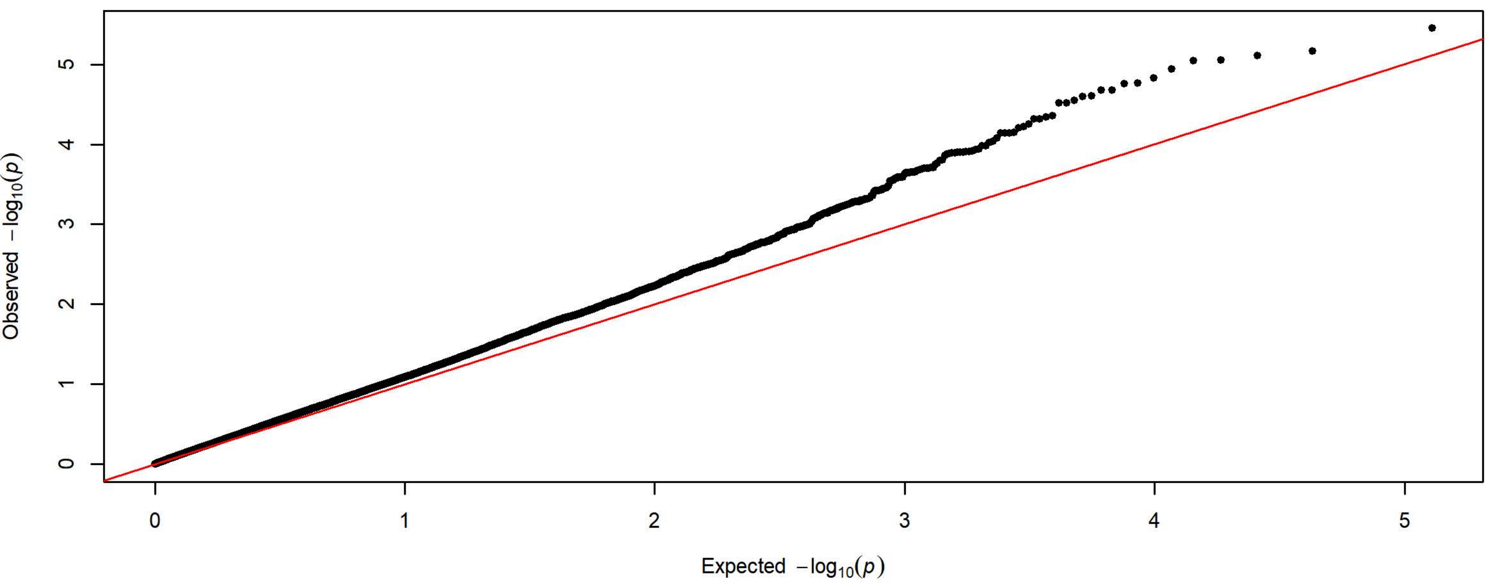


**Q-Q Plot Latency Score Day 5 - Charles River 4 Subgroups - 198k SNPs (n=1728)**

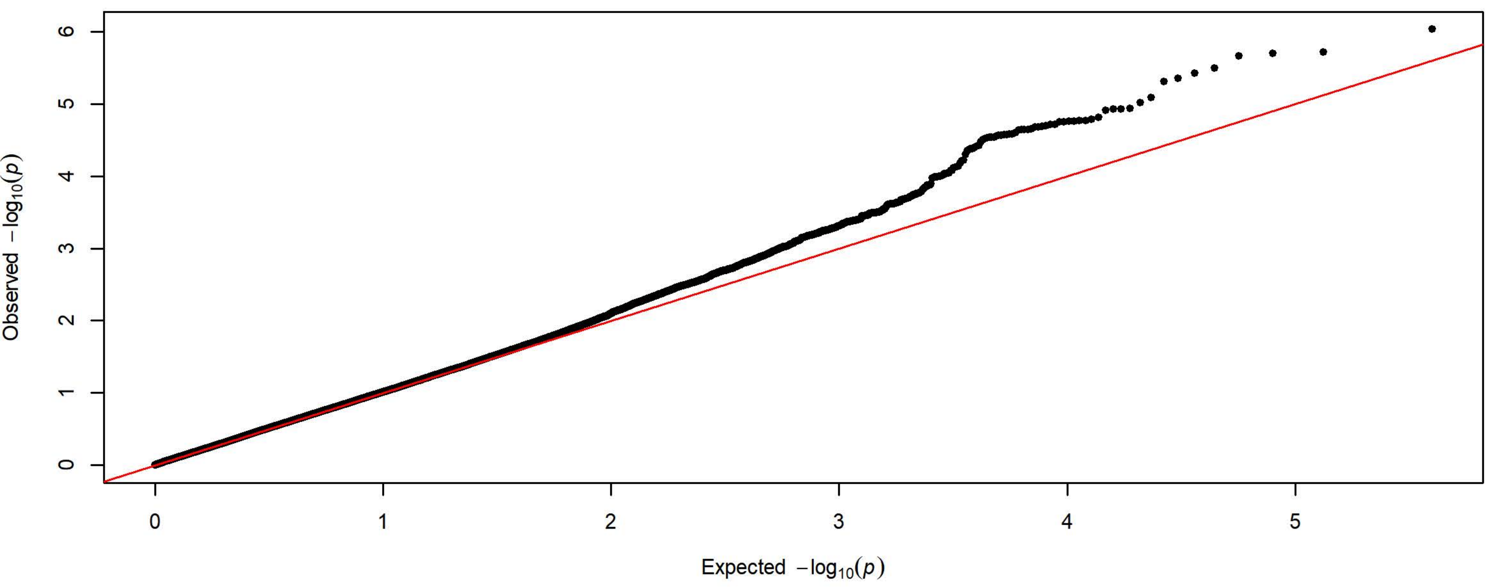


**Q-Q Plot Latency Score Day 5 - Harlan 3 Subgroups - 83k SNPs (n=2208)**

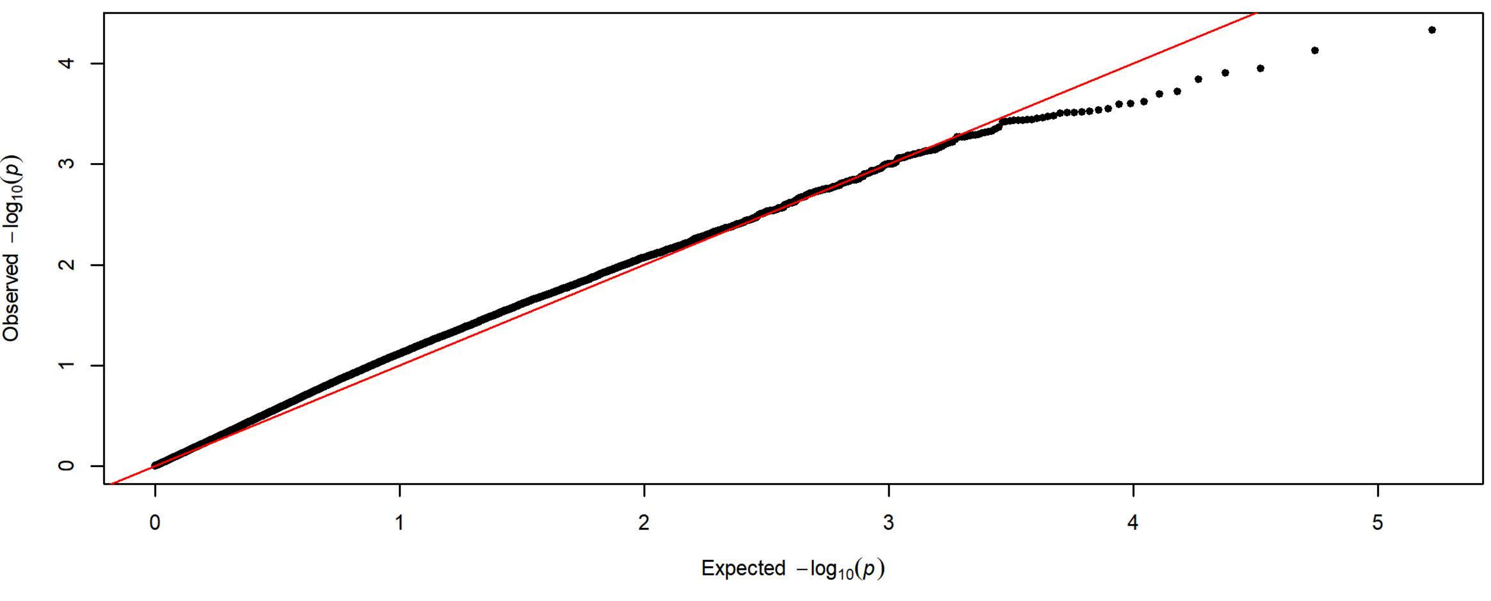

**Q-Q Plot Lever Presses Day 1 - Meta-analysis of 7 Subgroups - 64k SNPs (n=3933)**

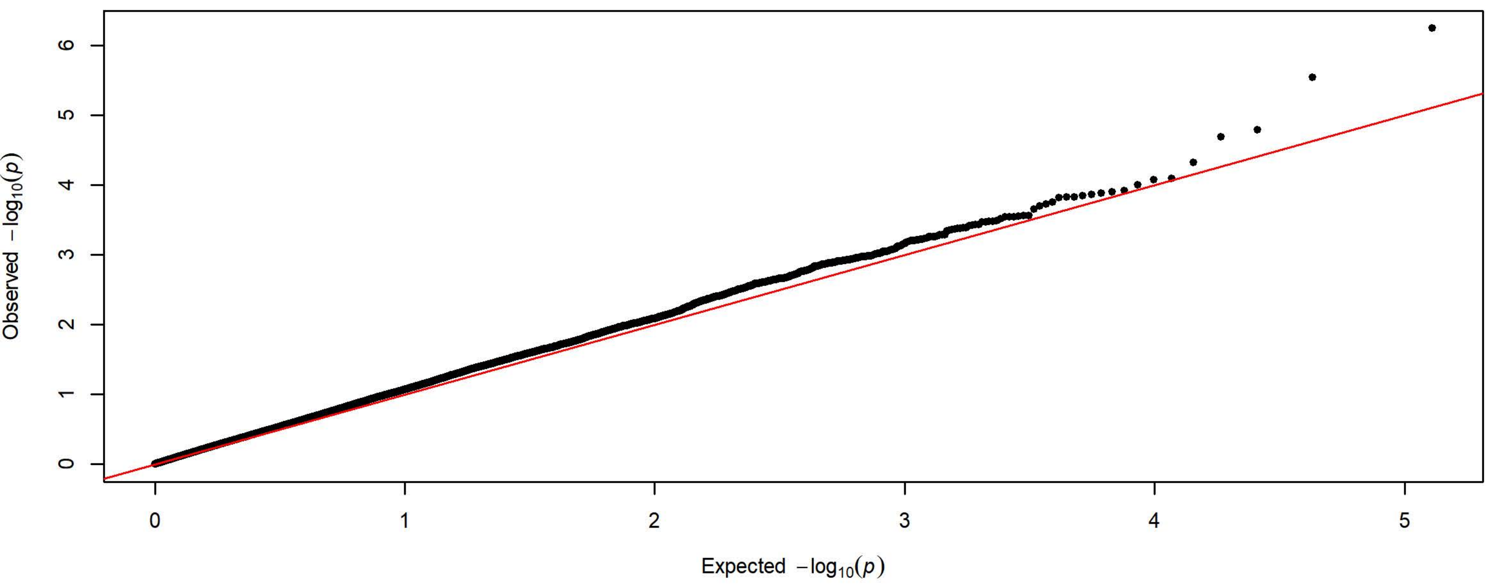


**Q-Q Plot Lever Presses Day 1 - Charles River 4 Subgroups - 198k SNPs (n=1727)**

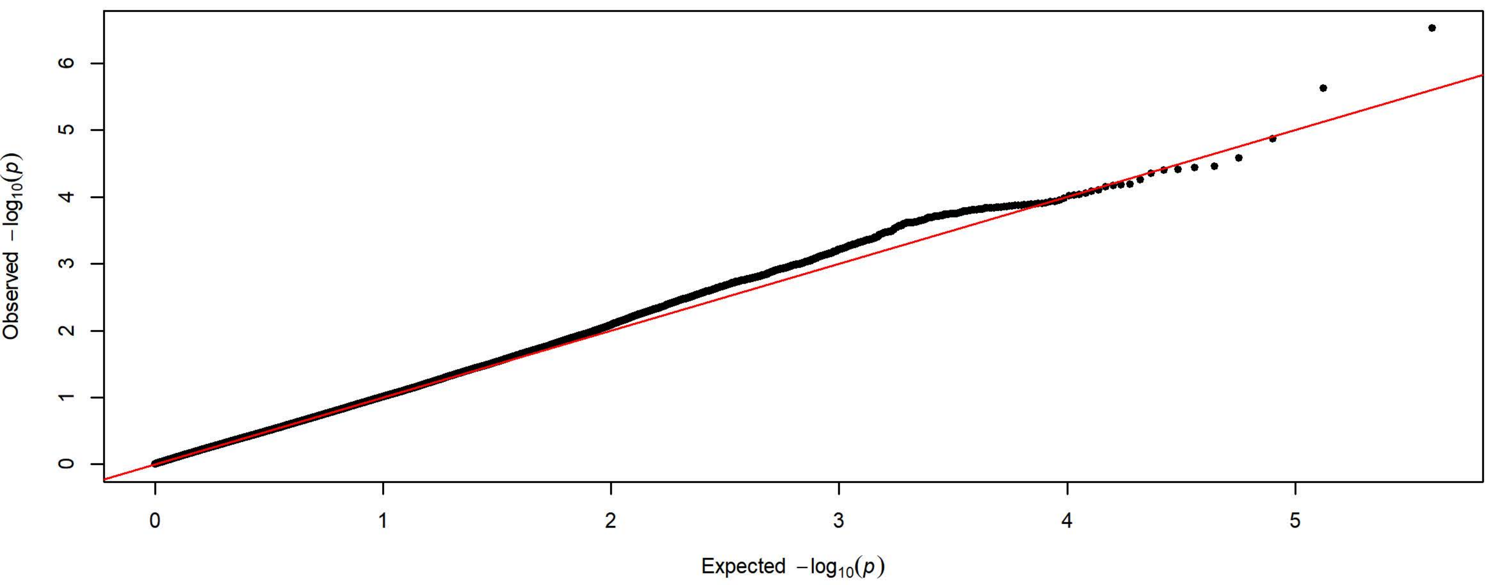


**Q-Q Plot Lever Presses Day 1 - Harlan 3 Subgroups - 83k SNPs (n=2206)**

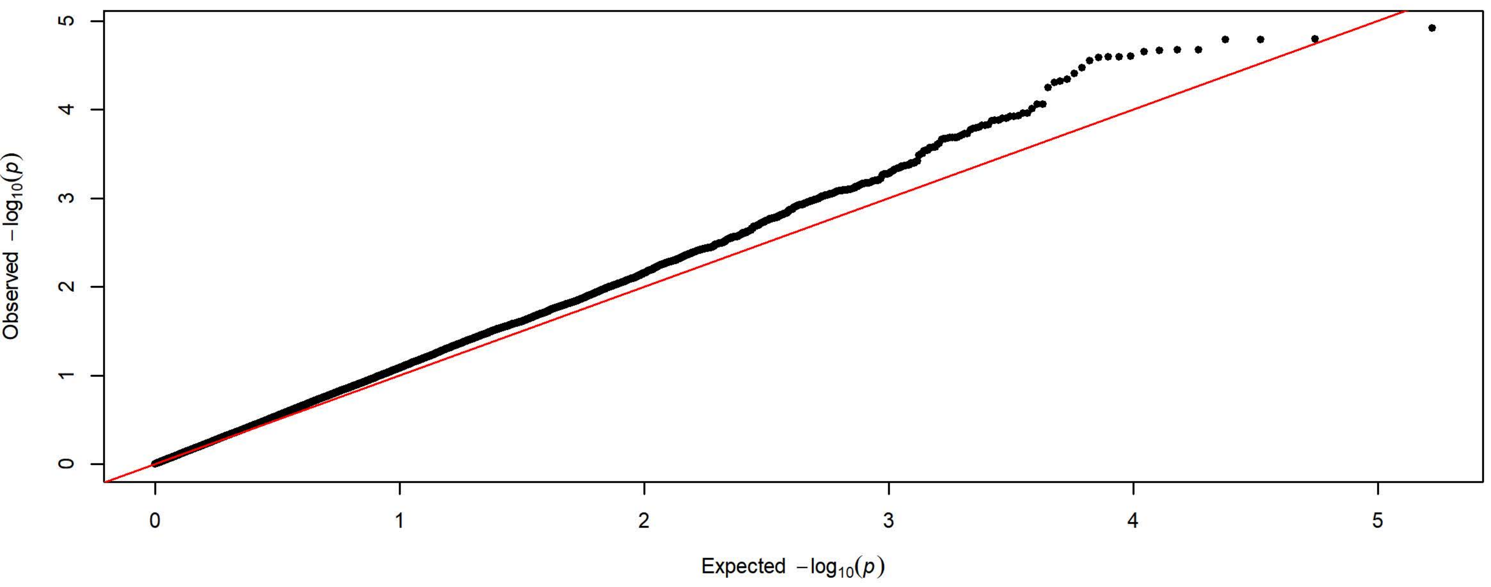

**Q-Q Plot Lever Presses Day 2 - Meta-analysis of 7 Subgroups - 64k SNPs (n=3931)**

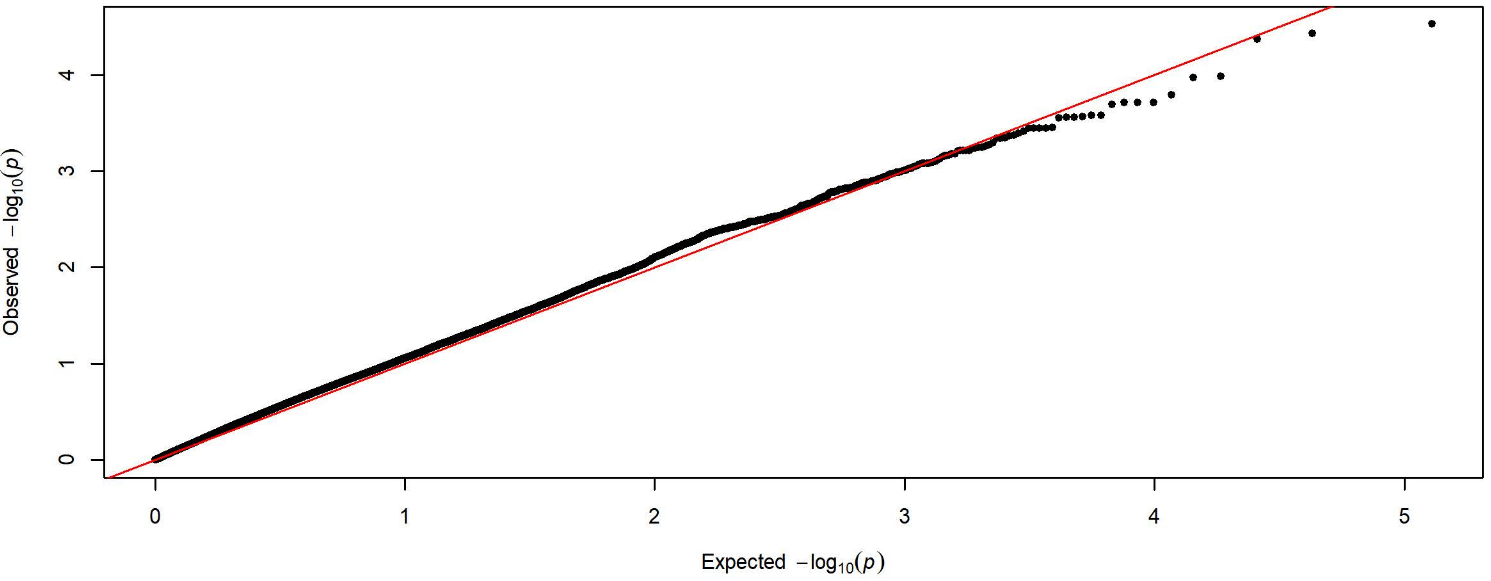


**Q-Q Plot Lever Presses Day 2 - Charles River 4 Subgroups - 198k SNPs (n=1726)**

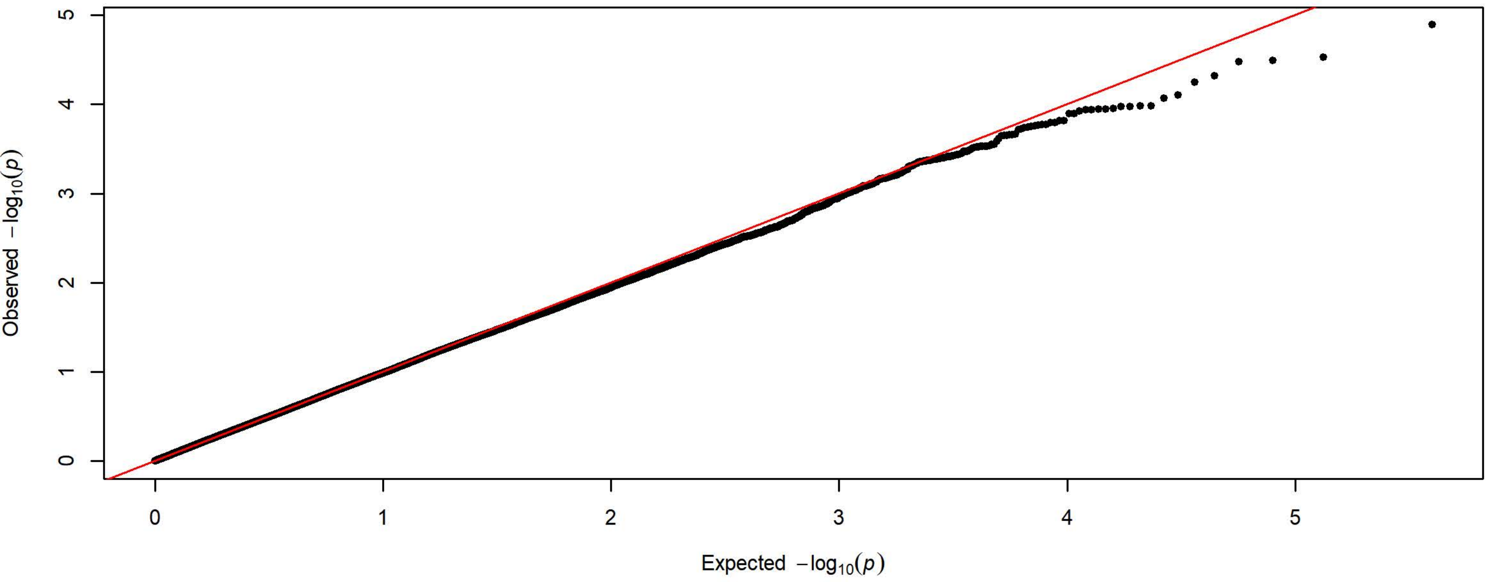


**Q-Q Plot Lever Presses Day 2 - Harlan 3 Subgroups - 83k SNPs (n=2205)**

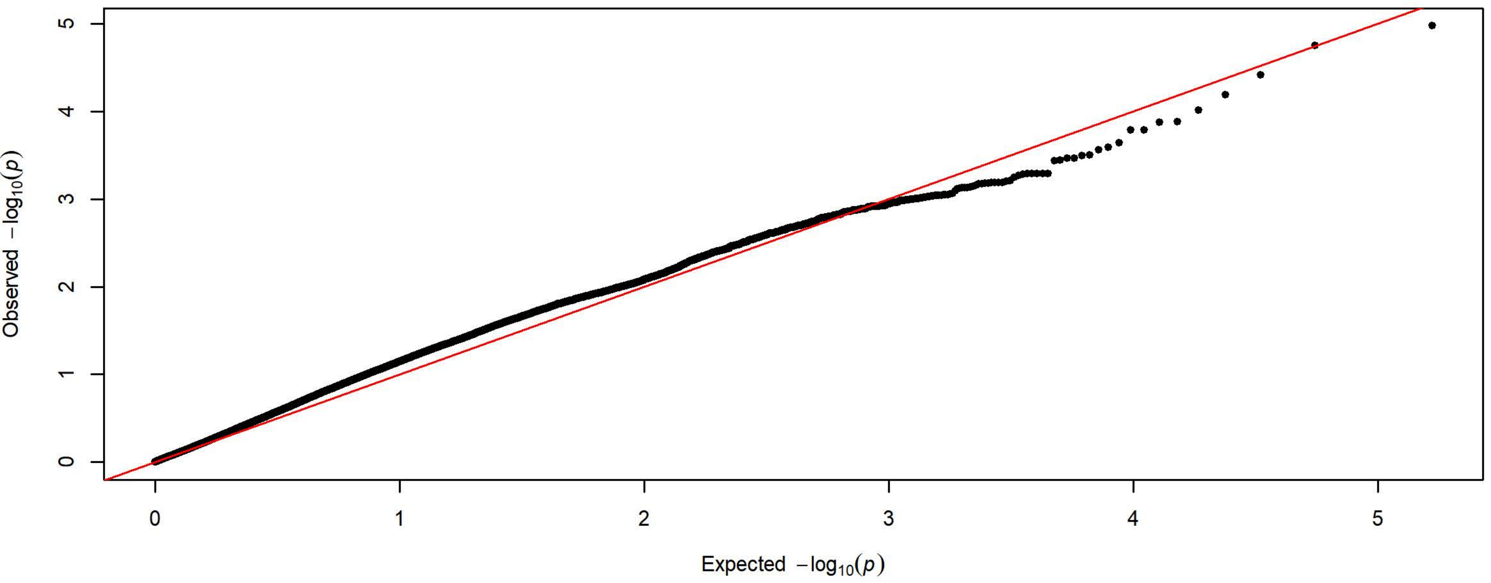

**Q-Q Plot Lever Presses Day 3 - Meta-analysis of 7 Subgroups - 64k SNPs (n=3931)**

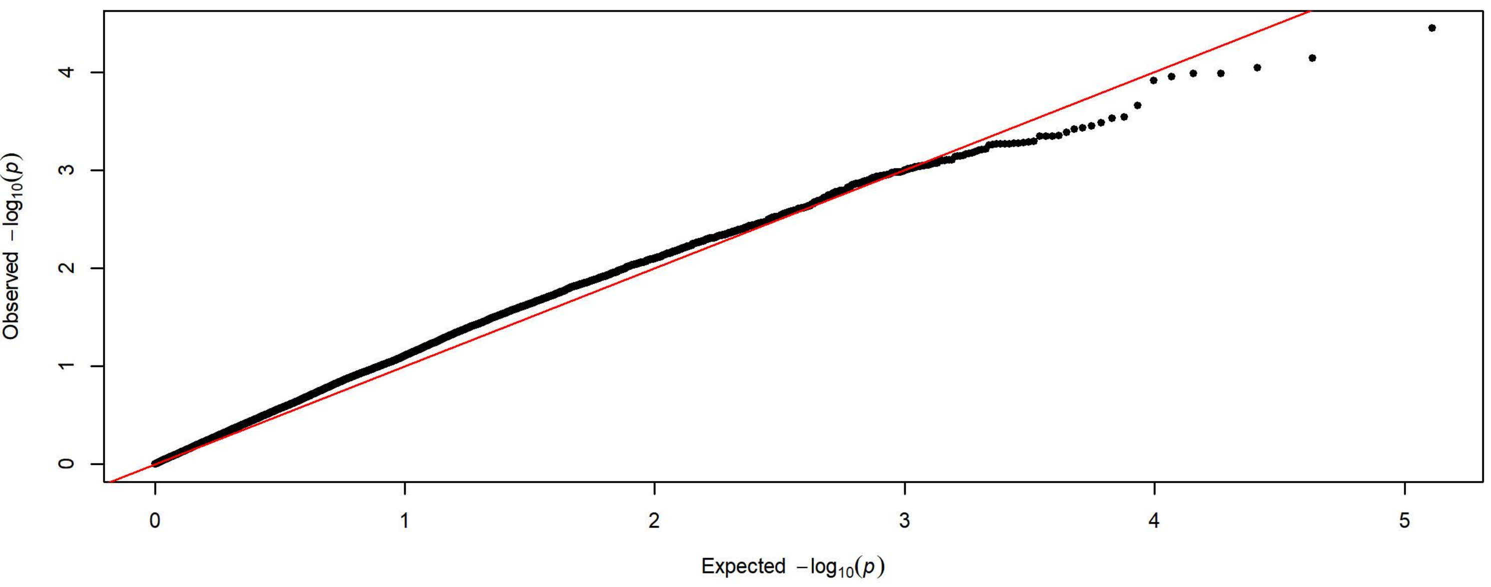


**Q-Q Plot Lever Presses Day 3 - Charles River 4 Subgroups - 198k SNPs (n=1727)**

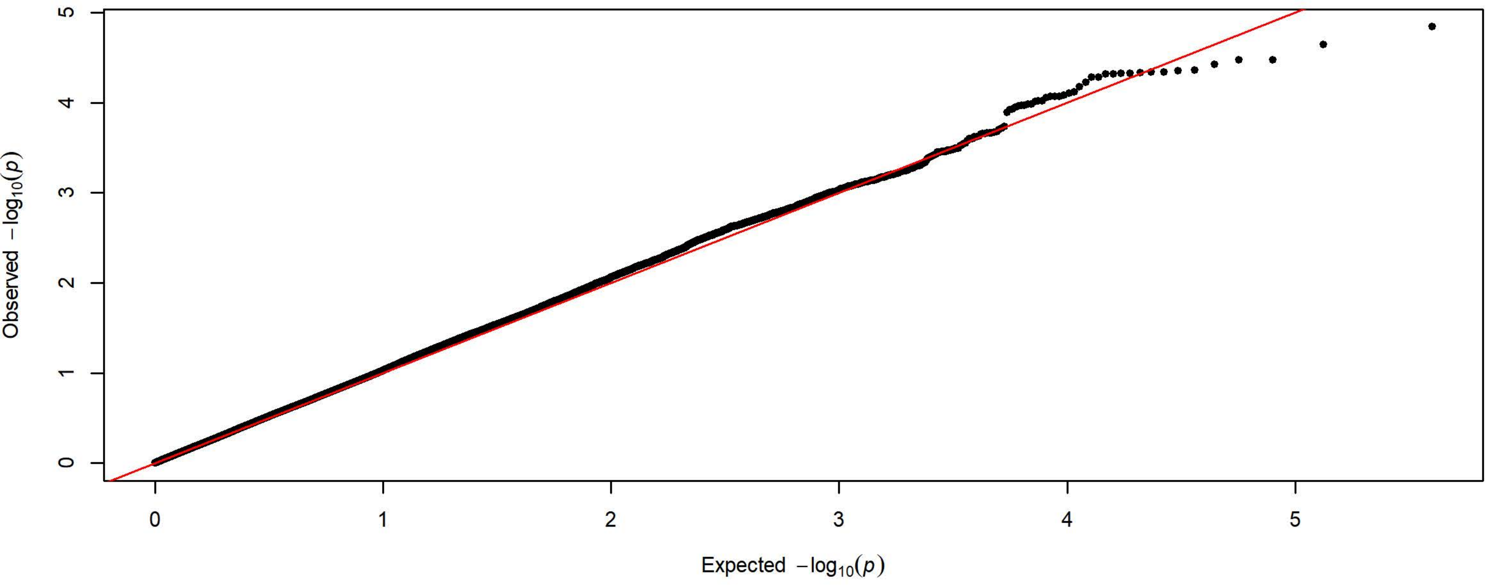


**Q-Q Plot Lever Presses Day 3 - Harlan 3 Subgroups - 83k SNPs (n=2204)**

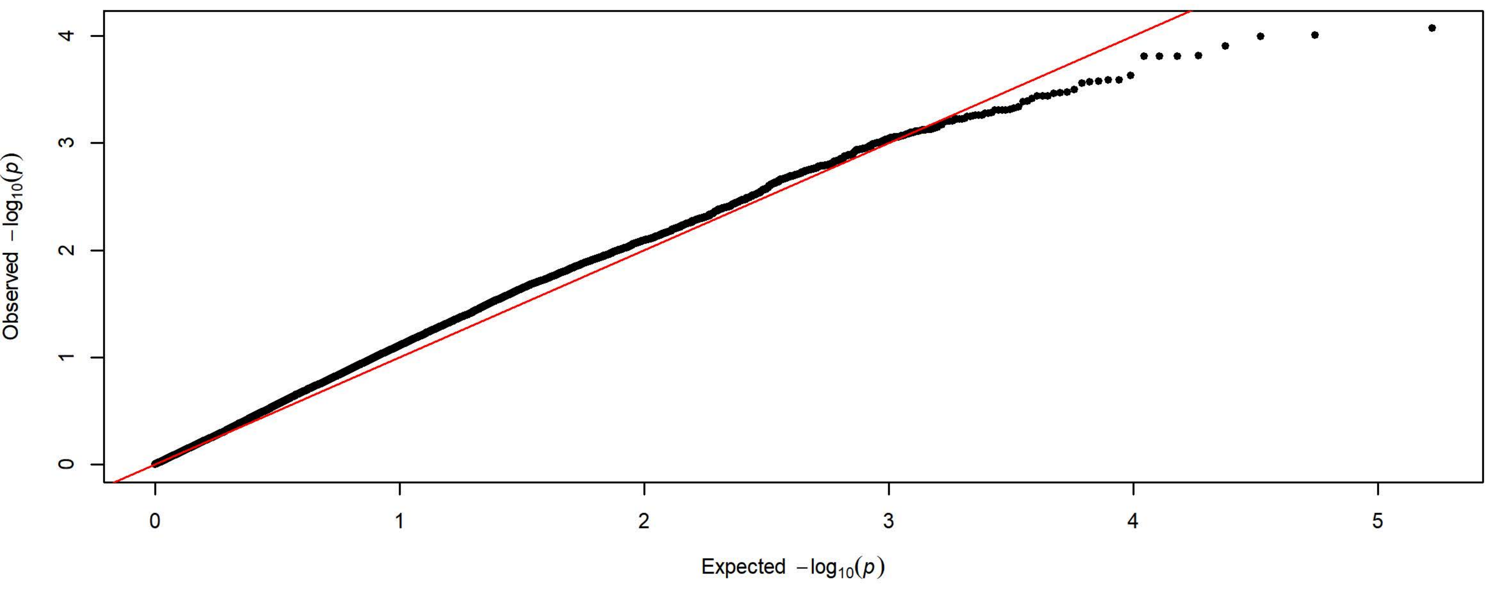

Q-Q Plot Lever Presses Day 4 - Meta-analysis of 7 Subgroups - 64k SNPs (n=3936)

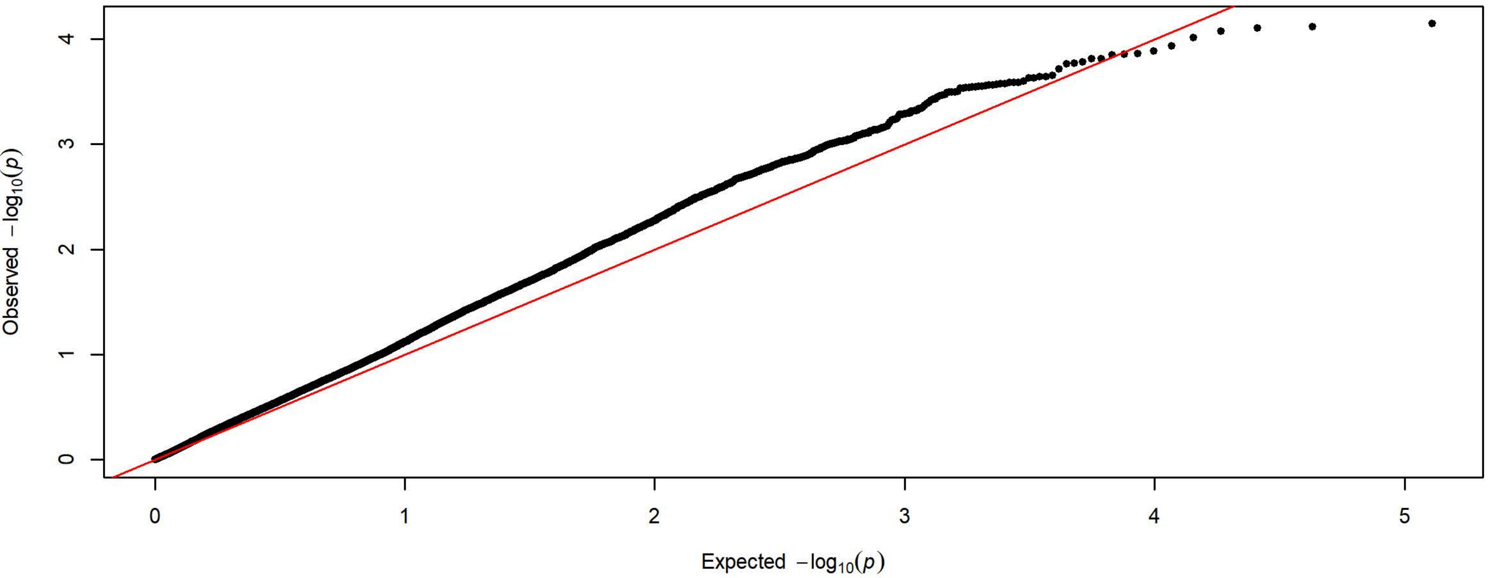


Q-Q Plot Lever Presses Day 4 - Charles River 4 Subgroups - 198k SNPs (n=1728)

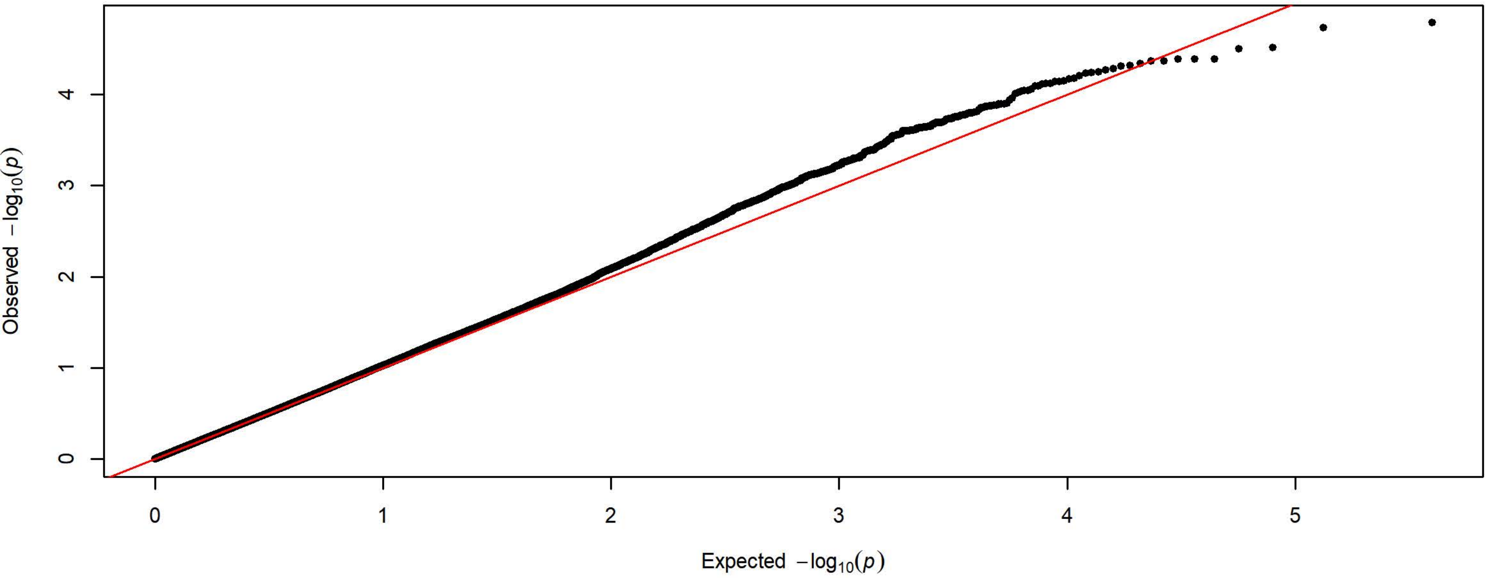


Q-Q Plot Lever Presses Day 4 - Harlan 3 Subgroups - 83k SNPs (n=2208)

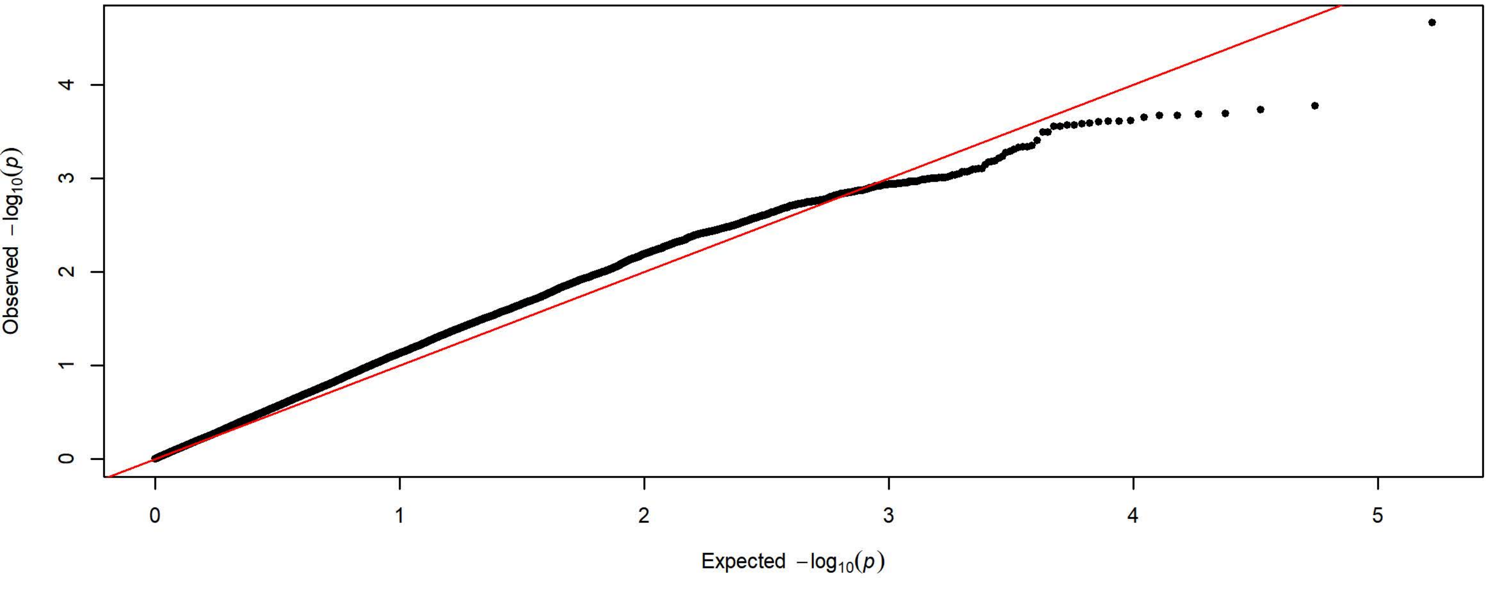

**Q-Q Plot Lever Presses Day 5 - Meta-analysis of 7 Subgroups - 64k SNPs (n=3936)**

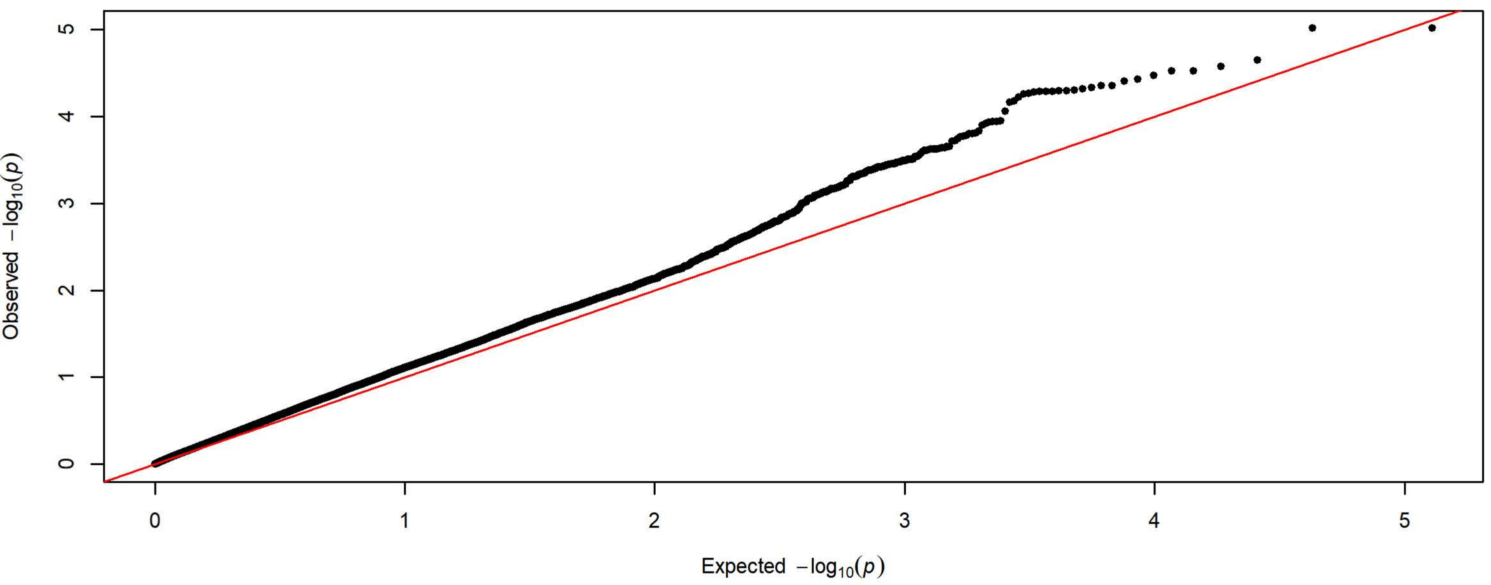


**Q-Q Plot Lever Presses Day 5 - Charles River 4 Subgroups - 198k SNPs (n=1728)**

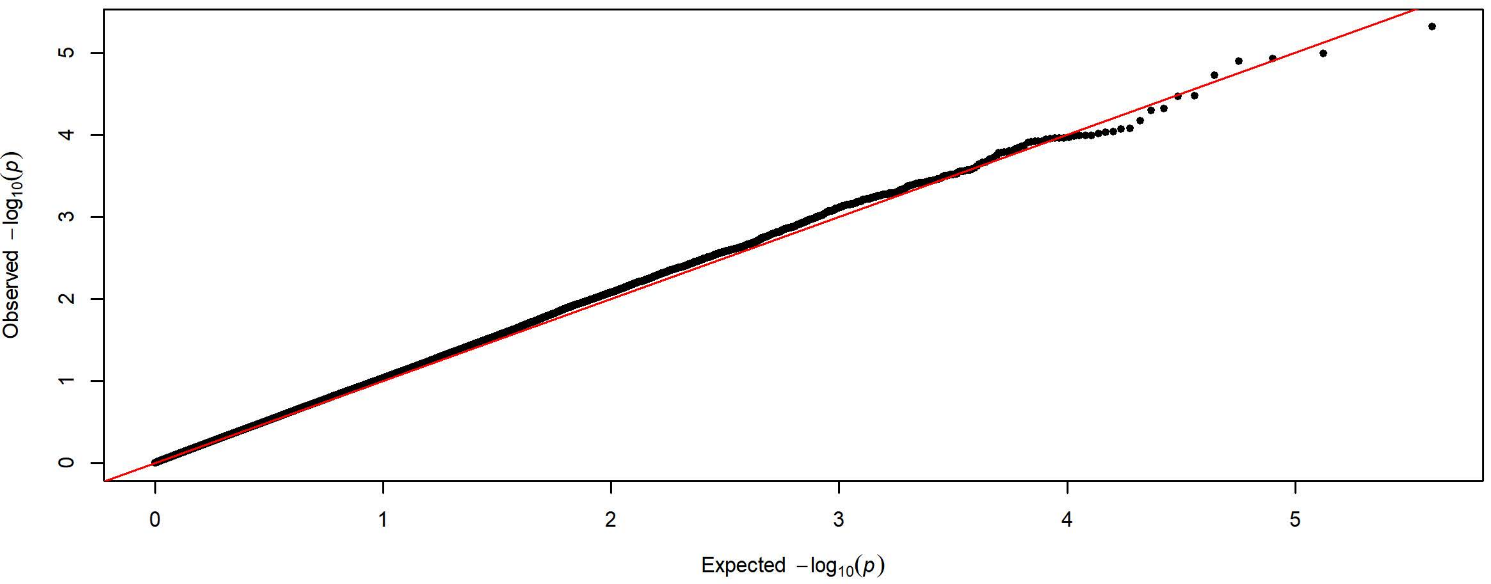


**Q-Q Plot Lever Presses Day 5 - Harlan 3 Subgroups - 83k SNPs (n=2208)**

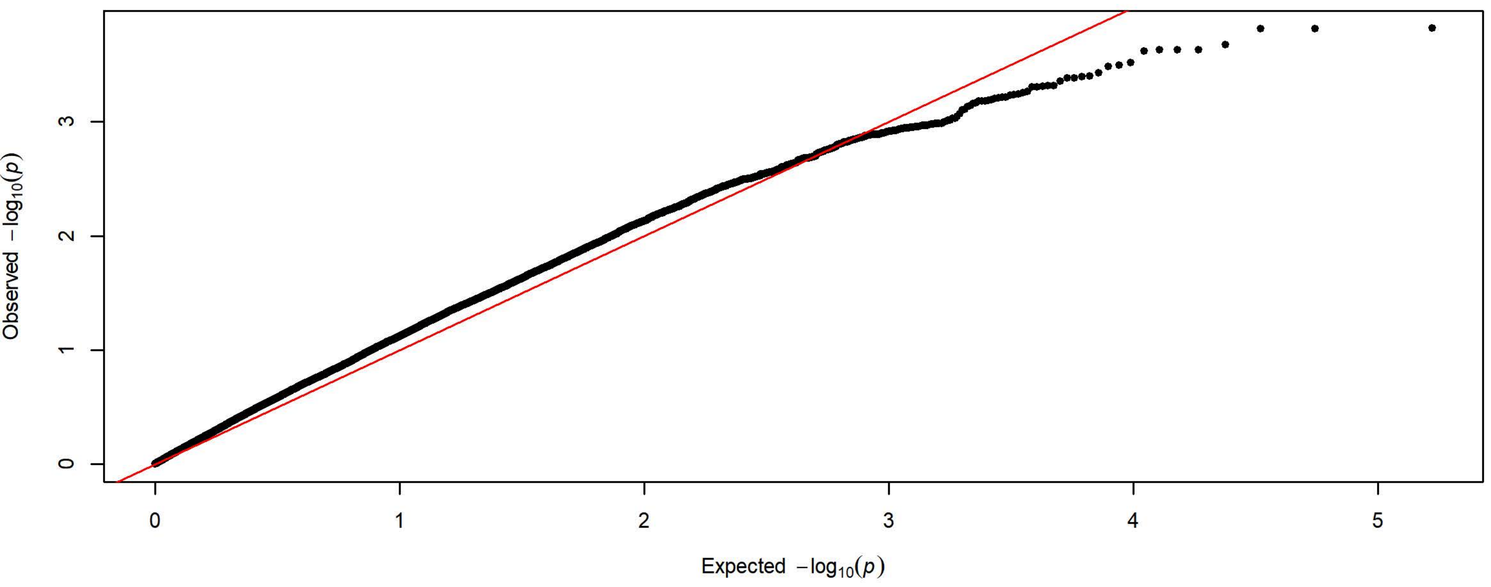

**Q-Q Plot Magazaine Entries Day 1 - Meta-analysis of 7 Subgroups - 64k SNPs (n=3933)**

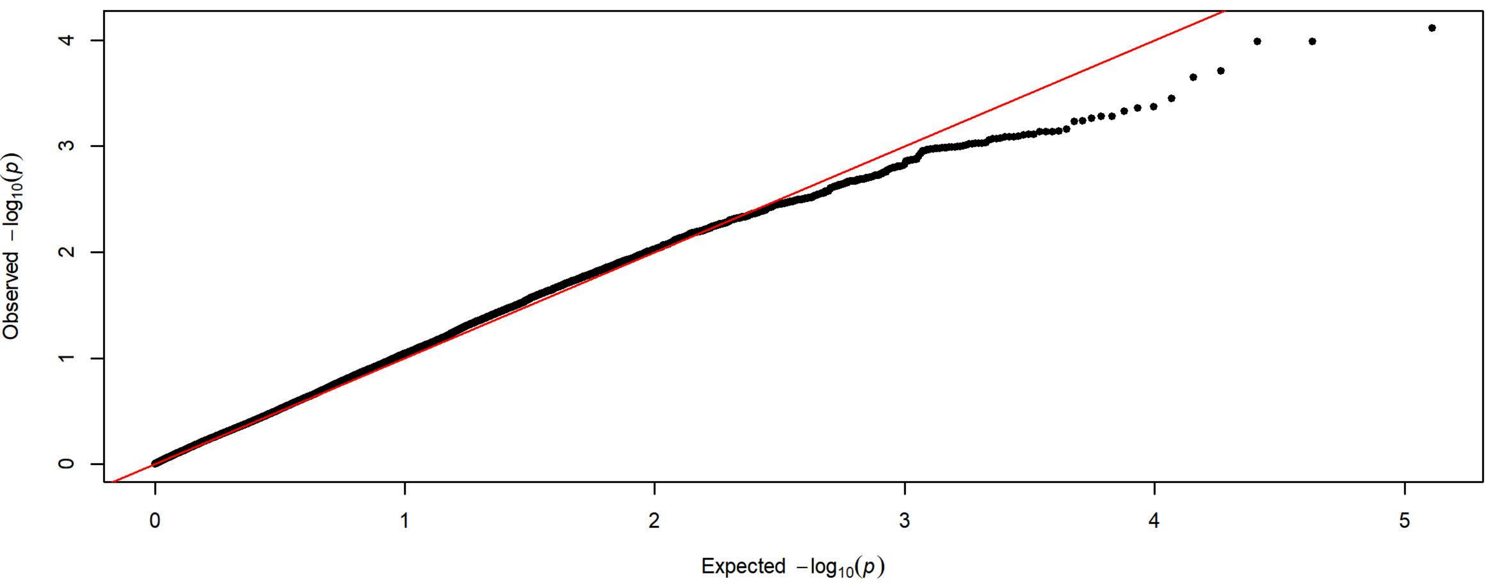


**Q-Q Plot Magazaine Entries Day 1 - Charles River 4 Subgroups - 198k SNPs (n=1728)**

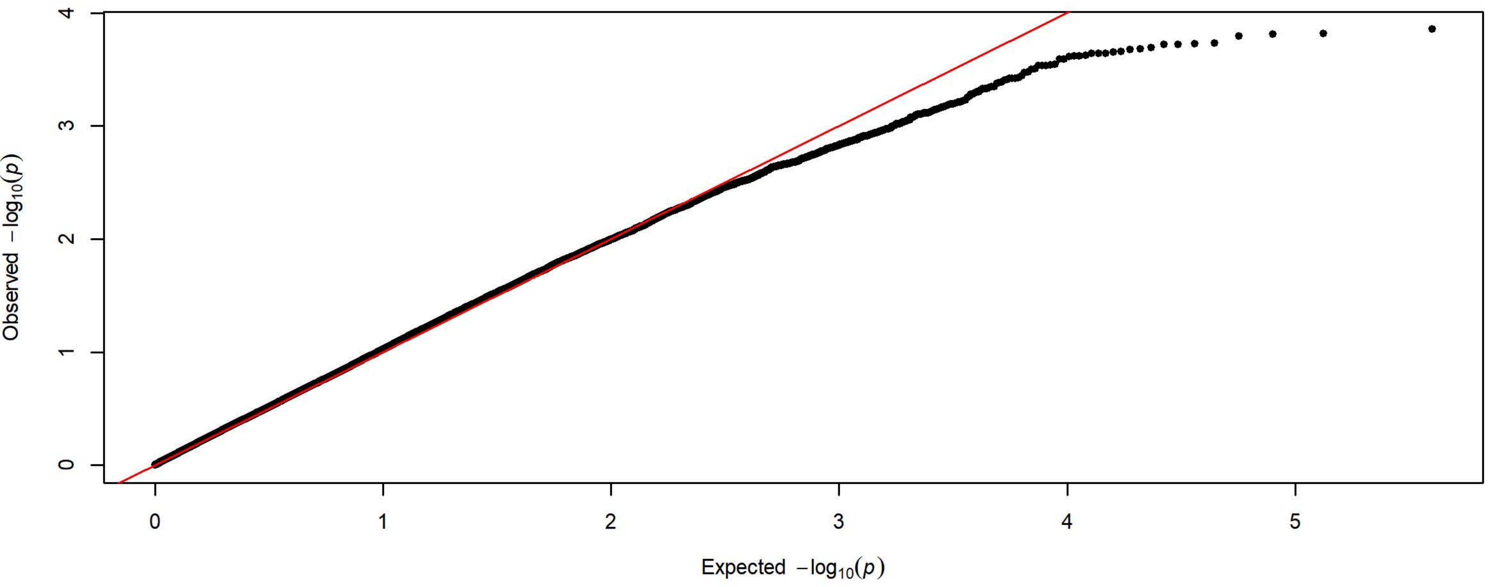


**Q-Q Plot Magazaine Entries Day 1 - Harlan 3 Subgroups - 83k SNPs (n=2205)**

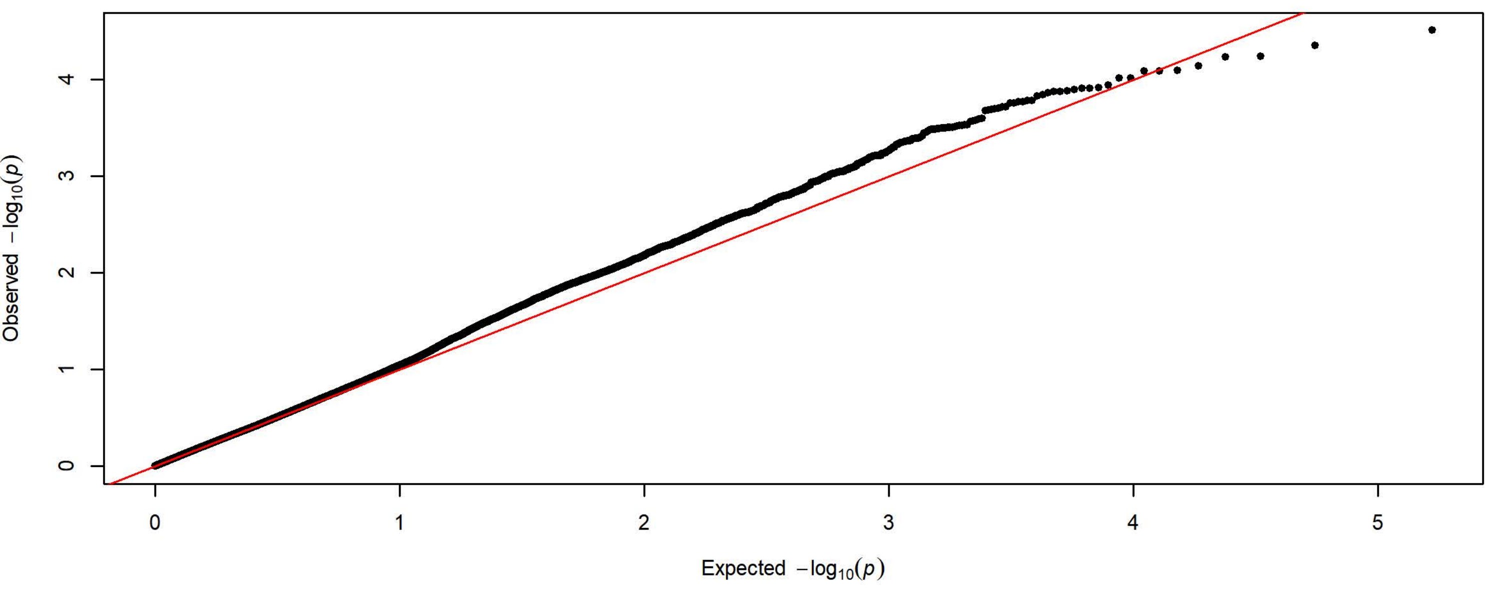

**Q-Q Plot Magazaine Entries Day 2 - Meta-analysis of 7 Subgroups - 64k SNPs (n=3933)**

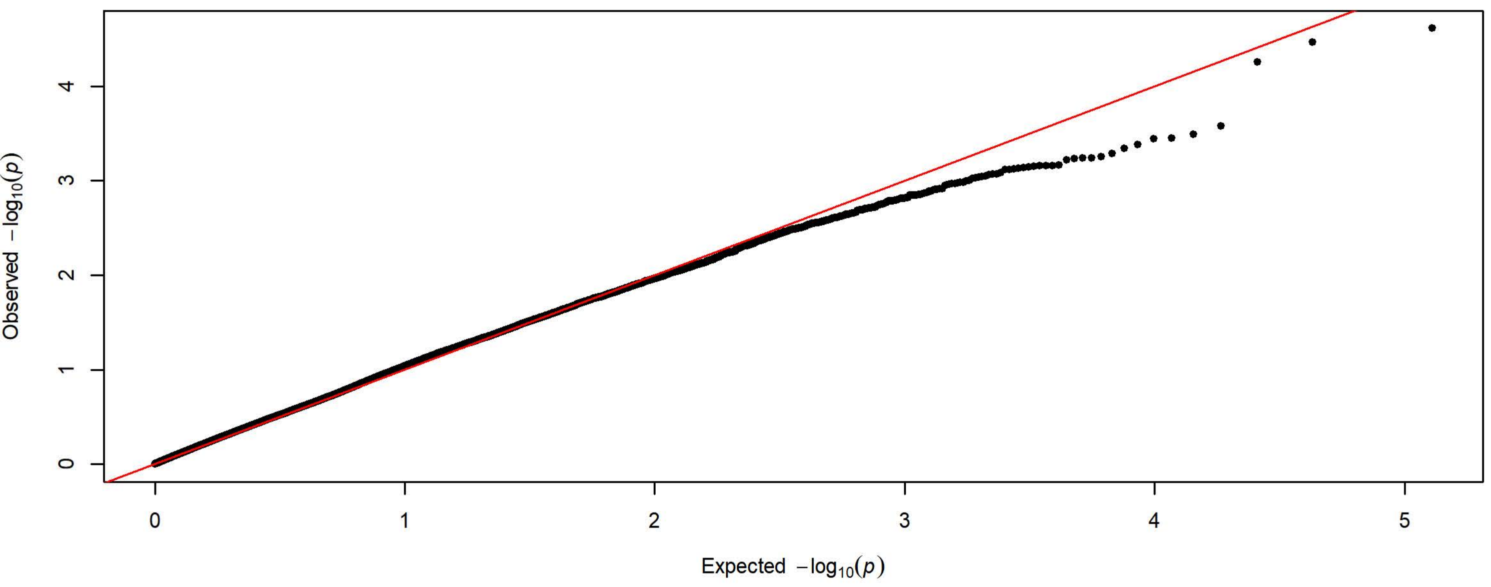


**Q-Q Plot Magazaine Entries Day 2 - Charles River 4 Subgroups - 198k SNPs (n=1726)**

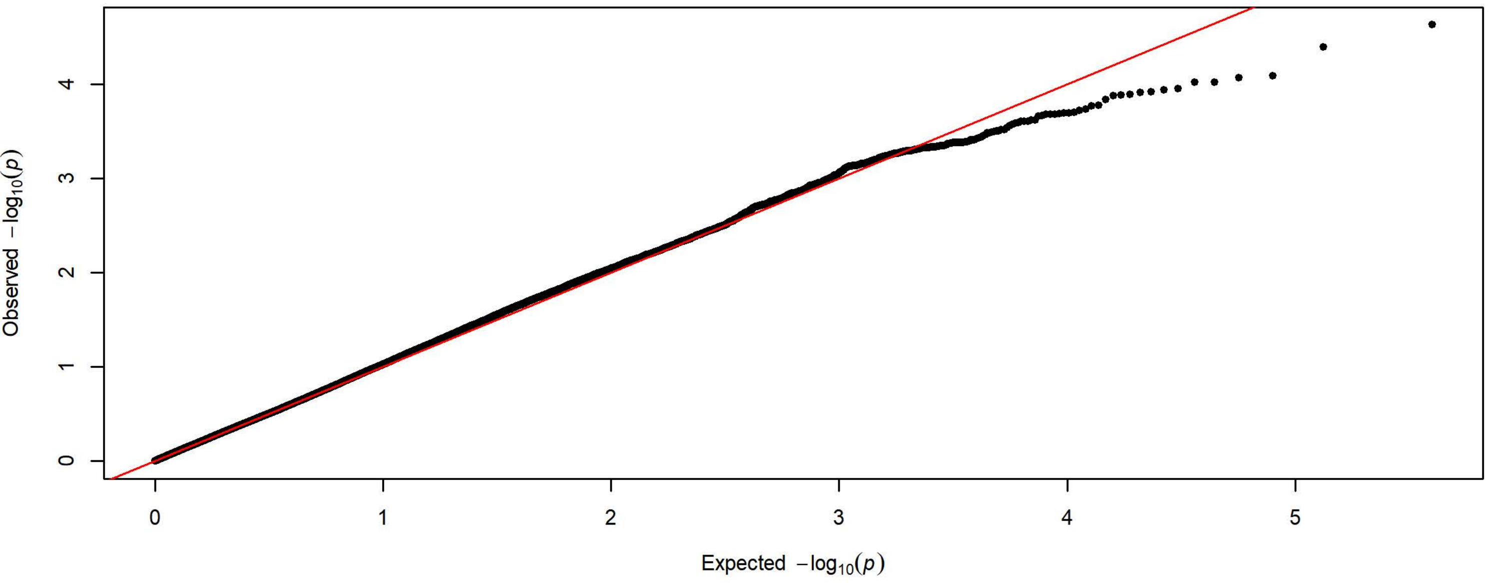


**Q-Q Plot Magazaine Entries Day 2 - Harlan 3 Subgroups - 83k SNPs (n=2207)**

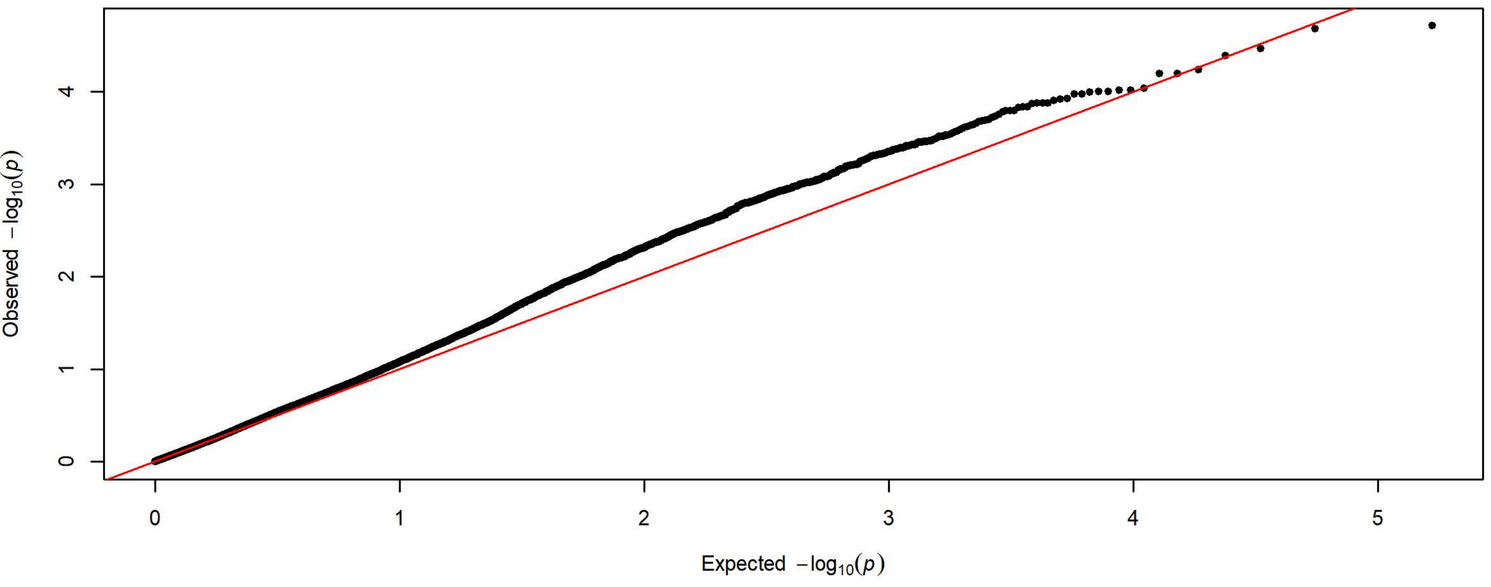

Q-Q Plot Magazaine Entries Day 3 - Meta-analysis of 7 Subgroups - 64k SNPs (n=3931)

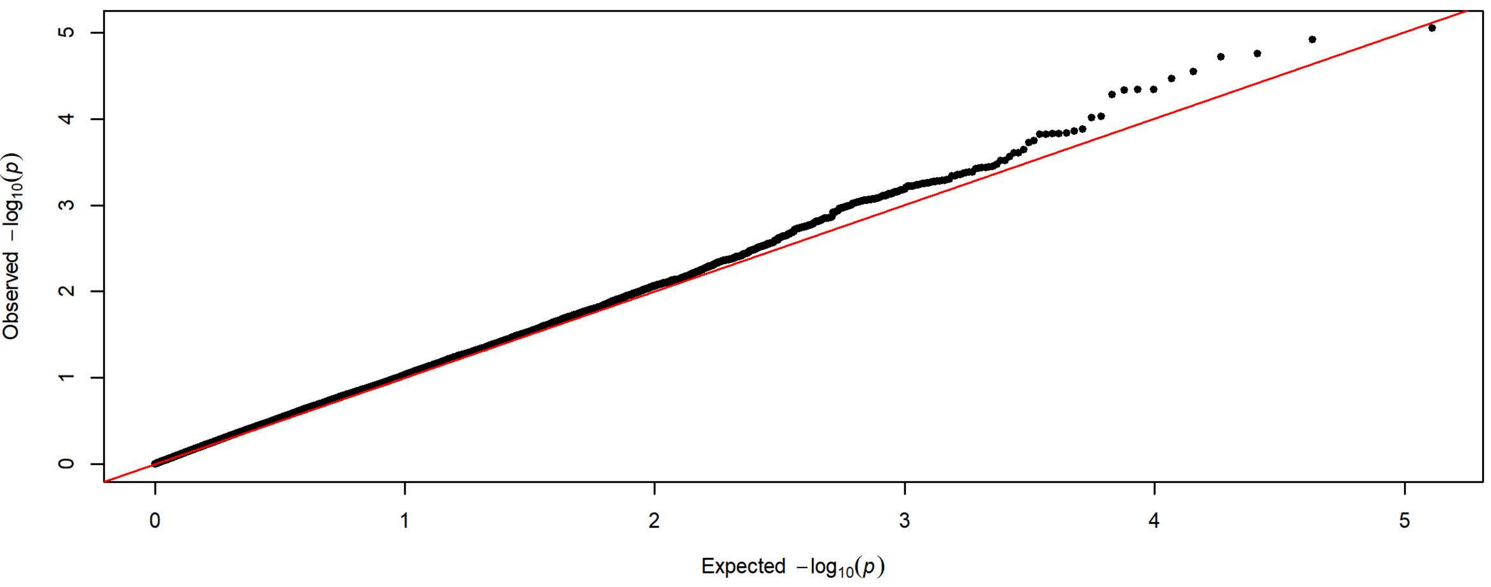


Q-Q Plot Magazaine Entries Day 3 - Charles River 4 Subgroups - 198k SNPs (n=1727)

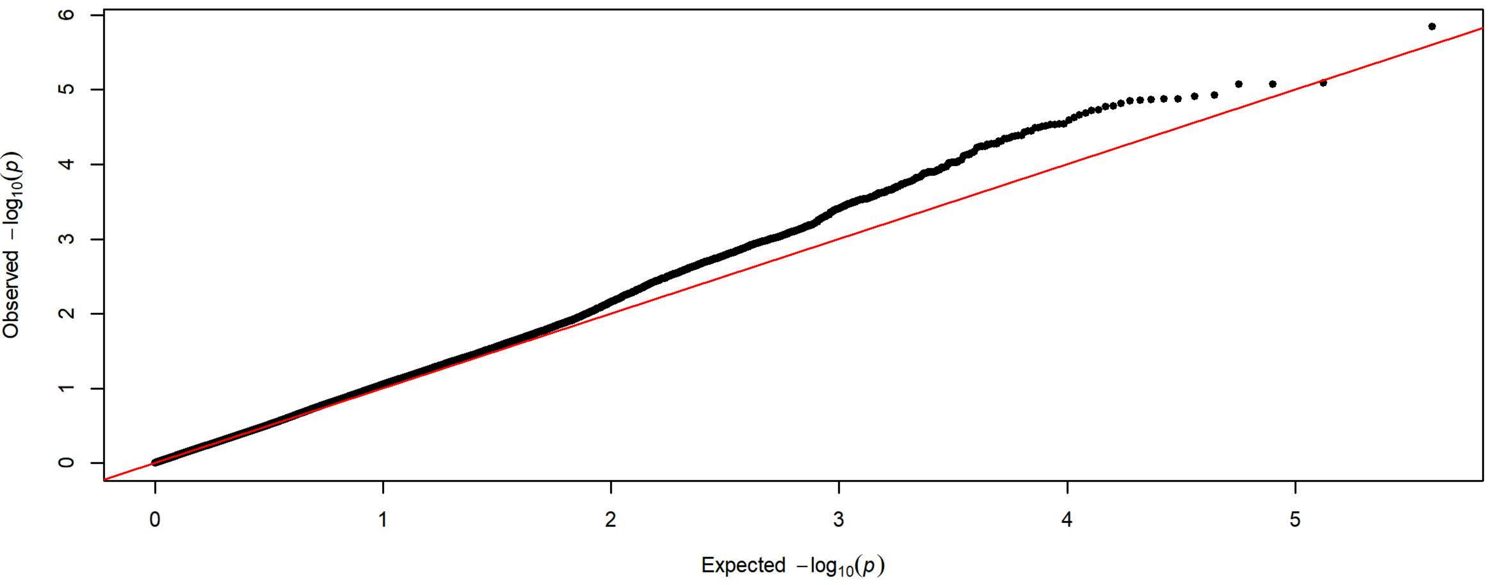


Q-Q Plot Magazaine Entries Day 3 - Harlan 3 Subgroups - 83k SNPs (n=2204)

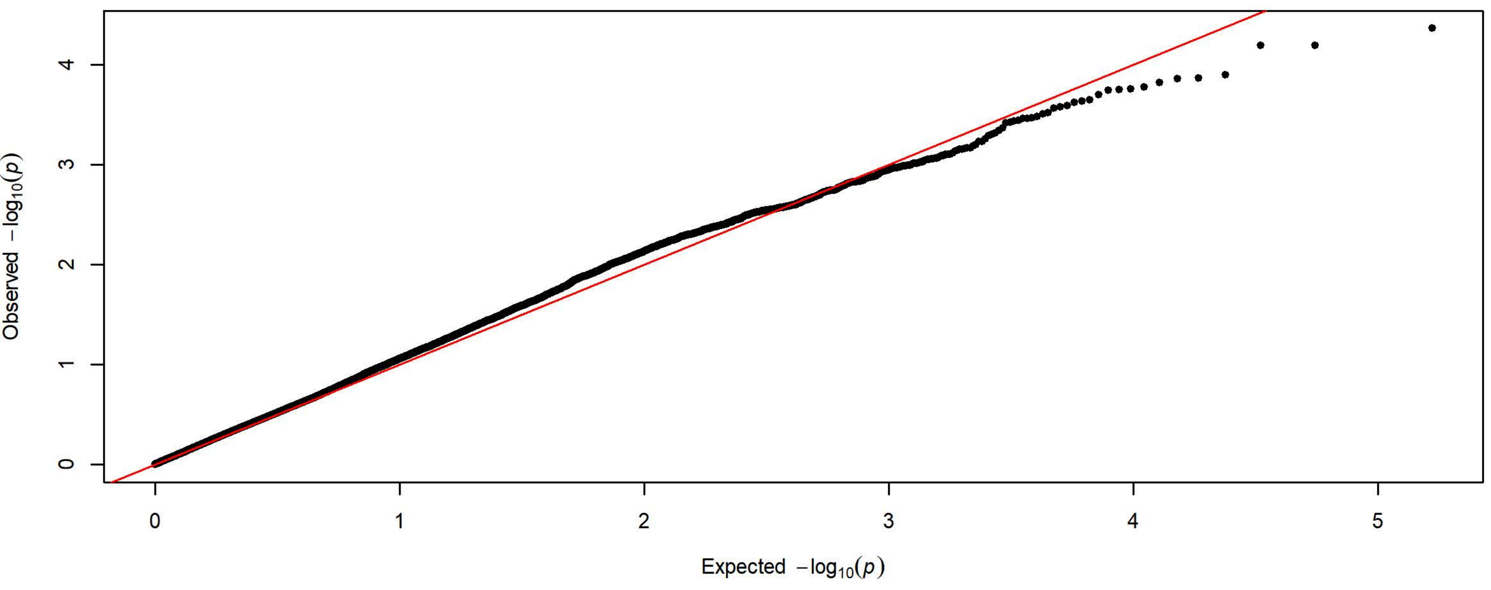

**Q-Q Plot Magazaine Entries Day 4 - Meta-analysis of 7 Subgroups - 64k SNPs (n=3936)**

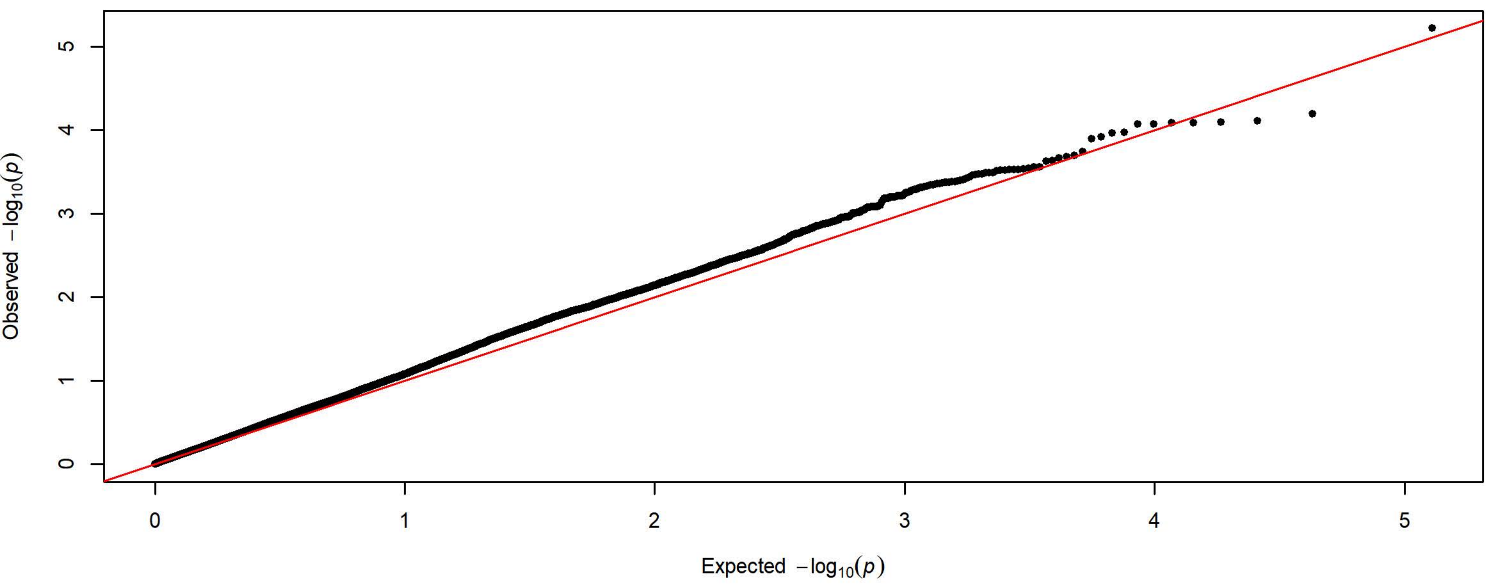


**Q-Q Plot Magazaine Entries Day 4 - Charles River 4 Subgroups - 198k SNPs (n=1728)**

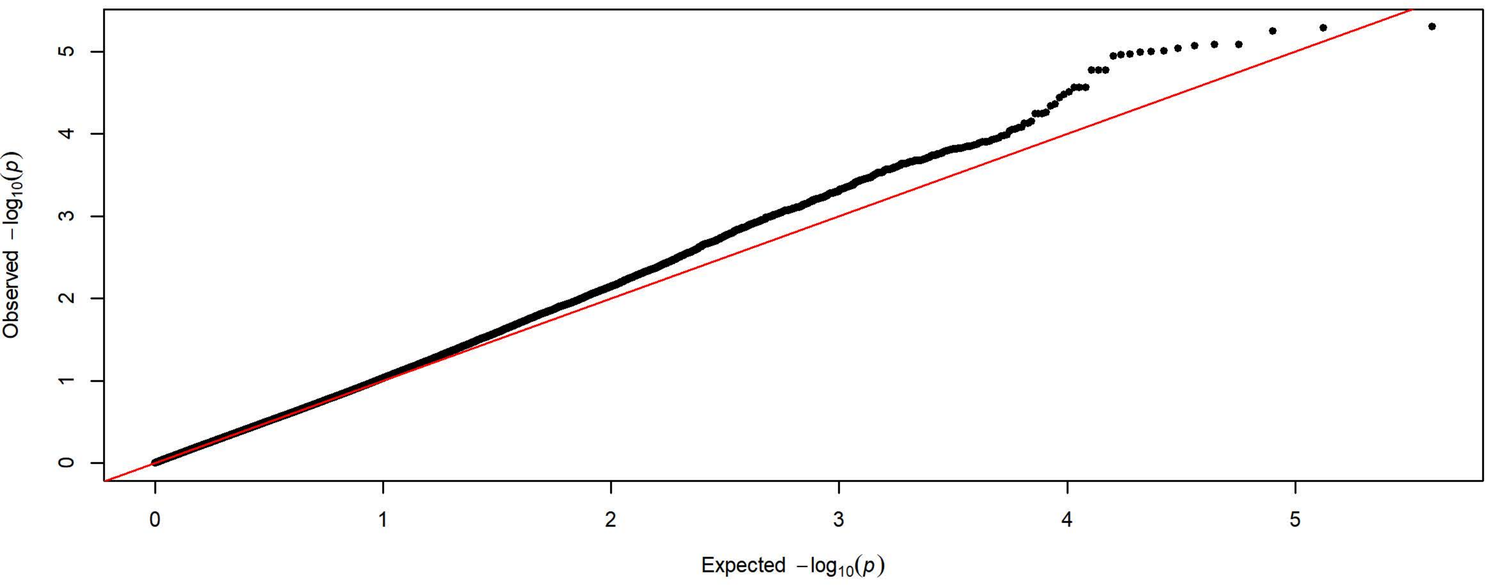


**Q-Q Plot Magazaine Entries Day 4 - Harlan 3 Subgroups - 83k SNPs (n=2208)**

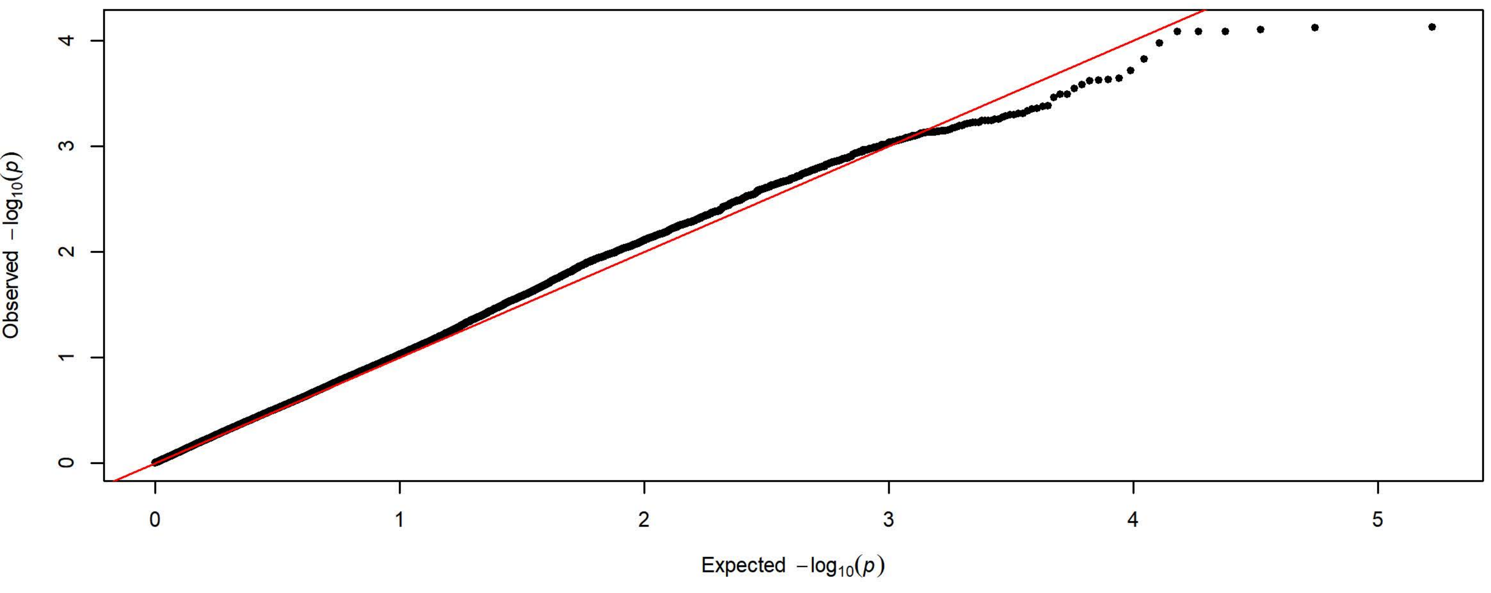

Q-Q Plot Magazaine Entries Day 5 - Meta-analysis of 7 Subgroups - 64k SNPs (n=3936)

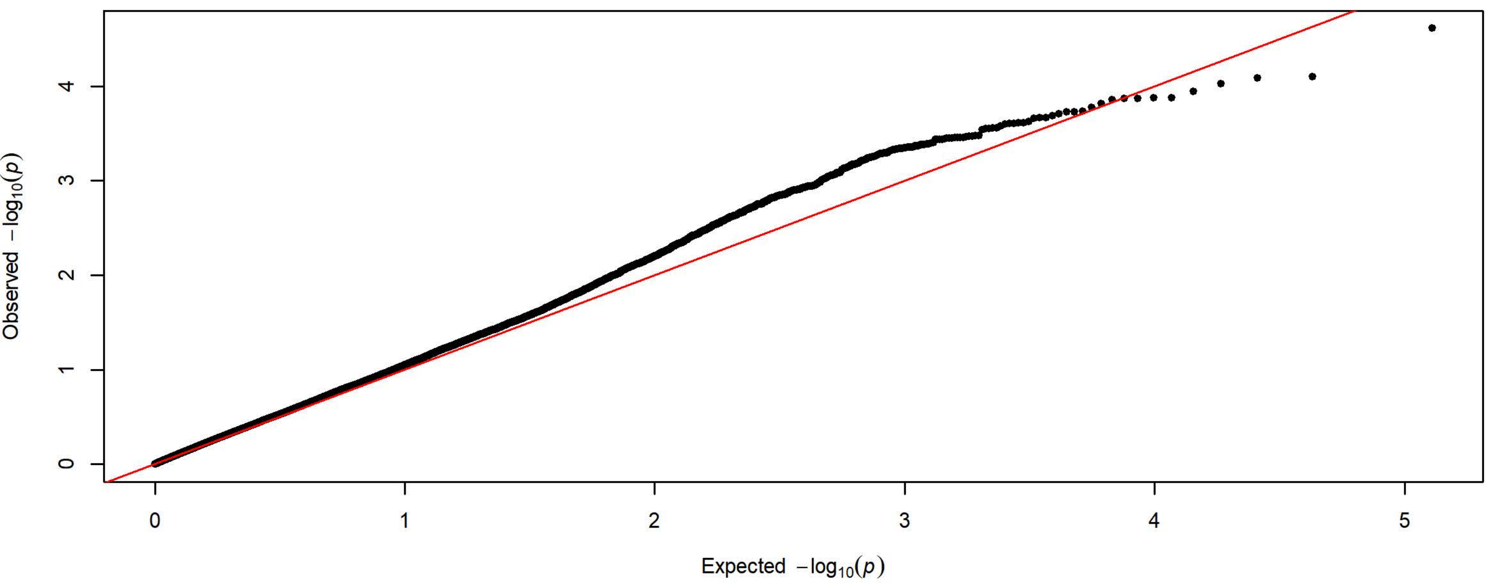


Q-Q Plot Magazaine Entries Day 5 - Charles River 4 Subgroups - 198k SNPs (n=1728)

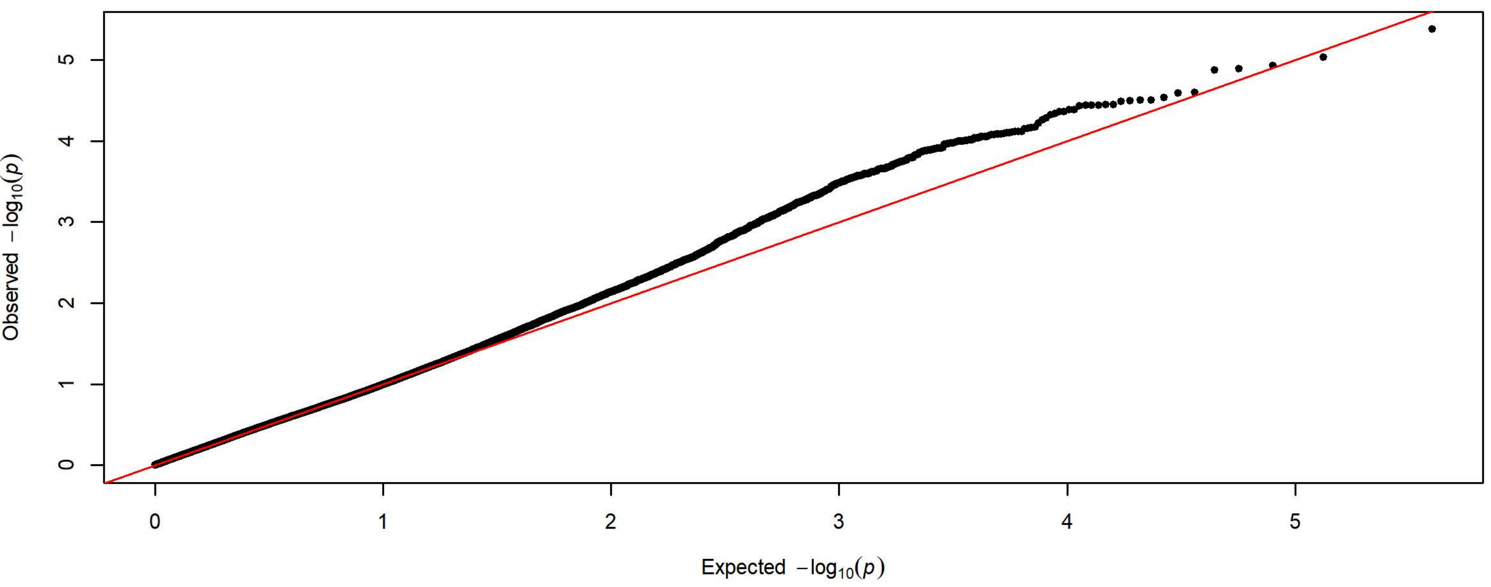


Q-Q Plot Magazaine Entries Day 5 - Harlan 3 Subgroups - 83k SNPs (n=2208)

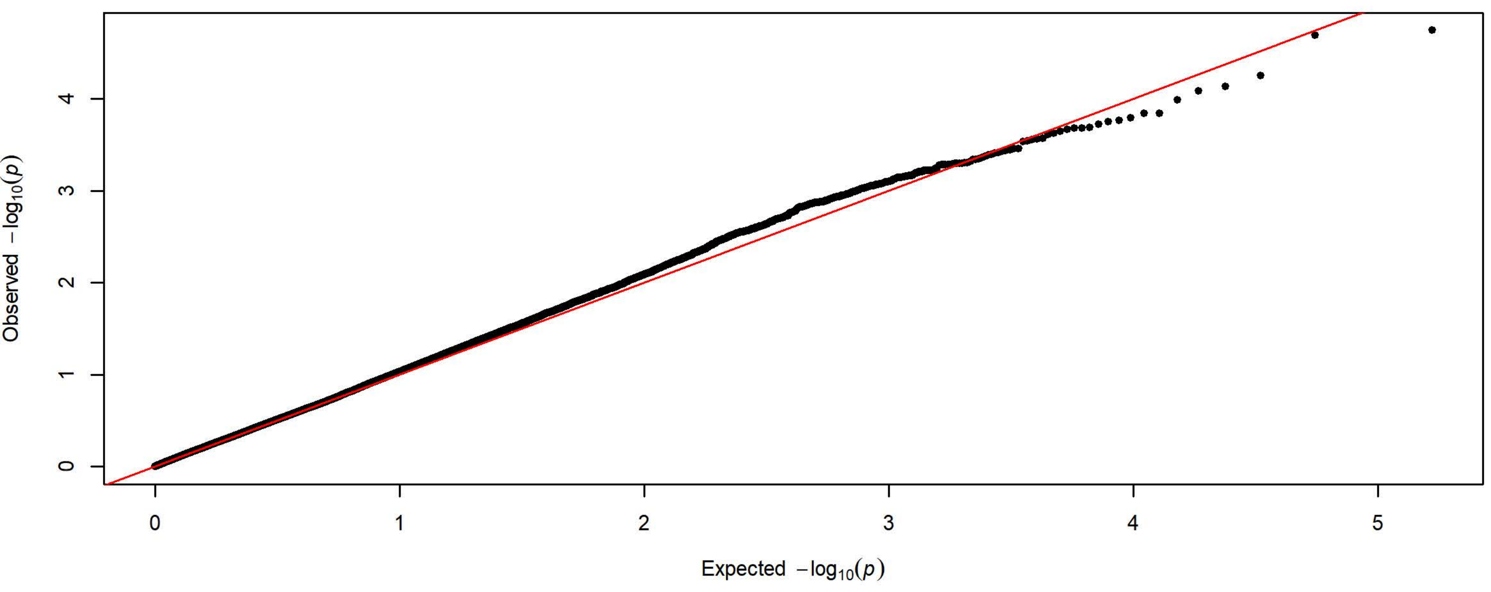

**Q-Q Plot Magazine Entries NCS Day 1 - Meta-analysis of 7 Subgroups - 64k SNPs (n=3933)**

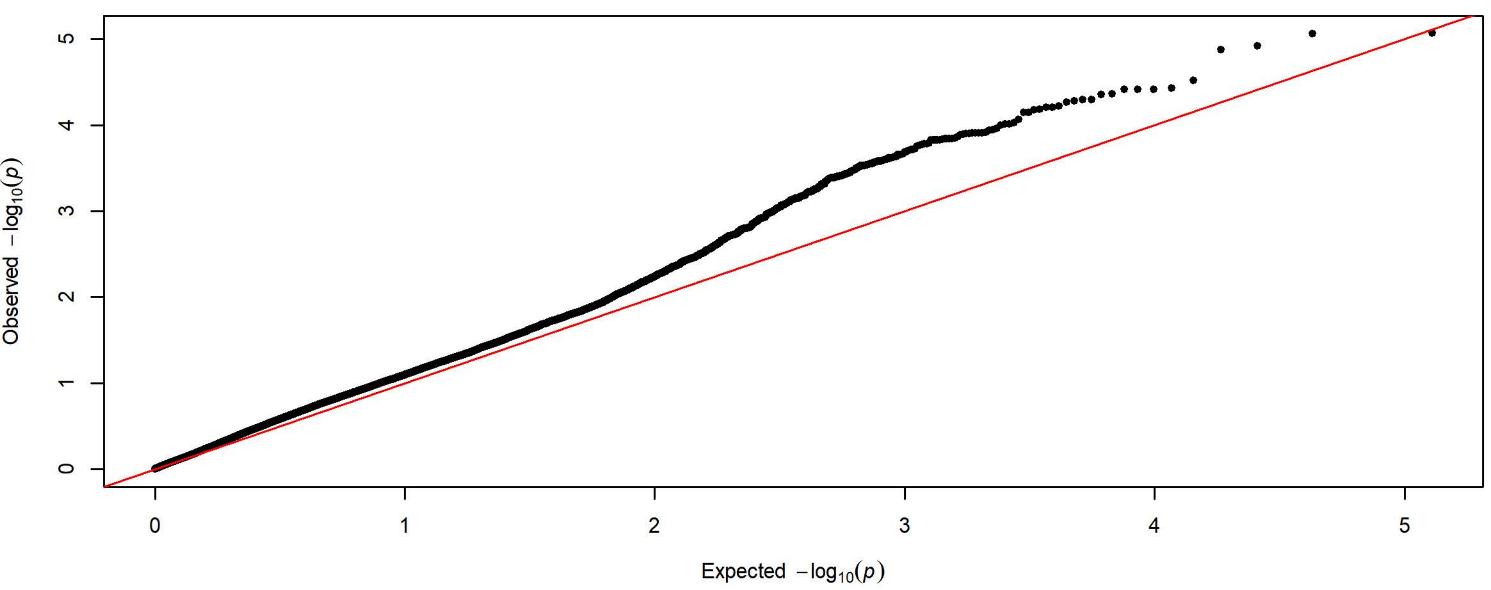


**Q-Q Plot Magazine Entries NCS Day 1 - Charles River 4 Subgroups - 198k SNPs (n=1728)**

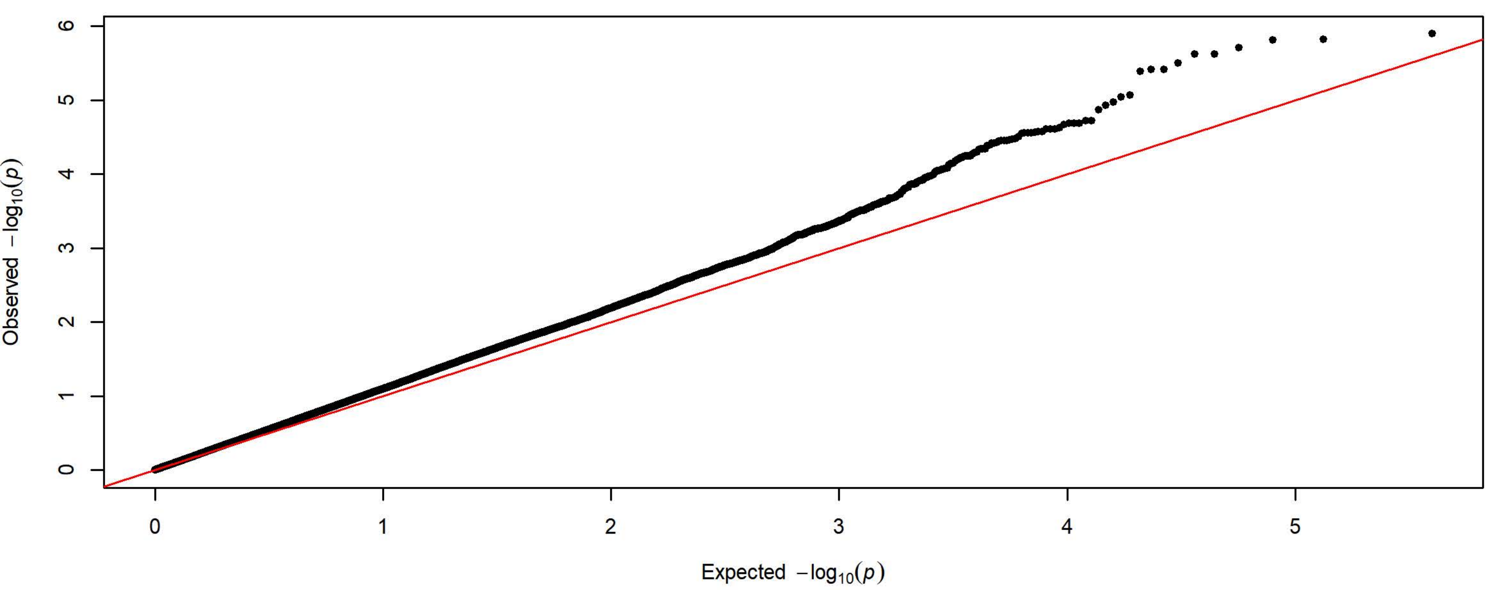


**Q-Q Plot Magazine Entries NCS Day 1 - Harlan 3 Subgroups - 83k SNPs (n=2205)**

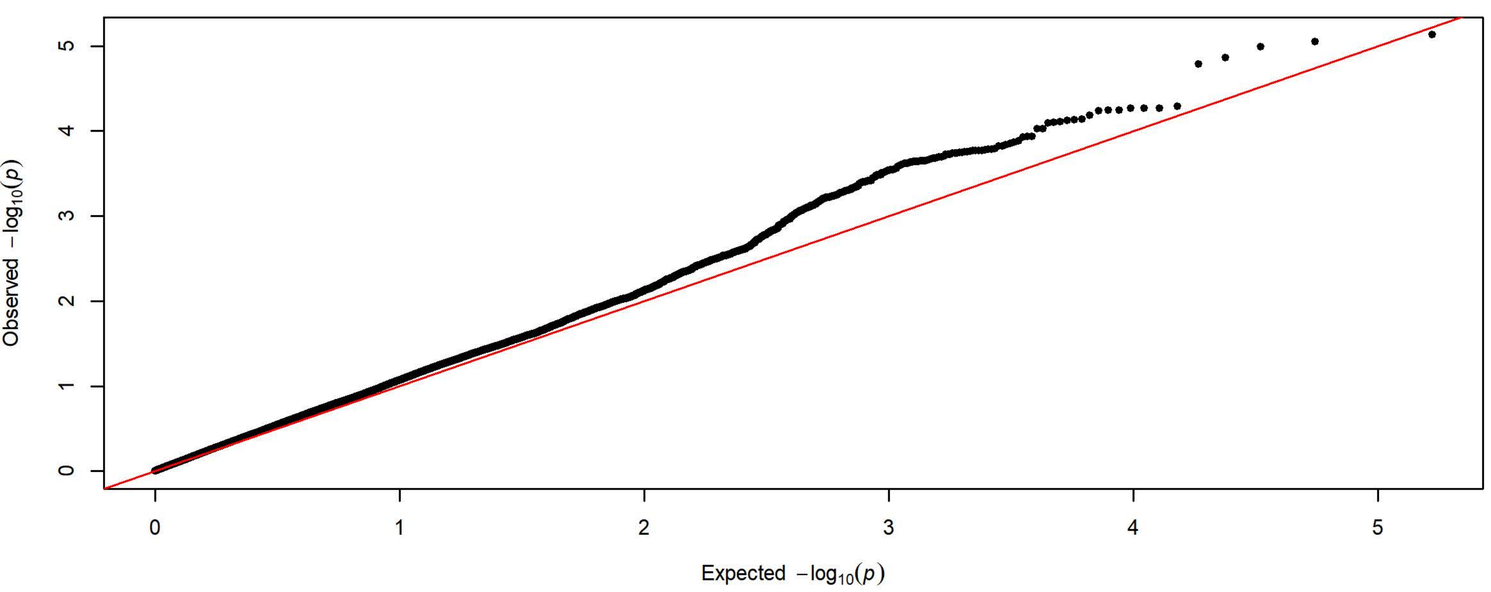

**Q-Q Plot Magazine Entries NCS Day 2 - Meta-analysis of 7 Subgroups - 64k SNPs (n=3933)**

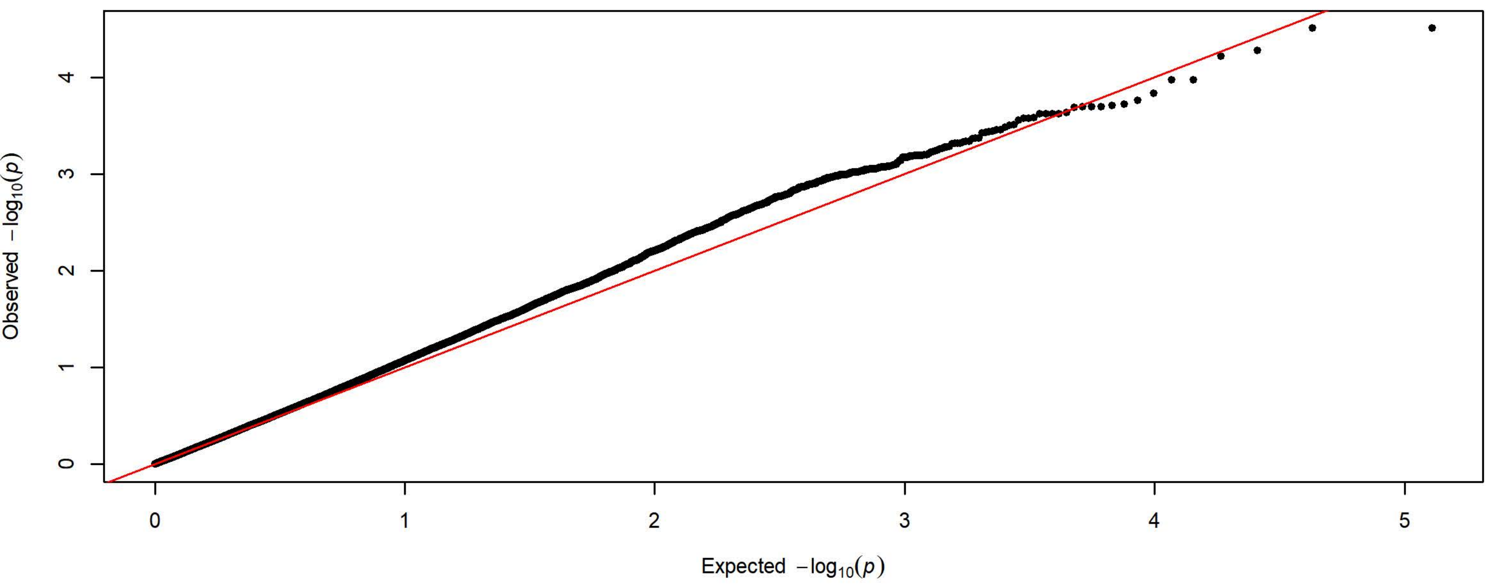


**Q-Q Plot Magazine Entries NCS Day 2 - Charles River 4 Subgroups - 198k SNPs (n=1726)**

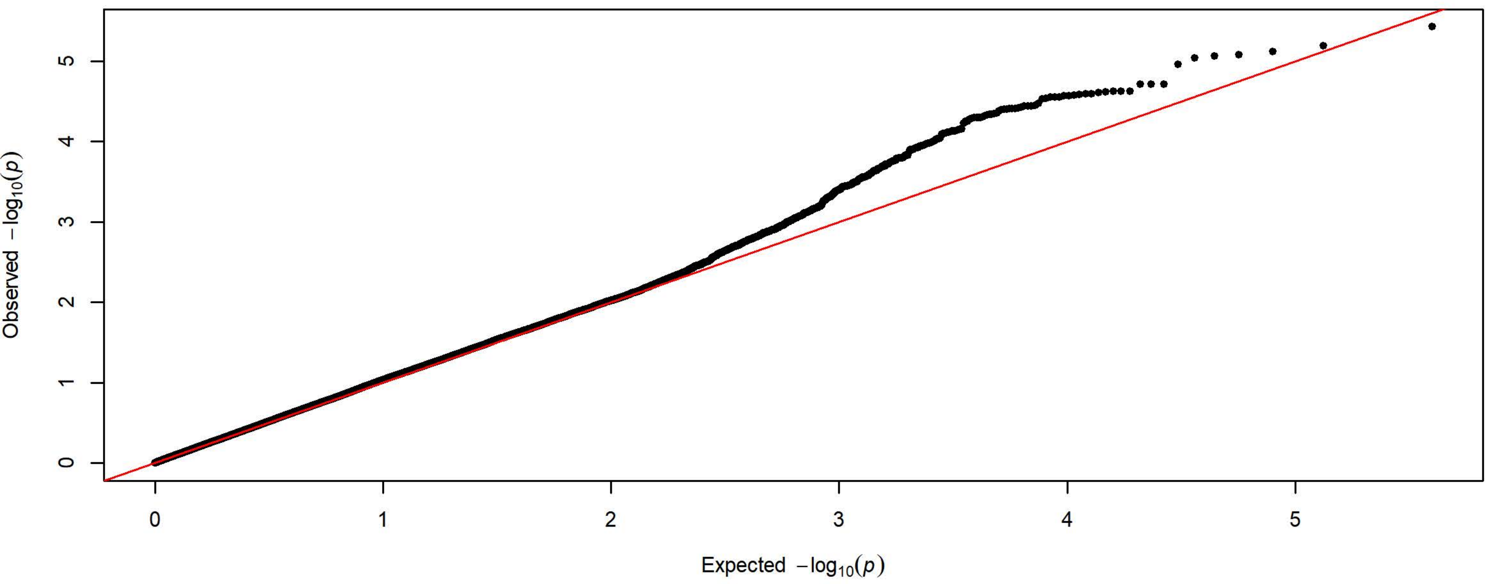


**Q-Q Plot Magazine Entries NCS Day 2 - Harlan 3 Subgroups - 83k SNPs (n=2207)**

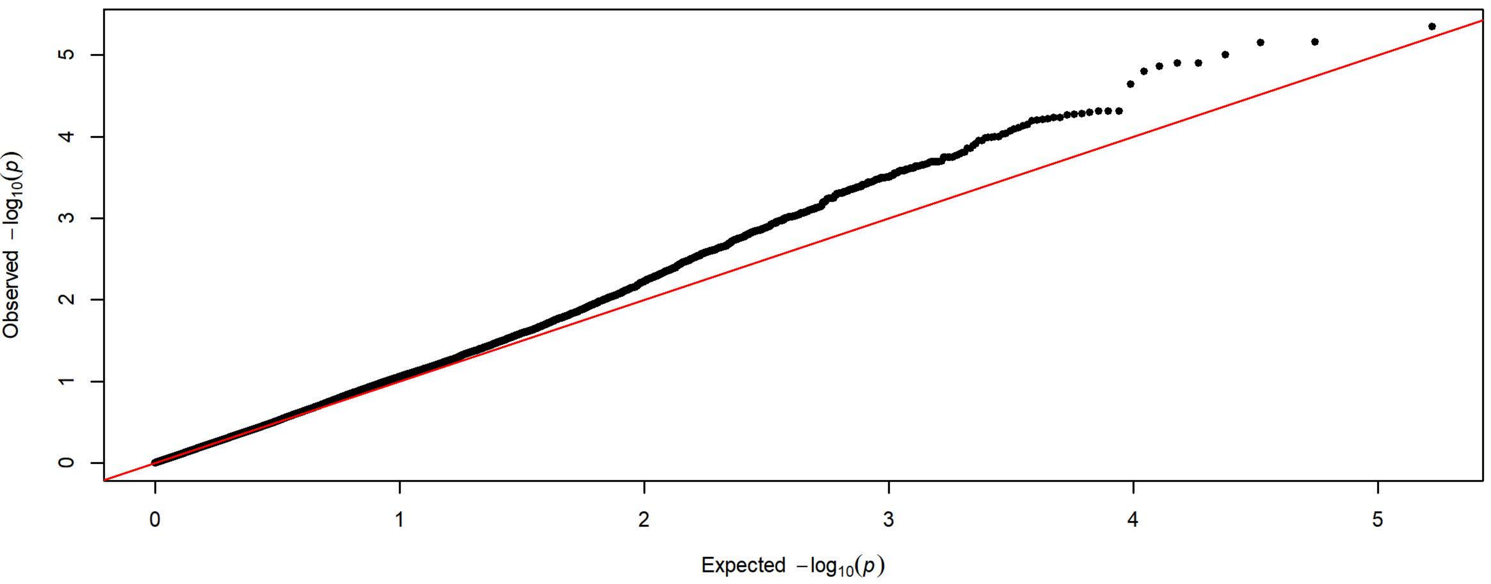

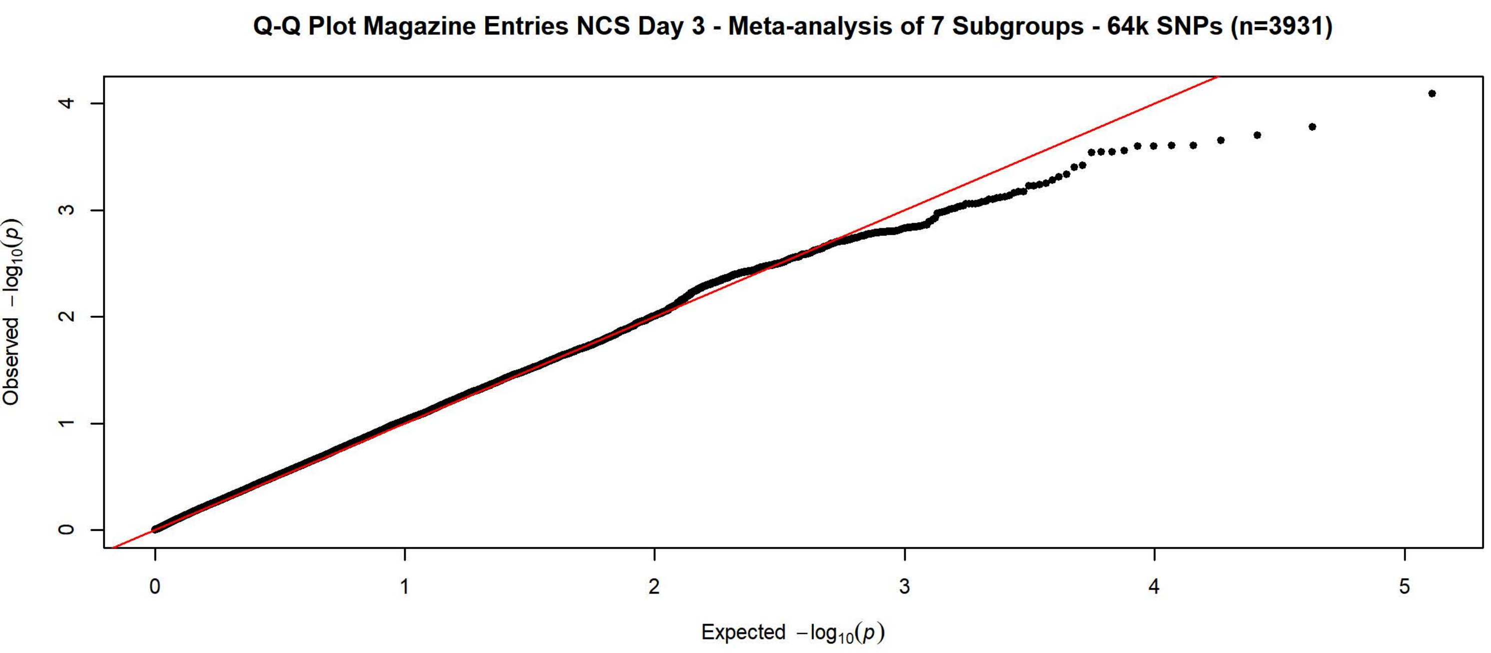

Q-Q Plot Magazine Entries NCS Day 3 - Meta-analysis of 7 Subgroups - 64k SNPs (n=3931)
Observed $-\log_{10}(p)$
0
1
2
3
4
0
1
2
3
4
5
Expected $-\log_{10}(p)$


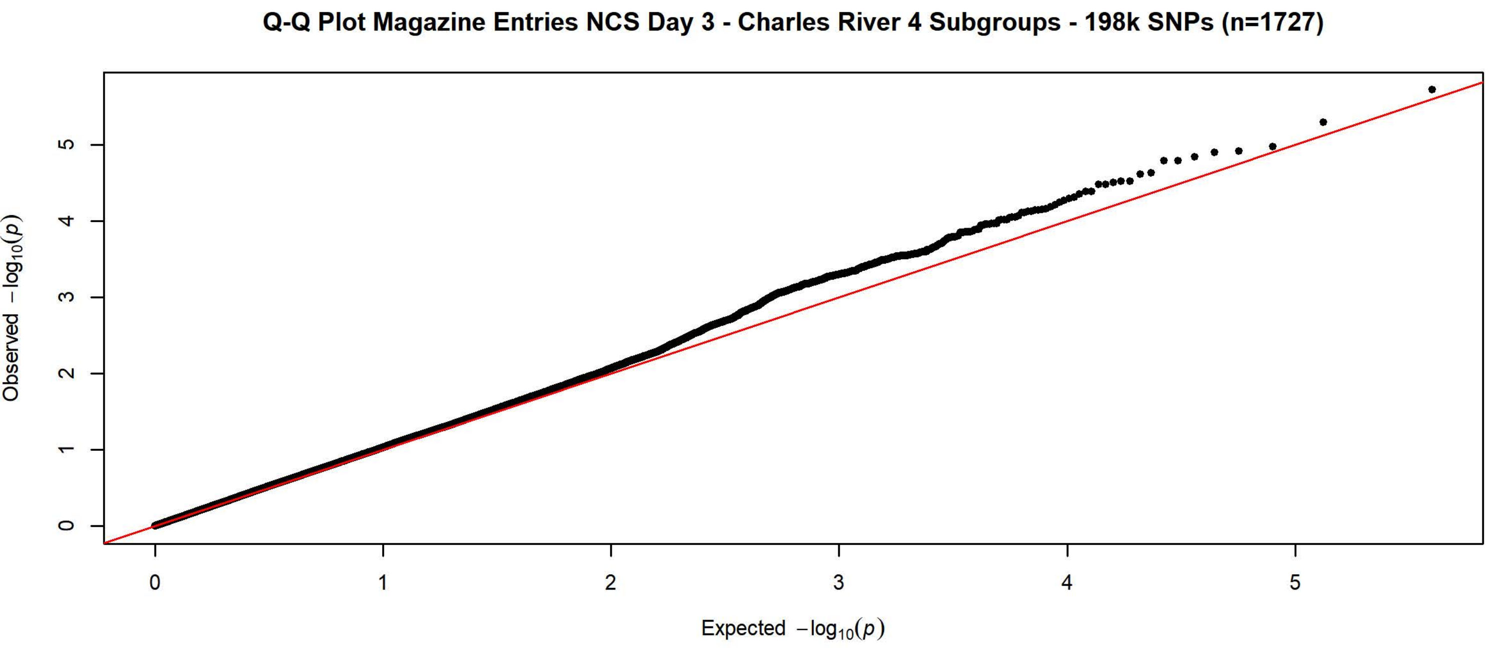

Q-Q Plot Magazine Entries NCS Day 3 - Charles River 4 Subgroups - 198k SNPs (n=1727)
Observed $-\log_{10}(p)$
0
1
2
3
4
5
0
1
2
3
4
5
Expected $-\log_{10}(p)$


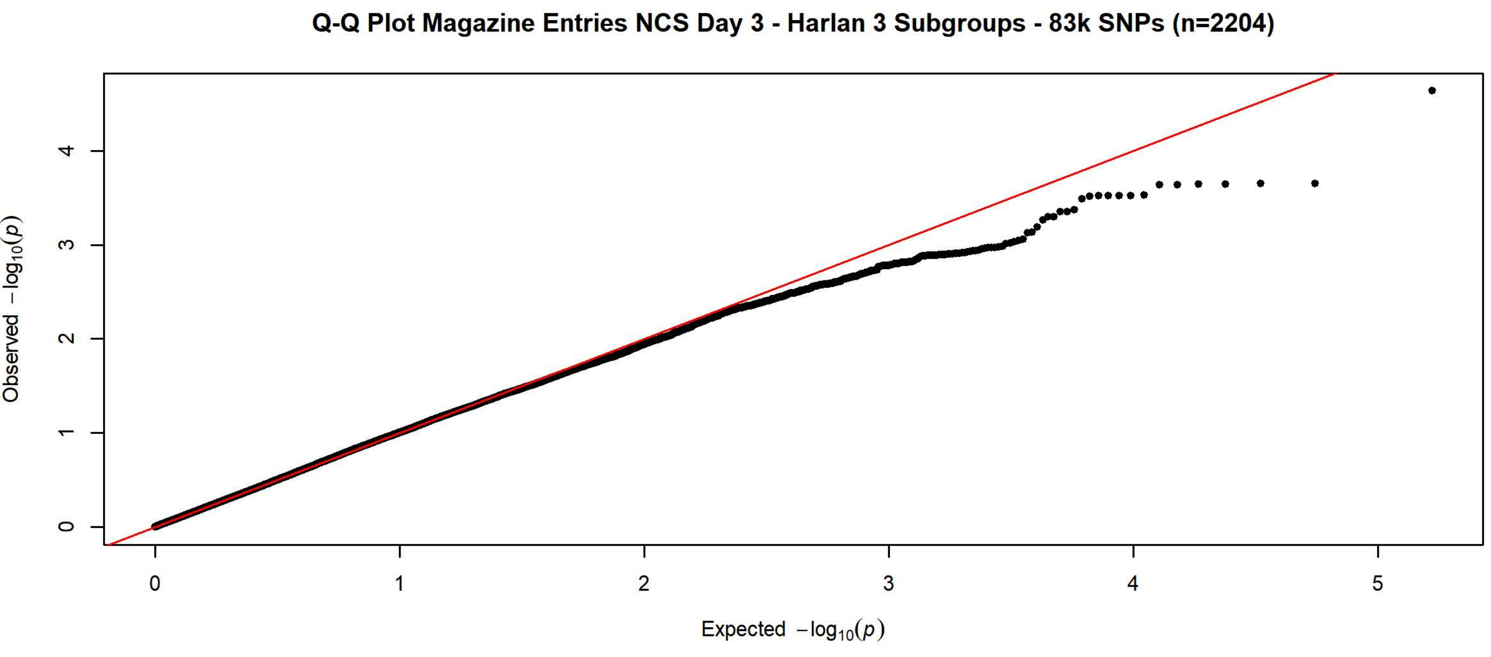

Q-Q Plot Magazine Entries NCS Day 3 - Harlan 3 Subgroups - 83k SNPs (n=2204)
Observed $-\log_{10}(p)$
0
1
2
3
4
0
1
2
3
4
5
Expected $-\log_{10}(p)$

Q-Q Plot Magazine Entries NCS Day 4 - Meta-analysis of 7 Subgroups - 64k SNPs (n=3936)

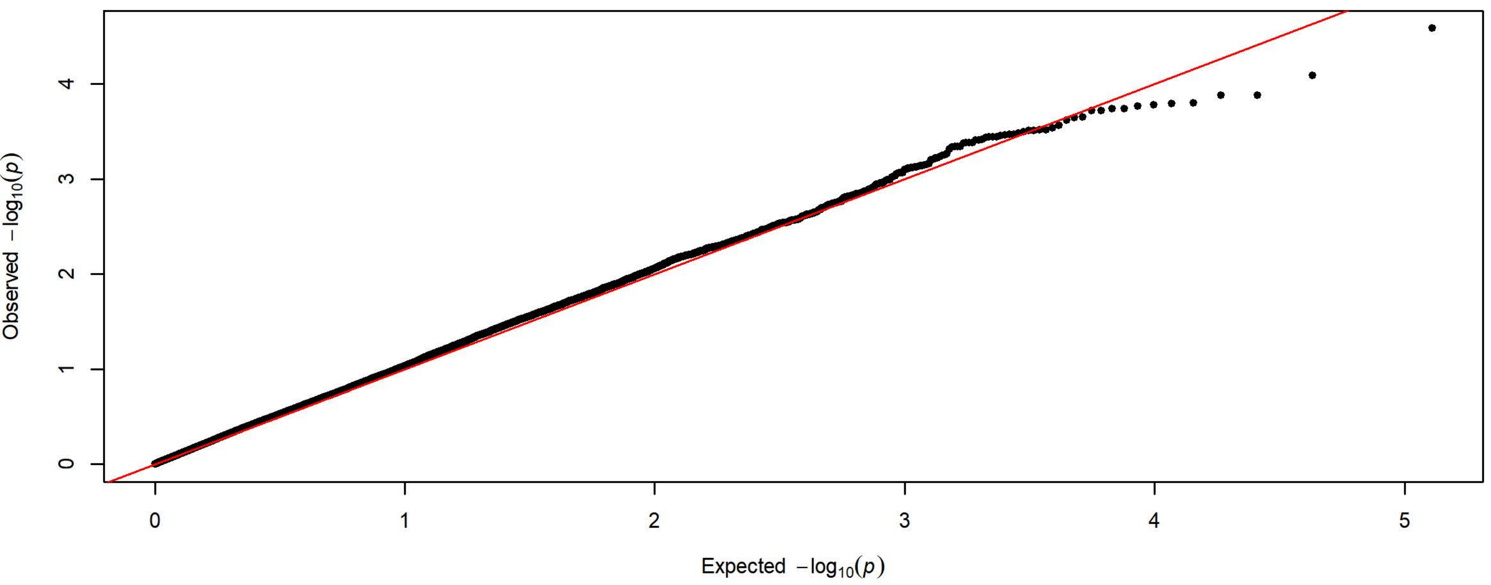


Q-Q Plot Magazine Entries NCS Day 4 - Charles River 4 Subgroups - 198k SNPs (n=1728)

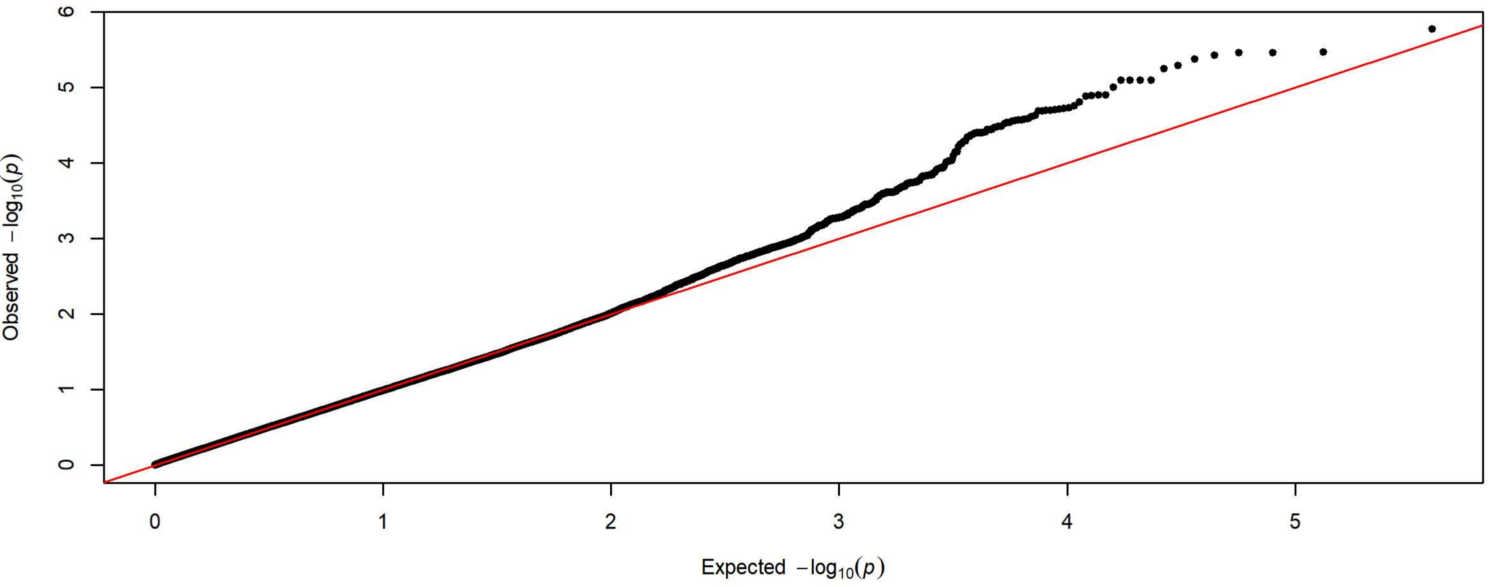


Q-Q Plot Magazine Entries NCS Day 4 - Harlan 3 Subgroups - 83k SNPs (n=2208)

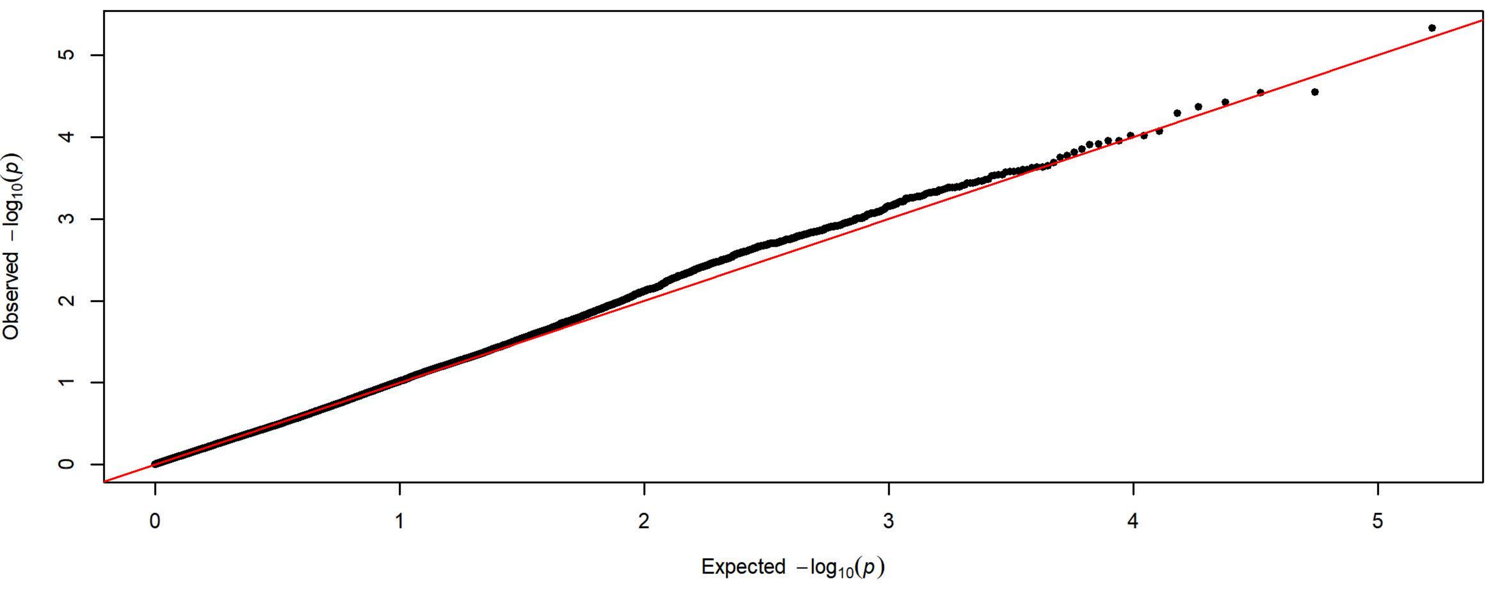

**Q-Q Plot Magazine Entries NCS Day 5 - Meta-analysis of 7 Subgroups - 64k SNPs (n=3936)**

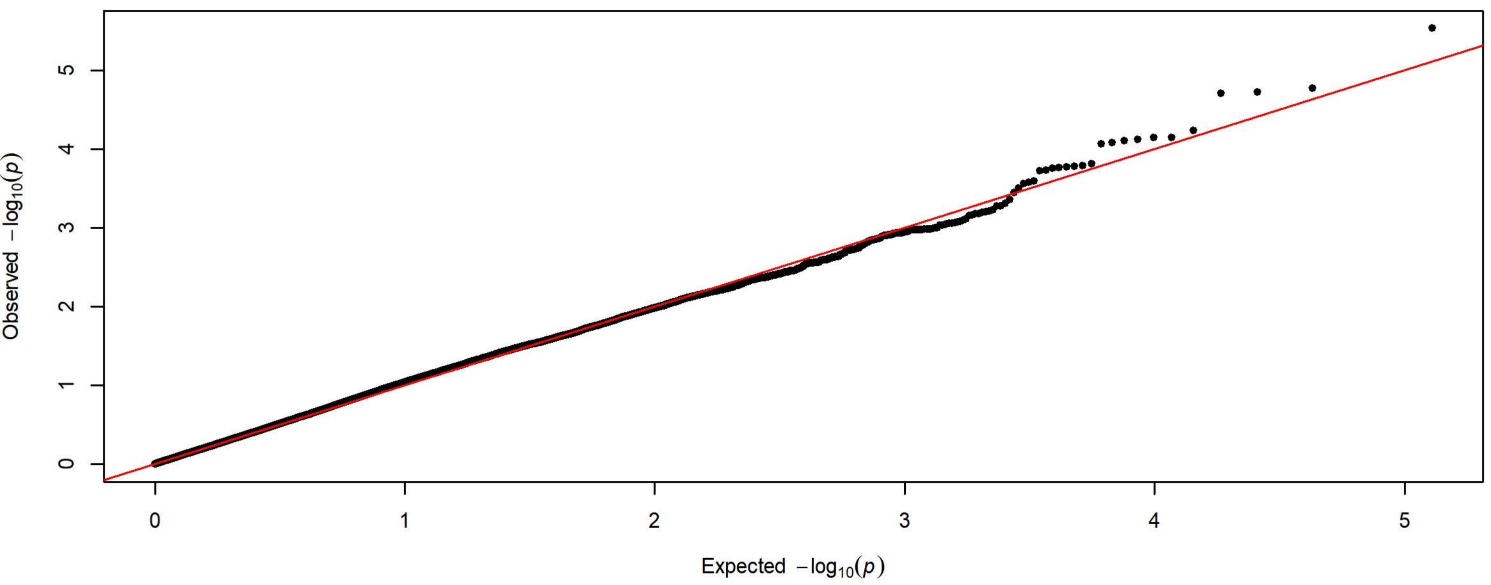


**Q-Q Plot Magazine Entries NCS Day 5 - Charles River 4 Subgroups - 198k SNPs (n=1728)**

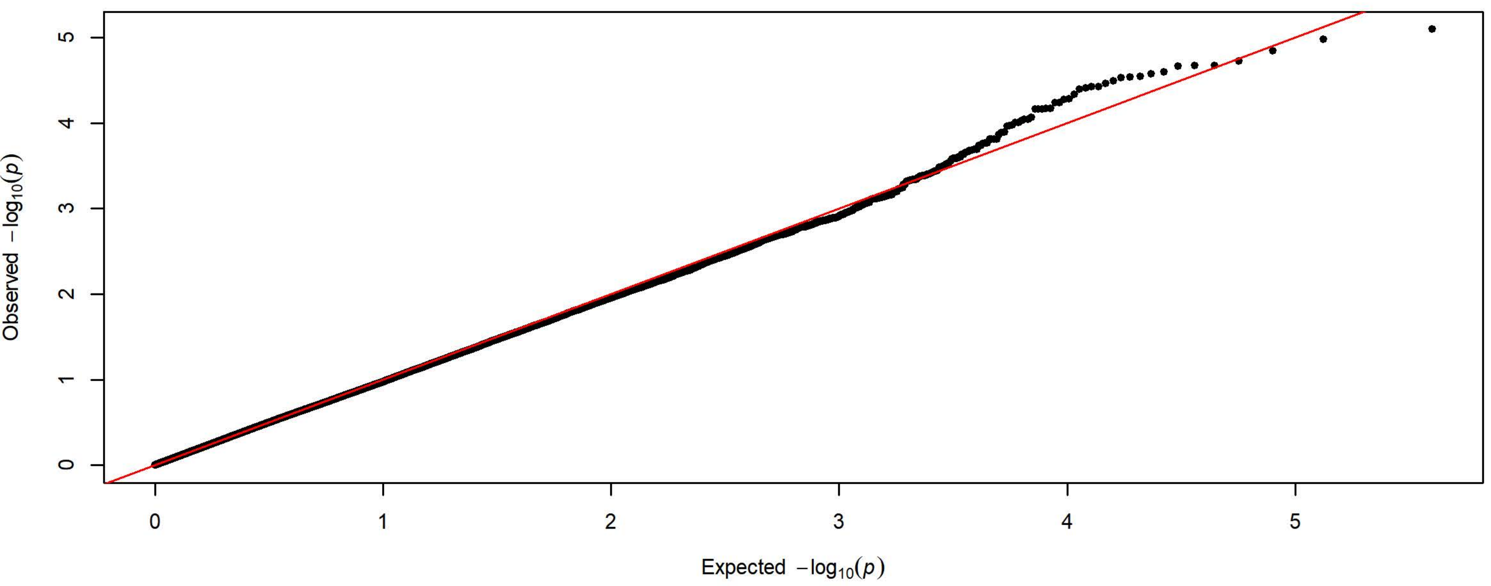


**Q-Q Plot Magazine Entries NCS Day 5 - Harlan 3 Subgroups - 83k SNPs (n=2208)**

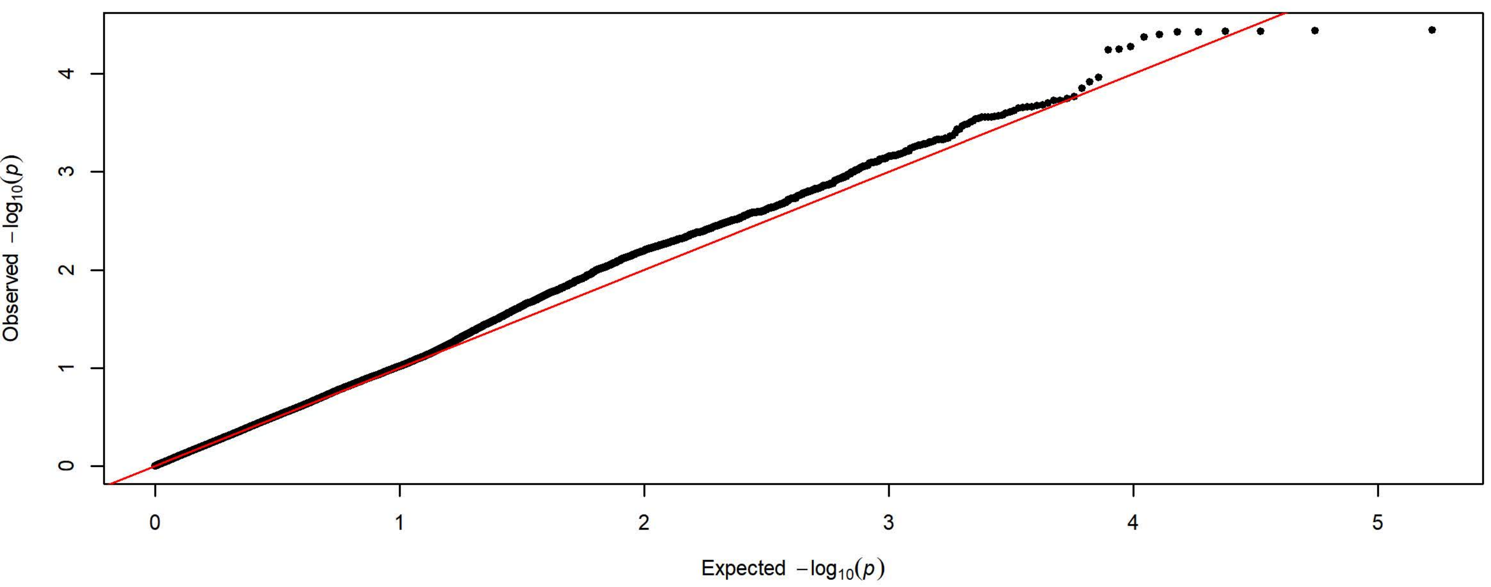

**Q-Q Plot Probability Difference Day 1 - Meta-analysis of 7 Subgroups - 64k SNPs (n=3903)**

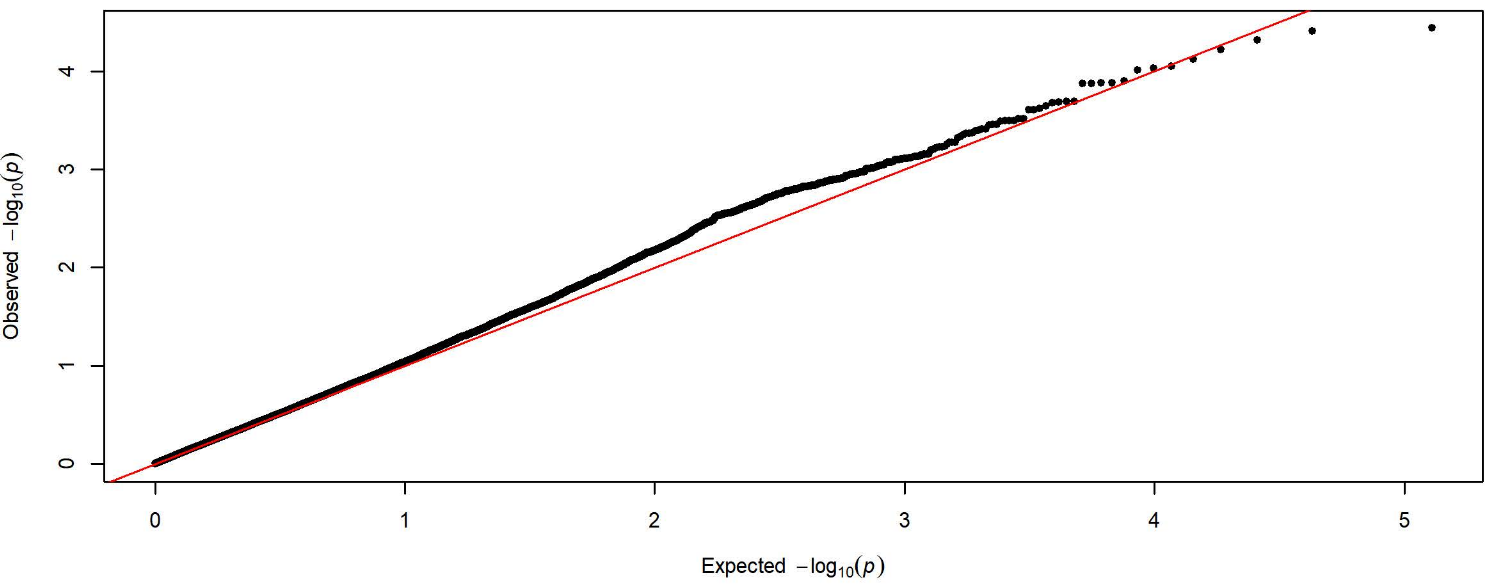


**Q-Q Plot Probability Difference Day 1 - Charles River 4 Subgroups - 198k SNPs (n=1728)**

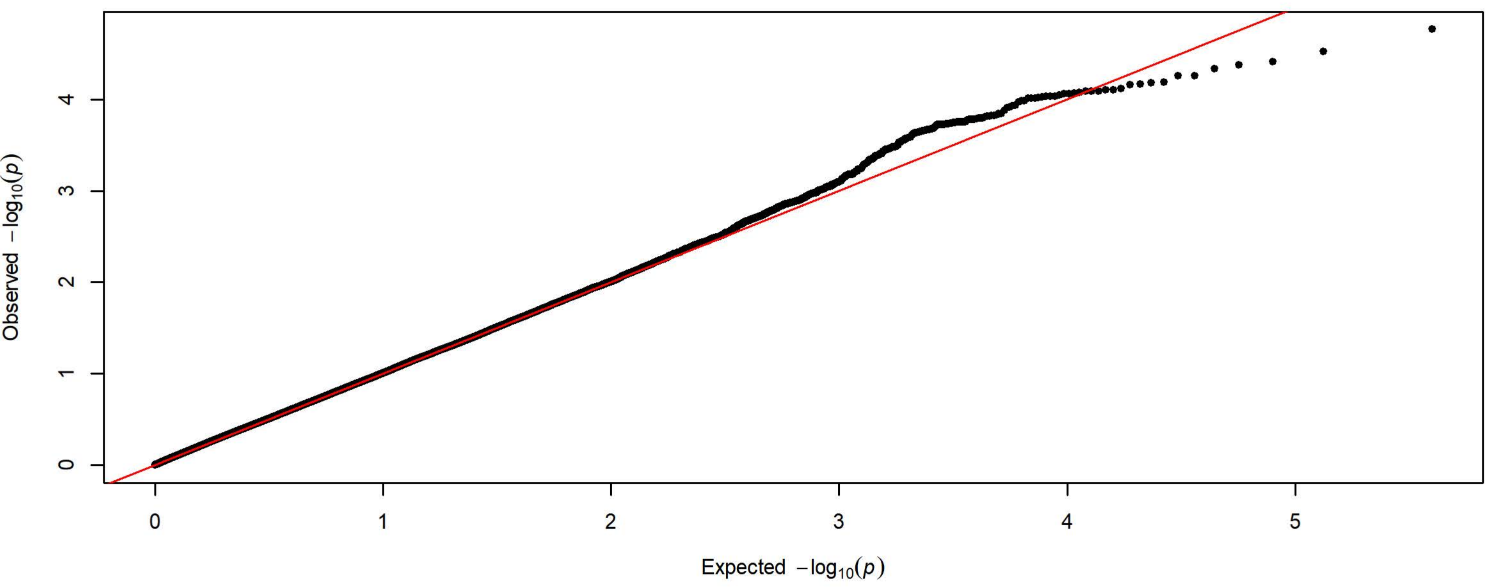


**Q-Q Plot Probability Difference Day 1 - Harlan 3 Subgroups - 83k SNPs (n=2175)**

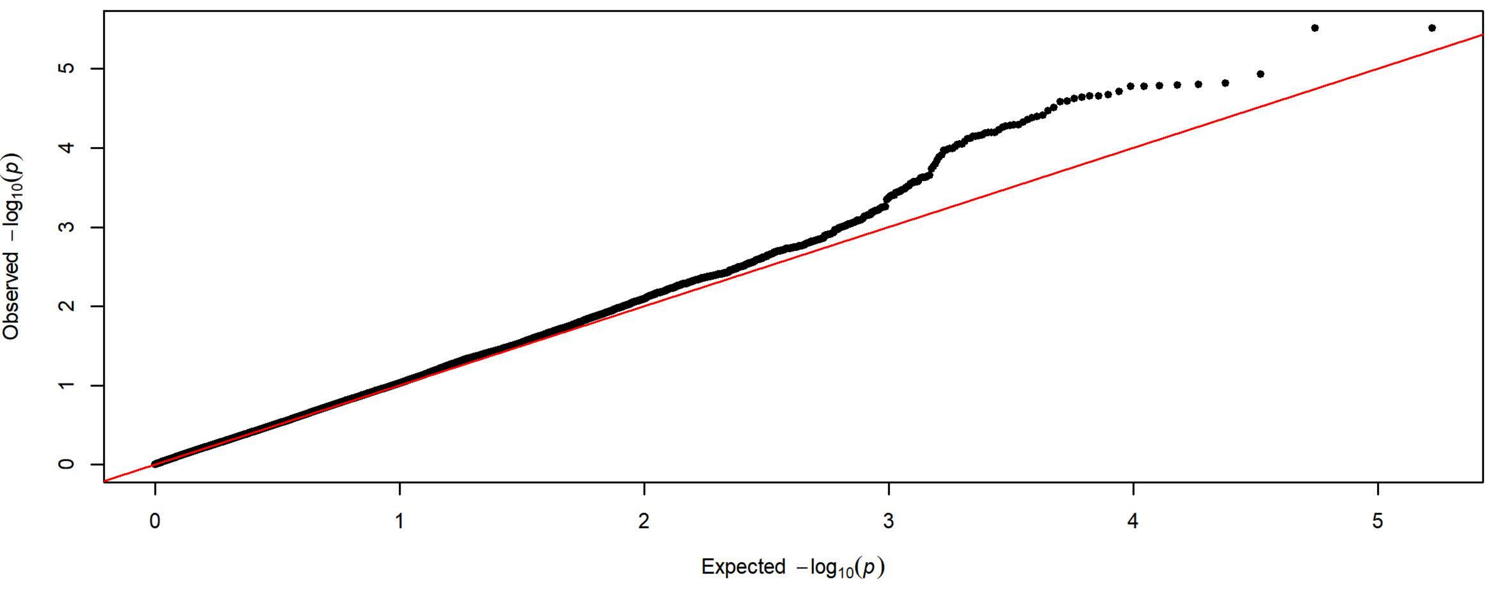

**Q-Q Plot Probability Difference Day 2 - Meta-analysis of 7 Subgroups - 64k SNPs (n=3934)**

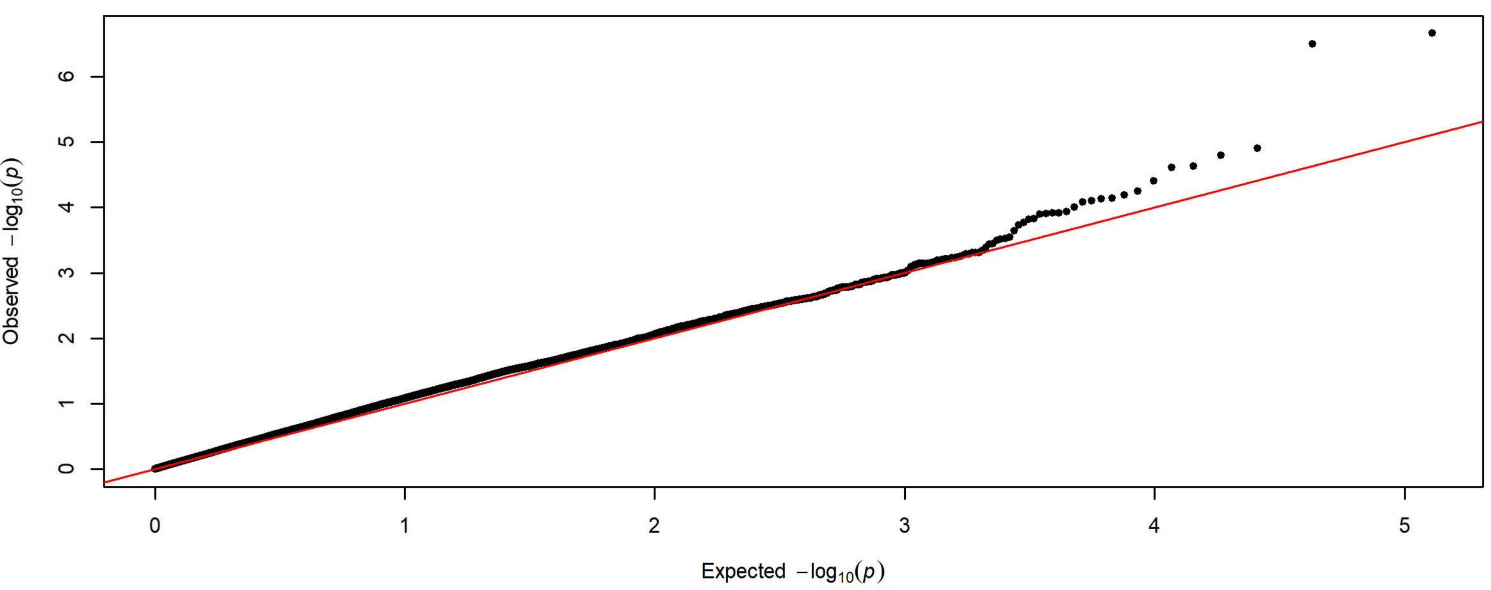


**Q-Q Plot Probability Difference Day 2 - Charles River 4 Subgroups - 198k SNPs (n=1726)**

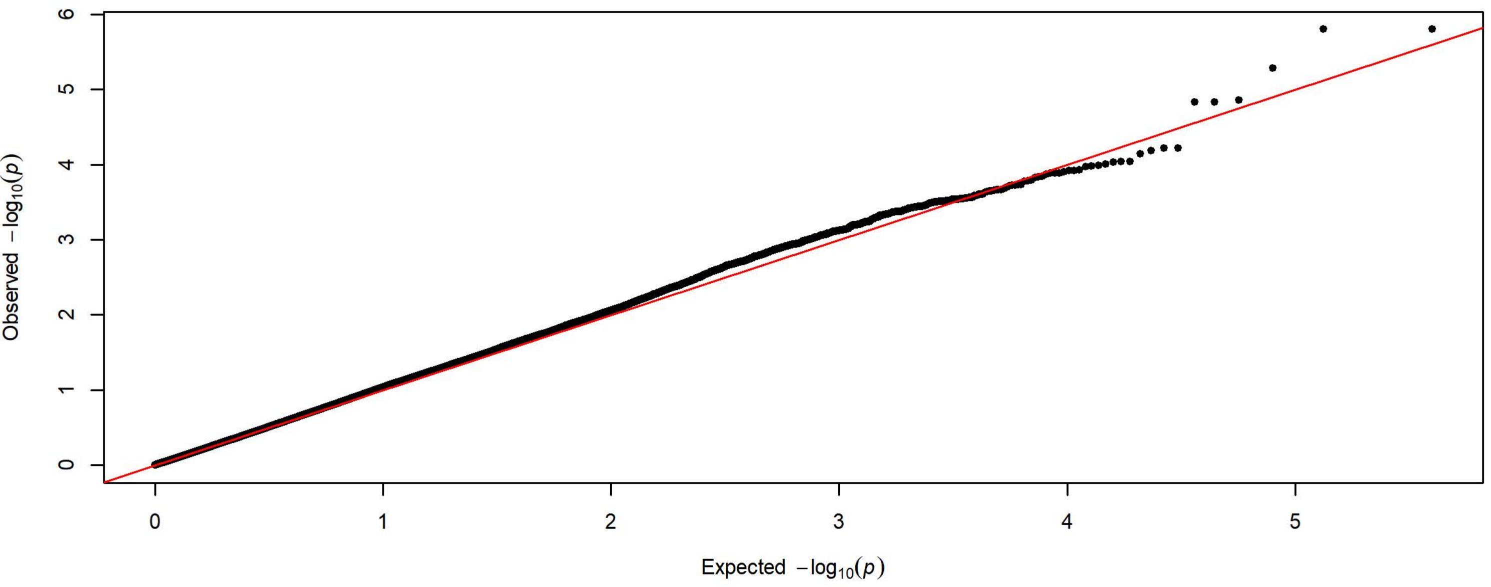


**Q-Q Plot Probability Difference Day 2 - Harlan 3 Subgroups - 83k SNPs (n=2208)**

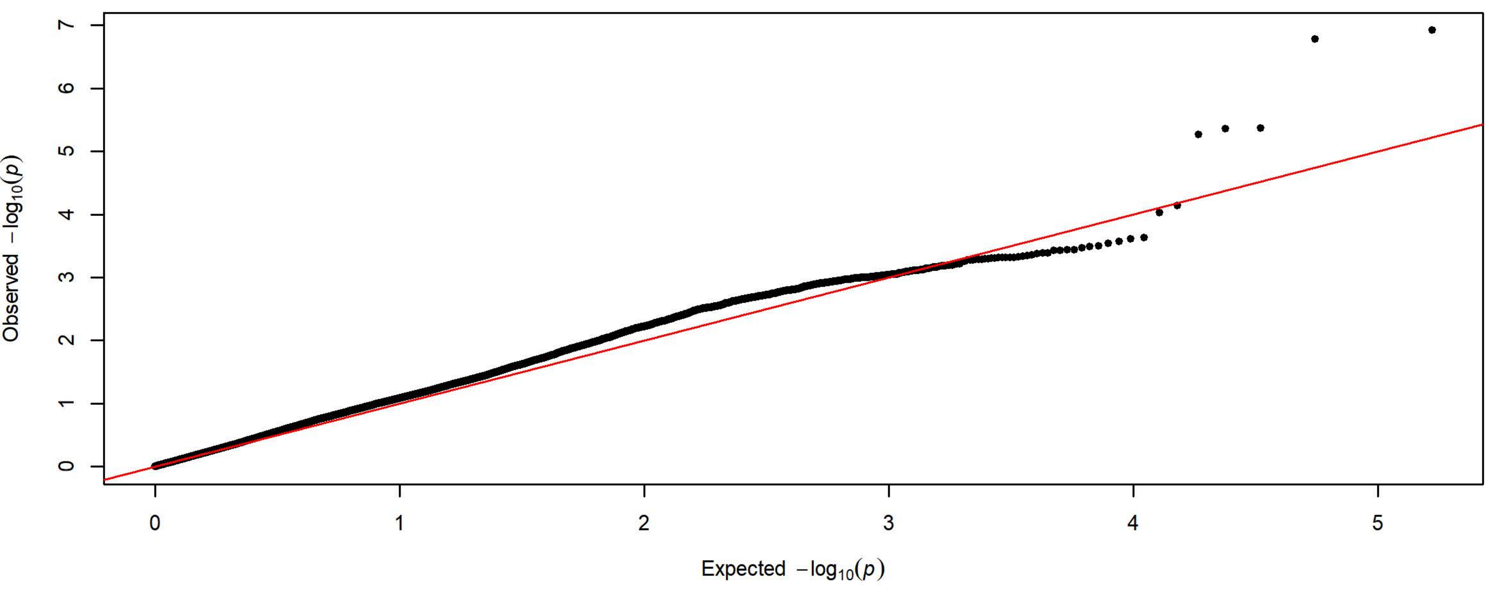

**Q-Q Plot Probability Difference Day 3 - Meta-analysis of 7 Subgroups - 64k SNPs (n=3932)**

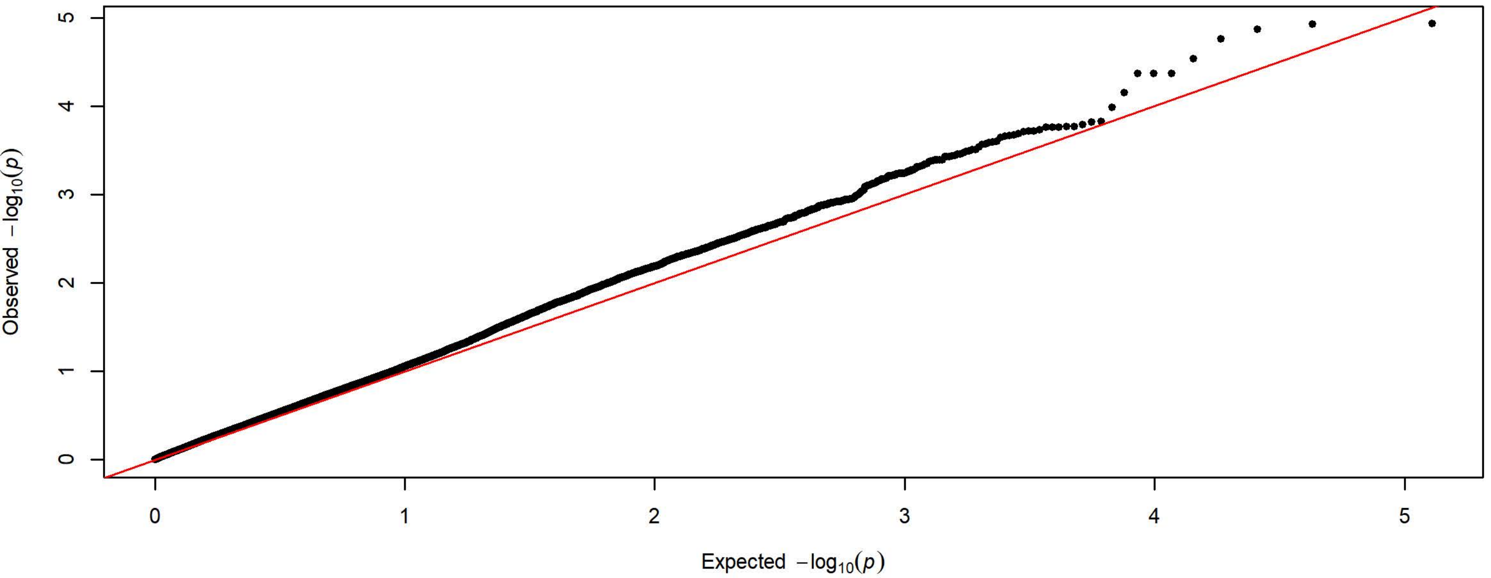


**Q-Q Plot Probability Difference Day 3 - Charles River 4 Subgroups - 198k SNPs (n=1727)**

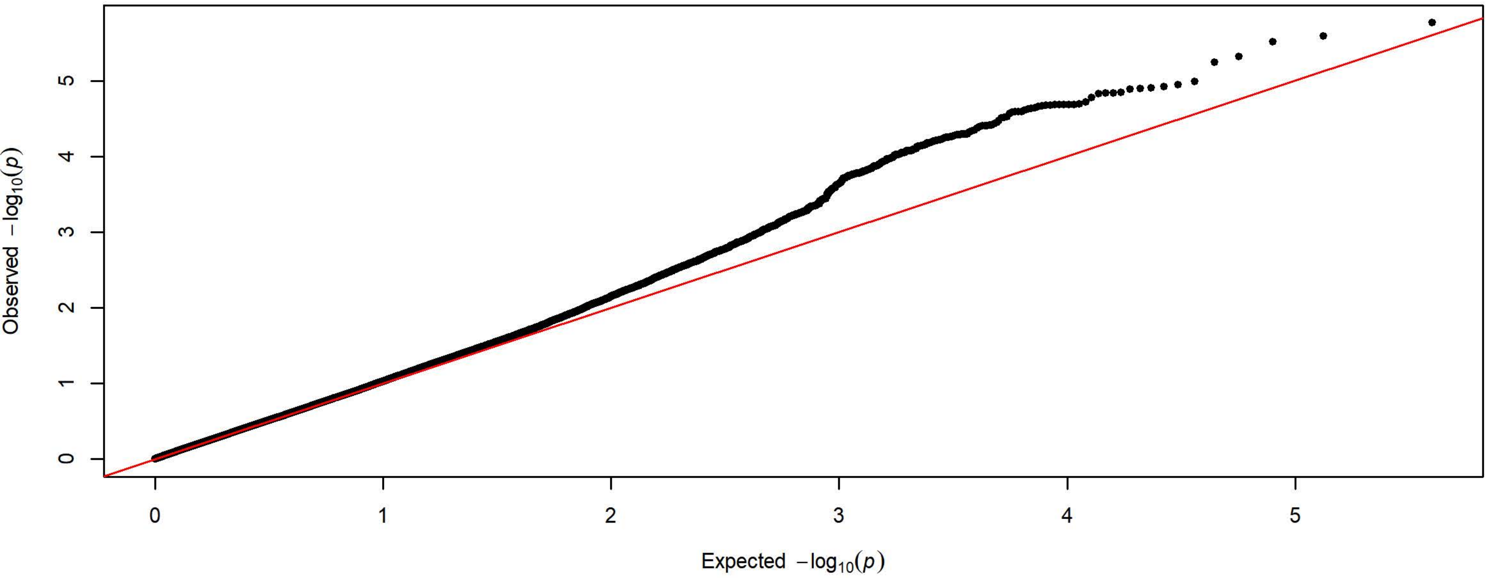


**Q-Q Plot Probability Difference Day 3 - Harlan 3 Subgroups - 83k SNPs (n=2205)**

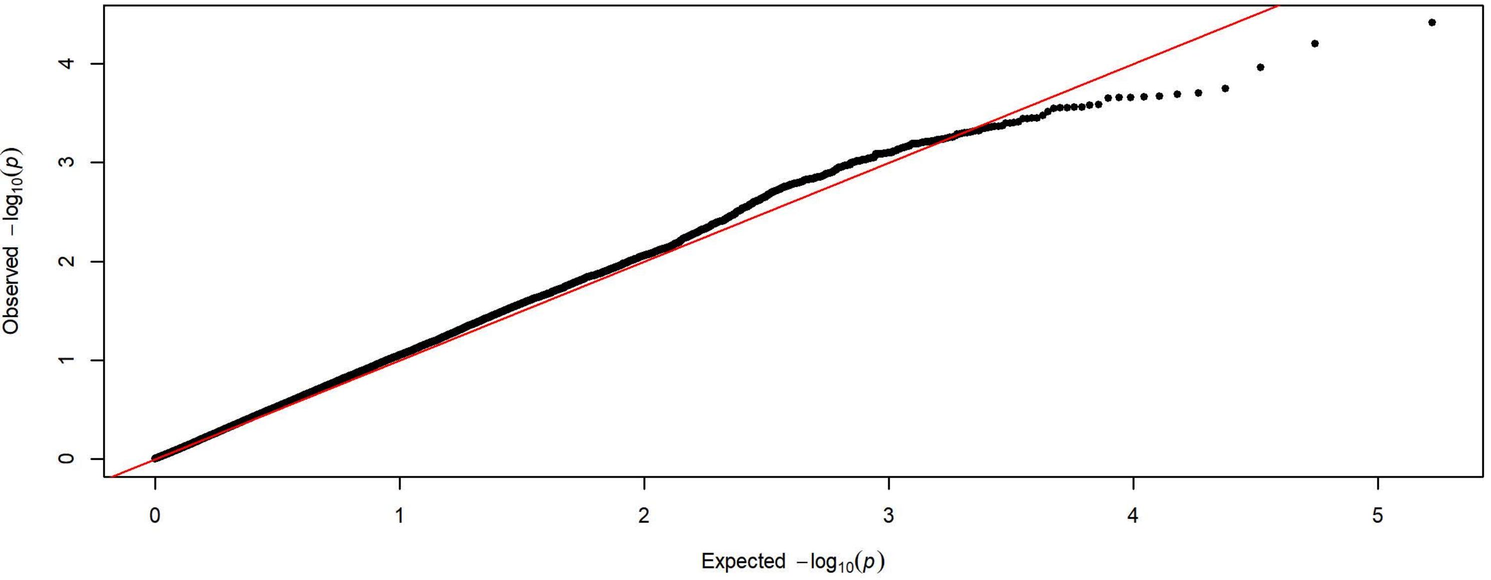

**Q-Q Plot Probability Difference Day 4 - Meta-analysis of 7 Subgroups - 64k SNPs (n=3936)**

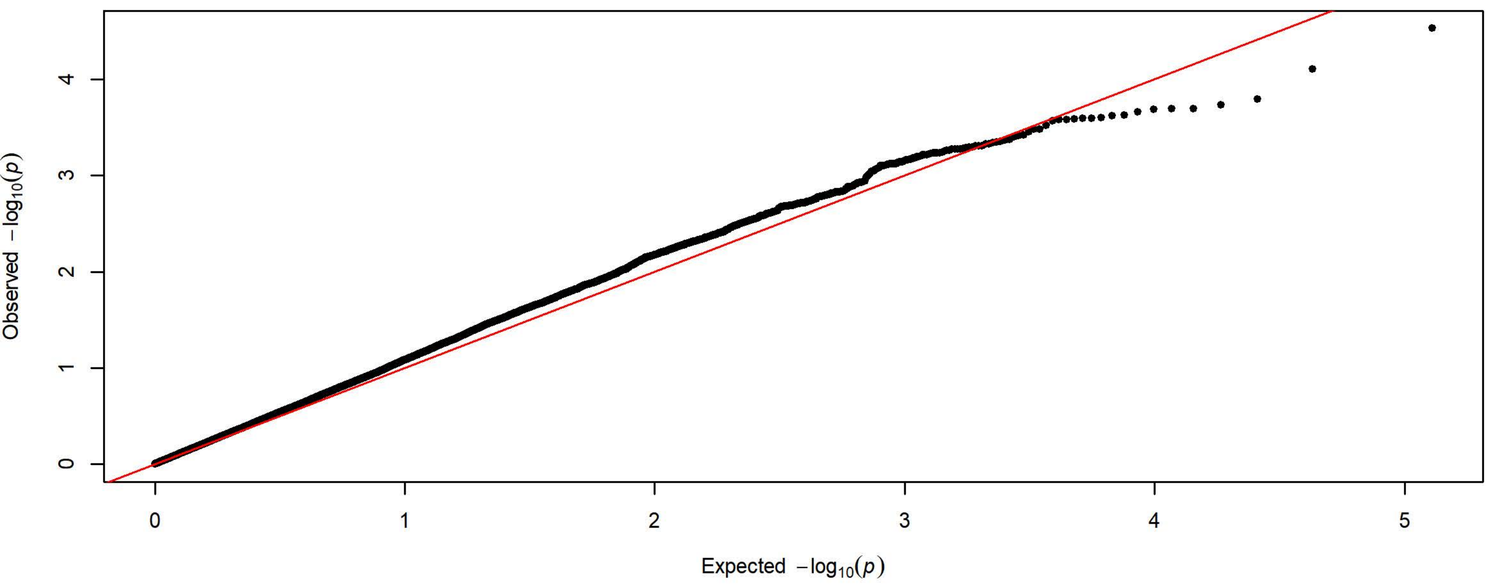


**Q-Q Plot Probability Difference Day 4 - Charles River 4 Subgroups - 198k SNPs (n=1728)**

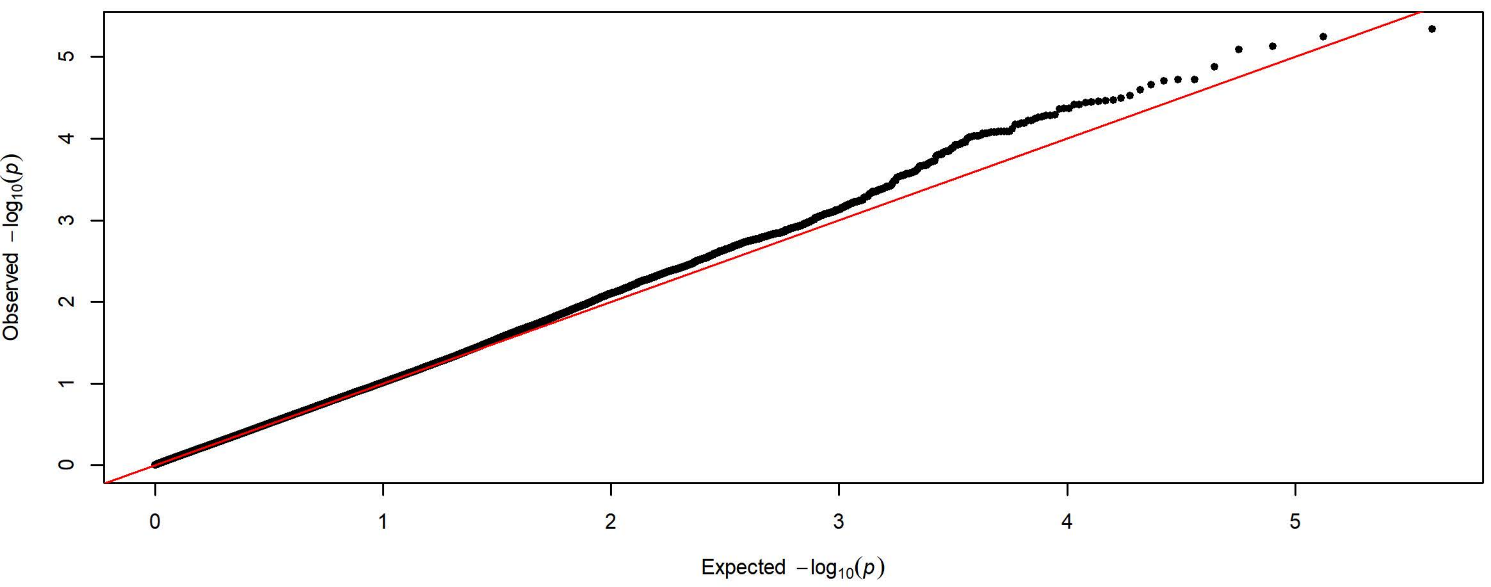


**Q-Q Plot Probability Difference Day 4 - Harlan 3 Subgroups - 83k SNPs (n=2208)**

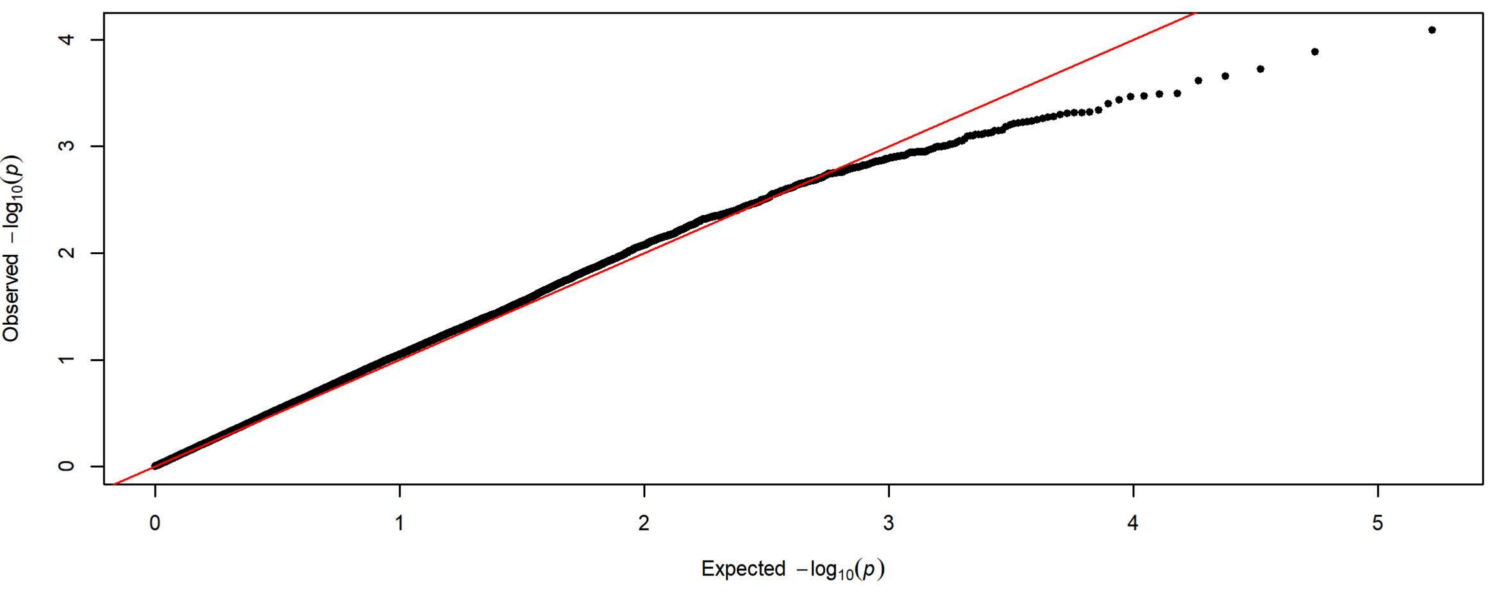

**Q-Q Plot Probability Difference Day 5 - Meta-analysis of 7 Subgroups - 64k SNPs (n=3936)**

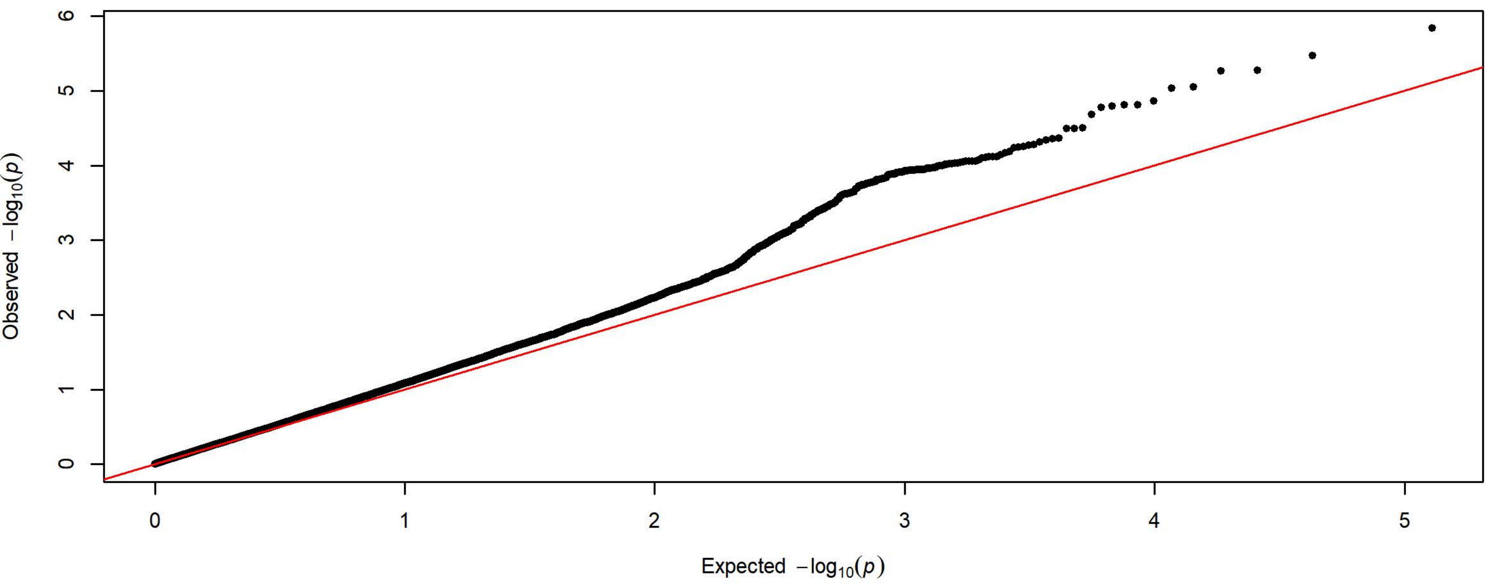


**Q-Q Plot Probability Difference Day 5 - Charles River 4 Subgroups - 198k SNPs (n=1728)**

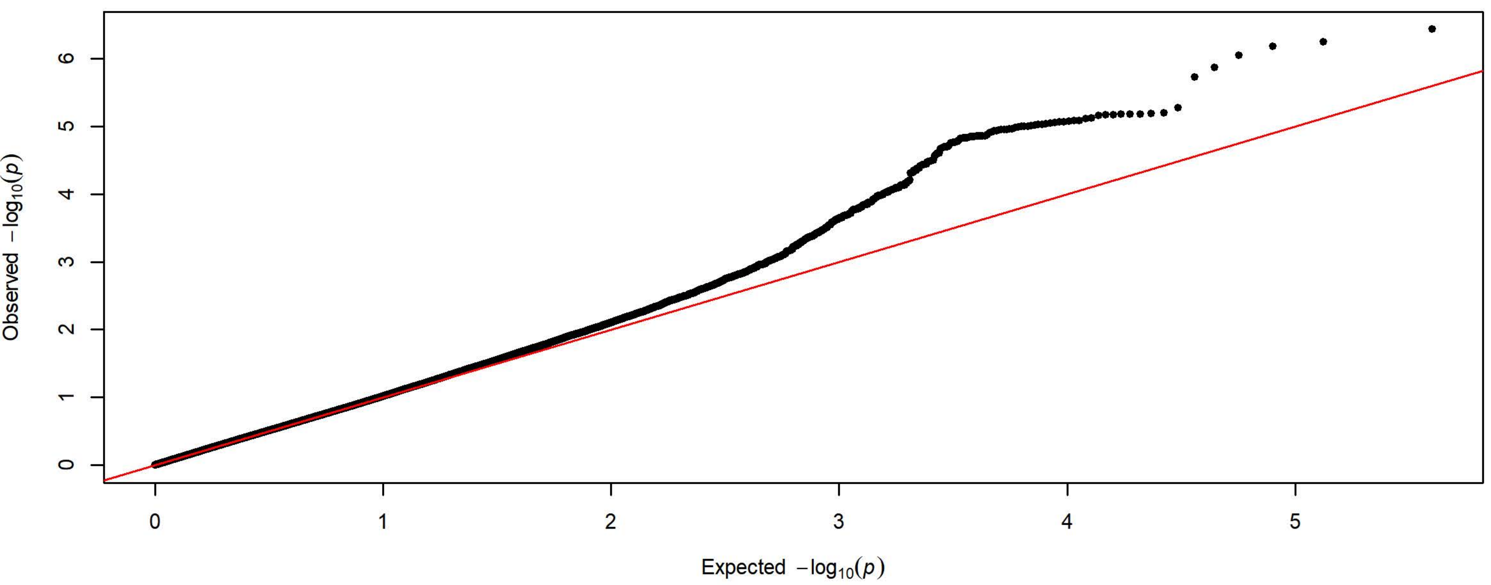


**Q-Q Plot Probability Difference Day 5 - Harlan 3 Subgroups - 83k SNPs (n=2208)**

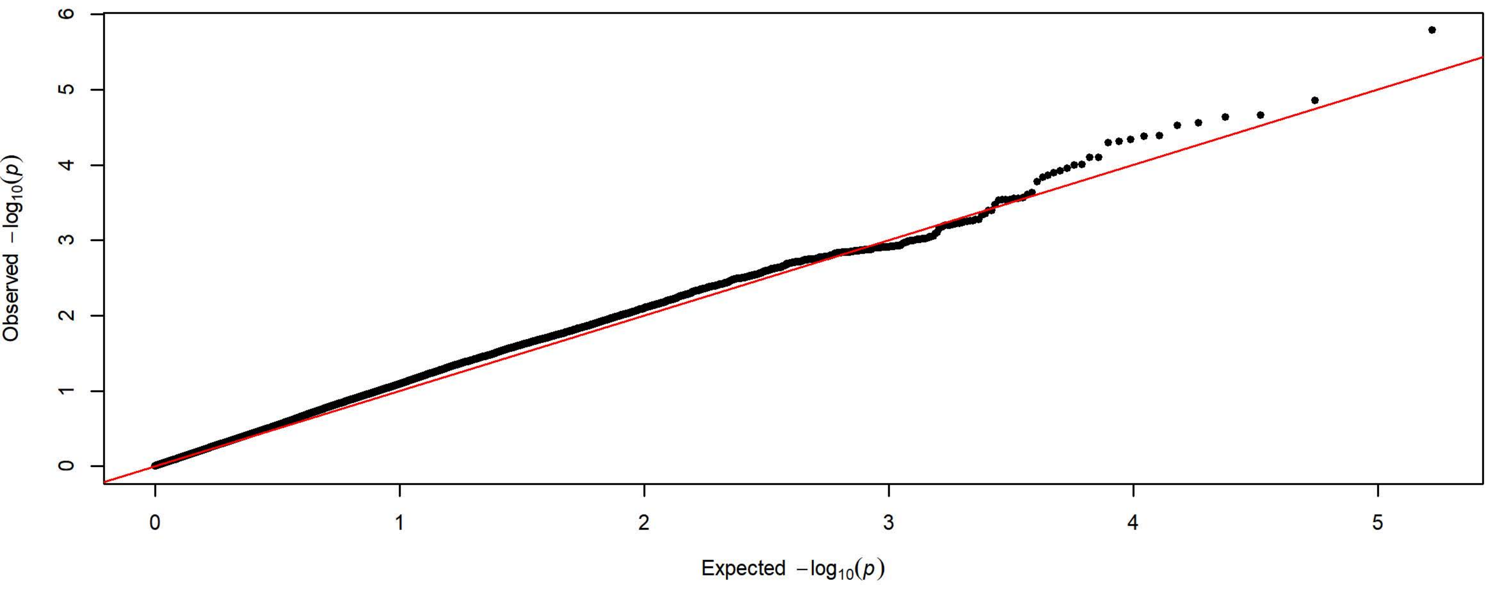

**Q-Q Plot Probability of Lever Press Day 1 - Meta-analysis of 7 Subgroups - 64k SNPs (n=3903)**

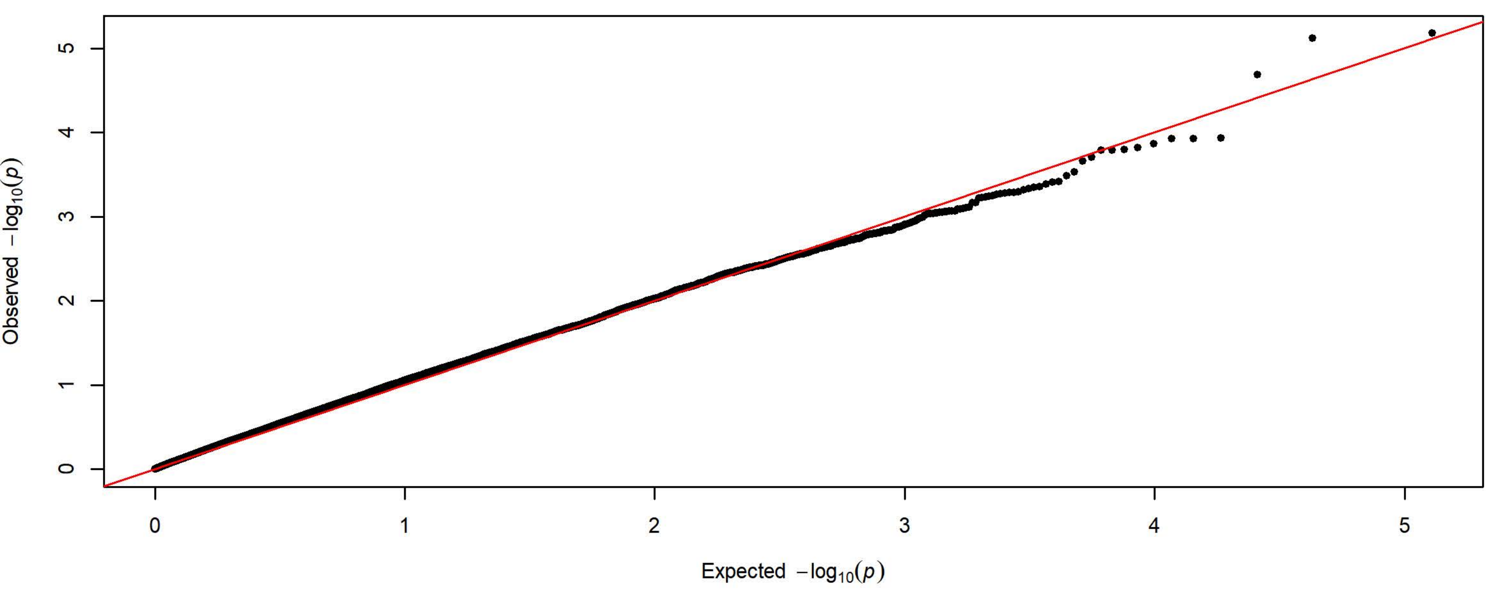


**Q-Q Plot Probability of Lever Press Day 1 - Charles River 4 Subgroups - 198k SNPs (n=1728)**

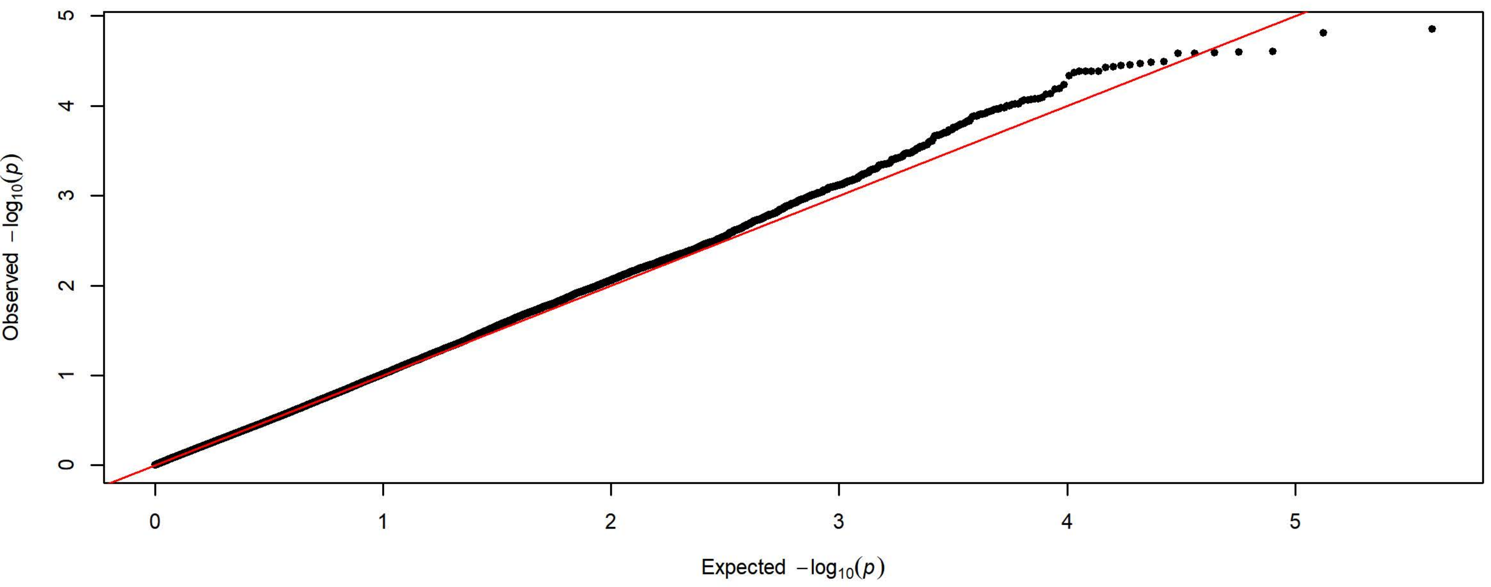


**Q-Q Plot Probability of Lever Press Day 1 - Harlan 3 Subgroups - 83k SNPs (n=2175)**

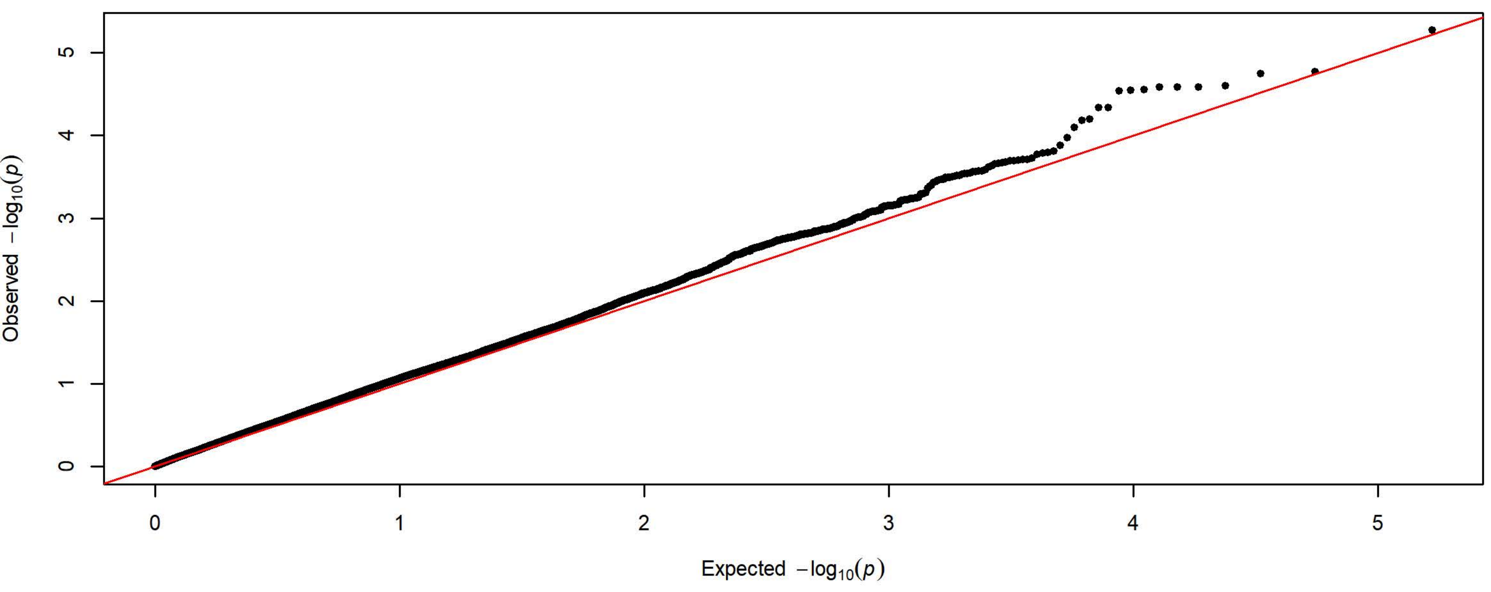

**Q-Q Plot Probability of Lever Press Day 2 - Meta-analysis of 7 Subgroups - 64k SNPs (n=3934)**

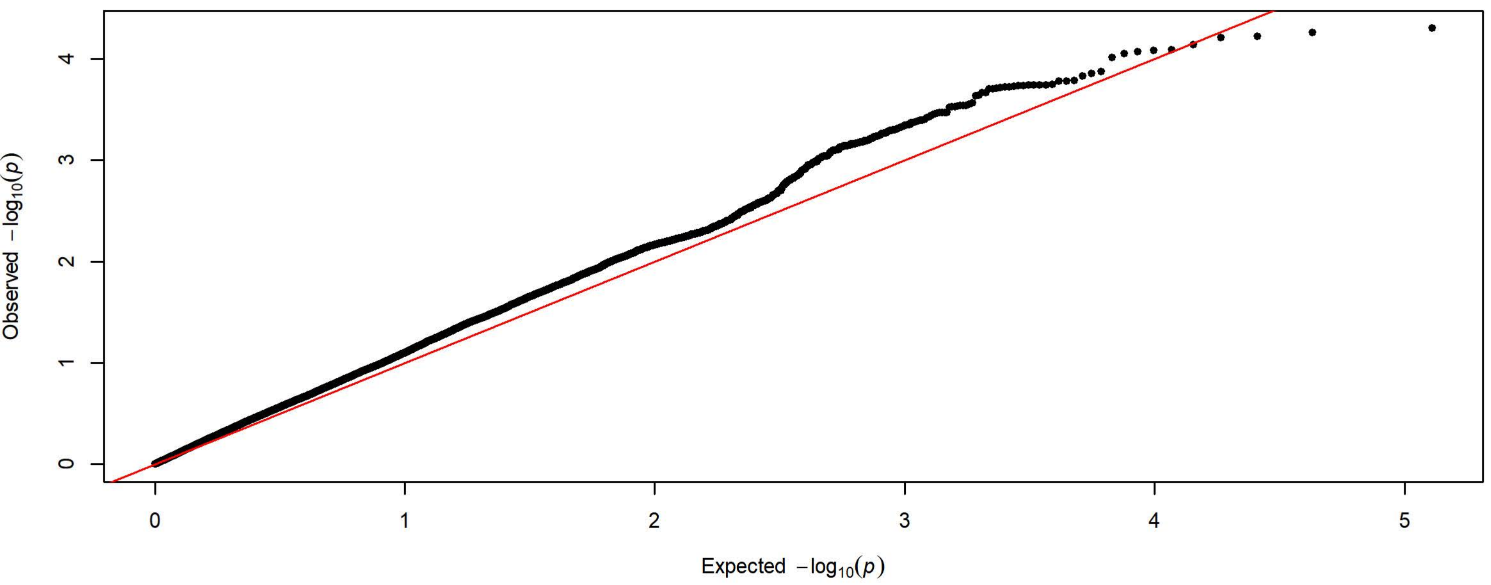


**Q-Q Plot Probability of Lever Press Day 2 - Charles River 4 Subgroups - 198k SNPs (n=1726)**

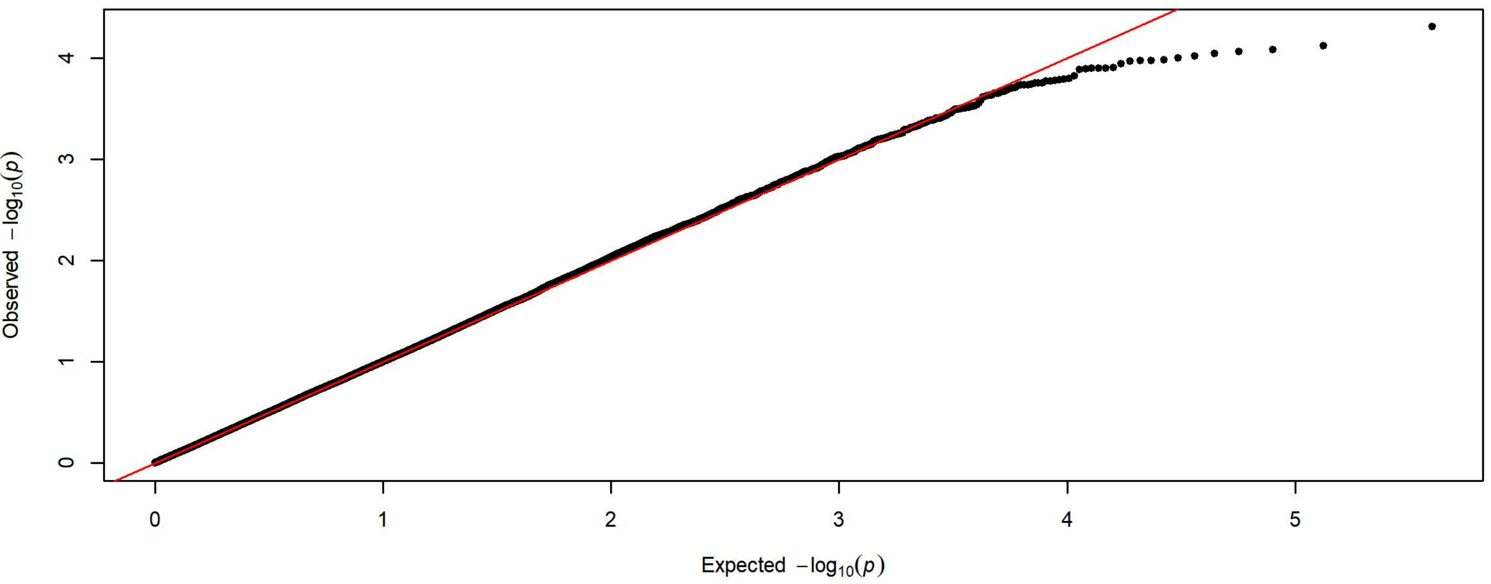


**Q-Q Plot Probability of Lever Press Day 2 - Harlan 3 Subgroups - 83k SNPs (n=2208)**

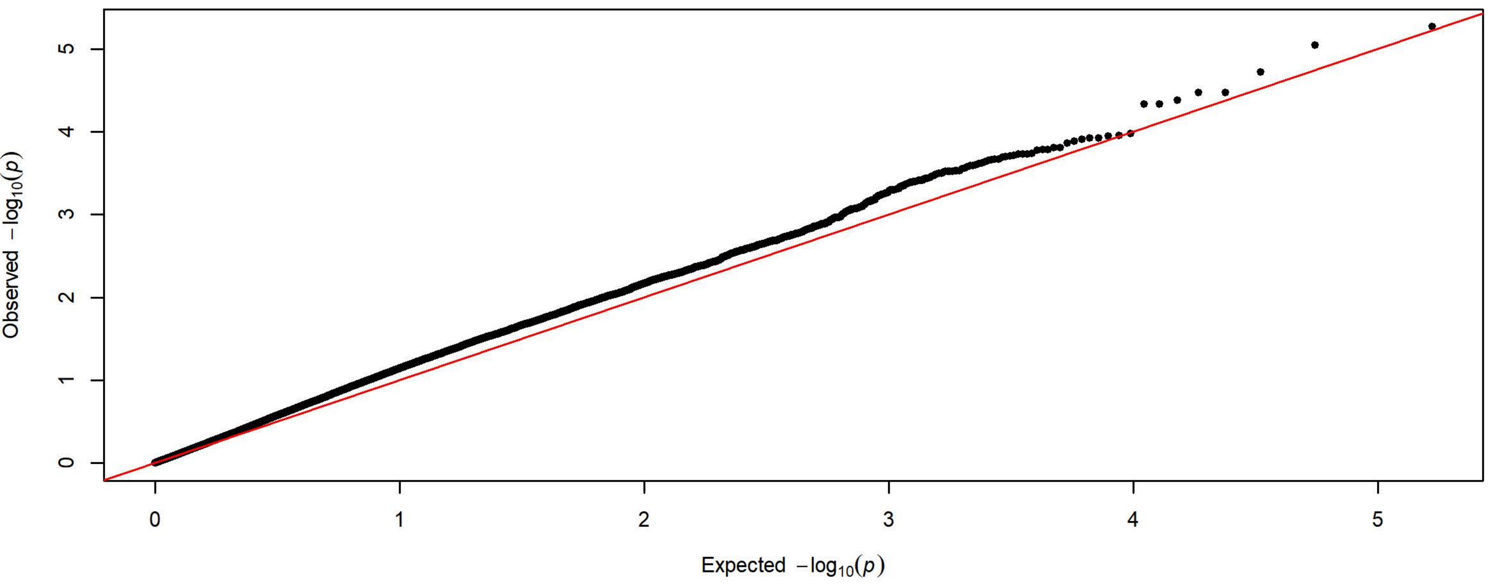

**Q-Q Plot Probability of Lever Press Day 3 - Meta-analysis of 7 Subgroups - 64k SNPs (n=3932)**

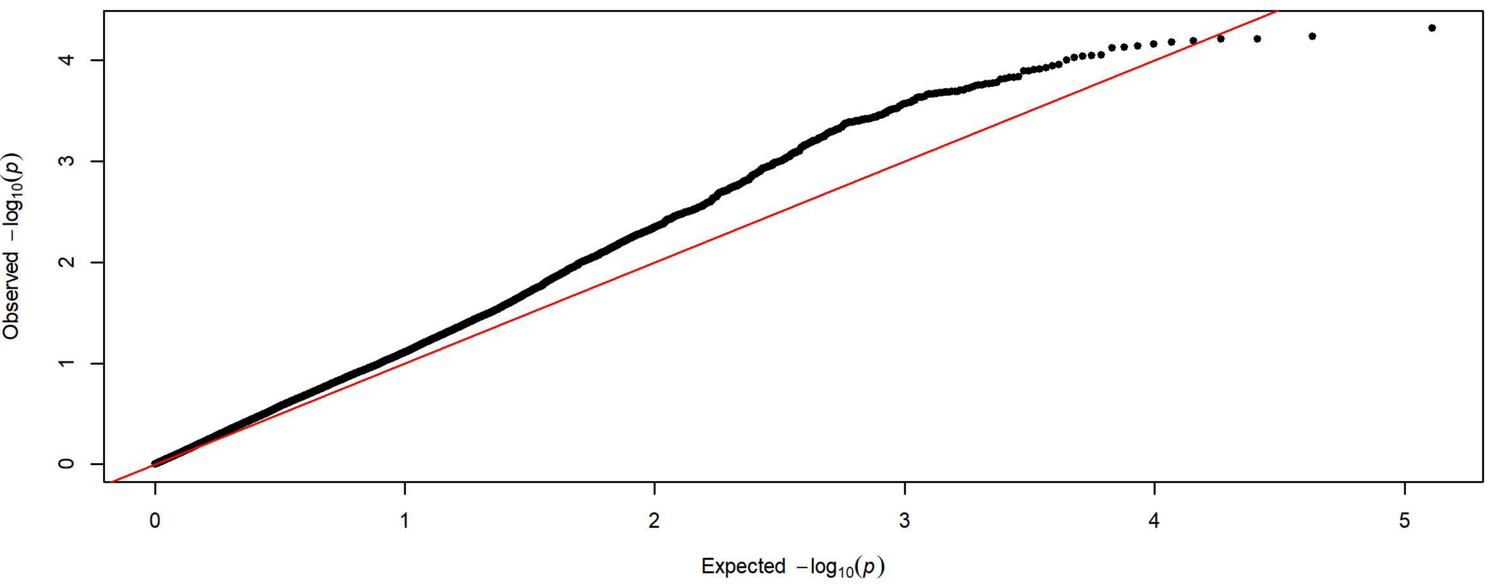


**Q-Q Plot Probability of Lever Press Day 3 - Charles River 4 Subgroups - 198k SNPs (n=1727)**

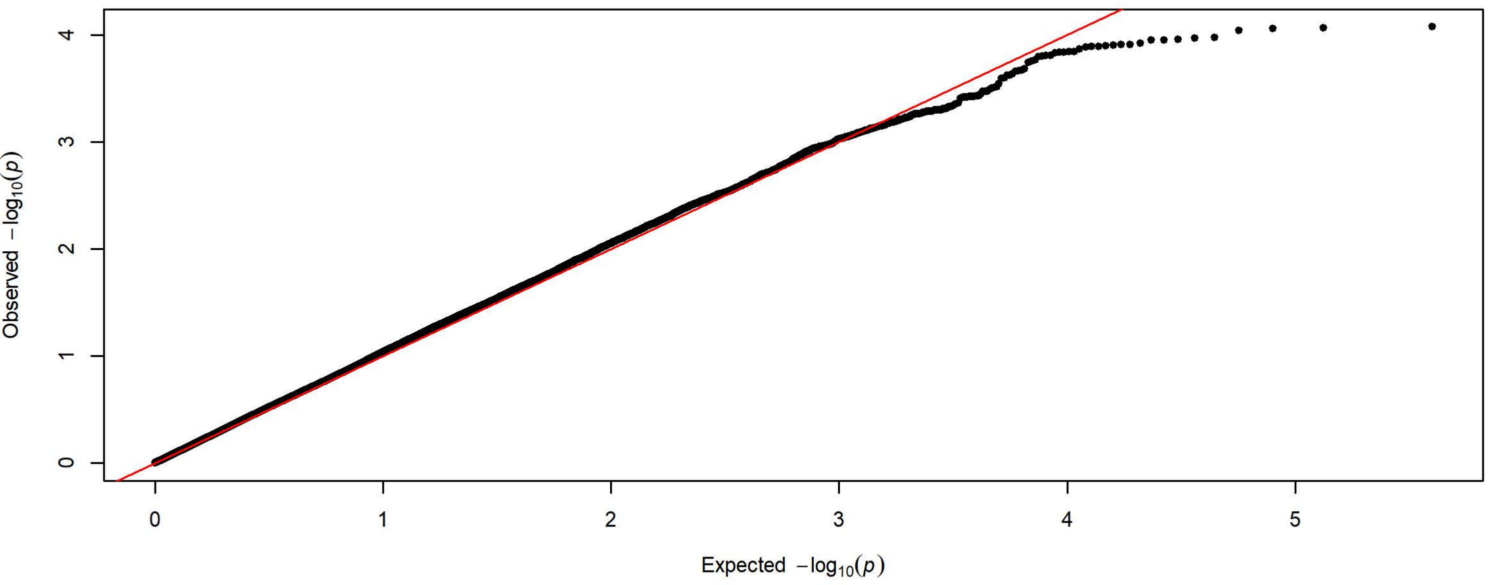


**Q-Q Plot Probability of Lever Press Day 3 - Harlan 3 Subgroups - 83k SNPs (n=2205)**

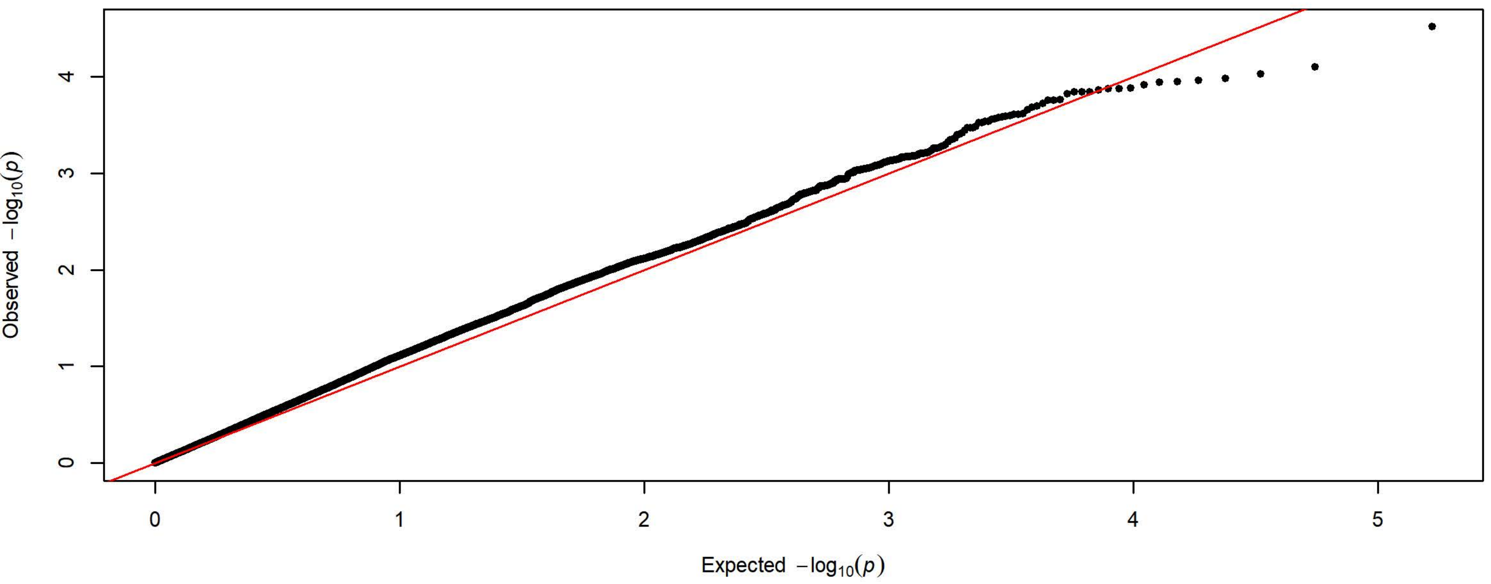

**Q-Q Plot Probability of Lever Press Day 4 - Meta-analysis of 7 Subgroups - 64k SNPs (n=3936)**

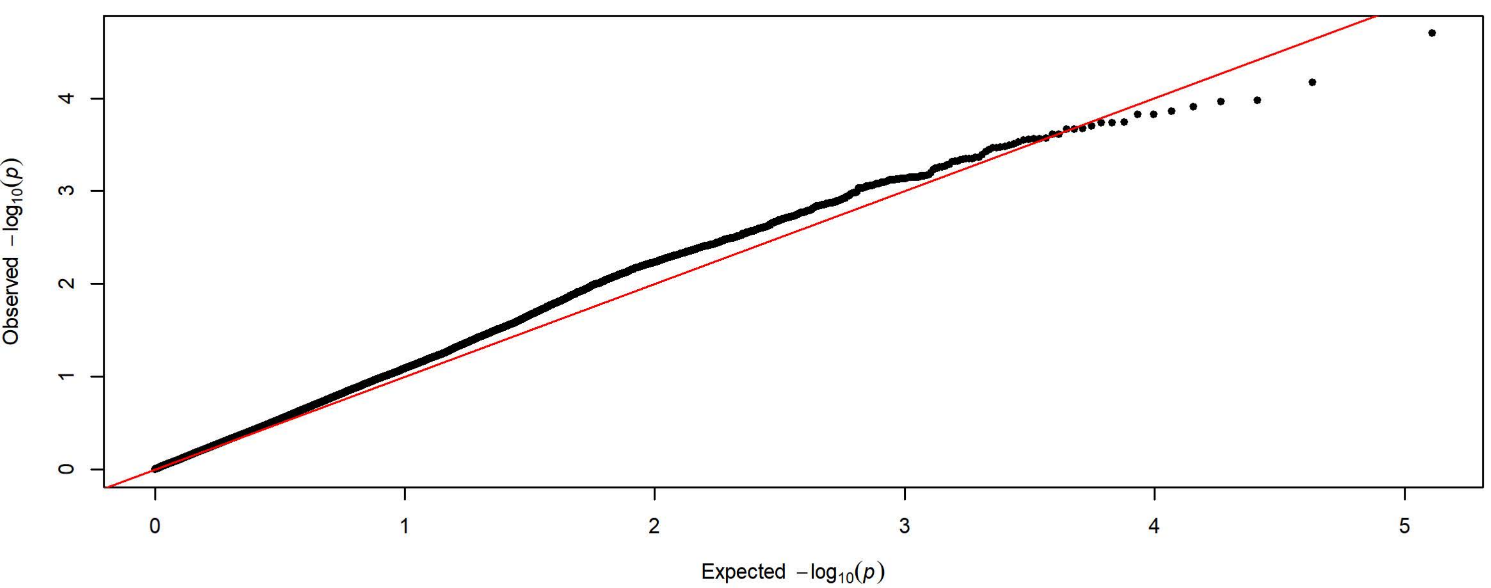


**Q-Q Plot Probability of Lever Press Day 4 - Charles River 4 Subgroups - 198k SNPs (n=1728)**

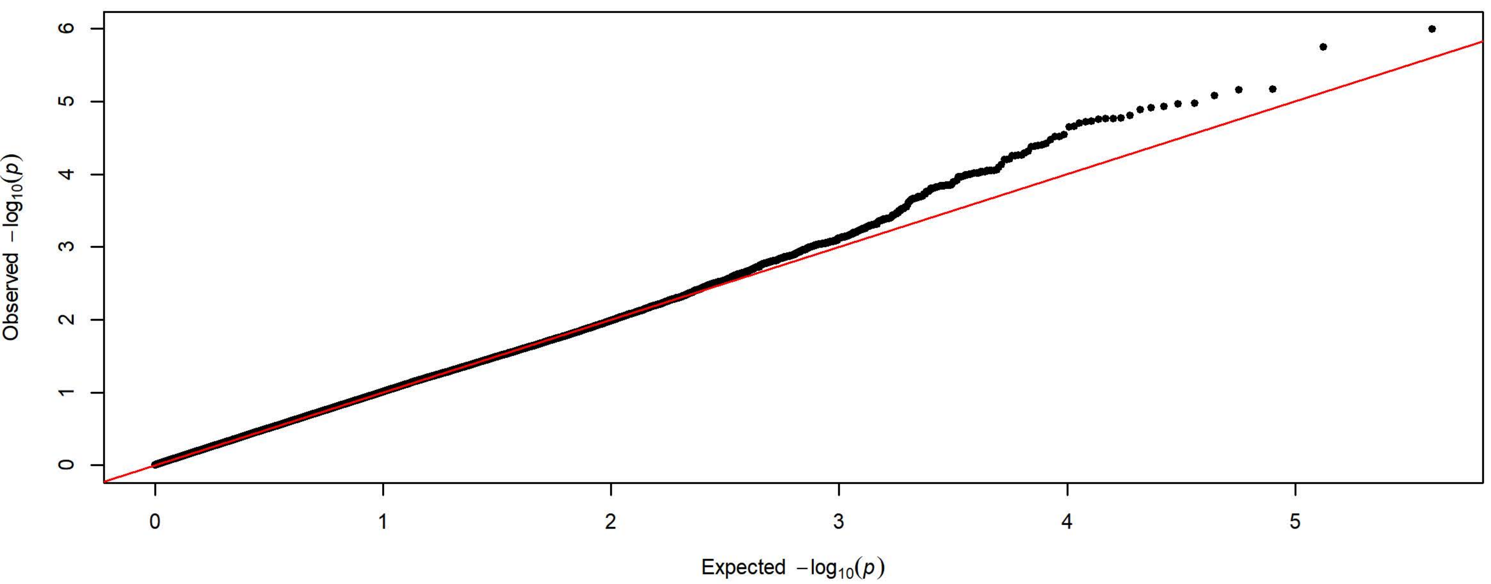


**Q-Q Plot Probability of Lever Press Day 4 - Harlan 3 Subgroups - 83k SNPs (n=2208)**

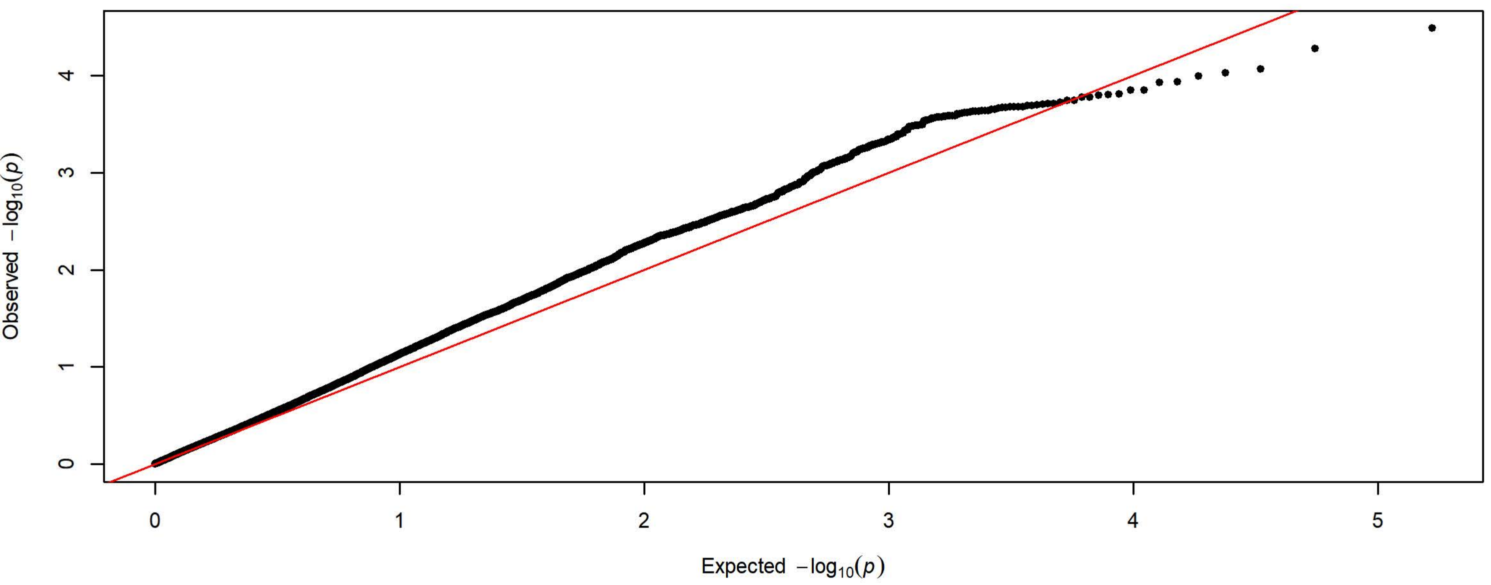

**Q-Q Plot Probability of Lever Press Day 5 - Meta-analysis of 7 Subgroups - 64k SNPs (n=3936)**

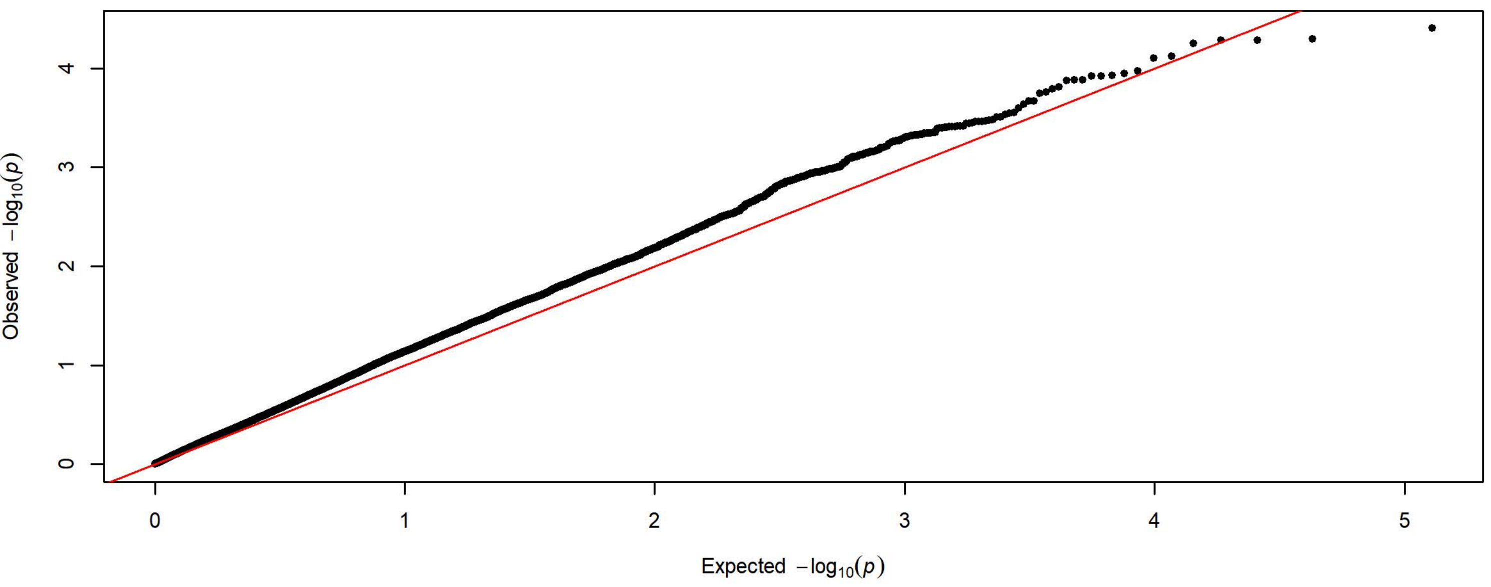


**Q-Q Plot Probability of Lever Press Day 5 - Charles River 4 Subgroups - 198k SNPs (n=1728)**

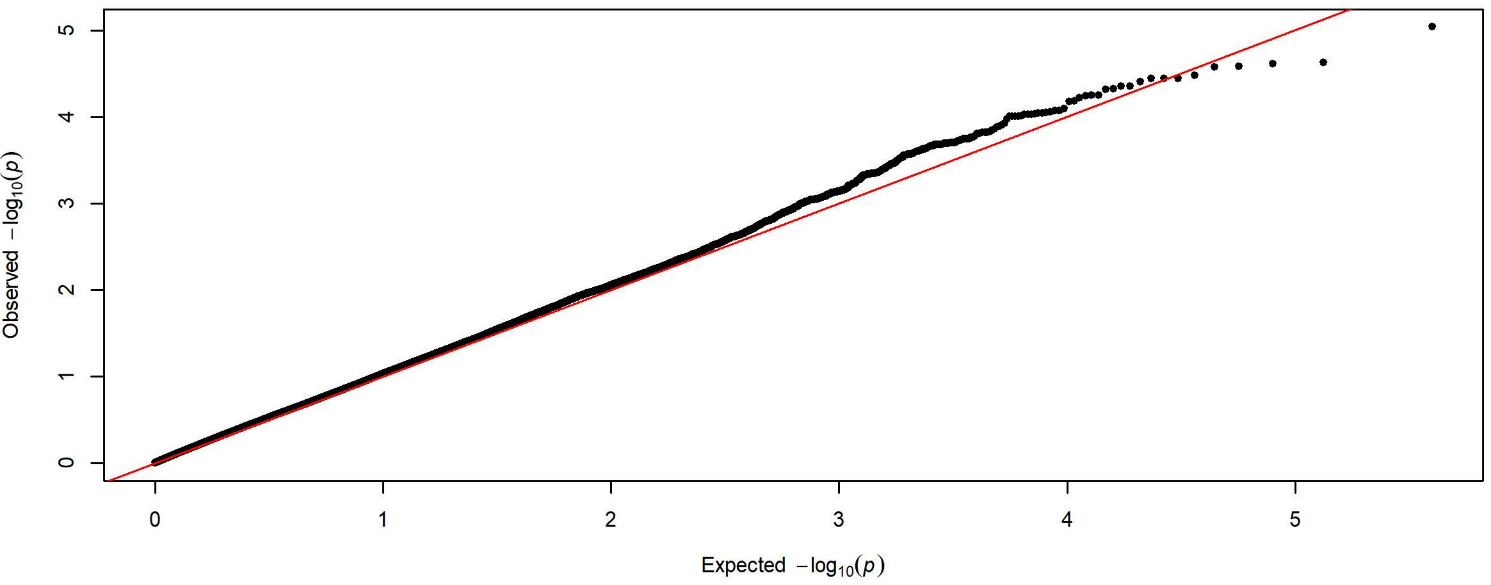


**Q-Q Plot Probability of Lever Press Day 5 - Harlan 3 Subgroups - 83k SNPs (n=2208)**

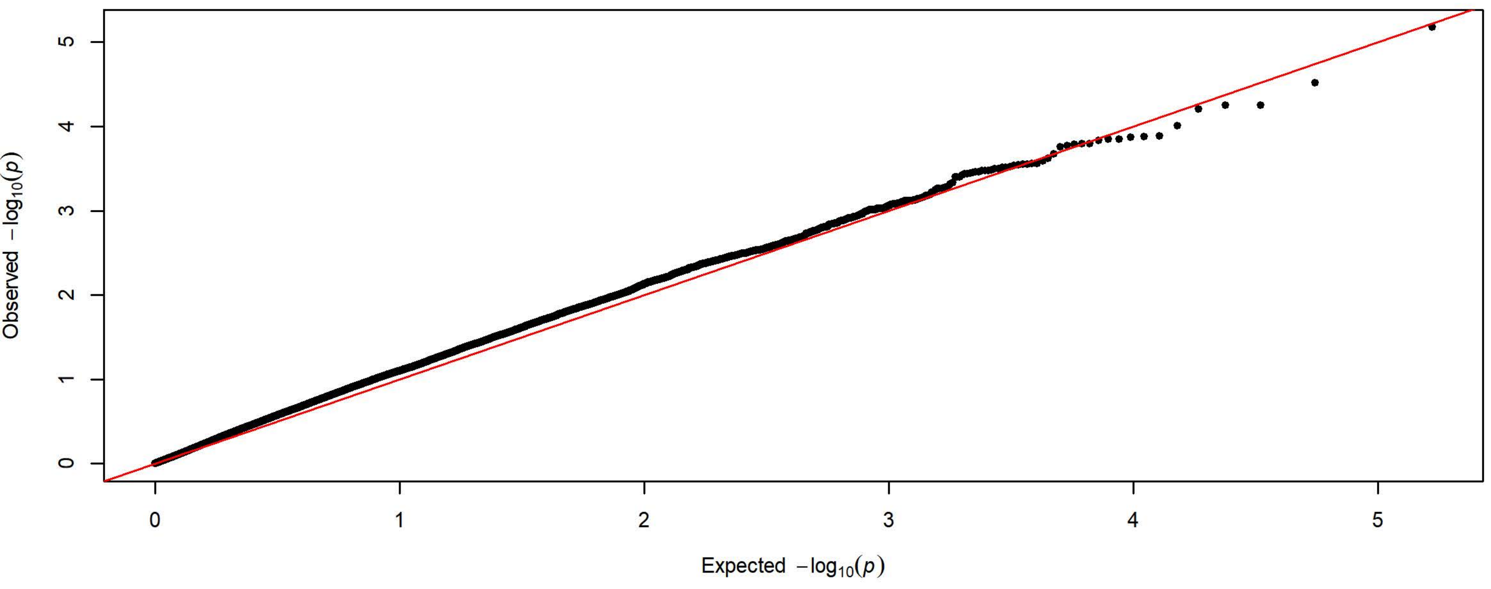

**Q-Q Plot Probability of Magazine Entry Day 1 - Meta-analysis of 7 Subgroups - 64k SNPs (n=3903)**

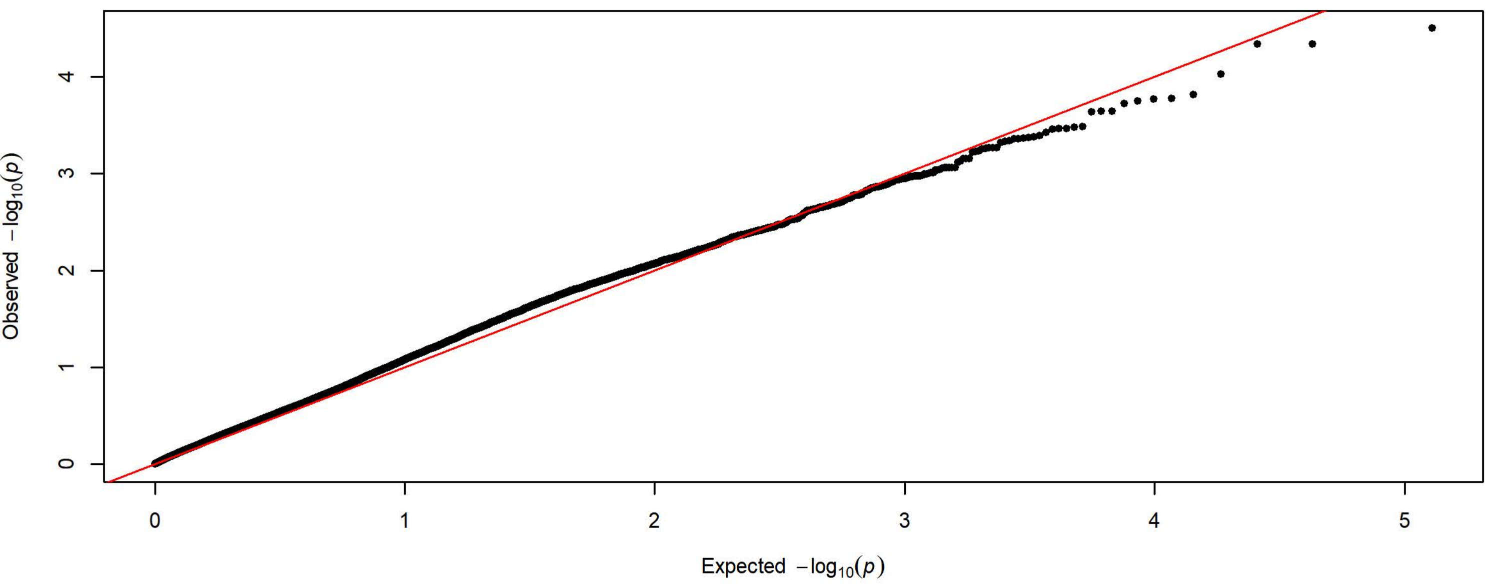


**Q-Q Plot Probability of Magazine Entry Day 1 - Charles River 4 Subgroups - 198k SNPs (n=1728)**

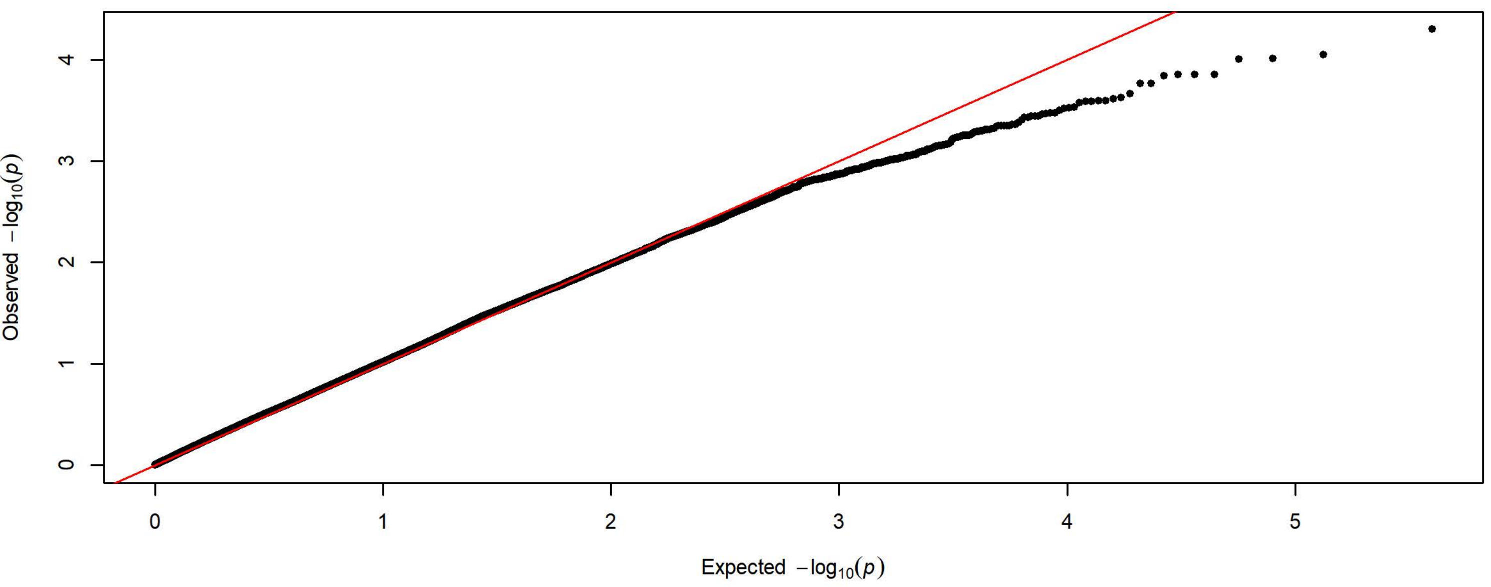


**Q-Q Plot Probability of Magazine Entry Day 1 - Harlan 3 Subgroups - 83k SNPs (n=2175)**

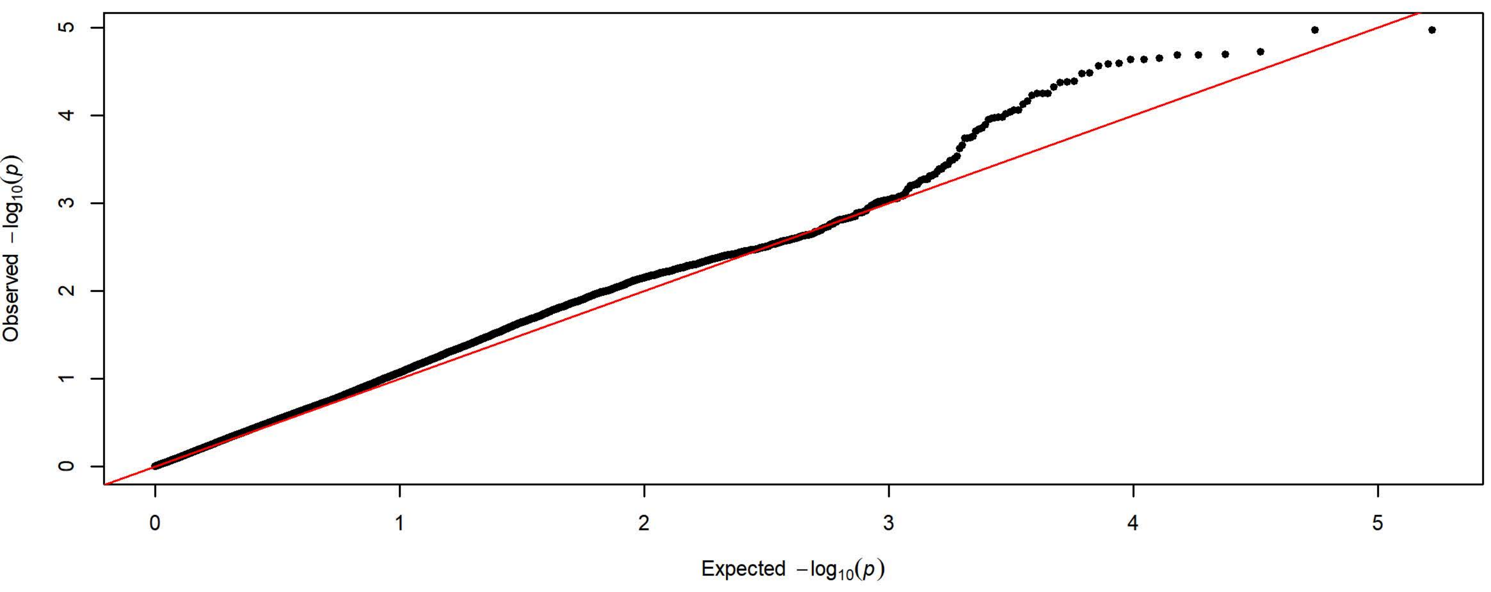

**Q-Q Plot Probability of Magazine Entry Day 2 - Meta-analysis of 7 Subgroups - 64k SNPs (n=3934)**

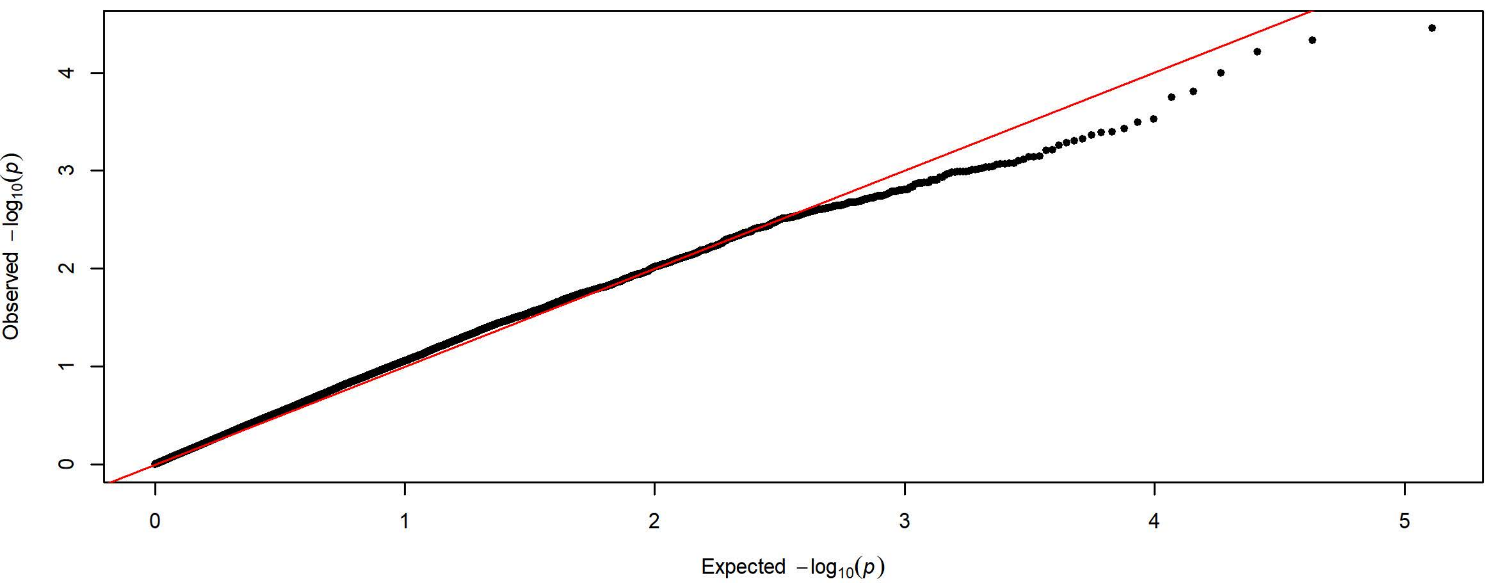


**Q-Q Plot Probability of Magazine Entry Day 2 - Charles River 4 Subgroups - 198k SNPs (n=1726)**

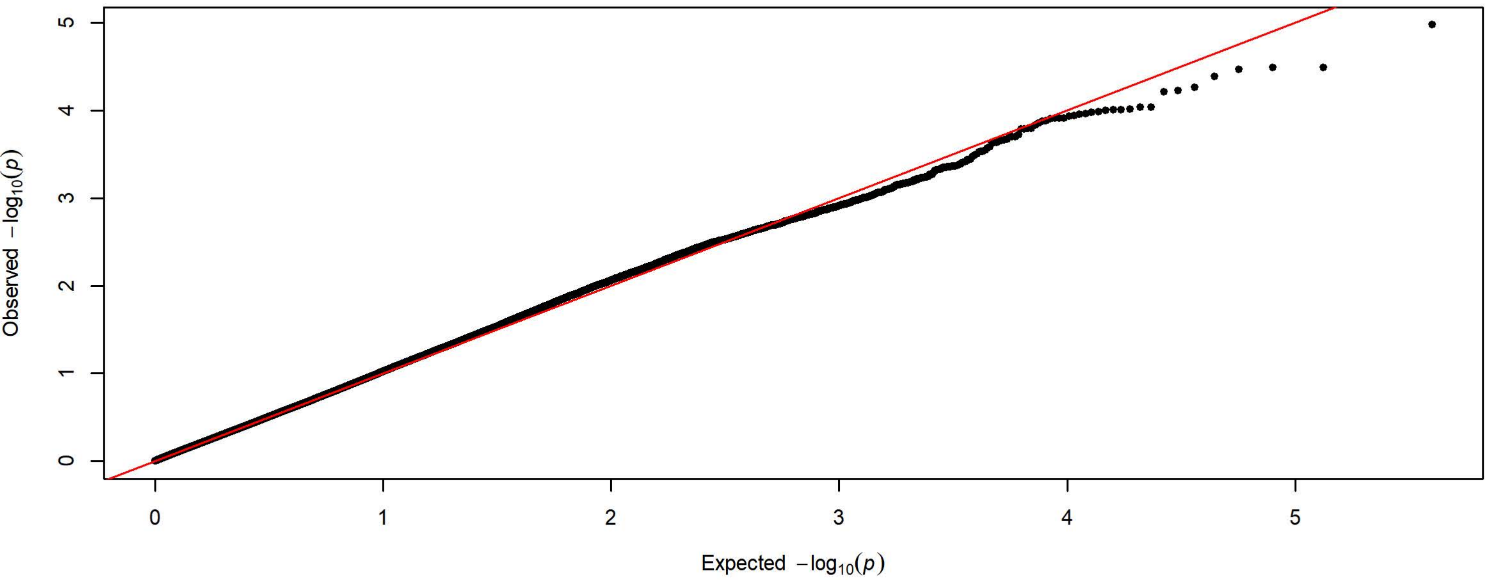


**Q-Q Plot Probability of Magazine Entry Day 2 - Harlan 3 Subgroups - 83k SNPs (n=2208)**

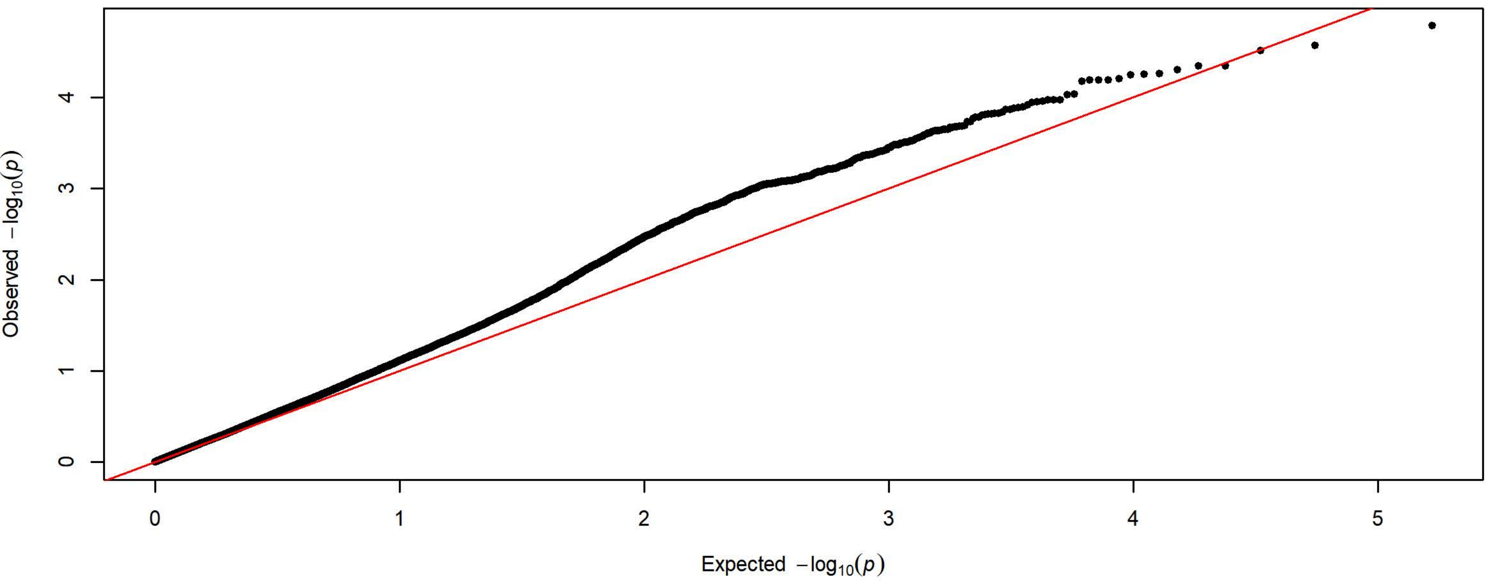

**Q-Q Plot Probability of Magazine Entry Day 3 - Meta-analysis of 7 Subgroups - 64k SNPs (n=3932)**

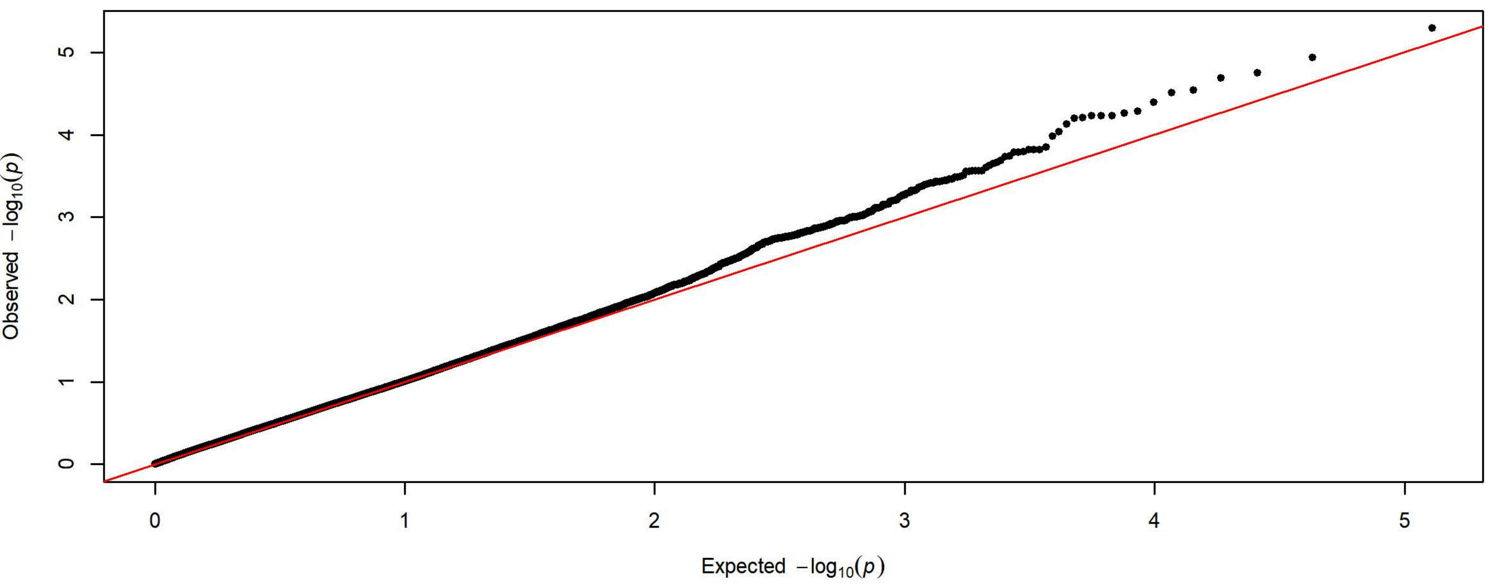


**Q-Q Plot Probability of Magazine Entry Day 3 - Charles River 4 Subgroups - 198k SNPs (n=1727)**

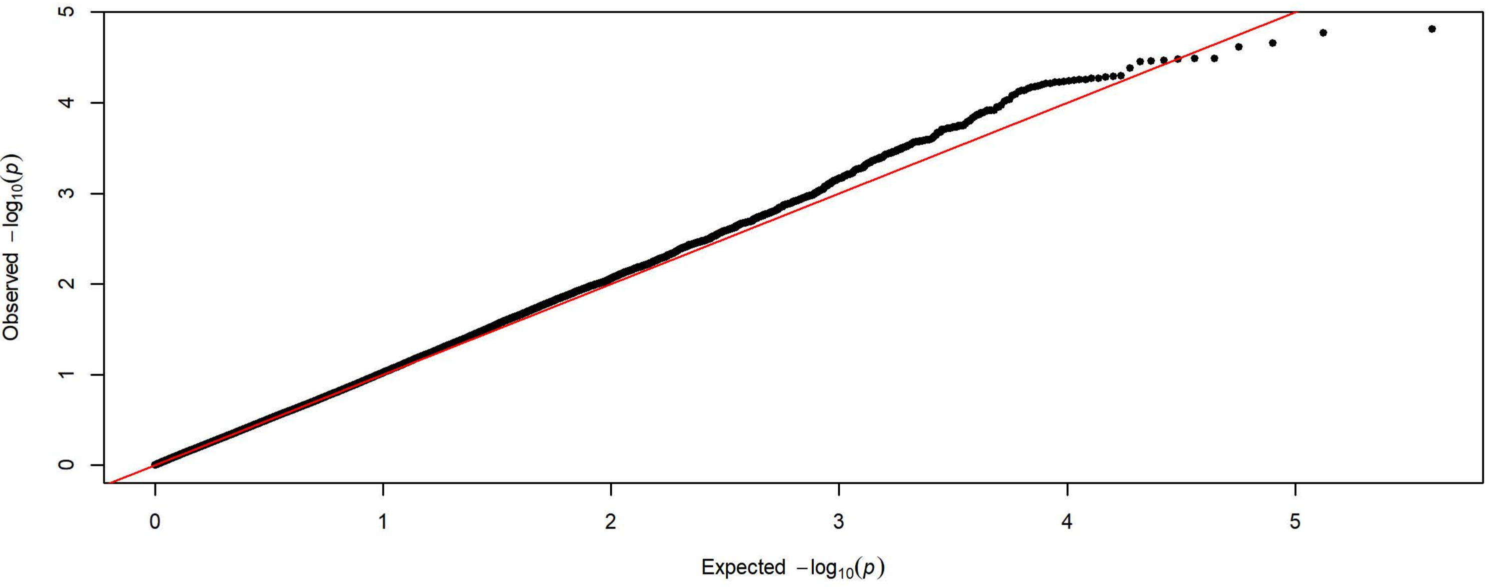


**Q-Q Plot Probability of Magazine Entry Day 3 - Harlan 3 Subgroups - 83k SNPs (n=2205)**

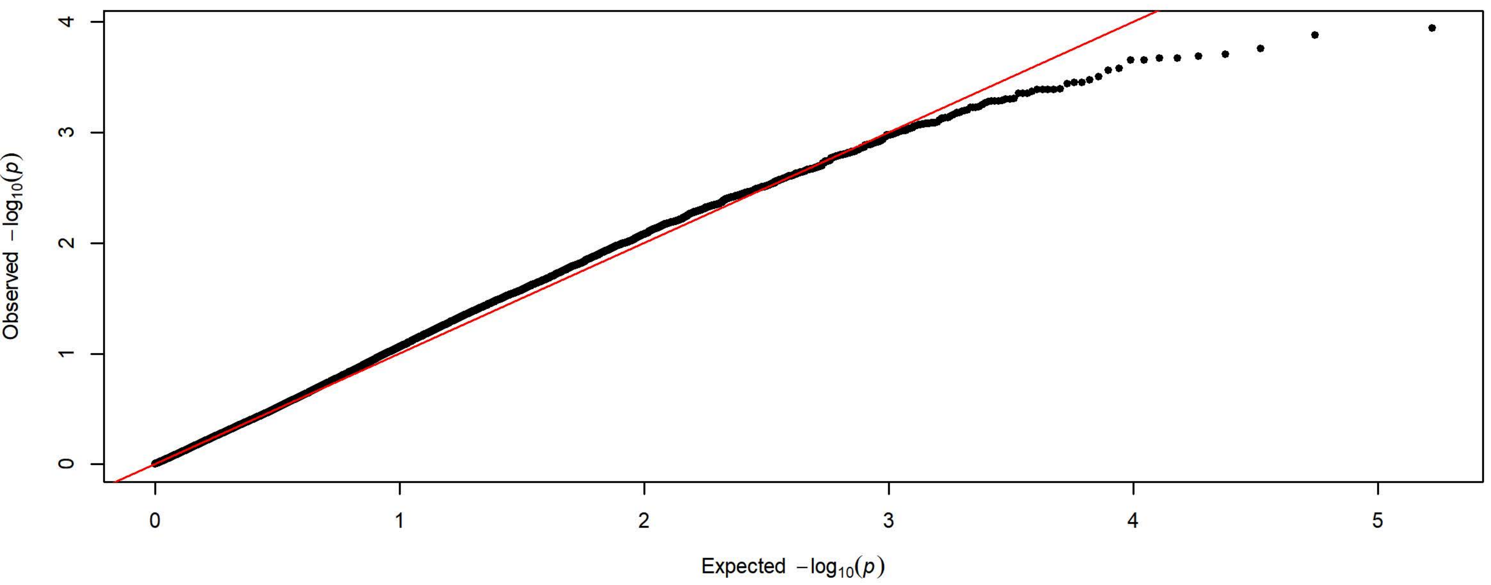

**Q-Q Plot Probability of Magazine Entry Day 4 - Meta-analysis of 7 Subgroups - 64k SNPs (n=3936)**

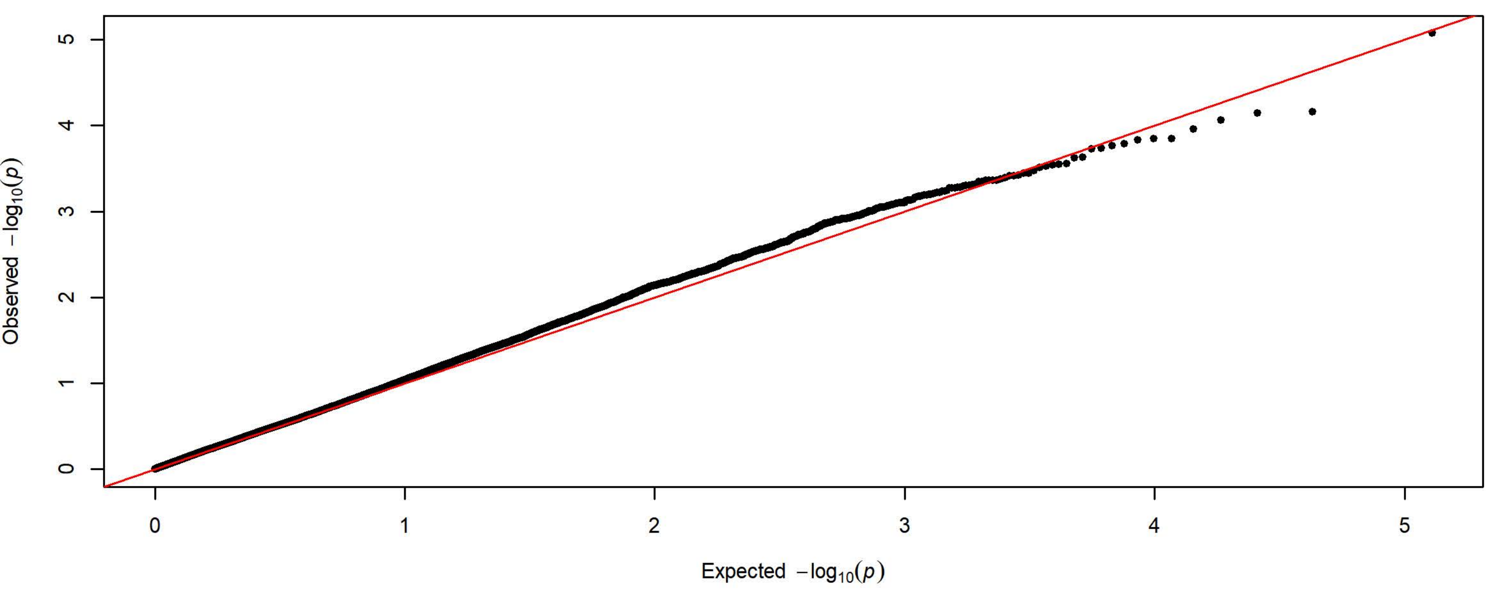


**Q-Q Plot Probability of Magazine Entry Day 4 - Charles River 4 Subgroups - 198k SNPs (n=1728)**

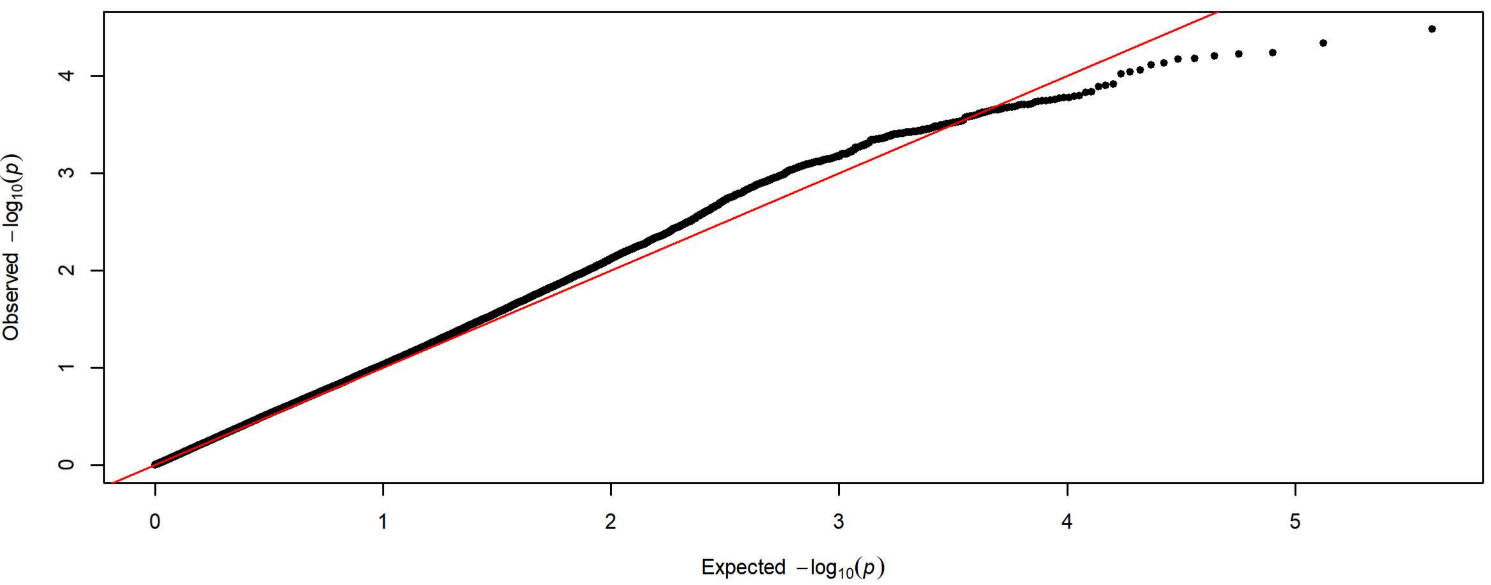


**Q-Q Plot Probability of Magazine Entry Day 4 - Harlan 3 Subgroups - 83k SNPs (n=2208)**

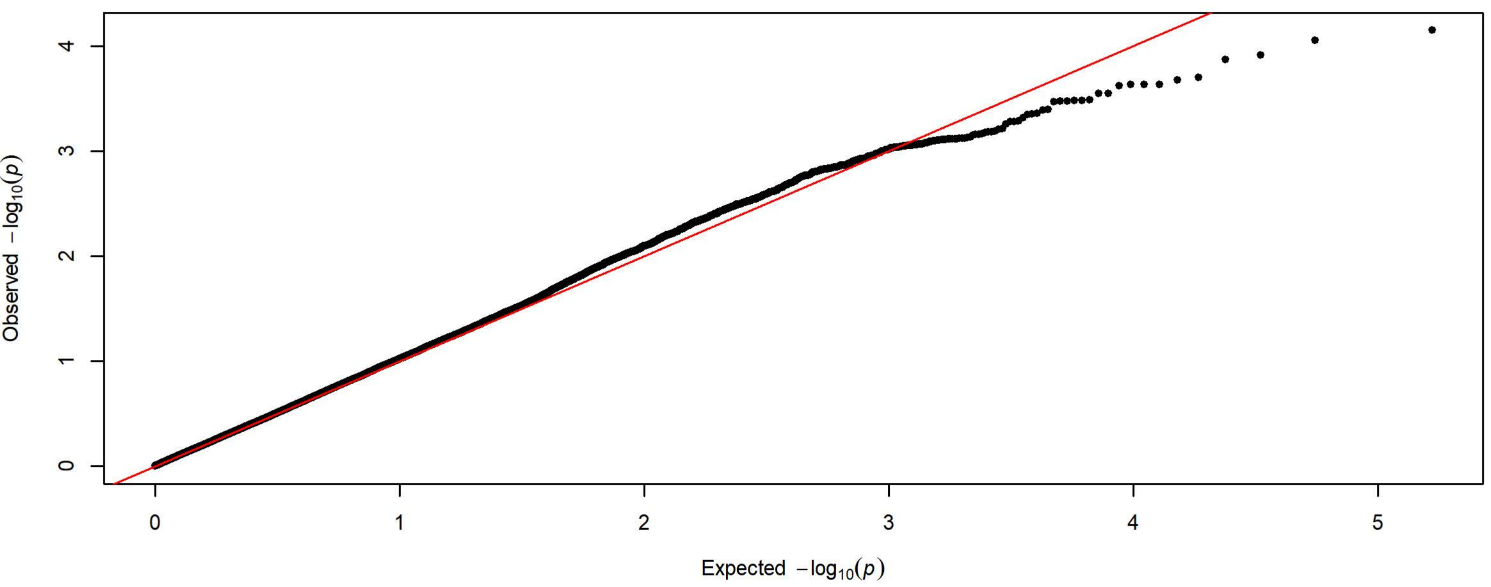

**Q-Q Plot Probability of Magazine Entry Day 5 - Meta-analysis of 7 Subgroups - 64k SNPs (n=3936)**

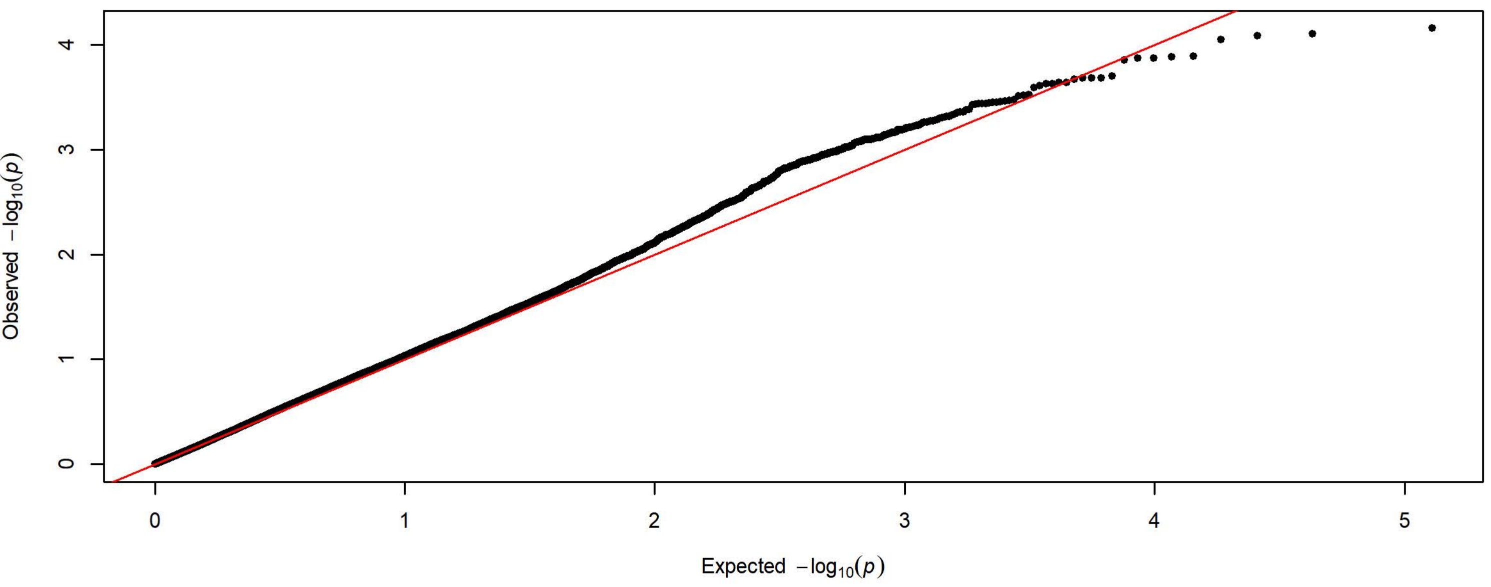


**Q-Q Plot Probability of Magazine Entry Day 5 - Charles River 4 Subgroups - 198k SNPs (n=1728)**

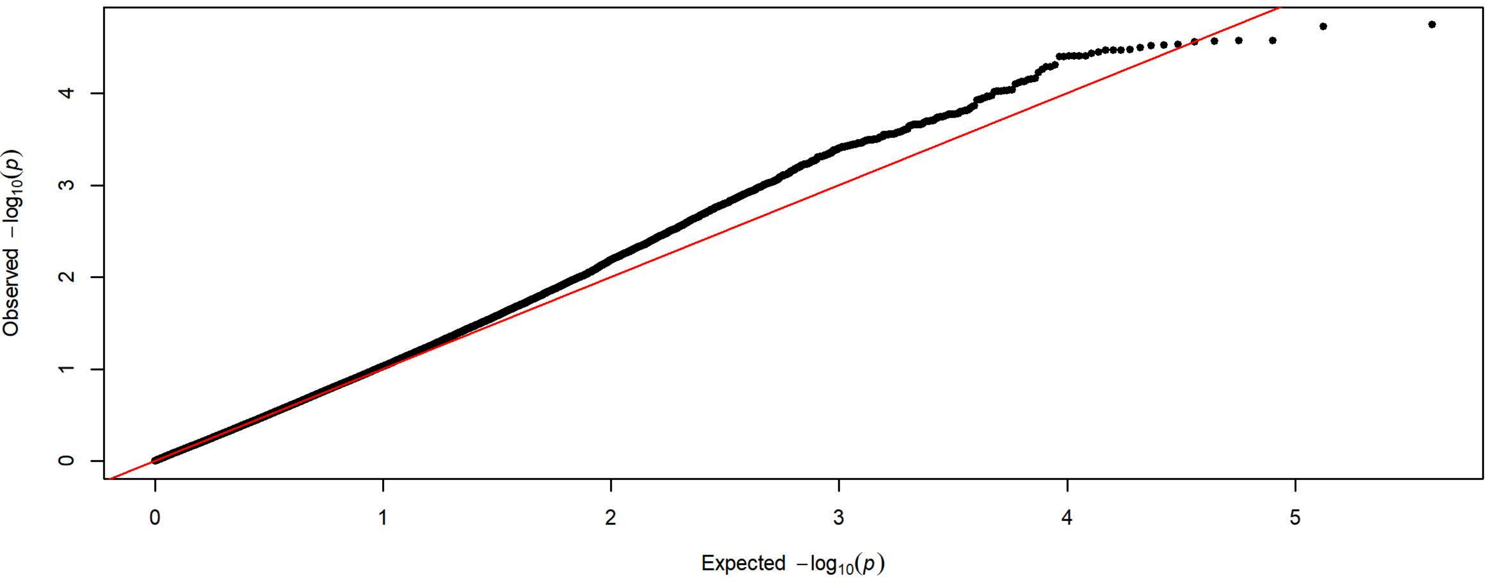


**Q-Q Plot Probability of Magazine Entry Day 5 - Harlan 3 Subgroups - 83k SNPs (n=2208)**

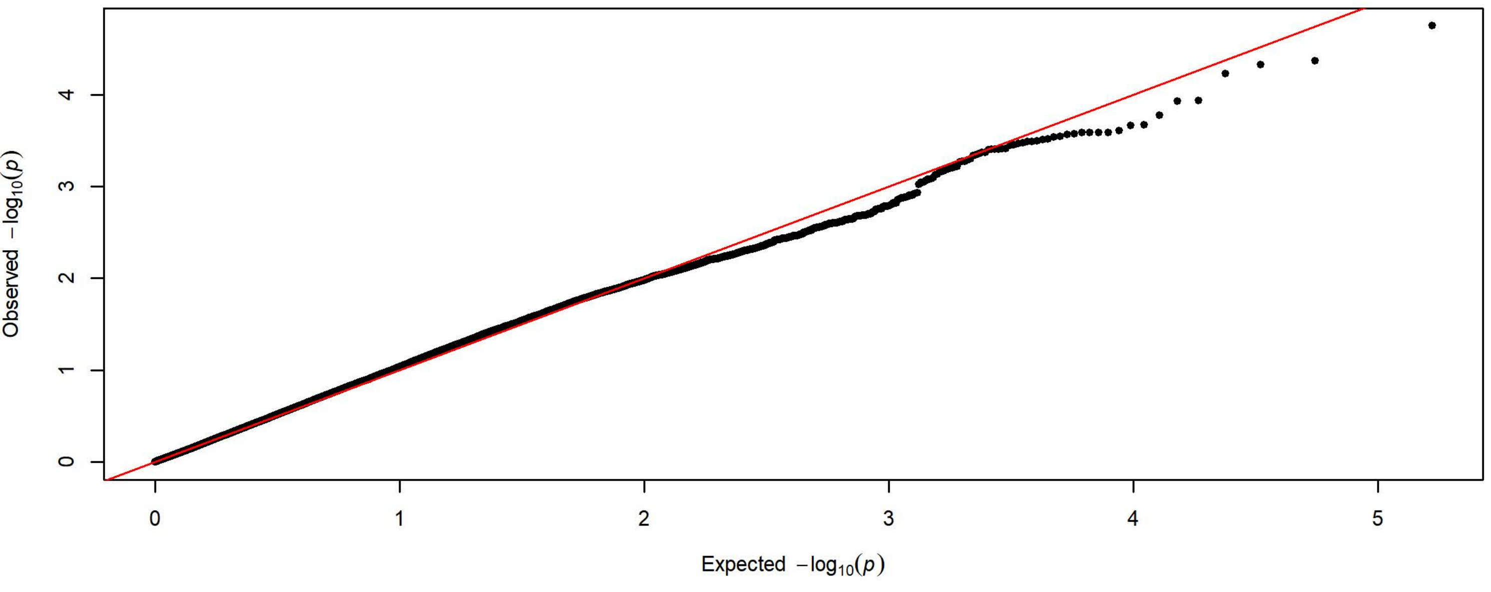

**Q-Q Plot Response Bias Day 1 - Meta-analysis of 7 Subgroups - 64k SNPs (n=3912)**

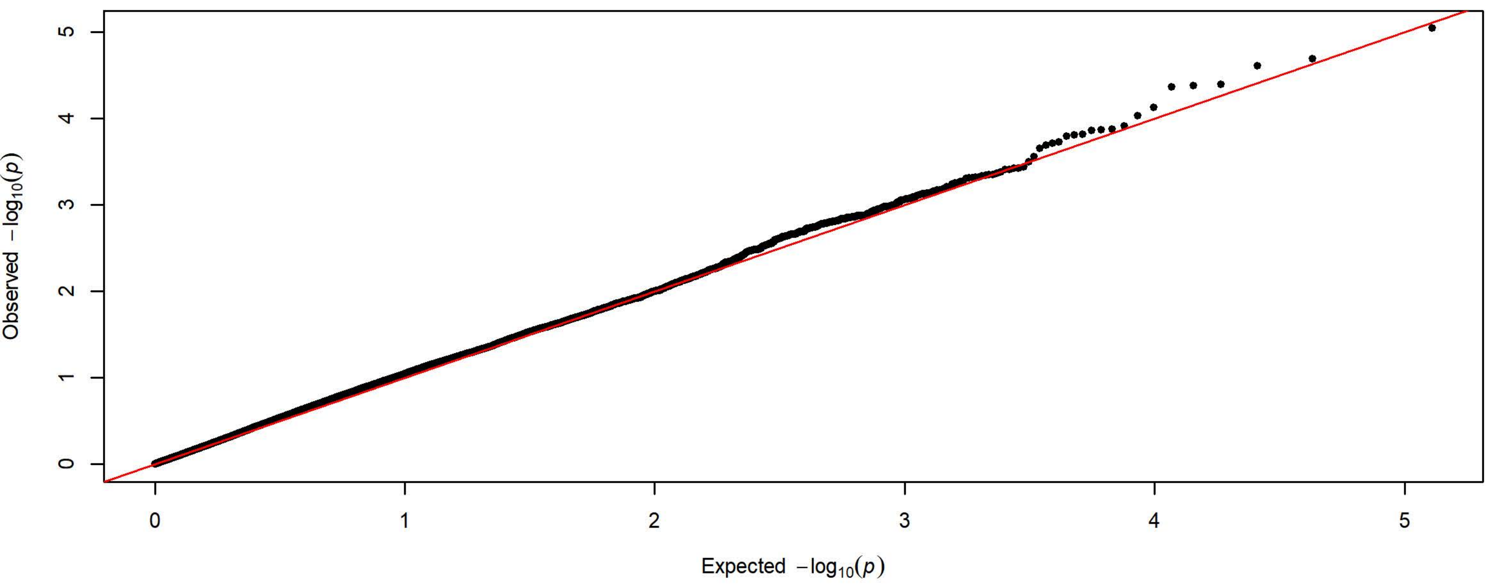


**Q-Q Plot Response Bias Day 1 - Charles River 4 Subgroups - 198k SNPs (n=1720)**

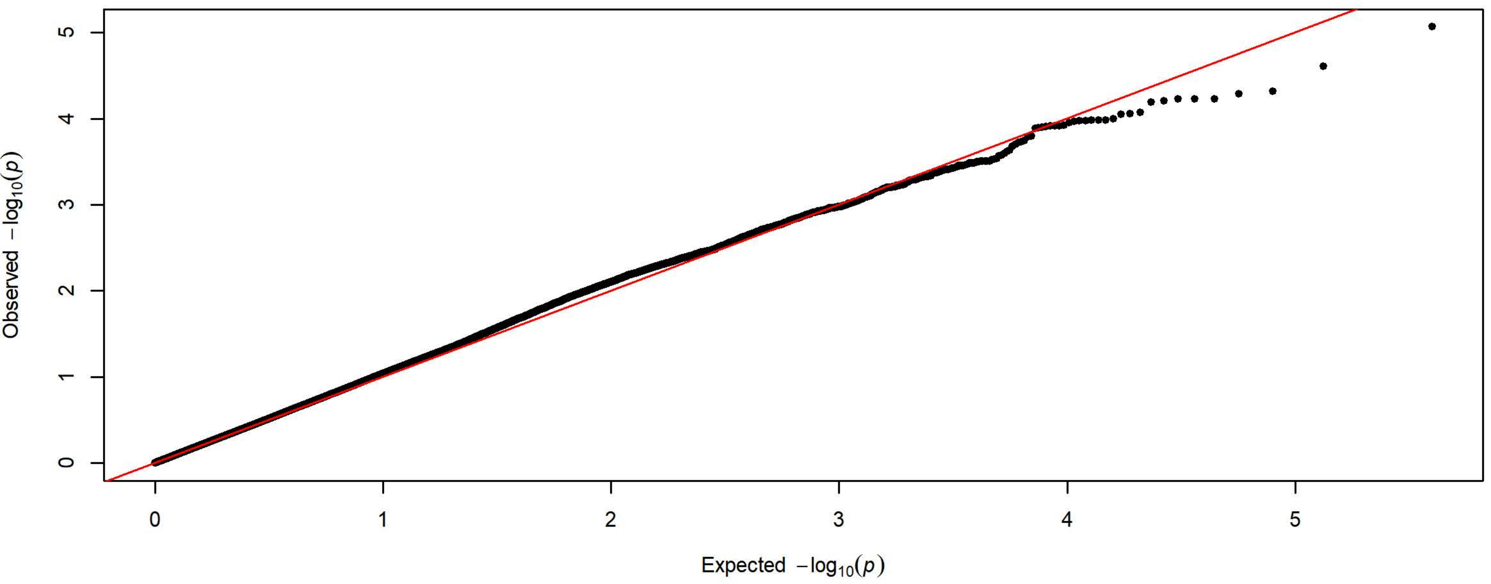


**Q-Q Plot Response Bias Day 1 - Harlan 3 Subgroups - 83k SNPs (n=2192)**

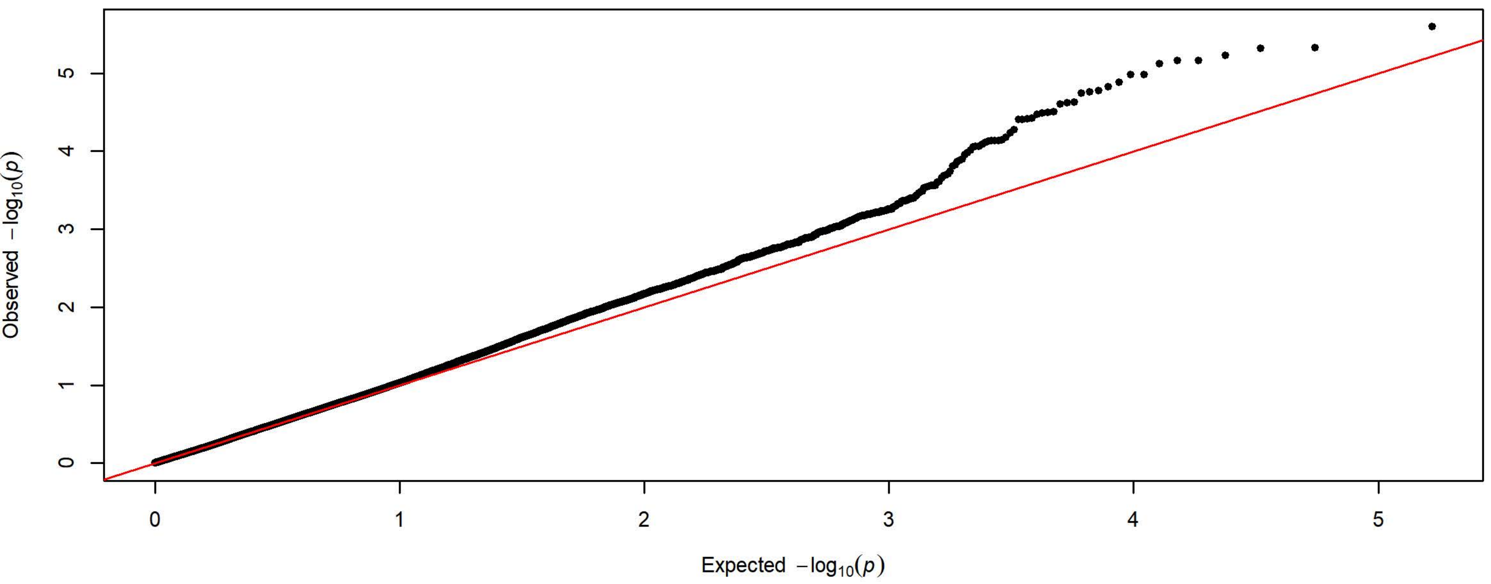

**Q-Q Plot Response Bias Day 2 - Meta-analysis of 7 Subgroups - 64k SNPs (n=3919)**

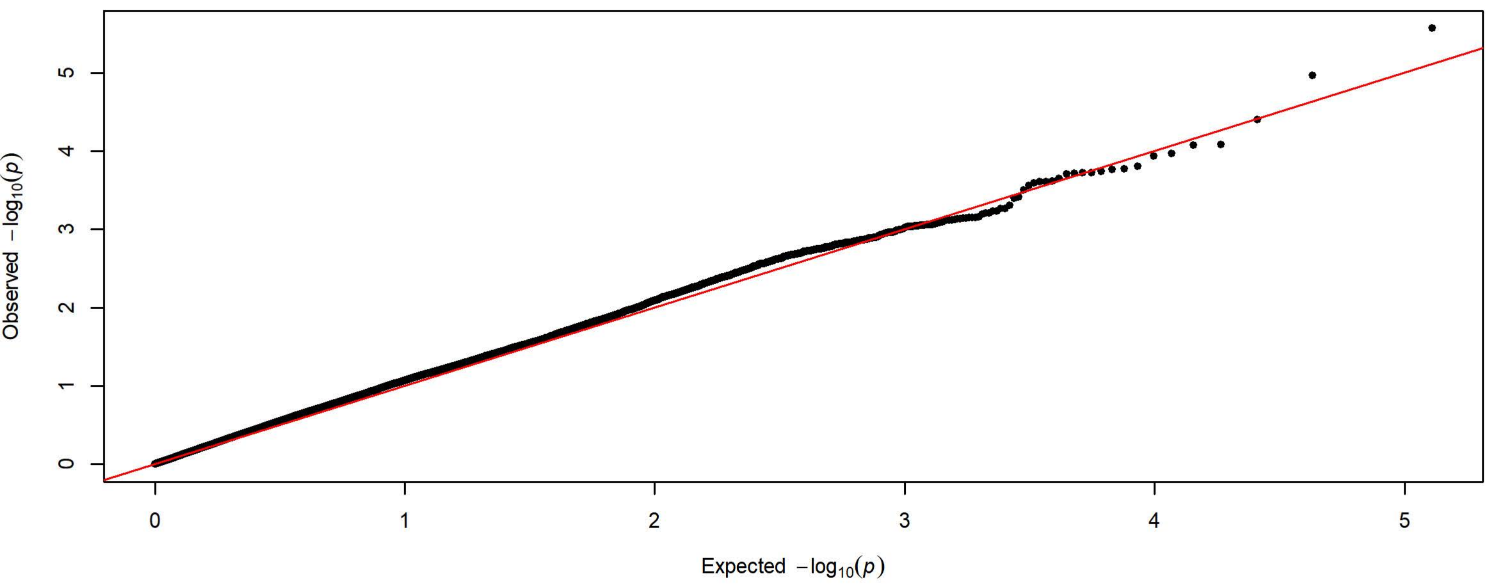


**Q-Q Plot Response Bias Day 2 - Charles River 4 Subgroups - 198k SNPs (n=1719)**

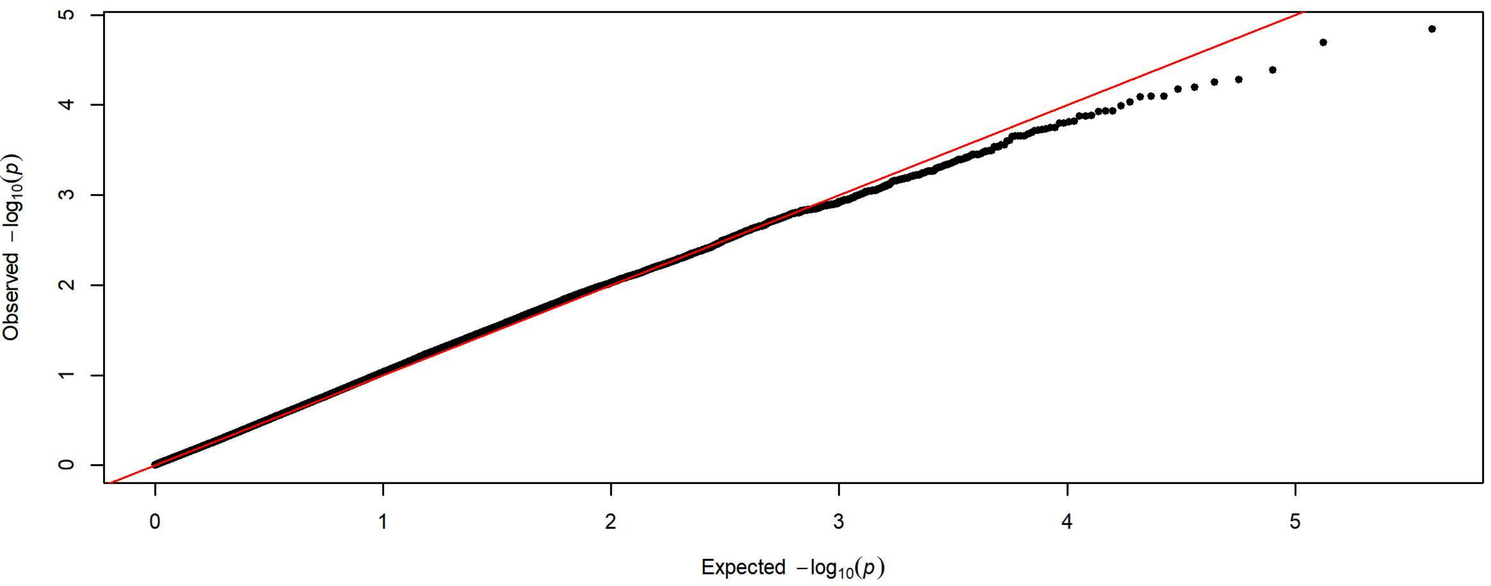


**Q-Q Plot Response Bias Day 2 - Harlan 3 Subgroups - 83k SNPs (n=2200)**

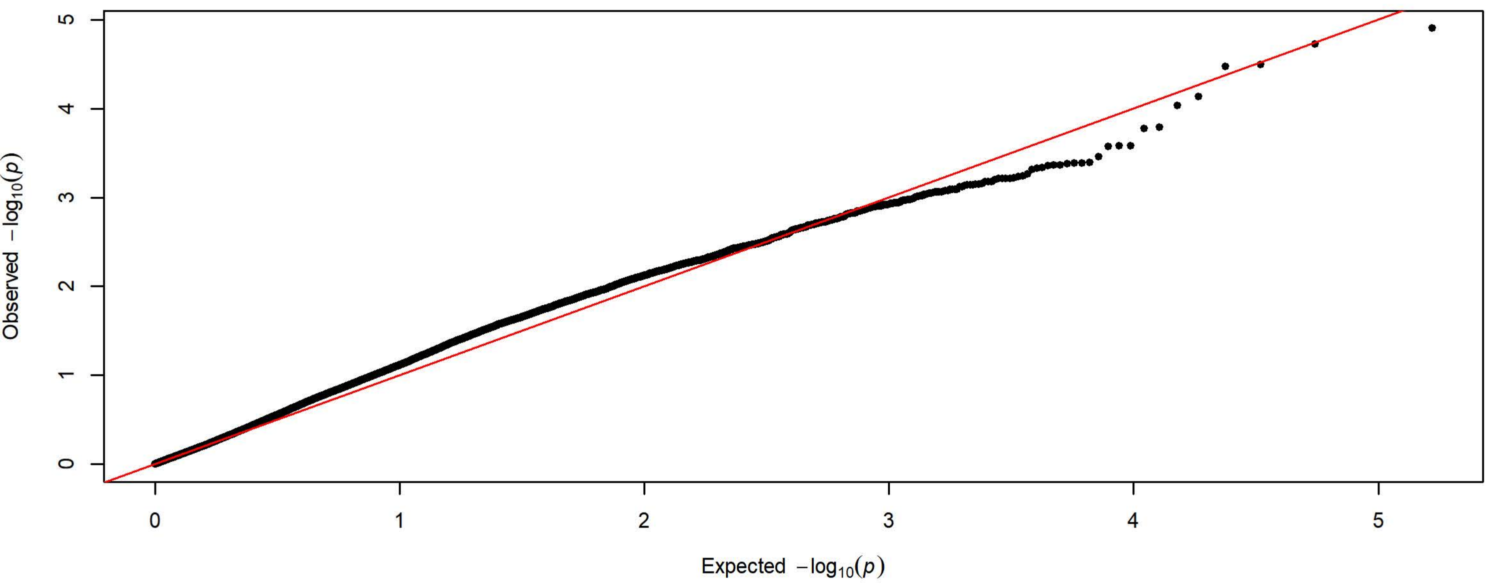

**Q-Q Plot Response Bias Day 3 - Meta-analysis of 7 Subgroups - 64k SNPs (n=3923)**

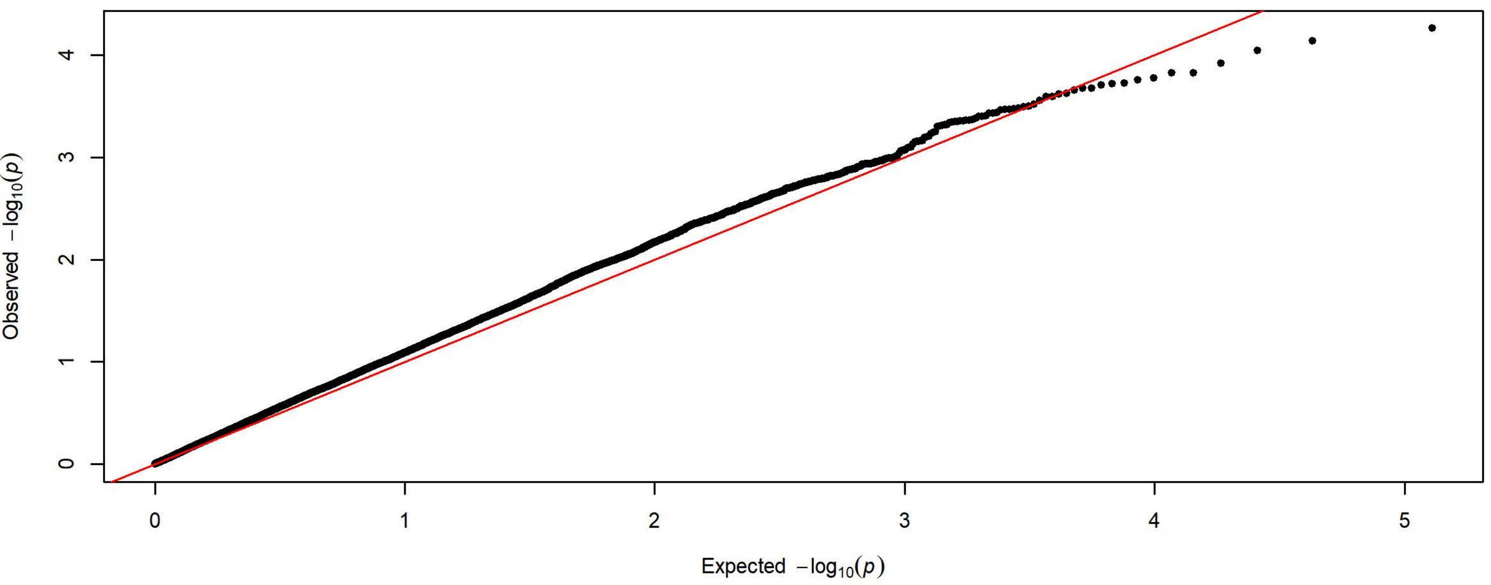


**Q-Q Plot Response Bias Day 3 - Charles River 4 Subgroups - 198k SNPs (n=1722)**

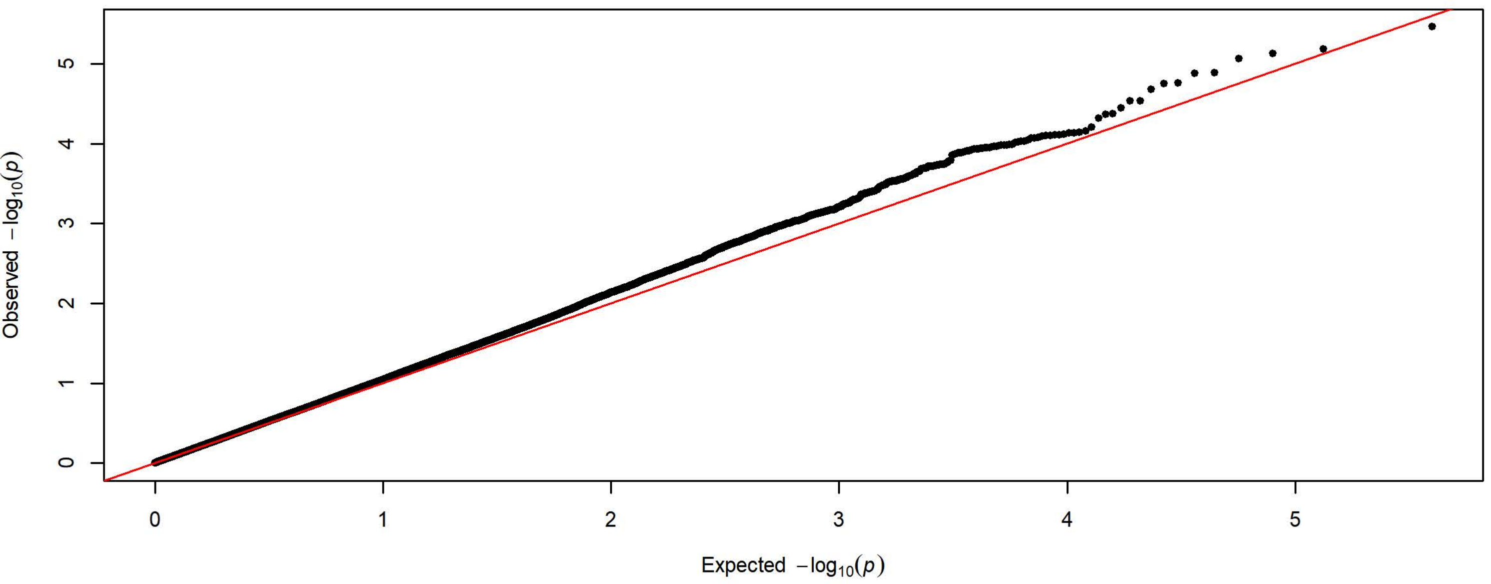


**Q-Q Plot Response Bias Day 3 - Harlan 3 Subgroups - 83k SNPs (n=2201)**

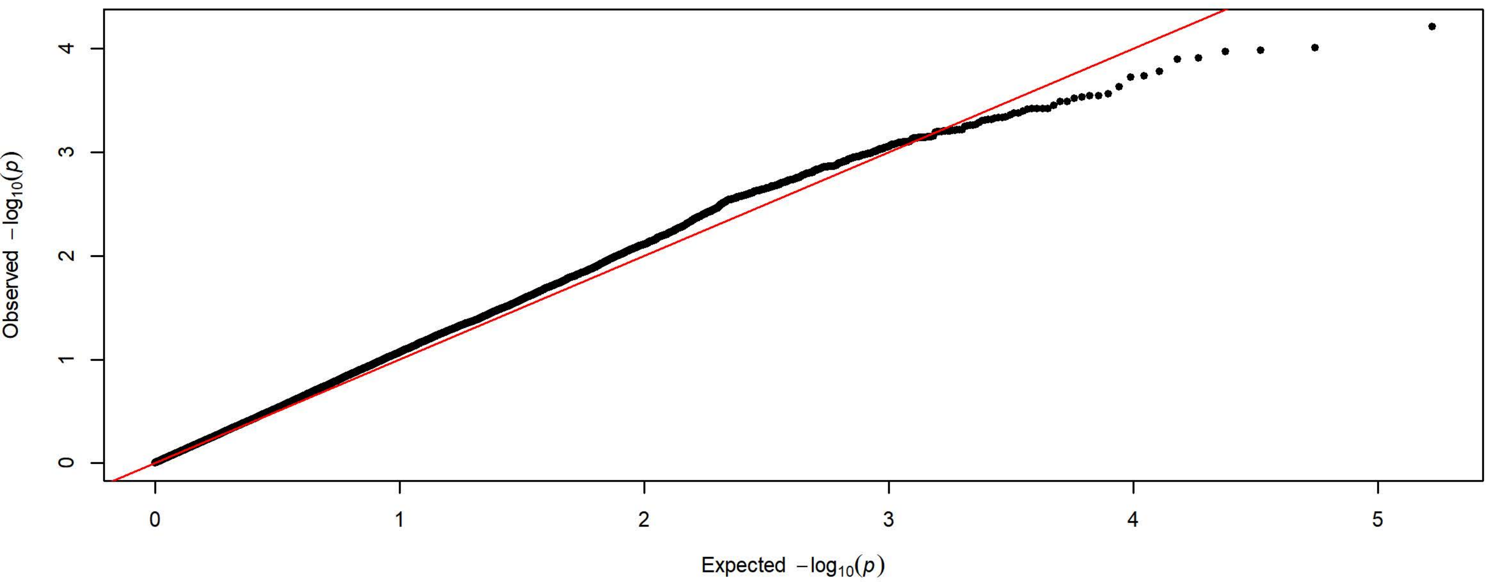

**Q-Q Plot Response Bias Day 4 - Meta-analysis of 7 Subgroups - 64k SNPs (n=3935)**

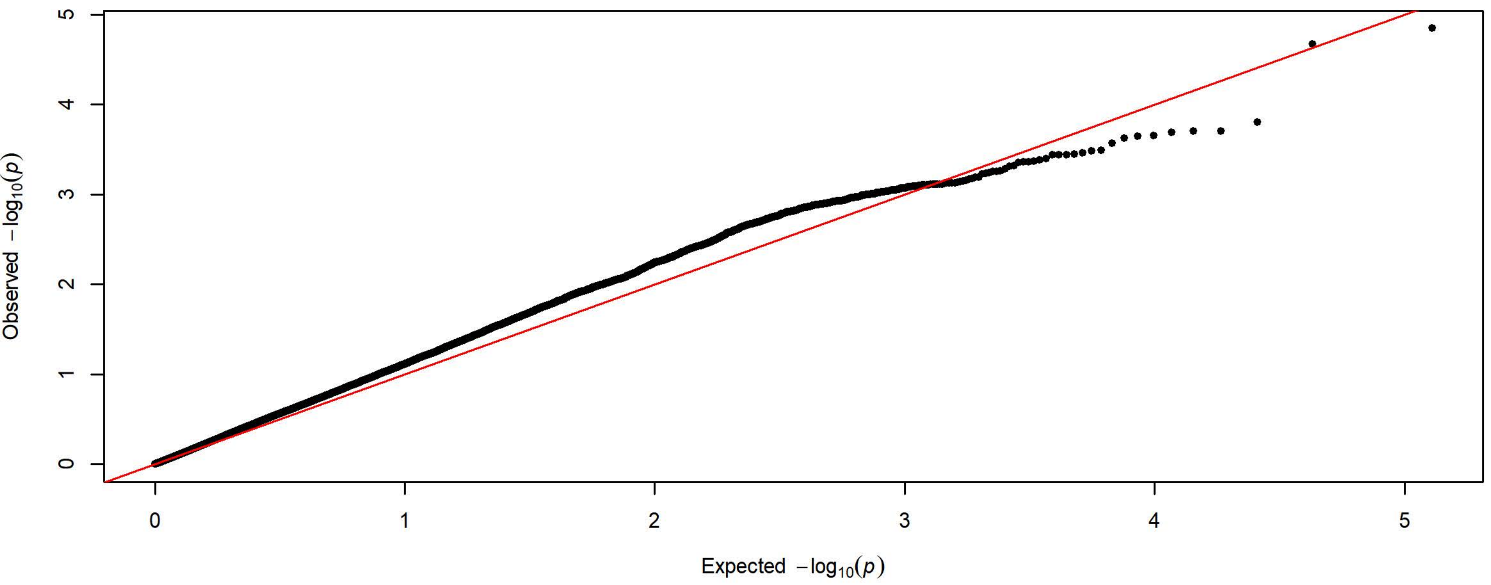


**Q-Q Plot Response Bias Day 4 - Charles River 4 Subgroups - 198k SNPs (n=1727)**

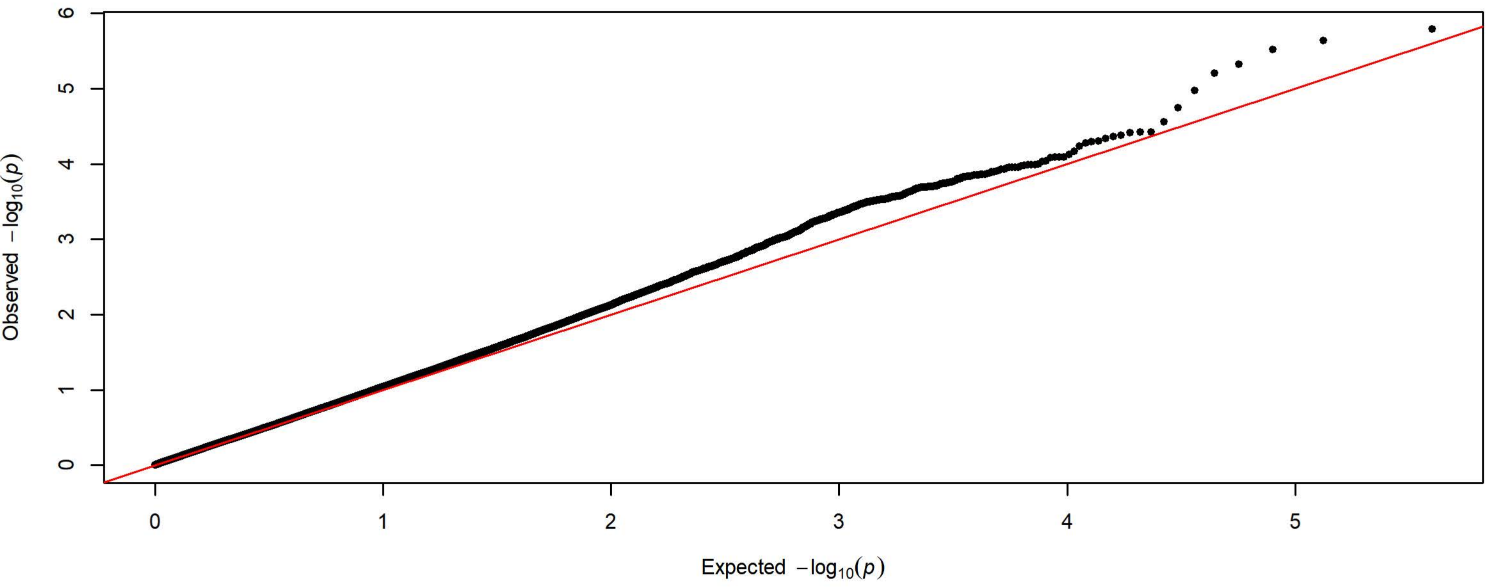


**Q-Q Plot Response Bias Day 4 - Harlan 3 Subgroups - 83k SNPs (n=2208)**

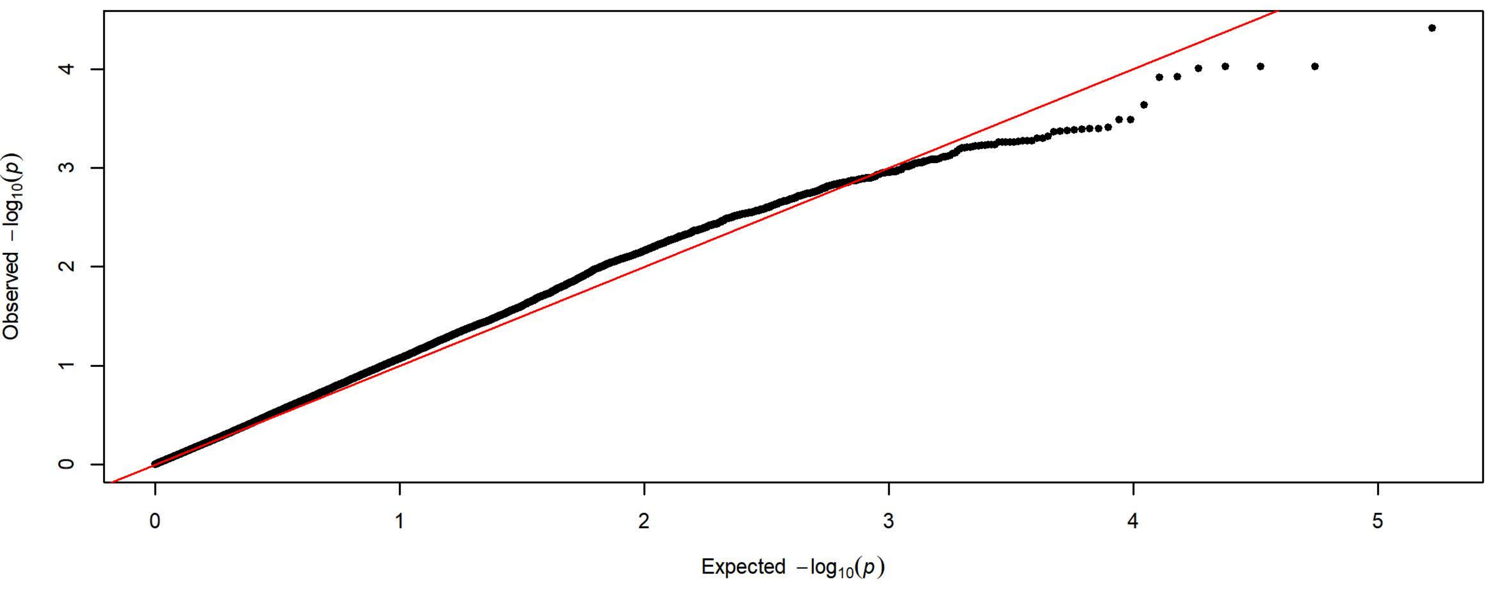

**Q-Q Plot Response Bias Day 5 - Meta-analysis of 7 Subgroups - 64k SNPs (n=3933)**

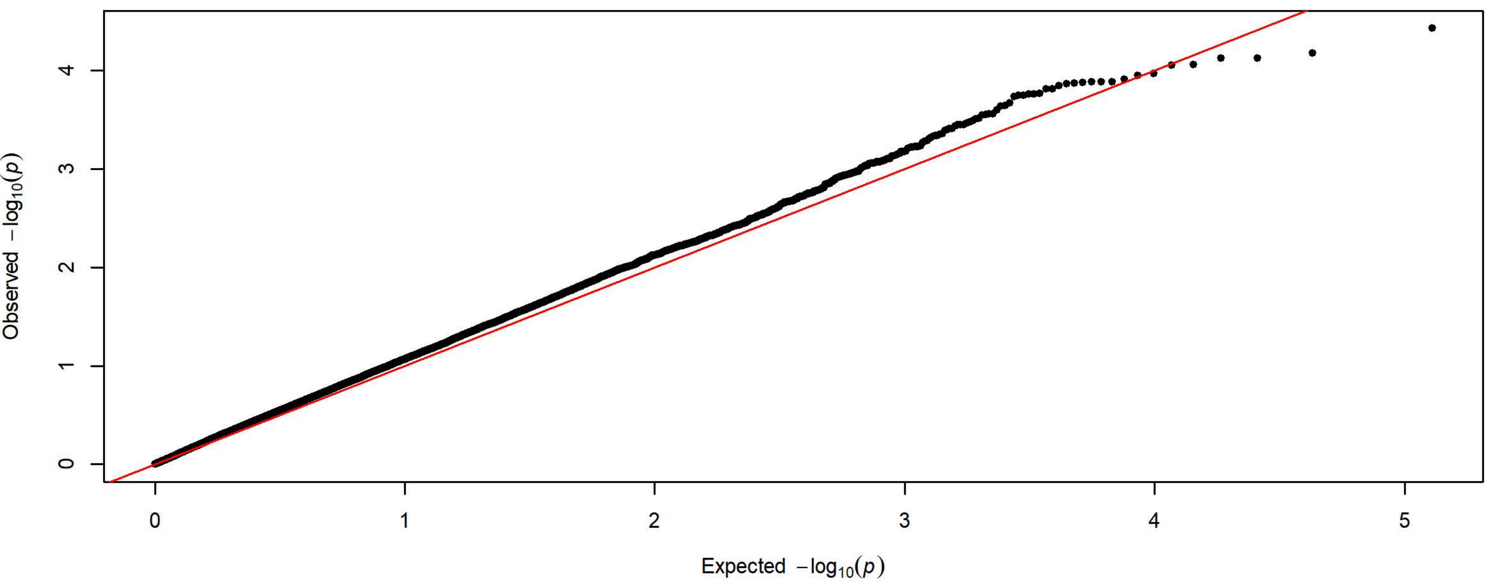


**Q-Q Plot Response Bias Day 5 - Charles River 4 Subgroups - 198k SNPs (n=1726)**

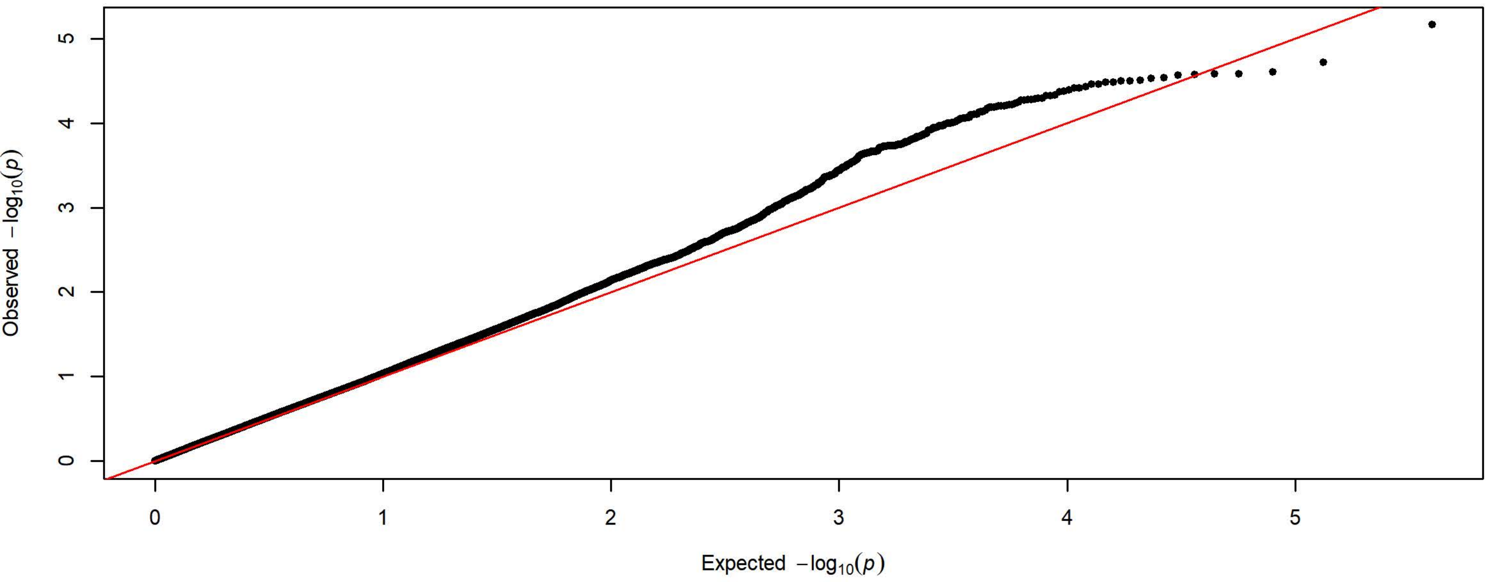


**Q-Q Plot Response Bias Day 5 - Harlan 3 Subgroups - 83k SNPs (n=2207)**

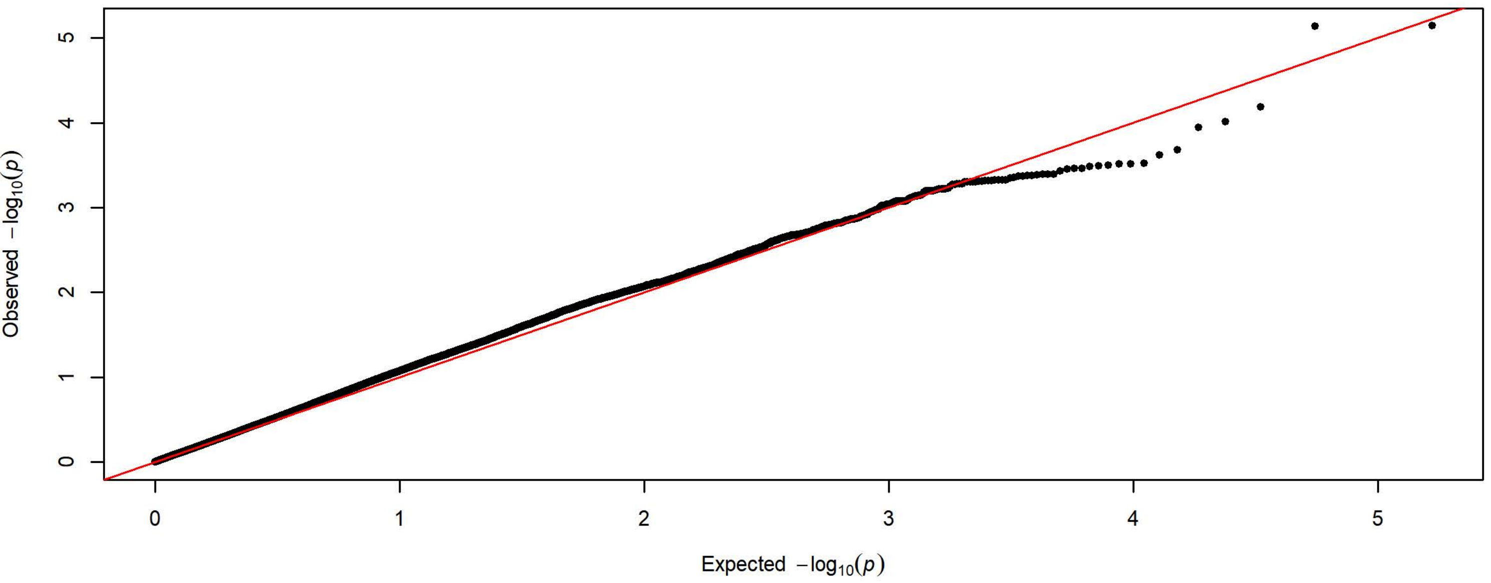
